# Investigating cocaine- and abstinence-induced effects on astrocyte gene expression in the nucleus accumbens

**DOI:** 10.1101/2024.08.05.606656

**Authors:** Janay P. Franklin, Anze Testen, Piotr A. Mieczkowski, Austin Hepperla, Gogce Crynen, Jeremy M. Simon, Jonathan D. Wood, Eden V. Harder, Tania J. Bellinger, Emily A. Witt, N. LaShae Powell, Jonathan W. VanRyzin, Kathryn J. Reissner

**Author notes:** Address correspondence to KJR, CB 3270 Department of Psychology & Neuroscience, UNC CH, 27599.

## Abstract

In recent years, astrocytes have been increasingly implicated in cellular mechanisms of substance use disorders (SUD). Astrocytes are structurally altered following exposure to drugs of abuse; specifically, astrocytes within the nucleus accumbens (NAc) exhibit significantly decreased surface area, volume, and synaptic colocalization after operant self-administration of cocaine and extinction or protracted abstinence (45 days). However, the mechanisms that elicit these morphological modifications are unknown. The current study aims to elucidate the molecular modifications that lead to observed astrocyte structural changes in rats across cocaine abstinence using astrocyte-specific RiboTag and RNA-seq, as an unbiased, comprehensive approach to identify genes whose transcription or translation change within NAc astrocytes following cocaine self-administration and extended abstinence. Using this method, our data reveal cellular processes including cholesterol biosynthesis that are altered specifically by cocaine self-administration and abstinence, suggesting that astrocyte involvement in these processes is changed in cocaine-abstinent rats. Overall, the results of this study provide insight into astrocyte functional adaptations that occur due to cocaine exposure or during cocaine withdrawal, which may pinpoint further mechanisms that contribute to cocaine-seeking behavior.

## INTRODUCTION

Astrocytes serve diverse roles in the brain, including blood-brain-barrier integrity (Cabezas et al., 2014), glutamate uptake (Mahmoud et al., 2019), synaptic regulation (Ota et al. 2013; Lui et al., 2021), immune responses (Burda et al., 2017), and metabolic homeostasis (Zhang et al., 2023). These roles are facilitated by a complex morphology, including GFAP-positive primary branches, as well as fine, membranous peripheral astrocyte processes (PAPs) (Derouiche and Frotscher, 2001; Zhou et al., 2019), which communicate with synapses and modulate synaptic function (Chung et al., 2015; Zhou et al., 2019). Synaptic regulation by astrocytes is accomplished in part through the governance of glutamate homeostasis via their abundant expression of the glutamate antiporter system xC- and glutamate transporter, glutamate transporter 1 (GLT-1) (Cho and Bannai, 1990; Mahmoud et al., 2019; Peterson and Binder, 2019). Disruptions in glutamate homeostasis within the nucleus accumbens (NAc) occur following rat self-administration of drugs including cocaine, nicotine, and heroin, and contribute to mechanisms of cocaine seeking behavior (Kalivas, 2009).

In addition to drug-dependent disruptions in astrocyte-mediated glutamate homeostasis, other investigations have revealed effects of rodent drug self-administration on structural features of astrocytes (Adermark and Bowers, 2016; Testen et al., 2018; Kruyer et al., 2019; Namba et al., 2020; Siemsen et al., 2023). Further, synaptic colocalization of NAc astrocytes with both pre- and post-synaptic markers is decreased following cocaine or heroin self-administration and extinction or extended abstinence (Scofield et al., 2016; Testen et al., 2018; Kruyer et al., 2019; Kim et al., 2022), suggesting that NAc astrocytes exhibit decreased proximity to synaptic elements, perhaps contributing to impaired regulation of glutamate homeostasis. Of relevance to the current study, Kim et al. reported that NAc astrocytes are approximately 40% smaller in structure (surface area and volume) following long-access cocaine self-administration (6h/day for 10 days) and 45 days of abstinence compared to astrocytes from saline-abstinent rats (Kim et al., 2022). However, the underlying mechanisms of this cocaine- and abstinence-induced retracted phenotype are unknown.

In order to address this question in an unbiased manner, we employed astrocyte-specific RiboTag RNA-Seq in rat NAc astrocytes following cocaine or saline self-administration across early and late abstinence (Sanz et al., 2019). This method employs astrocyte-specific AAV5 GfaABC1D-Rpl22-HA to enable immunoprecipitation of HA-tagged ribosome-associated mRNAs from NAc astrocytes, thus providing insight into how NAc astrocyte gene expression is affected by a history of cocaine self-administration and home cage abstinence and how cocaine-induced functional deficits in NAc astrocytes may promote cocaine-seeking behaviors. Several others (Boisvert et al., 2018; Yu et al., 2020; Diaz-Castro et al., 2021) have utilized this technique either with AAV or transgenic mice to examine astrocyte gene expression changes in diverse contexts, including development and aging, neuroinflammation, and pharmacological activation of astrocytes. In relation to cocaine, other previous studies have investigated the effects of cocaine on gene expression in varied brain regions including the ventral tegmental area, but these studies used bulk RNA-Seq (Walker et al., 2018; Campbell et al., 2021), which lacks ability to examine cell-specific gene expression. To our knowledge, the current study is the first to utilize RiboTag RNA-Seq to examine temporal and cell-specific effects of cocaine and abstinence on NAc astrocyte gene expression.

## MATERIALS AND METHODS

### Animals and Surgical Procedures

Male Sprague Dawley rats (225-250 g, ∼6 weeks of age) used in this study were purchased from Enivgo and were housed in a temperature- and humidity-controlled facility on a reverse light-dark cycle (lights on at 7 pm; lights off at 7 am). Upon arrival, rats were single-housed in Plexiglass cages and allowed to acclimate to the animal facility for one week, with food and water ad libitum. Prior to the start of behavioral experiments, rats were placed on a restricted food diet to maintain a steady weight gain of a few g per day, which lasted throughout pre- and post-surgical procedures, as well as food training procedures. Rats returned to an ad libitum food diet following the third day of operant self-administration. All procedures were approved by the Institutional Animal Care and Use Committee of the University of North Carolina at Chapel Hill and followed the Guide for the Care and Use of Laboratory Animals (Institute of Laboratory Animal Resources on Life Sciences, National Research Council, 1996).

Rats were anesthetized with ketamine (100 mg/kg) and xylazine (7 mg/kg) for all surgical procedures. For catheter implantation procedures, rats were surgically implanted with chronic indwelling jugular catheters, composed of guide cannulas (cat #C313G, Plastics One), silastic tubing (.025 ID, .047 OD Bio-sil), and subdermal surgical mesh (Atrium). Following catheter implantation, rats received bilateral 2 uL viral injections of rAAV5 pZac2.1-GfaABC1D-Rpl22-HA (Addgene cat #111811) into the nucleus accumbens (6° angle, coordinates (mm): +1.5 anterior/posterior, +2.6 medial/lateral, -7.2 dorsal/ventral). The virus was microinjected at an infusion rate of 0.1 uL/min, followed by a diffusion period of 15 minutes. Following diffusion, microinjectors were removed slowly over 1-2 minutes. Plasmids for AAV manufacturing used were pZac2.1.GfaABC1D-Rpl22-HA (Diaz-Castro et al., 2019; Diaz-Castro et al., 2021; Yu et al., 2020). The plasmid was provided by Addgene and packaged into an AAV at the UNC Viral Vector Core. After surgery, rats recovered for 5 days and received post-operative care, including ketofen (5 mg/kg, s.c.) for the first 3 days of recovery. Additionally, rats received daily catheter flushing of heparin (100 u/mL, i.v.) and gentamicin (5 mg/mL, i.v.), to ensure patency and minimize infection until the conclusion of the self-administration period. Before the start of self-administration procedures, additional confirmation of catheter patency was examined by administration of a subthreshold dose of propofol (10 mg/mL, 0.1 mL).

### Self-Administration and Behavioral Training

All behavioral training and self-administration procedures occurred in sound-attenuated operant conditioning chambers (Med Associates, Fairfax, Vermont). Prior to surgical procedures, rats received a single operant self-administration training session, where they received food pellets as a reward. The food training session lasted 6 hours, and to meet the criteria, rats must have at minimum 100 responses on the active lever. Following recovery from surgical procedures, rats began self-administration of saline or cocaine (6 hr/day for 10 days). Self-administration procedures operated on a FR1 schedule. Presses on the active lever resulted in an infusion of saline or cocaine (0.75 mg/kg/infusion), along with a 5-second light cue and tone and a 20-second timeout period. The number of infusions received was restricted to 80 on the first two days of self-administration but was increased to a maximum of 200 infusions on days 3-10. Cocaine was provided by the NIH Drug Supply Program. Following self-administration, rats underwent either 1 or 45 days of home cage abstinence and were handled twice weekly.

### Immunohistochemistry

To confirm the viral specificity of the RPL22-HA AAV, we first performed immunohistochemistry on naïve male Sprague Dawley rats approximately 3.5 weeks following bilateral viral microinjections of RPL22-HA AAV into the nucleus accumbens (NAc). For these procedures, rats were deeply anesthetized with pentobarbital and transcardially perfused with 4% paraformaldehyde (PFA). Brains were extracted, post-fixed for 24 hrs in PFA at 4°C, and then transferred to a 30% sucrose solution in 1x PB at 4°C for 3-4 days. Brains were coronally sliced at 45μm on a Leica cryostat and prepared for immunohistochemistry. Slices were washed three times in 1X PBS on an orbital shaker at room temperature (RT), followed by blocking incubation for 1 hour at RT in 5% BSA in 0.4% 1X PBST. Following blocking, slices were incubated in primary antibody solution (2.5% BSA in 0.4% 1X PBST) at 4 °C, which contained the following antibodies: mouse anti-HA (BioLegend cat #901513; 1:500), rabbit anti-GFAP (Dako cat #Z0334; 1:500), and chicken anti-Fox3 (NeuN) (Boster Bio cat #M11954-3; 1:1000). After an overnight primary antibody incubation, slices were washed three times in 1X PBS at RT and then incubated with secondary antibody solution (2.5% BSA in 0.4% 1X PBST) for 2 hrs at RT. Secondary antibodies conjugated to Alexa Fluorophores included: goat anti-mouse 488 (ThermoFisher #A-11001; 1:1000), goat anti-rabbit 647 (ThermoFisher #A21244; 1:1000), and goat anti-chicken 408 (ThermoFisher #A48260; 1:1000). After the secondary antibody incubation, the slices received a final wash in 1X PBS for three times at RT and were plated before imaging with SouthernBiotech DAPI Fluoromount-G mounting medium (Fisher Scientific: 0100-20). Images were acquired using a Zeiss LSM 800 confocal-scanning microscope with a 63× oil-immersed objective, two GaAsP detectors and 405, 488, 561, and 640 nm diode lasers at a frame size of 1,024 × 1,024 pixels, 4× averaging, and 1 μm z-step size. Subsequently, AutoQuant (v. X3.0.4, MediaCybernetics) deconvolution software was used and deconvolved optical stacks were imported into Imaris (v. 10) for colocalization analysis.

### RiboTag Immunoprecipitation and RNA Extraction

Following cocaine or saline self-administration and abstinence, rats were euthanized by rapid decapitation at two timepoints: withdrawal day 1 (24 hrs following the last self-administration session) or withdrawal day 45, based on their experimental group assignment. A 3-mm coronal section containing the nucleus accumbens (NAc) was isolated, followed by a 2-mm circular tissue excision to dissect the NAc region and immediately submerged in chilled homogenization buffer. Following homogenization of bilateral punches from each rat, the resulting lysate was incubated with mouse anti-HA (BioLegend cat #901513) while continuously rotating for 4 hours at 4°C. Subsequently, magnetic beads (Invitrogen Dynabeads #10004D) were added to the lysate containing anti-HA antibody for overnight incubation (16-18 hours) on a rotator at 4°C. Following overnight incubations, samples were washed four times with high salt buffer and supplemented with Buffer RLT (Qiagen #79216) before storing at -80°C. RNA purification of immunoprecipitation samples was conducted using Qiagen RNEasy Micro Kit (Qiagen #74004). RNA concentration and quality were tested using the Agilent 2100 Bioanalyzer. All RNA samples used in the study has a RIN number of ≥7.

For experiment 1, half of the total number of saline (n=12) and cocaine (n=11) abstinent rats were euthanized at either withdrawal (WD) 1 or WD45, which were the days in which immunoprecipitation experiments occurred (see Fig. 2A for timeline). RNA extractions and quality testing were performed for each group within 1 week following their immunoprecipitation and stored at -80°C until RNA sequencing.

**Figure 1:**
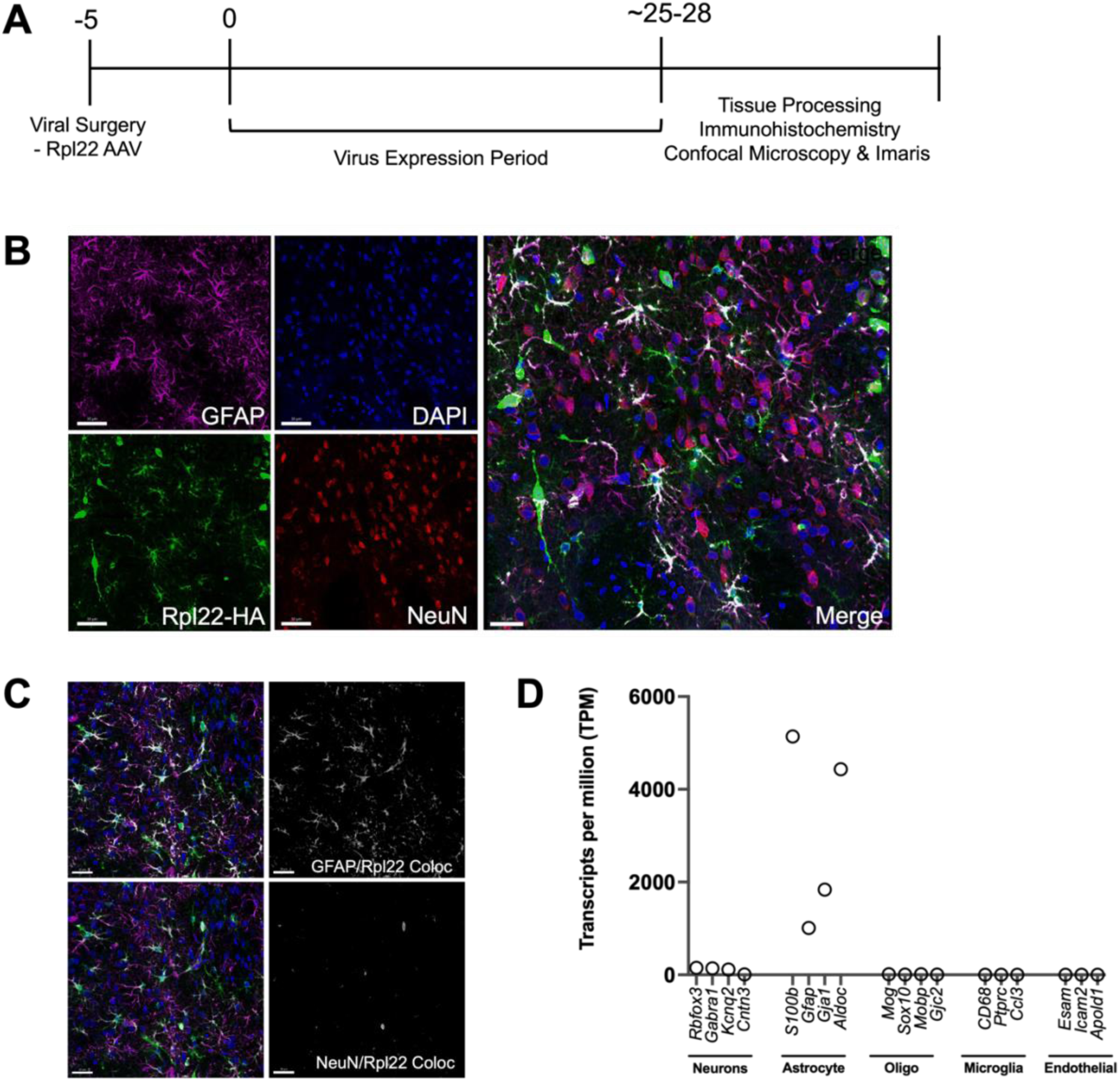
Validation of astrocyte-specific expression of RiboTag (AAV5-GfaABC1D-Rpl22-HA) and RNASeq. **A**) Experimental timeline. **B**) Virally transduced NAc slices stained for anti-HA to detect Rpl22 virus expression within astrocytes; GFAP and NeuN were used to detect astrocytes and neurons, respectively. Rpl22-transduced astrocytes are labeled in white in Merge image (magnification 20X; scale bar: 30 pm). **C**) Colocalization of GFAP + Rpl22 expression in top panel and colocalization of NeuN + Rpl22 expression in bottom panel (magnification 20X; scale bar: 30 pm). **D**) Confirmation of Rpl22 AAV specificity in NAc astrocytes of naive male rats using RNASeq to compare markers of other cell types (neurons, oligodendrocytes, microglia, and endothelial cells).

**Figure 2:**
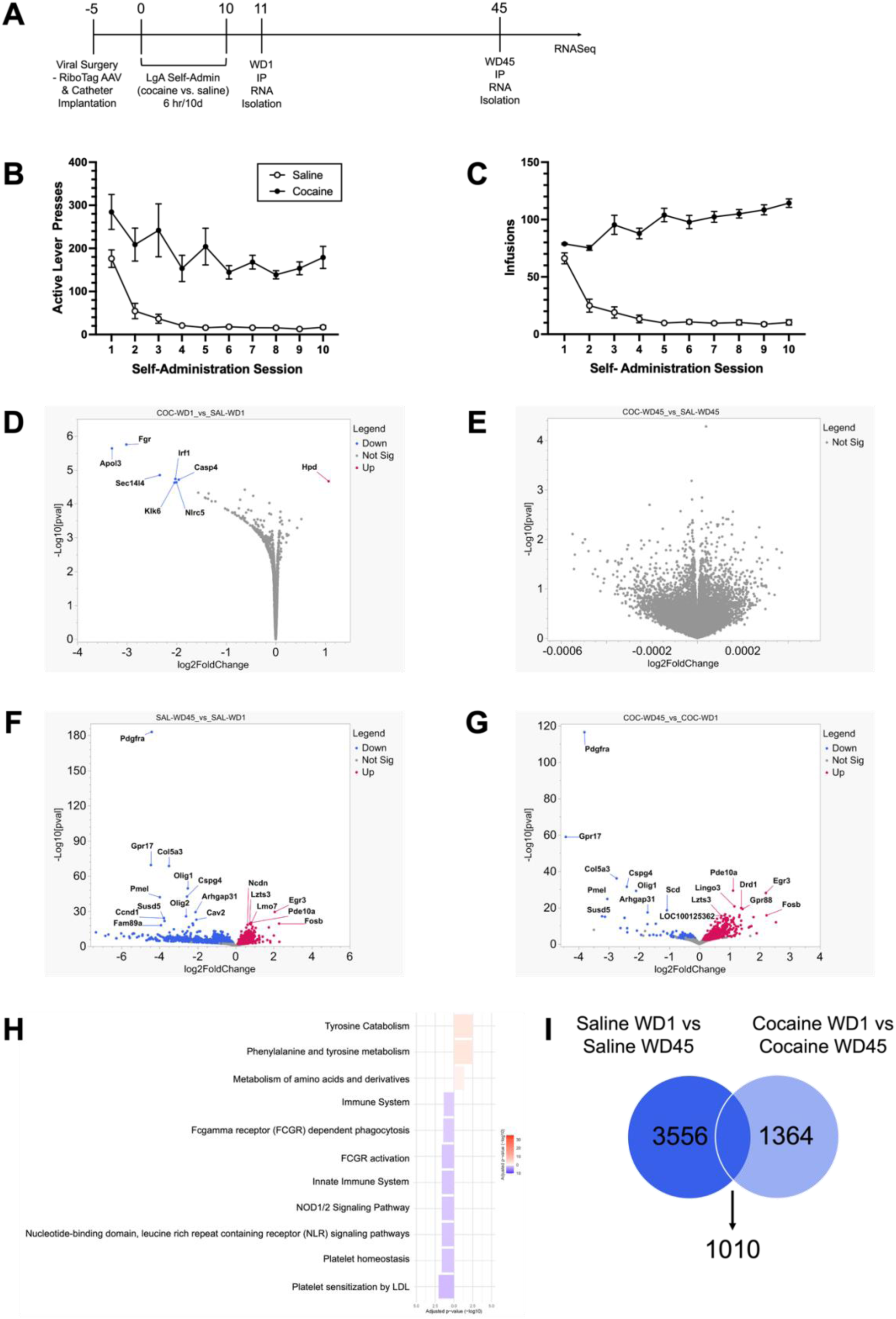
Experiment I. *Cocaine-administering rats displayed significantly increased drug-seeking behavior than saline controls; differential gene expression detected within experimental group comparisons.* A) Timeline for behavior and immunoprecipitation experiments. B) Active lever presses for both saline and cocaine groups shown across self-administration. C) Number of infusions shown for each treatment group throughout self-administration. (p-value < 0.05). Volcano plots showing number of significant differentially expressed genes (DEGs) in experimental comparisons including cocaine vs. saline at withdrawal day (WD1) (D), cocaine vs. saline at WD45 (E), saline WD45 vs. saline WD1 (F), cocaine WD45 vs. cocaine WD1 (G) (nonsignificant: gray, significant downregulated DEGs: blue, and significantly upregulated DEGs: red). No DEGs were detected between saline and cocaine groups at WD45 (FDR p-value cutoff of < 0.05). (H) Pathway enrichment analysis displaying most significant cellular pathways between saline- and cocaine-administering rats at withdrawal day 1. Downregulated pathways shown in blue and upregulated pathways shown in red (adjusted p-value < 0.05). (I) Venn diagram showing the total number of significant DEGs detected within saline and cocaine groups across time (WD45 vs. WD1).

For experiment 2, we divided the total number of rats (n=40) into 4 groups. We assigned them to either groups A, B, C, or D. Rats began self-administration in a staggered manner, in which groups C and D started self-administration 35 days into the 45-day abstinence period of groups A and B, to align immunoprecipitation experiments of withdrawal day 45 groups (A and B) and withdrawal day 1 groups (C and D) (e.g. A and C euthanized on the same day, and B and D euthanized on the same day) (see **Fig. 3A** for timeline). Following immunoprecipitation, RNA purification, and RNA quality testing was performed in several rounds, which included a balanced representation of all treatment groups (saline-WD1, saline-WD45, cocaine-WD1, cocaine-WD45) from each timepoint. All RNA samples included in this experiment had a minimum RIN of 7. Samples were then stored at -80°C until RNA sequencing.

**Figure 3:**
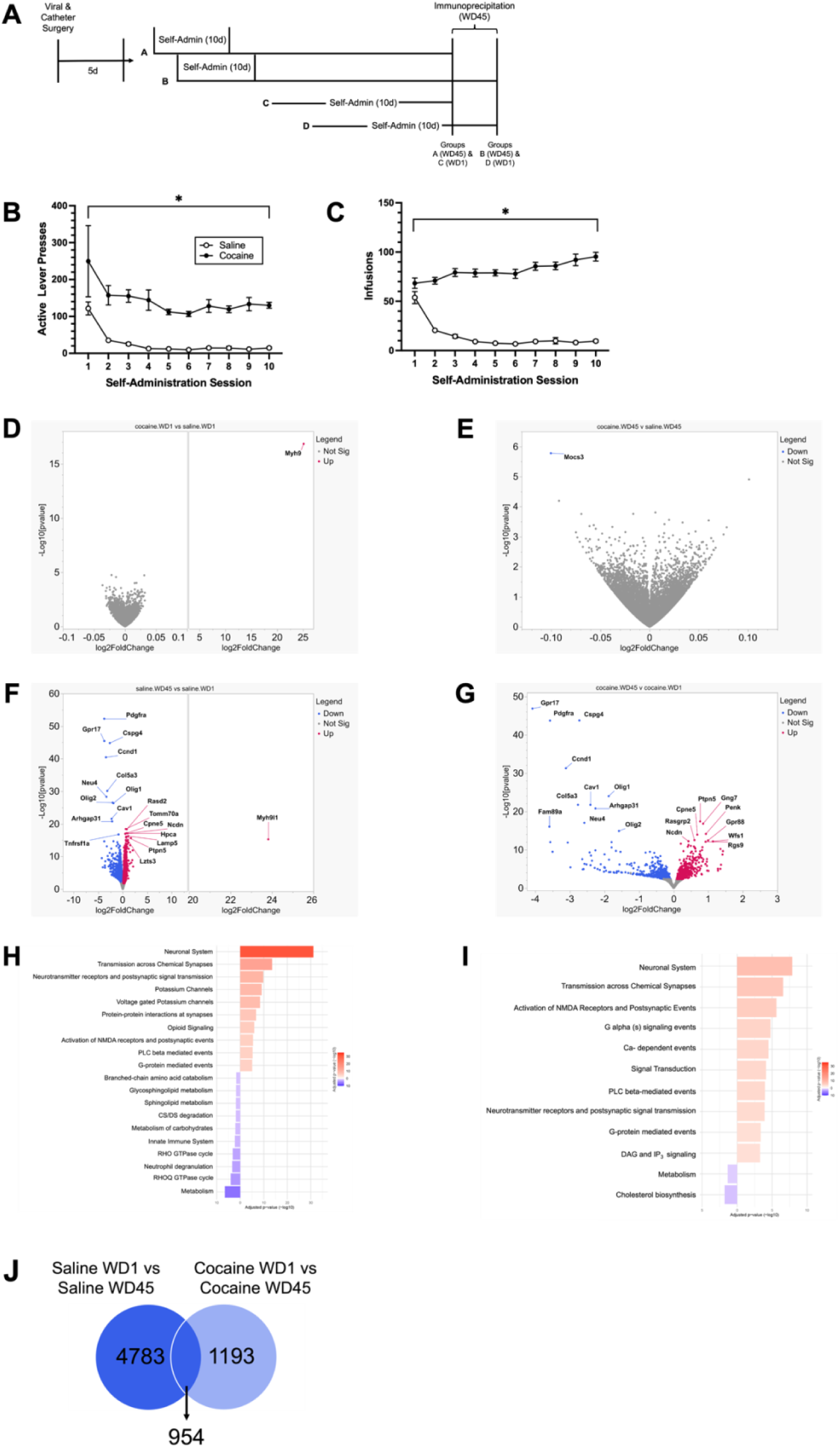
Experiment II. *Cocaine-administering rats received greater reinforcers than saline controls.* A) Experimental timeline for behavior, immunoprecipitation, and RNASeq. Rats were euthanized in a synchronized manner to align withdrawal days 1 and 45. B) Quantification of active lever presses for saline and cocaine groups across self-administration. C) Infusion counts shown for saline and cocaine groups throughout self-administration. (p < 0.05). *Differential gene expression in nucleus accumbens gene expression after saline or cocaine operant self-administration and 1 or 45 day(s) of home cage abstinence.* Volcano plots displaying the total number of significant differentially expressed genes (DEGs) in experimental group comparisons including cocaine vs. saline at withdrawal day (WD1) (D), cocaine vs. saline at WD45 (E), saline WD45 vs. saline WD1 (F), cocaine WD45 vs. cocaine WD1 (G) (nonsignificant: gray, significant downregulated DEGs: blue, and significantly upregulated DEGs: red) (FDR p-value cutoff of ≤ 0.05). Pathway enrichment analysis displaying most significant cellular pathways between saline-administering (H) and cocaine-administering (I) rats across abstinence (WD45 vs. WD1). Downregulated pathways are shown in blue and upregulated pathways shown in red (adjusted p-value < 0.05). (J) Venn diagram showing the total number of significant DEGs detected within saline and cocaine groups across time (WD45 vs. WD1).

### Library Preparation and RNA-Seq Analysis of NAc Astrocyte Transcriptomes Experiments 1 and 2

RNA from a RiboTag immunoprecipitation of nucleus accumbens astrocytes was analyzed in triplicates, where three sequences of each sample were sequenced. Libraries were prepared using Universal Plus Total RNA-Seq with NuQuant Custom AnyDeplete (cat# 30202148) according to the manufacturer’s instructions. Experiments 1 and 2 were separately sequenced at the UNC High-Throughput Sequencing Facility (HTSF) on an Illumina NovaSeq S4 with paired-end 2x100-bp reads. Reads were then filtered for adapter contamination using cutadapt (v2.9) (DOI: https://doi.org/10.14806/ej.17.1.200) and filtered using the FASTX-Toolkit (v0.0.14) (hannonlab.cshl.edu/fastx_toolkit/index.html) so that at least 90% of bases of each read had a Phred-33 score of >20. Reads were aligned to the reference genome (rn6) using STAR (v2.5.2b) by retaining only primary alignments (Dobin et al., 2013). Transcript abundance was estimated using salmon (v0.11.3) (Patro et al., 2017), and differential expression of genes was detected using DESeq2 (v1.14.1) (Love et al., 2014). Differentially expressed genes were called using lfcShrink and pairwise contrast models between all sample types.

### Statistical Analysis

For self-administration experiments, two-way ANOVA in Prism 10 was used to analyze differences in active lever presses and infusions between saline- and cocaine-administering rats. Sidak’s multiple comparisons test was used to determine if there were differences in the mean number of active lever presses and infusion counts between drug treatment groups on each day of self-administration. Data shown in self-administration figures are SEM, and significance was considered p ≤ 0.05. DESeq2 was used to detect differentially expressed genes between the following group comparisons: cocaine WD1 vs. saline WD1, saline WD45 vs. saline WD1, cocaine WD45 vs. cocaine WD1, and cocaine WD45 vs saline WD45. An FDR cutoff of 0.05 was set to isolate further and examine significantly differentially expressed genes from these experimental comparison groups. JMP® Pro, Version 15.0.0 (2019) SAS Institute Inc., Cary, NC, was used to create Volcano plots and Venn diagrams. For pathway enrichment analysis, only enriched pathways with a p-value or adjusted p-value of p < 0.05 were included, and only the top 15 upregulated and downregulated significantly enriched pathways were shown in the figures in this manuscript.

### Resource Availability

Requests for additional information and resources, including code availability should be directed to Dr. Kathryn Reissner.

## RESULTS

### AAV5 GfaABC1D-Rpl22-HA is highly specific and abundantly expressed in NAc astrocytes

Prior to beginning self-administration experiments, the specificity and expression of AAV5 GfaABC_1_D-Rpl22-HA (Rpl22-HA) in rat nucleus accumbens (NAc) astrocytes was investigated (**Fig. 1**). Figure 1B shows robust expression of Rpl22-HA in the NAc and overlapping expression with GFAP. Additionally, colocalization analyses using Bitplane Imaris software showed notable colocalization of Rpl22-HA and GFAP expression, with minimal colocalization observed with NeuN (**Fig. 1C**). As a secondary method of validation, a separate cohort of naïve male Sprague Dawley rats (n=3) received bilateral injections of Rpl22-HA into the NAc followed by immunoprecipitation and RNA-seq. Each sample used for RNA-seq validation studies shown in Fig. 1D were sequenced separately and each sample was sequenced in triplicates. The sequencing results demonstrated high levels of expression of astrocyte-specific genes (e.g. *S100b*, *Aldoc*) and nominal expression of common marker genes expressed in neurons, microglia, or oligodendrocytes (**Fig. 1D**).

*Cocaine self-administration and abstinence from cocaine induce differing degrees of differential gene expression in NAc astrocytes*

### Experiment 1: Effects of long-access cocaine self-administration on nucleus accumbens astrocyte gene expression – Part I

To identify the effects of cocaine self-administration and abstinence in the NAc, male Sprague Dawley rats were assigned to either saline (n=12) or cocaine (n=11) for long-access operant self-administration experiments (6hr/10d) (see timeline in **Fig. 2A**). As expected, cocaine-administering rats displayed greater self-administration behavior than saline controls, as the number of active lever presses (F_(1,21)_ = 42.11, p < 0.001) (**Fig. 2B**) and infusions (F_(1,21)_ = 436.2, p < 0.001) (**Fig. 2C**) were significantly increased. Two-way ANOVA showed a significant effect of time (F_(1.826, 38.35)_ = 13.24, p < 0.0001) and treatment (F_(1, 21)_ = 42.11, p < 0.0001) for active lever presses. Also, Sidak’s multiple comparisons test indicated that cocaine-administering rats received significantly more active lever presses than their saline controls on days 2 and 4-10 of self-administration (p < 0.05). Likewise, two-way ANOVA displayed a significant effect of time (F_(2.515, 52.82)_ = 7.810, p < 0.0001) and treatment (F_(1, 21)_ = 436.2, p < 0.0001) in infusions of cocaine-administering rats versus saline controls. Sidak’s multiple comparisons test showed a significant increase in infusions in cocaine-administering rats on days 2-10 of self-administration (p < 0.0001).

Following self-administration, half of the total number of rats in each treatment group, saline (n=12) and cocaine (n=11), were euthanized for immunoprecipitation and downstream RNA-seq at withdrawal day (WD) 1, and the remaining half was euthanized later on withdrawal day 45. To our surprise, differential expression analyses detected few significant differentially expressed genes (DEGs) (n=8 genes, FDR p-adjusted value <0.05) between saline- and cocaine-administering groups at WD1 (**Fig. 2D**) and detected none at WD45 (**Fig. 2E**). Of the identified significant DEGs at WD1, 7 out of 8 were downregulated by cocaine. In contrast, *Hpd* was the only upregulated DEG (**Fig. 2D**).

Despite the minimal effect on astrocyte gene expression, a robust time effect was observed within both saline and cocaine groups, as 3556 genes were identified as DEGs in saline-experienced rats and 1364 DEGs were observed in the cocaine-abstinent group (**Fig. 2I; Supplementary Data 1**). Approximately 1000 DEGs were shared between saline and cocaine groups over time (**Fig. 2I**). Pathway enrichment analysis (**Fig. 2H**) revealed that most of the pathways identified in the experimental group comparison of cocaine- and saline-administering rats at WD1 are downregulated as opposed to upregulated. The downregulated pathways identified include the NOD1/2 signaling pathway, Fc-gamma receptor-dependent phagocytosis, among others (**Fig. 2H**). Additionally, the upregulated pathways enriched in our dataset were primarily involved in metabolic processes such as tyrosine catabolism. Pathway enrichment analyses for saline (**Fig. S2A**) and cocaine (**Fig. S2A**) treated rats across abstinence are shown in the supplemental data (**Supplementary Data 5**). Principal component analysis (PCA) was used to investigate contributors to variance within these results. The PCA analysis showed that time was the largest contributor to gene expression variance as opposed to drug treatment (**Fig. S1A**).

### Experiment 2: Effects of long-access cocaine self-administration on nucleus accumbens astrocyte gene expression – Part II

Because experiment 1 largely revealed unexpected changes in astrocyte gene expression across time within each treatment group, rather than between drug/saline administration, we wished to repeat the study with a modified timeline in order to mitigate any possible confounds resulting from batch effects of time. To do this, we performed a subsequent experiment with identical experimental groups but a distinct temporal organization of self-administration and immunoprecipitation procedures. Rats were divided into four groups (n=10 per group) and assigned either withdrawal day 45 (groups A and B) or withdrawal day 1 (groups C and D). Immunoprecipitation of these groups was conducted such that each withdrawal day 1 group aligned with a withdrawal day 45 group (ex: group A and C euthanized on the same day). The experimental timeline for experiment 2 is shown in **Fig. 3A**. The surgical and self-administration procedures were identical to those for Experiment 1.

Two-way ANOVA showed that cocaine-administering rats had significantly higher active lever presses (F_(1, 29)_ = 57.46, p < 0.0001) (**Fig. 3B**) and infusions (F_(1, 29)_ = 805.3, p < 0.0001) (**Fig. 3C**) compared to the saline control group. Particularly, a significant effect of time (F_(9, 261)_ = 5.403, p < 0.0001) and treatment (F_(1,29)_ = 57.46, p < 0.0001) were observed. There was also a significant effect of time (F_(9, 261)_ = 5.958, p < 0.0001) and treatment (F_(1, 29)_ = 805.3, p < 0.0001) in number of infusions. Using Sidak’s multiple comparisons test, statistics showed that cocaine-administering had significantly more active lever presses and infusions than their saline controls on all 10 days of self-administration (p < 0.05).

We then compared gene expression profiles derived rat NAc astrocytes from saline- and cocaine-administering and found just one significant gene (padj < 0.05), *Myh9*, that was upregulated at withdrawal day 1 (**Fig. 3D**). Similarly, there was only one gene, *Mocs3*, which was downregulated in comparison to saline-WD45 to cocaine-WD45 (**Fig. 3E**). Nonetheless, both within drug group comparisons (saline and cocaine) across abstinence (WD1 vs. WD45) showed similar effects of time as reflected by an extensive number of genes identified between withdrawal days 1 and 45, as approximately 4700 and 1100 genes were detected in each group (**Supplementary Data 2)**, respectively. In the same experimental comparison of saline-administering rats across abstinence, pathway enrichment analysis identified canonical pathways as significantly upregulated, including *G-protein mediated events* and *Voltage-gated Potassium Channels*. In contrast, downregulated pathways included the *Rho GTPase Cycle,* and several pathways associated with metabolism regulation (**Fig. 3H and Supplementary Data 6**). Pathway enrichment detected the *Cholesterol Biosynthesis* pathway as downregulated in the cocaine treatment group across withdrawal, and conversely, more enriched pathways in the experimental comparison of cocaine treated rats across abstinence were upregulated, including *Calcium-Dependent Events* and *DAG and IP^3^ Signaling*, of which were also upregulated in saline controls across abstinence (**Fig. 3I and Supplementary Data 6**). Like experiment 1, there were more genes detected as significantly differentially expressed in the saline group across abstinence (WD45 vs. WD1) than in the cocaine-administering group (**Fig. 3J**). Additionally, there were approximately 1000 DEGs shared within comparisons of the same drug treatment (saline and cocaine) across time in both experiments 1 and 2 (**Figs. 2I and 3J; Supplementary Data 4**), further attesting to the reproducibility and validity of the RiboTag AAV method in astrocytes. Additionally, PCA analysis showed time as a greater variance contributor than drug treatment (**Fig. S1B**), as found in experiment 1, which further demonstrated the reproducibility and validity of these results.

### Cocaine self-administration uniquely affects diverse genes and cellular processes within the central nervous system

We next performed Pathway Enrichment Analyses to provide biological context and assess more broadly the effects of chronic cocaine exposure and abstinence on astrocytes. In both experiments, 1 and 2, pathway enrichment analyses pinpointed distinct cellular pathways that were only detected in cocaine- (but not saline) administering rats across abstinence (WD45 vs. WD1). In experiments 1 and 2, we identified 31 and 7 distinct pathways that changed over time, specifically in cocaine-administering groups, respectively. However, only 4 of these pathways were enriched in both cocaine-administering comparisons from both experiments 1 and 2, which included the *Cholesterol Biosynthesis* pathway (**Supplementary Data 5 and 6**). In our studies, the *Cholesterol Biosynthesis* pathway was significantly downregulated in cocaine-assigned rats in late abstinence (WD45) in comparison to early abstinence (WD1).

Alterations in cholesterol production and metabolism processes have been observed in other studies involving cocaine exposure and relapse in both rodents and humans (Buydens-Branchey and Branchey, 2003 and Ahooyi et al., 2018). However, to our knowledge, investigations into astrocyte involvement in modified cholesterol synthesis and regulation by cocaine have not been explored. Further, *Dhcr7*, a gene involved in cholesterol production (Suzuki et al., 2020), was downregulated in our cocaine WD45 group compared to cocaine WD1. Also, several other genes involved in cholesterol production were downregulated in the cocaine WD45 group, including *Sqle*, *Hmgcs1*, and *Fdf1*. These results further support that cocaine exposure affects cholesterol production processes and suggest that astrocyte-mediated regulation of cholesterol biosynthesis is further diminished by cocaine withdrawal.

## DISCUSSION

Akin to previous studies, results from the current study showed that RiboTag can be used as an effective method to isolate astrocyte-specific mRNAs for subsequent RNA-seq (Boisvert et al., 2018; Yu et al., 2021; Diaz-Castro et al., 2021), which can be used to elucidate cell-type specific transcriptional and translational changes. We hypothesized that cocaine-administering rats would display significant numbers of differentially expressed genes in comparison to saline at both withdrawal timepoints, day 1 and 45. However, our results detected few significant genes at both withdrawal timepoints between drug treatment groups, saline, and cocaine. In contrast, both experiments 1 and 2 reliably displayed robust differential gene expression within drug treatment groups across abstinence. However, treatment-specific pathway analysis across time yields insights into candidate processes and genes for further investigation.

### Minimal differential gene expression detected in NAc astrocytes of cocaine-administering rats

Published work from our lab and others have shown that NAc astrocytes display altered morphology, specifically decreased surface area, volume, and synaptic colocalization, following cocaine self-administration and extinction or protracted abstinence (Scofield et al., 2016; Testen et al., 2018; Kim et al., 2022). Due to these observed structural changes, we hypothesized that progressive differential gene expression would be observed in cocaine-abstinent rats in comparison to saline controls. Surprisingly, DESeq2 analyses did not detect any significant DEGs in experiment 1 in comparisons between saline and cocaine groups at WD45, and experiment 2 detected a single DEG in saline and cocaine comparisons at both WD1 and WD45.

Although relatively few published studies have employed RiboTag immunoprecipitation to investigate astrocyte gene expression changes, other studies have detected significant DEGs in astrocytes between experimental groups (Yu et al., 2020; Diaz-Castro et al., 2021; Bravo-Ferrer et al., 2022). This may be attributed to technical considerations of the RiboTag method and differences in experimental paradigms. In both experiments, 1 and 2, minimal detection of significant DEGs could be caused by technical features of RiboTag as it captures actively translating mRNAs by immunoprecipitating mRNAs that are physically attached to the ribosomal subunit, Rpl22, at the time of euthanasia. Despite prior studies including a different form of external stimuli (e.g., LPS-induced neuroinflammation or calcium signaling manipulation), the main difference between these studies and the current study is the amount of time elapsed between the stimulus and euthanasia for immunoprecipitation. For example, Yu et al. (Yu et al., 2020) used RiboTag to investigate the effects of varied experimental perturbations that evoke behaviors associated with neuropsychiatric and neurodegenerative disorders, on striatal astrocyte gene expression. In experimental manipulations that directly targeted astrocytes, the mice were euthanized 2-6 hours following the stimulus, and approximately 1100-3400 significant DEGs were detected between control and experimental conditions.

Boisvert et al. (Boisvert et al., 2018) have provided a developmental and aging transcriptome database from four brain regions (visual cortex, motor cortex, cerebellum, and hypothalamus) from Rpl22-HA transgenic mice. Of relevance, time points of a searchable database include P7, P14, P28, and P128. Rats used in the current study were approximately 4 months old at the time of euthanasia. While these studies are in mice versus rats, comparing differentially changed pathways or genes specific to cocaine abstinence in the current studies can be compared with developmental changes at P128 in the Boisvert searchable database. For example, while *P2ry1* is not significantly different between saline and cocaine at either withdrawal daytime point, it’s consistent and selective upregulation across cocaine (but not saline abstinence), combined with the observation that it is not developmentally upregulated at P128 compared to P28, yield it an enticing candidate for further investigation.

In the context of cocaine, Campbell et al. (Campbell et al., 2021) used bulk RNAseq to investigate the effects of varied cocaine administration paradigms on gene expression in the ventral tegmental area (VTA) and detected 389 significant DEGs between cocaine and saline mice following intravenous cocaine self-administration. In this study, tissue for RNA extraction and sequencing was collected 1 hour following the last self-administration session. Thus, the withdrawal day 1 timepoint used in the current study may surpass the point at which robust translational changes are occurring in NAc astrocytes as a result of cocaine. This hypothesis is supported by a study conducted by Walker et al. (Walker et al., 2018) in which the lowest amount of significant DEGs in the NAc were detected in saline vs. cocaine self-administering rats after 24 hours of home cage abstinence in comparison to experimental conditions that included a challenge dose of saline or cocaine followed by context re-exposure 1 hour before tissue collection. Although a direct comparison between our current study and the aforementioned investigations cannot be made, several postulations regarding the effect of the euthanasia timepoint on the degree of differentially expressed genes can be inferred. In addition to experimental influences, it is plausible that technical factors regarding the RiboTag method may not comprehensively encapsulate stimuli-induced translational and ribosomal dynamics in astrocytes.

As previously discussed in the introduction, astrocytes are primarily comprised of peripheral astrocyte processes (PAPs), which are the fine, membranous processes that ensheathe synaptic elements. Few studies have explored translational dynamics within astrocytes, but Mazaré et al. (Mazaré et al., 2020) demonstrated that there are intra-cellular differences in translational activity within the PAPs compared to the entire astrocyte in the dorsal hippocampus using transgenic Aldh1l1-eGFP/Rpl10a mice for characterization of ribosome-bound mRNAs. Using fluorescence *in situ* hybridization, this study found that mRNAs for glutamate transporter 1 (GLT-1), which is abundantly expressed in astrocytes, were almost exclusively found in the PAPs of dorsal hippocampus astrocytes (Mazaré et al., 2020). Further, published studies from our lab by Kim et al. (2018) showed that GLT-1 is significantly decreased in the NAc following cocaine self-administration and 45 days of home cage abstinence (Kim et al., 2018). However, *Slc1a2* (GLT-1) was not detected as a significant DEG in our experimental comparison of saline and cocaine at withdrawal day 45. This may be a result of the technical features of RiboTag, which provides a snapshot of gene expression at a given time by only “capturing” mRNAs that are bound to ribosomes, which suggests an increased likelihood of the gene being actively translated into protein (Sanz et al., 2019). There is also evidence that RiboTag provides less coverage than other methods for quantifying gene expression (Kronman et al., 2019).

Moreover, studies have shown no direct correlation between the mRNA and protein levels (Koussounadis et al., 2015; Edfors et al., 2016; Wegler et al., 2019). Therefore, at the 45-day timepoint, it is possible that GLT-1 is not being actively translated or ribosome-bound, thus leading to an inability to detect significant changes in its expression in NAc astrocytes of cocaine-abstinent rats. Further information on translational dynamics within astrocytes would be beneficial in elucidating these observations.

Unexpectedly, greater levels of significant DEGs were detected within the same drug treatment groups (saline and cocaine) across abstinence (WD45 vs. WD1). Experiments 1 and 2 identified similar differential gene expression patterns, as upwards of 3000 significant DEGs were detected in saline-administering groups over time. In contrast, approximately 1100-1400 DEGs were identified in cocaine-administering rats. A few hypotheses can be made regarding this pattern of differential gene expression, including 1) a more significant influence of aging may be occurring and increasing translational activity in late abstinence rats as opposed to early abstinence rats, 2) a lesser degree of significant DEGs in cocaine-abstinent rats may occur due to cocaine-induced influences on post-translational modifications which can cause gene repression, and 3) extended home cage abstinence may contribute to differential gene expression as it may evoke anxiety-like effects on gene expression. Ultimately, further investigation into the contribution of these factors is suggested, including categorizing genes potentially enriched in either of these scenarios.

### Prior history of cocaine exposure may exacerbate biological aging

In the current study, the *Cholesterol Biosynthesis* pathway was downregulated in only the cocaine group across abstinence in both experiments 1 and 2, suggesting that this is a cocaine-specific effect and reproducible finding. Also, various genes involved in cholesterol synthesis were detected as downregulated DEGs in cocaine-abstinent rats, such as *Dhcr7* and *Hmgcs1*. Our findings further support prior literature that has demonstrated astrocytes as the primary synthesizer and their role in cholesterol metabolism in the brain (Nieweg et al., 2009; Jin et al., 2019). Additionally, previous studies have shown that cholesterol levels are affected by cocaine. Buydens-Branchey et al. and others showed that individuals with a history of cocaine use display lower levels of blood cholesterol and that cocaine exposure causes alterations in lipid metabolism in the brain. Astrocyte-mediated cholesterol synthesis is important for glutamatergic synaptic transmission, as cholesterol depletion in hippocampal neurons has been exhibited in impaired synaptic transmission in various studies (Chen et al., 2023; Shin et al., 2024). Although both experiments in the current study suggest a greater effect of time as opposed to drug treatment, the cocaine-specific finding of the downregulated *Cholesterol Biosynthesis* pathway indicates that there may be a connection between cocaine exposure and premature aging of cellular pathways.

Previous literature has explored the connection between cholesterol synthesis and aging in healthy and diseased instances. For example, deficits in cholesterol production and metabolism have been discovered in clinical studies of Alzheimer’s Disease (AD) (Ahmed et al., 2024). Cholesterol levels are highest during development but decrease throughout normal aging (Thelen et al., 2006). In studies focused on AD, increased and reduced cholesterol levels have been observed (Feringa et al., 2021). Further, Boisvert et al. also identified the *Cholesterol Biosynthesis* pathway as downregulated in astrocytes of various brain regions in mice at 2 years of age compared to mice at 4 months (Boisvert et al., 2018). Our data show that this pathway is significantly downregulated at approximately 4 months of age in cocaine-abstinent rats who previously underwent cocaine self-administration 45 days prior to euthanasia, which suggests that a history of cocaine may accelerate biological aging by altering critical cellular processes involved in the maintenance the physiological state of the CNS. This hypothesis is supported by previously published literature that showed that cocaine addiction ages the human brain (Beheshti, 2023) by measuring DNA methylation (Poisel et al., 2023) levels to determine the biological aging of cells in individuals with cocaine use disorder. Few studies have investigated either relationship between cholesterol metabolism and cocaine (Buydens-Branchey et al., 2003) and also astrocytes and cholesterol (Pfreiger and Uneger, 2011; Ferris et al., 2017), but the current study suggests that there is a tri-relationship between these three factors. Taken together, our results indicate that cocaine-abstinent rats displayed premature aging by exhibiting alterations in important cellular processes that have been identified in pathophysiological aging investigations. To our knowledge, our study is the first to provide evidence that proposes a tri-part relationship of astrocyte involvement in the connection between cocaine use and aging. Nonetheless, additional studies into how astrocyte regulation of cholesterol metabolism is modified by cocaine exposure and how these changes may influence cocaine-seeking behavior would be beneficial.

### Conclusions and Future Directions

Results obtained from the current study advance our knowledge of how cocaine self-administration and abstinence affect gene expression in nucleus accumbens astrocytes. These results suggested that cocaine self-administration evoked minimal changes in NAc astrocyte gene expression early in abstinence but that cocaine-abstinent rats displayed substantially greater significant DEGs, suggesting that a history of cocaine may have delayed influences on NAc astrocyte gene expression. Further, these results may pinpoint gene expression differences that underly cocaine-induced astrocyte adaptations that promote relapse behaviors. In addition, important cellular pathways that are required for the normal functioning of neurons and the brain were identified as downregulated in cocaine-administering rats at late abstinence, suggesting that cocaine modulates NAc astrocyte functional regulation of diverse cellular pathways and the CNS, which is prompted by cocaine self-administration and withdrawal. Overall, the results described in the current study progress our knowledge of astrocyte functional mechanisms in the diseased brain and may also help elucidate astrocyte function in the healthy brain. Future studies will investigate changes in NAc astrocyte gene expression in rats with a history of cocaine after undergoing context re-exposure to discern if there are distinct dysregulated genes that contribute to various stages of the addiction cycle. Additionally, as Kim et al. did not observe morphological alterations in NAc astrocytes of females following cocaine self-administration and 45 days of abstinence, future studies will include female rats to investigate underlying genetics of potential protective effects of NAc astrocytes in female rats.

## Supporting information

Exp 1_significant DEGs_supplementary data

Exp 2_significant DEGs_supplementary data

Experiment 1_Common and Group Specific Significant DEGs

Experiment 2_Common and Group Specific Significant DEGs

Experiment 1_Pathway Enrichment Analysis

Experiment 2_Pathway Enrichment Analysis

## Acknowledgments

This work was supported by NIDA 1R21DA052447 (KJR), NIDA 1R01DA041455 (KJR), and NIDA 1F31DA057113-01A1 (JPF)

## References

Adermark, L., & Bowers, M. S. (2016). Disentangling the Role of Astrocytes in Alcohol Use Disorder. Alcoholism, clinical and experimental research, 40(9), 1802–1816.

Ahmed, H., Wang, Y., Griffiths, W. J., Levey, A. I., Pikuleva, I., Liang, S. H., & Haider, A. (2024). Brain cholesterol and Alzheimer’s disease: challenges and opportunities in probe and drug development. Brain : a journal of neurology, 147(5), 1622–1635.

Araque, A., & Perea, G. (2004). Glial modulation of synaptic transmission in culture. Glia, 47(3), 241–248.

Batiuk, M. Y., Martirosyan, A., Wahis, J., de Vin, F., Marneffe, C., Kusserow, C., Koeppen, J., Viana, J. F., Oliveira, J. F., Voet, T., Ponting, C. P., Belgard, T. G., & Holt, M. G. (2020). Identification of region-specific astrocyte subtypes at single cell resolution. Nature communications, 11(1), 1220.

Beheshti I. (2023). Cocaine Destroys Gray Matter Brain Cells and Accelerates Brain Aging. Biology, 12(5), 752.

Boisvert, M. M., Erikson, G. A., Shokhirev, M. N., & Allen, N. J. (2018). The Aging Astrocyte Transcriptome from Multiple Regions of the Mouse Brain. Cell reports, 22(1), 269–285.

Bravo-Ferrer, I., Khakh, B. S., & Díaz-Castro, B. (2022). Cell-specific RNA purification to study translatomes of mouse central nervous system. STAR protocols, 3(2), 101397.

Burda, J. E., Bernstein, A. M., & Sofroniew, M. V. (2016). Astrocyte roles in traumatic brain injury. Experimental neurology, 275 *Pt 3*(0 3), 305–315.

Buydens-Branchey, L., & Branchey, M. (2003). Association between low plasma levels of cholesterol and relapse in cocaine addicts. Psychosomatic medicine, 65(1), 86–91.

Cabezas, R., Avila, M., Gonzalez, J., El-Bachá, R. S., Báez, E., García-Segura, L. M., Jurado Coronel, J. C., Capani, F., Cardona-Gomez, G. P., & Barreto, G. E. (2014). Astrocytic modulation of blood brain barrier: perspectives on Parkinson’s disease. Frontiers in cellular neuroscience, 8, 211.

Campbell, R. R., Chen, S., Beardwood, J. H., López, A. J., Pham, L. V., Keiser, A. M., Childs, J. E., Matheos, D. P., Swarup, V., Baldi, P., & Wood, M. A. (2021). Cocaine induces paradigm-specific changes to the transcriptome within the ventral tegmental area. Neuropsychopharmacology : official publication of the American College of Neuropsychopharmacology, 46(10), 1768–1779.

Caracciolo, L., Marosi, M., Mazzitelli, J., Latifi, S., Sano, Y., Galvan, L., Kawaguchi, R., Holley, S., Levine, M. S., Coppola, G., Portera-Cailliau, C., Silva, A. J., & Carmichael, S. T. (2018). CREB controls cortical circuit plasticity and functional recovery after stroke. Nature communications, 9(1), 2250.

Carlezon, W. A., Jr, Thome, J., Olson, V. G., Lane-Ladd, S. B., Brodkin, E. S., Hiroi, N., Duman, R. S., Neve, R. L., & Nestler, E. J. (1998). Regulation of cocaine reward by CREB. *Science (New York*, N.Y.), 282(5397), 2272–2275.

Chen, Z., Yuan, Z., Yang, S., Zhu, Y., Xue, M., Zhang, J., & Leng, L. (2023). Brain Energy Metabolism: Astrocytes in Neurodegenerative Diseases. CNS neuroscience & therapeutics, 29(1), 24–36.

Cho, Y., & Bannai, S. (1990). Uptake of glutamate and cysteine in C-6 glioma cells and in cultured astrocytes. Journal of neurochemistry, 55(6), 2091–2097.

Chung, W. S., Allen, N. J., & Eroglu, C. (2015). Astrocytes Control Synapse Formation, Function, and Elimination. Cold Spring Harbor perspectives in biology, 7(9), a020370.

Clarke, T. K., Crist, R. C., Kampman, K. M., Dackis, C. A., Pettinati, H. M., O’Brien, C. P., Oslin, D. W., Ferraro, T. N., Lohoff, F. W., & Berrettini, W. H. (2013). Low frequency genetic variants in the μ-opioid receptor (OPRM1) affect risk for addiction to heroin and cocaine. Neuroscience letters, 542, 71–75.

Corkrum, M., Rothwell, P. E., Thomas, M. J., Kofuji, P., & Araque, A. (2019). Opioid-Mediated Astrocyte-Neuron Signaling in the Nucleus Accumbens. Cells, 8(6), 586.

Corkrum, M., Covelo, A., Lines, J., Bellocchio, L., Pisansky, M., Loke, K., Quintana, R., Rothwell, P. E., Lujan, R., Marsicano, G., Martin, E. D., Thomas, M. J., Kofuji, P., & Araque, A. (2020). Dopamine-Evoked Synaptic Regulation in the Nucleus Accumbens Requires Astrocyte Activity. Neuron, 105(6), 1036–1047.e5.

Cotto, B., Natarajaseenivasan, K., Ferrero, K., Wesley, L., Sayre, M., & Langford, D. (2018). Cocaine and HIV-1 Tat disrupt cholesterol homeostasis in astrocytes: Implications for HIV-associated neurocognitive disorders in cocaine user patients. Glia, 66(4), 889–902.

Dance, A., Fernandes, J., Toussaint, B., Vaillant, E., Boutry, R., Baron, M., Loiselle, H., Balkau, B., Charpentier, G., Franc, S., Ibberson, M., Marre, M., Gernay, M., Fadeur, M., Paquot, N., Vaxillaire, M., Boissel, M., Amanzougarene, S., Derhourhi, M., Khamis, A., … Bonnefond, A. (2024). Exploring the role of purinergic receptor P2RY1 in type 2 diabetes risk and pathophysiology: Insights from human functional genomics. Molecular metabolism, 79, 101867.

Dhaliwal A, Gupta M. Physiology, Opioid Receptor. [Updated 2023 Jul 24]. In: StatPearls [Internet]. Treasure Island (FL): StatPearls Publishing; 2024 Jan-.

Derouiche, A., & Frotscher, M. (2001). Peripheral astrocyte processes: monitoring by selective immunostaining for the actin-binding ERM proteins. Glia, 36(3), 330–341.

Diaz-Castro, B., Gangwani, M. R., Yu, X., Coppola, G., & Khakh, B. S. (2019). Astrocyte molecular signatures in Huntington’s disease. Science translational medicine, 11(514), eaaw8546.

Diaz-Castro, B., Bernstein, A. M., Coppola, G., Sofroniew, M. V., & Khakh, B. S. (2021). Molecular and functional properties of cortical astrocytes during peripherally induced neuroinflammation. Cell reports, 36(6), 109508.

Dobin, A., Davis, C. A., Schlesinger, F., Drenkow, J., Zaleski, C., Jha, S., Batut, P., Chaisson, M., & Gingeras, T. R. (2013). STAR: ultrafast universal RNA-seq aligner. *Bioinformatics (Oxford*, England), 29(1), 15–21.

Edfors, F., Danielsson, F., Hallström, B. M., Käll, L., Lundberg, E., Pontén, F., Forsström, B., & Uhlén, M. (2016). Gene-specific correlation of RNA and protein levels in human cells and tissues. Molecular systems biology, 12(10), 883.

Ferris, H. A., Perry, R. J., Moreira, G. V., Shulman, G. I., Horton, J. D., & Kahn, C. R. (2017). Loss of astrocyte cholesterol synthesis disrupts neuronal function and alters whole-body metabolism. Proceedings of the National Academy of Sciences of the United States of America, 114(5), 1189–1194.

Garrido, E., Pérez-García, C., Alguacil, L. F., & Díez-Fernández, C. (2005). The alpha2-adrenoceptor antagonist yohimbine reduces glial fibrillary acidic protein upregulation induced by chronic morphine administration. Neuroscience letters, 383(1-2), 141–144.

Goenaga, J., Araque, A., Kofuji, P., & Herrera Moro Chao, D. (2023). Calcium signaling in astrocytes and gliotransmitter release. Frontiers in synaptic neuroscience, 15, 1138577.

Gorelick, D. A., Kim, Y. K., Bencherif, B., Boyd, S. J., Nelson, R., Copersino, M., Endres, C. J., Dannals, R. F., & Frost, J. J. (2005). Imaging brain mu-opioid receptors in abstinent cocaine users: time course and relation to cocaine craving. Biological psychiatry, 57(12), 1573–1582.

Guebel D. V. (2023). Human hippocampal astrocytes: Computational dissection of their transcriptome, sexual differences and exosomes across ageing and mild-cognitive impairment. The European journal of neuroscience, 58(3), 2677–2707.

Guerra-Gomes, S., Sousa, N., Pinto, L., & Oliveira, J. F. (2018). Functional Roles of Astrocyte Calcium Elevations: From Synapses to Behavior. Frontiers in cellular neuroscience, 11, 427.

Fellin, T., Pozzan, T., & Carmignoto, G. (2006). Purinergic receptors mediate two distinct glutamate release pathways in hippocampal astrocytes. The Journal of biological chemistry, 281(7), 4274–4284.

Fernàndez-Castillo, N., Cabana-Domínguez, J., Corominas, R., & Cormand, B. (2022). Molecular genetics of cocaine use disorders in humans. Molecular psychiatry, 27(1), 624–639.

Feringa, F. M., & van der Kant, R. (2021). Cholesterol and Alzheimer’s Disease; From Risk Genes to Pathological Effects. Frontiers in aging neuroscience, 13, 690372.

Kim, H. B., Lu, Y., Oh, S. C., Morris, J., Miyashiro, K., Kim, J., Eberwine, J., & Sul, J. Y. (2022). Astrocyte ethanol exposure reveals persistent and defined calcium response subtypes and associated gene signatures. The Journal of biological chemistry, 298(8), 102147.

Kim, R., Sepulveda-Orengo, M. T., Healey, K. L., Williams, E. A., & Reissner, K. J. (2018). Regulation of glutamate transporter 1 (GLT-1) gene expression by cocaine self-administration and withdrawal. Neuropharmacology, 128, 1–10.

Kim, R., Testen, A., Harder, E. V., Brown, N. E., Witt, E. A., Bellinger, T. J., Franklin, J. P., & Reissner, K. J. (2022). Abstinence-Dependent Effects of Long-Access Cocaine Self-Administration on Nucleus Accumbens Astrocytes Are Observed in Male, But Not Female, Rats. eNeuro, 9(5), ENEURO.0310-22.2022.

Koussounadis, A., Langdon, S. P., Um, I. H., Harrison, D. J., & Smith, V. A. (2015). Relationship between differentially expressed mRNA and mRNA-protein correlations in a xenograft model system. Scientific reports, 5, 10775.

Kronman, H., Richter, F., Labonté, B., Chandra, R., Zhao, S., Hoffman, G., Lobo, M. K., Schadt, E. E., & Nestler, E. J. (2019). Biology and Bias in Cell Type-Specific RNAseq of Nucleus Accumbens Medium Spiny Neurons. Scientific reports, 9(1), 8350.

Jin, J., Daniel, J. L., & Kunapuli, S. P. (1998). Molecular basis for ADP-induced platelet activation. II. The P2Y1 receptor mediates ADP-induced intracellular calcium mobilization and shape change in platelets. The Journal of biological chemistry, 273(4), 2030–2034.

Jin, U., Park, S. J., & Park, S. M. (2019). Cholesterol Metabolism in the Brain and Its Association with Parkinson’s Disease. Experimental neurobiology, 28(5), 554–567.

Li, D., Agulhon, C., Schmidt, E., Oheim, M., & Ropert, N. (2013). New tools for investigating astrocyte-to-neuron communication. Frontiers in cellular neuroscience, 7, 193.

Liu, X., Ying, J., Wang, X., Zheng, Q., Zhao, T., Yoon, S., Yu, W., Yang, D., Fang, Y., & Hua, F. (2021). Astrocytes in Neural Circuits: Key Factors in Synaptic Regulation and Potential Targets for Neurodevelopmental Disorders. Frontiers in molecular neuroscience, 14, 729273.

Love, M. I., Huber, W., & Anders, S. (2014). Moderated estimation of fold change and dispersion for RNA-seq data with DESeq2. Genome biology, 15(12), 550.

Mahmoud, S., Gharagozloo, M., Simard, C., & Gris, D. (2019). Astrocytes Maintain Glutamate Homeostasis in the CNS by Controlling the Balance between Glutamate Uptake and Release. Cells, 8(2), 184.

Mazaré, N., Oudart, M., Moulard, J., Cheung, G., Tortuyaux, R., Mailly, P., Mazaud, D., Bemelmans, A. P., Boulay, A. C., Blugeon, C., Jourdren, L., Le Crom, S., Rouach, N., & Cohen-Salmon, M. (2020). Local Translation in Perisynaptic Astrocytic Processes Is Specific and Changes after Fear Conditioning. Cell reports, 32(8), 108076.

McClung, C. A., & Nestler, E. J. (2003). Regulation of gene expression and cocaine reward by CREB and DeltaFosB. Nature neuroscience, 6(11), 1208–1215.

Mews, P., Cunningham, A. M., Scarpa, J., Ramakrishnan, A., Hicks, E. M., Bolnick, S., Garamszegi, S., Shen, L., Mash, D. C., & Nestler, E. J. (2023). Convergent abnormalities in striatal gene networks in human cocaine use disorder and mouse cocaine administration models. Science advances, 9(6), eadd8946.

Mohseni Ahooyi, T., Shekarabi, M., Torkzaban, B., Langford, T. D., Burdo, T. H., Gordon, J., Datta, P. K., Amini, S., & Khalili, K. (2018). Dysregulation of Neuronal Cholesterol Homeostasis upon Exposure to HIV-1 Tat and Cocaine Revealed by RNA-Sequencing. Scientific reports, 8(1), 16300.

Namba, M. D., Kupchik, Y. M., Spencer, S. M., Garcia-Keller, C., Goenaga, J. G., Powell, G. L., Vicino, I. A., Hogue, I. B., & Gipson, C. D. (2020). Accumbens neuroimmune signaling and dysregulation of astrocytic glutamate transport underlie conditioned nicotine-seeking behavior. Addiction biology, 25(5), e12797.

Newell-Litwa, K. A., Horwitz, R., & Lamers, M. L. (2015). Non-muscle myosin II in disease: mechanisms and therapeutic opportunities. Disease models & mechanisms, 8(12), 1495–1515.

Nieweg, K., Schaller, H., & Pfrieger, F. W. (2009). Marked differences in cholesterol synthesis between neurons and glial cells from postnatal rats. Journal of neurochemistry, 109(1), 125–134.

O’Donovan, B., Neugornet, A., Neogi, R., Xia, M., & Ortinski, P. (2021). Cocaine experience induces functional adaptations in astrocytes: Implications for synaptic plasticity in the nucleus accumbens shell. Addiction biology, 26(6), e13042.

Ota, Y., Zanetti, A. T., & Hallock, R. M. (2013). The role of astrocytes in the regulation of synaptic plasticity and memory formation. Neural plasticity, 2013, 185463.

Patro, R., Duggal, G., Love, M. I., Irizarry, R. A., & Kingsford, C. (2017). Salmon provides fast and bias-aware quantification of transcript expression. Nature methods, 14(4), 417–419.

Peterson, A. R., & Binder, D. K. (2019). Post-translational Regulation of GLT-1 in Neurological Diseases and Its Potential as an Effective Therapeutic Target. Frontiers in molecular neuroscience, 12, 164.

Pfrieger, F. W., & Ungerer, N. (2011). Cholesterol metabolism in neurons and astrocytes. Progress in lipid research, 50(4), 357–371.

Poisel, E., Zillich, L., Streit, F., Frank, J., Friske, M. M., Foo, J. C., Mechawar, N., Turecki, G., Hansson, A. C., Nöthen, M. M., Rietschel, M., Spanagel, R., & Witt, S. H. (2023). DNA methylation in cocaine use disorder-An epigenome-wide approach in the human prefrontal cortex. Frontiers in psychiatry, 14, 1075250.

Sakamoto, K., Karelina, K., & Obrietan, K. (2011). CREB: a multifaceted regulator of neuronal plasticity and protection. Journal of neurochemistry, 116(1), 1–9. 10.1111/j.1471-4159.2010.07080.x

Sanz, E., Bean, J. C., Carey, D. P., Quintana, A., & McKnight, G. S. (2019). RiboTag: Ribosomal Tagging Strategy to Analyze Cell-Type-Specific mRNA Expression In Vivo. Current protocols in neuroscience, 88(1), e77.

Savell, K. E., Tuscher, J. J., Zipperly, M. E., Duke, C. G., Phillips, R. A., 3rd, Bauman, A. J., Thukral, S., Sultan, F. A., Goska, N. A., Ianov, L., & Day, J. J. (2020). A dopamine-induced gene expression signature regulates neuronal function and cocaine response. Science advances, 6(26), eaba4221.

Scofield, M. D., Li, H., Siemsen, B. M., Healey, K. L., Tran, P. K., Woronoff, N., Boger, H. A., Kalivas, P. W., & Reissner, K. J. (2016). Cocaine Self-Administration and Extinction Leads to Reduced Glial Fibrillary Acidic Protein Expression and Morphometric Features of Astrocytes in the Nucleus Accumbens Core. Biological psychiatry, 80(3), 207–215.

Shin, K.C., Ali Moussa, H.Y. & Park, Y. (2024). Cholesterol imbalance and neurotransmission defects in neurodegeneration. Exp Mol Med.

Siemsen, B. M., Denton, A. R., Parrila-Carrero, J., Hooker, K. N., Carpenter, E. A., Prescot, M. E., Brock, A. G., Westphal, A. M., Leath, M. N., McFaddin, J. A., Jhou, T. C., McGinty, J. F., & Scofield, M. D. (2023). Heroin Self-Administration and Extinction Increase Prelimbic Cortical Astrocyte-Synapse Proximity and Alter Dendritic Spine Morphometrics That Are Reversed by N-Acetylcysteine. Cells, 12(14), 1812.

Song, P., & Zhao, Z. Q. (2001). The involvement of glial cells in the development of morphine tolerance. Neuroscience research, 39(3), 281–286.

Spurgat, M. S., & Tang, S. J. (2022). Single-Cell RNA-Sequencing: Astrocyte and Microglial Heterogeneity in Health and Disease. Cells, 11(13), 2021.

Suzuki, A., Ogata, K., Yoshioka, H., Shim, J., Wassif, C. A., Porter, F. D., & Iwata, J. (2020). Disruption of *Dhcr7* and *Insig1/2* in cholesterol metabolism causes defects in bone formation and homeostasis through primary cilium formation. Bone research, 8, 1.

Testen, A., Sepulveda-Orengo, M. T., Gaines, C. H., & Reissner, K. J. (2018). Region-Specific Reductions in Morphometric Properties and Synaptic Colocalization of Astrocytes Following Cocaine Self-Administration and Extinction. Frontiers in cellular neuroscience, 12, 246.

Thelen, K. M., Falkai, P., Bayer, T. A., & Lütjohann, D. (2006). Cholesterol synthesis rate in human hippocampus declines with aging. Neuroscience letters, 403(1-2), 15–19.

Verkhratsky, A., Parpura, V., Li, B., & Scuderi, C. (2021). Astrocytes: The Housekeepers and Guardians of the CNS. Advances in neurobiology, 26, 21–53. 10.1007/978-3-030-77375-5_2

Volterra, A., & Meldolesi, J. (2005). Astrocytes, from brain glue to communication elements: the revolution continues. Nature reviews. Neuroscience, 6(8), 626–640.

Walker, D. M., Cates, H. M., Loh, Y. E., Purushothaman, I., Ramakrishnan, A., Cahill, K. M., Lardner, C. K., Godino, A., Kronman, H. G., Rabkin, J., Lorsch, Z. S., Mews, P., Doyle, M. A., Feng, J., Labonté, B., Koo, J. W., Bagot, R. C., Logan, R. W., Seney, M. L., Calipari, E. S., … Nestler, E. J. (2018). Cocaine Self-administration Alters Transcriptome-wide Responses in the Brain’s Reward Circuitry. Biological psychiatry, 84(12), 867–880.

Wegler, C., Ölander, M., Wiśniewski, J. R., Lundquist, P., Zettl, K., Åsberg, A., Hjelmesæth, J., Andersson, T. B., & Artursson, P. (2019). Global variability analysis of mRNA and protein concentrations across and within human tissues. NAR genomics and bioinformatics, 2(1), lqz010.

Wen, A. Y., Sakamoto, K. M., & Miller, L. S. (2010). The role of the transcription factor CREB in immune function. Journal of immunology (Baltimore, Md. : 1950), 185(11), 6413–6419.

Yu, X., Nagai, J., Marti-Solano, M., Soto, J. S., Coppola, G., Babu, M. M., & Khakh, B. S. (2020). Context-Specific Striatal Astrocyte Molecular Responses Are Phenotypically Exploitable. Neuron, 108(6), 1146–1162.e10.

Zhang, Y. M., Qi, Y. B., Gao, Y. N., Chen, W. G., Zhou, T., Zang, Y., & Li, J. (2023). Astrocyte metabolism and signaling pathways in the CNS. Frontiers in neuroscience, 17, 1217451.

Zhou, B., Zuo, Y. X., & Jiang, R. T. (2019). Astrocyte morphology: Diversity, plasticity, and role in neurological diseases. CNS neuroscience & therapeutics, 25(6), 665–673.

