## Supplementary material for "Investigating cocaine- and abstinence-induced effects on astrocyte gene expression in the nucleus accumbens": Exp 1_significant DEGs_supplementary data

| Gene | baseMean | log2FoldChange | adjusted p-value |
|---|---|---|---|
| Aak1 | 21465.629 | 0.232798604 | 0.045298375 |
| Aamp | 6203.1962 | -0.169669064 | 0.046118057 |
| Abcd4 | 141.75697 | -0.505309619 | 0.004140553 |
| Abhd13 | 1518.974 | 0.19462207 | 0.023005459 |
| Abhd4 | 13404.84 | -0.520972417 | 0.000137428 |
| Abl2 | 4313.8603 | 0.221414615 | 0.003250108 |
| Ablim2 | 2775.4094 | 0.396133178 | 5.67E-05 |
| Ablim3 | 1274.4488 | 0.462592531 | 8.06E-05 |
| Acaa1 | 978.71981 | -0.295898616 | 0.03549784 |
| Acadvl | 3015.3776 | -0.402087365 | 0.005769639 |
| Acbd4 | 1330.6 | -0.271334023 | 0.005674122 |
| Acbd5 | 3864.2197 | -0.235227656 | 0.037246685 |
| Acot11 | 1096.0484 | -0.422932612 | 0.012925115 |
| Acot13 | 578.79342 | -0.39857019 | 0.000495024 |
| Acot3 | 243.40992 | -0.31784391 | 0.028005732 |
| Acsbg1 | 38622.628 | -0.377138892 | 0.007764526 |
| Acss2 | 3047.3237 | -0.495923916 | 0.000527053 |
| Actl6b | 1341.167 | 0.284690316 | 0.036791471 |
| Actn1 | 2628.8085 | 0.781160076 | 7.61E-11 |
| Actn2 | 1015.8165 | 0.828131997 | 4.84E-08 |
| Actr2 | 10459.926 | 0.192898346 | 0.012177679 |
| Actr3b | 587.15346 | 0.479463555 | 0.005769639 |
| Acvr1c | 202.67684 | 0.587003869 | 0.00237863 |
| Acvr2a | 1639.2622 | 0.304784096 | 0.005431424 |
| Adam17 | 1357.6301 | -0.341097905 | 0.015872285 |
| Adamts12 | 123.31438 | -0.381192391 | 0.024940213 |
| Adamts15 | 300.38443 | 0.576448244 | 0.008160252 |
| Adamts3 | 647.22192 | 0.737173853 | 3.54E-05 |
| Adcy5 | 11046.332 | 0.77831363 | 9.90E-11 |
| Adcy9 | 4331.7421 | 0.399931547 | 0.010629693 |
| Add2 | 11222.546 | 0.317192187 | 0.020182402 |
| Add3 | 20193.081 | -0.256307338 | 0.03549784 |
| Adgra3 | 1473.2405 | -0.299845035 | 0.01352317 |
| Adora2a | 1308.1356 | 0.991954678 | 2.25E-06 |
| Adra2b | 40.995063 | 0.941674322 | 0.001693324 |

| | | | |
|---|---|---|---|
| Afap1l1 | 96.910156 | 0.547803791 | 0.008869862 |
| Afap1l2 | 626.43037 | -0.88287822 | 1.90E-05 |
| Afg3l1 | 987.82127 | -0.231323782 | 0.042835196 |
| Aga | 798.98421 | -0.259384965 | 0.044188495 |
| Agap2 | 27785.813 | 0.508166178 | 0.001313694 |
| Agk | 960.05669 | 0.239803066 | 0.039240934 |
| Agl | 8109.5236 | -0.270409955 | 0.019671737 |
| Ago2 | 3413.0017 | 0.427028201 | 4.84E-06 |
| Ajap1 | 961.97626 | 0.34272753 | 0.015322734 |
| Ak3 | 1596.8306 | -0.240109779 | 0.049362201 |
| Akap1 | 2157.1715 | 0.224297686 | 0.038522421 |
| Akap11 | 17150.422 | 0.40286302 | 9.64E-05 |
| Akap3 | 29.34912 | -0.462559515 | 0.024405849 |
| Akap5 | 3627.1545 | 1.086515137 | 9.03E-11 |
| Akap9 | 6848.024 | 0.301393703 | 0.017206321 |
| Alcam | 7558.4226 | 0.281562546 | 0.002694371 |
| Aldh1l1 | 2761.4329 | -0.406748686 | 0.00537545 |
| Aldh4a1 | 2904.5617 | -0.560195304 | 3.01E-05 |
| Aldh7a1 | 4839.8044 | -0.336687157 | 0.020537625 |
| Aldh9a1 | 2807.56 | -0.309537193 | 0.022437424 |
| Amotl2 | 2010.7968 | -0.350369712 | 0.014595571 |
| Amph | 1137.2357 | 0.275870233 | 0.030130096 |
| Amt | 741.21247 | -0.347500193 | 0.004505082 |
| Ankrd17 | 14224.419 | 0.144121382 | 0.043361782 |
| Ankrd33b | 326.89485 | 0.754866418 | 0.004320059 |
| Ankrd34a | 3217.7487 | 0.332842441 | 0.015648161 |
| Ankrd40 | 4482.2459 | -0.244963542 | 0.042409834 |
| Ankrd45 | 1349.7023 | 0.339960967 | 0.009570463 |
| Ankrd50 | 3769.651 | 0.245595306 | 0.006246528 |
| Ankrd63 | 2947.4251 | 0.940888195 | 3.69E-09 |
| Anks1b | 6849.876 | 0.516352047 | 0.001277879 |
| Ano3 | 1293.2344 | 0.874875716 | 2.72E-05 |
| Anp32e | 6616.3618 | -0.186203084 | 0.041128265 |
| Antxr1 | 505.92542 | -0.631266534 | 0.005271476 |
| Anxa3 | 63.013371 | -1.5264502 | 0.000409906 |
| Aox1 | 1872.3466 | -0.380273066 | 0.005271476 |

| | | | |
|---|---|---|---|
| Ap1s1 | 1756.4642 | 0.371565204 | 1.70E-05 |
| Ap3b2 | 6147.3883 | 0.230732237 | 0.012805122 |
| Apobec1 | 106.37231 | -0.409277959 | 0.033077453 |
| Appl1 | 4360.9634 | 0.158455715 | 0.039227522 |
| Aprt | 415.518 | -0.260364168 | 0.03549784 |
| Arap1 | 3050.2727 | -0.30084162 | 0.012898095 |
| Arc | 2588.739 | 1.333754477 | 0.000143995 |
| Arel1 | 6743.6707 | 0.316229932 | 0.000456051 |
| Arf3 | 5723.6727 | 0.460595941 | 0.000966509 |
| Arfgef2 | 4407.2023 | 0.198457643 | 0.034354216 |
| Arhgap10 | 97.056051 | 0.373117063 | 0.044157351 |
| Arhgap20 | 2706.9486 | 0.403055785 | 0.001100178 |
| Arhgap31 | 233.84449 | -1.718040512 | 4.61E-15 |
| Arhgap32 | 19845.091 | 0.35791862 | 0.004570198 |
| Arhgap33 | 4584.8536 | 0.291801017 | 0.049287215 |
| Arhgap6 | 912.00503 | 0.647131893 | 4.84E-05 |
| Arhgef26 | 2752.4852 | -0.39299533 | 0.010629693 |
| Arhgef3 | 3167.8202 | 0.378572672 | 0.019268382 |
| Arhgef9 | 6624.0355 | 0.293524484 | 0.010236886 |
| Arid1b | 6798.9737 | 0.195260521 | 0.00548891 |
| Arid4a | 1848.4764 | 0.341453351 | 0.003660647 |
| Arl13b | 504.43022 | -0.255279495 | 0.045123399 |
| Arl15 | 793.68625 | 0.502110058 | 0.000469707 |
| Arl4d | 401.38896 | 0.448120886 | 0.013191632 |
| Armcx2 | 3370.0602 | 0.268117505 | 0.03138697 |
| Armcx3 | 4066.7046 | 0.171352677 | 0.039550841 |
| Arpc1a | 2809.3542 | 0.287031101 | 0.002498022 |
| Arpp19 | 5988.3288 | 0.308603024 | 0.010940557 |
| Arpp21 | 9034.8647 | 0.777061911 | 9.07E-08 |
| Arse | 53.353358 | -0.426385259 | 0.029673981 |
| Asb2 | 56.284389 | 0.734113752 | 0.003809914 |
| Asf1a | 813.66601 | -0.31432322 | 0.020759211 |
| Ash1l | 13999.756 | 0.271781935 | 5.24E-06 |
| Asic4 | 1032.6035 | 0.891966466 | 1.16E-05 |
| Asrgl1 | 9751.0309 | -0.31926639 | 0.018480694 |
| Atf2 | 3847.9158 | 0.223860621 | 0.038270086 |

| | | | |
|---|---|---|---|
| Atg16l1 | 1423.8243 | 0.238444602 | 0.039240934 |
| Atl1 | 4056.8271 | 0.299049898 | 0.025776188 |
| Atox1 | 424.77033 | -0.239395882 | 0.027025083 |
| Atp1a1 | 17379.823 | 0.421516866 | 0.003973079 |
| Atp1a2 | 235586.72 | -0.289665609 | 0.046922194 |
| Atp2b1 | 17736.114 | 0.756915259 | 2.87E-07 |
| Atp6ap1l | 167.7801 | 0.574356252 | 0.010955008 |
| Atpaf2 | 497.56121 | -0.217717686 | 0.027879331 |
| Atraid | 646.26551 | -0.316318015 | 0.018480694 |
| Atrip | 69.834014 | 0.322988348 | 0.049806825 |
| Atxn7l1 | 1698.8683 | 0.233890765 | 0.038230165 |
| Axl | 3993.664 | -0.506889309 | 0.000911711 |
| B3gnt2 | 565.77104 | 0.802258277 | 7.43E-12 |
| B3gnt7 | 87.933222 | -0.525155575 | 0.013549639 |
| B4galnt1 | 1769.6791 | 0.370752971 | 0.012659599 |
| Bag4 | 1204.9041 | 0.267973206 | 0.026819388 |
| Bahd1 | 2360.7023 | 0.276836475 | 0.012100166 |
| Baiap2 | 4848.9977 | 0.907913906 | 7.35E-08 |
| Bax | 585.45782 | -0.38517346 | 0.006722757 |
| Bckdk | 1819.2891 | -0.256109964 | 0.011073887 |
| Bcl11b | 6893.7829 | 0.943097327 | 1.88E-11 |
| Bcor | 1253.1912 | 0.229331507 | 0.049237383 |
| Bcr | 8939.6624 | 0.266063461 | 0.000372986 |
| Bdnf | 316.68192 | 0.435750717 | 0.026483753 |
| Bgn | 397.16544 | -0.464957171 | 0.015243052 |
| Bhlhe23 | 45.427192 | 1.898674141 | 9.03E-05 |
| Bmp1 | 1540.3741 | -0.347382709 | 0.02019114 |
| Bmp2 | 110.81907 | 0.637428987 | 0.001255779 |
| Bmp4 | 50.724857 | -2.07219741 | 4.78E-06 |
| Braf | 4186.2544 | 0.327866536 | 0.001442267 |
| Brinp1 | 2916.7106 | 0.497077552 | 0.004465577 |
| Brox | 3372.8806 | -0.157416619 | 0.039257348 |
| Brpf1 | 1506.6937 | 0.206894124 | 0.017176333 |
| Btbd7 | 2511.3275 | -0.263892078 | 0.024174188 |
| Btbd8 | 4044.5436 | 0.382619887 | 0.013568107 |
| Btd | 808.95097 | -0.301147273 | 0.032605019 |

| | | | |
|---|---|---|---|
| C11H22orf2 | 226.17953 | -0.28426926 | 0.032865699 |
| C1qa | 43.704088 | -0.620722057 | 0.012333639 |
| C1ql1 | 421.0991 | -0.442373374 | 0.014053505 |
| C1qtnf4 | 3188.9241 | 0.350648726 | 0.036811212 |
| C2cd2l | 8273.3769 | 0.501415917 | 8.69E-05 |
| Cacna1c | 3268.1263 | 0.541348232 | 3.12E-05 |
| Cacna1e | 13118.69 | 0.437410669 | 0.007595487 |
| Cacna1g | 3525.4581 | 0.342450713 | 0.012015802 |
| Cacna1h | 1577.613 | 0.37959851 | 0.013314998 |
| Cacna1i | 2856.3257 | 0.482251644 | 0.003132644 |
| Cacna2d1 | 4125.2278 | 0.707757295 | 0.000602577 |
| Cacna2d3 | 4889.3167 | 0.330750193 | 0.022028973 |
| Cacnb1 | 4502.3325 | 0.522702286 | 4.12E-06 |
| Cacnb2 | 1705.1914 | 0.496776639 | 1.42E-06 |
| Cacnb4 | 6784.6269 | 0.71328512 | 3.22E-06 |
| Cadm4 | 8320.8311 | -0.18812083 | 0.036603574 |
| Calb1 | 1671.7342 | 0.70698333 | 0.00069684 |
| Calcrl | 209.84071 | -1.193847919 | 1.17E-05 |
| Calm1 | 29440.801 | 0.266349862 | 0.036763715 |
| Caln1 | 5097.6633 | 0.412255187 | 0.010872811 |
| Camk1 | 2273.7429 | 0.254256782 | 0.045343479 |
| Camk2a | 6562.438 | 0.67957311 | 0.000125976 |
| Camk2b | 22178.926 | 0.573217021 | 1.70E-05 |
| Camk4 | 4664.3377 | 0.796504937 | 9.64E-09 |
| Camkk1 | 3175.2116 | 0.257217656 | 0.046637114 |
| Camkk2 | 891.99626 | 0.455832702 | 0.01321865 |
| Camkv | 7498.155 | 0.77436825 | 3.75E-06 |
| Camsap1 | 7256.6312 | 0.246723468 | 0.018282944 |
| Cap2 | 4683.2153 | 0.358131497 | 0.012951229 |
| Car11 | 1639.6267 | 0.337678175 | 0.006979483 |
| Car8 | 1643.9581 | -0.346366394 | 0.024390808 |
| Card10 | 53.780187 | -0.489497685 | 0.021394488 |
| Carm1 | 3054.4565 | 0.188259065 | 0.017533185 |
| Cartpt | 356.03138 | 1.723636724 | 2.93E-08 |
| Casd1 | 2314.8749 | 0.309367534 | 0.000920984 |
| Caskin2 | 4211.0016 | -0.356428076 | 0.024600206 |

| | | | |
|---|---|---|---|
| Casz1 | 397.8713 | 0.829813938 | 0.000132574 |
| Cat | 4410.8859 | -0.295862759 | 0.006246528 |
| Catsper2 | 92.410796 | -0.349718525 | 0.036811212 |
| Cav1 | 211.17634 | -2.482902931 | 2.34E-12 |
| Cav2 | 159.1871 | -1.445272297 | 1.82E-09 |
| Cbfa2t3 | 1653.9162 | 0.439247053 | 0.000866427 |
| Cbr3 | 261.86778 | 0.561654991 | 0.001972228 |
| Cbx6 | 12049.23 | 0.276020106 | 0.006431804 |
| Ccdc106 | 519.58126 | 0.314866143 | 0.018735326 |
| Ccdc17 | 83.59648 | -0.319084369 | 0.04844972 |
| Ccdc28b | 372.61441 | -0.290623313 | 0.042986698 |
| Ccdc61 | 223.63947 | -0.276400442 | 0.038199374 |
| Ccdc77 | 142.19292 | -0.85274843 | 0.001588134 |
| Ccdc91 | 1452.5237 | -0.256850436 | 0.017931876 |
| Ccnd1 | 337.95816 | -3.132242868 | 6.47E-13 |
| Ccng1 | 8144.9721 | -0.296592289 | 0.042231073 |
| Ccnh | 572.93621 | 0.187100093 | 0.0483335 |
| Ccs | 1161.4614 | -0.24206645 | 0.024390808 |
| Ccsap | 1720.5123 | 0.870301188 | 6.74E-09 |
| Ccser1 | 403.62752 | 0.241587558 | 0.048194963 |
| Cd2bp2 | 4825.9381 | -0.145171777 | 0.028428243 |
| Cd302 | 535.0766 | -0.262594924 | 0.048936478 |
| Cd38 | 1563.3094 | -0.610357367 | 0.00040416 |
| Cd59 | 3109.0881 | -0.347915693 | 0.021983493 |
| Cd81 | 25910.029 | -0.341629587 | 0.016954915 |
| Cdc42ep3 | 340.19372 | 0.77800172 | 2.34E-08 |
| Cdc42se2 | 2184.58 | 0.291745146 | 0.013314998 |
| Cdh2 | 13302.928 | -0.189968449 | 0.018584008 |
| Cdh3 | 26.90289 | -1.652658892 | 0.00069684 |
| Cdk17 | 5451.3576 | 0.609100207 | 3.88E-10 |
| Cdk5r1 | 2652.1757 | 0.431431623 | 0.00069684 |
| Cdkl5 | 3633.0986 | 0.59306417 | 2.13E-05 |
| Cdkn1a | 3020.3927 | -0.559063831 | 0.007104799 |
| Cdyl2 | 1497.1288 | 0.43817859 | 7.35E-05 |
| Celf1 | 8019.7742 | 0.327640205 | 0.000110439 |
| Celf2 | 10923.47 | 0.241850943 | 0.016898989 |

| | | | |
|---|---|---|---|
| Celf3 | 2633.8504 | 0.476088081 | 0.00244492 |
| Celf4 | 12474.801 | 0.289538726 | 0.020182402 |
| Celf5 | 7596.1582 | 0.662383255 | 1.51E-05 |
| Cep120 | 1421.84 | 0.193608419 | 0.042409834 |
| Cercam | 151.93921 | -0.345909606 | 0.031791198 |
| Cers4 | 2664.9717 | -0.31333585 | 0.000742278 |
| Cherp | 3351.2155 | 0.139636773 | 0.026002044 |
| Chl1 | 6607.2921 | 0.2833923 | 0.047620083 |
| Chmp1b | 1380.8226 | -0.265293992 | 0.014877776 |
| Chn1 | 10971.827 | 0.675521596 | 7.26E-05 |
| Chpf | 5267.6887 | 0.187057606 | 0.027150309 |
| Chrm1 | 3019.0467 | 0.734081404 | 0.001018586 |
| Chrm4 | 1175.4704 | 0.898528827 | 4.46E-11 |
| Chst15 | 1098.8552 | 1.125412415 | 1.51E-11 |
| Chst3 | 91.664578 | -0.927152695 | 0.000460637 |
| Chtf8 | 2551.6488 | 0.188870111 | 0.022220003 |
| Cinp | 614.21908 | 0.337218472 | 0.001450538 |
| Cipc | 2310.8955 | 0.253438524 | 0.011496112 |
| Cited2 | 1887.5703 | 0.302734953 | 0.013683356 |
| Clcn4 | 6400.4329 | 0.253384932 | 0.015618639 |
| Cln5 | 294.64363 | -0.290777045 | 0.042867959 |
| Clock | 3766.6636 | 0.222338698 | 0.000719908 |
| Clspn | 40.07488 | 0.419317371 | 0.0285696 |
| Clvs2 | 276.6946 | 0.348217889 | 0.015589607 |
| Cnksr2 | 6918.2452 | 0.828569912 | 6.76E-06 |
| Cnnm1 | 4371.4954 | 0.288131823 | 0.026743361 |
| Cnot1 | 6537.0174 | 0.214791992 | 0.002539547 |
| Cnot7 | 1587.6661 | 0.187747621 | 0.004570198 |
| Cnp | 7190.2787 | -0.373542156 | 0.03549784 |
| Cnst | 1611.1457 | 0.256096838 | 0.002568252 |
| Cntn3 | 509.68448 | 0.370689208 | 0.007569609 |
| Cntn5 | 425.77195 | 0.695639589 | 3.76E-05 |
| Cntnap3b | 530.51048 | 0.440484006 | 0.002462658 |
| Cobl | 3027.4028 | 0.669120494 | 0.003097913 |
| Coch | 321.26318 | 0.408682058 | 0.012964143 |
| Col14a1 | 112.75266 | 0.916704292 | 0.000311561 |

| | | | |
|---|---|---|---|
| Col5a3 | 408.28257 | -2.745547949 | 3.84E-33 |
| Col9a3 | 265.80387 | -0.325337966 | 0.028827331 |
| Coq2 | 1433.8951 | 0.261671749 | 0.018026722 |
| Coro1a | 1066.6673 | 0.336509283 | 0.048439437 |
| Cpne5 | 3765.7562 | 0.933262181 | 2.76E-12 |
| Cpne6 | 1655.9106 | 0.471796924 | 0.000851225 |
| Cpxm1 | 184.6107 | -0.527141076 | 0.00973374 |
| Crabp1 | 1324.908 | 0.423796413 | 0.020550957 |
| Cramp1 | 1630.0519 | 0.233866392 | 0.007795763 |
| Creb5 | 509.71592 | -0.410788004 | 0.03318044 |
| Crebbp | 12620.658 | 0.195997942 | 0.006896307 |
| Crot | 1559.4189 | -0.339336772 | 0.018532204 |
| Crtac1 | 3530.1203 | 0.3053283 | 0.04223939 |
| Crtc1 | 6526.2855 | 0.521791553 | 1.62E-10 |
| Csad | 2124.7867 | -0.252847786 | 0.047434032 |
| Csnk1g1 | 602.55693 | 0.323967782 | 0.015105299 |
| Cspg4 | 841.69117 | -2.406496964 | 1.14E-28 |
| Csrnp1 | 225.87041 | 0.354848376 | 0.036446513 |
| Csrnp2 | 1153.1639 | 0.204465357 | 0.046932183 |
| Csrnp3 | 1315.6469 | 0.368849096 | 0.01245146 |
| Csrp1 | 15219.355 | -0.332539317 | 0.023552299 |
| Cst3 | 60496.34 | -0.284741175 | 0.029248715 |
| Ctdsp1 | 2140.6967 | -0.304663584 | 0.020424947 |
| Ctnnd1 | 5165.4889 | -0.239529684 | 0.015157997 |
| Ctsd | 10647.241 | -0.182712091 | 0.03370062 |
| Ctso | 713.83292 | -0.372782514 | 0.017847483 |
| Ctss | 45.578256 | -1.417342648 | 0.000782129 |
| Ctxn1 | 6889.5097 | 0.389067396 | 0.018683923 |
| Cuedc1 | 1159.0162 | -0.284284327 | 0.003733418 |
| Cx3cl1 | 14786.325 | 0.650308856 | 0.000602923 |
| Cyfip2 | 30595.469 | 0.528932686 | 0.000696986 |
| Cyld | 2811.7938 | 0.444902895 | 6.89E-05 |
| Cyp2j3 | 2311.6781 | -0.277209913 | 0.045066313 |
| Cyp51 | 12731.593 | -0.514335378 | 0.000897967 |
| Cyth4 | 32.190193 | -0.52072673 | 0.018685706 |
| Dab2 | 142.19643 | -0.519741503 | 0.012331824 |

| | | | |
|---|---|---|---|
| Dach1 | 365.56771 | 0.356295485 | 0.032234217 |
| Dact2 | 884.26306 | 0.450546792 | 0.00030096 |
| Dag1 | 15977.038 | -0.232751294 | 0.039894344 |
| Dapk1 | 4568.7753 | 0.399074213 | 0.00244492 |
| Dars | 2161.7305 | -0.344330118 | 0.001883729 |
| Dbi | 2132.7675 | -0.472430335 | 0.000616929 |
| Dbn1 | 4091.7909 | 0.296103779 | 0.018404824 |
| Dbx2 | 734.78197 | -0.516924939 | 0.000820817 |
| Dcaf1 | 1536.4776 | 0.185840231 | 0.034864201 |
| Dcbld2 | 1770.6083 | 0.371000951 | 0.000614172 |
| Dclk3 | 1369.9379 | 0.793530326 | 4.20E-06 |
| Ddah1 | 9103.3482 | -0.161073157 | 0.040782596 |
| Ddah2 | 4339.2635 | -0.343809915 | 0.018735326 |
| Ddhd1 | 2631.1664 | -0.238719708 | 0.00063977 |
| Ddit4l | 67.561541 | -0.436739585 | 0.028488006 |
| Ddt | 1890.1748 | -0.276187581 | 0.02848178 |
| Deptor | 1726.7712 | 0.376524275 | 0.009351421 |
| Dgat2 | 948.82886 | 0.436440575 | 0.003433683 |
| Dgkb | 10721.748 | 0.572179936 | 3.05E-06 |
| Dgkg | 3154.8664 | 0.469003115 | 0.010582209 |
| Dgkh | 3759.4818 | 0.661964311 | 0.000114597 |
| Dgki | 5383.7946 | 0.394567192 | 0.001400675 |
| Dhcr7 | 1182.6012 | -0.454634979 | 0.001838124 |
| Dhrs11 | 178.35779 | -0.36255455 | 0.042409834 |
| Dip2c | 5739.4196 | 0.295665063 | 0.00497343 |
| Diras2 | 9765.1419 | 0.353687518 | 0.044776792 |
| Dixdc1 | 3011.3585 | 0.429387015 | 0.00347036 |
| Dlg2 | 15716.354 | 0.283738326 | 0.011553088 |
| Dlg3 | 7362.4524 | 0.407224833 | 0.000686636 |
| Dlg4 | 17979.98 | 0.285757253 | 0.026294557 |
| Dlgap2 | 2201.6476 | 0.72766993 | 5.74E-06 |
| Dlgap3 | 7903.1363 | 0.34239389 | 0.024854151 |
| Dlgap4 | 6441.263 | 0.351205167 | 0.006229891 |
| Dll1 | 134.02511 | -0.535630498 | 0.002651128 |
| Dlx5 | 627.44166 | 0.429894946 | 0.015872285 |
| Dmd | 6822.5383 | -0.256969719 | 0.047593971 |

| | | | |
|---|---|---|---|
| Dmpk | 1810.0897 | -0.449450594 | 0.009199578 |
| Dmtf1 | 2801.0408 | 0.24747999 | 0.015360263 |
| Dmtn | 5604.1373 | 0.307881089 | 0.034043595 |
| Dmxl1 | 4784.286 | 0.1871845 | 0.021670745 |
| Dmxl2 | 12177.579 | 0.25824052 | 0.045112143 |
| Dnaja2 | 5656.0843 | 0.204567201 | 0.002802801 |
| Dnajb5 | 2962.8019 | 0.316610573 | 0.019845917 |
| Dnase2 | 1070.6878 | -0.287303949 | 0.040646506 |
| Dnm2 | 3537.6472 | -0.236044911 | 0.049287215 |
| Dnph1 | 67.492362 | -0.420186752 | 0.024935642 |
| Doc2b | 1210.3101 | 0.5511092 | 0.000135611 |
| Dock1 | 5228.1877 | -0.249285724 | 0.04762472 |
| Dock3 | 12314.879 | 0.322458553 | 0.042852907 |
| Dok6 | 1262.2241 | 0.305557385 | 0.039387677 |
| Dopey2 | 2792.0247 | 0.25797831 | 0.035160224 |
| Dpf1 | 1659.7661 | 0.721508144 | 1.58E-06 |
| Dpp7 | 1837.1088 | -0.194770443 | 0.038917388 |
| Dpysl3 | 2265.5897 | -0.253887215 | 0.029981841 |
| Drd1 | 2832.1985 | 1.384778373 | 2.18E-17 |
| Drd2 | 1770.1274 | 0.528362569 | 0.001312673 |
| Dscr3 | 806.85722 | -0.250868794 | 0.029822876 |
| Dtnb | 1865.979 | 0.280500437 | 0.026123697 |
| Dusp1 | 683.05131 | 0.913733227 | 0.000174779 |
| Dusp14 | 356.34914 | 0.596202517 | 0.001667804 |
| Dusp2 | 68.239131 | 0.509734739 | 0.020175353 |
| Dusp4 | 234.1546 | 0.68274889 | 0.003754791 |
| Dusp5 | 95.703926 | 0.761159601 | 0.002463922 |
| Dusp8 | 2505.9312 | 0.342346337 | 0.003292966 |
| Dync1h1 | 37819.47 | 0.230886701 | 0.042939001 |
| Ebf1 | 125.67092 | 0.392686198 | 0.036446513 |
| Ebi3 | 29.175372 | 0.361132481 | 0.047062871 |
| Ebpl | 217.84253 | -0.492599516 | 0.003513553 |
| Echdc1 | 2717.4439 | -0.33527722 | 0.010189128 |
| Edc3 | 651.64132 | 0.317141912 | 0.006442325 |
| Efnb2 | 567.42073 | 0.353525292 | 0.011710748 |
| Efs | 1521.5469 | -0.291523757 | 0.032534499 |

| | | | |
|---|---|---|---|
| Eftud2 | 3408.299 | 0.183218079 | 0.025503348 |
| Egr1 | 3912.6232 | 1.239250907 | 6.48E-05 |
| Egr2 | 252.42617 | 2.536809704 | 3.32E-10 |
| Egr3 | 2019.2873 | 2.202608198 | 1.87E-25 |
| Egr4 | 809.14397 | 1.795179144 | 9.64E-13 |
| Eif2d | 524.96651 | -0.324517172 | 0.00671353 |
| Eif4ebp1 | 428.50146 | -0.334346107 | 0.033932848 |
| Eif4enif1 | 1742.2735 | 0.30823788 | 0.004567654 |
| Elf1 | 380.85087 | -0.348629178 | 0.036985136 |
| Elk1 | 1134.9283 | 0.204115146 | 0.038230165 |
| Elmod1 | 5212.6325 | 0.664403387 | 1.08E-06 |
| Elovl5 | 7011.6111 | -0.316530902 | 0.012389117 |
| Emc7 | 2824.9198 | -0.19389911 | 0.034973979 |
| Emd | 616.73846 | 0.249379699 | 0.03370062 |
| Emp2 | 225.4498 | -0.715783871 | 0.00548891 |
| Endou | 59.373421 | -0.349357153 | 0.04485385 |
| Enox2 | 189.04841 | 0.426661572 | 0.023968682 |
| Enpp5 | 9886.9076 | 0.182305894 | 0.020500863 |
| Entpd2 | 5508.31 | -0.277829906 | 0.0483335 |
| Epb41l4b | 1257.364 | 0.382652481 | 0.010868767 |
| Epc2 | 1844.2798 | 0.236620231 | 0.015322734 |
| Epha4 | 5968.6539 | 0.280536954 | 0.043361782 |
| Ephx1 | 4151.0953 | -0.633807602 | 0.000598175 |
| Ephx4 | 354.30025 | 0.784687489 | 0.001157993 |
| Erbb3 | 431.17417 | -0.953354622 | 0.000754626 |
| Erbb4 | 3679.0532 | 0.609845509 | 1.34E-07 |
| Ercc6 | 1314.7064 | 0.173094686 | 0.031470763 |
| Erich3 | 3226.0582 | 0.387551931 | 0.019764361 |
| Ermp1 | 4327.737 | -0.267091006 | 0.008763887 |
| Ets2 | 1501.2478 | 0.821899398 | 2.35E-05 |
| Etv3 | 1055.5769 | 0.25880234 | 0.021983493 |
| Etv4 | 184.44882 | -0.497500029 | 0.020941302 |
| Eva1b | 134.24867 | -0.452122619 | 0.027335936 |
| Evc2 | 287.87832 | -0.327566733 | 0.027150309 |
| Exoc6 | 1174.8664 | 0.295692185 | 0.027087168 |
| Eya2 | 524.44741 | 0.391597015 | 0.033715252 |

| | | | |
|---|---|---|---|
| Ezh1 | 2775.9366 | 0.214213606 | 0.045298375 |
| Fabp5 | 2497.3036 | -0.370489195 | 0.005905612 |
| Fads1 | 26488.445 | -0.367259455 | 0.014525964 |
| Fads2 | 13991.999 | -0.367599486 | 0.00973277 |
| Fam102b | 1248.9537 | 0.61909119 | 0.000643429 |
| Fam110b | 1986.5761 | 0.377006394 | 0.005224357 |
| Fam117b | 3920.2284 | 0.222084341 | 0.029248715 |
| Fam124a | 984.96928 | 0.24221518 | 0.031883206 |
| Fam126b | 2474.5256 | 0.341965213 | 0.019845917 |
| Fam129b | 5660.3856 | -0.270747118 | 0.042231073 |
| Fam13b | 4628.3216 | 0.297439859 | 0.001723498 |
| Fam167a | 412.41325 | -0.331235226 | 0.03370062 |
| Fam171a2 | 4368.5521 | 0.261942929 | 0.043764586 |
| Fam184b | 250.78013 | 0.383194145 | 0.033193209 |
| Fam189a2 | 847.70809 | -0.368268949 | 0.012995176 |
| Fam189b | 1456.6406 | 0.396821293 | 0.004570198 |
| Fam213a | 508.35198 | -0.304102575 | 0.033564712 |
| Fam49a | 3252.6859 | 0.468138614 | 0.001354384 |
| Fam65a | 4336.5124 | 0.256252512 | 0.042986698 |
| Fam65b | 3848.1202 | 0.527459689 | 0.000167068 |
| Fam78b | 259.62786 | 0.319476434 | 0.036811212 |
| Fam84a | 4411.3909 | 0.435868388 | 0.000128775 |
| Fam89a | 43.966734 | -2.61724798 | 2.82E-07 |
| Fam8a1 | 2481.8903 | 0.200034439 | 0.023597059 |
| Fas | 457.06411 | -0.353859574 | 0.033346153 |
| Fasn | 28685.589 | -0.236111515 | 0.017670365 |
| Fbln2 | 4594.997 | -0.398833871 | 0.027150309 |
| Fbxl16 | 20549.852 | 0.644835226 | 5.40E-06 |
| Fbxl5 | 2727.6297 | -0.183655915 | 0.039643268 |
| Fbxo22 | 1611.0388 | -0.165639231 | 0.037531598 |
| Fbxo3 | 3682.379 | -0.18707619 | 0.022015376 |
| Fbxo34 | 1815.7612 | 0.403667825 | 0.003176112 |
| Fbxo41 | 3071.5875 | 0.26735456 | 0.03205964 |
| Fbxw4 | 611.10667 | 0.27893487 | 0.01922338 |
| Fdft1 | 4514.2981 | -0.259110447 | 0.03370062 |
| Fech | 1948.743 | -0.1856322 | 0.041388391 |

| | | | |
|---|---|---|---|
| Fermt2 | 10249.354 | -0.409311527 | 0.003391352 |
| Fgf13 | 2975.3108 | 0.294369088 | 0.012964143 |
| Fgfbp3 | 490.0716 | -0.330382348 | 0.048325325 |
| Fhl4 | 78.70235 | -0.358748786 | 0.045857663 |
| Filip1 | 894.24985 | 1.016800695 | 3.94E-07 |
| Fkbp1a | 7726.5764 | 0.564285469 | 3.33E-05 |
| Fkbp5 | 1653.0172 | 0.373979361 | 0.014337762 |
| Fmnl1 | 2586.4391 | 0.365421515 | 0.028128109 |
| Fnta | 2801.4123 | -0.215237202 | 0.049689815 |
| Folh1 | 3695.6829 | -0.330674919 | 0.022507561 |
| Fos | 852.38705 | 0.807005431 | 0.002396355 |
| Fosb | 479.01568 | 2.225772732 | 1.48E-13 |
| Fosl2 | 226.52311 | 0.530216334 | 0.017931876 |
| Foxp1 | 3780.1264 | 1.008528906 | 3.29E-13 |
| Foxp2 | 1493.2406 | 0.85915086 | 1.12E-05 |
| Fras1 | 1052.999 | 0.457098143 | 0.001201849 |
| Frem3 | 143.65783 | 0.608490111 | 0.009758988 |
| Frmd6 | 695.75381 | 0.483836157 | 0.000539393 |
| Frmd8 | 1606.2634 | -0.281503179 | 0.043361782 |
| Frmpd4 | 5992.8026 | 0.291431806 | 0.038230165 |
| Fry | 14772.896 | 0.213406881 | 0.039172988 |
| Fst | 25.81722 | 0.486263883 | 0.02256412 |
| Fstl4 | 836.12216 | 0.796242133 | 1.56E-06 |
| Fubp3 | 2323.1101 | -0.25628624 | 0.046118057 |
| Fuca1 | 1968.4271 | -0.240964581 | 0.040863716 |
| Fuca2 | 893.53534 | -0.267677001 | 0.033346153 |
| Fut2 | 179.03329 | -0.491095585 | 0.010189128 |
| Fut4 | 78.75923 | -0.615491682 | 0.008272224 |
| Fyn | 6227.0403 | -0.170261501 | 0.046940386 |
| G6pd | 3656.361 | -0.180140394 | 0.021960795 |
| Gabra4 | 2964.9175 | 0.490540205 | 5.31E-06 |
| Gabrb3 | 2772.615 | 0.458444324 | 0.002282162 |
| Gabrd | 514.88144 | 1.206975211 | 3.02E-08 |
| Gad1 | 15330.948 | 0.379357332 | 0.001946231 |
| Gal3st3 | 808.43629 | 0.491286558 | 0.003011329 |
| Galnt13 | 1075.1191 | 0.409045228 | 0.002657219 |

| | | | |
|---|---|---|---|
| Gas1 | 1534.9331 | -0.553907445 | 4.41E-05 |
| Gas7 | 19154.239 | 0.475378833 | 0.008009515 |
| Gcc2 | 3344.7749 | 0.274550786 | 0.006119053 |
| Gcnt2 | 171.23853 | 0.870368014 | 2.25E-05 |
| Gcsh | 3392.4277 | -0.245398925 | 0.048325325 |
| Gfod1 | 4952.9649 | 0.412486664 | 0.011553088 |
| Gga3 | 1946.869 | 0.24823781 | 0.048216151 |
| Ginm1 | 732.92171 | -0.3067296 | 0.03370062 |
| Glcci1 | 802.53805 | 0.264966294 | 0.02326195 |
| Glce | 2131.0604 | 0.670186263 | 5.43E-06 |
| Glis3 | 331.75381 | -0.393458845 | 0.013678882 |
| Glmp | 1161.7223 | -0.281153515 | 0.013966268 |
| Gltp | 808.96757 | -0.423323786 | 0.010617211 |
| Gltscr1 | 1395.7339 | 0.169414409 | 0.026050827 |
| Glul | 162252.08 | -0.262053284 | 0.039770336 |
| Glyctk | 295.84562 | -0.270202521 | 0.023078009 |
| Gmeb2 | 390.45168 | 0.314856198 | 0.018464042 |
| Gna12 | 7201.2447 | -0.299976243 | 0.017847483 |
| Gnal | 3701.7271 | 0.643440263 | 2.73E-05 |
| Gng4 | 2956.4374 | 0.323838748 | 0.020424947 |
| Gng7 | 4367.576 | 0.851200125 | 9.80E-07 |
| Golm1 | 660.82519 | -0.295623425 | 0.038538916 |
| Gpcpd1 | 996.21549 | 0.365776401 | 0.001312673 |
| Gpr12 | 401.14632 | 0.75200551 | 1.26E-06 |
| Gpr149 | 327.95941 | 0.704915109 | 0.001624173 |
| Gpr158 | 9158.697 | 0.216913284 | 0.019735852 |
| Gpr17 | 429.64546 | -4.429102474 | 9.26E-56 |
| Gpr176 | 475.20097 | 0.517105377 | 0.00357761 |
| Gpr22 | 883.12048 | 0.592957291 | 0.0067133 |
| Gpr37l1 | 31884.675 | -0.391752012 | 0.008410043 |
| Gpr52 | 262.72499 | 0.571277162 | 0.003093041 |
| Gpr6 | 508.5797 | 1.670107204 | 5.92E-08 |
| Gpr63 | 451.48902 | 0.877735297 | 3.83E-07 |
| Gpr83 | 1028.2914 | 0.347334401 | 0.029260344 |
| Gpr88 | 3714.6306 | 1.431287767 | 7.05E-17 |
| Gprin3 | 1061.2653 | 0.608746096 | 0.000680964 |

| | | | |
|---|---|---|---|
| Grasp | 509.62033 | 0.901746284 | 0.000171298 |
| Gria1 | 6627.9514 | 0.545891968 | 2.65E-08 |
| Gria2 | 14720.619 | 0.425414014 | 0.000834608 |
| Grin1 | 19839.778 | 0.332491797 | 0.015872285 |
| Grin2a | 6602.188 | 0.60278945 | 0.002165031 |
| Grin2b | 26375.56 | 0.46992811 | 0.003587937 |
| Grip1 | 1473.7028 | 0.335165543 | 0.002539547 |
| Gripap1 | 5653.4316 | 0.246884073 | 0.032534499 |
| Grm4 | 1250.1528 | 0.695587133 | 0.000146579 |
| Grm5 | 6954.5082 | 0.488141716 | 0.000179587 |
| Grm7 | 1415.3962 | 0.386171134 | 0.0116449 |
| Gstk1 | 718.56548 | -0.287575476 | 0.042986698 |
| Gtf2h1 | 1040.6448 | 0.216215648 | 0.045745704 |
| Gtf3c1 | 7812.4019 | 0.211285245 | 0.007500626 |
| Gtse1 | 545.3399 | -0.320997828 | 0.044180233 |
| Gucy1a2 | 1398.3456 | 0.428630004 | 0.000109267 |
| Gucy1a3 | 2766.3526 | 0.506723023 | 0.00025688 |
| Gucy1b3 | 5076.243 | 0.406181914 | 2.25E-05 |
| Gusb | 368.58307 | -0.403912395 | 0.005529975 |
| Hadha | 6897.0805 | -0.212917699 | 0.027195834 |
| Hadhb | 6309.6814 | -0.234734942 | 0.03549784 |
| Hapln3 | 119.33222 | -0.598684815 | 0.004667816 |
| Has3 | 131.33074 | 0.510641319 | 0.010850735 |
| Hdac4 | 4704.0607 | 0.295233204 | 0.000985124 |
| Hecw2 | 3309.6749 | 0.283588918 | 0.042231073 |
| Hepacam | 15070.23 | -0.260532967 | 0.046922194 |
| Hes6 | 1206.232 | -0.446662132 | 0.000719908 |
| Hibadh | 5250.093 | -0.27734405 | 0.024346105 |
| Hipk3 | 6026.9559 | 0.20632095 | 0.038998945 |
| Hist1h1d | 1701.3269 | -0.262103355 | 0.001480975 |
| Hist1h2bcl1 | 59.166933 | -0.505123103 | 0.02006917 |
| Hist1h3a | 39.1733 | -0.392733696 | 0.035542643 |
| Hist3h2ba | 382.97514 | 0.277586627 | 0.04087785 |
| Hmg20b | 736.86463 | -0.302980662 | 0.021958962 |
| Hmgcs1 | 18809.073 | -0.356592418 | 0.01245146 |
| Hmgn5b | 148.84184 | -0.411879728 | 0.015322734 |

| | | | |
|---|---|---|---|
| Hmgxb3 | 1608.6942 | 0.274705726 | 0.009806403 |
| Homer1 | 3421.0791 | 0.661073506 | 0.000336292 |
| Hpca | 6064.3862 | 0.995149043 | 3.64E-09 |
| Hpcal4 | 17135.339 | 0.62076705 | 0.00184316 |
| Hs6st2 | 2777.4586 | 0.295930532 | 0.028809824 |
| Hsd17b11 | 1262.0731 | -0.283575279 | 0.030185408 |
| Hsd17b12 | 4509.7341 | -0.244609281 | 0.043502062 |
| Hsd17b4 | 4234.2126 | -0.306009882 | 0.02162047 |
| Hsdl2 | 3032.2063 | -0.271715638 | 0.036731549 |
| Hspa13 | 2472.1308 | -0.165713585 | 0.027729474 |
| Hsph1 | 9735.4042 | 0.284496735 | 0.038230165 |
| Htr1b | 373.06482 | 0.702569164 | 1.28E-05 |
| Htr2c | 3599.8481 | 0.439611091 | 0.009570463 |
| Htt | 7589.7845 | 0.209646456 | 0.011761936 |
| Hyal1 | 4760.7745 | -0.470730996 | 0.000674895 |
| Icam5 | 4448.5583 | 0.989729077 | 1.23E-06 |
| Ide | 3289.0188 | -0.258315231 | 0.022437424 |
| Idh1 | 3003.5906 | -0.290611269 | 0.017569089 |
| Ids | 6005.1322 | 0.224617504 | 0.028929763 |
| Iffo1 | 1557.2274 | 0.296696801 | 0.002408826 |
| Ifi30 | 292.79848 | -0.362989207 | 0.027150309 |
| Igfbp4 | 1403.7803 | 0.711478528 | 0.001321987 |
| Ikbip | 1231.8528 | -0.300548511 | 0.016347312 |
| Il10ra | 140.98032 | 0.415227432 | 0.029181897 |
| Il11ra1 | 1655.4459 | -0.274444468 | 0.044491828 |
| Il1rapl2 | 441.14099 | 0.511033771 | 0.002829485 |
| Inf2 | 2600.4077 | 0.316211633 | 0.042409834 |
| Ing2 | 521.27819 | 0.298599832 | 0.035542643 |
| Ints7 | 960.23482 | 0.284029143 | 0.012898095 |
| Ip6k2 | 1794.3262 | 0.240751115 | 0.027150309 |
| Iqgap3 | 169.83602 | 0.377137211 | 0.037731263 |
| Irs2 | 6028.9138 | 0.358845698 | 0.004667816 |
| Irx1 | 50.73018 | -1.288751962 | 0.000632451 |
| Isca2 | 1068.3821 | -0.228432637 | 0.043014325 |
| Islr | 26.798503 | 0.345622788 | 0.048131614 |
| Ism1 | 43.710002 | 0.374494531 | 0.042419469 |

| | | | |
|---|---|---|---|
| Itga9 | 197.58521 | -0.431736493 | 0.023968682 |
| Itgam | 35.705255 | -0.349964659 | 0.043014325 |
| Itgav | 7867.149 | -0.289932799 | 0.019046528 |
| Itgb5 | 4263.0448 | -0.303892254 | 0.020424947 |
| Itgb8 | 14223.843 | -0.254454132 | 0.049694547 |
| Itpa | 652.61477 | 0.307442964 | 0.01138543 |
| Itpka | 1488.7092 | 0.961354982 | 2.27E-06 |
| Itpr1 | 18327.403 | 0.579976253 | 0.00395093 |
| Jam3 | 1510.3674 | -0.298843346 | 0.04762472 |
| Josd1 | 1290.9145 | 0.373995038 | 0.001065936 |
| Jph1 | 268.83503 | 0.645162042 | 0.003761373 |
| Jph3 | 8079.9464 | 0.288194164 | 0.028097423 |
| Jph4 | 8618.6815 | 0.543832037 | 0.000125135 |
| Junb | 1377.2429 | 1.228373847 | 1.12E-06 |
| Kalrn | 7719.4708 | 0.309520349 | 0.042359371 |
| Kank1 | 1416.6237 | -0.43813854 | 0.005895842 |
| Kank2 | 1828.1833 | -0.27720461 | 0.030713861 |
| Kantr | 96.416357 | 0.638292867 | 0.001588134 |
| Kcna1 | 1524.9523 | 0.368941668 | 0.03984196 |
| Kcna4 | 1963.6211 | 0.654344175 | 5.73E-06 |
| Kcna5 | 396.34488 | 0.568856584 | 0.000674895 |
| Kcnab1 | 4144.7618 | 0.916881769 | 1.38E-09 |
| Kcnb1 | 3545.7525 | 0.308857445 | 0.042322861 |
| Kcnd2 | 4071.6221 | 0.239515544 | 0.012995176 |
| Kcne5 | 275.26893 | -0.35528623 | 0.042361177 |
| Kcnf1 | 2779.3736 | 0.920235407 | 2.08E-05 |
| Kcnh1 | 3568.826 | 0.84567598 | 7.89E-06 |
| Kcnh3 | 1491.7734 | 0.411944099 | 0.029224061 |
| Kcnh4 | 213.03777 | 0.789337392 | 5.18E-05 |
| Kcnj2 | 432.34082 | 1.067488084 | 1.30E-06 |
| Kcnj4 | 1596.1601 | 0.827061578 | 0.00025688 |
| Kcnk13 | 101.15821 | 0.376579637 | 0.042348444 |
| Kcnk2 | 1306.6124 | 0.538631176 | 0.000193915 |
| Kcnq5 | 1788.7072 | 0.789797873 | 0.000152712 |
| Kcns2 | 318.22839 | 0.550198334 | 0.006729732 |
| Kcnt1 | 1760.3737 | 0.443968629 | 8.05E-07 |

| | | | |
|---|---|---|---|
| Kcnt2 | 657.51993 | 0.306309022 | 0.037485042 |
| Kcnv1 | 985.41151 | 0.480879208 | 0.02060561 |
| Kctd1 | 2811.1874 | 0.344512426 | 0.001513953 |
| Kctd12 | 2574.2886 | 0.69775261 | 2.12E-05 |
| Kctd13 | 1910.7933 | 0.389872836 | 0.001946231 |
| Kctd16 | 1440.1463 | 0.599567767 | 0.000113491 |
| Kctd8 | 479.46871 | 0.423922235 | 0.002441699 |
| Kdelc2 | 1083.2538 | -0.241563997 | 0.045351476 |
| Kif11 | 44.099202 | -0.452069163 | 0.025567429 |
| Kif1c | 5943.9005 | -0.267133608 | 0.042409834 |
| Kifc3 | 3849.0283 | -0.339736213 | 0.004196452 |
| Kit | 4619.8675 | 0.602746711 | 0.001442267 |
| Kitlg | 453.77164 | 0.563136962 | 0.00033699 |
| Kl | 102.16572 | 0.527652181 | 0.014561277 |
| Klf10 | 516.86773 | 0.802451439 | 0.001563953 |
| Klf16 | 1878.0123 | 0.568116795 | 8.26E-06 |
| Klf2 | 32.54995 | 1.169278449 | 0.001442267 |
| Klf3 | 1565.3535 | -0.316445357 | 0.011553088 |
| Klf5 | 818.89103 | 0.845861087 | 2.45E-06 |
| Klf7 | 1029.448 | 0.282302526 | 0.045853193 |
| Klhl2 | 3761.7623 | 0.503986885 | 0.000726668 |
| Klhl29 | 2786.9619 | 0.277650476 | 0.02185164 |
| Klhl3 | 423.89481 | 0.351376932 | 0.03370062 |
| Kmt2d | 14025.582 | 0.303774148 | 0.004301218 |
| Krcc1 | 980.04322 | -0.331294706 | 0.02630111 |
| Kremen1 | 403.05387 | 0.272113156 | 0.039598359 |
| Krt71 | 170.41152 | 1.175844158 | 1.57E-05 |
| L1cam | 9451.9449 | 0.277056853 | 0.027645705 |
| Lamb1 | 1537.6097 | 0.578621803 | 0.000767877 |
| Lamp1 | 21289.493 | -0.230496837 | 0.037015005 |
| Lamp5 | 691.21146 | 1.329558192 | 1.19E-13 |
| Lancl1 | 2350.91 | 0.586081839 | 7.57E-07 |
| Lca5 | 881.84699 | -0.302867593 | 0.030855903 |
| Lcn2 | 57.147315 | -0.230777246 | 0.02277008 |
| Lcor | 1465.4785 | 0.238420648 | 0.018362068 |
| Ldhd | 1099.3364 | -0.347711467 | 0.001920085 |

| | | | |
|---|---|---|---|
| Lgi1 | 1904.3893 | 0.575889519 | 4.72E-08 |
| Lgi4 | 13635.611 | -0.384022223 | 0.009125973 |
| Lgr6 | 390.45081 | 1.227313481 | 5.51E-07 |
| Lhpp | 3978.1227 | -0.243275673 | 0.03949779 |
| Lims1 | 1784.216 | -0.310028229 | 0.029891713 |
| Lingo3 | 682.17158 | 1.160514811 | 3.08E-18 |
| Lmbrd2 | 2778.0787 | 0.200599695 | 0.04087785 |
| Lmo4 | 1484.214 | 0.250868006 | 0.036166859 |
| Lmo7 | 1863.3702 | 0.684046796 | 5.03E-10 |
| Lmtk2 | 9717.9364 | 0.340099873 | 0.018743412 |
| LOC100125 | 4647.8902 | 0.834119174 | 1.71E-13 |
| LOC100359 | 344.45965 | 0.267823162 | 0.017670365 |
| LOC100909 | 64.474929 | 0.55331756 | 0.012995176 |
| LOC100909 | 357.77994 | 0.278578312 | 0.03839648 |
| LOC100910 | 41.532557 | 0.485294871 | 0.022460173 |
| LOC100910 | 485.11981 | -0.354023331 | 0.013574023 |
| LOC100910 | 1051.0021 | 0.270493379 | 0.023955603 |
| LOC100911 | 4091.992 | 0.2033468 | 0.017569089 |
| LOC100911 | 1513.588 | 0.219912804 | 0.0199464 |
| LOC100911 | 171.04073 | -0.328428557 | 0.041894635 |
| LOC100911 | 2340.1579 | -0.17151365 | 0.041493703 |
| LOC100911 | 248.45114 | 0.364564339 | 0.023422381 |
| LOC100912 | 2266.3104 | 0.854950759 | 2.50E-07 |
| LOC102546 | 110.794 | -0.406993133 | 0.028817381 |
| LOC102546 | 136.46557 | 0.781480373 | 0.0005425 |
| LOC102547 | 88.735529 | 0.375047088 | 0.038377108 |
| LOC102547 | 113.51947 | -0.484724126 | 0.021299288 |
| LOC102547 | 24.256805 | 0.652751496 | 0.009739963 |
| LOC102549 | 258.92513 | 0.560036228 | 0.004882826 |
| LOC102550 | 518.81087 | -0.341025047 | 0.030097449 |
| LOC102550 | 41.582435 | -0.699934169 | 0.008960193 |
| LOC102551 | 368.31807 | 0.522302302 | 0.000379361 |
| LOC102551 | 1404.999 | 0.432729632 | 0.007527668 |
| LOC102551 | 79.13393 | -0.369407945 | 0.045038699 |
| LOC102552 | 87.634001 | 0.935952811 | 4.60E-05 |
| LOC102553 | 50.115716 | 0.407426684 | 0.035660344 |

| | | | |
|---|---|---|---|
| LOC102557 | 38.001079 | 1.139991208 | 0.000777069 |
| LOC103691 | 494.85616 | -0.318526044 | 0.048799918 |
| LOC103692 | 3641.6919 | -0.353866164 | 0.023552299 |
| LOC103693 | 499.43849 | 0.492884276 | 0.010940557 |
| LOC108348 | 5680.3811 | -0.313997796 | 0.042359371 |
| LOC108348 | 99.7172 | 0.476313057 | 0.023862019 |
| LOC108350 | 514.70901 | -0.327094827 | 0.026270301 |
| LOC361635 | 2492.9246 | -0.171003401 | 0.049806825 |
| LOC365985 | 3978.0825 | 0.416954467 | 0.017780352 |
| LOC501038 | 332.73825 | 0.377997044 | 0.030340145 |
| LOC680142 | 135.83428 | 0.400160175 | 0.025811912 |
| LOC684270 | 451.93769 | -0.427703431 | 0.002985894 |
| LOC687399 | 78.601579 | 0.453511859 | 0.021247261 |
| LOC689561 | 37.536587 | 0.570148657 | 0.015105299 |
| LOC689986 | 646.60476 | 0.45850205 | 0.00504516 |
| Lonp2 | 5114.6103 | -0.178656623 | 0.023755448 |
| Lpl | 1588.3741 | 1.371490891 | 9.42E-09 |
| Lrig1 | 4424.5871 | -0.325044385 | 0.015322734 |
| Lrp12 | 1826.7659 | 0.417104748 | 0.00044862 |
| Lrp5 | 1626.9037 | -0.278978599 | 0.048866243 |
| Lrrc10b | 357.42966 | 0.43784285 | 0.002787883 |
| Lrrc4 | 2634.586 | 0.237042532 | 0.029891713 |
| Lrrc4c | 2885.9053 | 0.297807407 | 0.043172776 |
| Lrrc7 | 4171.5132 | 0.517891091 | 3.01E-05 |
| Lrrc73 | 553.69549 | 0.438970199 | 0.005146944 |
| Lrrc8b | 2491.2163 | 0.408789923 | 0.012995176 |
| Lrrk2 | 3435.1606 | 0.591041603 | 9.79E-07 |
| Lrrtm3 | 1716.6047 | 0.526041893 | 0.00128427 |
| Lsamp | 25873.201 | -0.151330361 | 0.047062871 |
| Lss | 1288.698 | -0.308872155 | 0.014473699 |
| Ltbp3 | 4860.9506 | -0.245513961 | 0.032707788 |
| Luc7l | 3323.1646 | 0.137789128 | 0.038944272 |
| Lypd1 | 2691.4551 | 0.361795669 | 0.016464863 |
| Lyplal1 | 415.59649 | -0.304435254 | 0.006521067 |
| Lyst | 2198.7656 | 0.372666274 | 6.65E-05 |
| Lzts1 | 4551.1585 | 0.706445878 | 8.49E-05 |

| | | | |
|---|---|---|---|
| Lzts3 | 5645.2874 | 0.729267285 | 2.59E-13 |
| Macrod2 | 1034.625 | 0.232028443 | 0.046118057 |
| Madd | 6959.3882 | 0.293491851 | 0.016516661 |
| Mafb | 703.08562 | 0.47961358 | 0.010549698 |
| Maged2 | 1976.8318 | -0.217618588 | 0.04555874 |
| Man1a1 | 1133.7553 | 0.554161124 | 0.000289197 |
| Man1c1 | 1208.0604 | 0.596606424 | 0.000146579 |
| Map2k1 | 10686.703 | 0.490360352 | 5.36E-08 |
| Map3k13 | 1190.9166 | 0.394773091 | 0.000294239 |
| Map3k3 | 481.24468 | 0.286156648 | 0.01170859 |
| Map4k4 | 10099.908 | -0.290538916 | 0.018528008 |
| Map9 | 5642.0858 | 0.29717617 | 0.029385721 |
| Mapk1 | 17196.442 | 0.38716492 | 6.79E-05 |
| Mapk12 | 350.76671 | -0.322183898 | 0.032883247 |
| Mapkap1 | 1764.1579 | -0.14595217 | 0.049015258 |
| Mapkbp1 | 1593.3617 | 0.400541012 | 2.06E-05 |
| Mapre1 | 3921.4617 | -0.226147548 | 0.020948585 |
| Mapre2 | 8859.3577 | 0.21751625 | 0.00062075 |
| March4 | 733.12042 | 0.410525989 | 0.004647909 |
| March8 | 2209.4629 | -0.410165523 | 0.004424841 |
| Mark2 | 4725.6229 | 0.427675848 | 0.000288322 |
| Mast3 | 6228.8961 | 0.931050476 | 3.93E-11 |
| Mat2b | 2798.1791 | 0.293676065 | 0.047593971 |
| Matn4 | 38.999591 | -2.392134517 | 3.00E-05 |
| Matr3 | 10894.775 | 0.260885363 | 0.015105299 |
| Mavs | 1049.0637 | -0.227333241 | 0.04087785 |
| Mblac2 | 526.71153 | 0.244982963 | 0.0285696 |
| Mbnl1 | 2461.8549 | 0.232746099 | 0.008160252 |
| Mbnl2 | 6439.5096 | 0.23479813 | 0.015872285 |
| Mchr1 | 377.8082 | 1.154974103 | 3.48E-09 |
| Mctp1 | 1756.2085 | 0.769274123 | 1.53E-08 |
| Mdk | 1447.0972 | -0.333857036 | 0.008350772 |
| Me1 | 2878.8025 | -0.315200926 | 0.007287532 |
| Mecp2 | 8565.1271 | 0.161077057 | 0.035892978 |
| Mef2a | 5773.9338 | 0.571660553 | 2.29E-05 |
| Mef2d | 8661.9722 | 0.475939707 | 3.13E-07 |

| | | | |
|---|---|---|---|
| Megf11 | 2044.9449 | 0.343300474 | 0.033866092 |
| Meis1 | 737.53357 | -0.272594663 | 0.034511766 |
| Meis2 | 6008.1556 | 0.609811395 | 1.67E-09 |
| Mfsd1 | 1809.5956 | -0.247784999 | 0.042867959 |
| Mgat4a | 1330.2276 | 0.289272631 | 0.024854151 |
| Mgme1 | 121.0343 | -0.300879347 | 0.04087785 |
| Micall2 | 191.88548 | -0.638014053 | 0.000937671 |
| Micu3 | 2256.3402 | 0.385914331 | 0.000921288 |
| Mid1ip1 | 6539.1922 | -0.494682355 | 0.000332221 |
| Mier2 | 790.49549 | 0.23130389 | 0.0483335 |
| Mink1 | 16183.795 | 0.298464281 | 0.000756379 |
| Mkl1 | 2171.1755 | 0.32632223 | 0.005332545 |
| Mkl2 | 783.34295 | 0.598354677 | 0.000379993 |
| Mlc1 | 49782.738 | -0.289676252 | 0.033955394 |
| Mmab | 727.30098 | -0.291916629 | 0.011496112 |
| Mmd | 3374.0611 | 0.342126679 | 0.029989459 |
| Mmp15 | 5238.3363 | -0.277258244 | 0.046272378 |
| Mmp2 | 233.06772 | -1.070570216 | 8.52E-05 |
| Mmp24 | 1268.7372 | 0.27867687 | 0.032534499 |
| Mpeg1 | 82.575375 | -0.348158823 | 0.047062871 |
| Mpp3 | 1043.341 | 0.378838382 | 0.009256237 |
| Mpv17l2 | 568.92176 | -0.354218465 | 0.013864846 |
| Mrpl32 | 309.17691 | 0.26118119 | 0.04688646 |
| Mrpl4 | 1619.5542 | -0.176579228 | 0.04555874 |
| Mrpl45 | 1691.0445 | -0.216985142 | 0.031470763 |
| Msmo1 | 6298.8676 | -0.530440736 | 0.000211327 |
| Mthfd1 | 3243.4727 | -0.264628924 | 0.018537955 |
| Mthfs | 344.51777 | -0.359004939 | 0.015203891 |
| Mtmr12 | 760.82564 | 0.37291535 | 0.001562073 |
| Mtmr7 | 1936.225 | 0.311464328 | 0.028824169 |
| Mtus2 | 1064.8957 | 0.463830256 | 8.67E-07 |
| Myh14 | 6287.3279 | -0.251041714 | 0.048372633 |
| Myh7 | 281.51199 | 0.571092631 | 0.008763887 |
| Myo5b | 4263.3457 | 0.35246039 | 0.028885951 |
| Nab2 | 706.24549 | 0.921949813 | 4.40E-08 |
| Nanos1 | 602.0353 | 0.398583128 | 0.002932558 |

| | | | |
|---|---|---|---|
| Napa | 5009.8114 | 0.232188792 | 0.013314998 |
| Nat6 | 1872.7613 | -0.41323325 | 0.003421864 |
| Nbea | 10414.937 | 0.412731015 | 0.000162268 |
| Ncapd2 | 625.73426 | -0.265926809 | 0.042409834 |
| Ncdn | 49443.727 | 0.584353058 | 6.62E-12 |
| Ncoa5 | 2199.0498 | 0.233510963 | 0.006817652 |
| ND6 | 414.62981 | -0.32524032 | 0.04685399 |
| Ndel1 | 2814.695 | 0.297033553 | 0.005171146 |
| Ndrg2 | 160446.41 | -0.317425534 | 0.026076232 |
| Necab1 | 858.28078 | 0.871690506 | 1.86E-11 |
| Necab2 | 1950.8365 | 0.621196041 | 0.000525457 |
| Necap2 | 1603.1361 | -0.273006172 | 0.041842731 |
| Nedd4l | 3527.2712 | 0.373892586 | 0.000368199 |
| Nell2 | 9424.1939 | 0.445301415 | 0.016801061 |
| Neto1 | 3274.9704 | 0.656202668 | 0.000246483 |
| Neu2 | 148.50761 | 0.481721475 | 0.015322734 |
| Neurl4 | 3982.2891 | 0.196245394 | 0.048194963 |
| NEWGENE_ | 2760.6377 | 0.220688442 | 0.049689815 |
| Nexn | 478.24065 | 0.519446587 | 0.006089657 |
| Nfe2l2 | 3193.0611 | -0.271717788 | 0.045298375 |
| Nfx1 | 3408.798 | 0.365002627 | 6.83E-06 |
| Ngef | 8600.2103 | 0.616246039 | 4.84E-05 |
| Nhsl2 | 3250.8852 | 0.394800071 | 0.007888407 |
| Ninl | 150.63164 | -0.30763499 | 0.032108601 |
| Nit2 | 1034.004 | -0.33852419 | 0.000631616 |
| Nkiras1 | 2509.2588 | 0.574377003 | 0.000586146 |
| Nkpd1 | 179.01376 | 1.038137909 | 2.34E-08 |
| Noct | 1481.0011 | 0.346978976 | 0.00310838 |
| Nol4 | 1203.3648 | 0.361786889 | 0.007595487 |
| Nol6 | 5821.835 | 0.156630268 | 0.045123399 |
| Nova2 | 2806.165 | 0.217716872 | 0.048214429 |
| Npas4 | 576.23358 | 1.188088363 | 0.000328459 |
| Npc2 | 1748.6939 | -0.351458361 | 0.006817652 |
| Npepl1 | 897.16768 | -0.270139343 | 0.042986698 |
| Npr1 | 621.89716 | -0.401828605 | 0.030259654 |
| Nptn | 10834.326 | 0.407638442 | 0.000146579 |

| | | | |
|---|---|---|---|
| Nr3c1 | 28.257176 | 0.988079456 | 0.001723498 |
| Nr4a1 | 2189.5358 | 1.400221003 | 6.25E-07 |
| Nr4a3 | 2200.1121 | 0.488344412 | 0.013385261 |
| Nsg2 | 23950.691 | 0.343573603 | 0.001501262 |
| Nt5dc3 | 2400.2765 | 0.50419571 | 0.003242835 |
| Nthl1 | 96.241525 | -0.897653836 | 6.75E-05 |
| Ntng1 | 674.31448 | 0.495076855 | 0.005146944 |
| Nudt12 | 796.02693 | -0.210112627 | 0.02848178 |
| Nudt5 | 414.7213 | -0.262898596 | 0.021790856 |
| Nup210 | 1187.4659 | 0.314979736 | 0.012295122 |
| Nxph1 | 2934.9096 | 0.458647019 | 0.001079558 |
| Obsl1 | 496.39182 | -0.362762131 | 0.026057776 |
| Ofd1 | 506.99992 | 0.237461325 | 0.035660344 |
| Ogdh | 7090.974 | -0.199499209 | 0.019845917 |
| Olfml1 | 3414.6737 | -0.292258831 | 0.041440168 |
| Olig1 | 1419.3994 | -2.099982214 | 1.38E-26 |
| Olig2 | 459.56987 | -1.721991214 | 3.02E-09 |
| Oplah | 2158.2251 | -0.278946954 | 0.02060561 |
| Opn3 | 412.3528 | 0.531714286 | 0.006657137 |
| Oprk1 | 351.01652 | 0.695010777 | 0.00062075 |
| Oprm1 | 771.28749 | 0.321721792 | 0.023147251 |
| Osbpl3 | 662.99994 | 0.452025309 | 0.019735852 |
| Osbpl8 | 4074.5185 | 0.315357164 | 0.000726668 |
| Otof | 1343.5306 | 1.135387455 | 2.39E-07 |
| Otud1 | 934.2657 | 0.341981559 | 0.038187064 |
| P2ry1 | 323.12094 | 0.331097246 | 0.046118057 |
| P4htm | 2371.0079 | 0.243557545 | 0.036811212 |
| Padi2 | 3068.4919 | -0.464543155 | 0.012659599 |
| Pafah2 | 687.70492 | -0.309454738 | 0.012553347 |
| Paip1 | 3611.7499 | -0.159684332 | 0.035892978 |
| Pak6 | 2507.6833 | 0.44777168 | 0.009758988 |
| Paqr9 | 631.02999 | 0.378193437 | 0.020500863 |
| Park7 | 5370.08 | -0.146836681 | 0.029989459 |
| Pbx2 | 1590.9846 | 0.256461748 | 0.04097385 |
| Pbxip1 | 6099.959 | -0.388430187 | 0.005905612 |
| Pccb | 945.40427 | -0.344324276 | 0.010189128 |

| | | | |
|---|---|---|---|
| Pcdh1 | 7338.57 | 0.292777186 | 0.028097423 |
| Pcdha4 | 8269.3781 | 0.393173144 | 0.002606118 |
| Pcdhgb4 | 653.65508 | -0.280782155 | 0.043361782 |
| Pcp4l1 | 2290.5043 | 0.435470132 | 0.024346105 |
| Pcsk1 | 225.98339 | 0.60664225 | 0.000488778 |
| Pcsk2 | 5047.0714 | 0.498691689 | 0.002885686 |
| Pde10a | 3294.3726 | 1.115927206 | 1.25E-26 |
| Pde1b | 7933.3939 | 0.801947329 | 1.44E-11 |
| Pde1c | 1168.2314 | 0.604427386 | 0.000670741 |
| Pde2a | 8062.4126 | 0.779784567 | 7.84E-08 |
| Pde7b | 3728.4148 | 0.903254492 | 2.34E-12 |
| Pdgfb | 164.56532 | 0.402284839 | 0.036811212 |
| Pdgfra | 1155.6333 | -3.813436799 | 5.18E-113 |
| Pdlim5 | 5497.2017 | -0.25512722 | 0.048799918 |
| Pdp1 | 1542.2967 | 0.591902058 | 0.000244838 |
| Pdp2 | 687.15976 | -0.266594748 | 0.044487649 |
| Pdpk1 | 7395.9473 | 0.30356458 | 0.00152522 |
| Pdyn | 2096.7656 | 1.319352825 | 8.86E-13 |
| Pdzd2 | 2560.0199 | 0.885265447 | 1.23E-10 |
| Pdzrn3 | 3854.9608 | -0.289547372 | 0.012632788 |
| Pdzrn4 | 171.35541 | -0.591597755 | 0.001000708 |
| Penk | 7963.7989 | 0.860908663 | 1.54E-07 |
| Per1 | 2550.3434 | 0.311758428 | 0.043861839 |
| Per2 | 1924.1821 | 0.315235313 | 0.01352317 |
| Pex3 | 1026.7132 | -0.24279467 | 0.001170368 |
| Pfkm | 27237.806 | -0.167646207 | 0.027746436 |
| Pgbd5 | 2888.5792 | 0.405736257 | 0.017204694 |
| Pgm3 | 1479.1246 | -0.304806294 | 0.010491142 |
| Phactr1 | 6881.1745 | 0.655306535 | 3.94E-07 |
| Phka1 | 5287.4439 | -0.281402177 | 0.042951471 |
| Phlda1 | 821.23762 | 0.564885362 | 0.00244492 |
| Phlda3 | 3194.9455 | -0.557326281 | 0.002813367 |
| Phldb1 | 2053.6376 | -0.349264111 | 0.034401146 |
| Phyh | 3506.7473 | -0.293464389 | 0.03217283 |
| Pi4ka | 15534.892 | 0.244034263 | 0.049214614 |
| Pigk | 2014.5766 | -0.215493055 | 0.024648457 |

| | | | |
|---|---|---|---|
| Pik3cg | 32.370087 | 0.423100683 | 0.032534499 |
| Pip5k1a | 2323.4452 | 0.358768584 | 0.002205747 |
| Pir | 878.98905 | -0.345579611 | 0.007750658 |
| Pirt | 134.27268 | -0.401859059 | 0.0344978 |
| Pitpnm2 | 14158.933 | 0.242411695 | 0.024648457 |
| Pitpnm3 | 2179.2584 | 0.530855581 | 9.20E-06 |
| Pja2 | 31899.941 | 0.286590615 | 0.004095648 |
| Pkia | 3619.8763 | 0.258863785 | 0.03549784 |
| Pla2g2c | 402.5631 | -0.362199459 | 0.031902767 |
| Plbd2 | 2904.684 | -0.204109425 | 0.001961251 |
| Plcb1 | 10647.182 | 0.206146157 | 0.049806825 |
| Plcd1 | 2272.8078 | -0.346246197 | 0.008410043 |
| Plcl2 | 2239.2224 | 0.419183616 | 0.006561137 |
| Plcxd3 | 1183.1576 | 0.395622954 | 0.018687114 |
| Plekha7 | 163.08757 | -0.333573518 | 0.03549784 |
| Plekhd1 | 1721.308 | -0.435909151 | 0.005209594 |
| Plk2 | 5313.5384 | 0.559218753 | 0.004045521 |
| Pllp | 400.7429 | -0.521033828 | 0.018584008 |
| Plppr1 | 381.44777 | 0.629122486 | 0.001312673 |
| Plppr4 | 7182.4385 | 0.665892556 | 2.19E-06 |
| Plppr5 | 2194.2705 | 0.357577107 | 0.003949358 |
| Plxdc1 | 287.11467 | 0.44221802 | 0.020157314 |
| Plxna2 | 9219.3663 | 0.451747728 | 0.000249244 |
| Plxnb3 | 670.8467 | -0.451815517 | 0.008135492 |
| Pmel | 92.006651 | -3.054131583 | 3.05E-22 |
| Pmm1 | 5503.041 | -0.226113959 | 0.045120097 |
| Pmp22 | 640.14771 | -0.351334567 | 0.042867959 |
| Pnlip | 35.402641 | -2.396633039 | 3.17E-07 |
| Pnp | 8886.6893 | -0.384127414 | 0.01245146 |
| Poglut1 | 877.76232 | -0.241777577 | 0.029174354 |
| Pogz | 3953.5165 | 0.201223555 | 0.020424947 |
| Polm | 312.59315 | -0.27826064 | 0.046028852 |
| Pou3f2 | 2458.0374 | -0.298653887 | 0.023979765 |
| Ppdpf | 1391.8639 | -0.314905051 | 0.000674895 |
| Ppfia2 | 3082.3696 | 0.318207755 | 0.013385261 |
| Ppm1h | 1936.3072 | 0.292979491 | 0.004233849 |

| | | | |
|---|---|---|---|
| Ppox | 334.28243 | -0.275004502 | 0.042409834 |
| Ppp1r13b | 3330.0046 | 0.286931148 | 0.01184379 |
| Ppp1r1a | 1177.4336 | 0.365123159 | 0.005271476 |
| Ppp1r3c | 7597.6027 | -0.393286119 | 0.004209806 |
| Ppp1r7 | 4214.7581 | 0.264401206 | 0.027500606 |
| Ppp1r9a | 10006.416 | 0.526213311 | 4.95E-05 |
| Ppp3ca | 15224.672 | 0.855700115 | 4.72E-14 |
| Ppp3cb | 6655.0315 | 0.389208496 | 0.007144518 |
| Ppp3r1 | 12418.259 | 0.450828038 | 0.000275477 |
| Ppp4r4 | 382.86847 | 0.534954696 | 0.001645874 |
| Prdm15 | 415.62106 | 0.243413878 | 0.046118057 |
| Prickle2 | 8830.2267 | 0.258880672 | 0.038230165 |
| Prkacb | 19035.759 | 0.315844616 | 0.001684115 |
| Prkar2b | 2848.9507 | 0.463431277 | 0.000108495 |
| Prkcb | 11811.901 | 0.410233043 | 0.015105299 |
| Prkcg | 6152.1009 | 0.687138706 | 0.001145512 |
| Prkch | 401.54544 | 0.985004236 | 4.35E-05 |
| Prkcz | 5042.2628 | 0.4322436 | 0.001358012 |
| Prkg2 | 260.43983 | 0.478318489 | 0.005780364 |
| Prodh1 | 4525.2072 | -0.365562053 | 0.018735326 |
| Prokr2 | 467.29147 | 0.525931667 | 0.01352317 |
| Pros1 | 594.23454 | -0.571638113 | 0.000568292 |
| Prp2l1 | 2047.6588 | 0.219033202 | 0.040627784 |
| Prpf8 | 17332.152 | 0.148534259 | 0.037245499 |
| Prr12 | 5814.4933 | 0.196005546 | 0.032744823 |
| Prrc2c | 9129.5031 | 0.180241116 | 0.037309969 |
| Prrt2 | 3392.0001 | 0.427324312 | 0.006246528 |
| Prrx1 | 1207.9263 | -0.368515807 | 0.005062133 |
| Psd | 5479.7924 | 0.548522805 | 1.72E-05 |
| Psenen | 820.43383 | -0.219501917 | 0.035414682 |
| Psmc3ip | 241.22799 | -0.313428357 | 0.041266518 |
| Psph | 816.08369 | -0.250089949 | 0.032194197 |
| Pstk | 172.31993 | -0.30193009 | 0.031467121 |
| Ptgr2 | 4834.0785 | -0.36411017 | 0.009642473 |
| Ptgs2 | 103.08017 | 0.780500392 | 0.002165031 |
| Pthlh | 29.623435 | 1.072080832 | 0.002472741 |

| | | | |
|---|---|---|---|
| Ptk2b | 15160.343 | 0.426824096 | 0.00025341 |
| Ptpn5 | 3400.4617 | 0.862802752 | 2.34E-12 |
| Ptpn7 | 31.374547 | 0.450755676 | 0.027195834 |
| Ptprj | 4442.1658 | 0.364924465 | 0.013666768 |
| Ptprn | 19725.008 | 0.256126923 | 0.041043424 |
| Ptprn2 | 15040.519 | 0.281469106 | 0.025042201 |
| Pum2 | 9210.5774 | 0.156542396 | 0.044157351 |
| Purg | 4220.1991 | 0.236775479 | 0.032605019 |
| Pygm | 10500.953 | -0.297120711 | 0.033025544 |
| Rab21 | 4727.5252 | -0.255042576 | 0.006550686 |
| Rab27b | 1962.0376 | 0.282848167 | 0.018735326 |
| Rab31 | 4808.6409 | -0.258079655 | 0.0285696 |
| Rab34 | 496.73721 | -0.30860485 | 0.031186715 |
| Rab40b | 815.08258 | 1.139251925 | 2.34E-12 |
| Rad9a | 482.8288 | -0.331763666 | 0.028005102 |
| Rad9b | 68.80945 | -0.540413822 | 0.007788093 |
| Rai1 | 4927.6876 | 0.241662386 | 0.012572006 |
| Ranbp2 | 8691.5846 | 0.217476588 | 0.002139052 |
| Rap1gap | 9388.5669 | 0.383938692 | 0.001088701 |
| Rapgef2 | 9478.831 | 0.295721815 | 0.021088806 |
| Rapgef4 | 3070.084 | 0.316015875 | 0.021983493 |
| Rapgef5 | 3536.024 | 0.630983156 | 0.000115364 |
| Rarb | 1311.1569 | 0.490233791 | 0.00665364 |
| Rasal2 | 2325.5355 | 0.319600466 | 0.034450487 |
| Rasd2 | 4113.6031 | 0.597190896 | 0.000124013 |
| Rasgef1a | 3638.1318 | 0.350169987 | 0.029510215 |
| Rasgrp1 | 14762.215 | 0.6801341 | 0.002645775 |
| Rasgrp2 | 872.96 | 0.296994137 | 0.023552299 |
| Rassf10 | 39.480204 | -0.42605035 | 0.027333715 |
| Rbfox1 | 7146.4893 | 0.77886016 | 3.94E-07 |
| Rbfox3 | 3121.6623 | 0.401828674 | 0.014845417 |
| Rbm24 | 206.31028 | 0.402149763 | 0.021958962 |
| Rbp4 | 579.04558 | 0.580096537 | 0.001160071 |
| Rc3h2 | 6255.1245 | 0.187715222 | 0.001583006 |
| Rcbtb2 | 1092.4454 | -0.245022826 | 0.049694547 |
| Rcl1 | 720.45469 | 0.26472586 | 0.023955603 |

| | | | |
|---|---|---|---|
| Rcor1 | 767.33546 | 0.296371666 | 0.006481941 |
| Rdx | 4647.3634 | -0.27843711 | 0.017498303 |
| Reep3 | 6288.8169 | -0.257984476 | 0.045038699 |
| Rem2 | 251.06741 | 0.98958813 | 4.20E-06 |
| Reps2 | 5387.0897 | 0.362224381 | 0.012995176 |
| Rft1 | 417.70693 | -0.302367383 | 0.033796151 |
| Rftn2 | 4134.6403 | -0.302291271 | 0.032865699 |
| Rfx4 | 2095.8136 | -0.287667083 | 0.018745489 |
| Rfx5 | 2070.8344 | -0.267584012 | 0.008160252 |
| Rgcc | 2254.9809 | -0.3021018 | 0.0387779 |
| RGD130300 | 4259.9525 | -0.191658778 | 0.043014325 |
| RGD130488 | 17582.501 | 0.284711841 | 0.032883247 |
| RGD130710 | 14251.036 | 0.215732857 | 0.024854151 |
| RGD130967 | 3039.309 | -0.285706468 | 0.015363767 |
| RGD131020 | 668.4748 | 0.611961967 | 0.000215463 |
| RGD131081 | 3841.9446 | 1.05860357 | 2.59E-11 |
| RGD131134 | 1021.0492 | -0.208486698 | 0.016850006 |
| RGD131173 | 1746.2362 | 0.343526275 | 0.010940557 |
| RGD131175 | 529.86641 | -0.406461967 | 0.008160252 |
| RGD131189 | 27.617082 | -1.931742097 | 0.000478194 |
| RGD156047 | 4763.3982 | 0.272801147 | 0.017218974 |
| RGD156193 | 2482.0812 | 0.305083317 | 0.036446513 |
| RGD156203 | 3694.8694 | 0.842738766 | 2.05E-06 |
| RGD156466 | 1129.3994 | 0.446560665 | 0.000686278 |
| RGD156553 | 971.71256 | -0.280532088 | 0.044816828 |
| RGD156577 | 1136.8223 | 0.229631115 | 0.035660344 |
| RGD156626 | 455.48586 | 0.302709104 | 0.022412337 |
| RGD156635 | 3427.7697 | 0.383344447 | 9.42E-09 |
| RGD621098 | 613.58286 | 0.241484298 | 0.023552299 |
| Rgs10 | 171.49859 | 0.318168445 | 0.049806825 |
| Rgs14 | 1310.0119 | 0.755694673 | 1.25E-05 |
| Rgs2 | 497.16301 | 0.602924191 | 3.53E-05 |
| Rgs6 | 906.57758 | -0.284590772 | 0.042867959 |
| Rgs7bp | 7798.9428 | 0.361353477 | 0.020302923 |
| Rgs8 | 3558.4927 | 0.251790997 | 0.02162047 |
| Rgs9 | 1459.0484 | 0.846984491 | 1.45E-09 |

| | | | |
|---|---|---|---|
| Rhoa | 10545.02 | -0.180458472 | 0.030326768 |
| Rhoc | 1179.6331 | -0.374484741 | 0.016609571 |
| Rictor | 4064.8834 | 0.18791946 | 0.028005102 |
| Rida | 1998.4714 | -0.388095891 | 0.00529415 |
| Ripk4 | 46.501908 | 1.030083371 | 0.001443865 |
| Rnf13 | 564.44524 | -0.352311476 | 0.005769639 |
| Rnf144b | 406.00247 | 0.575298293 | 8.64E-06 |
| Rnf150 | 969.96509 | 0.642157551 | 3.82E-05 |
| Rnf170 | 1254.4598 | -0.227484087 | 0.007388248 |
| Rnf180 | 2078.0018 | -0.219670509 | 0.032614756 |
| Rnf208 | 1719.4912 | 0.288382376 | 0.036731549 |
| Rnf215 | 829.03057 | -0.237777427 | 0.045298375 |
| Rock1 | 3271.7236 | -0.253832378 | 0.035780988 |
| Rock2 | 9952.4053 | 0.260581185 | 0.049111233 |
| Rph3al | 359.73006 | -0.4404492 | 0.008561381 |
| Rpp25 | 456.0303 | 0.476115224 | 0.004098051 |
| Rprd1a | 3846.616 | 0.149172968 | 0.024935642 |
| Rps27l | 393.95511 | -0.306565538 | 0.034157344 |
| Rps6ka5 | 2439.9488 | 0.25743463 | 0.040683587 |
| Rps6kl1 | 836.88715 | 0.253210803 | 0.042409834 |
| Rrm2 | 23.521267 | -0.607395308 | 0.012556283 |
| Runx1t1 | 1244.953 | 0.409429828 | 0.00341102 |
| Rxrg | 515.07639 | 0.442600333 | 0.018569156 |
| Rybp | 1840.7836 | 0.453236201 | 0.001016241 |
| Ryr2 | 6284.6214 | 0.280337266 | 0.042409834 |
| Sardh | 3125.9672 | -0.371037445 | 0.012615715 |
| Sash1 | 12393.601 | -0.381972035 | 0.000885358 |
| Sat1 | 2316.019 | -0.56781733 | 0.001354384 |
| Sbds | 3738.2385 | -0.286156259 | 0.005695151 |
| Sc5d | 9343.9116 | -0.29885272 | 0.012139612 |
| Scai | 1759.4572 | 0.220229638 | 0.042409834 |
| Scarb2 | 17419.086 | -0.274466637 | 0.026369562 |
| Scd | 2090.4977 | -1.0788752 | 3.13E-16 |
| Scd2 | 388621.48 | -0.439044119 | 0.003093041 |
| Scg2 | 14647.053 | 0.248273657 | 0.044901735 |
| Scn2b | 11764.251 | 0.328161013 | 0.009826484 |

| | | | |
|---|---|---|---|
| Scn3a | 3392.8778 | 0.249944639 | 0.048744284 |
| Scn3b | 5932.617 | 0.377102407 | 0.007849599 |
| Scn4b | 2014.9889 | 0.857490516 | 9.63E-05 |
| Scp2 | 10757.967 | -0.231266042 | 0.044776792 |
| Sdhb | 4372.6533 | -0.139703118 | 0.012995176 |
| Sdhc | 5691.104 | -0.223525011 | 0.029561749 |
| Sema3d | 502.55426 | -0.545802459 | 0.012572006 |
| Sema6d | 8924.0495 | -0.218538056 | 0.023779001 |
| Senp2 | 2358.5751 | 0.197915045 | 0.0483335 |
| Sepp1 | 24816.067 | -0.262389386 | 0.037411618 |
| Sepsecs | 559.46301 | -0.287919007 | 0.016667447 |
| Sept10 | 361.15331 | -0.346080205 | 0.024005389 |
| Sept2 | 10645.76 | -0.292668819 | 0.020181508 |
| Serac1 | 657.06572 | 0.196631381 | 0.042986698 |
| Serinc5 | 1414.2066 | -0.512327156 | 0.003614423 |
| Serpina3n | 32.908857 | -0.183713799 | 0.00493211 |
| Sertad4 | 480.33835 | 0.498986353 | 0.000166758 |
| Sesn2 | 1642.5966 | -0.428109698 | 0.003508329 |
| Setd1b | 2680.0393 | 0.15493195 | 0.04485385 |
| Setd4 | 243.73518 | -0.305310341 | 0.033077453 |
| Sez6 | 22950.099 | 0.290812183 | 0.01352317 |
| Sf1 | 7435.1477 | 0.14588143 | 0.012295122 |
| Sf3a1 | 4486.6548 | 0.138411951 | 0.0387779 |
| Sgip1 | 12824.342 | 0.258335737 | 0.035063489 |
| Sgpl1 | 1646.504 | -0.267359492 | 0.021762248 |
| Sgsm2 | 4225.4127 | 0.318495109 | 0.000539496 |
| Sgsm3 | 2158.654 | 0.254581876 | 0.013466378 |
| Sgtb | 6266.7139 | 0.298327144 | 0.026870318 |
| Sh2d5 | 1196.9292 | 0.81106454 | 5.91E-08 |
| Sh3bgrl | 8210.6397 | -0.269406166 | 0.024600206 |
| Sh3rf2 | 225.30407 | 1.203421904 | 3.34E-06 |
| Sh3rf3 | 1316.6404 | 0.448699419 | 0.000651735 |
| Shank3 | 5422.2097 | 0.362092741 | 0.008160252 |
| Shf | 446.38203 | 0.307758825 | 0.021805449 |
| Shisa4 | 2910.6445 | -0.326525621 | 0.025014064 |
| Shisa7 | 4650.3772 | 0.498869712 | 0.000306761 |

| | | | |
|---|---|---|---|
| Sik1 | 604.67329 | 0.459616783 | 0.007074336 |
| Sik3 | 10449.975 | 0.233131689 | 0.000736244 |
| Sipa1l1 | 9618.9252 | 0.492136033 | 4.43E-07 |
| Sirt2 | 6512.6913 | -0.299967469 | 0.00501884 |
| Skil | 3632.1194 | 0.525135935 | 1.72E-05 |
| Slc12a5 | 15073.371 | 0.296451294 | 0.023862019 |
| Slc1a3 | 125401.91 | -0.309665424 | 0.022811289 |
| Slc20a2 | 3574.0734 | -0.289907778 | 0.014034654 |
| Slc22a5 | 503.53029 | -0.443035266 | 0.006755965 |
| Slc24a4 | 1467.3622 | 0.862981399 | 0.000100949 |
| Slc25a1 | 2519.4625 | -0.331896785 | 0.016954915 |
| Slc25a13 | 208.93298 | -0.341538501 | 0.035284553 |
| Slc27a3 | 1055.0031 | -0.425880455 | 0.025567429 |
| Slc2a13 | 7083.1926 | 0.432529527 | 6.18E-05 |
| Slc31a1 | 683.87684 | -0.290979541 | 0.022478749 |
| Slc35d3 | 778.98143 | 0.406434512 | 0.009125973 |
| Slc35f3 | 690.16261 | 0.606455961 | 0.000446194 |
| Slc39a7 | 2820.9573 | -0.205647698 | 0.002802801 |
| Slc46a1 | 570.11426 | -0.313083994 | 0.028809824 |
| Slc4a10 | 7682.5487 | 0.380349864 | 0.012964143 |
| Slc4a11 | 154.97809 | 1.638710818 | 1.44E-07 |
| Slc6a11 | 11243.206 | -0.450983091 | 0.006817768 |
| Slc7a14 | 4591.7097 | 0.234061143 | 0.034157344 |
| Slc7a8 | 2178.6425 | 0.346507883 | 0.032534499 |
| Slit3 | 1732.5015 | 0.548689536 | 0.005146944 |
| Slitrk3 | 2228.756 | 0.309825256 | 0.014136511 |
| Slitrk5 | 2738.1196 | 0.317948969 | 0.014136511 |
| Slk | 4934.2524 | 0.219320878 | 0.009125973 |
| Slmap | 5597.66 | 0.273379307 | 0.000444688 |
| Smad3 | 1974.8942 | 0.582171856 | 0.000204176 |
| Smarcd1 | 3729.849 | 0.22370129 | 0.02082844 |
| Smo | 1356.8672 | -0.312631223 | 0.038522421 |
| Smoc1 | 1030.3276 | -0.401851248 | 0.011406973 |
| Smpdl3b | 266.23601 | 0.307082927 | 0.048679638 |
| Snap23 | 354.26261 | -0.368195041 | 0.008135492 |
| Snx18 | 1467.5728 | -0.327283551 | 0.002202231 |

| | | | |
|---|---|---|---|
| Snx22 | 30.719577 | -0.865058562 | 0.00504516 |
| Snx5 | 3618.1141 | -0.219487194 | 0.046344975 |
| Snx7 | 647.2753 | 0.580242443 | 0.000444688 |
| Sod3 | 3419.2051 | -0.353404731 | 0.026819388 |
| Soga1 | 2545.7651 | -0.264083033 | 0.040683587 |
| Son | 12109.783 | 0.189188654 | 0.009758988 |
| Sorbs2 | 4913.4493 | 0.521802072 | 0.001562402 |
| Sowaha | 1237.0933 | 0.752098269 | 4.72E-08 |
| Sowahc | 636.79581 | -0.32568804 | 0.011553088 |
| Sox10 | 631.76934 | -0.929181352 | 0.002885171 |
| Sox12 | 326.27687 | -0.341221692 | 0.015105299 |
| Sox2 | 6686.4449 | -0.322036588 | 0.012056077 |
| Sox21 | 1407.5069 | -0.359007499 | 0.021352829 |
| Sox4 | 878.70813 | -0.354506896 | 0.005882358 |
| Sox9 | 15629.858 | -0.349618311 | 0.00973374 |
| Sp4 | 1397.3802 | 0.226557915 | 0.008135492 |
| Sp9 | 616.4355 | 0.471942559 | 0.005283394 |
| Spata13 | 2348.1111 | 0.44205828 | 0.017014954 |
| Spata2L | 1339.5364 | 0.554838653 | 0.000627132 |
| Spock3 | 3001.5078 | 0.368380506 | 0.000687226 |
| Srf | 2145.2309 | 0.540969558 | 6.81E-07 |
| Ss18 | 1933.7463 | -0.271660922 | 0.028929763 |
| Ssbp2 | 3897.049 | 0.2327248 | 0.031681468 |
| Ssbp4 | 2237.2786 | 0.229652078 | 0.035660344 |
| Ssc5d | 421.27268 | -0.360931517 | 0.046294528 |
| Ssfa2 | 5660.8602 | -0.280329611 | 0.038763835 |
| Sspn | 1899.3857 | -0.475885515 | 0.002767089 |
| Sstr4 | 249.57199 | 0.525044221 | 0.009037893 |
| Ssx2ip | 2089.8629 | 0.205186585 | 0.034157344 |
| St6galnac4 | 796.26451 | -0.251864428 | 0.042409834 |
| St8sia3 | 1946.0583 | 0.362095858 | 0.022769774 |
| Stambpl1 | 956.7317 | 0.301176974 | 0.035935143 |
| Stap2 | 124.63297 | -0.470144363 | 0.02019114 |
| Steap3 | 1114.5178 | -0.369967717 | 0.01994124 |
| Stk10 | 269.66997 | -0.336816954 | 0.046118057 |
| Stk3 | 337.17626 | -0.316308091 | 0.041128265 |

| | | | |
|---|---|---|---|
| Stk32c | 2772.0851 | 0.661132219 | 4.20E-06 |
| Strip2 | 1665.1215 | 1.125512609 | 2.87E-10 |
| Strn4 | 4855.5801 | 0.31374698 | 0.006089657 |
| Stx16 | 2305.3216 | 0.145386747 | 0.043545615 |
| Stxbp5l | 1929.2225 | 0.385505112 | 0.011835246 |
| Suclg2 | 5661.535 | -0.346077132 | 0.021983493 |
| Sugp2 | 4682.7008 | 0.185823767 | 0.018735326 |
| Sumf2 | 641.07845 | -0.311563299 | 0.023779001 |
| Supt6h | 12193.274 | 0.209563446 | 0.009174755 |
| Susd5 | 64.885765 | -3.230407665 | 3.97E-13 |
| Sv2c | 5954.7528 | 0.4498408 | 0.001771635 |
| Sympk | 4140.2146 | 0.209532633 | 0.002229202 |
| Syn2 | 10429.232 | 0.308010564 | 0.049214614 |
| Syndig1 | 698.81159 | 0.512974192 | 0.00244492 |
| Syndig1l | 760.25662 | 0.590256252 | 2.25E-05 |
| Syne1 | 20158.938 | 0.432395766 | 0.00184316 |
| Syngap1 | 9138.192 | 0.463855836 | 0.004196452 |
| Synpo | 5643.9747 | 0.657342822 | 0.002165031 |
| Synpr | 3045.2986 | 0.680600038 | 2.27E-06 |
| Syt1 | 53488.614 | 0.333188927 | 0.026058845 |
| Syt10 | 462.58894 | 0.719696886 | 0.001016241 |
| Syt12 | 689.73998 | 0.415186352 | 0.007104799 |
| Syt16 | 890.99084 | 0.383260524 | 0.019576919 |
| Syt4 | 6365.5735 | 0.456156648 | 0.00184316 |
| Syt5 | 2432.5195 | 0.317888177 | 0.045256001 |
| Sytl5 | 526.32436 | 0.360276051 | 0.009199578 |
| Tac1 | 848.75202 | 1.283639028 | 4.36E-08 |
| Tac3 | 433.37899 | 1.092855469 | 1.23E-06 |
| Tanc2 | 13116.811 | 0.367910728 | 0.002139052 |
| Tbc1d8 | 686.06424 | 0.324332927 | 0.014337762 |
| Tbc1d9b | 8790.9717 | -0.144534464 | 0.040627784 |
| Tbcel | 2944.744 | -0.266141925 | 0.04087785 |
| Tcaf1 | 6747.8583 | 0.288584984 | 0.008045276 |
| Tcaim | 1355.3864 | -0.286980689 | 0.006332826 |
| Tcf20 | 8770.2316 | 0.257265175 | 0.002598587 |
| Tcf3 | 795.5515 | -0.279926875 | 0.015105299 |

| | | | |
|---|---|---|---|
| Tcf7l1 | 866.26427 | -0.274136533 | 0.038230165 |
| Tcf7l2 | 833.73116 | -0.317678781 | 0.029506935 |
| Tdg | 592.52803 | 0.240236963 | 0.03370062 |
| Tec | 132.21943 | 0.521222586 | 0.013468744 |
| Tecta | 225.04461 | -0.372440941 | 0.033494053 |
| Tesc | 384.05891 | 0.349802751 | 0.049474132 |
| Tet3 | 2356.4551 | 0.25238279 | 0.017900316 |
| Tex15 | 314.96223 | 0.418612254 | 0.015322734 |
| Tex30 | 51.144237 | 0.352756099 | 0.043014325 |
| Tfe3 | 1771.1003 | -0.285153145 | 0.017656703 |
| Thnsl2 | 587.56994 | -0.28314118 | 0.03549784 |
| Thrb | 3161.6113 | 0.246979928 | 0.041266518 |
| Thsd7a | 2283.7682 | 0.462384427 | 7.71E-06 |
| Tiam1 | 4399.8957 | 0.714840225 | 2.34E-12 |
| Tiam2 | 1842.6223 | 1.062702749 | 3.13E-10 |
| Timp4 | 1305.6719 | -0.511407664 | 0.003433683 |
| Tjp1 | 11662.506 | -0.306909437 | 0.012056077 |
| Tjp2 | 3706.4642 | -0.383625343 | 0.013314998 |
| Tkt | 10807.145 | -0.304675514 | 0.000132618 |
| Tlcd1 | 842.30187 | -0.515487864 | 0.009125973 |
| Tll2 | 38.727887 | 0.444663937 | 0.027746436 |
| Tm2d1 | 614.26107 | 0.287440919 | 0.011877151 |
| Tmc6 | 256.14111 | -0.367377091 | 0.032582414 |
| Tmed3 | 470.4484 | -0.259230349 | 0.039227522 |
| Tmed5 | 1258.5561 | -0.288692398 | 0.019845917 |
| Tmem134 | 232.27374 | -0.332732457 | 0.016850006 |
| Tmem158 | 1135.7868 | 0.704473257 | 2.04E-06 |
| Tmem181 | 1072.3352 | 0.250995719 | 0.045112143 |
| Tmem184b | 2649.2839 | 0.236308088 | 0.048372633 |
| Tmem198 | 1134.3845 | 0.337285819 | 0.017249043 |
| Tmem241 | 134.05608 | -0.32635005 | 0.048372633 |
| Tmem242 | 656.29318 | -0.220099473 | 0.02019114 |
| Tmem255b | 93.093339 | -1.061432865 | 0.000135274 |
| Tmem30a | 10189.733 | 0.170395105 | 0.020548211 |
| Tmem38a | 1273.855 | 0.282980555 | 0.017850733 |
| Tmem41a | 536.04534 | 0.527678904 | 0.003176112 |

| | | | |
|---|---|---|---|
| Tmod1 | 707.11525 | 0.329770271 | 0.046940386 |
| Tnfaip8 | 386.41849 | -0.522412294 | 0.000523443 |
| Tnfsf12 | 699.33877 | -0.298850025 | 0.048679638 |
| Tnpo3 | 2554.8693 | 0.163105601 | 0.043014325 |
| Tns1 | 4662.8582 | -0.272336763 | 0.046118057 |
| Tomm20 | 2837.4087 | 0.301342642 | 0.017931876 |
| Tomm70 | 6339.263 | 0.59648954 | 1.54E-08 |
| Top2a | 55.994137 | -0.728295408 | 0.008763887 |
| Tp53inp1 | 1327.0154 | -0.559100319 | 0.000204527 |
| Tpd52l1 | 625.93439 | 0.883559095 | 6.93E-08 |
| Tpm1 | 3440.809 | 0.324739532 | 0.035129295 |
| Tpp1 | 6472.4766 | -0.273319292 | 0.037015005 |
| Traf4 | 411.70288 | -0.765668574 | 6.83E-06 |
| Traf7 | 2442.0564 | -0.368763001 | 0.002568252 |
| Trappc2 | 956.88167 | 0.213854844 | 0.042926341 |
| Trerf1 | 1624.0566 | 0.503717632 | 3.27E-06 |
| Trim23 | 2155.5016 | 0.201571751 | 0.042867959 |
| Trim32 | 4461.7581 | 0.217365732 | 0.014473699 |
| Trim46 | 1103.4017 | 0.46027057 | 0.000808528 |
| Trim65 | 253.81143 | -0.295939382 | 0.048216151 |
| Trim66 | 1973.5964 | 0.26695915 | 0.041493703 |
| Trim7 | 310.99557 | -0.52357523 | 0.000484565 |
| Trrap | 8733.0154 | 0.251827441 | 0.005064182 |
| Tsc22d1 | 12580.725 | 0.252417892 | 0.011374676 |
| Tshz1 | 3892.2424 | 0.260371072 | 0.045123399 |
| Tspyl1 | 7243.0697 | 0.178836814 | 0.013485763 |
| Ttc12 | 433.73435 | -0.382051803 | 0.037101286 |
| Ttc19 | 2595.0162 | 0.270545168 | 0.03549784 |
| Ttc28 | 2688.1178 | -0.281099055 | 0.027150309 |
| Ttll12 | 932.31895 | -0.178956456 | 0.015053426 |
| Tubb2b | 10667.059 | -0.465015352 | 0.002640023 |
| Tubgcp5 | 709.60608 | 0.193055508 | 0.026007197 |
| Tvp23b | 1138.6895 | -0.275775405 | 0.032773528 |
| Tyms | 420.19679 | -0.213266814 | 0.047062871 |
| Tyro3 | 7490.6027 | 0.299101306 | 0.046294528 |
| Ube2q2l | 272.50384 | 0.422665133 | 0.000214478 |

| | | | |
|---|---|---|---|
| Ube2w | 1460.1971 | 0.293356075 | 0.016333805 |
| Ubr7 | 2964.1296 | -0.304814644 | 0.018745489 |
| Unc5c | 1537.4128 | 0.261827407 | 0.046922194 |
| Unc79 | 3523.8108 | 0.17443094 | 0.028195865 |
| Usp22 | 5436.3958 | 0.221768486 | 0.033878858 |
| Usp33 | 3946.911 | 0.179176235 | 0.04981703 |
| Usp5 | 2247.0338 | 0.187752367 | 0.046294528 |
| Usp6nl | 1971.4845 | -0.331297044 | 0.003761373 |
| Usp7 | 7346.5673 | 0.121313267 | 0.042409834 |
| Usp8 | 5639.8677 | -0.177630622 | 0.029981841 |
| Vangl2 | 1579.1646 | -0.475596066 | 0.000148188 |
| Vcan | 587.54216 | -0.336756358 | 0.045113165 |
| Vdac2 | 7513.1943 | -0.17985137 | 0.028885951 |
| Vegfb | 3059.4005 | -0.272349687 | 0.037015005 |
| Vimp | 1194.3371 | -0.250772459 | 0.033130748 |
| Vipr1 | 185.64924 | 0.356219877 | 0.048787673 |
| Vof16 | 76.733897 | -0.406468535 | 0.032534499 |
| Vps13c | 4360.561 | 0.185417651 | 0.044244665 |
| Vps29 | 1784.4659 | 0.215418417 | 0.011496112 |
| Vps36 | 560.7417 | -0.201991464 | 0.046294528 |
| Vps39 | 3579.8989 | 0.175094312 | 0.019086955 |
| Vps50 | 1172.8696 | 0.351255664 | 0.002492069 |
| Vsir | 3159.0412 | -0.333026322 | 0.02277008 |
| Vstm2a | 2743.4958 | 0.238794914 | 0.023968682 |
| Vstm5 | 184.89301 | 0.489198455 | 0.010956998 |
| Wasf1 | 5089.2266 | 0.57530685 | 0.0001068 |
| Wasl | 4551.5792 | 0.266613834 | 0.007849599 |
| Wdr17 | 859.13231 | 0.329614608 | 0.044886693 |
| Wdr46 | 812.16682 | 0.267569001 | 0.046377391 |
| Wdr47 | 5205.8645 | 0.272343591 | 0.027879331 |
| Wdr7 | 10718.902 | 0.28475357 | 0.010113729 |
| Wfdc1 | 313.90705 | -0.512220099 | 0.009938888 |
| Wfs1 | 4350.6227 | 1.386003607 | 5.56E-11 |
| Wipf3 | 4163.4712 | 0.881543619 | 4.70E-11 |
| Wipi1 | 1601.6287 | -0.302234865 | 0.021872246 |
| Wnt10a | 63.764898 | 0.408095728 | 0.032582414 |

| | | | |
|---|---|---|---|
| Wscd2 | 1135.9811 | 0.451608397 | 0.001358012 |
| Xbp1 | 3094.9583 | 0.228984965 | 0.047062871 |
| Xrcc6 | 1233.3885 | -0.373793546 | 0.001667804 |
| Yipf1 | 1425.0492 | -0.257748872 | 0.02584492 |
| Yipf3 | 2846.4464 | -0.159978215 | 0.047062871 |
| Ypel2 | 1433.0878 | 0.565777817 | 0.002357642 |
| Ywhaz | 52976.135 | 0.291085631 | 0.003409259 |
| Zbtb16 | 2230.5283 | 0.353459696 | 0.043014325 |
| Zbtb8a | 36.072165 | -0.466714039 | 0.024600206 |
| Zc3h12c | 1044.7435 | 0.283805223 | 0.013400572 |
| Zcchc14 | 4150.971 | 0.262796286 | 0.003718598 |
| Zdhhc14 | 1215.9705 | 0.357082415 | 0.011060835 |
| Zdhhc23 | 914.61827 | 0.736101008 | 5.40E-06 |
| Zfand3 | 3528.1621 | -0.262118235 | 0.02162047 |
| Zfhx2 | 2504.5617 | 0.448080867 | 0.010012392 |
| Zfp189 | 456.06276 | 0.375446691 | 0.000108801 |
| Zfp280d | 1837.8501 | 0.314578973 | 0.004682098 |
| Zfp281 | 1074.1631 | 0.257669852 | 0.038309023 |
| Zfp292 | 3159.3512 | 0.264240876 | 0.01027919 |
| Zfp385b | 1525.6002 | 0.370683116 | 0.005600125 |
| Zfp488 | 41.032872 | -0.495537278 | 0.021590955 |
| Zfp496 | 817.62017 | -0.264442867 | 0.034309374 |
| Zfp532 | 2609.9679 | 0.183058313 | 0.036442356 |
| Zfp536 | 2148.8074 | 0.353861282 | 0.01943578 |
| Zfp644 | 3577.8776 | 0.181970437 | 0.019571908 |
| Zfp704 | 2276.5746 | 0.195640997 | 0.046118057 |
| Zfp786 | 285.35838 | 0.516185687 | 0.000202208 |
| Zfp831 | 1030.8671 | 0.550487812 | 0.00554991 |
| Zfyve28 | 988.81069 | 0.512456093 | 0.000162766 |
| Zhx2 | 3568.477 | -0.338922532 | 0.012964143 |
| Zmat4 | 661.08589 | 0.330665349 | 0.004115071 |
| Zmpste24 | 2074.3177 | -0.212226583 | 0.043361782 |
| Zswim6 | 1347.7789 | 0.463881012 | 8.44E-05 |

| Gene | baseMean | log2FoldChange | adjusted p-value |
|---|---|---|---|
| Pdgfra | 1155.63326 | -4.414848112 | 2.02E-179 |
| Gpr17 | 429.645464 | -4.457141913 | 4.41E-66 |
| Col5a3 | 408.282566 | -3.506994464 | 1.87E-65 |
| Olig1 | 1419.39941 | -2.514965252 | 1.82E-46 |
| Cspg4 | 841.691171 | -2.560510877 | 1.36E-39 |
| Pmel | 92.0066508 | -3.988248706 | 4.05E-39 |
| Egr3 | 2019.2873 | 2.054010057 | 1.27E-26 |
| Arhgap31 | 233.844491 | -2.113204212 | 1.69E-26 |
| Olig2 | 459.569869 | -2.610700199 | 5.64E-23 |
| Ccnd1 | 337.958155 | -3.77058724 | 1.69E-21 |
| Cav2 | 159.187099 | -2.067296565 | 3.15E-20 |
| Susd5 | 64.8857647 | -3.742069049 | 3.84E-19 |
| Pde10a | 3294.37261 | 0.816464737 | 9.33E-18 |
| Calcrl | 209.840715 | -2.263499516 | 4.08E-17 |
| Fosb | 479.015685 | 2.284747647 | 5.43E-17 |
| Lzts3 | 5645.28743 | 0.741257804 | 6.41E-17 |
| Lmo7 | 1863.37024 | 0.802316729 | 2.43E-16 |
| Ncdn | 49443.7274 | 0.621504247 | 5.70E-16 |
| Mmp2 | 233.067716 | -2.237230299 | 8.07E-16 |
| Fam89a | 43.9667338 | -3.926773252 | 9.93E-16 |
| Necab1 | 858.280775 | 0.889602171 | 1.71E-14 |
| Cav1 | 211.17634 | -2.460099065 | 3.50E-14 |
| Egr4 | 809.143968 | 1.744600961 | 6.54E-14 |
| Crtc1 | 6526.28545 | 0.554725338 | 9.26E-14 |
| RGD131081 | 3841.94463 | 1.056431171 | 1.52E-13 |
| Wipf3 | 4163.47124 | 0.866207463 | 5.23E-13 |
| Anxa3 | 63.0133709 | -3.251770571 | 2.05E-12 |
| Ppp3ca | 15224.6719 | 0.701291871 | 9.23E-12 |
| Htr1b | 373.06482 | 0.970515503 | 2.32E-11 |
| Map3k13 | 1190.91659 | 0.660171706 | 2.49E-11 |
| LOC100125 | 4647.89023 | 0.67632984 | 5.14E-11 |
| Serinc5 | 1414.20656 | -1.144023635 | 5.51E-11 |
| Pllp | 400.742895 | -2.539563509 | 6.96E-11 |
| Per2 | 1924.18214 | 0.780887918 | 7.78E-11 |
| Sowaha | 1237.09334 | 0.802033934 | 7.89E-11 |

| | | | |
|---|---|---|---|
| Chst3 | 91.6645779 | -1.865861373 | 7.96E-11 |
| LOC103692 | 3641.69186 | -1.070958156 | 7.96E-11 |
| Sema3d | 502.554264 | -1.927864138 | 8.37E-11 |
| Mtus2 | 1064.89567 | 0.557058845 | 8.51E-11 |
| Sf1 | 7435.14772 | 0.302621893 | 9.24E-11 |
| Scd | 2090.49766 | -0.778126381 | 9.43E-11 |
| Sox10 | 631.769345 | -2.788446771 | 9.71E-11 |
| Lingo3 | 682.171578 | 0.778348226 | 1.05E-10 |
| Adcy5 | 11046.3316 | 0.694667036 | 2.00E-10 |
| Psd | 5479.79235 | 0.732757789 | 2.19E-10 |
| Slc20a2 | 3574.07343 | -0.687866644 | 2.31E-10 |
| Baiap2 | 4848.99774 | 0.95323288 | 2.78E-10 |
| Erbb3 | 431.174172 | -2.026275561 | 3.15E-10 |
| Camk2a | 6562.43796 | 1.014751413 | 3.15E-10 |
| Tiam1 | 4399.89572 | 0.582254486 | 3.93E-10 |
| Lamp5 | 691.211461 | 0.992240485 | 3.93E-10 |
| Cxcl10 | 1166.09689 | -7.349057247 | 4.17E-10 |
| Slfn13 | 121.300094 | -3.480629597 | 5.67E-10 |
| Mast3 | 6228.89606 | 0.784934879 | 6.72E-10 |
| Mink1 | 16183.7949 | 0.486558499 | 6.86E-10 |
| Lgi1 | 1904.38929 | 0.590705588 | 8.06E-10 |
| Bmp4 | 50.7248572 | -2.61055559 | 1.25E-09 |
| Cacnb1 | 4502.33246 | 0.622181225 | 1.86E-09 |
| Akap5 | 3627.1545 | 0.907433371 | 1.86E-09 |
| Mef2d | 8661.97222 | 0.513612658 | 1.88E-09 |
| Fabp4 | 21.1691981 | -2.356193772 | 1.89E-09 |
| Actn1 | 2628.80854 | 0.654814519 | 1.99E-09 |
| Cd74 | 399.900263 | -5.380195901 | 2.49E-09 |
| Pde2a | 8062.41256 | 0.77573134 | 2.49E-09 |
| Rasd2 | 4113.60306 | 0.838622319 | 2.58E-09 |
| Mctp1 | 1756.20847 | 0.725426051 | 2.92E-09 |
| C3 | 759.480914 | -5.381943368 | 3.04E-09 |
| RT1-Da | 89.7604895 | -6.019714383 | 3.11E-09 |
| Gria1 | 6627.95139 | 0.533631794 | 3.11E-09 |
| Fas | 457.064108 | -1.137246561 | 3.43E-09 |
| Gbp2 | 3314.88875 | -5.340124952 | 3.48E-09 |

| | | | |
|---|---|---|---|
| Drd1 | 2832.19853 | 0.861065087 | 3.55E-09 |
| Bcl11b | 6893.78295 | 0.74886912 | 3.73E-09 |
| Camk4 | 4664.33773 | 0.733774834 | 3.84E-09 |
| Phlda3 | 3194.94549 | -1.074178218 | 4.22E-09 |
| Fbxl16 | 20549.8516 | 0.744981067 | 5.04E-09 |
| Rsad2 | 1552.59826 | -5.42523779 | 6.82E-09 |
| Mid1ip1 | 6539.19217 | -0.730306024 | 6.82E-09 |
| Tomm70 | 6339.26295 | 0.558237978 | 7.31E-09 |
| Penk | 7963.79891 | 0.845074009 | 7.52E-09 |
| Rph3al | 359.730056 | -0.954905203 | 7.72E-09 |
| Hist1h1d | 1701.32692 | -0.415586359 | 9.67E-09 |
| Ptk2b | 15160.3425 | 0.611955987 | 9.81E-09 |
| Pdpk1 | 7395.94732 | 0.488781684 | 1.26E-08 |
| Phactr1 | 6881.17449 | 0.66475112 | 1.26E-08 |
| Lzts1 | 4551.15855 | 0.917613455 | 1.44E-08 |
| Mpeg1 | 82.5753746 | -2.689887241 | 1.45E-08 |
| Rab40b | 815.082579 | 0.824599042 | 1.45E-08 |
| Slamf8 | 128.785257 | -4.100169933 | 1.67E-08 |
| Isg15 | 824.884195 | -4.898094187 | 1.76E-08 |
| Ciita | 23.1925724 | -6.69127957 | 1.83E-08 |
| RT1-T24-4 | 160.960753 | -2.365727929 | 1.93E-08 |
| Chrm4 | 1175.47037 | 0.694848777 | 1.93E-08 |
| Ngef | 8600.21031 | 0.76801869 | 1.93E-08 |
| Pdzd2 | 2560.01989 | 0.698031776 | 2.10E-08 |
| Pde1b | 7933.39393 | 0.605778248 | 2.31E-08 |
| Rgs14 | 1310.0119 | 0.861282217 | 2.35E-08 |
| Map2k1 | 10686.7034 | 0.464012542 | 2.44E-08 |
| Dlgap2 | 2201.64765 | 0.798412243 | 2.52E-08 |
| Rbfox1 | 7146.48932 | 0.759294785 | 3.14E-08 |
| Phldb1 | 2053.63763 | -1.038644663 | 3.18E-08 |
| Kank1 | 1416.62366 | -0.848034657 | 3.18E-08 |
| Sipa1l1 | 9618.92521 | 0.495991476 | 3.18E-08 |
| Chst15 | 1098.85524 | 0.826642804 | 3.18E-08 |
| Tecpr2 | 3551.44199 | 0.295375604 | 3.31E-08 |
| Camk2b | 22178.9257 | 0.666125802 | 3.55E-08 |
| Tp53inp1 | 1327.01543 | -0.753525631 | 3.77E-08 |

| | | | |
|---|---|---|---|
| Fkbp5 | 1653.01721 | 0.833922348 | 3.96E-08 |
| Atp2b1 | 17736.1137 | 0.723502376 | 4.17E-08 |
| Gltp | 808.967573 | -0.901687306 | 4.57E-08 |
| LOC100910 | 870.493789 | -4.61664113 | 5.09E-08 |
| Matn4 | 38.9995914 | -3.063986855 | 5.28E-08 |
| Tnfrsf1a | 1149.25099 | -1.161951965 | 5.32E-08 |
| Foxp1 | 3780.12637 | 0.683157335 | 5.32E-08 |
| Kcnt1 | 1760.37371 | 0.448647299 | 5.40E-08 |
| Gabra4 | 2964.91745 | 0.537046407 | 5.48E-08 |
| Itpka | 1488.70924 | 0.985384545 | 5.93E-08 |
| Sat1 | 2316.01901 | -0.894876242 | 6.65E-08 |
| Gbp5 | 852.429277 | -4.016853572 | 6.73E-08 |
| Sh2d5 | 1196.92924 | 0.720507156 | 7.24E-08 |
| Dbi | 2132.76747 | -0.682317687 | 7.61E-08 |
| Fam167a | 412.41325 | -0.894441326 | 8.14E-08 |
| Nab2 | 706.245489 | 0.800185224 | 8.86E-08 |
| Rgs2 | 497.163008 | 0.701237081 | 9.21E-08 |
| RGD131189 | 27.6170823 | -3.445851633 | 9.36E-08 |
| Tubb2b | 10667.0586 | -0.77724682 | 9.36E-08 |
| Gpr158 | 9158.69696 | 0.434276562 | 9.36E-08 |
| Hpca | 6064.38617 | 0.804589221 | 9.36E-08 |
| Egr2 | 252.42617 | 2.009210136 | 9.39E-08 |
| Afap1l2 | 626.43037 | -0.988503541 | 9.96E-08 |
| Sesn2 | 1642.59659 | -0.734342026 | 1.10E-07 |
| Agap2 | 27785.813 | 0.779166738 | 1.13E-07 |
| Camkv | 7498.15501 | 0.789325099 | 1.13E-07 |
| Grasp | 509.620334 | 1.226833671 | 1.13E-07 |
| Usp18 | 184.589169 | -4.903536668 | 1.34E-07 |
| Ncoa5 | 2199.04975 | 0.394979237 | 1.43E-07 |
| Gpr88 | 3714.63061 | 0.800051596 | 1.44E-07 |
| Actn2 | 1015.81647 | 0.714273184 | 1.48E-07 |
| Lyst | 2198.76563 | 0.444892396 | 1.54E-07 |
| Gng12 | 2519.79952 | -0.760553628 | 1.68E-07 |
| Ifit3 | 654.081821 | -6.853423997 | 1.77E-07 |
| Tpd52l1 | 625.934392 | 0.757105032 | 1.93E-07 |
| Ppm1h | 1936.30721 | 0.479924211 | 1.97E-07 |

| | | | |
|---|---|---|---|
| Arpp21 | 9034.8647 | 0.678846937 | 2.05E-07 |
| Aldh4a1 | 2904.56169 | -0.63484293 | 2.07E-07 |
| Pitpnm3 | 2179.25842 | 0.56782538 | 2.14E-07 |
| Rnf150 | 969.965091 | 0.728001219 | 2.22E-07 |
| Rps27l | 393.955113 | -0.76623143 | 2.45E-07 |
| Irf1 | 1126.89521 | -3.446535964 | 2.62E-07 |
| Cdk17 | 5451.35761 | 0.463607097 | 2.62E-07 |
| Fabp7 | 2546.45363 | -2.091372243 | 2.73E-07 |
| Muc1 | 69.0915728 | -1.600681237 | 2.93E-07 |
| Ctss | 45.5782557 | -2.574409402 | 3.44E-07 |
| Clec2g | 43.5150843 | -5.876469857 | 3.55E-07 |
| LOC360231 | 127.92318 | -2.385868512 | 3.58E-07 |
| Laptm5 | 35.1879249 | -2.012152971 | 3.61E-07 |
| MGC10882 | 331.800842 | -5.090897186 | 3.70E-07 |
| C2cd2l | 8273.37693 | 0.594739653 | 3.77E-07 |
| Plppr4 | 7182.4385 | 0.643778285 | 3.86E-07 |
| Cacnb4 | 6784.6269 | 0.695892436 | 3.97E-07 |
| Irf7 | 549.632518 | -3.951913004 | 4.00E-07 |
| Irgm | 1052.40794 | -2.966527596 | 4.09E-07 |
| Dll1 | 134.025108 | -0.85322845 | 4.16E-07 |
| Igtp | 749.325396 | -3.577235097 | 4.33E-07 |
| Scn4b | 2014.9889 | 1.010366762 | 4.41E-07 |
| Dmtf1 | 2801.04084 | 0.462426353 | 4.43E-07 |
| RGD156024 | 5266.66707 | -0.652001598 | 4.68E-07 |
| Cdkn1a | 3020.39275 | -1.122801179 | 4.74E-07 |
| Apol9a | 183.925402 | -5.599057122 | 5.01E-07 |
| Shank3 | 5422.20968 | 0.653474055 | 5.04E-07 |
| Fgfrl1 | 1339.5509 | -0.936913944 | 5.35E-07 |
| Snx22 | 30.7195771 | -2.657421625 | 5.66E-07 |
| Ash1l | 13999.7561 | 0.268331245 | 5.83E-07 |
| Wfs1 | 4350.62275 | 0.922091029 | 5.83E-07 |
| Birc3 | 112.743039 | -2.670662417 | 6.34E-07 |
| Ptgs2 | 103.080167 | 1.419146857 | 6.34E-07 |
| Rb1 | 2210.32106 | -0.459840275 | 6.38E-07 |
| Trhde | 1041.9358 | 0.691167761 | 6.58E-07 |
| Pde7b | 3728.41482 | 0.584521111 | 6.93E-07 |

| | | | |
|---|---|---|---|
| Thrb | 3161.61132 | 0.573863773 | 7.51E-07 |
| Jph4 | 8618.68149 | 0.639571903 | 7.84E-07 |
| C1qa | 43.7040878 | -2.386926687 | 7.88E-07 |
| RGD156203 | 3694.86941 | 0.776108333 | 8.31E-07 |
| Cpne5 | 3765.75619 | 0.601623747 | 8.37E-07 |
| Bcas1 | 1155.19355 | -1.210308454 | 8.45E-07 |
| Parp12 | 893.317408 | -1.532487497 | 8.70E-07 |
| Tmem158 | 1135.78677 | 0.653825858 | 8.99E-07 |
| Dopey2 | 2792.02472 | 0.575991171 | 9.05E-07 |
| Irgm2 | 2107.19683 | -3.493910524 | 9.08E-07 |
| Carm1 | 3054.45648 | 0.32904419 | 9.22E-07 |
| Josd1 | 1290.91452 | 0.515433468 | 9.26E-07 |
| C1qb | 28.8617985 | -3.302271179 | 9.31E-07 |
| Dact2 | 884.263064 | 0.561353233 | 9.34E-07 |
| Nlrc5 | 371.143833 | -3.600423873 | 9.70E-07 |
| Dgki | 5383.79456 | 0.559428848 | 9.87E-07 |
| Cdc42ep1 | 577.920085 | -1.117939452 | 1.08E-06 |
| Gucy1b3 | 5076.24298 | 0.428144917 | 1.08E-06 |
| Dbx2 | 734.781971 | -0.696668877 | 1.09E-06 |
| LOC100912 | 2266.31043 | 0.716830522 | 1.15E-06 |
| Icam5 | 4448.55828 | 0.875382398 | 1.16E-06 |
| Oasl2 | 1278.80993 | -3.850439228 | 1.16E-06 |
| Lrrc7 | 4171.51316 | 0.552997664 | 1.16E-06 |
| Shisa7 | 4650.37723 | 0.61695335 | 1.16E-06 |
| Abcb1b | 271.899365 | -1.944142044 | 1.17E-06 |
| Ccsap | 1720.51226 | 0.655294419 | 1.17E-06 |
| Kcnab1 | 4144.76181 | 0.660474757 | 1.27E-06 |
| Cd38 | 1563.30939 | -0.759115965 | 1.29E-06 |
| Tap1 | 1243.78536 | -3.345317696 | 1.36E-06 |
| Cxcl16 | 263.329375 | -2.610485689 | 1.37E-06 |
| C1r | 2221.79643 | -2.885196229 | 1.39E-06 |
| Mki67 | 45.0411507 | -2.542119788 | 1.44E-06 |
| Rab7b | 294.476193 | -0.876701479 | 1.54E-06 |
| Cdk5r1 | 2652.17572 | 0.56517739 | 1.55E-06 |
| Cyfip2 | 30595.4694 | 0.691650131 | 1.55E-06 |
| MGC10556 | 173.936211 | -4.433261026 | 1.59E-06 |

| | | | |
|---|---|---|---|
| Sp110 | 247.371773 | -3.156530966 | 1.60E-06 |
| Shc4 | 137.120764 | -1.112428358 | 1.60E-06 |
| Rbp1 | 1268.66998 | -0.976407042 | 1.61E-06 |
| Cacna1g | 3525.45814 | 0.625035127 | 1.61E-06 |
| Psmb9 | 660.88528 | -3.518794003 | 1.61E-06 |
| Ddah1 | 9103.34815 | -0.314781378 | 1.63E-06 |
| Lims2 | 14.9810916 | -2.785708243 | 1.83E-06 |
| RGD156466 | 1129.39944 | 0.579788552 | 1.83E-06 |
| RGD130518 | 116.252053 | -5.190693767 | 1.95E-06 |
| Nkiras1 | 2509.25884 | 0.725892672 | 2.03E-06 |
| Kctd13 | 1910.79333 | 0.553377944 | 2.17E-06 |
| Man1a1 | 1133.75534 | 0.660261964 | 2.20E-06 |
| Adcy9 | 4331.74213 | 0.725530732 | 2.24E-06 |
| Serping1 | 3137.37294 | -3.651754175 | 2.25E-06 |
| Dhcr7 | 1182.60125 | -0.645129425 | 2.26E-06 |
| B3gnt2 | 565.771041 | 0.503452611 | 2.26E-06 |
| Cnksr2 | 6918.24524 | 0.770846181 | 2.26E-06 |
| Abhd4 | 13404.8402 | -0.591647388 | 2.31E-06 |
| C1s | 1421.41381 | -3.082648956 | 2.36E-06 |
| LOC103689 | 181.172736 | -3.78851827 | 2.37E-06 |
| Kctd1 | 2811.18741 | 0.468002027 | 2.41E-06 |
| Grm4 | 1250.15283 | 0.775357802 | 2.42E-06 |
| Homer1 | 3421.07906 | 0.785325412 | 2.46E-06 |
| Tlr2 | 45.3117248 | -2.938387468 | 2.58E-06 |
| Parp14 | 1329.1177 | -3.016410895 | 2.60E-06 |
| Jph3 | 8079.94635 | 0.601028179 | 2.66E-06 |
| Tiam2 | 1842.6223 | 0.709616498 | 2.66E-06 |
| Tead3 | 290.260309 | -0.701806032 | 2.79E-06 |
| Trim46 | 1103.40165 | 0.597061129 | 2.79E-06 |
| Chn1 | 10971.8271 | 0.714873631 | 2.79E-06 |
| Stat1 | 7129.89804 | -2.759502123 | 2.91E-06 |
| Ankrd63 | 2947.42508 | 0.668353682 | 2.99E-06 |
| Traf7 | 2442.0564 | -0.529383337 | 3.05E-06 |
| Cnp | 7190.27867 | -1.006428371 | 3.08E-06 |
| Rem2 | 251.067412 | 0.874822961 | 3.08E-06 |
| Ptpn5 | 3400.46169 | 0.524742911 | 3.24E-06 |

| | | | |
|---|---|---|---|
| Slc4a11 | 154.978089 | 1.246569465 | 3.27E-06 |
| Top2a | 55.9941371 | -2.586732782 | 3.32E-06 |
| Pdpn | 1679.10095 | -0.963711392 | 3.53E-06 |
| Ifit2 | 656.553832 | -4.213770584 | 3.60E-06 |
| Rnf144b | 406.00247 | 0.542264744 | 3.60E-06 |
| Carhsp1 | 1683.644 | -0.969872298 | 3.67E-06 |
| Slc35f4 | 192.482114 | 0.921320959 | 3.88E-06 |
| Clvs1 | 645.502845 | 0.624436308 | 3.93E-06 |
| Ebpl | 217.842535 | -0.742537971 | 3.94E-06 |
| Casp4 | 238.590496 | -2.989599141 | 4.03E-06 |
| Aim2 | 31.2109835 | -3.249859804 | 4.12E-06 |
| Ano3 | 1293.23444 | 0.847713947 | 4.12E-06 |
| Actr3b | 587.153456 | 0.775851109 | 4.17E-06 |
| Dynlt1 | 322.246218 | -0.733381141 | 4.34E-06 |
| Camta2 | 7881.12512 | 0.521915605 | 4.36E-06 |
| Psmb10 | 908.504571 | -3.361325391 | 4.39E-06 |
| Cers4 | 2664.9717 | -0.38359062 | 4.39E-06 |
| Klhl29 | 2786.96189 | 0.524183494 | 4.39E-06 |
| Spata2L | 1339.5364 | 0.682384977 | 4.39E-06 |
| Kcnj4 | 1596.16011 | 0.943482485 | 4.39E-06 |
| Dlg4 | 17979.9797 | 0.569638959 | 4.49E-06 |
| Asic4 | 1032.60352 | 0.820598614 | 4.49E-06 |
| Tmcc3 | 2276.49403 | -0.662182889 | 4.67E-06 |
| Stk32c | 2772.08507 | 0.592090436 | 4.82E-06 |
| Gal3st1 | 129.466225 | -0.923162794 | 4.98E-06 |
| Neurl1 | 3591.15568 | 0.683552082 | 5.00E-06 |
| Rsu1 | 714.729452 | -0.600883634 | 5.04E-06 |
| Fnbp1 | 2804.79596 | -0.561787141 | 5.12E-06 |
| Nt5dc3 | 2400.27653 | 0.74083246 | 5.19E-06 |
| Mx1 | 1335.25996 | -5.5706949 | 5.20E-06 |
| Kcnj2 | 432.34082 | 0.872573738 | 5.28E-06 |
| Tubgcp5 | 709.606084 | 0.341822638 | 5.34E-06 |
| Col22a1 | 48.6550413 | -2.929767427 | 5.38E-06 |
| Ppp3r1 | 12418.2594 | 0.517878314 | 5.54E-06 |
| Braf | 4186.25443 | 0.42256834 | 5.72E-06 |
| Gtse1 | 545.339897 | -0.815228866 | 5.81E-06 |

| | | | |
|---|---|---|---|
| Kcnh1 | 3568.82604 | 0.756389283 | 5.98E-06 |
| Bcl3 | 78.3067026 | -3.7752537 | 6.02E-06 |
| Hdac4 | 4704.06074 | 0.361617369 | 6.04E-06 |
| Otof | 1343.53061 | 0.864170179 | 6.13E-06 |
| Nova2 | 2806.16497 | 0.463006209 | 6.29E-06 |
| Eva1b | 134.248673 | -1.878094578 | 6.31E-06 |
| Gabrd | 514.881439 | 0.84827519 | 6.59E-06 |
| Rhbdf2 | 96.3642197 | -2.166624824 | 6.78E-06 |
| Mcam | 330.118888 | -2.552588561 | 6.95E-06 |
| Dhx58 | 426.3628 | -2.712171828 | 7.05E-06 |
| Ube2l6 | 219.010259 | -2.132757537 | 7.05E-06 |
| Sh3bp4 | 1122.17172 | -0.508325099 | 7.11E-06 |
| Gpr12 | 401.146316 | 0.617135542 | 7.17E-06 |
| Gpr155 | 1318.16476 | 0.397549867 | 7.25E-06 |
| Man2b1 | 901.561406 | -0.449720592 | 7.34E-06 |
| Card11 | 32.8407152 | -2.858922182 | 7.35E-06 |
| Arel1 | 6743.67069 | 0.361961753 | 7.35E-06 |
| Nbea | 10414.9369 | 0.449298221 | 7.39E-06 |
| Fa2h | 271.975178 | -2.399253532 | 7.47E-06 |
| LOC685067 | 171.398917 | -4.527491298 | 7.66E-06 |
| Tmem176b | 794.675961 | -1.032687021 | 7.66E-06 |
| Kcnq5 | 1788.70716 | 0.831714162 | 7.66E-06 |
| Camkk1 | 3175.21156 | 0.575932489 | 7.88E-06 |
| Ppdpf | 1391.86389 | -0.370019719 | 8.21E-06 |
| Rab13 | 972.533862 | -1.016623935 | 8.28E-06 |
| Kcnf1 | 2779.3736 | 0.845363018 | 8.49E-06 |
| Tjp2 | 3706.46418 | -0.682434964 | 8.49E-06 |
| Chst5 | 18.9045563 | -3.166026697 | 8.75E-06 |
| Cdkl5 | 3633.09861 | 0.562210314 | 9.03E-06 |
| Synpo | 5643.9747 | 0.907633608 | 9.08E-06 |
| Cxcl11 | 54.8380141 | -6.282918439 | 9.23E-06 |
| Cxcl9 | 291.163003 | -6.267105777 | 9.36E-06 |
| Glipr2 | 203.874919 | -3.286645235 | 9.40E-06 |
| Rnf213 | 3255.83592 | -2.657501889 | 9.46E-06 |
| Vcam1 | 6508.29208 | -0.837337712 | 9.77E-06 |
| Apol3 | 64.7870671 | -4.400545194 | 1.00E-05 |

| | | | |
|---|---|---|---|
| Agmo | 59.9576502 | -1.164533954 | 1.06E-05 |
| Rtp4 | 293.410823 | -3.055187428 | 1.07E-05 |
| Mov10 | 309.332013 | -1.518885132 | 1.07E-05 |
| Srgap1 | 1650.27244 | -0.440418602 | 1.18E-05 |
| Dusp8 | 2505.93121 | 0.470008714 | 1.19E-05 |
| Irf9 | 778.035126 | -2.067122033 | 1.25E-05 |
| Gcc2 | 3344.77494 | 0.391185074 | 1.25E-05 |
| Cfh | 49.2542547 | -1.911971073 | 1.26E-05 |
| Mpa2l | 1582.68135 | -2.790714587 | 1.27E-05 |
| Iqsec2 | 4548.33833 | 0.510998786 | 1.27E-05 |
| Slmap | 5597.66003 | 0.300455947 | 1.31E-05 |
| Slc1a4 | 3751.21065 | -0.585627679 | 1.31E-05 |
| Hyal1 | 4760.77445 | -0.557103161 | 1.37E-05 |
| Srsf5 | 2973.87176 | -0.468589899 | 1.37E-05 |
| Myo5b | 4263.34573 | 0.745980276 | 1.39E-05 |
| Kctd16 | 1440.14634 | 0.610487329 | 1.41E-05 |
| Akap11 | 17150.4224 | 0.409257816 | 1.46E-05 |
| Celf5 | 7596.15815 | 0.596663646 | 1.46E-05 |
| Sp1 | 3271.21692 | -0.486629903 | 1.48E-05 |
| Cp | 2211.75167 | -3.004687301 | 1.49E-05 |
| Arpc1b | 94.7367069 | -1.808653957 | 1.53E-05 |
| Zfp189 | 456.06276 | 0.381629663 | 1.54E-05 |
| Syngap1 | 9138.19201 | 0.66786344 | 1.55E-05 |
| Lgals3bp | 1408.8857 | -2.498890802 | 1.62E-05 |
| Frmd8 | 1606.26342 | -0.618253031 | 1.68E-05 |
| C1qtnf9 | 23.3986125 | -3.428829493 | 1.69E-05 |
| Gucy1a2 | 1398.34559 | 0.437105822 | 1.69E-05 |
| Rasgrp2 | 872.960003 | 0.546522727 | 1.74E-05 |
| Gabrb3 | 2772.61503 | 0.605901217 | 1.74E-05 |
| Rapgef4 | 3070.08399 | 0.583447402 | 1.75E-05 |
| Itgam | 35.7052546 | -2.350191677 | 1.76E-05 |
| Pros1 | 594.23454 | -0.648721043 | 1.78E-05 |
| Fabp5 | 2497.30362 | -0.545256825 | 1.78E-05 |
| Cacna1c | 3268.12629 | 0.50967694 | 1.81E-05 |
| Cnnm1 | 4371.49542 | 0.540820286 | 1.82E-05 |
| Trim25 | 327.077918 | -2.388924711 | 1.85E-05 |

| | | | |
|---|---|---|---|
| Slc39a1 | 4357.56986 | -0.651785836 | 1.85E-05 |
| Gria2 | 14720.6192 | 0.504524685 | 1.90E-05 |
| Sp140 | 60.7167378 | -3.213427382 | 1.90E-05 |
| Creb5 | 509.715924 | -1.160421406 | 1.94E-05 |
| Ttyh2 | 2088.55984 | -0.706957547 | 1.95E-05 |
| Uba7 | 293.421499 | -3.577268166 | 1.96E-05 |
| Fam189b | 1456.64055 | 0.563996431 | 1.96E-05 |
| RT1-CE7 | 198.241296 | -2.964464621 | 2.19E-05 |
| Skil | 3632.11936 | 0.474273662 | 2.25E-05 |
| Srebf1 | 15225.0694 | -0.588748233 | 2.30E-05 |
| Tsc22d1 | 12580.725 | 0.375319974 | 2.31E-05 |
| Hnrnpdl | 5067.41606 | -0.337570483 | 2.32E-05 |
| Bcr | 8939.66236 | 0.28066151 | 2.32E-05 |
| RGD131117 | 1746.23625 | 0.544037986 | 2.33E-05 |
| Lcor | 1465.47852 | 0.380520876 | 2.34E-05 |
| Mcm10 | 11.1545189 | -3.511005056 | 2.37E-05 |
| Adgrb3 | 5829.30327 | 0.256077994 | 2.48E-05 |
| LOC102555 | 147.318055 | -3.403399696 | 2.50E-05 |
| Psmb8 | 390.994672 | -3.079289236 | 2.51E-05 |
| Fkbp1a | 7726.57637 | 0.522546395 | 2.51E-05 |
| C1qc | 29.3807099 | -2.77025068 | 2.55E-05 |
| Pla2g16 | 1994.94125 | -0.911798424 | 2.55E-05 |
| Hist1h2ac | 20.3126398 | -1.954405378 | 2.57E-05 |
| Ankfy1 | 5791.85052 | -0.435701849 | 2.58E-05 |
| Dcbld2 | 1770.60826 | 0.415086957 | 2.64E-05 |
| Tmem100 | 1593.25354 | -0.634006249 | 2.64E-05 |
| Il18bp | 676.577858 | -3.010956045 | 2.67E-05 |
| Csrnp2 | 1153.16386 | 0.390745458 | 2.70E-05 |
| Xbp1 | 3094.95831 | 0.458619905 | 2.70E-05 |
| C2 | 382.836401 | -3.312110264 | 2.72E-05 |
| Mapk1 | 17196.4421 | 0.371610194 | 2.72E-05 |
| Trerf1 | 1624.05657 | 0.416221815 | 2.72E-05 |
| Dlg2 | 15716.3543 | 0.430280496 | 2.74E-05 |
| Bax | 585.457821 | -0.56687269 | 2.75E-05 |
| Parp10 | 656.479017 | -1.32529714 | 2.75E-05 |
| Npepl1 | 897.167684 | -0.56356244 | 2.79E-05 |

| | | | |
|---|---|---|---|
| Cacna1e | 13118.69 | 0.666843382 | 2.80E-05 |
| Ppp1r12b | 6978.43972 | 0.452443463 | 2.85E-05 |
| Syndig1l | 760.256625 | 0.528762981 | 2.86E-05 |
| Steap4 | 12.5748474 | -4.479912541 | 2.86E-05 |
| Pld4 | 19.0072661 | -3.32918448 | 2.87E-05 |
| Ripk1 | 767.374413 | -0.772132289 | 2.87E-05 |
| Ppp1r13b | 3330.00459 | 0.436581883 | 2.87E-05 |
| Tln2 | 11408.3663 | 0.491499554 | 2.87E-05 |
| Prkcz | 5042.26278 | 0.524580304 | 2.88E-05 |
| Ap3b2 | 6147.38834 | 0.339000989 | 2.91E-05 |
| Csf1r | 98.3156864 | -1.328912243 | 2.99E-05 |
| Rhbg | 15.8397086 | -4.532944867 | 3.03E-05 |
| Bahd1 | 2360.70228 | 0.417598068 | 3.07E-05 |
| Gnb5 | 2735.18453 | 0.371017059 | 3.15E-05 |
| Socs3 | 156.923705 | -3.321429469 | 3.19E-05 |
| Lap3 | 2642.78793 | -0.967698403 | 3.21E-05 |
| Gjb1 | 58.7264429 | -2.651576872 | 3.22E-05 |
| LOC102549 | 37.5943061 | -2.217153752 | 3.22E-05 |
| Rgs9 | 1459.04836 | 0.52636572 | 3.22E-05 |
| Ifitm3 | 358.296968 | -4.205046003 | 3.24E-05 |
| Mchr1 | 377.808195 | 0.706899989 | 3.26E-05 |
| Pak6 | 2507.68327 | 0.711413387 | 3.31E-05 |
| Chadl | 2261.37009 | -0.708601555 | 3.37E-05 |
| Abtb2 | 307.325191 | -0.950103953 | 3.42E-05 |
| Gfod1 | 4952.96489 | 0.668547584 | 3.42E-05 |
| Slc39a10 | 5025.60065 | 0.42206031 | 3.44E-05 |
| Tmem98 | 284.756604 | -0.692021379 | 3.47E-05 |
| Nptn | 10834.3263 | 0.406529988 | 3.47E-05 |
| Mog | 591.995931 | -2.424671529 | 3.48E-05 |
| Fam65b | 3848.12021 | 0.530485114 | 3.48E-05 |
| Deptor | 1726.77119 | 0.574910901 | 3.56E-05 |
| Icam1 | 272.771319 | -1.519415032 | 3.67E-05 |
| Pgpep1 | 866.84484 | -0.45377849 | 3.68E-05 |
| Psme2 | 791.345592 | -1.187966523 | 3.78E-05 |
| Opalin | 586.628325 | -1.968027208 | 3.82E-05 |
| Traf4 | 411.702877 | -0.622290592 | 3.82E-05 |

| | | | |
|---|---|---|---|
| Nr4a3 | 2200.1121 | 0.86320903 | 3.84E-05 |
| Meis2 | 6008.15556 | 0.382808731 | 3.85E-05 |
| Vsir | 3159.04122 | -0.603417528 | 3.88E-05 |
| Tspan15 | 186.553243 | -0.758213084 | 3.88E-05 |
| Arhgap42 | 1008.27243 | -0.645418648 | 3.89E-05 |
| Fam84a | 4411.39087 | 0.428745988 | 3.89E-05 |
| S1pr3 | 877.11175 | -2.729090695 | 3.91E-05 |
| Hsph1 | 9735.40421 | 0.568514118 | 4.01E-05 |
| Ptx3 | 122.109925 | -3.088864852 | 4.01E-05 |
| Grin2b | 26375.5598 | 0.628026639 | 4.01E-05 |
| Lgi4 | 13635.611 | -0.581695369 | 4.02E-05 |
| Ablim2 | 2775.40937 | 0.367402943 | 4.02E-05 |
| Egr1 | 3912.62317 | 1.147231426 | 4.02E-05 |
| Wasl | 4551.57918 | 0.366560967 | 4.04E-05 |
| Oprk1 | 351.016518 | 0.749626438 | 4.06E-05 |
| Csf1 | 1070.23774 | -1.569176211 | 4.08E-05 |
| Slc12a7 | 540.737861 | -1.112194363 | 4.08E-05 |
| Cald1 | 290.147017 | -0.870957571 | 4.09E-05 |
| Fxyd1 | 2311.50733 | -0.605550799 | 4.10E-05 |
| Mmp14 | 1030.98672 | -0.835651439 | 4.12E-05 |
| Slc4a1 | 55.7714254 | -4.3382929 | 4.12E-05 |
| Snapc2 | 1848.80637 | -0.469722977 | 4.12E-05 |
| Tyms | 420.196786 | -0.406869379 | 4.23E-05 |
| RGD156635 | 3427.76968 | 0.253514088 | 4.33E-05 |
| Myo1f | 11.9247039 | -2.922505032 | 4.36E-05 |
| Lmtk2 | 9717.93639 | 0.583692321 | 4.38E-05 |
| Cfb | 343.138001 | -4.536283398 | 4.43E-05 |
| Fn1 | 282.236898 | -1.368794407 | 4.43E-05 |
| Lrrk2 | 3435.16056 | 0.448319907 | 4.43E-05 |
| Dgkh | 3759.48178 | 0.626714082 | 4.43E-05 |
| Sh3rf2 | 225.304071 | 0.888890446 | 4.43E-05 |
| Itpr1 | 18327.4028 | 0.787728276 | 4.44E-05 |
| Parp9 | 714.011554 | -2.07172054 | 4.45E-05 |
| Cbx6 | 12049.2298 | 0.369121125 | 4.50E-05 |
| Slc35b2 | 990.898898 | -0.34789816 | 4.57E-05 |
| Elf1 | 380.850875 | -0.76015811 | 4.58E-05 |

| | | | |
|---|---|---|---|
| Cds2 | 5574.76797 | 0.298346598 | 4.60E-05 |
| Gga3 | 1946.86904 | 0.502839525 | 4.60E-05 |
| Kif14 | 6.21570306 | -4.575014358 | 4.65E-05 |
| XAF1 | 559.382927 | -2.280253582 | 4.65E-05 |
| Ltbr | 1528.69623 | -0.651176752 | 4.69E-05 |
| Gtf3c1 | 7812.40192 | 0.276642045 | 4.69E-05 |
| Pdyn | 2096.76556 | 0.655272251 | 4.69E-05 |
| Slc2a13 | 7083.19258 | 0.40071778 | 4.72E-05 |
| Tac1 | 848.752018 | 0.803322773 | 4.72E-05 |
| Tnfaip8 | 386.418492 | -0.561467695 | 4.76E-05 |
| March11 | 75.9253572 | 0.794563516 | 4.80E-05 |
| Ggta1 | 789.101624 | -0.86509351 | 4.80E-05 |
| Cdh3 | 26.9028896 | -2.397510815 | 4.85E-05 |
| Lpcat3 | 1190.8808 | -0.532241036 | 4.87E-05 |
| Ids | 6005.13221 | 0.373367764 | 4.94E-05 |
| RGD130842 | 1428.91202 | -0.325116337 | 4.97E-05 |
| Sgpl1 | 1646.50404 | -0.437857647 | 4.97E-05 |
| Srrm2 | 28423.6294 | 0.217541981 | 5.06E-05 |
| Misp | 22.6854738 | -3.537428771 | 5.24E-05 |
| Tmem198 | 1134.38453 | 0.558383854 | 5.24E-05 |
| LOC100910 | 1051.00207 | 0.454204262 | 5.25E-05 |
| RT1-S3 | 3072.07033 | -2.135625553 | 5.26E-05 |
| Zmat4 | 661.085886 | 0.426269783 | 5.29E-05 |
| Cacng8 | 2180.09735 | 0.60783118 | 5.31E-05 |
| Lpl | 1588.37408 | 0.804272861 | 5.35E-05 |
| Abr | 22417.3982 | 0.293421654 | 5.36E-05 |
| Ctnnd1 | 5165.48885 | -0.352168408 | 5.37E-05 |
| LOC102553 | 1612.25303 | 0.399025526 | 5.41E-05 |
| Mt2A | 2216.10718 | -1.749474202 | 5.42E-05 |
| Fgr | 35.87201 | -3.076507736 | 5.54E-05 |
| Camkk2 | 891.996259 | 0.757657658 | 5.54E-05 |
| Zfhx2 | 2504.56175 | 0.691220667 | 5.58E-05 |
| Litaf | 1206.58605 | -0.844149112 | 5.59E-05 |
| Slc24a4 | 1467.3622 | 0.778940309 | 5.59E-05 |
| Gpnmb | 701.40992 | -1.617913734 | 5.60E-05 |
| Sympk | 4140.21458 | 0.23914069 | 5.70E-05 |

| | | | |
|---|---|---|---|
| Casp7 | 369.580979 | -0.990083349 | 5.72E-05 |
| Fry | 14772.8963 | 0.373517644 | 5.79E-05 |
| Hdlbp | 13994.2645 | 0.267550483 | 5.83E-05 |
| Rnpepl1 | 786.451357 | -0.511211343 | 5.84E-05 |
| Gab1 | 3456.14222 | -0.639066364 | 5.88E-05 |
| Hectd1 | 12366.9439 | 0.160465407 | 5.88E-05 |
| Fah | 266.310473 | -0.820711074 | 5.89E-05 |
| Ankrd33b | 326.894854 | 1.23893142 | 5.90E-05 |
| Lgals9 | 218.964303 | -3.709165942 | 5.93E-05 |
| Oas1b | 182.049289 | -4.131159905 | 5.95E-05 |
| LOC103690 | 1321.82408 | -1.018317965 | 6.00E-05 |
| Akap1 | 2157.17146 | 0.39590085 | 6.04E-05 |
| Adam17 | 1357.63012 | -0.550750975 | 6.06E-05 |
| RGD156047 | 4763.39824 | 0.420089709 | 6.19E-05 |
| Slc12a5 | 15073.3711 | 0.506782456 | 6.19E-05 |
| Ryr2 | 6284.62138 | 0.562157164 | 6.19E-05 |
| Strip2 | 1665.12148 | 0.629789776 | 6.19E-05 |
| Fnta | 2801.41227 | -0.405207649 | 6.33E-05 |
| Mthfd1l | 756.657934 | 0.482722885 | 6.34E-05 |
| Rnaset2 | 784.805706 | -0.625795851 | 6.37E-05 |
| Pias3 | 1113.711 | -0.357328699 | 6.37E-05 |
| Wasf1 | 5089.22657 | 0.535187099 | 6.55E-05 |
| Nr4a1 | 2189.53578 | 0.920151488 | 6.66E-05 |
| Mark2 | 4725.62294 | 0.429053802 | 6.69E-05 |
| Pth1r | 403.361646 | -0.824355426 | 6.72E-05 |
| Npr1 | 621.897162 | -0.866828888 | 6.75E-05 |
| Rab5c | 8780.16645 | -0.333066766 | 6.77E-05 |
| Rela | 1413.98144 | -0.584901718 | 6.77E-05 |
| Igsf10 | 483.079311 | -1.028875782 | 6.79E-05 |
| Eda2r | 201.055426 | -0.961303015 | 6.79E-05 |
| Neurl4 | 3982.28914 | 0.351452803 | 6.87E-05 |
| Klf16 | 1878.01235 | 0.459328844 | 6.90E-05 |
| Syne1 | 20158.9381 | 0.512925748 | 6.96E-05 |
| Chgb | 13582.4413 | 0.451356342 | 7.02E-05 |
| Parp11 | 705.842237 | -0.7671003 | 7.03E-05 |
| Kif18b | 19.4568919 | -2.461387974 | 7.24E-05 |

| | | | |
|---|---|---|---|
| Osmr | 132.773046 | -3.534390354 | 7.33E-05 |
| Camsap2 | 11751.4794 | 0.261782063 | 7.33E-05 |
| Cnst | 1611.14567 | 0.295588173 | 7.33E-05 |
| Mtmr12 | 760.825636 | 0.424861558 | 7.36E-05 |
| Mag | 1855.6262 | -2.170954621 | 7.41E-05 |
| Plcd1 | 2272.80777 | -0.488430628 | 7.55E-05 |
| Rasa4 | 74.2898909 | -2.64828136 | 7.63E-05 |
| Kcnh3 | 1491.77336 | 0.890152066 | 7.65E-05 |
| Map3k12 | 2889.02188 | 0.370793045 | 7.70E-05 |
| Lif | 39.454686 | -1.446442689 | 7.73E-05 |
| Ptpn21 | 443.245481 | -0.523114115 | 7.73E-05 |
| Map4k4 | 10099.908 | -0.456373499 | 7.73E-05 |
| Adrb1 | 476.53026 | 0.611846363 | 7.73E-05 |
| Dgkg | 3154.86636 | 0.720943735 | 7.75E-05 |
| Mkl2 | 783.34295 | 0.59869508 | 7.88E-05 |
| Dlgap3 | 7903.13635 | 0.612946997 | 7.92E-05 |
| Adamts4 | 207.870949 | -2.167999856 | 7.94E-05 |
| Cmpk2 | 2020.17099 | -1.92046411 | 7.94E-05 |
| Vwa5b2 | 584.473484 | 0.642259874 | 8.41E-05 |
| Tgfbr1 | 317.375143 | -0.535065164 | 8.48E-05 |
| Cdyl2 | 1497.12876 | 0.394486349 | 8.61E-05 |
| Emp2 | 225.449804 | -1.18572847 | 8.73E-05 |
| Wscd1 | 4579.72106 | -0.574426441 | 8.76E-05 |
| Cdk5rap2 | 1181.06685 | -0.613160142 | 8.85E-05 |
| Cacnb2 | 1705.1914 | 0.368091123 | 8.86E-05 |
| Nsmce1 | 799.207551 | -0.530617556 | 8.93E-05 |
| Lrp12 | 1826.7659 | 0.424553734 | 8.93E-05 |
| LOC689986 | 646.604761 | 0.607894379 | 8.96E-05 |
| Ddn | 10986.119 | 0.667073668 | 9.01E-05 |
| Arc | 2588.73899 | 1.260460528 | 9.39E-05 |
| Nckap1l | 29.0998222 | -2.049132156 | 9.39E-05 |
| Dock4 | 4192.5631 | 0.316621691 | 9.39E-05 |
| Marcksl1 | 473.050315 | -0.525426864 | 9.39E-05 |
| Ppp1r9a | 10006.4163 | 0.459782315 | 9.53E-05 |
| Nkx2-2 | 368.231316 | -0.804618941 | 9.64E-05 |
| Lama4 | 250.06387 | -0.58431071 | 9.64E-05 |

| | | | |
|---|---|---|---|
| LOC108348 | 1490.02715 | -0.515967846 | 9.64E-05 |
| Tmem43 | 1533.70685 | -0.481787672 | 9.64E-05 |
| Pcsk2 | 5047.07143 | 0.612144647 | 9.64E-05 |
| Prr18 | 363.123698 | -0.895310689 | 9.73E-05 |
| Micall2 | 191.885484 | -0.683495611 | 9.73E-05 |
| Stat3 | 6176.24295 | -0.784317099 | 9.89E-05 |
| Rdx | 4647.3634 | -0.419453583 | 9.98E-05 |
| Abl2 | 4313.86025 | 0.25410852 | 9.98E-05 |
| Trim5 | 769.329998 | -0.940206488 | 0.000101568 |
| Rps6ka2 | 3013.11609 | 0.642386222 | 0.000102991 |
| Mapkbp1 | 1593.36174 | 0.330314459 | 0.000106361 |
| Adgrb2 | 15249.0687 | 0.385906115 | 0.000108776 |
| Arf3 | 5723.67272 | 0.49788393 | 0.000109033 |
| Afg3l1 | 987.821266 | -0.410926553 | 0.000109422 |
| Zfp36 | 187.017097 | -1.689205151 | 0.000109494 |
| Dlg3 | 7362.4524 | 0.424072402 | 0.000109966 |
| Gfap | 91950.4251 | -1.944947254 | 0.000110641 |
| Inpp4a | 9242.93123 | 0.424961944 | 0.000110641 |
| Acy1 | 2058.42368 | -0.462533466 | 0.000112376 |
| Pitpnm2 | 14158.933 | 0.377099006 | 0.000112376 |
| Plekha5 | 1768.58454 | 0.343969591 | 0.000112865 |
| Cldn11 | 1019.88416 | -2.238004612 | 0.00011324 |
| Dpf1 | 1659.76608 | 0.515979287 | 0.000114487 |
| Trim21 | 1040.5433 | -1.649879055 | 0.000115582 |
| RGD130488 | 17582.5014 | 0.504171787 | 0.000117153 |
| Atp1a1 | 17379.8235 | 0.529666328 | 0.00011776 |
| Gm2a | 3264.02948 | -0.582409752 | 0.000118745 |
| Atf3 | 37.3267099 | -3.066830539 | 0.000119206 |
| Tmem132b | 2547.07685 | 0.524490277 | 0.000119206 |
| Ezh1 | 2775.93664 | 0.374660785 | 0.000120496 |
| Ifi35 | 157.032411 | -1.537979538 | 0.000121523 |
| Sec16a | 6467.48999 | 0.216144471 | 0.00012227 |
| Syt4 | 6365.5735 | 0.522374456 | 0.00012227 |
| Slc27a3 | 1055.0031 | -0.838989499 | 0.00012337 |
| Slit3 | 1732.50154 | 0.72176078 | 0.00012337 |
| Osbpl8 | 4074.51848 | 0.320523775 | 0.000123705 |

| | | | |
|---|---|---|---|
| Csmd1 | 3438.4947 | 0.491151704 | 0.000124173 |
| Sybu | 3369.55548 | 0.301798884 | 0.000126115 |
| Trim34 | 325.403141 | -1.018930591 | 0.000126856 |
| Hpcal4 | 17135.3389 | 0.702488052 | 0.000128517 |
| Sgip1 | 12824.3418 | 0.445488257 | 0.000128579 |
| Tmem176a | 231.432772 | -1.317334393 | 0.000130989 |
| Cirbp | 2100.53167 | -0.368305236 | 0.000131888 |
| Mapre1 | 3921.46173 | -0.329906627 | 0.000132761 |
| Gsdmd | 687.382227 | -0.88343292 | 0.000133076 |
| Wscd2 | 1135.98108 | 0.497303593 | 0.000133076 |
| LOC100910 | 74.4190126 | -3.756460938 | 0.000135674 |
| Nhsl2 | 3250.88516 | 0.541048128 | 0.000135674 |
| Chst1 | 5801.11861 | 0.322105686 | 0.000136009 |
| Man2a1 | 1200.79897 | 0.374613388 | 0.000138294 |
| Dnajc5 | 14738.9275 | 0.345051702 | 0.000139174 |
| Fam57a | 84.6997057 | -1.268586341 | 0.000140477 |
| LOC690155 | 221.494444 | -0.464298502 | 0.000140876 |
| C1ql1 | 421.0991 | -0.696766657 | 0.000140981 |
| Gpr63 | 451.489016 | 0.576926337 | 0.000142491 |
| Vamp3 | 1520.50399 | -0.580411802 | 0.000142719 |
| Myzap | 58.0798112 | -0.963882776 | 0.000142807 |
| Bhlhe23 | 45.427192 | 1.662383152 | 0.000142807 |
| Odc1 | 3257.83811 | -0.415140302 | 0.000143383 |
| Ikbip | 1231.85283 | -0.445691232 | 0.000143471 |
| Zfyve28 | 988.810688 | 0.468949006 | 0.0001465 |
| Slc14a1 | 7354.07666 | -0.535999863 | 0.000147615 |
| Rgma | 15221.3926 | -0.481586141 | 0.000147923 |
| Dip2c | 5739.41963 | 0.357756937 | 0.000148355 |
| Il13ra1 | 134.860667 | -1.172988683 | 0.000148666 |
| Zdhhc23 | 914.618269 | 0.542362336 | 0.000148666 |
| Ccdc106 | 519.58126 | 0.485396539 | 0.000150282 |
| Tanc2 | 13116.8106 | 0.41516876 | 0.000151075 |
| Ephx1 | 4151.09532 | -0.627570004 | 0.000152058 |
| Rnf43 | 296.909385 | -0.774901267 | 0.000152723 |
| Parp3 | 537.322916 | -1.826948132 | 0.000153029 |
| Frmd4b | 544.217039 | -0.551801907 | 0.000154258 |

| | | | |
|---|---|---|---|
| Lpar1 | 436.610788 | -1.206867157 | 0.000154939 |
| Tvp23b | 1138.68953 | -0.472102652 | 0.000155283 |
| Trrap | 8733.01538 | 0.297970356 | 0.000155283 |
| Zfp638 | 3138.86468 | 0.26793671 | 0.000155624 |
| Slc44a1 | 2880.78521 | -0.757018615 | 0.00015678 |
| Chrm1 | 3019.04665 | 0.760432168 | 0.000156826 |
| Usp31 | 2709.40645 | 0.512665039 | 0.000157895 |
| Oas1a | 268.627664 | -3.716734527 | 0.000158127 |
| Cobl | 3027.40277 | 0.80928275 | 0.000158524 |
| Herc6 | 803.374144 | -1.934902403 | 0.000160156 |
| Wdtc1 | 3344.17512 | 0.30733774 | 0.000160156 |
| Kcns2 | 318.22839 | 0.748446302 | 0.000160578 |
| Npas4 | 576.233582 | 1.145583329 | 0.000160578 |
| Eps8 | 7569.86947 | -0.426327965 | 0.0001608 |
| Ncf1 | 21.359332 | -2.484269428 | 0.000162679 |
| Ifi44 | 416.273728 | -2.125189398 | 0.000163417 |
| Cacna1a | 6818.75519 | 0.582483743 | 0.000163417 |
| Kcnk1 | 1914.0785 | 0.530622749 | 0.000163684 |
| Lamb1 | 1537.60973 | 0.588277521 | 0.000165176 |
| Abcc8 | 559.445841 | 0.616395427 | 0.000165584 |
| Cox20 | 255.045739 | -0.488997172 | 0.000167108 |
| Capn15 | 1834.50727 | 0.354513678 | 0.000167754 |
| Rai1 | 4927.68762 | 0.31965285 | 0.000169407 |
| Manba | 682.417332 | -0.433620765 | 0.000170593 |
| Tmem117 | 283.788568 | -0.590856267 | 0.000173833 |
| Prrt2 | 3392.00008 | 0.558310248 | 0.000173833 |
| Nudt12 | 796.026927 | -0.314911354 | 0.000176408 |
| Aida | 2348.01958 | -0.597246225 | 0.00017674 |
| Abca8a | 882.32285 | -1.567992245 | 0.000177025 |
| Enpp2 | 1521.40155 | -0.853071895 | 0.000177412 |
| Rrm2 | 23.5212669 | -1.426142124 | 0.000182387 |
| Megf10 | 4223.0317 | -0.575970946 | 0.000182605 |
| Pcolce | 32.1471508 | -3.6789484 | 0.000183466 |
| Necap2 | 1603.13611 | -0.498577369 | 0.000183466 |
| Rapgef2 | 9478.83098 | 0.454950875 | 0.000183466 |
| Kif5c | 29368.102 | 0.503024481 | 0.000183466 |

| | | | |
|---|---|---|---|
| Tor3a | 624.919546 | -0.725473323 | 0.000184157 |
| Mtcl1 | 4207.29953 | 0.462743608 | 0.000184279 |
| Dnajb5 | 2962.80186 | 0.489543593 | 0.000184617 |
| Kcna4 | 1963.6211 | 0.481391065 | 0.000184746 |
| Ppp2r3a | 1012.23562 | -0.344667726 | 0.00018488 |
| Neto1 | 3274.97036 | 0.594999681 | 0.000185199 |
| Tp53 | 1068.14699 | -0.617753664 | 0.000186128 |
| Ehmt2 | 5998.67237 | 0.270810524 | 0.000186223 |
| Serac1 | 657.065716 | 0.316838607 | 0.00018837 |
| Mtor | 7072.33632 | 0.290746208 | 0.000188454 |
| Klhl2 | 3761.76228 | 0.509315081 | 0.000188472 |
| Dcakd | 2869.56843 | -0.675637981 | 0.000188807 |
| Usp7 | 7346.56729 | 0.180170658 | 0.000189134 |
| Prkcg | 6152.10085 | 0.710116529 | 0.000189134 |
| Jak3 | 258.063718 | -0.917272436 | 0.000192116 |
| Pdp1 | 1542.29667 | 0.539787915 | 0.000192116 |
| Tshr | 233.095198 | -1.30629283 | 0.000194474 |
| Celf2 | 10923.4704 | 0.333687755 | 0.000194474 |
| Lamp2 | 7485.34568 | -0.518544515 | 0.000198865 |
| Cacng3 | 685.254267 | 0.682828424 | 0.000198955 |
| Plpp1 | 934.265878 | -0.521407864 | 0.000201187 |
| Nln | 2910.74265 | -0.402117343 | 0.000202922 |
| Grin1 | 19839.7783 | 0.491292687 | 0.000202922 |
| Sorbs2 | 4913.44928 | 0.564151551 | 0.000202972 |
| Plekhg1 | 1827.66097 | -0.624456966 | 0.000206703 |
| Klk6 | 198.537556 | -2.989290465 | 0.000207264 |
| Gnal | 3701.72707 | 0.507070185 | 0.000207387 |
| Zmynd8 | 4565.90514 | 0.216165157 | 0.000207873 |
| Unc80 | 13687.1486 | 0.481325919 | 0.000208416 |
| Slfn4 | 33.7233311 | -4.664456185 | 0.000208741 |
| Fbxo41 | 3071.58753 | 0.43812995 | 0.000210165 |
| Palmd | 131.361908 | 1.088914117 | 0.000211225 |
| Ubd | 97.1959714 | -4.694316395 | 0.000217389 |
| Aif1 | 9.44192073 | -3.649479672 | 0.000222034 |
| Rabl6 | 6616.47229 | 0.261285571 | 0.000223665 |
| Rims4 | 2336.87456 | 0.48146195 | 0.000224964 |

| | | | |
|---|---|---|---|
| Nsg2 | 23950.6906 | 0.360426197 | 0.000225068 |
| Fam189a1 | 850.977194 | 0.566570636 | 0.000229022 |
| Prkcb | 11811.9007 | 0.623247856 | 0.000229022 |
| Dock8 | 68.3779397 | -1.368422519 | 0.000229586 |
| Enpp6 | 46.0294011 | -2.015455779 | 0.00023028 |
| Ubqln2 | 16275.0756 | 0.254259312 | 0.000230957 |
| Ppp1r14a | 125.106264 | -1.821936216 | 0.000235612 |
| Tmem255b | 93.0933394 | -0.854067704 | 0.000235612 |
| Cttnbp2 | 3278.31641 | 0.347258622 | 0.000235612 |
| Elmod1 | 5212.63251 | 0.447527314 | 0.000235612 |
| Myl9 | 355.076492 | -1.07031384 | 0.000237565 |
| Lsm6 | 762.386261 | -0.44281926 | 0.000237565 |
| Zfand1 | 460.769171 | -0.395655516 | 0.00023779 |
| Vangl2 | 1579.16461 | -0.41863976 | 0.000238182 |
| Cd82 | 1294.96162 | -0.564808665 | 0.000239095 |
| Tmco4 | 395.287315 | -0.689925093 | 0.000241695 |
| Cdk6 | 250.897677 | -1.259884657 | 0.000246788 |
| Ylpm1 | 7650.86518 | 0.311092545 | 0.000246788 |
| Socs1 | 70.9763192 | -2.057071562 | 0.000250636 |
| Tecta | 225.04461 | -0.716107484 | 0.000252188 |
| Elavl3 | 3721.83572 | 0.331236474 | 0.000253096 |
| Aen | 1341.84859 | -0.539931027 | 0.000253475 |
| Doc2b | 1210.31006 | 0.475571684 | 0.000253475 |
| Slc7a5 | 3896.12909 | -0.497010492 | 0.000254469 |
| Stxbp5l | 1929.22247 | 0.541084809 | 0.000255693 |
| Lgals3 | 107.780202 | -2.564092281 | 0.00025611 |
| Adgrd1 | 222.620654 | -0.977291797 | 0.000257551 |
| Cpne6 | 1655.91058 | 0.473637222 | 0.000257551 |
| Rab11fip3 | 5143.03405 | 0.366239558 | 0.00025906 |
| Trafd1 | 1220.43526 | -0.78148403 | 0.000259395 |
| Epb41l1 | 14338.4689 | 0.413347984 | 0.000259395 |
| Slc12a2 | 3670.08489 | -0.530071288 | 0.000260211 |
| Pla2g4a | 55.5312563 | -0.891280229 | 0.000260375 |
| Nrgn | 5710.56803 | 0.811877831 | 0.00026125 |
| Lpcat2 | 66.5843801 | -0.835514677 | 0.000262039 |
| Eif2ak2 | 2300.87881 | -1.032608084 | 0.000262717 |

| | | | |
|---|---|---|---|
| Aqp4 | 133176.721 | -0.57847792 | 0.000263193 |
| Sh3rf3 | 1316.64044 | 0.436820718 | 0.000263626 |
| Tnfrsf1b | 17.4674174 | -3.727780152 | 0.000264578 |
| Arhgef10 | 1654.13855 | -0.524002481 | 0.000266032 |
| Psme1 | 2409.90406 | -1.083155343 | 0.000266765 |
| Plcb1 | 10647.1817 | 0.346160264 | 0.000271294 |
| Btbd8 | 4044.54363 | 0.551853347 | 0.000272009 |
| Caln1 | 5097.66334 | 0.571190868 | 0.000273874 |
| Rab21 | 4727.52522 | -0.299739277 | 0.000274884 |
| Ddx58 | 975.968323 | -1.38220633 | 0.000275274 |
| Sirt2 | 6512.69132 | -0.348130711 | 0.000275274 |
| Car11 | 1639.62666 | 0.419979923 | 0.000275274 |
| Neurl1b | 373.196458 | 0.632547973 | 0.000275274 |
| Sptbn2 | 20520.6589 | 0.662567831 | 0.000275274 |
| Madd | 6959.38822 | 0.412213723 | 0.000276285 |
| Zfyve9 | 3895.74142 | 0.315373283 | 0.000279121 |
| Pnp | 8886.68933 | -0.540758302 | 0.000279627 |
| Nedd4l | 3527.2712 | 0.343085587 | 0.000280436 |
| Scn3b | 5932.61699 | 0.485399171 | 0.000280893 |
| Tf | 4866.11748 | -2.358798468 | 0.000285234 |
| Apod | 801.006759 | -2.350637497 | 0.000285234 |
| Jag1 | 274.672069 | -0.54832075 | 0.000285548 |
| Mpp3 | 1043.341 | 0.501858323 | 0.000285548 |
| Adgre1 | 12.3339956 | -2.685887711 | 0.000286314 |
| Pdlim4 | 7294.89883 | -0.572390451 | 0.000287866 |
| Ubr7 | 2964.12962 | -0.444583435 | 0.000289142 |
| Serpinb9 | 798.796689 | -0.781287485 | 0.000289723 |
| Ablim3 | 1274.44882 | 0.384121175 | 0.000289723 |
| Slc4a10 | 7682.54873 | 0.537863832 | 0.000290395 |
| Col16a1 | 847.403629 | -0.795598786 | 0.000293259 |
| Mypop | 1614.70352 | 0.38453813 | 0.000293486 |
| Krcc1 | 980.043225 | -0.542066992 | 0.000294759 |
| Helz2 | 190.921573 | -2.271006387 | 0.000294963 |
| Micu3 | 2256.34019 | 0.382170095 | 0.000295482 |
| LOC102554 | 24.4003089 | -2.852149621 | 0.000295885 |
| Zfp385b | 1525.60024 | 0.450119459 | 0.000298506 |

| | | | |
|---|---|---|---|
| Sash1 | 12393.6011 | -0.376652373 | 0.000300168 |
| Lrrc8b | 2491.21626 | 0.58454494 | 0.000302093 |
| Rap1gap | 9388.56685 | 0.385185376 | 0.000308162 |
| Tagln | 450.138978 | -2.600170927 | 0.000309555 |
| Arhgef3 | 3167.82023 | 0.586830712 | 0.000310711 |
| Celf1 | 8019.77421 | 0.273973564 | 0.000312079 |
| Zfp365 | 5419.51089 | 0.625384922 | 0.00031348 |
| RGD130627 | 2779.13899 | 0.440266333 | 0.00031574 |
| Scn2b | 11764.2509 | 0.423670143 | 0.000316454 |
| Sntb1 | 5193.428 | -0.517378096 | 0.000316813 |
| Ssx2ip | 2089.86293 | 0.30507532 | 0.000319675 |
| Itgal | 19.9411109 | -2.929751905 | 0.000323363 |
| Plp1 | 12027.5336 | -1.530653267 | 0.000324252 |
| Arfgef3 | 5198.91154 | 0.406912315 | 0.000324252 |
| Mvp | 1239.20688 | -1.170666636 | 0.000324791 |
| Sema6a | 5794.03459 | -0.482224558 | 0.000324791 |
| Batf2 | 41.8672041 | -4.16847642 | 0.000324887 |
| Fuca1 | 1968.42708 | -0.395082259 | 0.000324887 |
| Gp1bb | 170.784338 | 0.683755837 | 0.000333638 |
| Kmt2d | 14025.5824 | 0.341427544 | 0.000338218 |
| Add2 | 11222.5458 | 0.470290873 | 0.000338792 |
| Kcng2 | 312.933262 | 0.840003678 | 0.000348322 |
| Erbin | 7219.38445 | -0.452953117 | 0.000348447 |
| Kalrn | 7719.47078 | 0.576552266 | 0.000348447 |
| LOC691807 | 2467.41382 | -0.40716019 | 0.000352831 |
| Prokr2 | 467.291465 | 0.814344503 | 0.000352831 |
| Cenpj | 535.062646 | -0.820014621 | 0.000354922 |
| Casd1 | 2314.8749 | 0.297131218 | 0.000354922 |
| Lmtk3 | 6833.765 | 0.434775996 | 0.000355554 |
| Ets2 | 1501.24783 | 0.599189063 | 0.000355745 |
| Evi2a | 313.308493 | -1.0539564 | 0.000358831 |
| Gsap | 33.4978859 | -2.850111511 | 0.000366531 |
| Rasgef1a | 3638.13185 | 0.597895744 | 0.0003693 |
| Sstr2 | 640.331784 | 0.895977309 | 0.0003693 |
| Apbb1ip | 45.8091199 | -0.923572011 | 0.00037066 |
| Ifih1 | 609.779182 | -1.319840708 | 0.000377161 |

| | | | |
|---|---|---|---|
| Ccl19 | 37.2243565 | -4.110596856 | 0.000378549 |
| Armt1 | 757.117376 | 0.285737749 | 0.000383099 |
| Dmxl2 | 12177.5794 | 0.447593184 | 0.000383099 |
| Ctnna1 | 9066.60132 | -0.452451327 | 0.00038372 |
| Btbd7 | 2511.32749 | -0.382171079 | 0.000391199 |
| Chpf | 5267.6887 | 0.256884642 | 0.000393677 |
| Clic1 | 817.472796 | -0.673865587 | 0.000401741 |
| Irs2 | 6028.91384 | 0.412778884 | 0.000402203 |
| Pcdh1 | 7338.56999 | 0.455904627 | 0.000402203 |
| Htt | 7589.78453 | 0.25325302 | 0.000403778 |
| Kdm7a | 1378.87982 | 0.435460331 | 0.000403778 |
| Arhgap32 | 19845.091 | 0.409837983 | 0.000408523 |
| Setx | 10227.4037 | 0.28907466 | 0.000408865 |
| Paip1 | 3611.74993 | -0.225638622 | 0.000411198 |
| Sptan1 | 41053.2011 | 0.410603296 | 0.000413478 |
| Ezr | 13702.0804 | -0.483335217 | 0.000414741 |
| Erbb4 | 3679.05323 | 0.367803024 | 0.000415457 |
| Bsn | 25557.3617 | 0.693440656 | 0.000418403 |
| Zfp488 | 41.032872 | -1.163014057 | 0.000422609 |
| LOC100362 | 244.542028 | 0.865391206 | 0.000426003 |
| Adamts15 | 300.384431 | 0.766933561 | 0.000426228 |
| Has2 | 9.34315991 | -3.65005839 | 0.00042876 |
| Dusp1 | 683.051307 | 0.727533386 | 0.000430579 |
| Stk3 | 337.176258 | -0.576819381 | 0.00043132 |
| Kif11 | 44.0992016 | -0.888254274 | 0.00043635 |
| Rapgef5 | 3536.02405 | 0.510473442 | 0.000438308 |
| Hjurp | 18.2592267 | -2.075091849 | 0.000438935 |
| Gpr6 | 508.579698 | 0.812224721 | 0.00044085 |
| Cnn2 | 502.112495 | -0.879037213 | 0.00044138 |
| B3gnt7 | 87.9332219 | -0.79280509 | 0.000442432 |
| Ckap2l | 20.9153337 | -1.272945966 | 0.00044466 |
| Apobec1 | 106.372309 | -0.826538552 | 0.00044466 |
| Atp2a2 | 54818.4199 | 0.284830777 | 0.00044466 |
| Hmgxb3 | 1608.69418 | 0.33280798 | 0.00044466 |
| Dbn1 | 4091.79092 | 0.411144379 | 0.00044489 |
| Aak1 | 21465.6287 | 0.381045067 | 0.000450231 |

| | | | |
|---|---|---|---|
| P2ry2 | 30.8001311 | -1.315584063 | 0.000453089 |
| Tcirg1 | 762.513618 | -0.62328211 | 0.000453089 |
| Dlgap4 | 6441.26304 | 0.415312108 | 0.000453145 |
| Sertad4 | 480.338351 | 0.415559892 | 0.000457303 |
| C4b | 25.4518395 | -4.054617304 | 0.000457801 |
| Mtmr4 | 4244.67958 | 0.355632697 | 0.000457801 |
| P2rx7 | 290.774844 | -0.89765237 | 0.000460687 |
| Mbnl2 | 6439.50959 | 0.299340608 | 0.000463666 |
| Rexo1 | 2433.8822 | 0.315252371 | 0.000474106 |
| Rangap1 | 7498.6104 | 0.352566249 | 0.000474106 |
| Lcn2 | 57.1473151 | -4.021611286 | 0.000474862 |
| Galnt9 | 1416.76012 | 0.751292802 | 0.00047738 |
| Dtx3l | 87.4167474 | -1.81204308 | 0.000477572 |
| Grm5 | 6954.50825 | 0.411480517 | 0.000479018 |
| Fmnl1 | 2586.43908 | 0.610368476 | 0.000479362 |
| Mitd1 | 158.992541 | -0.923169122 | 0.000480254 |
| Fubp3 | 2323.1101 | -0.438251534 | 0.000480373 |
| Necab2 | 1950.83654 | 0.556177566 | 0.00048359 |
| Grin2a | 6602.18798 | 0.62563283 | 0.000483776 |
| Mlkl | 29.3963929 | -3.998854177 | 0.000484646 |
| LOC100363 | 1360.8131 | -0.336447271 | 0.000485901 |
| Aspa | 133.991513 | -1.095781618 | 0.000486432 |
| RGD130936 | 116.045585 | -4.107922284 | 0.000486799 |
| Rhoc | 1179.63311 | -0.536248283 | 0.000489397 |
| Hgf | 486.043045 | -0.702836429 | 0.000491592 |
| Rpusd1 | 1284.9967 | 0.365947234 | 0.000496184 |
| Ncoa7 | 4487.71232 | 0.471213438 | 0.000496184 |
| Foxp2 | 1493.24055 | 0.584268122 | 0.00050666 |
| Srsf9 | 280.922792 | -0.527510753 | 0.000508856 |
| Plin2 | 873.612194 | -0.649912521 | 0.000509278 |
| Ypel2 | 1433.08779 | 0.592150952 | 0.000510605 |
| Arhgef9 | 6624.03554 | 0.358593068 | 0.000513019 |
| Elk1 | 1134.92828 | 0.300504095 | 0.000514944 |
| Elovl5 | 7011.61107 | -0.406239354 | 0.000521142 |
| Rab18 | 3436.11358 | -0.234652225 | 0.000525368 |
| Itpkb | 4923.64226 | -0.576764301 | 0.000528733 |

| | | | |
|---|---|---|---|
| Chd7 | 1980.36896 | -0.480709728 | 0.000531231 |
| Mzt2b | 1209.85578 | -0.443804642 | 0.000531231 |
| Dstn | 2692.58216 | -0.362977346 | 0.000531231 |
| Isoc2b | 289.251171 | -0.754156019 | 0.000531693 |
| Ifi27l2b | 36.1318441 | -2.968722492 | 0.000532774 |
| Parp8 | 676.372487 | -0.426354894 | 0.000532774 |
| Gas7 | 19154.2393 | 0.59610874 | 0.000539953 |
| Pja2 | 31899.9405 | 0.306573675 | 0.000541167 |
| Tspoap1 | 3738.49526 | 0.409932594 | 0.000547917 |
| Plxnb3 | 670.846704 | -0.566173666 | 0.000548056 |
| Samhd1 | 509.143274 | -0.715593758 | 0.000551073 |
| Irx1 | 50.7301799 | -1.15774498 | 0.000552646 |
| Cacna1b | 5397.50923 | 0.448521983 | 0.000552646 |
| Acss2 | 3047.32368 | -0.446990177 | 0.000552833 |
| Rnf44 | 5288.42105 | 0.274862451 | 0.000554392 |
| Idh1 | 3003.59065 | -0.390158608 | 0.000555395 |
| Sh3glb1 | 7077.67045 | -0.446030946 | 0.000559965 |
| Tmem140 | 62.3233536 | -1.791783322 | 0.00056121 |
| Prkca | 13303.2573 | 0.314938219 | 0.000566601 |
| Phf11 | 10.3038836 | -3.156869431 | 0.000567789 |
| Pgbd5 | 2888.57921 | 0.588449269 | 0.000574496 |
| Pbxip1 | 6099.95903 | -0.453450189 | 0.000577395 |
| Tspyl1 | 7243.06973 | 0.212065745 | 0.000577693 |
| Vcl | 3955.3258 | -0.551315106 | 0.000578337 |
| Nrarp | 2047.06298 | -0.655177634 | 0.000581755 |
| Map2 | 78695.666 | 0.413294021 | 0.000584395 |
| Anks1b | 6849.87599 | 0.501449241 | 0.000592833 |
| Mef2a | 5773.93376 | 0.414792934 | 0.000594902 |
| Map1a | 72036.141 | 0.460947593 | 0.000594902 |
| LOC100910 | 41.5325574 | 1.048713116 | 0.000596948 |
| Irf8 | 212.59425 | -2.343203746 | 0.000599934 |
| RT1-N3 | 99.9907188 | -2.805906549 | 0.00060031 |
| Prdm2 | 4653.44906 | 0.283131558 | 0.00060031 |
| Hrh3 | 6433.9629 | 0.758506899 | 0.00060031 |
| Atxn1 | 3948.48333 | 0.399874641 | 0.000601386 |
| Rbfox3 | 3121.66228 | 0.558334889 | 0.000602071 |

| | | | |
|---|---|---|---|
| Ranbp2 | 8691.58458 | 0.212764668 | 0.000606778 |
| Mcm3 | 51.0669972 | -0.922756493 | 0.000607611 |
| Shisa5 | 1902.07493 | -0.489206885 | 0.000608391 |
| Rgcc | 2254.98094 | -0.512003788 | 0.000615612 |
| Cx3cl1 | 14786.325 | 0.573805099 | 0.000619662 |
| Gopc | 2507.78667 | 0.298339267 | 0.000621821 |
| Cntnap1 | 8697.6415 | 0.487727997 | 0.000623592 |
| Fam126b | 2474.52557 | 0.490024704 | 0.000629563 |
| Abi2 | 4214.45147 | 0.243785245 | 0.000629616 |
| Aspg | 74.2940416 | -2.193639097 | 0.000630561 |
| Gpam | 26274.7451 | -0.517685395 | 0.000630561 |
| Srf | 2145.23087 | 0.335440699 | 0.000630561 |
| Tmem106a | 48.8178093 | -3.221293626 | 0.000631955 |
| LOC103694 | 38.3004197 | -1.850645539 | 0.000631955 |
| Fam13c | 1311.20675 | 0.429762925 | 0.000632141 |
| Prp2l1 | 2047.65878 | 0.328787454 | 0.000633726 |
| Naprt | 1170.14318 | -0.641869912 | 0.000639525 |
| Ubxn2b | 2266.74519 | 0.254456582 | 0.000639525 |
| Ptpru | 1567.13483 | 0.668464191 | 0.000639525 |
| Prpf40a | 2681.63999 | -0.234262906 | 0.000639785 |
| LOC103691 | 679.610516 | -0.645924353 | 0.000643035 |
| Wipf1 | 780.676495 | -0.571169117 | 0.000647479 |
| Als2cl | 550.602322 | -0.637891403 | 0.000648193 |
| Gps2 | 663.299785 | -0.287853774 | 0.000653844 |
| Bdnf | 316.681921 | 1.814785063 | 0.000653844 |
| Snx4 | 1530.59772 | -0.277629967 | 0.000658215 |
| Tapbp | 1282.74729 | -1.082142067 | 0.000659359 |
| Trim32 | 4461.75814 | 0.262434843 | 0.000659359 |
| Adcy1 | 16787.8503 | 0.612843722 | 0.000659488 |
| Ranbp3 | 2849.88621 | 0.207608373 | 0.000660741 |
| Elfn2 | 7188.36578 | 0.420754506 | 0.000660741 |
| Plekhg2 | 268.500658 | -0.695135427 | 0.000662533 |
| Otud7a | 2384.66122 | 0.487838446 | 0.000665451 |
| Cldn9 | 185.295531 | -0.824244246 | 0.00066812 |
| Pgghg | 531.183935 | -0.563434468 | 0.000668259 |
| Gng7 | 4367.57599 | 0.512547995 | 0.000668259 |

| | | | |
|---|---|---|---|
| RGD131134 | 1021.04916 | -0.255896294 | 0.000672486 |
| Ddhd2 | 1686.77397 | 0.305368709 | 0.000672486 |
| Prkd2 | 306.263029 | -0.509895832 | 0.000674478 |
| S100a16 | 6749.88738 | -0.494958438 | 0.000675533 |
| Plek | 14.2811053 | -2.212990148 | 0.000677321 |
| Calm1 | 29440.8012 | 0.414071459 | 0.000683255 |
| Ago2 | 3413.00173 | 0.288156262 | 0.000683419 |
| Nek6 | 1995.28558 | -0.633862245 | 0.000683737 |
| Samd9l | 1352.49432 | -0.962554618 | 0.000688369 |
| Bhlhe40 | 4178.18626 | -0.501839421 | 0.0006887 |
| Tlr3 | 353.274072 | -0.700351295 | 0.00069653 |
| Gna12 | 7201.24469 | -0.399469991 | 0.00069653 |
| Slc16a1 | 3304.00922 | -0.580986864 | 0.000698203 |
| Mtmr1 | 1786.03681 | 0.285985658 | 0.000707097 |
| Myo9b | 2511.97131 | -0.407607906 | 0.000707513 |
| Klhl3 | 423.894809 | 0.595525316 | 0.000708722 |
| Scamp2 | 816.74655 | -0.517752592 | 0.000709986 |
| Tspan12 | 4251.67166 | -0.363218149 | 0.000716255 |
| Tom1 | 2273.18213 | -0.290772588 | 0.000718212 |
| Fbxo34 | 1815.76117 | 0.425502237 | 0.000723481 |
| Tspan2 | 1257.09505 | -0.59684107 | 0.000724381 |
| Tdrd7 | 2847.65192 | -0.467440944 | 0.000726188 |
| Hpgd | 69.023852 | -1.526873515 | 0.000726514 |
| Ppic | 45.4235789 | -1.428930484 | 0.000726764 |
| Unc79 | 3523.81084 | 0.227980367 | 0.000729292 |
| Pcdha4 | 8269.37812 | 0.403146131 | 0.000737621 |
| Rffl | 418.291432 | -0.667389452 | 0.000740683 |
| Celf4 | 12474.8011 | 0.390849057 | 0.00074181 |
| Timp1 | 111.457446 | -2.590221301 | 0.000741908 |
| Lrrtm3 | 1716.60473 | 0.499331185 | 0.000743723 |
| Cebpd | 1038.11971 | -1.765005688 | 0.000744206 |
| Tmem150a | 219.000514 | -0.592019237 | 0.000744206 |
| Arfgef2 | 4407.20234 | 0.275018476 | 0.000744206 |
| Ppp4r4 | 382.868471 | 0.519403784 | 0.000744206 |
| Afap1 | 1774.42289 | -0.362218829 | 0.000748295 |
| B4galnt1 | 1769.67915 | 0.478508756 | 0.000754076 |

| | | | |
|---|---|---|---|
| Ralgapb | 7426.11874 | 0.225643818 | 0.000764406 |
| Lancl1 | 2350.90995 | 0.356763132 | 0.000764406 |
| Glce | 2131.06036 | 0.436718919 | 0.000764406 |
| LOC108348 | 5680.38113 | -0.549982194 | 0.000770846 |
| Ddi2 | 4407.00244 | 0.315301127 | 0.000772303 |
| Relb | 133.736733 | -0.972050731 | 0.000772393 |
| Nkx6-2 | 504.879249 | -0.767691353 | 0.000772393 |
| Avl9 | 4455.97542 | 0.213157744 | 0.000773244 |
| Kcnj12 | 826.703038 | 0.475211767 | 0.000773244 |
| Plekhh1 | 580.761132 | -0.802309184 | 0.000776782 |
| Dmxl1 | 4784.28598 | 0.234059757 | 0.000778699 |
| Kitlg | 453.771645 | 0.469168799 | 0.000787067 |
| Dgat2 | 948.828865 | 0.461640675 | 0.000792035 |
| Myrf | 1587.62413 | -1.432258931 | 0.00081273 |
| Hipk3 | 6026.95586 | 0.294902762 | 0.000823462 |
| Fgfbp3 | 490.071604 | -0.64411275 | 0.000829225 |
| Pfkfb3 | 4907.23517 | -0.575881181 | 0.000830715 |
| Kpna1 | 4420.14586 | 0.256957908 | 0.000834687 |
| Camsap1 | 7256.63121 | 0.309779515 | 0.000835798 |
| Emsy | 2356.24363 | 0.258219755 | 0.0008398 |
| Ppip5k1 | 3448.67365 | 0.350398056 | 0.000844414 |
| Nampt | 2191.52345 | -0.697397402 | 0.000845543 |
| Limk2 | 3358.40022 | -0.371558707 | 0.000845543 |
| Arhgap20 | 2706.94859 | 0.373312157 | 0.000845543 |
| LOC100362 | 6479.51207 | 0.616903441 | 0.000845543 |
| Arhgap10 | 97.0560507 | 0.876502006 | 0.000845543 |
| Tmem30a | 10189.7331 | 0.208121753 | 0.000848721 |
| Lmnb1 | 382.755529 | -0.383383634 | 0.000860394 |
| Ifi27 | 1937.82495 | -1.232877763 | 0.000861184 |
| Rras | 663.656931 | -0.607785213 | 0.000861329 |
| Sez6l2 | 13401.5329 | 0.42720752 | 0.000861329 |
| Cacna2d1 | 4125.22776 | 0.598637389 | 0.000861329 |
| Cap2 | 4683.21526 | 0.455734687 | 0.000862929 |
| Fbxw17 | 191.092764 | -0.920225305 | 0.000872291 |
| Pi4ka | 15534.8924 | 0.396709302 | 0.000879856 |
| Gatsl2 | 9526.52263 | 0.305992711 | 0.00088486 |

| | | | |
|---|---|---|---|
| Pik3r2 | 5301.48916 | 0.35716373 | 0.000898086 |
| Klf5 | 818.891032 | 0.512224691 | 0.000898252 |
| LOC102551 | 221.45099 | -0.846253514 | 0.000902012 |
| Llgl1 | 6036.47585 | -0.409211532 | 0.000903125 |
| Dgkb | 10721.7484 | 0.363837325 | 0.000903125 |
| Muc15 | 10.9073033 | -3.555645366 | 0.000904062 |
| Entpd3 | 655.177722 | 0.464344374 | 0.000906705 |
| Fam65a | 4336.51236 | 0.402850636 | 0.000914093 |
| Cst3 | 60496.34 | -0.413055128 | 0.000921488 |
| Arhgef1 | 938.795748 | -0.368979149 | 0.000922425 |
| Cnn3 | 15749.0421 | -0.808429012 | 0.000924469 |
| Cacna1i | 2856.32568 | 0.499312739 | 0.000925616 |
| Dgka | 808.895429 | 0.26917576 | 0.000928879 |
| Kcna5 | 396.344878 | 0.494681811 | 0.000930609 |
| Lyrm5 | 326.997521 | -0.566221934 | 0.000937216 |
| Sgsm3 | 2158.654 | 0.301647046 | 0.000937615 |
| Atp2b2 | 23771.2355 | 0.525336478 | 0.000944584 |
| Trio | 5078.80283 | 0.425166456 | 0.000946067 |
| Frmpd4 | 5992.80258 | 0.463821449 | 0.000946811 |
| Gpr61 | 501.230504 | 0.487433441 | 0.00095162 |
| Ap2a2 | 17152.4748 | 0.381752205 | 0.000952766 |
| Mkl1 | 2171.17548 | 0.349429318 | 0.00095383 |
| Rnf208 | 1719.4912 | 0.449344671 | 0.000954697 |
| Pcnx2 | 3099.34174 | 0.560740041 | 0.000956153 |
| Rc3h2 | 6255.12452 | 0.172609197 | 0.000957766 |
| Cd99 | 311.465564 | -0.591200753 | 0.000959956 |
| Kcnj3 | 913.834432 | 0.649592035 | 0.000964682 |
| Atmin | 2704.56299 | 0.327604175 | 0.000967457 |
| Gbp4 | 31.8847738 | -3.353657028 | 0.000971251 |
| Rap1a | 1271.79135 | -0.393697449 | 0.000972338 |
| Hapln3 | 119.332222 | -0.652250752 | 0.000973226 |
| Cd63 | 3023.44387 | -0.506543446 | 0.000977561 |
| Rab31 | 4808.64088 | -0.357264213 | 0.000978181 |
| Drap1 | 2269.72082 | -0.235065729 | 0.000978181 |
| Zc3hav1 | 1283.72355 | -1.142492459 | 0.000983423 |
| Nras | 1988.84412 | -0.41123927 | 0.00098673 |

| | | | |
|---|---|---|---|
| Lhfpl2 | 1362.97349 | -0.550755204 | 0.000993878 |
| Prkce | 7337.15598 | 0.580827205 | 0.000996953 |
| Grip1 | 1473.70282 | 0.327233753 | 0.001004792 |
| Dapk1 | 4568.77528 | 0.39453595 | 0.001010738 |
| Msn | 724.412162 | -0.802149596 | 0.001012262 |
| Glb1l | 743.834728 | -0.507624443 | 0.001013863 |
| Snx24 | 1027.29313 | -0.357739351 | 0.001013863 |
| Ptprn | 19725.0085 | 0.391420686 | 0.001015246 |
| Tapbpl | 1177.20467 | -0.899461655 | 0.001015597 |
| Kcnk2 | 1306.61243 | 0.42408469 | 0.001017183 |
| Acvr1c | 202.676836 | 0.574661815 | 0.001017183 |
| Fosl2 | 226.523114 | 0.97804832 | 0.001017183 |
| Gpr176 | 475.200967 | 0.537376523 | 0.00103204 |
| Fam8a1 | 2481.89027 | 0.250279857 | 0.001037873 |
| Mylk | 132.507538 | -0.932122577 | 0.001042603 |
| Dcaf4 | 437.657882 | -0.375439878 | 0.001045534 |
| Rasgrp1 | 14762.2147 | 0.669540801 | 0.001047293 |
| Ccser1 | 403.627523 | 0.378241274 | 0.00104763 |
| Ncs1 | 3161.20544 | 0.467411328 | 0.001049413 |
| Nell2 | 9424.19386 | 0.617514339 | 0.00105304 |
| Fcer1g | 24.5905615 | -1.567642621 | 0.001055899 |
| Kif13a | 2109.86834 | -0.414576174 | 0.001055899 |
| Bst2 | 106.08308 | -2.565527148 | 0.001058048 |
| Usp32 | 9332.19486 | 0.233435669 | 0.001062754 |
| Cdc42ep3 | 340.193716 | 0.39765653 | 0.001062754 |
| Mal | 1025.1267 | -1.108431051 | 0.001065841 |
| Erap1 | 1626.11538 | -0.598076799 | 0.00106948 |
| LOC500475 | 45.2323433 | -0.892789545 | 0.00106996 |
| Dbndd2 | 4972.33445 | -0.55536875 | 0.001078931 |
| Myt1l | 6301.08689 | 0.494816505 | 0.001078931 |
| Noct | 1481.00112 | 0.345934953 | 0.001097456 |
| Nudcd2 | 779.105589 | -0.405414772 | 0.001103243 |
| Ube4b | 5867.03488 | 0.274857129 | 0.001103243 |
| Rfng | 1036.7777 | 0.330974117 | 0.001103243 |
| Ss18 | 1933.74628 | -0.379517998 | 0.001106339 |
| Adcy3 | 3400.02217 | 0.47455495 | 0.001106723 |

| | | | |
|---|---|---|---|
| Lrig3 | 253.255275 | -0.46155527 | 0.001106955 |
| Shf | 446.382034 | 0.413562138 | 0.001109719 |
| Ybx1 | 3216.18672 | -0.377495409 | 0.001112928 |
| F10 | 5.3053012 | -3.399526525 | 0.00111828 |
| Cdk4 | 846.0409 | -0.381821725 | 0.001131478 |
| Hcn1 | 1869.97398 | 0.566931922 | 0.001131478 |
| Wdr7 | 10718.9024 | 0.321667206 | 0.001142132 |
| Gal3st3 | 808.436285 | 0.495070852 | 0.001142132 |
| Kif13b | 1970.68865 | -0.477971759 | 0.00115104 |
| Tesc | 384.058908 | 0.735688793 | 0.001156342 |
| Ctsb | 22796.0394 | -0.246933749 | 0.001165609 |
| Tsc1 | 4142.05366 | 0.22167877 | 0.001165744 |
| Plk2 | 5313.53838 | 0.582482142 | 0.001168077 |
| Nod1 | 938.713488 | -0.601694768 | 0.001177931 |
| Cntn3 | 509.684477 | 0.416996509 | 0.001183344 |
| LOC108352 | 7.18701882 | -3.272232422 | 0.001192813 |
| Slc25a13 | 208.932984 | -0.55618792 | 0.001196634 |
| Stat2 | 3303.72948 | -1.084320701 | 0.001197743 |
| Synpr | 3045.29859 | 0.409641793 | 0.001197743 |
| Hes6 | 1206.23197 | -0.38672378 | 0.001201777 |
| Nmi | 160.936569 | -0.790667294 | 0.001205148 |
| Sptbn4 | 5992.73212 | 0.459658552 | 0.001205148 |
| Sema6b | 4053.48599 | 0.446013904 | 0.001210516 |
| Htra3 | 429.082744 | -0.639799679 | 0.00121468 |
| Drd2 | 1770.12736 | 0.479946327 | 0.00121468 |
| Mecp2 | 8565.12712 | 0.209686288 | 0.001217193 |
| Srpk2 | 7595.01841 | 0.230296914 | 0.001220864 |
| Slc35f3 | 690.162606 | 0.491316391 | 0.001232768 |
| Nos1ap | 2197.50272 | 0.576848877 | 0.001252538 |
| Mcm5 | 42.9365707 | -0.918905525 | 0.001254629 |
| Rhog | 665.704284 | -0.700436632 | 0.001255438 |
| Syt5 | 2432.5195 | 0.554225541 | 0.001263308 |
| Csk | 1727.50786 | -0.271312787 | 0.001268024 |
| Clec16a | 4846.42412 | 0.181946223 | 0.001302647 |
| Fmn1 | 444.321212 | 0.499047115 | 0.001310789 |
| Adamts1 | 202.121457 | -0.716743302 | 0.001310904 |

| | | | |
|---|---|---|---|
| Epha6 | 2319.67391 | 0.379604988 | 0.001315012 |
| Hectd4 | 28354.489 | 0.305259366 | 0.001315809 |
| Fgf13 | 2975.31076 | 0.345153701 | 0.001315809 |
| LOC501038 | 332.738251 | 0.597940728 | 0.001324537 |
| Loxl4 | 506.339745 | -0.697323239 | 0.001325747 |
| Nrg2 | 263.91529 | 0.471922878 | 0.001340578 |
| Fuz | 782.470212 | -0.356680323 | 0.001344126 |
| Man1c1 | 1208.06036 | 0.443932041 | 0.001346453 |
| Soga1 | 2545.76507 | -0.396838352 | 0.0013523 |
| Spi1 | 8.45817988 | -2.508229961 | 0.001366259 |
| Aldh1l1 | 2761.43287 | -0.434050228 | 0.001366919 |
| Ankrd17 | 14224.4192 | 0.19104751 | 0.001367154 |
| Dusp16 | 919.830289 | -0.406698181 | 0.001372325 |
| Pml | 1716.847 | -0.69352173 | 0.001386773 |
| Col27a1 | 248.035757 | -0.627091116 | 0.001387816 |
| Zfp148 | 5364.03572 | 0.180146925 | 0.001387816 |
| Sik3 | 10449.9751 | 0.196969316 | 0.001387816 |
| Ptprj | 4442.16581 | 0.4519612 | 0.001387816 |
| B2m | 29124.4848 | -1.309066141 | 0.001390978 |
| Rbm43 | 192.465987 | -0.825294821 | 0.001390978 |
| Fam175a | 46.2289264 | -0.72928364 | 0.001390978 |
| Lcp1 | 152.943138 | -0.633757523 | 0.001390978 |
| Dmtn | 5604.13726 | 0.46392423 | 0.001390978 |
| Sv2c | 5954.75283 | 0.418739781 | 0.001392162 |
| Cby1 | 1387.03515 | -0.294898305 | 0.00139262 |
| Ifi47 | 128.159057 | -2.676044531 | 0.001398117 |
| Rimbp2 | 5096.17792 | 0.534071444 | 0.001398117 |
| Susd6 | 3766.82048 | -0.4684003 | 0.001398512 |
| Ttc19 | 2595.01617 | 0.391224706 | 0.001398512 |
| Antxr1 | 505.925421 | -0.67495862 | 0.001404133 |
| Zbtb7a | 5647.34855 | 0.263048168 | 0.001406443 |
| Kcnj11 | 795.096349 | 0.468526448 | 0.001406443 |
| Cers6 | 724.036086 | 0.460099213 | 0.00140822 |
| Nat6 | 1872.76134 | -0.413584179 | 0.001408762 |
| Chi3l1 | 7958.20538 | -0.499428332 | 0.001409489 |
| Ap2a1 | 10350.9171 | 0.296121663 | 0.001418039 |

| | | | |
|---|---|---|---|
| Car12 | 2669.06057 | 0.597579108 | 0.001419521 |
| Ankrd13b | 1657.35412 | 0.390538392 | 0.001421319 |
| Cd302 | 535.076598 | -0.421563129 | 0.001421681 |
| Sept3 | 2578.84877 | 0.466963957 | 0.001433349 |
| Zbp1 | 16.9933986 | -2.641314417 | 0.00143432 |
| Rnd3 | 1575.24991 | -0.528193275 | 0.001445695 |
| Adamts3 | 647.22192 | 0.488316962 | 0.001445695 |
| Itga7 | 1240.00764 | -0.51116081 | 0.001446633 |
| Bid | 191.592172 | -0.758577763 | 0.001447572 |
| Sgsm2 | 4225.41273 | 0.26106487 | 0.001454509 |
| Dgkz | 9372.7721 | 0.576955787 | 0.001454509 |
| Per1 | 2550.34343 | 0.521472683 | 0.001456609 |
| Ppp2r2c | 2635.34604 | 0.490893369 | 0.001457634 |
| Ap1s1 | 1756.46422 | 0.249079115 | 0.001462996 |
| Abcc5 | 2704.53433 | 0.356427484 | 0.001468009 |
| Prkar2b | 2848.95071 | 0.340834822 | 0.001472315 |
| Szrd1 | 2521.15213 | -0.3369568 | 0.001473097 |
| Dpp7 | 1837.10881 | -0.260819204 | 0.001473097 |
| Cd276 | 551.826301 | -0.436315064 | 0.001474113 |
| Celsr2 | 22825.6752 | 0.436340368 | 0.001476183 |
| P4ha2 | 537.909437 | 0.474563376 | 0.001479017 |
| Rhoa | 10545.0199 | -0.226542054 | 0.001497513 |
| Akap9 | 6848.02396 | 0.370776856 | 0.001497513 |
| Rida | 1998.47143 | -0.406852975 | 0.001501676 |
| Pstpip1 | 34.907926 | -0.778248198 | 0.001505383 |
| Ydjc | 411.366187 | 0.426495191 | 0.001508929 |
| Rnf122 | 34.9728829 | -0.997671166 | 0.001523865 |
| Actr2 | 10459.9263 | 0.209370024 | 0.001536692 |
| Mycbp2 | 10417.286 | 0.370568984 | 0.001542975 |
| Mfsd14b | 2070.51948 | -0.266240225 | 0.001545058 |
| Slitrk5 | 2738.11957 | 0.379927173 | 0.001545058 |
| Stx1b | 21355.0753 | 0.393067016 | 0.001545058 |
| Usp33 | 3946.911 | 0.251101343 | 0.001545454 |
| Adgrl1 | 21418.514 | 0.352674937 | 0.001545454 |
| Carmil2 | 1727.64121 | 0.481144435 | 0.001545454 |
| Trpm3 | 7456.53447 | -0.396313565 | 0.001561837 |

| | | | |
|---|---|---|---|
| Mt1 | 294.309712 | -0.870644865 | 0.001562457 |
| Fam171a2 | 4368.55207 | 0.397334781 | 0.00156343 |
| Nhsl1 | 4041.40575 | -0.415859434 | 0.001566416 |
| Zfp831 | 1030.86712 | 0.583478629 | 0.001566416 |
| Slc39a6 | 2708.68935 | -0.286248353 | 0.00157442 |
| Gad1 | 15330.948 | 0.349353893 | 0.00157442 |
| Lxn | 1592.45503 | -0.414185599 | 0.001580911 |
| Trim59 | 131.056956 | -0.869186326 | 0.001586302 |
| C1qtnf4 | 3188.92413 | 0.56978991 | 0.001600251 |
| Col5a1 | 78.1918562 | -0.769992119 | 0.001607619 |
| Spock2 | 21535.6187 | 0.37020561 | 0.001611169 |
| Zzef1 | 3733.32078 | 0.27307587 | 0.001611896 |
| Calb1 | 1671.7342 | 0.568322995 | 0.001618704 |
| Tmem88b | 226.901264 | -1.491656836 | 0.001620355 |
| Plcb3 | 1719.73616 | -0.455553659 | 0.001633692 |
| RGD156500 | 2463.39346 | -0.43342704 | 0.001635367 |
| Ampd3 | 6060.60214 | -0.431774473 | 0.001635367 |
| Ankrd34a | 3217.74866 | 0.408476906 | 0.001635367 |
| Mia3 | 4311.21066 | 0.275749203 | 0.001640833 |
| Pde8a | 142.026403 | -0.986094057 | 0.001665306 |
| Slc2a5 | 6.01016988 | -2.238704948 | 0.001670835 |
| Mafg | 3562.4201 | 0.362841053 | 0.001672876 |
| Plppr2 | 5878.66094 | 0.382600823 | 0.001672876 |
| Cd24 | 3549.01162 | -0.384186055 | 0.001680499 |
| Maml2 | 2614.41373 | -0.407757391 | 0.001686737 |
| Rhoj | 33.7407552 | -0.973829656 | 0.001691948 |
| Mcm2 | 470.201502 | -0.484207646 | 0.001691948 |
| Hoga1 | 635.770315 | -0.442741715 | 0.001693198 |
| Mmd | 3374.06108 | 0.503710986 | 0.00169565 |
| Ppp2r1b | 491.641171 | -0.45216802 | 0.001703586 |
| Myo5a | 17904.8612 | 0.430611974 | 0.001704434 |
| Col19a1 | 816.9535 | 0.677081792 | 0.001706541 |
| Sorbs1 | 11635.7419 | -0.362864785 | 0.001715667 |
| Cdk2 | 115.04033 | -0.605716439 | 0.001715853 |
| Isg20 | 21.6984482 | -2.144793147 | 0.001717385 |
| Bcl7a | 2910.84118 | 0.322328627 | 0.001717385 |

| | | | |
|---|---|---|---|
| Scn2a | 16092.7198 | 0.467045992 | 0.001718558 |
| Fam198a | 155.690871 | -0.626038662 | 0.001723057 |
| Soat1 | 983.054712 | -0.38853575 | 0.001723057 |
| LOC102546 | 136.465567 | 0.598151853 | 0.001723057 |
| Asl | 738.429744 | -0.322333 | 0.001733431 |
| RGD156111 | 40.4500449 | -1.120173928 | 0.001736445 |
| Slc2a4 | 60.9884215 | 1.436700499 | 0.001747131 |
| RT1-CE4 | 44.56381 | -2.076629458 | 0.001775402 |
| Ddr2 | 781.096522 | -0.957215659 | 0.001777486 |
| Kcnma1 | 3989.73018 | 0.41407218 | 0.001777867 |
| Rarres2 | 698.827785 | -0.531473489 | 0.001787649 |
| Ulk1 | 3452.43826 | 0.251733155 | 0.001794574 |
| Vcan | 587.542161 | -0.585743833 | 0.001795751 |
| Myh11 | 586.499928 | -0.536425513 | 0.00180232 |
| Dars | 2161.73049 | -0.309651724 | 0.001807996 |
| Stambpl1 | 956.731696 | 0.445691471 | 0.001807996 |
| Map9 | 5642.08585 | 0.409576785 | 0.001808782 |
| Plekha4 | 97.4405846 | -1.911167916 | 0.00181449 |
| Itgb2 | 14.1329808 | -1.770606297 | 0.001828561 |
| Itgb5 | 4263.04482 | -0.382533204 | 0.001828561 |
| Ppp3cb | 6655.0315 | 0.418171691 | 0.001828561 |
| Sult1a1 | 388.452602 | 0.727229578 | 0.001828561 |
| Pdzd4 | 3237.56746 | 0.384576551 | 0.001850321 |
| Cckbr | 667.720171 | 0.694401581 | 0.001866549 |
| Parp4 | 2125.81785 | -0.383194413 | 0.001867998 |
| Zmpste24 | 2074.31771 | -0.29226884 | 0.001873494 |
| Dpp9 | 2732.12264 | 0.286625735 | 0.00187473 |
| Banp | 956.015179 | 0.3657701 | 0.00188123 |
| Vps28 | 2729.40564 | -0.173188664 | 0.00188653 |
| Mob3a | 521.382981 | -0.466294817 | 0.001888122 |
| Ifitm2 | 68.3952482 | -1.0583106 | 0.001890496 |
| Tgfbr3 | 1106.80723 | 0.407475794 | 0.001892911 |
| Plcl2 | 2239.22238 | 0.445392004 | 0.001892911 |
| RT1-M3-1 | 719.940153 | -0.598492044 | 0.001909481 |
| Nol6 | 5821.83496 | 0.205425642 | 0.001909481 |
| Asb1 | 2963.37382 | 0.362250531 | 0.001909481 |

| | | | |
|---|---|---|---|
| Arhgap17 | 296.726527 | -0.540501253 | 0.001912905 |
| Wwtr1 | 1406.10376 | -0.492802663 | 0.00193195 |
| Cbfa2t3 | 1653.9162 | 0.368159179 | 0.001932781 |
| Myo1e | 755.921593 | -0.49496411 | 0.001932841 |
| Ube2c | 8.77396791 | -2.801877621 | 0.001935947 |
| Adgrg6 | 218.482478 | -0.960481105 | 0.001938169 |
| Arap1 | 3050.27272 | -0.340123928 | 0.001938169 |
| Ctxn1 | 6889.50966 | 0.505587015 | 0.001951621 |
| Tet1 | 613.025253 | -0.296370075 | 0.001953823 |
| Ermn | 758.760974 | -1.371154076 | 0.00195569 |
| Ahdc1 | 5714.77981 | 0.286810588 | 0.001962767 |
| Scn3a | 3392.87776 | 0.377704178 | 0.00196556 |
| Rnf165 | 916.041675 | 0.544170277 | 0.00196556 |
| Csrnp3 | 1315.64694 | 0.430062947 | 0.001967632 |
| Cnpy3 | 2400.44319 | -0.189237767 | 0.001970094 |
| LOC102551 | 368.318067 | 0.403853637 | 0.001971256 |
| Rnf114 | 1616.61353 | -0.6013203 | 0.001979228 |
| Leprotl1 | 858.595046 | -0.259188261 | 0.002004518 |
| Lpin2 | 3045.40515 | 0.241267013 | 0.002009162 |
| Pde1c | 1168.23135 | 0.481091152 | 0.002015118 |
| Rpe | 1298.27607 | -0.340250214 | 0.002019555 |
| Epcam | 31.9584705 | 0.906679537 | 0.002020693 |
| RGD130710 | 14251.036 | 0.259742424 | 0.002023655 |
| Shc2 | 906.333205 | 0.428106119 | 0.002026254 |
| Car13 | 74.9826034 | -0.915762041 | 0.002052698 |
| C1galt1 | 1283.25538 | -0.385775402 | 0.002059615 |
| Maged2 | 1976.83182 | -0.302622091 | 0.002087464 |
| Ubn1 | 2203.39554 | 0.266727047 | 0.002096253 |
| Dctn1 | 20570.0776 | 0.335163355 | 0.002096253 |
| Armcx3 | 4066.7046 | 0.218690344 | 0.002102262 |
| Josd2 | 462.251265 | -0.433669534 | 0.002110683 |
| Slc26a7 | 6.68471967 | 2.296981067 | 0.00211853 |
| Tpm1 | 3440.80898 | 0.482810639 | 0.002122782 |
| Chst7 | 240.917111 | -0.550418286 | 0.002126855 |
| Pogk | 1887.92341 | -0.261190596 | 0.002136028 |
| Plxdc1 | 287.114672 | 0.601673281 | 0.002140961 |

| | | | |
|---|---|---|---|
| Fkbp9 | 1675.81528 | -0.445615588 | 0.002156536 |
| Brsk2 | 2222.32466 | 0.37562509 | 0.002171152 |
| Clvs2 | 276.694595 | 0.415618947 | 0.002171152 |
| Tcf20 | 8770.23164 | 0.231236482 | 0.002175307 |
| Sema4f | 1499.2788 | 0.451098021 | 0.002180347 |
| Kcnc3 | 3382.94598 | 0.553074799 | 0.002194587 |
| Clu | 129970.444 | -0.464906816 | 0.002194889 |
| Nr1d1 | 6510.25471 | -0.467775357 | 0.002203671 |
| Ccdc92 | 2864.98141 | 0.369098006 | 0.002213969 |
| Trip6 | 246.151372 | -0.614590221 | 0.002215032 |
| Fam131b | 3659.12057 | 0.487479035 | 0.002215032 |
| Kif21a | 12159.2881 | 0.247809615 | 0.002217554 |
| Flot2 | 4517.93276 | -0.298524588 | 0.002231412 |
| Sobp | 3831.97377 | 0.359047406 | 0.002233991 |
| Slc6a9 | 5736.88601 | -0.516972079 | 0.002237771 |
| Ifi44l | 36.1017352 | -2.265885527 | 0.002241078 |
| Pus7l | 95.9712754 | -0.449774432 | 0.002241078 |
| Pip5k1c | 10129.6198 | 0.444823601 | 0.00224384 |
| Ppp1r12c | 3850.61949 | 0.333824216 | 0.002253216 |
| Syt7 | 7628.36694 | 0.455518468 | 0.002253216 |
| Car2 | 2636.83675 | -0.564624118 | 0.002253536 |
| Kcng4 | 2318.96005 | -0.604023362 | 0.002258156 |
| Ripk2 | 540.323922 | -0.510376744 | 0.002259425 |
| Vps13b | 4415.37548 | 0.220895185 | 0.002262089 |
| Cc2d1b | 1102.51771 | -0.271392627 | 0.00227866 |
| Srek1 | 2169.32168 | 0.192730013 | 0.002281972 |
| Dennd3 | 291.519877 | -0.566242472 | 0.002286183 |
| Vgf | 2259.18298 | 0.500204291 | 0.002300134 |
| Tln1 | 4324.98785 | -0.404748464 | 0.002301327 |
| Ajap1 | 961.976264 | 0.405990102 | 0.002301327 |
| Aga | 798.984213 | -0.377320311 | 0.002305587 |
| Cgrrf1 | 604.942786 | -0.407868025 | 0.002308111 |
| Mical2 | 5613.85725 | 0.563844902 | 0.002311455 |
| Chrm2 | 835.963004 | 0.389970352 | 0.002323937 |
| Shisa6 | 1979.45707 | 0.632876386 | 0.002323937 |
| Celf3 | 2633.85041 | 0.433509302 | 0.002325569 |

| | | | |
|---|---|---|---|
| Sh3bp1 | 287.918771 | 0.488413913 | 0.002325569 |
| Wdr13 | 5741.34869 | 0.321307733 | 0.00232952 |
| Rab15 | 3132.28548 | 0.520943639 | 0.00232952 |
| Gpr84 | 9.86978245 | -2.460302285 | 0.002332104 |
| Srpk1 | 2226.50282 | 0.216882011 | 0.002332104 |
| Socs7 | 6435.21479 | 0.438515137 | 0.002334118 |
| LOC100910 | 222.267406 | 0.502761843 | 0.002334118 |
| Bgn | 397.165443 | -0.579907386 | 0.002339218 |
| Osbp2 | 1968.10752 | 0.506139866 | 0.002356594 |
| Ttbk1 | 2590.40515 | 0.450884731 | 0.00235833 |
| Syde1 | 371.063783 | -0.406232466 | 0.002364427 |
| Shc1 | 456.731619 | -0.394568905 | 0.002367298 |
| Ocrl | 1806.47877 | 0.297877415 | 0.002367298 |
| Sil1 | 886.260891 | -0.306174086 | 0.002374663 |
| Bak1 | 295.767974 | -0.647303523 | 0.002386844 |
| Evc2 | 287.878317 | -0.4424179 | 0.002386844 |
| Sfpq | 5081.84445 | 0.251333039 | 0.002386844 |
| Frmd6 | 695.753808 | 0.376704672 | 0.002386844 |
| Pcsk1 | 225.983386 | 0.456265607 | 0.002386844 |
| Lmbrd2 | 2778.07867 | 0.261057406 | 0.002392641 |
| Brsk1 | 7451.45519 | 0.474798247 | 0.002392641 |
| Jakmip1 | 6626.04345 | 0.337154346 | 0.002403003 |
| Cplx2 | 9732.54524 | 0.459651382 | 0.002403003 |
| Clic4 | 872.856037 | -0.615856449 | 0.00241229 |
| Aamdc | 235.557319 | -0.486925021 | 0.002417935 |
| Golga7 | 1707.25701 | -0.32016395 | 0.002421458 |
| Adra2b | 40.9950631 | 0.743432779 | 0.002446309 |
| Gas1 | 1534.93308 | -0.363136786 | 0.002458467 |
| Arfgef1 | 5184.30267 | 0.251525233 | 0.002481373 |
| LOC102557 | 13.1253023 | -1.615480412 | 0.00249479 |
| Rbfox2 | 3762.95452 | 0.310205859 | 0.00249479 |
| Tulp4 | 13849.7702 | 0.291789465 | 0.002495506 |
| Ppfia2 | 3082.36963 | 0.357134217 | 0.002495506 |
| Slc8a1 | 20035.8511 | 0.516956224 | 0.002498742 |
| S100a1 | 2334.18403 | -0.364626863 | 0.002500266 |
| Stard9 | 191.691368 | -0.592090265 | 0.002513802 |

| | | | |
|---|---|---|---|
| Cdk5r2 | 4600.03691 | 0.525646324 | 0.002525219 |
| Strn3 | 2542.14368 | 0.179131416 | 0.002526559 |
| Ube2o | 7092.65985 | 0.323935702 | 0.00253835 |
| LOC365985 | 3978.08253 | 0.524993466 | 0.00253835 |
| Ifi30 | 292.798482 | -0.504080745 | 0.002547279 |
| LOC100909 | 39.1892775 | -0.728812911 | 0.002552799 |
| Padi2 | 3068.49194 | -0.543493029 | 0.002585712 |
| Actl6b | 1341.16704 | 0.397744863 | 0.002596573 |
| Rest | 893.680414 | -0.445031752 | 0.002604099 |
| Ctnnal1 | 787.657268 | -0.348890054 | 0.002604839 |
| Mvb12a | 330.580334 | -0.44283735 | 0.002605037 |
| LOC108348 | 207.994652 | -0.384621918 | 0.002608165 |
| Ncapd2 | 625.734257 | -0.379500436 | 0.002627627 |
| Syt12 | 689.739981 | 0.430286401 | 0.002627627 |
| Sparc | 32295.8804 | -0.578834423 | 0.002637628 |
| Arhgef26 | 2752.48521 | -0.434049071 | 0.002640929 |
| Dok6 | 1262.22409 | 0.450824793 | 0.002641582 |
| LOC108350 | 32.3277616 | 0.85864238 | 0.002644275 |
| Efs | 1521.54693 | -0.393088773 | 0.002649493 |
| Hmgxb4 | 650.562681 | -0.295125523 | 0.002650957 |
| Ephx4 | 354.300254 | 0.61260634 | 0.002650957 |
| Fyn | 6227.04031 | -0.221440638 | 0.002666573 |
| Ckap2 | 80.8557539 | -0.712587524 | 0.00267989 |
| Ptgds | 477.394205 | -1.417063829 | 0.002699098 |
| Gpr37l1 | 31884.6752 | -0.414174922 | 0.002699098 |
| Atl1 | 4056.82708 | 0.380369948 | 0.002699098 |
| Rapgefl1 | 4513.05391 | 0.514230798 | 0.002699154 |
| Sp9 | 616.435496 | 0.466918105 | 0.002700377 |
| Mxd1 | 1764.91507 | -0.389949795 | 0.002714273 |
| Pgap2 | 506.255583 | -0.343340096 | 0.002716288 |
| Fam46c | 31.2341607 | -0.76033905 | 0.00271937 |
| Dlx6 | 167.852873 | 0.594620041 | 0.002720024 |
| Pdgfc | 693.711116 | -0.517747867 | 0.002720866 |
| Zfp280d | 1837.85006 | 0.29718158 | 0.002722509 |
| Arih1 | 4178.84719 | 0.184621758 | 0.002732834 |
| LOC102549 | 258.925129 | 0.542412954 | 0.002736232 |

| | | | |
|---|---|---|---|
| Tmed5 | 1258.55615 | -0.340274591 | 0.00274051 |
| Herc1 | 16149.3561 | 0.328743124 | 0.00274051 |
| Klf3 | 1565.35348 | -0.340857317 | 0.002763184 |
| Hnrnpf | 6423.39919 | -0.315826664 | 0.002766005 |
| Pgf | 104.331572 | -0.663704916 | 0.002775051 |
| Tnfaip3 | 675.408074 | -0.533710687 | 0.002789991 |
| Mcl1 | 948.39738 | -0.511507253 | 0.002790681 |
| Tubgcp6 | 1555.06485 | 0.223527331 | 0.002790681 |
| Frrs1l | 4413.46365 | 0.463173907 | 0.002821251 |
| Rarg | 381.912748 | -0.386330803 | 0.002841396 |
| Rnf38 | 2143.7101 | -0.240257813 | 0.002848058 |
| Pip5k1a | 2323.44523 | 0.312860225 | 0.002848058 |
| Reps2 | 5387.0897 | 0.408380329 | 0.002855196 |
| Tmem63a | 175.490959 | -1.3030177 | 0.002869729 |
| Slc26a2 | 138.811318 | -0.731350653 | 0.002876076 |
| Senp2 | 2358.57512 | 0.264606762 | 0.002877056 |
| LOC102554 | 67.4267007 | 0.603800228 | 0.002877056 |
| Agap3 | 6711.72934 | 0.368733744 | 0.002888416 |
| Prtfdc1 | 1910.23478 | -0.432545753 | 0.002899035 |
| Sept2 | 10645.7601 | -0.345455617 | 0.002899187 |
| Arhgap30 | 7.21091581 | -2.333392314 | 0.002899969 |
| Sox2 | 6686.44489 | -0.348390024 | 0.002905687 |
| Kif1b | 44958.1844 | 0.216419063 | 0.002905687 |
| Fam117b | 3920.22843 | 0.267619565 | 0.002905687 |
| Atp9a | 13480.9678 | 0.365427155 | 0.002919574 |
| Chrnb2 | 2143.71875 | 0.38930191 | 0.002930628 |
| Nuak2 | 458.252366 | -0.606519718 | 0.002959667 |
| Ankrd45 | 1349.70227 | 0.354967035 | 0.002965596 |
| LOC103693 | 499.43849 | 0.549096135 | 0.002979654 |
| Tep1 | 1516.93597 | -0.423329733 | 0.002984785 |
| LRRTM1 | 1655.034 | 0.417417591 | 0.002984785 |
| Nexn | 478.240649 | 0.518651524 | 0.002984785 |
| Xpo1 | 3815.50542 | 0.255102279 | 0.002988255 |
| Kazn | 2562.74391 | 0.453843594 | 0.002988255 |
| St6galnac3 | 192.31781 | -0.675871345 | 0.003021856 |
| Tgif1 | 184.217708 | -0.646128975 | 0.003023425 |

| | | | |
|---|---|---|---|
| Stk10 | 269.669972 | -0.556457649 | 0.003024791 |
| Ttc7a | 1111.30452 | -0.432119896 | 0.003029282 |
| St6gal1 | 647.247581 | -0.329150281 | 0.003029282 |
| Sfrp2 | 169.692004 | -1.004148712 | 0.003031637 |
| Armc1 | 2218.03477 | 0.191240208 | 0.003046968 |
| Ociad2 | 446.741658 | 0.538509379 | 0.003046968 |
| Cd2ap | 274.584973 | -0.35278469 | 0.003056224 |
| C1rl | 44.6203562 | -0.916050477 | 0.003082202 |
| Slitrk4 | 1380.5221 | 0.47032455 | 0.003084882 |
| Tcf7l1 | 866.264274 | -0.374905928 | 0.003100373 |
| Tnpo2 | 7327.25108 | 0.254039165 | 0.00310347 |
| Htr1d | 82.329257 | 0.585825456 | 0.00310347 |
| Vstm2a | 2743.49583 | 0.277728813 | 0.003116388 |
| Chaf1b | 14.4901131 | -0.955246155 | 0.003119376 |
| S100a13 | 1766.2412 | -0.42482187 | 0.003119376 |
| Nup58 | 1773.3682 | 0.242731945 | 0.003119376 |
| Atp6v0a1 | 19222.2605 | 0.277125467 | 0.003119376 |
| Prickle2 | 8830.2267 | 0.345548309 | 0.003119376 |
| Arhgap6 | 912.005027 | 0.405124915 | 0.003119376 |
| Tmem220 | 69.2705647 | -0.538805509 | 0.003124008 |
| Ogfod1 | 1565.81372 | 0.400491051 | 0.003124008 |
| Fam174b | 480.609962 | 0.483142731 | 0.003124008 |
| Prcp | 379.331146 | -0.525974559 | 0.003132483 |
| Ugt8 | 462.279346 | -0.88832127 | 0.003132666 |
| Man1a2 | 2430.59337 | 0.237863916 | 0.003139722 |
| Sox4 | 878.708128 | -0.344723337 | 0.003163396 |
| Cacnb3 | 3151.01462 | 0.426868502 | 0.00316865 |
| Msmo1 | 6298.86756 | -0.372821751 | 0.003168674 |
| Snph | 8333.23516 | 0.403241747 | 0.003168674 |
| Syt10 | 462.588942 | 0.548594432 | 0.003170135 |
| Nckap5 | 920.950746 | -0.455299048 | 0.003213605 |
| Atrnl1 | 5441.99181 | 0.533192921 | 0.003213605 |
| Smap2 | 6811.51429 | 0.387355888 | 0.003236023 |
| Hace1 | 1440.12621 | 0.373251758 | 0.003275554 |
| Hspa8 | 66673.3003 | 0.212741764 | 0.003287156 |
| Glmp | 1161.72231 | -0.301461724 | 0.00329916 |

| | | | |
|---|---|---|---|
| Socs5 | 2970.2093 | 0.274987206 | 0.003299565 |
| St8sia3 | 1946.05831 | 0.459916739 | 0.003300674 |
| P4htm | 2371.00786 | 0.313119692 | 0.003300761 |
| Tmbim1 | 1350.44562 | -0.564269148 | 0.003315279 |
| Tyro3 | 7490.60269 | 0.455938718 | 0.003342084 |
| Sox9 | 15629.8576 | -0.3633956 | 0.003342319 |
| Ptafr | 19.8428314 | -1.343234313 | 0.003353557 |
| Unc93b1 | 126.770981 | -0.8644395 | 0.003353557 |
| Nuak1 | 897.514767 | 0.490109642 | 0.003385348 |
| Pdzrn3 | 3854.96075 | -0.305073802 | 0.003386176 |
| Plxna2 | 9219.36625 | 0.322038452 | 0.003386176 |
| Ptpn3 | 531.642841 | 0.797334752 | 0.003386176 |
| Hmox1 | 521.830933 | -0.663292085 | 0.003388931 |
| Diras2 | 9765.14194 | 0.587615211 | 0.003410444 |
| LOC108350 | 1467.20593 | 0.374922435 | 0.003412417 |
| Psd3 | 19898.6302 | 0.451451088 | 0.003412417 |
| Anxa1 | 126.664012 | -1.459299366 | 0.003416118 |
| Golga4 | 5096.79137 | 0.287215015 | 0.003429284 |
| Snrpn | 3881.17101 | 0.388709798 | 0.003432377 |
| Dram1 | 26.4125051 | -1.23530112 | 0.003437372 |
| Gpr37 | 5705.32992 | -0.507891316 | 0.003447222 |
| Dennd5b | 2947.11207 | 0.23317501 | 0.003451842 |
| Nfx1 | 3408.79797 | 0.217144614 | 0.00348007 |
| Palm | 6631.36951 | 0.274010594 | 0.003480747 |
| LOC102550 | 41.5824348 | -0.879223058 | 0.003483394 |
| Adss | 2741.2785 | -0.212001215 | 0.003489271 |
| Ccar1 | 1221.65261 | 0.259658362 | 0.003491675 |
| Ybx3 | 547.602135 | -0.502454 | 0.003504247 |
| Zmym2 | 3750.57549 | 0.176797092 | 0.003504247 |
| Usp20 | 3712.25023 | 0.233063056 | 0.003507383 |
| Ogfr | 1557.31096 | -0.468456764 | 0.003510762 |
| Opn3 | 412.352799 | 0.527061457 | 0.003520146 |
| Tubb6 | 229.357495 | -1.124267679 | 0.003528262 |
| Sumf2 | 641.078449 | -0.380047592 | 0.003531643 |
| Kifc2 | 4180.81102 | 0.508048142 | 0.003552358 |
| Smpdl3a | 1177.59342 | -0.438520335 | 0.003554878 |

| | | | |
|---|---|---|---|
| Ankrd13a | 2223.44293 | -0.286156522 | 0.003554878 |
| Mib2 | 4963.8627 | 0.220691245 | 0.003556427 |
| Echdc1 | 2717.44393 | -0.346564158 | 0.003557024 |
| Pom121 | 5905.26866 | 0.291951214 | 0.003574836 |
| Vezf1 | 3352.09678 | -0.289382897 | 0.003575624 |
| Tpr | 6852.00654 | 0.203767501 | 0.003575624 |
| Abca5 | 1309.51749 | 0.346655204 | 0.003575624 |
| Npas2 | 3102.53297 | 0.40506541 | 0.003580209 |
| Ppargc1b | 742.763418 | 0.386978478 | 0.003582135 |
| Cacna1h | 1577.61301 | 0.421256701 | 0.003582641 |
| Creb3l1 | 308.765537 | -0.511632851 | 0.003592299 |
| Rtn4rl1 | 2002.95961 | 0.357846796 | 0.003617497 |
| Cnih2 | 3812.80504 | 0.431732786 | 0.003618279 |
| Efna3 | 496.175894 | 0.551851884 | 0.003623299 |
| Rassf4 | 171.344436 | -0.714250627 | 0.003628446 |
| Wdr17 | 859.132306 | 0.515251369 | 0.003648914 |
| Havcr2 | 14.4541115 | -1.487290729 | 0.003652371 |
| Bdp1 | 3064.8808 | 0.229918749 | 0.003652371 |
| Angptl2 | 51.4957081 | -0.724085731 | 0.003658318 |
| Slk | 4934.25245 | 0.21271941 | 0.003658318 |
| Ubxn1 | 3749.06052 | -0.279453575 | 0.00366088 |
| Ttll11 | 793.244421 | 0.400265181 | 0.00366088 |
| Nanos1 | 602.035305 | 0.349706051 | 0.003665531 |
| Lrrc42 | 1011.77432 | -0.271358906 | 0.003671277 |
| Ehd1 | 3231.05608 | -0.187260932 | 0.003688712 |
| Slc30a4 | 2855.63495 | 0.24852529 | 0.003688712 |
| Fnip1 | 2570.6986 | 0.188971659 | 0.003694198 |
| Prkch | 401.545442 | 0.55459901 | 0.00370866 |
| Oasl | 81.437562 | -2.180184 | 0.003710356 |
| Ctsz | 891.223608 | -0.32747897 | 0.003712465 |
| Ndrg4 | 57236.4689 | 0.387881157 | 0.003723278 |
| Elf2 | 1089.06135 | -0.290768058 | 0.003755842 |
| Ift140 | 2787.56373 | -0.284300234 | 0.003755842 |
| Kdm4a | 1857.41355 | -0.214616197 | 0.003755842 |
| Tac3 | 433.378995 | 0.522502032 | 0.003755842 |
| Slc16a14 | 560.555566 | 0.574986598 | 0.003755842 |

| | | | |
|---|---|---|---|
| Snap23 | 354.262614 | -0.369095747 | 0.003776095 |
| Opa1 | 7692.21438 | 0.238548093 | 0.003776095 |
| Tbk1 | 1100.7956 | 0.296437887 | 0.003776095 |
| Islr2 | 2689.55049 | 0.510112703 | 0.003776095 |
| Ppp1r12a | 4977.52487 | 0.19083726 | 0.003779459 |
| Tmem132a | 10046.6868 | 0.316753279 | 0.003794238 |
| Ywhaz | 52976.1347 | 0.254730037 | 0.003810902 |
| Arhgap33 | 4584.85356 | 0.444331771 | 0.003816169 |
| Lsm14b | 3919.08867 | 0.193969234 | 0.003852007 |
| Tcf12 | 3624.95992 | -0.303245212 | 0.003856117 |
| Ubtf | 5140.28697 | 0.180385191 | 0.003856117 |
| Tmem143 | 334.679091 | 0.325305049 | 0.003863059 |
| Sacs | 6074.02549 | 0.390334687 | 0.003863059 |
| Glcci1 | 802.53805 | 0.304981824 | 0.003868916 |
| Acot1 | 114.852599 | -0.791090156 | 0.003871452 |
| Bend3 | 469.280019 | -0.360845622 | 0.003876705 |
| Trak2 | 5778.27199 | 0.314040671 | 0.003883342 |
| Nxph1 | 2934.90962 | 0.361200752 | 0.003888154 |
| Rgs7bp | 7798.94282 | 0.434913319 | 0.003888154 |
| Triobp | 1174.73337 | -0.407690864 | 0.003916839 |
| Sugp2 | 4682.70083 | 0.196226718 | 0.003924866 |
| Prdm15 | 415.62106 | 0.330803093 | 0.003927399 |
| Myo9a | 6179.86117 | 0.215986738 | 0.003934991 |
| Far2 | 790.644234 | 0.494137885 | 0.003934991 |
| Nalcn | 5303.85878 | 0.364492153 | 0.00395534 |
| LOC102547 | 466.171526 | 0.426688773 | 0.003966806 |
| Gyg1 | 1364.44844 | -0.301099956 | 0.003970038 |
| Tsc22d2 | 2308.87326 | 0.327677098 | 0.00398263 |
| Chsy1 | 1748.81705 | -0.314613437 | 0.00400702 |
| Twf1 | 2329.65274 | -0.298455563 | 0.00400702 |
| Rhot2 | 1533.2657 | 0.23414361 | 0.004015406 |
| Rhobtb2 | 4566.42362 | 0.217361258 | 0.004015482 |
| Rprd1a | 3846.61604 | 0.16176688 | 0.004025909 |
| Shank2 | 947.108084 | 0.450674812 | 0.004031524 |
| Akap13 | 1310.70566 | -0.220213727 | 0.004095887 |
| Aprt | 415.518 | -0.330012543 | 0.004109521 |

| | | | |
|---|---|---|---|
| Myh10 | 17748.6221 | 0.368970144 | 0.004109521 |
| Golgb1 | 4632.89315 | 0.298255499 | 0.004110893 |
| Snap91 | 13723.7235 | 0.419565952 | 0.004114547 |
| Prr12 | 5814.49335 | 0.228593027 | 0.004117492 |
| Nol4 | 1203.36485 | 0.353024116 | 0.004119556 |
| Cflar | 1559.34814 | -0.425560947 | 0.004121043 |
| Arl4d | 401.388958 | 0.495601513 | 0.004122848 |
| Herc2 | 15367.5808 | 0.30116577 | 0.004160633 |
| Asah1 | 1898.8825 | -0.299398189 | 0.004184647 |
| Plekhd1 | 1721.30795 | -0.406869739 | 0.004232569 |
| Lyz2 | 94.0853064 | -1.551798079 | 0.00423324 |
| Lmo1 | 355.072657 | -0.456136227 | 0.00423324 |
| Zdhhc14 | 1215.97046 | 0.369366742 | 0.004248662 |
| Mcf2l | 4441.68867 | 0.275100449 | 0.004256067 |
| Dnm1 | 33760.8949 | 0.411784718 | 0.00429201 |
| Ighmbp2 | 929.734068 | 0.208463965 | 0.004308864 |
| Arhgap39 | 2112.70654 | 0.292650121 | 0.004309336 |
| Pcdh15 | 199.482639 | -0.434766752 | 0.004317491 |
| Atf2 | 3847.91581 | 0.275848435 | 0.004317491 |
| Ppp1r7 | 4214.75809 | 0.312110222 | 0.004329224 |
| Wdr47 | 5205.86452 | 0.325154086 | 0.004329224 |
| Gse1 | 4687.19152 | 0.284110315 | 0.004339629 |
| Irak3 | 128.681383 | -0.673125274 | 0.004342623 |
| Scg2 | 14647.053 | 0.330987634 | 0.004346536 |
| Dbnl | 2840.80047 | -0.309514106 | 0.004353331 |
| LOC103691 | 1346.57795 | 0.360455291 | 0.004358959 |
| Cpne3 | 1384.8304 | -0.394305499 | 0.004378201 |
| Usp9x | 25019.9193 | 0.198107821 | 0.004385761 |
| Kif3b | 3506.52881 | 0.228206432 | 0.004385761 |
| LOC100911 | 1513.58799 | 0.234749242 | 0.004385761 |
| Atg16l1 | 1423.82434 | 0.299847649 | 0.004385761 |
| Fam160b2 | 1290.8544 | 0.251488581 | 0.00438674 |
| Map1b | 96190.6672 | 0.445855676 | 0.004410194 |
| Clec11a | 228.618686 | 0.575103065 | 0.004420324 |
| Pdgfb | 164.565319 | 0.669982156 | 0.004422322 |
| Cpxm1 | 184.610695 | -0.543511638 | 0.004440144 |

| | | | |
|---|---|---|---|
| LOC100362 | 120.801504 | -0.689941185 | 0.00444117 |
| Zyg11b | 8766.14896 | 0.209707058 | 0.00445737 |
| Parvb | 5606.88837 | -0.388435719 | 0.004462181 |
| Spin4 | 66.6046758 | -0.655067361 | 0.004469485 |
| Emp1 | 136.220952 | -0.649477954 | 0.004469485 |
| Pomt1 | 1220.82039 | -0.296659076 | 0.004483583 |
| Stat5a | 257.437709 | -0.475726429 | 0.004483619 |
| Wnk2 | 16277.5028 | 0.254964646 | 0.004483619 |
| H3f3b | 6915.03657 | -0.338568185 | 0.004494068 |
| Sri | 2756.89914 | -0.186743494 | 0.004519467 |
| Asb2 | 56.2843886 | 0.61694185 | 0.00452613 |
| March6 | 11167.493 | 0.223077148 | 0.004543307 |
| Qser1 | 2980.19094 | -0.294239308 | 0.004558724 |
| Mfsd13a | 708.074334 | -0.297120511 | 0.004568882 |
| Ip6k2 | 1794.32625 | 0.275334543 | 0.004568882 |
| Snx5 | 3618.11415 | -0.282833365 | 0.004569253 |
| Phpt1 | 1074.44195 | -0.237136448 | 0.004573383 |
| Casp12 | 11.2088372 | -2.062634552 | 0.004573766 |
| Galns | 846.628125 | -0.289858022 | 0.004582751 |
| G6pd | 3656.36103 | -0.192135216 | 0.004582751 |
| Abhd8 | 6091.85365 | 0.344258508 | 0.00459924 |
| Hspa5 | 23185.6885 | 0.366843012 | 0.004604891 |
| Srrt | 1219.01008 | 0.248271272 | 0.004610649 |
| L1cam | 9451.94493 | 0.328876435 | 0.00461243 |
| Dlgap1 | 13982.0564 | 0.439863747 | 0.00461243 |
| Alpk3 | 53.2517205 | -0.696495074 | 0.004638083 |
| Pxn | 860.121166 | -0.33350721 | 0.004655042 |
| Trpc1 | 1037.8758 | 0.225514142 | 0.004655042 |
| Kmt2a | 5907.79603 | 0.301530661 | 0.004655042 |
| Prr36 | 2860.69102 | 0.324343166 | 0.004655042 |
| Sec14l4 | 20.3061946 | -1.57584913 | 0.004661986 |
| Lyn | 141.676018 | -0.741906215 | 0.004675043 |
| Stim2 | 2342.72494 | 0.293703987 | 0.004675043 |
| Phyhd1 | 778.664268 | -0.552884804 | 0.004698837 |
| Cx3cr1 | 49.5632185 | -0.760904852 | 0.004704851 |
| Zmiz1 | 15024.6463 | 0.192273863 | 0.004726322 |

| | | | |
|---|---|---|---|
| Jam3 | 1510.36741 | -0.440231859 | 0.004740384 |
| Lmf2 | 1143.4557 | -0.335485063 | 0.004755878 |
| Dera | 470.900273 | -0.385588319 | 0.004781731 |
| Tram1 | 1976.65772 | -0.256361356 | 0.004781731 |
| Macrod2 | 1034.62498 | 0.300970709 | 0.004781731 |
| Zwint | 93450.7005 | 0.307026467 | 0.004781731 |
| RGD131020 | 668.474803 | 0.400799644 | 0.004796359 |
| Crtac1 | 3530.12034 | 0.427573939 | 0.00485785 |
| LOC102547 | 8.94553879 | -1.101575812 | 0.004867881 |
| Foxs1 | 13.6531012 | -1.789381923 | 0.004904846 |
| March3 | 313.702905 | -0.458745059 | 0.004904846 |
| Kifap3 | 8711.47758 | 0.320288385 | 0.004921191 |
| Lgals8 | 1154.9492 | -0.261912255 | 0.004921215 |
| S1pr5 | 329.003887 | -0.837767133 | 0.004921676 |
| Nsf | 22567.048 | 0.404902853 | 0.004924384 |
| Noa1 | 848.273407 | 0.218121232 | 0.004932318 |
| Snurf | 9531.41748 | 0.353820835 | 0.004961204 |
| Eef2k | 2877.3097 | -0.345492747 | 0.004984785 |
| Flna | 2291.04904 | -0.753782442 | 0.004990904 |
| Slc12a4 | 1628.96909 | -0.435293791 | 0.004990904 |
| Dnajc6 | 13286.2925 | 0.415529671 | 0.004997232 |
| Hyou1 | 9467.27818 | 0.279629802 | 0.005001023 |
| Nrn1 | 524.955105 | 0.866441589 | 0.005002757 |
| Skap2 | 556.388567 | -0.372454667 | 0.005003483 |
| Lin7b | 284.856573 | 0.468616582 | 0.005019511 |
| Wdr1 | 10681.1439 | -0.246079127 | 0.00502505 |
| Cd164 | 5277.96392 | -0.272978917 | 0.005044422 |
| Strn4 | 4855.58014 | 0.285754831 | 0.005049077 |
| Plekhg5 | 1324.82908 | 0.554548658 | 0.005049077 |
| Raly | 4025.54599 | -0.236588461 | 0.005069141 |
| Rac2 | 10.9026071 | -1.950291913 | 0.005069936 |
| Vipr1 | 185.649238 | 0.630809833 | 0.00507038 |
| Kcna1 | 1524.95225 | 0.553986973 | 0.005083578 |
| Gprin1 | 4119.86483 | 0.426617289 | 0.005095928 |
| Lonrf1 | 606.363394 | 0.477640291 | 0.005097137 |
| Birc2 | 1741.08296 | -0.299772899 | 0.005101894 |

| | | | |
|---|---|---|---|
| Unc13a | 5587.51388 | 0.34730732 | 0.005130891 |
| Atf5 | 3211.4272 | -0.475189775 | 0.005133256 |
| Tmem260 | 1163.37664 | 0.204354536 | 0.005133256 |
| Zfyve16 | 821.55172 | -0.355642995 | 0.005136953 |
| Ptprc | 18.0719457 | -1.59317337 | 0.005148287 |
| Ctdsp1 | 2140.69675 | -0.339896774 | 0.005148287 |
| Adck3 | 915.838685 | 0.276932936 | 0.005148287 |
| Lpcat4 | 1598.91488 | 0.434672122 | 0.00515158 |
| Tsku | 280.342784 | -0.452258921 | 0.005172268 |
| Lactb | 849.350072 | -0.291730885 | 0.005172937 |
| Rusc2 | 7415.6381 | 0.206427924 | 0.005172937 |
| Rtcd1 | 1514.62333 | -0.203803416 | 0.005205413 |
| Ppp1r37 | 3607.59367 | 0.303904801 | 0.005212867 |
| Unc5a | 2162.85863 | 0.521041053 | 0.005212867 |
| Map1s | 3525.6496 | 0.232108474 | 0.005220531 |
| Phgdh | 13489.9868 | -0.396628669 | 0.005230276 |
| Cyp2j3 | 2311.67806 | -0.378375897 | 0.005236218 |
| Pttg1ip | 5099.97846 | -0.322327708 | 0.005236218 |
| Irf2 | 946.964065 | -0.365652441 | 0.005243775 |
| Fermt2 | 10249.3541 | -0.350802372 | 0.005243775 |
| Slc8a2 | 7179.91225 | 0.424804003 | 0.005256577 |
| Atp1b2 | 68698.7752 | -0.369644802 | 0.005260224 |
| Scnn1a | 168.356334 | -0.610762423 | 0.00529959 |
| Ppfia3 | 5370.28267 | 0.41122584 | 0.005301582 |
| Slc7a11 | 20968.5453 | -0.442847898 | 0.005308997 |
| Camk1d | 4822.65973 | 0.330638582 | 0.005319037 |
| Slc6a17 | 17012.0371 | 0.469759029 | 0.005321359 |
| Dtnb | 1865.97905 | 0.323381483 | 0.005322772 |
| Naip6 | 9.02153514 | -1.246305974 | 0.00533757 |
| Ap3m1 | 1673.18897 | -0.311218365 | 0.005342844 |
| Kif1a | 85821.7268 | 0.253533297 | 0.005342844 |
| Adprhl2 | 1660.85532 | -0.299421933 | 0.005366244 |
| Pdxk | 1417.01239 | 0.303296276 | 0.005367759 |
| Parp6 | 1428.03488 | 0.365630608 | 0.005368815 |
| Nr1h3 | 178.66705 | -0.548129976 | 0.005384789 |
| Pmp22 | 640.147713 | -0.526958867 | 0.005384789 |

| | | | |
|---|---|---|---|
| Sh3bgrl | 8210.63966 | -0.303046258 | 0.005384789 |
| Nthl1 | 96.2415251 | -0.498027431 | 0.005391408 |
| Shroom1 | 149.101491 | 0.467514967 | 0.005396242 |
| Nolc1 | 6693.12214 | 0.19846128 | 0.005408945 |
| Tbc1d8 | 686.064243 | 0.336977721 | 0.005408945 |
| LOC102548 | 9.55147122 | 1.24146673 | 0.005414801 |
| Lrrcc1 | 180.020676 | -0.40196528 | 0.005424477 |
| Mdk | 1447.09721 | -0.316875048 | 0.005424477 |
| Cd9 | 1298.44496 | -0.738185071 | 0.005429095 |
| Snapin | 2186.41791 | -0.289527011 | 0.005433281 |
| Hmgcs1 | 18809.0728 | -0.365223842 | 0.005452946 |
| Ptprz1 | 34381.1867 | -0.43673342 | 0.005454778 |
| RGD130546 | 822.968676 | -0.364671498 | 0.005484048 |
| S100a4 | 315.375133 | -0.707174202 | 0.005486116 |
| Uspl1 | 1122.11471 | 0.292417088 | 0.005509008 |
| Dach1 | 365.567708 | 0.467947972 | 0.005541928 |
| Sbds | 3738.23852 | -0.253985625 | 0.005554738 |
| Calu | 5211.14999 | -0.277715528 | 0.005567776 |
| Rpl19 | 6086.64252 | -0.282208775 | 0.005581539 |
| Fam168b | 9756.11489 | 0.193692501 | 0.005581539 |
| Ttc28 | 2688.11783 | -0.325427301 | 0.005590997 |
| Odf2l | 313.724739 | -0.421122171 | 0.005596567 |
| Pip4k2b | 7378.72587 | 0.304613454 | 0.005598307 |
| Arpp19 | 5988.3288 | 0.301050438 | 0.005599804 |
| Hecw1 | 2168.51044 | 0.440750912 | 0.005622474 |
| Ednrb | 19583.8956 | -0.413417324 | 0.00563445 |
| Fdft1 | 4514.29808 | -0.309849325 | 0.005648878 |
| Gpr62 | 135.377354 | -0.678205259 | 0.005657524 |
| P2ry12 | 15.7706903 | -0.846683055 | 0.005677588 |
| Rcbtb2 | 1092.44544 | -0.325628943 | 0.0056795 |
| Myl12a | 1018.93885 | -0.501849494 | 0.005711744 |
| Npas3 | 1442.52693 | -0.439972494 | 0.005717129 |
| Foxk2 | 3807.7549 | 0.211915867 | 0.005718236 |
| Kcnt2 | 657.519927 | 0.401314634 | 0.005718236 |
| Kcnab2 | 9381.10227 | 0.459685605 | 0.005718236 |
| Scrib | 3247.29335 | -0.286768712 | 0.005727912 |

| | | | |
|---|---|---|---|
| Prdx1 | 3288.34503 | -0.316437417 | 0.005729325 |
| Frmpd3 | 2267.06448 | 0.388368134 | 0.005738163 |
| Cacng5 | 1219.2591 | 0.504845815 | 0.005744172 |
| Slit1 | 4868.48331 | 0.384007966 | 0.005748282 |
| Prrx1 | 1207.92634 | -0.326501639 | 0.005757121 |
| Paip2b | 1617.54868 | 0.212447709 | 0.005757121 |
| Vamp2 | 21139.1876 | 0.339455521 | 0.005766888 |
| Srd5a1 | 434.687877 | -0.453011157 | 0.005779516 |
| Cuedc2 | 273.72815 | -0.419738024 | 0.005788592 |
| Orai2 | 1381.02274 | 0.509985444 | 0.005788738 |
| LOC102547 | 59.8400916 | 1.823316393 | 0.005793821 |
| Gdf15 | 5.97635157 | -1.291364183 | 0.005798002 |
| Sec11a | 640.366396 | -0.279152775 | 0.005798002 |
| Hspa4 | 11731.6702 | 0.183323658 | 0.005803654 |
| Cmtm3 | 33.5475504 | -0.663456809 | 0.005831238 |
| Csrp1 | 15219.3546 | -0.386015132 | 0.005863274 |
| Huwe1 | 18135.6768 | 0.221255623 | 0.005896129 |
| Vps50 | 1172.8696 | 0.284453351 | 0.005896129 |
| Itm2b | 29225.9587 | -0.323843001 | 0.005902769 |
| Tmem243 | 186.452813 | -0.505126401 | 0.005946368 |
| Pim2 | 1343.69508 | 0.40490983 | 0.005972851 |
| Kctd8 | 479.468707 | 0.342769573 | 0.005973244 |
| Runx1t1 | 1244.95299 | 0.344787959 | 0.005973244 |
| LOC680142 | 135.834277 | 0.499132916 | 0.005973244 |
| Lrrc73 | 553.695485 | 0.389366398 | 0.006007757 |
| Pex3 | 1026.71325 | -0.183578312 | 0.006016671 |
| Ube2q2l | 272.503838 | 0.276222125 | 0.006016671 |
| Scara3 | 7665.36267 | -0.375603696 | 0.006018348 |
| Ddr1 | 6190.32036 | -0.37992409 | 0.006019273 |
| Pcp4 | 5218.71214 | 0.463293919 | 0.006021328 |
| LOC102547 | 113.519474 | -0.608095792 | 0.006030643 |
| Cd274 | 399.731508 | -0.999360273 | 0.006031302 |
| Nphp3 | 201.749366 | -0.40295098 | 0.006031302 |
| Pced1b | 260.437078 | -0.398976708 | 0.006038778 |
| Hs6st2 | 2777.45865 | 0.349321307 | 0.006038778 |
| Gria3 | 5091.67566 | 0.510689011 | 0.006038778 |

| | | | |
|---|---|---|---|
| Btbd3 | 3832.26271 | 0.46125521 | 0.006055854 |
| Kif5a | 22877.0567 | 0.354038251 | 0.006064418 |
| Entpd2 | 5508.30997 | -0.383266525 | 0.006070591 |
| Hsp90ab1 | 66007.5354 | 0.211362772 | 0.006103546 |
| Pnlip | 35.4026409 | -0.697898541 | 0.006108694 |
| Zfp236 | 1431.74662 | 0.208417027 | 0.006140258 |
| Sowahc | 636.795811 | -0.318943937 | 0.006147991 |
| Fyco1 | 1228.76285 | -0.311355946 | 0.006147991 |
| Prr5l | 45.3807849 | -0.85337795 | 0.006150179 |
| RT1-CE5 | 312.437038 | -0.901326841 | 0.006208576 |
| Thumpd3 | 1292.74284 | -0.223566161 | 0.006228363 |
| RGD15638 | 168.157557 | -0.630602043 | 0.006229068 |
| Clock | 3766.66356 | 0.162181424 | 0.006229913 |
| Gdap2 | 882.706206 | 0.284969567 | 0.006267394 |
| Hecw2 | 3309.67495 | 0.370223118 | 0.006267394 |
| Pigk | 2014.57662 | -0.229503526 | 0.006286967 |
| Rock2 | 9952.40531 | 0.349444618 | 0.006293444 |
| Hspa12a | 10839.4823 | 0.420492621 | 0.006293444 |
| Mgst1 | 958.40271 | -0.48905534 | 0.006296187 |
| Shank1 | 26216.0303 | 0.483935084 | 0.006296187 |
| Lcp2 | 18.2394962 | -0.946894807 | 0.006312945 |
| Cited2 | 1887.57033 | 0.302367693 | 0.006344325 |
| Mpp2 | 6794.76858 | 0.236326151 | 0.006346833 |
| Adra2c | 436.160931 | 0.486056317 | 0.0063476 |
| Ano10 | 1598.9102 | -0.210319118 | 0.006406384 |
| Micall1 | 1271.76714 | -0.346730125 | 0.006416725 |
| Rmnd5a | 4007.40158 | 0.18702542 | 0.006416725 |
| Mapre2 | 8859.35769 | 0.156478901 | 0.006426016 |
| Mapk10 | 7028.8419 | 0.42092107 | 0.006426016 |
| Ccl12 | 16.0681256 | -1.115505755 | 0.006441906 |
| Cmya5 | 185.77649 | 0.488880967 | 0.006441906 |
| RT1-T24-3 | 17.2236333 | -1.162969402 | 0.006445232 |
| Lamp1 | 21289.493 | -0.269858518 | 0.006503222 |
| Mocos | 82.9700913 | -0.759483041 | 0.006520116 |
| Arl15 | 793.686247 | 0.340954028 | 0.006541681 |
| Coq2 | 1433.89509 | 0.266648317 | 0.006575184 |

| | | | |
|---|---|---|---|
| Dnaja2 | 5656.08431 | 0.165143344 | 0.006586694 |
| Cdh13 | 8977.10565 | 0.216518925 | 0.006586694 |
| Syngr3 | 3737.81434 | 0.346108913 | 0.006586694 |
| Bicc1 | 140.929009 | -0.630032916 | 0.006595957 |
| Dock6 | 191.395414 | -0.537536425 | 0.006600684 |
| LOC100362 | 1650.37348 | -0.360858922 | 0.006600684 |
| Synj1 | 15039.7659 | 0.327163682 | 0.006620973 |
| Zfp317 | 982.479283 | -0.212476408 | 0.006667481 |
| Trex1 | 394.298416 | -0.539266007 | 0.006670823 |
| Lrrc24 | 929.895141 | 0.38368623 | 0.006677347 |
| LOC103692 | 1439.97942 | -0.438710655 | 0.006680215 |
| Ubr3 | 7370.02494 | 0.264435965 | 0.006680215 |
| Azin1 | 5070.47214 | 0.208208025 | 0.006684134 |
| Mbd5 | 3276.39246 | 0.231139237 | 0.006705205 |
| Acvr1b | 1855.8027 | 0.257647615 | 0.006709589 |
| Tnpo3 | 2554.86927 | 0.187135378 | 0.006720698 |
| Vsnl1 | 9289.29846 | 0.387237009 | 0.006795307 |
| Pou3f2 | 2458.03739 | -0.33228037 | 0.006798208 |
| Plekhf2 | 1403.07268 | -0.392042544 | 0.00680482 |
| Armcx2 | 3370.06019 | 0.309406239 | 0.00680482 |
| LOC100909 | 655.700124 | 0.353751081 | 0.006814015 |
| Syt1 | 53488.614 | 0.389657427 | 0.006817579 |
| RGD13595( | 131.195811 | -0.410269456 | 0.006841098 |
| Tex14 | 27.7076925 | -1.711876216 | 0.006851859 |
| Myo1d | 413.287251 | -0.72601711 | 0.006851859 |
| Wrn | 887.592738 | -0.371944328 | 0.006893334 |
| Nlrp3 | 19.5113774 | -0.836049302 | 0.006913889 |
| Bbc3 | 91.5729879 | -0.541200255 | 0.006913889 |
| Prrc1 | 957.513394 | -0.268397865 | 0.006936394 |
| Nsmf | 8317.4803 | 0.503246727 | 0.006938605 |
| Tns3 | 13915.1939 | -0.339899288 | 0.006954365 |
| Hipk2 | 25416.4762 | -0.269159768 | 0.006954365 |
| Ssr4 | 624.560727 | -0.221102653 | 0.006954365 |
| Mapk3 | 3531.59323 | -0.185687937 | 0.006954365 |
| Nav3 | 3853.47711 | 0.430940723 | 0.006954365 |
| Rassf2 | 5042.70143 | -0.379099337 | 0.006960133 |

| | | | |
|---|---|---|---|
| Coro2a | 839.307702 | 0.412981188 | 0.006960133 |
| Baz1b | 6767.08887 | 0.147457702 | 0.007005701 |
| Slc7a8 | 2178.64247 | 0.437410214 | 0.007009202 |
| Magi2 | 5765.08718 | 0.235122379 | 0.007009575 |
| Nus1 | 2062.18469 | 0.251988179 | 0.007009575 |
| Sytl5 | 526.32436 | 0.3387386 | 0.007024854 |
| Knstrn | 8.89632733 | -1.352149137 | 0.007027956 |
| Clstn3 | 15142.3146 | 0.290787455 | 0.007047683 |
| Nat1 | 121.092726 | -0.518898099 | 0.007049041 |
| Acap3 | 2092.22375 | 0.423541638 | 0.007049041 |
| Sept5 | 11267.4517 | 0.503740643 | 0.007085277 |
| Src | 6565.12518 | -0.226774773 | 0.007085572 |
| Ncald | 9247.82139 | 0.51055327 | 0.007085572 |
| Gpr108 | 864.445797 | -0.3019784 | 0.007105401 |
| Phf24 | 11146.9636 | 0.357598986 | 0.00713094 |
| Atg13 | 2332.55958 | 0.277887398 | 0.007144653 |
| Itgb1 | 8473.45885 | -0.35337601 | 0.0071742 |
| Smcr8 | 1934.01211 | 0.190102608 | 0.00719385 |
| Peli3 | 1202.88966 | 0.264858913 | 0.00719385 |
| Dcaf6 | 2699.19328 | 0.347250661 | 0.00719401 |
| Epb41l2 | 8783.30442 | -0.393409986 | 0.007235529 |
| Arpc1a | 2809.35423 | 0.226062576 | 0.007235529 |
| Msc | 4.54092985 | -1.544417194 | 0.007239419 |
| Pradc1 | 250.723326 | 0.416282595 | 0.007252851 |
| Nectin2 | 698.9484 | -0.317127623 | 0.007280282 |
| Rbp4 | 579.045579 | 0.41200081 | 0.007290447 |
| Cldnd1 | 2870.68524 | -0.171224937 | 0.007329458 |
| Mnt | 1937.9606 | 0.301758974 | 0.007339416 |
| Kcnn4 | 68.6770893 | -0.607683437 | 0.007340707 |
| Hsd17b4 | 4234.21261 | -0.329934939 | 0.007340707 |
| Fam110b | 1986.57605 | 0.324639131 | 0.007340707 |
| Hif3a | 121.879731 | 0.698739081 | 0.007342182 |
| Galnt6 | 183.955436 | -0.717287787 | 0.007349192 |
| Faap100 | 349.099401 | -0.29562832 | 0.007349192 |
| Tgm2 | 515.424595 | -0.96067863 | 0.007359207 |
| Klhdc7a | 2905.36771 | -0.421414489 | 0.007370895 |

| | | | |
|---|---|---|---|
| Fam29a | 440.921985 | -0.297187079 | 0.00738164 |
| Rapgef1 | 4211.2238 | 0.193227563 | 0.007394154 |
| Kcnq3 | 3168.01196 | 0.364490057 | 0.00741493 |
| Ppp1r3c | 7597.60266 | -0.329784581 | 0.007431884 |
| Arid1b | 6798.9737 | 0.165602051 | 0.007439363 |
| Mapk8ip2 | 11680.2169 | 0.339612761 | 0.007439363 |
| Napa | 5009.8114 | 0.219695824 | 0.007482929 |
| Wasf2 | 958.362087 | -0.383382429 | 0.007484691 |
| Iffo1 | 1557.22737 | 0.23137898 | 0.007484691 |
| Mapk8ip1 | 12919.1535 | 0.26112336 | 0.007487814 |
| Pds5b | 8078.70585 | 0.251644913 | 0.00754805 |
| Kcnv1 | 985.411509 | 0.566234219 | 0.00754805 |
| Gjc2 | 274.332714 | -0.855170593 | 0.00756869 |
| Neil2 | 64.6834934 | -0.517972281 | 0.007571562 |
| Cyp46a1 | 6701.2575 | 0.316217167 | 0.007572917 |
| Rcan1 | 2221.64043 | -0.253670082 | 0.007576365 |
| Vasn | 1037.85575 | 0.327857596 | 0.007577535 |
| Atl3 | 677.484875 | -0.318299122 | 0.00757781 |
| Spen | 5562.19614 | 0.222946004 | 0.007643584 |
| Axl | 3993.66403 | -0.356745387 | 0.00770443 |
| Miga1 | 3904.92457 | 0.221056952 | 0.007711348 |
| Dusp2 | 68.2391315 | 0.667999657 | 0.007719454 |
| Clcc1 | 2333.57733 | -0.289416391 | 0.007725454 |
| Cyld | 2811.79379 | 0.264490162 | 0.007757435 |
| Syn1 | 21254.492 | 0.403665061 | 0.007757435 |
| Ccl2 | 87.2385799 | -1.079369507 | 0.007777577 |
| Begain | 2458.25498 | 0.342287776 | 0.007813292 |
| Ralgapa1 | 3560.13272 | 0.224829687 | 0.007823884 |
| Prx | 60.10276 | -0.578834939 | 0.007825845 |
| Dock5 | 413.833758 | -0.500017574 | 0.007825845 |
| Akap6 | 12889.0588 | 0.399000043 | 0.007832969 |
| Atp6ap2 | 3541.24692 | 0.307273416 | 0.007862115 |
| Slitrk3 | 2228.75598 | 0.303386919 | 0.007865555 |
| Lrpprc | 4954.12514 | 0.167764279 | 0.007893282 |
| Tcf7l2 | 833.731165 | -0.372053309 | 0.007894393 |
| Xkrx | 14.7322135 | 0.784638313 | 0.007909064 |

| | | | |
|---|---|---|---|
| Dennd4c | 2568.59793 | -0.341171769 | 0.007991648 |
| Mapre3 | 5984.36181 | 0.286757756 | 0.007998805 |
| Car8 | 1643.95808 | -0.39253964 | 0.008018622 |
| Ppil4 | 1411.78868 | 0.202913803 | 0.008020914 |
| LOC680121 | 2547.1948 | 0.279961023 | 0.008029274 |
| Mipol1 | 418.593907 | -0.371152822 | 0.008045209 |
| Gpr180 | 1325.00501 | 0.221359502 | 0.008060663 |
| Tmem132d | 734.350307 | 0.555710142 | 0.008061333 |
| Tlr1 | 7.38063374 | -1.554836597 | 0.008063113 |
| Rad51b | 29.1599072 | -0.669188535 | 0.008063113 |
| Znfx1 | 4514.60495 | -0.584369011 | 0.008063113 |
| Tbc1d9 | 5135.68298 | 0.37568476 | 0.008063113 |
| Col9a3 | 265.803872 | -0.38047403 | 0.008075996 |
| LOC681355 | 374.345366 | -0.6033045 | 0.008087113 |
| Hmgb1 | 1695.85926 | -0.24939977 | 0.008087113 |
| Prkag2 | 1504.66471 | 0.352266815 | 0.008087113 |
| LOC108350 | 715.124349 | 0.356683744 | 0.008087113 |
| Stk11ip | 1071.23444 | 0.219503734 | 0.008095509 |
| Mrpl52 | 597.32584 | -0.234226189 | 0.008114496 |
| LOC100912 | 124.530974 | 0.515228352 | 0.008121853 |
| Gjc3 | 87.8279713 | -0.706796144 | 0.008164925 |
| Ywhae | 40295.4864 | -0.148130295 | 0.008168153 |
| Kank2 | 1828.18331 | -0.313780772 | 0.008191711 |
| Flii | 5829.14572 | -0.187845988 | 0.008227242 |
| Pten | 2547.5965 | 0.256237869 | 0.008227242 |
| Ppp1r1b | 15534.4888 | 0.415888578 | 0.008227242 |
| Amph | 1137.23571 | 0.309450053 | 0.008233357 |
| RGD131174 | 323.670505 | -0.295912657 | 0.008235663 |
| Pcnx1 | 9437.67394 | 0.219426269 | 0.008235663 |
| Tspan31 | 1301.83095 | -0.268741501 | 0.008249522 |
| Dusp4 | 234.1546 | 0.5294221 | 0.008249522 |
| Hmgb2 | 394.777729 | -0.42706873 | 0.008277926 |
| Zfand5 | 4808.88427 | 0.248469835 | 0.008277926 |
| Acss1 | 6926.20901 | -0.381859589 | 0.008285162 |
| Dusp15 | 936.607495 | -0.35941068 | 0.008285162 |
| Syvn1 | 2143.07129 | 0.183907271 | 0.008285162 |

| | | | |
|---|---|---|---|
| Mapk8ip3 | 21146.9991 | 0.284172339 | 0.008300092 |
| Tmem147 | 767.938518 | -0.238888153 | 0.008307328 |
| Cntnap2 | 8672.47568 | 0.385221031 | 0.008321256 |
| LOC102553 | 170.057689 | -0.50787548 | 0.008327591 |
| Gprin3 | 1061.26531 | 0.401921635 | 0.008327591 |
| Inhba | 394.032297 | 0.480566867 | 0.008327591 |
| Tyrobp | 12.0333288 | -1.032775458 | 0.008335209 |
| Stbd1 | 123.393759 | -0.614801191 | 0.008348523 |
| Scg5 | 9128.57741 | 0.328172544 | 0.008348523 |
| Rbm24 | 206.310281 | 0.451973133 | 0.008350329 |
| Cyba | 149.964262 | -0.568434886 | 0.0083933 |
| Inpp5j | 1739.95716 | 0.428337041 | 0.008406751 |
| Msi2 | 12947.0645 | -0.295300289 | 0.008468665 |
| Cmtr1 | 5021.61349 | -0.249880044 | 0.008476128 |
| Parl | 475.376643 | -0.229546901 | 0.008476128 |
| Sub1 | 8600.81494 | 0.141763943 | 0.008476128 |
| Gpr52 | 262.72499 | 0.443451301 | 0.008476128 |
| Arse | 53.3533579 | -0.54422916 | 0.008506523 |
| Nceh1 | 3540.67325 | 0.206723582 | 0.008506523 |
| Trim66 | 1973.59635 | 0.323914967 | 0.008528498 |
| Mtmr7 | 1936.22505 | 0.355433718 | 0.008552049 |
| Pex6 | 979.485361 | 0.262607708 | 0.008566122 |
| Plpp7 | 467.499675 | 0.334407816 | 0.008575198 |
| Tenm2 | 4582.49599 | 0.374985434 | 0.008605189 |
| Epg5 | 3583.67109 | 0.217926803 | 0.008605604 |
| Atp6v1e1 | 7246.22008 | 0.248413952 | 0.008623425 |
| Rab27b | 1962.03764 | 0.283991145 | 0.008625167 |
| Mbp | 13529.9816 | -0.655526827 | 0.008644622 |
| Mxra8 | 1026.34339 | -0.399632419 | 0.008644622 |
| Ahsa2 | 335.825871 | 0.451575512 | 0.008644622 |
| Pycard | 11.2056895 | -1.462850247 | 0.008648171 |
| Galnt16 | 4912.2771 | 0.405981043 | 0.008648171 |
| Prr7 | 555.063252 | 0.360198414 | 0.008654695 |
| St8sia5 | 2450.50468 | 0.513134519 | 0.008654695 |
| RGD130501 | 328.011926 | 0.333604736 | 0.008662577 |
| Spred2 | 1381.67738 | 0.324539916 | 0.008662712 |

| | | | |
|---|---|---|---|
| Mcm3ap | 3237.67478 | 0.222438904 | 0.008692676 |
| Rb1cc1 | 4954.43822 | 0.168468235 | 0.008724038 |
| Fancl | 213.544199 | -0.41428861 | 0.008739213 |
| RGD155989 | 22568.1341 | -0.390110879 | 0.008739213 |
| RGD130999 | 2116.12996 | -0.412283565 | 0.008748503 |
| Cd81 | 25910.0291 | -0.346581 | 0.008751225 |
| Rab11fip2 | 880.687277 | 0.259893226 | 0.00875132 |
| Pde4dip | 14335.4179 | 0.223689118 | 0.008762328 |
| Plce1 | 1277.19085 | -0.504385081 | 0.008785056 |
| Zfp445 | 3072.48237 | 0.142270344 | 0.00882684 |
| Coch | 321.263184 | 0.398823318 | 0.00884207 |
| Srgn | 6.39518785 | -0.992125732 | 0.008871675 |
| Ola1 | 2635.48275 | 0.247321343 | 0.008890639 |
| Celsr3 | 2390.65119 | 0.257288295 | 0.008890783 |
| Slc6a11 | 11243.2062 | -0.393907616 | 0.0089533 |
| Impa1 | 2278.05834 | 0.197392646 | 0.0089533 |
| Plxna4 | 8645.3721 | 0.348727637 | 0.0089533 |
| Cdc25a | 339.138085 | -0.337746708 | 0.009008052 |
| Pde6d | 1251.15925 | -0.233071709 | 0.009008052 |
| Snrnp25 | 387.668341 | -0.305342145 | 0.009010772 |
| Pld6 | 100.933065 | -0.50218951 | 0.009016547 |
| Syn2 | 10429.2323 | 0.425886015 | 0.009018304 |
| Heatr5a | 919.753454 | -0.363397917 | 0.009033262 |
| Lrig1 | 4424.5871 | -0.319178803 | 0.009049633 |
| Atg2a | 3918.02427 | 0.155453309 | 0.009049633 |
| Rims2 | 3208.44754 | 0.300523994 | 0.009049633 |
| Dagla | 5242.70897 | 0.304290454 | 0.009049633 |
| Pcmt1 | 3383.92159 | 0.248761614 | 0.009067122 |
| Brinp1 | 2916.71063 | 0.404921341 | 0.009067122 |
| Rims1 | 8368.24141 | 0.452345626 | 0.009114502 |
| Kifc1 | 27.537633 | -0.655014279 | 0.009114635 |
| Tspyl4 | 10125.6442 | 0.255949135 | 0.009114635 |
| Psmf1 | 1802.94132 | -0.464542983 | 0.009129373 |
| Nipal4 | 63.1344243 | -0.784263652 | 0.009137382 |
| Slc5a11 | 66.6939952 | -0.575791362 | 0.009137382 |
| Wipi1 | 1601.62871 | -0.315733941 | 0.009137382 |

| | | | |
|---|---|---|---|
| Notch1 | 13453.4807 | -0.379377579 | 0.009138969 |
| Cltc | 48253.0554 | 0.1763481 | 0.009174984 |
| Atg2b | 6886.22753 | 0.208101649 | 0.009174984 |
| Fam109a | 643.042187 | 0.233755044 | 0.009199066 |
| Stk40 | 1354.59193 | -0.42156179 | 0.009213157 |
| Tmed10 | 3505.48851 | -0.237222387 | 0.009213157 |
| Hsd17b12 | 4509.73412 | -0.290634325 | 0.0092217 |
| Rnf13 | 564.44524 | -0.295807091 | 0.009251011 |
| Pnmal2 | 22761.6112 | 0.291677816 | 0.009264585 |
| Zfp551 | 426.544674 | 0.348565027 | 0.009270446 |
| Gdpd2 | 560.233797 | -0.510270978 | 0.009272485 |
| Lyplal1 | 415.596494 | -0.260396241 | 0.009272485 |
| Ltn1 | 2496.64188 | 0.176800308 | 0.009275366 |
| Fus | 3510.15699 | -0.237778516 | 0.00931774 |
| LOC102547 | 28.1905078 | -0.816938992 | 0.009333501 |
| Cyp51 | 12731.5926 | -0.350208151 | 0.009333501 |
| Ikbkap | 3614.84483 | 0.18116139 | 0.009357831 |
| Prdm10 | 582.10104 | 0.342301618 | 0.009366492 |
| RGD156309 | 3227.2762 | -0.69901719 | 0.009367337 |
| Slc25a1 | 2519.46248 | -0.331701736 | 0.009368737 |
| Arfip2 | 1837.18837 | 0.280889055 | 0.009383407 |
| P2ry6 | 17.1524179 | -0.844343384 | 0.009415682 |
| Ap1b1 | 6071.24764 | 0.189840836 | 0.009497635 |
| Stx17 | 899.056294 | -0.247847963 | 0.009532564 |
| Il17rb | 13.4873804 | -1.018538667 | 0.009544772 |
| Nedd1 | 177.09977 | -0.479793277 | 0.009549766 |
| Vax1 | 126.322314 | -0.530806598 | 0.009590714 |
| Gatsl3 | 158.26237 | -0.50676729 | 0.009590714 |
| Zfp597 | 813.742481 | 0.26057787 | 0.00961388 |
| Chd3 | 17394.1449 | 0.359626303 | 0.009642206 |
| Hist1h3a | 39.1732999 | -0.682104271 | 0.009650735 |
| Notch2 | 7507.72415 | -0.336729589 | 0.009650735 |
| Rnf6 | 3274.68212 | 0.19799021 | 0.009653612 |
| Lmo3 | 2425.47058 | 0.481966833 | 0.009658469 |
| LOC100910 | 398.883616 | 0.549703724 | 0.009661165 |
| Slfn2 | 20.0548499 | -1.290688033 | 0.009673485 |

| | | | |
|---|---|---|---|
| Nup210 | 1187.46588 | 0.290873247 | 0.009673548 |
| LOC499469 | 13.8167513 | -0.749603781 | 0.009682348 |
| LOC100910 | 6657.86914 | 0.453979087 | 0.009690676 |
| Slc22a17 | 25102.8452 | 0.232797697 | 0.009711609 |
| Ahsa1 | 3734.7016 | 0.242220809 | 0.009739031 |
| Chsy3 | 277.122575 | 0.498210165 | 0.009752622 |
| Frmpd1 | 2980.14505 | -0.394823866 | 0.009758445 |
| Prr14l | 3812.51826 | 0.213152428 | 0.009763443 |
| Rnf144a | 2837.6261 | -0.338324821 | 0.009783758 |
| Il7r | 9.93214737 | -0.815793706 | 0.00980288 |
| Arid5a | 932.847059 | -0.369323558 | 0.009818541 |
| Usf3 | 3268.01516 | 0.306858808 | 0.009872107 |
| LOC103690 | 552.067908 | 0.333100849 | 0.009872107 |
| Dlk2 | 351.64922 | 0.382926107 | 0.009901076 |
| LOC102551 | 5.83668976 | -1.227611455 | 0.00993407 |
| Me3 | 1005.18325 | 0.416515039 | 0.009939894 |
| Wnt10b | 23.9895415 | 0.724190025 | 0.009995937 |
| Atp1a3 | 113775.028 | 0.35771344 | 0.010014262 |
| Fez2 | 1188.77243 | -0.324833924 | 0.010037274 |
| Mfsd11 | 830.149915 | 0.22069341 | 0.010037274 |
| Prrt3 | 2899.32418 | 0.482903958 | 0.01004816 |
| Atp6v1a | 23361.7192 | 0.26587292 | 0.010048646 |
| Arf1 | 11678.1988 | 0.210823909 | 0.010054917 |
| B3gat1 | 6806.14556 | 0.389183308 | 0.010058009 |
| LOC690871 | 417.814788 | -0.267666649 | 0.010121286 |
| Dap | 326.344292 | -0.428430721 | 0.010140008 |
| Ddx60 | 39.7456003 | -1.301335447 | 0.010161696 |
| Lamtor4 | 833.327994 | -0.289666356 | 0.010161696 |
| Ehd2 | 527.166269 | -0.496670647 | 0.010192131 |
| Plcd4 | 2921.03977 | -0.387527178 | 0.010192131 |
| Mesdc2 | 4180.52417 | -0.208341603 | 0.010192131 |
| Nde1 | 423.641752 | -0.339931352 | 0.010196166 |
| Kctd15 | 989.197575 | -0.316673248 | 0.010199103 |
| Cpne4 | 2233.08576 | 0.482905248 | 0.010242541 |
| Spg21 | 1231.0818 | -0.32568054 | 0.010250016 |
| Ccdc88c | 2000.92213 | -0.314395311 | 0.010257011 |

| | | | |
|---|---|---|---|
| Mapkapk2 | 949.355704 | -0.311811928 | 0.01029063 |
| Clgn | 167.953511 | 0.452157845 | 0.01029063 |
| Tlr7 | 12.5446452 | -0.838816083 | 0.010307479 |
| Smpdl3b | 266.236014 | 0.413208211 | 0.010309703 |
| Ppib | 2145.51102 | -0.182068911 | 0.010329743 |
| Vps13d | 11039.4105 | 0.186239316 | 0.010402739 |
| Cyp2u1 | 728.349103 | -0.279300565 | 0.010435494 |
| Fgfr1 | 11017.637 | -0.254967379 | 0.010435494 |
| Adam22 | 8638.08659 | 0.357503921 | 0.010453338 |
| Atp6v1b2 | 17812.1983 | 0.345716273 | 0.010458214 |
| RGD156405 | 1392.80933 | 0.447285798 | 0.010499627 |
| Prrc2b | 37572.4726 | 0.208602245 | 0.01054541 |
| Myh14 | 6287.3279 | -0.306032508 | 0.010551715 |
| Nrxn3 | 13820.5232 | 0.267769208 | 0.010551715 |
| Pdzrn4 | 171.35541 | -0.392057291 | 0.010621104 |
| Grk2 | 4342.12757 | 0.276807254 | 0.010621104 |
| ST7 | 598.618862 | 0.277693476 | 0.01063751 |
| Zfp644 | 3577.87759 | 0.17231749 | 0.010647846 |
| Chordc1 | 1732.11311 | 0.28600399 | 0.010647846 |
| Zfp608 | 1899.27095 | -0.306968059 | 0.010652193 |
| LOC108353 | 506.08793 | 0.379714584 | 0.010652193 |
| Gnpda1 | 562.402391 | 0.259365688 | 0.010663062 |
| Limd1 | 914.266548 | -0.324059836 | 0.010672912 |
| Wfikkn1 | 20.6894324 | 0.64707327 | 0.010684555 |
| Tet3 | 2356.45508 | 0.24017064 | 0.010694233 |
| LOC108351 | 32.7057143 | 0.580904297 | 0.010713612 |
| Lima1 | 1200.34966 | -0.384845988 | 0.010722152 |
| Cttnbp2nl | 1575.00583 | -0.353001083 | 0.010722152 |
| Chmp5 | 3474.38492 | -0.222119851 | 0.010722152 |
| Ttll12 | 932.318948 | -0.162179949 | 0.010722152 |
| Atp2b3 | 6875.39932 | 0.334880609 | 0.010722152 |
| Hepacam | 15070.2297 | -0.317233023 | 0.010734195 |
| Dag1 | 15977.0383 | -0.260109785 | 0.010734195 |
| Pdlim5 | 5497.20166 | -0.312841462 | 0.01076658 |
| Fmnl3 | 182.693955 | -0.462844939 | 0.010787882 |
| Mkks | 1243.37926 | 0.214878159 | 0.010799172 |

| | | | |
|---|---|---|---|
| Krt71 | 170.411519 | 0.498652257 | 0.010803534 |
| Ppm1e | 4605.45742 | 0.439529467 | 0.010827857 |
| Abcc3 | 68.0262417 | -0.635287556 | 0.010846232 |
| Cntn4 | 1825.1496 | 0.389996089 | 0.010889218 |
| Fras1 | 1052.99897 | 0.314555736 | 0.010908281 |
| S100a6 | 79.9889664 | -0.572943173 | 0.010936981 |
| Nedd9 | 1416.23308 | -0.335239543 | 0.010992538 |
| Fgl2 | 404.886555 | -0.471952529 | 0.010997815 |
| Rprml | 471.249365 | 0.499907091 | 0.010997815 |
| Nme6 | 141.087968 | -0.370188064 | 0.010998575 |
| Stx16 | 2305.32163 | 0.155806203 | 0.010998575 |
| Sash3 | 6.16864063 | -0.921450544 | 0.011019407 |
| Ddx49 | 769.747285 | -0.251880515 | 0.011019407 |
| Tub | 4734.21564 | 0.34411135 | 0.011019407 |
| Ccdc88a | 9243.33448 | -0.214224792 | 0.011037008 |
| Panx1 | 698.759059 | 0.394225548 | 0.011087188 |
| LOC691153 | 1096.27621 | 0.528998434 | 0.011118231 |
| Lims1 | 1784.21603 | -0.342992075 | 0.011124102 |
| Ick | 662.64262 | -0.247039069 | 0.011201435 |
| Foxm1 | 138.191332 | -0.40378534 | 0.011255892 |
| Mef2c | 6383.43807 | 0.534816952 | 0.011255892 |
| Kcnh7 | 2806.96504 | 0.541622834 | 0.011255892 |
| Zfp652 | 1480.94288 | -0.230798596 | 0.011331469 |
| Ezh2 | 86.7092667 | -0.450005467 | 0.011336175 |
| Actr1b | 8932.9793 | 0.200282184 | 0.011356777 |
| Gria4 | 3256.34591 | 0.262900331 | 0.011357517 |
| Cdh19 | 52.0588374 | -0.655806107 | 0.011366868 |
| RGD156193 | 2482.08119 | 0.35581346 | 0.011366868 |
| Cercam | 151.939206 | -0.397758124 | 0.011382674 |
| LOC108351 | 13.2301142 | 0.607596348 | 0.011391836 |
| Ppp1r9b | 27366.2428 | 0.303923893 | 0.01139623 |
| Rps6kl1 | 836.88715 | 0.291087325 | 0.011478959 |
| Stum | 1380.37755 | 0.505469352 | 0.011491783 |
| Luc7l | 3323.16461 | 0.142461396 | 0.011498626 |
| Thra | 12093.2405 | 0.284673573 | 0.011530975 |
| Rtn2 | 1178.26169 | 0.338494615 | 0.011534801 |

| | | | |
|---|---|---|---|
| Gpr20 | 7.09418153 | -1.256508092 | 0.011544221 |
| RGD130614 | 6475.88546 | 0.189953593 | 0.011617473 |
| Cacna1d | 1995.03843 | 0.313989632 | 0.011630763 |
| Akt2 | 3557.13321 | -0.295668731 | 0.01164779 |
| Senp5 | 66.1061078 | 0.469130955 | 0.011715295 |
| Wdr91 | 2285.8084 | -0.273166063 | 0.011718885 |
| Capg | 113.62627 | -0.716081944 | 0.011725416 |
| Mybl1 | 352.985761 | -0.443003897 | 0.011792875 |
| Col5a2 | 1431.65183 | -0.391054365 | 0.011792875 |
| Slc44a2 | 2759.89455 | -0.381702726 | 0.011792875 |
| Xrcc6 | 1233.38847 | -0.265713034 | 0.011792875 |
| Ptbp1 | 2715.4761 | -0.367156399 | 0.011841756 |
| Ube2w | 1460.19707 | 0.274948259 | 0.011857104 |
| Usp3 | 305.140033 | -0.31412276 | 0.011857859 |
| Cdh4 | 7167.46744 | -0.270055178 | 0.011873958 |
| Ccs | 1161.46136 | -0.239985807 | 0.011893309 |
| Mat2b | 2798.17907 | 0.372341887 | 0.011936443 |
| Tmem41a | 536.045344 | 0.390999515 | 0.011941663 |
| Atg3 | 2133.10111 | -0.20611993 | 0.011952285 |
| Sbf1 | 10346.3445 | 0.226216334 | 0.011952285 |
| Gorasp2 | 508.907844 | 0.301476633 | 0.011952285 |
| Trip12 | 11025.0818 | 0.167360784 | 0.011964452 |
| Pdp2 | 687.159762 | -0.315902816 | 0.011973195 |
| Scai | 1759.45716 | 0.243529762 | 0.012012834 |
| Pclo | 15357.5746 | 0.458817139 | 0.012012834 |
| Pcnx4 | 2900.18238 | 0.220869312 | 0.012015406 |
| Kcnn1 | 413.555164 | 0.485351062 | 0.012015406 |
| Slc12a9 | 791.644514 | -0.27647609 | 0.012092725 |
| Rufy1 | 1123.68556 | -0.23582524 | 0.012093919 |
| LOC100362 | 29.9971698 | 0.701125444 | 0.012093919 |
| Cdk14 | 4831.71715 | 0.244112993 | 0.012100195 |
| Atp2b4 | 22086.2056 | 0.451033914 | 0.012108044 |
| Focad | 1966.50364 | 0.256882535 | 0.012159465 |
| Tmc6 | 256.141109 | -0.4330729 | 0.012197967 |
| Fam217b | 1556.58798 | 0.378592839 | 0.012197967 |
| Sez6 | 22950.0985 | 0.263207259 | 0.012209553 |

| | | | |
|---|---|---|---|
| Osgin1 | 120.131543 | -0.453428735 | 0.012234846 |
| Pkm | 43717.821 | 0.246076143 | 0.012311801 |
| Iqgap1 | 574.776866 | -0.382024958 | 0.012315736 |
| Trove2 | 3299.23855 | -0.245427748 | 0.012315736 |
| Zbtb16 | 2230.52833 | 0.465057594 | 0.012315736 |
| Carmil1 | 5113.10593 | -0.334658886 | 0.012319135 |
| Ppme1 | 5243.97633 | 0.309078801 | 0.012336915 |
| Soga3 | 8583.79791 | 0.217468618 | 0.012353411 |
| Stxbp3 | 1613.22458 | -0.298423091 | 0.012357871 |
| Npc1 | 2991.31898 | -0.265659428 | 0.012382033 |
| Sf3a1 | 4486.6548 | 0.141984797 | 0.01239262 |
| Fam179b | 3240.24915 | 0.219925394 | 0.012392632 |
| Ugcg | 2755.88419 | 0.322134071 | 0.012399645 |
| March8 | 2209.46286 | -0.320276054 | 0.012424172 |
| Hdac7 | 725.250064 | -0.361890893 | 0.012427159 |
| Pikfyve | 6099.84875 | 0.167927135 | 0.012434102 |
| Bicdl1 | 989.036233 | 0.33237881 | 0.012450814 |
| Vwf | 64.4819305 | -0.594125326 | 0.012450996 |
| Epha7 | 1725.54806 | 0.46776118 | 0.012450996 |
| Zp2 | 3.87115923 | -1.09761079 | 0.012522741 |
| Kif1c | 5943.90052 | -0.3084924 | 0.012596113 |
| R3hdm2 | 7553.34163 | 0.306676504 | 0.012596113 |
| Ldlrap1 | 158.609967 | -0.699454102 | 0.0126175 |
| Sec11c | 1098.28865 | -0.381607527 | 0.012619225 |
| Zfp518b | 1084.38074 | -0.196394568 | 0.012652852 |
| Nfe2l2 | 3193.06114 | -0.323364881 | 0.012732055 |
| Lhpp | 3978.12275 | -0.267349723 | 0.012735106 |
| Nomo1 | 7372.73656 | 0.270436412 | 0.012735106 |
| LOC103692 | 3.91983825 | -1.196302752 | 0.012743929 |
| LOC100912 | 16.4070579 | -0.672326027 | 0.012746236 |
| Hk3 | 5.96620046 | -1.061085163 | 0.012748332 |
| Adamts6 | 126.973347 | -0.451193337 | 0.012768006 |
| Mmel1 | 278.249344 | -0.435018961 | 0.012768006 |
| S1pr2 | 306.930358 | -0.421207142 | 0.012768006 |
| Maob | 10745.1179 | -0.375615358 | 0.012768006 |
| Mbnl1 | 2461.85494 | 0.192696692 | 0.012768006 |

| | | | |
|---|---|---|---|
| Trappc13 | 1599.52581 | 0.240805334 | 0.012768006 |
| Ube2ql1 | 4973.28493 | 0.447686692 | 0.012768006 |
| LOC108352 | 8.75798092 | -0.805436315 | 0.012787007 |
| Cat | 4410.88595 | -0.238255771 | 0.012874815 |
| Mtmr11 | 326.818207 | -0.330863177 | 0.012876391 |
| Zc3h12b | 796.549138 | 0.263917794 | 0.01289748 |
| Rftn2 | 4134.64029 | -0.332987052 | 0.012902989 |
| Sav1 | 634.145506 | -0.332672481 | 0.012902989 |
| Antxr2 | 255.947337 | -0.469220034 | 0.012909637 |
| Dlat | 3559.65693 | 0.20747555 | 0.012909637 |
| Phrf1 | 3299.07448 | 0.17643349 | 0.012993371 |
| Sh3d19 | 710.326084 | -0.324753803 | 0.012994097 |
| Prmt8 | 801.782697 | 0.44629619 | 0.012994097 |
| Snca | 3446.77638 | 0.43686025 | 0.013057985 |
| Sgtb | 6266.71387 | 0.308986799 | 0.013064473 |
| Il10rb | 151.407727 | -0.530138744 | 0.013077321 |
| Itfg1 | 9856.35568 | 0.168685025 | 0.013097108 |
| Serpina3n | 32.9088569 | -0.875728783 | 0.01316046 |
| Ttc3 | 22681.8151 | 0.294193712 | 0.013269586 |
| Fam81a | 2214.11039 | 0.456203408 | 0.013269586 |
| Efhd1 | 4159.57176 | -0.35683939 | 0.01327024 |
| Cln8 | 476.627061 | -0.267878941 | 0.01327024 |
| Syndig1 | 698.811594 | 0.363406946 | 0.013304364 |
| Erbb2 | 259.498506 | -0.470361724 | 0.013315614 |
| Cdkn1c | 79.3316181 | -0.541895357 | 0.01332276 |
| Wdfy3 | 26705.5799 | 0.196261674 | 0.013507796 |
| Ano2 | 57.7822159 | 1.029284523 | 0.013515624 |
| Prkar2a | 4272.73417 | 0.182687012 | 0.013531368 |
| Slc9a5 | 491.596099 | 0.303708334 | 0.013531368 |
| Ptcd1 | 813.933589 | -0.253810362 | 0.013563141 |
| Mdm2 | 1208.51906 | -0.274922226 | 0.013590878 |
| Arhgap45 | 25.4171711 | -0.712665022 | 0.01364385 |
| RGD130967 | 3039.30904 | -0.259796873 | 0.013668826 |
| Opcml | 9058.73026 | 0.462853698 | 0.013668826 |
| Slco2b1 | 49.2229477 | -0.567650053 | 0.013681129 |
| Il6st | 8955.61697 | -0.269515865 | 0.013681129 |

| | | | |
|---|---|---|---|
| Otud1 | 934.265702 | 0.408990131 | 0.013681129 |
| Dhx57 | 1584.34624 | 0.232862183 | 0.013689571 |
| Rnf157 | 5228.47152 | 0.334087681 | 0.013689571 |
| Hs1bp3 | 760.155971 | -0.278494587 | 0.013689971 |
| Slc35d3 | 778.981429 | 0.346440172 | 0.013689971 |
| Lrrc8a | 11991.8203 | -0.29496557 | 0.013706709 |
| Enox1 | 691.558841 | 0.389219747 | 0.013775707 |
| Rora | 4453.59425 | -0.282902691 | 0.013812463 |
| Alkbh3 | 1023.65215 | -0.265853053 | 0.013892806 |
| Atcay | 7012.52582 | 0.345042396 | 0.013892806 |
| Atp6v1g2 | 3685.65782 | 0.308404915 | 0.013923142 |
| Tbc1d30 | 1039.51578 | 0.313933224 | 0.013936812 |
| Aqp9 | 245.449 | -0.413790796 | 0.013950831 |
| Rasal3 | 8.39227415 | -0.995781064 | 0.013955884 |
| Gripap1 | 5653.43163 | 0.254714365 | 0.013956092 |
| Slc22a5 | 503.530292 | -0.359234171 | 0.013977225 |
| Cdc20 | 44.8936692 | -0.527610052 | 0.013979465 |
| Syt16 | 890.990841 | 0.379031873 | 0.013990393 |
| Hint3 | 452.831624 | -0.263134535 | 0.014020248 |
| Doc2g | 433.383655 | 0.476281281 | 0.01406604 |
| LOC100911 | 19.9538032 | -1.001328793 | 0.014124035 |
| Cib1 | 607.738999 | -0.362013363 | 0.014150469 |
| Rundc3a | 5087.89815 | 0.313960306 | 0.014154364 |
| Tspo | 137.108862 | -0.707925767 | 0.014213176 |
| Slc32a1 | 8896.57016 | 0.314632346 | 0.014213176 |
| Tspan5 | 2945.91146 | 0.365615111 | 0.014239521 |
| Clip1 | 2853.16297 | 0.299718104 | 0.014277371 |
| Snrpc | 661.185228 | -0.292190857 | 0.014294167 |
| Kpnb1 | 7660.93115 | -0.1410362 | 0.014294167 |
| Rassf1 | 328.523598 | -0.361591048 | 0.014307457 |
| Acbd5 | 3864.21969 | -0.248605702 | 0.014333038 |
| Wdr37 | 2110.38893 | 0.278698852 | 0.014333038 |
| LOC102553 | 50.1157156 | 0.528303532 | 0.014363124 |
| Ptprn2 | 15040.5188 | 0.279538173 | 0.014370778 |
| Clstn1 | 43776.4733 | 0.294557369 | 0.014391891 |
| Mthfd1 | 3243.47271 | -0.244410226 | 0.014393814 |

| | | | |
|---|---|---|---|
| Il12rb1 | 31.1999794 | -0.81578021 | 0.014414781 |
| LOC102555 | 58.1123033 | -0.488301578 | 0.014414781 |
| Tnxa-ps1 | 159.426961 | 0.523972828 | 0.014485177 |
| Gtpbp10 | 375.816579 | -0.32196108 | 0.014498487 |
| Gls | 6810.90319 | 0.369767445 | 0.014498487 |
| Sbno2 | 832.342492 | -0.510507684 | 0.014512433 |
| Ginm1 | 732.921711 | -0.335446475 | 0.014529775 |
| Jazf1 | 1667.66338 | 0.211996025 | 0.014538691 |
| LOC501110 | 1075.42081 | -0.363198341 | 0.014583189 |
| Gpcpd1 | 996.215489 | 0.24589211 | 0.01458866 |
| Tax1bp3 | 411.193148 | -0.409350071 | 0.014611083 |
| Ksr2 | 2836.5831 | 0.332034618 | 0.014613679 |
| Lars | 3713.7982 | -0.190185255 | 0.014631941 |
| Epsti1 | 14.275913 | -0.91344508 | 0.014733271 |
| Scn8a | 9691.63311 | 0.428987502 | 0.014739211 |
| Hist1h1c | 2810.2513 | -0.314499762 | 0.014772955 |
| Fcgr3a | 4.32850807 | -1.089051979 | 0.014822535 |
| Plppr5 | 2194.27046 | 0.267845187 | 0.014846804 |
| Tp53bp1 | 4978.11811 | 0.293474595 | 0.014860275 |
| Hivep2 | 11331.5881 | 0.324569093 | 0.014862489 |
| Klf9 | 5217.70668 | 0.30695321 | 0.014864727 |
| Ank3 | 13846.3038 | 0.311715183 | 0.014867057 |
| Dbndd1 | 490.422103 | 0.30944878 | 0.014879127 |
| Nptxr | 15092.6553 | 0.517595376 | 0.014879127 |
| Sdc4 | 9546.18708 | -0.33032645 | 0.014894089 |
| Adgra3 | 1473.2405 | -0.26466906 | 0.014932241 |
| Fbxw8 | 2271.45101 | -0.22164869 | 0.014958743 |
| Kcnq2 | 13398.0658 | 0.357125251 | 0.014969306 |
| Zfp426 | 546.889016 | -0.274666362 | 0.014969641 |
| LOC100909 | 298.421483 | 0.285149306 | 0.014969641 |
| Btd | 808.950969 | -0.322423769 | 0.015018686 |
| Kcnb1 | 3545.75249 | 0.363900136 | 0.015074321 |
| Atpaf1 | 1285.9779 | 0.297409697 | 0.01509728 |
| Kansl1l | 637.452364 | -0.284136198 | 0.015136788 |
| Rarres1 | 64.4096282 | -0.513017309 | 0.015168047 |
| Safb | 5212.28369 | 0.205109378 | 0.015179626 |

| | | | |
|---|---|---|---|
| Sdhb | 4372.65331 | -0.1181699 | 0.015210983 |
| Wnt2 | 455.422106 | 0.526431822 | 0.015210983 |
| Myd88 | 471.682852 | -0.363575908 | 0.015224499 |
| Rap1b | 2568.19604 | -0.258612969 | 0.015224499 |
| Slc39a8 | 19.6448038 | -0.577381188 | 0.015244777 |
| Asrgl1 | 9751.03089 | -0.299246178 | 0.015244777 |
| Thoc5 | 1580.61823 | 0.224935834 | 0.015251297 |
| Scd2 | 388621.482 | -0.316776846 | 0.015283779 |
| Ykt6 | 2413.65329 | -0.23624157 | 0.015283779 |
| Ckmt1b | 6292.73714 | 0.302967573 | 0.01529493 |
| LOC102553 | 8.44384952 | -0.876738423 | 0.015306502 |
| Slfn5 | 46.8342609 | -0.78659415 | 0.015306502 |
| Tmem159 | 56.105933 | -0.541525472 | 0.015306502 |
| Mpst | 957.054823 | -0.354489763 | 0.015307796 |
| Rap1gap2 | 6919.60768 | 0.433716363 | 0.015307796 |
| Ap2b1 | 6018.03664 | 0.267591755 | 0.015320845 |
| RGD156568 | 368.086022 | -0.341661383 | 0.015354258 |
| Ddhd1 | 2631.16638 | -0.154419641 | 0.015364208 |
| Cnppd1 | 798.475066 | -0.254862942 | 0.015367175 |
| Nptx2 | 1524.49266 | 0.60143563 | 0.015374612 |
| Ryr3 | 1865.21279 | 0.285542185 | 0.015387782 |
| Cenpe | 118.965672 | -0.405042124 | 0.015391082 |
| Dclk3 | 1369.93795 | 0.34168838 | 0.0154117 |
| Srrm3 | 1561.9481 | 0.329364006 | 0.015415729 |
| Rpap3 | 1427.81923 | 0.201354255 | 0.015456184 |
| Zfyve26 | 1512.52862 | -0.211331871 | 0.015483721 |
| Fam155a | 2501.90535 | 0.370437147 | 0.015483721 |
| Npc2 | 1748.69389 | -0.279934023 | 0.015486279 |
| Hs3st2 | 1880.10678 | 0.514244485 | 0.015576376 |
| R3hdm4 | 2782.2867 | 0.295770184 | 0.015613862 |
| Epb41l4b | 1257.36398 | 0.327177438 | 0.015613862 |
| Edc4 | 2933.38577 | 0.177734671 | 0.015618561 |
| Ephb4 | 486.671882 | -0.377656756 | 0.015651238 |
| Tpm4 | 175.142673 | -0.50142537 | 0.015657799 |
| Gpr22 | 883.120481 | 0.45520649 | 0.01569718 |
| Scarb2 | 17419.0856 | -0.270348755 | 0.015715311 |

| | | | |
|---|---|---|---|
| Elovl1 | 507.582476 | -0.464740818 | 0.015769983 |
| Znf408 | 267.126557 | -0.3724432 | 0.015769983 |
| Comt | 1404.74582 | -0.226606384 | 0.015778011 |
| Zfp341 | 545.694145 | 0.317074417 | 0.015781186 |
| Usp25 | 1739.85992 | -0.328614685 | 0.015784327 |
| Tomm20 | 2837.40873 | 0.276498489 | 0.015823236 |
| Zbtb11 | 2204.67748 | 0.275215513 | 0.015862582 |
| Canx | 34777.2058 | -0.150169923 | 0.015872571 |
| Whsc1l1 | 5445.34729 | 0.151330384 | 0.015929207 |
| Rnf115 | 2668.90385 | -0.254969843 | 0.015939425 |
| Pdcl3 | 771.228707 | -0.223331524 | 0.015939425 |
| Cherp | 3351.21554 | 0.129185557 | 0.015939425 |
| Gstk1 | 718.565478 | -0.329619356 | 0.015939443 |
| Tspan6 | 927.409865 | -0.325926784 | 0.016026248 |
| LOC100151 | 783.973105 | -0.292724033 | 0.01606099 |
| Pwwp2b | 495.817209 | 0.2482379 | 0.016086392 |
| Prkcd | 678.930456 | 0.479954147 | 0.016086392 |
| Nat8l | 8929.31417 | 0.354371853 | 0.016105338 |
| Gna15 | 8.36460275 | -0.914060992 | 0.016112597 |
| Ldlrad3 | 180.201372 | -0.421604348 | 0.016141946 |
| Cd2bp2 | 4825.93807 | -0.13674461 | 0.016141946 |
| Lrp11 | 5523.39974 | 0.256826641 | 0.016141946 |
| Smim1 | 715.3476 | -0.352456647 | 0.016197425 |
| Dynll2 | 15129.2559 | 0.209071206 | 0.016197425 |
| Egln2 | 2227.98868 | 0.225939798 | 0.016201171 |
| Large1 | 6743.58478 | 0.335511736 | 0.016201171 |
| Pde3b | 397.235727 | -0.304468252 | 0.016201629 |
| Smc4 | 701.589573 | -0.299972894 | 0.016270684 |
| Paqr9 | 631.029985 | 0.368241854 | 0.016270684 |
| Kcnj6 | 3674.89572 | 0.5127559 | 0.016299041 |
| Ribc1 | 246.879092 | -0.498149754 | 0.016302036 |
| Grik5 | 6631.79672 | 0.27472331 | 0.016311109 |
| Pik3ip1 | 1800.56933 | -0.289643692 | 0.016313163 |
| Gpd1l | 1474.35093 | 0.241439902 | 0.01637376 |
| Ccdc189 | 172.76529 | -0.473009737 | 0.016408113 |
| Slc38a3 | 7943.85437 | -0.34959891 | 0.016451451 |

| | | | |
|---|---|---|---|
| RGD130813 | 525.145022 | -0.2371724 | 0.016495078 |
| Olfm1 | 15704.5393 | 0.417732202 | 0.016511533 |
| Zdbf2 | 5815.12663 | 0.446264823 | 0.016511533 |
| LOC102552 | 87.6340011 | 0.416981997 | 0.016526335 |
| Pum2 | 9210.57737 | 0.160692559 | 0.016541705 |
| Cds1 | 886.991708 | 0.375045612 | 0.016576956 |
| LOC100911 | 4731.07474 | -0.348537389 | 0.016594941 |
| Kcnk4 | 226.063196 | 0.489006367 | 0.016720176 |
| Cobll1 | 240.701327 | -0.487312423 | 0.016723055 |
| Col9a2 | 76.1060553 | -0.495033873 | 0.016723128 |
| Prkd3 | 2390.0652 | -0.30196607 | 0.016723128 |
| Sepp1 | 24816.0671 | -0.277148562 | 0.016724278 |
| Tcaf1 | 6747.8583 | 0.230207515 | 0.01674098 |
| Gga1 | 3133.96638 | 0.201236994 | 0.016822963 |
| Helb | 622.385249 | -0.249219096 | 0.016851338 |
| Hps1 | 436.77327 | -0.372820711 | 0.016886027 |
| Cers2 | 963.382512 | -0.346098389 | 0.016887162 |
| Lrrc10b | 357.429656 | 0.308463514 | 0.016927157 |
| Hid1 | 5131.17431 | 0.222574826 | 0.016941472 |
| Zfp318 | 1750.70654 | 0.228128791 | 0.01694148 |
| Prkab1 | 449.113089 | -0.292760323 | 0.016945205 |
| Gch1 | 13.9716618 | -0.797243222 | 0.016948745 |
| Gbas | 4282.28387 | -0.155042796 | 0.016978821 |
| Gcn1l1 | 6602.68331 | 0.158770315 | 0.016978821 |
| Rps6ka1 | 352.582043 | -0.35054664 | 0.017000949 |
| Plcg2 | 71.9313901 | -0.481697738 | 0.017064263 |
| Rasl10a | 184.210723 | 0.520767679 | 0.017066383 |
| Gsn | 4236.84785 | -0.509799897 | 0.01710248 |
| Gpr3 | 206.023369 | 0.510692934 | 0.017110798 |
| LOC100910 | 52.9376036 | -0.57136894 | 0.017116056 |
| RGD131095 | 692.206128 | -0.318305981 | 0.017116056 |
| Ide | 3289.01879 | -0.240532008 | 0.017116056 |
| Med14 | 2428.30166 | 0.205488011 | 0.017116056 |
| RGD130723 | 2317.67345 | 0.246483529 | 0.017116056 |
| Prkar1b | 6663.94223 | 0.326273763 | 0.017116056 |
| Kcnh4 | 213.037768 | 0.370124936 | 0.017116056 |

| | | | |
|---|---|---|---|
| Nit2 | 1034.00399 | -0.21166506 | 0.017129754 |
| Plk4 | 39.2400331 | -0.491655504 | 0.017201811 |
| Qdpr | 2277.20267 | -0.437989363 | 0.017201811 |
| Mn1 | 3585.49033 | 0.216587621 | 0.017259239 |
| LOC303140 | 64.8257743 | -0.451917071 | 0.017298992 |
| Map3k10 | 3902.32124 | 0.349255678 | 0.017306811 |
| Fbxl18 | 205.793981 | 0.342672036 | 0.017386514 |
| Lypla2 | 1333.63538 | 0.247455525 | 0.017424286 |
| Mtss1l | 45628.382 | -0.27346867 | 0.0175131 |
| Hdac1 | 1256.62137 | -0.230181293 | 0.017774786 |
| Clint1 | 3715.99963 | -0.127156973 | 0.017774786 |
| Rnf126 | 1537.47532 | 0.278291816 | 0.017774786 |
| Vstm4 | 321.706928 | -0.410781016 | 0.017823096 |
| Dhx8 | 1853.44178 | 0.161327543 | 0.017898376 |
| Prnp | 30658.3356 | -0.170413254 | 0.018026197 |
| Ercc6 | 1314.70636 | 0.165045335 | 0.01813108 |
| Chst14 | 79.3233933 | -0.48506956 | 0.018181577 |
| Sin3b | 2244.20905 | 0.202511334 | 0.018186865 |
| Prdx4 | 669.108438 | -0.268166973 | 0.018247342 |
| Tbc1d16 | 1092.73367 | 0.26034922 | 0.01828773 |
| Seli | 1426.77781 | 0.193667543 | 0.018378114 |
| Irf6 | 112.577429 | 0.425114821 | 0.018387119 |
| Lactb2 | 780.004521 | -0.197942215 | 0.018448832 |
| Cntnap3b | 530.510476 | 0.299937699 | 0.018452896 |
| Mllt6 | 5634.91327 | 0.192636707 | 0.018481915 |
| Slc25a27 | 729.353622 | 0.217153489 | 0.018481915 |
| Kars | 3322.1924 | -0.319664886 | 0.018488718 |
| Cep290 | 1152.0922 | 0.282560971 | 0.018488718 |
| LOC100910 | 21.6463371 | -0.544853343 | 0.018516235 |
| Smarcd1 | 3729.849 | 0.200766304 | 0.018581439 |
| Prpf8 | 17332.152 | 0.144372831 | 0.018604782 |
| Crtc2 | 688.101157 | -0.240642493 | 0.0186164 |
| Mpnd | 1313.82366 | -0.245161213 | 0.018646269 |
| Itgav | 7867.14903 | -0.261539513 | 0.01865957 |
| Irf2bpl | 3100.14704 | 0.221668438 | 0.018666197 |
| Mllt11 | 4390.84468 | 0.348202995 | 0.018666197 |

| | | | |
|---|---|---|---|
| Bin2 | 12.6764955 | -0.626808514 | 0.018672241 |
| Rnf169 | 962.901848 | -0.231164642 | 0.018685526 |
| Tprn | 465.52959 | -0.437587257 | 0.018711185 |
| Swap70 | 2833.79805 | -0.319823893 | 0.018847807 |
| Chp1 | 5093.19139 | 0.193313755 | 0.018920497 |
| Rtn4rl2 | 686.854725 | 0.601542973 | 0.01895655 |
| Itih3 | 30467.2861 | -0.425769133 | 0.018958523 |
| Nrip3 | 3774.04258 | 0.417755415 | 0.018958523 |
| Ap1ar | 986.452216 | 0.242251881 | 0.01896204 |
| Diablo | 823.886184 | -0.248247896 | 0.019023032 |
| Ddx50 | 1386.64965 | 0.184365396 | 0.019052829 |
| Ophn1 | 1826.98179 | 0.209662612 | 0.019052829 |
| Lilrb3 | 6.70568657 | -0.905254926 | 0.019082883 |
| Mx2 | 2844.07438 | -0.804511575 | 0.019088628 |
| Rasa3 | 3315.83102 | -0.214024615 | 0.019141909 |
| Fgd3 | 257.205282 | -0.444755761 | 0.019145268 |
| Brpf3 | 2005.24494 | 0.262852719 | 0.019153524 |
| Scp2 | 10757.9665 | -0.244572386 | 0.019205889 |
| Man2a2 | 8957.9329 | 0.194911989 | 0.019214511 |
| Rab6b | 26415.4268 | 0.330579013 | 0.019214511 |
| Sult4a1 | 8921.68958 | 0.33737314 | 0.019214511 |
| Cmtm6 | 394.556232 | -0.433746713 | 0.019237012 |
| Megf8 | 15457.0518 | 0.233044548 | 0.01924366 |
| Pianp | 4760.7075 | 0.312916682 | 0.019257358 |
| Cyth2 | 2444.69565 | 0.266810907 | 0.019260408 |
| Arhgap4 | 38.7825185 | -0.489253888 | 0.01929032 |
| Sphkap | 4318.68801 | 0.279016648 | 0.019324322 |
| Slc44a3 | 59.6482674 | -0.484755086 | 0.019373437 |
| Ttc12 | 433.734347 | -0.441289731 | 0.019392867 |
| Rbm12b | 56.3703443 | 0.450723411 | 0.019392867 |
| Nek9 | 3559.21016 | -0.25255762 | 0.019490881 |
| Lrp8 | 3847.24779 | 0.192445693 | 0.019495267 |
| Klf10 | 516.867728 | 0.453429522 | 0.019495267 |
| Tle1 | 952.919229 | 0.249604831 | 0.019582548 |
| Stac2 | 1409.72424 | 0.410660135 | 0.019678031 |
| Rmi2 | 6.25347594 | -0.729229849 | 0.019720243 |

| | | | |
|---|---|---|---|
| Kcnd2 | 4071.62212 | 0.198434206 | 0.019772576 |
| Erich5 | 87.1778198 | 0.493370319 | 0.019776752 |
| Mmp17 | 1210.21054 | 0.455606778 | 0.01978463 |
| Ap2m1 | 15921.092 | 0.24236018 | 0.019822331 |
| Cpeb1 | 710.469269 | 0.253237485 | 0.019910516 |
| Irgq | 6638.55819 | 0.224276122 | 0.019912054 |
| Filip1 | 894.249852 | 0.36609079 | 0.019912054 |
| Col12a1 | 77.1119302 | -0.450835371 | 0.019936385 |
| Tspan11 | 174.291646 | -0.502207318 | 0.019952701 |
| Snrpb2 | 505.053671 | -0.280179984 | 0.020048785 |
| Clca1 | 19.0651857 | -0.816097051 | 0.02005689 |
| Hcls1 | 39.1042156 | -0.508933228 | 0.02005689 |
| Coro1a | 1066.66729 | 0.418803729 | 0.02005689 |
| Tfeb | 600.124615 | -0.299865307 | 0.020091216 |
| Kif7 | 1011.21657 | -0.300956988 | 0.020094093 |
| Ndel1 | 2814.69502 | 0.21874875 | 0.020126587 |
| Cxcr4 | 136.25929 | -0.496746948 | 0.02017391 |
| Slc25a18 | 4249.58627 | -0.422225643 | 0.02017391 |
| LOC100909 | 5.75943452 | -0.780214816 | 0.020175475 |
| Supt6h | 12193.274 | 0.165148855 | 0.020177536 |
| Tbc1d4 | 347.742784 | 0.338989599 | 0.020239393 |
| Napb | 12835.5798 | 0.374918973 | 0.020290423 |
| Asap3 | 304.798182 | -0.459489103 | 0.020400948 |
| Grk1 | 4.01685737 | 0.894325661 | 0.020400948 |
| Gabarap | 4090.97313 | -0.17492624 | 0.020416949 |
| Nfatc1 | 1053.8374 | -0.350653462 | 0.020484078 |
| Rnf123 | 3462.94148 | 0.190135365 | 0.020493809 |
| Otud7b | 2326.32421 | -0.300370449 | 0.02054188 |
| Ctsc | 46.5083854 | -0.582611601 | 0.020593243 |
| Sox13 | 860.430694 | -0.36161887 | 0.020684834 |
| Eftud2 | 3408.29903 | 0.165891516 | 0.020709896 |
| Selt | 4235.78786 | -0.134521158 | 0.020745313 |
| Calm2 | 5880.64426 | 0.250106654 | 0.020745313 |
| Csad | 2124.78674 | -0.275385008 | 0.020805882 |
| Tpp2 | 1218.52756 | 0.229904906 | 0.020805882 |
| A3galt2 | 58.9231498 | -0.524874491 | 0.020846597 |

| | | | |
|---|---|---|---|
| Ncan | 29085.7098 | -0.334214646 | 0.0209183 |
| Dpysl3 | 2265.58969 | -0.243090772 | 0.020954485 |
| Manf | 1716.32988 | 0.259017524 | 0.020954485 |
| Mgp | 10.4379901 | -0.802123499 | 0.020962781 |
| Pfas | 2663.00543 | -0.228723859 | 0.020962781 |
| Rtn1 | 28444.529 | 0.249159754 | 0.020962781 |
| Neu2 | 148.50761 | 0.421749337 | 0.020987744 |
| Abcd3 | 3653.81801 | -0.236564734 | 0.021019715 |
| Wdr83os | 965.866991 | -0.173507419 | 0.021019715 |
| Dusp5 | 95.7039256 | 0.44703631 | 0.021019715 |
| Bag6 | 7836.03508 | 0.158538941 | 0.021046215 |
| Hnrnpd | 3472.01866 | -0.183630444 | 0.021048598 |
| R3hdm1 | 5978.45144 | 0.29666793 | 0.021053342 |
| Srxn1 | 1674.75361 | 0.334082018 | 0.021095181 |
| Tmem181 | 1072.33522 | 0.266632605 | 0.021113433 |
| Cd84 | 17.2137744 | -0.58493897 | 0.021114985 |
| Gspt2 | 1103.15485 | 0.308585104 | 0.021148642 |
| Prrc2c | 9129.50306 | 0.174551867 | 0.021187565 |
| Lingo1 | 6826.99358 | 0.469192353 | 0.021192507 |
| Sipa1 | 164.18744 | -0.459980424 | 0.021193105 |
| Mlc1 | 49782.7383 | -0.293003188 | 0.02123828 |
| Sod2 | 7811.73759 | -0.244136281 | 0.02123828 |
| Zfp217 | 209.328511 | -0.446205622 | 0.021259036 |
| Ninj2 | 33.394119 | -0.606328873 | 0.021263781 |
| Spcs1 | 1717.79254 | -0.219804154 | 0.021264564 |
| Gtf2a1 | 2560.83406 | -0.215425744 | 0.021274558 |
| Napg | 5261.89366 | 0.274523075 | 0.021291662 |
| Tgfb3 | 140.892888 | 0.424274228 | 0.021330349 |
| Rab7a | 13161.5901 | -0.157378674 | 0.021416202 |
| Dopey1 | 1964.79327 | 0.252140886 | 0.021416202 |
| Cntrob | 143.370568 | -0.380264067 | 0.021461162 |
| Mtmr6 | 6053.98864 | 0.135661714 | 0.021461162 |
| RGD130573 | 6808.0365 | 0.303538846 | 0.021499205 |
| Gng4 | 2956.43742 | 0.293575283 | 0.021501365 |
| Fhdc1 | 328.560017 | -0.421615445 | 0.021507138 |
| Tspan13 | 2599.88117 | 0.258299818 | 0.021525401 |

| | | | |
|---|---|---|---|
| Gla | 641.349185 | 0.196449963 | 0.021537801 |
| Chm | 681.661845 | 0.261569975 | 0.021626224 |
| Arid4a | 1848.47636 | 0.239101537 | 0.021632899 |
| Nmnat2 | 2079.8434 | 0.296544663 | 0.021632899 |
| Aldh9a1 | 2807.56004 | -0.284637967 | 0.02166663 |
| Svep1 | 78.6083936 | -0.485483099 | 0.021714031 |
| Usp6nl | 1971.48452 | -0.232974466 | 0.021714031 |
| Prickle1 | 2646.79297 | 0.42669306 | 0.021714031 |
| Clcnkb | 14.5604515 | 0.51606685 | 0.021761755 |
| Dnm1l | 7164.53705 | 0.167546844 | 0.021810803 |
| Rgs17 | 2767.35711 | 0.273555752 | 0.021854297 |
| Pop1 | 249.352861 | -0.323264795 | 0.021893815 |
| Pisd | 1097.25066 | -0.210814551 | 0.021893815 |
| Golm1 | 660.825191 | -0.311227151 | 0.021898282 |
| Iqsec1 | 11474.2579 | 0.248181901 | 0.021903309 |
| Il4i1 | 16.3771576 | -0.519448588 | 0.021903983 |
| LOC102548 | 1010.22375 | 0.296245258 | 0.021903983 |
| Mapk12 | 350.766713 | -0.32859213 | 0.021905251 |
| Cep295nl | 95.5870904 | -0.449761265 | 0.021909812 |
| Crhr2 | 63.4793585 | 0.525286855 | 0.022048119 |
| Grn | 1507.99356 | -0.371864918 | 0.022061625 |
| Xkr4 | 2043.3482 | 0.356944314 | 0.022078623 |
| Lrp10 | 3464.76142 | -0.311279275 | 0.022084206 |
| Nlrp1a | 17.9867543 | -0.694672681 | 0.022094436 |
| Tpx2 | 162.71452 | -0.376912675 | 0.022140107 |
| Gnl1 | 2846.9065 | 0.222467931 | 0.022140107 |
| Zfp804a | 752.965224 | 0.429676388 | 0.022140107 |
| Sstr3 | 236.318774 | 0.481161884 | 0.022194612 |
| Ggt7 | 3296.07763 | 0.325875189 | 0.02223942 |
| Car14 | 40.3448694 | -0.532142028 | 0.022341177 |
| Hibch | 971.291853 | -0.260874838 | 0.022369944 |
| Scamp5 | 7139.66006 | 0.29008616 | 0.022370378 |
| Pacsin1 | 6238.06169 | 0.292391997 | 0.022380014 |
| Apold1 | 32.7945558 | 0.54839715 | 0.022384692 |
| Mark1 | 3768.028 | 0.25660274 | 0.022444653 |
| Znf660 | 189.720913 | -0.330548913 | 0.022465577 |

| | | | |
|---|---|---|---|
| Zfp641 | 174.8154 | -0.339532345 | 0.022504628 |
| Slc15a3 | 15.8925644 | -0.810545155 | 0.022506656 |
| Nr3c2 | 2172.16591 | 0.190777369 | 0.022506656 |
| Phospho2 | 615.572333 | 0.229959083 | 0.022506656 |
| Gnai3 | 2939.10485 | -0.275637223 | 0.022575417 |
| Slc7a2 | 3535.52993 | -0.354094613 | 0.022632678 |
| Uchl1 | 21140.7295 | 0.296042832 | 0.022660866 |
| Parm1 | 712.371575 | -0.333019988 | 0.022667169 |
| Ctif | 577.484322 | 0.226366184 | 0.022690992 |
| Acbd3 | 2211.71958 | -0.190386131 | 0.022725243 |
| Strbp | 8383.72267 | 0.302895386 | 0.022749616 |
| Troap | 4.18925557 | -0.825076983 | 0.02277893 |
| Zfp496 | 817.620172 | -0.259028149 | 0.022801779 |
| Zfp41 | 437.162417 | -0.296606936 | 0.022802798 |
| Adra1d | 107.467508 | 0.510193433 | 0.022911853 |
| Igsf8 | 4553.39864 | 0.180950399 | 0.022944644 |
| Fam134a | 6512.67899 | 0.184364029 | 0.022974306 |
| Camk2n1 | 1985.37101 | 0.33547981 | 0.022987962 |
| LOC108351 | 8.15475942 | -0.603809026 | 0.023026373 |
| Cry2 | 6207.71738 | 0.25363509 | 0.023026373 |
| Dock1 | 5228.18775 | -0.265723872 | 0.02306781 |
| Smarce1 | 2918.08347 | -0.160127885 | 0.023075584 |
| Asphd2 | 1517.43332 | 0.205996972 | 0.023121886 |
| Gadd45a | 538.105596 | -0.300179419 | 0.023129378 |
| Fam160a2 | 1884.96183 | 0.270779439 | 0.023224326 |
| Numbl | 2669.16088 | 0.345381198 | 0.023234337 |
| Tbl1xr1 | 4819.25682 | 0.143621273 | 0.023271623 |
| Zfp335 | 1229.94842 | 0.253344181 | 0.023286286 |
| Sel1l | 10626.3761 | 0.154350562 | 0.023303176 |
| Tmem178a | 1623.11928 | 0.464635221 | 0.023351978 |
| Ing2 | 521.278192 | 0.301882285 | 0.023456964 |
| Tnr | 4736.05279 | 0.303735634 | 0.023461493 |
| Dync1h1 | 37819.4698 | 0.233400484 | 0.023492122 |
| Zbtb40 | 624.254244 | 0.205249468 | 0.023494436 |
| Rassf10 | 39.4802038 | -0.535749838 | 0.023498555 |
| Hip1r | 2979.18805 | -0.232026481 | 0.023510841 |

| | | | |
|---|---|---|---|
| RGD156039 | 2255.11326 | 0.23898064 | 0.023516344 |
| Prps1 | 3549.30894 | 0.233435576 | 0.023591294 |
| Hras | 1528.35766 | 0.282028337 | 0.023669047 |
| Bmper | 564.604759 | 0.462290426 | 0.023685572 |
| Lat2 | 6.21783676 | -0.784655823 | 0.023694817 |
| Zfp36l1 | 2989.79867 | -0.342296115 | 0.023698187 |
| Ptn | 6933.8122 | -0.314506571 | 0.023698187 |
| Itga9 | 197.585214 | -0.413616318 | 0.023701864 |
| Atf7 | 1236.90836 | -0.253190808 | 0.023701864 |
| Trim23 | 2155.50157 | 0.199251095 | 0.023701864 |
| Vwc2 | 522.828118 | 0.31781804 | 0.023701864 |
| Fam19a2 | 1151.61845 | 0.385052955 | 0.023701864 |
| Ppp1r18 | 486.103578 | -0.358536451 | 0.023720613 |
| Zer1 | 10577.5066 | 0.20424153 | 0.023725962 |
| Cry1 | 1708.64546 | 0.286921464 | 0.023750499 |
| Inafm2 | 1614.9944 | 0.254159219 | 0.023835439 |
| Dhrs13 | 377.110115 | -0.313969661 | 0.023862387 |
| Chmp1b | 1380.82258 | -0.217814757 | 0.02397836 |
| Ccdc17 | 83.5964797 | -0.375805775 | 0.023979257 |
| Sod1 | 4611.32797 | -0.126083113 | 0.023988322 |
| Zfp24 | 3026.13346 | -0.207924816 | 0.024029357 |
| Cuedc1 | 1159.01624 | -0.197427836 | 0.024029357 |
| Snap25 | 21079.3212 | 0.364535681 | 0.02409302 |
| Zfp444 | 2331.70086 | 0.143464578 | 0.024115537 |
| Ap4e1 | 399.288562 | -0.237356621 | 0.024232428 |
| Sfmbt2 | 176.447583 | -0.397361293 | 0.024232923 |
| Bcat1 | 46388.6057 | 0.258086069 | 0.024232923 |
| Agtpbp1 | 7752.38403 | 0.311696708 | 0.024269155 |
| Zbtb25 | 1024.46961 | -0.231703605 | 0.024303048 |
| Gars | 3916.82546 | 0.205089727 | 0.024307233 |
| Smpd3 | 5699.49576 | 0.239316944 | 0.024344713 |
| Eps8l1 | 8.39149955 | 0.560855063 | 0.024381778 |
| Jtb | 677.731599 | -0.263287404 | 0.024464651 |
| Cadm3 | 10822.414 | 0.288978228 | 0.024464651 |
| Slc35e2b | 3216.99972 | 0.164825151 | 0.024473316 |
| Pdxdc1 | 1124.07231 | -0.177696201 | 0.02447611 |

| | | | |
|---|---|---|---|
| Ak2 | 489.441265 | -0.318437432 | 0.024503732 |
| Sorbs3 | 3650.92866 | -0.269731887 | 0.024504136 |
| Unc5b | 3237.52042 | -0.285371357 | 0.024508609 |
| Neo1 | 6281.21728 | -0.220412484 | 0.02458842 |
| Psmc3ip | 241.22799 | -0.334723489 | 0.024618802 |
| Gramd1b | 7263.84096 | 0.219732556 | 0.024654235 |
| Nudt2 | 1031.01859 | -0.223571472 | 0.024719425 |
| Atp5g1 | 2356.61353 | 0.232291642 | 0.024761623 |
| Gramd1a | 2023.8442 | 0.237320112 | 0.02484757 |
| Exoc2 | 1555.25747 | 0.17035807 | 0.02490573 |
| Amigo1 | 6134.75518 | 0.231263398 | 0.024941236 |
| Insc | 31.028878 | -0.505800228 | 0.024951238 |
| Map4k3 | 3105.57487 | 0.222369889 | 0.025032338 |
| Atp6v0e2 | 2825.97593 | 0.215305865 | 0.025164293 |
| Ctso | 713.832918 | -0.32258474 | 0.025189852 |
| Matk | 1696.12754 | 0.302260053 | 0.025189852 |
| Mad2l2 | 249.447072 | -0.302595818 | 0.025204204 |
| Stxbp1 | 30398.6778 | 0.29986682 | 0.025310381 |
| Slc9a3r2 | 817.673755 | 0.314051486 | 0.02537829 |
| Kcnj9 | 1130.45038 | 0.421058656 | 0.02537829 |
| Fat3 | 5472.22643 | 0.269569678 | 0.025438723 |
| Usp2 | 2929.32989 | 0.319555395 | 0.025438723 |
| Spsb1 | 957.374213 | -0.287061213 | 0.025450182 |
| Kif5b | 11538.8646 | -0.162862096 | 0.025450182 |
| Stip1 | 7466.88865 | 0.212172662 | 0.025450182 |
| Ttll7 | 13466.3442 | 0.30382349 | 0.025450182 |
| Aff3 | 3851.1607 | 0.267839338 | 0.025468057 |
| Gpr162 | 3583.85844 | 0.291105497 | 0.025485469 |
| Rab3a | 12754.8778 | 0.301510467 | 0.025485469 |
| Shc3 | 3081.02363 | 0.246080185 | 0.02548684 |
| Tmod3 | 807.906208 | -0.348434188 | 0.025587153 |
| Msl3 | 502.571424 | 0.244474764 | 0.025587153 |
| Greb1 | 361.182691 | 0.440542472 | 0.025613688 |
| Dcaf1 | 1536.47755 | 0.172557713 | 0.025616908 |
| Ddah2 | 4339.26346 | -0.297957178 | 0.025623557 |
| Ice1 | 4172.56688 | 0.174791971 | 0.025623557 |

| | | | |
|---|---|---|---|
| Bet1 | 765.324616 | -0.282208946 | 0.025628737 |
| Atp1b1 | 38156.7332 | 0.31619698 | 0.025648143 |
| Rmrp | 1178.06304 | -0.602883071 | 0.025651256 |
| St3gal4 | 2047.54855 | -0.270742633 | 0.025651256 |
| Rnf112 | 2058.772 | 0.29293356 | 0.025651256 |
| Fbxo6 | 1250.29461 | -0.241561346 | 0.02572228 |
| Sla2 | 27.0945174 | 0.501302844 | 0.025799436 |
| Fam102b | 1248.95372 | 0.331458568 | 0.025807293 |
| Pja1 | 14057.9855 | 0.2174601 | 0.025844244 |
| Upp1 | 361.104892 | -0.371795606 | 0.025853334 |
| Sspn | 1899.38568 | -0.30605329 | 0.025872886 |
| Ube2k | 3597.74925 | 0.183475393 | 0.025872886 |
| Pcyox1l | 1090.07122 | 0.248352371 | 0.025872886 |
| Usp21 | 1206.53981 | 0.175244308 | 0.02587362 |
| Ift81 | 1525.82047 | -0.295051828 | 0.025933832 |
| RGD156403 | 212.97197 | -0.29030788 | 0.025933832 |
| Uggt1 | 6839.13678 | 0.15241382 | 0.025933832 |
| Bri3bp | 969.095598 | 0.249208023 | 0.025962575 |
| Nfia | 6211.33756 | -0.244634176 | 0.025970913 |
| Adora2a | 1308.13561 | 0.357927463 | 0.025973577 |
| Cmtm7 | 10.9367681 | -0.669263364 | 0.026030935 |
| Fam83f | 57.0340908 | -0.463338982 | 0.026030935 |
| Ankrd26 | 1054.64141 | 0.189650992 | 0.026034604 |
| Fhl2 | 452.797303 | 0.42594324 | 0.026057677 |
| LOC108352 | 9.22044139 | 0.547239104 | 0.026067144 |
| Cd37 | 7.70500132 | -0.647957703 | 0.026121684 |
| Anapc13 | 724.05548 | -0.256280274 | 0.026121684 |
| Pomt2 | 866.449805 | -0.207071458 | 0.026149821 |
| Usp13 | 1877.17742 | 0.245157548 | 0.026199384 |
| Hapln2 | 160.968333 | -0.509904321 | 0.026254916 |
| Apc2 | 13988.0738 | 0.19538324 | 0.026255993 |
| Sdk2 | 7963.5055 | 0.333072719 | 0.026269396 |
| Atg101 | 1071.90298 | 0.18907356 | 0.026296761 |
| LOC100911 | 7.66032763 | -0.494772088 | 0.026298336 |
| Pi4k2a | 1618.76958 | 0.218103586 | 0.02631562 |
| Pabpc4 | 2262.64199 | 0.169577613 | 0.026379017 |

| | | | |
|---|---|---|---|
| Cebpb | 477.152838 | -0.372198719 | 0.026412401 |
| Gnptab | 3397.43898 | 0.144003922 | 0.026423375 |
| Morn5 | 18.8995373 | -0.475824541 | 0.026442306 |
| Sik1 | 604.673294 | 0.33096569 | 0.026447354 |
| Ccr5 | 12.1468955 | -0.566218009 | 0.026555962 |
| Dennd5a | 22635.4681 | -0.197389287 | 0.026555962 |
| Trnp1 | 2613.29901 | 0.357138477 | 0.026582173 |
| Ccne2 | 69.3113882 | -0.391449986 | 0.026641632 |
| Folh1 | 3695.68285 | -0.295319917 | 0.026641632 |
| Snx18 | 1467.57283 | -0.21104898 | 0.026641632 |
| Sh3bp5 | 841.509205 | 0.249885939 | 0.026641632 |
| Dclk1 | 31363.1365 | 0.373608214 | 0.026641632 |
| Rrm1 | 1506.46432 | -0.16946083 | 0.02669251 |
| Ogfod3 | 275.728204 | -0.298409026 | 0.026695273 |
| Ubxn4 | 4950.7469 | -0.199303206 | 0.026695273 |
| Ndufaf4 | 505.031964 | 0.236816831 | 0.026760966 |
| Pkdcc | 172.018883 | -0.375100821 | 0.026826997 |
| Bod1l1 | 4223.95644 | 0.240753556 | 0.026895097 |
| Anxa6 | 4645.2481 | -0.287645816 | 0.026898527 |
| LOC102546 | 110.793997 | -0.40197919 | 0.026922814 |
| Mylk2 | 7.06830161 | -0.734285537 | 0.02698383 |
| Maml3 | 323.995389 | 0.309316406 | 0.026995982 |
| Prkg2 | 260.439831 | 0.330010633 | 0.027029624 |
| Xiap | 2361.90312 | -0.207636955 | 0.027164507 |
| Cnot7 | 1587.66614 | 0.131047511 | 0.027164507 |
| Ctdsp2 | 1730.13379 | -0.295544139 | 0.027197922 |
| Fmc1 | 273.940894 | -0.293204635 | 0.027197922 |
| Rhebl1 | 34.1143456 | 0.454592133 | 0.027205518 |
| Pcnt | 1883.18256 | 0.252904254 | 0.02721214 |
| Cabin1 | 3869.05641 | 0.169293336 | 0.027231477 |
| Nectin3 | 1188.04792 | -0.28082701 | 0.027363679 |
| Dynll1 | 4570.94311 | 0.250538887 | 0.027363679 |
| LOC108349 | 1423.18038 | -0.288914024 | 0.027530878 |
| Tmem63c | 1644.48955 | 0.31675341 | 0.027530878 |
| Cys1 | 208.715705 | 0.348017003 | 0.027533464 |
| Efemp2 | 276.795535 | -0.390670134 | 0.02760467 |

| | | | |
|---|---|---|---|
| Sugp1 | 1792.21581 | -0.164935601 | 0.02760467 |
| Plin4 | 1721.71102 | 0.376365615 | 0.02760467 |
| Adat2 | 86.2498263 | 0.357885284 | 0.027630429 |
| Fbxw4 | 611.106668 | 0.233313966 | 0.027631783 |
| Gss | 348.05356 | -0.29094664 | 0.027768683 |
| Smad1 | 1425.57078 | -0.159149977 | 0.027768683 |
| Rnf20 | 2588.73836 | 0.13168319 | 0.027768683 |
| Eif4enif1 | 1742.27346 | 0.21194425 | 0.027771355 |
| Specc1 | 2289.10471 | 0.260659441 | 0.02782279 |
| Arhgap26 | 5950.71053 | 0.222880628 | 0.027833867 |
| Srrm4 | 1662.55473 | 0.341072677 | 0.027833867 |
| Foxk1 | 1805.16544 | 0.169225963 | 0.027872703 |
| Scaf1 | 10902.491 | 0.177049283 | 0.028014116 |
| Gstp1 | 1523.97404 | -0.195999111 | 0.028093773 |
| Zfp703 | 3308.76919 | -0.284944512 | 0.028125366 |
| Yae1d1 | 1090.42652 | -0.233953307 | 0.028125448 |
| Meis1 | 737.533569 | -0.259328753 | 0.028187632 |
| Atxn7l3 | 3533.58003 | 0.291658958 | 0.028187632 |
| Tuft1 | 151.526527 | -0.372163871 | 0.028222005 |
| Rab8b | 575.201761 | -0.271021206 | 0.028222005 |
| Rasgef1b | 1101.36224 | 0.325389404 | 0.028222005 |
| Alcam | 7558.42259 | 0.184549244 | 0.028222177 |
| Pbx4 | 68.2389893 | 0.426069715 | 0.028222177 |
| LOC108350 | 7.45643692 | -0.526973624 | 0.028286445 |
| Lyzl4 | 55.5367854 | 0.501483039 | 0.028286445 |
| Eif5 | 12252.4091 | -0.144063721 | 0.028302683 |
| Doc2a | 765.429911 | 0.431704877 | 0.028308054 |
| Usp22 | 5436.39582 | 0.203674239 | 0.02833702 |
| Dpp6 | 13910.7326 | 0.246985106 | 0.02833702 |
| Arfip1 | 1214.24726 | -0.265838973 | 0.028341331 |
| Fbxl2 | 862.275021 | 0.284588949 | 0.028374066 |
| Usf1 | 1825.6706 | -0.244498089 | 0.028462429 |
| Prelp | 7924.67989 | -0.3077524 | 0.028477134 |
| Camta1 | 7404.97289 | 0.232139829 | 0.028477134 |
| B4galt6 | 3624.87321 | 0.308296565 | 0.028491882 |
| Cdkn1b | 2318.40396 | -0.222623133 | 0.028540172 |

| | | | |
|---|---|---|---|
| Sorcs3 | 2116.7491 | 0.397233345 | 0.028608994 |
| Rspo2 | 364.239908 | 0.483596071 | 0.028608994 |
| Wnk3 | 2281.99719 | 0.219918722 | 0.028628853 |
| LOC108351 | 15.0258981 | 0.500091215 | 0.02863053 |
| Fyttd1 | 2057.26248 | 0.237974693 | 0.028634386 |
| Renbp | 660.194749 | -0.304366532 | 0.028636544 |
| Tmem251 | 187.330373 | -0.282009428 | 0.028636892 |
| Gcsh | 3392.42765 | -0.251198062 | 0.028668109 |
| Mlip | 567.071956 | 0.421781281 | 0.028767564 |
| Zmym3 | 4016.3696 | 0.209559969 | 0.02878344 |
| Reep3 | 6288.81691 | -0.260843238 | 0.028790695 |
| Tesk1 | 1657.45862 | 0.247654038 | 0.028790695 |
| Parva | 1600.15329 | -0.281358584 | 0.028808749 |
| Glrb | 3678.10643 | 0.25919451 | 0.028808749 |
| Htr1a | 298.970125 | 0.424895787 | 0.028808749 |
| Nhej1 | 63.4084616 | -0.381998776 | 0.028815955 |
| Tab1 | 1405.1901 | -0.186392162 | 0.028831623 |
| Rab10 | 9986.1275 | -0.180792385 | 0.028837581 |
| Pts | 133.567755 | -0.292558277 | 0.02886929 |
| Pam | 12735.5756 | 0.187815046 | 0.028896374 |
| Mex3a | 201.181266 | -0.319731496 | 0.028938962 |
| Tmprss5 | 27.5366595 | -0.532852793 | 0.029036357 |
| Tomm34 | 2443.64405 | 0.240962716 | 0.029077635 |
| Ppp1r14c | 475.389303 | -0.329174146 | 0.029089943 |
| Mphosph8 | 2376.60302 | 0.226155132 | 0.029089943 |
| Mgat3 | 4929.7326 | 0.247402484 | 0.029089943 |
| Nisch | 28583.199 | 0.173186116 | 0.029098268 |
| LOC102548 | 137.538192 | 0.400494391 | 0.029098268 |
| Nfe2l1 | 19879.3947 | 0.166556504 | 0.029120979 |
| Heatr5b | 3293.52253 | 0.188519762 | 0.029135469 |
| LOC102553 | 18.8820702 | -0.471257803 | 0.029146484 |
| Lrfn2 | 816.252049 | 0.378970474 | 0.029149693 |
| Insm1 | 105.168853 | -0.351119003 | 0.029187528 |
| Slc25a22 | 92.7732462 | 0.422127064 | 0.029187528 |
| Ptpn18 | 7.83234489 | 0.580606965 | 0.029187528 |
| LOC100362 | 8.91368267 | -0.546246493 | 0.029197969 |

| | | | |
|---|---|---|---|
| Prrc2a | 29585.8798 | 0.145025973 | 0.029203275 |
| Klhl8 | 2167.01353 | 0.205041694 | 0.029203275 |
| Habp4 | 5482.54847 | 0.296094524 | 0.029203275 |
| Gtf3c6 | 467.203083 | -0.28167747 | 0.029256675 |
| Zfp692 | 156.649764 | -0.258822606 | 0.029267232 |
| Plpp6 | 1012.79485 | 0.381300169 | 0.02927386 |
| LOC103690 | 1608.33721 | 0.320210461 | 0.029316187 |
| Lonrf2 | 17770.8495 | 0.280053196 | 0.029330019 |
| Cbfb | 762.945422 | -0.248692813 | 0.029368178 |
| Impdh1 | 1308.68453 | 0.267161626 | 0.029368178 |
| Sppl2a | 1914.23521 | -0.272557168 | 0.029369682 |
| LOC102550 | 41.7076181 | -0.435904415 | 0.029382661 |
| Cyp27a1 | 106.20672 | -0.462765869 | 0.029395151 |
| Ubald2 | 1064.50547 | -0.232403328 | 0.029398959 |
| Espl1 | 27.8833439 | -0.459247296 | 0.029408322 |
| Rfc3 | 143.669293 | -0.304754583 | 0.029408322 |
| LOC691254 | 16.3636546 | 0.472469149 | 0.029412971 |
| Msl1 | 3825.63326 | 0.212684949 | 0.029476243 |
| Npdc1 | 4238.4203 | 0.276789768 | 0.029662339 |
| E2f5 | 289.408978 | -0.35680536 | 0.029698278 |
| Sbno1 | 4471.02445 | 0.236152717 | 0.029698278 |
| Mb21d1 | 3.85747858 | -0.628921519 | 0.029809255 |
| Trim7 | 310.995566 | -0.275694925 | 0.02989208 |
| Lysmd2 | 1155.00811 | 0.212134404 | 0.02989208 |
| Slc26a8 | 433.521465 | 0.299904434 | 0.029921637 |
| Adamts7 | 30.5883528 | -0.462701283 | 0.029938555 |
| LOC103694 | 7.04563535 | -0.646417791 | 0.029963662 |
| Slc35f1 | 3077.16797 | 0.301380273 | 0.029963662 |
| Tmem50a | 2346.41098 | -0.185943586 | 0.030032001 |
| Pld1 | 2398.35204 | -0.274284798 | 0.030078228 |
| Iqsec3 | 10044.281 | 0.298377769 | 0.030129636 |
| Sfrp1 | 1256.16439 | -0.435321452 | 0.030147279 |
| Atad1 | 5541.25852 | -0.207215755 | 0.030172477 |
| Arhgef7 | 5291.32797 | 0.210244565 | 0.03019166 |
| Otub2 | 202.663266 | 0.351625952 | 0.030261054 |
| LOC294154 | 5511.51096 | 0.193158031 | 0.030307119 |

| | | | |
|---|---|---|---|
| Adam23 | 9458.3778 | 0.308334786 | 0.030307119 |
| Fn3k | 1038.78327 | 0.248149284 | 0.03031464 |
| NEWGENE_ | 21.0338122 | 0.449619607 | 0.03031464 |
| Rtn3 | 36099.7843 | 0.211053185 | 0.030371045 |
| LOC102552 | 351.864109 | 0.337073391 | 0.030371045 |
| Rreb1 | 1132.05921 | -0.266094922 | 0.030391361 |
| Ktn1 | 6273.14372 | -0.252776518 | 0.030422226 |
| Gkap1 | 1495.21654 | -0.216854595 | 0.030435363 |
| Spata24 | 120.082138 | -0.372307952 | 0.030470709 |
| Ebf1 | 125.67092 | 0.40965801 | 0.030499967 |
| Enc1 | 13600.6768 | 0.418242277 | 0.030503755 |
| Pfn1 | 3825.51942 | -0.195408166 | 0.030552503 |
| Park7 | 5370.07996 | -0.126890813 | 0.030552503 |
| Lrrn2 | 7407.4567 | 0.193794871 | 0.030552503 |
| Slc2a3 | 52.1721673 | 0.435680542 | 0.030552503 |
| Sh2d1b2 | 39.2715136 | 0.517116406 | 0.030552503 |
| Tpd52 | 990.866581 | 0.225720295 | 0.030619566 |
| Lrrc20 | 315.269963 | 0.260257409 | 0.030619566 |
| Cln3 | 348.676906 | -0.239759337 | 0.030682363 |
| Adamts8 | 41.6165873 | 0.464835347 | 0.030682363 |
| Homez | 254.925192 | -0.298137473 | 0.030692904 |
| Wnt10a | 63.7648981 | 0.458423488 | 0.030735887 |
| Mtch1 | 10906.4731 | 0.183068331 | 0.030859812 |
| Ccdc9 | 546.828682 | 0.214982256 | 0.030937259 |
| Ddit4l | 67.5615407 | -0.419978992 | 0.030978719 |
| Ccdc8 | 598.592673 | -0.360866378 | 0.030978719 |
| LOC108352 | 36.6521361 | 0.438482837 | 0.03110164 |
| Plekhb2 | 16033.5104 | 0.150748192 | 0.031184428 |
| Gmeb1 | 430.466933 | 0.258266617 | 0.031184428 |
| Mfsd4 | 805.132565 | 0.388876651 | 0.031184428 |
| Rasgrp3 | 138.463694 | -0.451839294 | 0.031208449 |
| Erc1 | 5266.82086 | 0.152183109 | 0.031215239 |
| Sprn | 1847.9968 | 0.367264094 | 0.031215239 |
| Dscr3 | 806.85722 | -0.22310275 | 0.031274568 |
| Ptpn4 | 5324.72963 | 0.190386759 | 0.031317703 |
| Cdk5 | 1166.3459 | 0.282310163 | 0.031377913 |

| | | | |
|---|---|---|---|
| Yes1 | 1012.19352 | -0.234546199 | 0.03142045 |
| Igf2r | 4376.14739 | 0.134933791 | 0.03150424 |
| Ica1 | 952.065552 | 0.30775654 | 0.03153854 |
| Osbp | 3731.83935 | 0.132902966 | 0.031581506 |
| Tceal8 | 1666.92088 | -0.196481491 | 0.031620855 |
| Gsk3a | 4798.67569 | 0.218951086 | 0.031642391 |
| LOC103690 | 43.0160611 | 0.419144423 | 0.031664815 |
| Rnf135 | 148.010308 | -0.335124426 | 0.031682441 |
| Hyal2 | 483.735032 | -0.232261406 | 0.031682441 |
| Gnai2 | 18775.3358 | -0.246293889 | 0.031802316 |
| Ermp1 | 4327.73701 | -0.193959573 | 0.031802316 |
| Tagln2 | 266.423253 | -0.500507571 | 0.031809938 |
| Spata6 | 554.50479 | -0.326362138 | 0.031843618 |
| Cspg5 | 23689.8046 | -0.256672812 | 0.031843618 |
| Cul3 | 6279.08662 | 0.127913052 | 0.031843618 |
| Enthd1 | 4.92015241 | 0.65316273 | 0.031854934 |
| Rcn3 | 294.720991 | -0.371513324 | 0.031876292 |
| Mamdc2 | 58.2436751 | -0.396644235 | 0.032015555 |
| Cyp4v3 | 2308.64106 | -0.387970339 | 0.032015555 |
| Arhgef40 | 1860.81621 | -0.26378527 | 0.032066843 |
| Zfp180 | 1203.03927 | 0.324038993 | 0.032078505 |
| Mmp24 | 1268.73722 | 0.254425827 | 0.032092634 |
| Cadm4 | 8320.83107 | -0.169856529 | 0.032105089 |
| Sox18 | 46.5027373 | -0.443862954 | 0.032165881 |
| Arl13b | 504.430223 | -0.252014328 | 0.032165881 |
| Atpaf2 | 497.56121 | -0.188346595 | 0.032165881 |
| Cpeb3 | 11363.2119 | 0.235012683 | 0.032165881 |
| Uhmk1 | 4217.49477 | 0.281986195 | 0.032165881 |
| Afap1l1 | 96.9101558 | 0.381246269 | 0.032165881 |
| Sec23b | 1880.91617 | -0.291109626 | 0.032233723 |
| Cand2 | 1255.3384 | -0.247316569 | 0.032266059 |
| Fam19a4 | 89.7443091 | -0.445631483 | 0.032382622 |
| Sgsm1 | 5667.15339 | 0.322832833 | 0.032413436 |
| Wrnip1 | 1720.60341 | 0.192469965 | 0.03243276 |
| Pde4d | 3102.23795 | 0.231525154 | 0.032438605 |
| Zfp414 | 660.503431 | -0.210008695 | 0.032442543 |

| | | | |
|---|---|---|---|
| Stx2 | 474.114485 | -0.321264192 | 0.032601508 |
| Khnyn | 419.146275 | -0.31898243 | 0.032601508 |
| Ccdc141 | 3841.33631 | -0.289098613 | 0.032601508 |
| Mipep | 2363.70843 | -0.207368745 | 0.032601508 |
| Nptx1 | 12834.6291 | 0.423173455 | 0.032601508 |
| Blnk | 13.8527989 | -0.554247621 | 0.032637239 |
| Iqgap3 | 169.836017 | 0.38501037 | 0.032637239 |
| Klhl34 | 30.9176897 | 0.43866984 | 0.032637239 |
| Eci1 | 1037.00161 | -0.306263017 | 0.03272133 |
| Col4a3bp | 3782.65533 | 0.157616313 | 0.03272133 |
| Fbxo7 | 1992.03436 | -0.215356069 | 0.032755806 |
| LOC108349 | 11.7031869 | -0.624113726 | 0.032772352 |
| Timm10 | 554.176552 | 0.208386129 | 0.032795607 |
| Apba1 | 8160.31002 | 0.283730128 | 0.032795607 |
| Pafah1b1 | 24136.8936 | 0.169985999 | 0.032811801 |
| Sall1 | 2847.79809 | -0.281919912 | 0.032832727 |
| Alkbh7 | 419.082234 | -0.214795221 | 0.032832727 |
| Cpne7 | 825.83389 | 0.311773221 | 0.032841449 |
| Rps17 | 2205.15348 | -0.188612528 | 0.032874828 |
| Pycrl | 850.039791 | -0.172441204 | 0.032874828 |
| Paqr7 | 7104.63719 | -0.251712099 | 0.032897648 |
| Prex1 | 13650.6651 | -0.248764473 | 0.032897648 |
| Pxmp2 | 279.010199 | -0.282778808 | 0.033026475 |
| Dnajb4 | 2143.15498 | 0.227460457 | 0.033026475 |
| Nefl | 9506.5926 | 0.380711713 | 0.033026475 |
| Tjp1 | 11662.5055 | -0.23080106 | 0.033106056 |
| Acsbg1 | 38622.628 | -0.265885323 | 0.033132056 |
| Arhgef19 | 2299.92068 | -0.277949774 | 0.033136506 |
| Cd33 | 8.95012236 | -0.534263139 | 0.03315866 |
| LOC108353 | 6.62122927 | -0.565373603 | 0.033173004 |
| Copz1 | 2521.1684 | -0.1582236 | 0.033173004 |
| Prkacb | 19035.7591 | 0.191662074 | 0.033173004 |
| Rnh1 | 2054.03254 | -0.27232369 | 0.033177212 |
| Psmg1 | 767.077317 | -0.186231384 | 0.033188341 |
| Btrc | 4421.8781 | 0.211298474 | 0.03323117 |
| LOC108348 | 11.5104373 | -0.567427934 | 0.033248016 |

| | | | |
|---|---|---|---|
| Lrrn1 | 3205.94073 | -0.304569161 | 0.03335862 |
| Trappc12 | 2199.41337 | 0.178295984 | 0.03335862 |
| Edil3 | 5115.25949 | -0.273593422 | 0.033372224 |
| Etl4 | 10250.1886 | 0.360410331 | 0.033522785 |
| Dusp14 | 356.349135 | 0.328139304 | 0.033557327 |
| Vamp5 | 10.0878478 | -0.505377971 | 0.03360072 |
| Mea1 | 2509.59231 | -0.170650592 | 0.033642607 |
| Pdlim2 | 63.566402 | -0.425982789 | 0.033647047 |
| Angptl4 | 301.81338 | -0.41673911 | 0.033647047 |
| Ankib1 | 3612.04763 | -0.178180377 | 0.033647047 |
| Angpt1 | 259.976384 | -0.398763842 | 0.033648662 |
| Rab24 | 1452.8805 | 0.174843769 | 0.033650381 |
| Slc38a5 | 5.36211412 | 0.610021711 | 0.033732581 |
| Tbc1d5 | 3395.37471 | -0.136845468 | 0.03378086 |
| Ddx31 | 402.602487 | -0.268825429 | 0.033784126 |
| Cpped1 | 674.913947 | -0.249900538 | 0.033818168 |
| Mtg1 | 415.330434 | -0.219684413 | 0.033818168 |
| Pabpc1 | 4522.90378 | -0.233302496 | 0.033883843 |
| Xpnpep1 | 2762.88499 | -0.181322812 | 0.033883843 |
| Fam63b | 17277.1759 | -0.193294462 | 0.033897596 |
| Ube2j1 | 3063.68205 | 0.202739754 | 0.033923306 |
| Glis3 | 331.753809 | -0.302680308 | 0.034037499 |
| Ubr1 | 3391.47457 | 0.182053608 | 0.034103421 |
| Tnfaip2 | 386.738633 | -0.437789322 | 0.034112882 |
| Ints2 | 645.536334 | 0.188556994 | 0.034154113 |
| Slc30a3 | 489.611382 | 0.451260672 | 0.034169255 |
| Cacul1 | 2778.10496 | 0.130342473 | 0.034200542 |
| Krtcap2 | 390.319067 | -0.237079157 | 0.034209706 |
| Ric1 | 2531.85029 | 0.164682671 | 0.034266747 |
| Popdc2 | 109.270994 | -0.322529621 | 0.034321903 |
| Ap3d1 | 12299.1753 | 0.133614414 | 0.034398395 |
| Pfkfb2 | 1526.65121 | 0.208628724 | 0.034398395 |
| Pipox | 61.6788433 | -0.418129537 | 0.034409456 |
| Ina | 6149.56004 | 0.313613543 | 0.034419456 |
| Ptchd1 | 350.73924 | 0.360105776 | 0.034419456 |
| LOC108349 | 46.8535446 | 0.386767332 | 0.034419456 |

| | | | |
|---|---|---|---|
| Tmod4 | 53.2281019 | -0.375949911 | 0.034483527 |
| Tspan14 | 535.747047 | -0.305692802 | 0.034483527 |
| Ttl | 3123.46228 | 0.295646619 | 0.034483527 |
| Fam163b | 3288.73882 | 0.375538723 | 0.034483527 |
| Apobr | 76.8627293 | -0.381356513 | 0.034490085 |
| Slc8b1 | 464.258591 | -0.309151523 | 0.034490085 |
| Emc7 | 2824.91981 | -0.171337455 | 0.034549274 |
| Scrt1 | 1464.20653 | 0.393803878 | 0.034557044 |
| Cnot1 | 6537.01744 | 0.136612876 | 0.034591452 |
| Mapk9 | 6479.0608 | 0.239212007 | 0.034591452 |
| Cacng2 | 1596.50094 | 0.340101394 | 0.034591452 |
| Tmed3 | 470.448399 | -0.241784728 | 0.034617963 |
| Ctnnd2 | 29388.8802 | -0.212426605 | 0.034699612 |
| Fam111a | 10.8617206 | -0.501863411 | 0.034708628 |
| Mrpl43 | 1039.06074 | 0.202051965 | 0.034736209 |
| Crhbp | 640.58479 | 0.390812811 | 0.03487054 |
| Snx19 | 3563.24514 | 0.175682825 | 0.034896154 |
| RGD156127 | 554.422207 | 0.223672488 | 0.034979629 |
| Fads2 | 13991.9989 | -0.264507625 | 0.035043044 |
| Fbln2 | 4594.997 | -0.35852616 | 0.035150883 |
| Kit | 4619.86749 | 0.32356982 | 0.035162617 |
| Abhd17a | 2653.2303 | 0.218606582 | 0.035181753 |
| Epha5 | 5078.15613 | 0.279558128 | 0.035181753 |
| Bcam | 588.388851 | -0.394143146 | 0.035262014 |
| Pcp4l1 | 2290.50431 | 0.381210149 | 0.035271181 |
| Cyb561a3 | 178.129599 | -0.345400041 | 0.035300722 |
| Gm5471 | 106.550405 | -0.291805664 | 0.035334719 |
| Lsm14a | 2797.39389 | -0.21194936 | 0.035355867 |
| Znrf1 | 1059.5479 | 0.227370037 | 0.035380389 |
| Dscaml1 | 3071.79006 | 0.293190858 | 0.035470693 |
| Ankh | 4059.19545 | 0.194054741 | 0.035532241 |
| Lsp1 | 30.8482148 | -0.431799029 | 0.035630493 |
| Appl1 | 4360.96341 | 0.140617685 | 0.035646159 |
| Fra10ac1 | 437.670099 | 0.241121175 | 0.035661221 |
| Ikzf2 | 341.612622 | -0.281211516 | 0.035716769 |
| Scamp1 | 5910.71007 | 0.163552736 | 0.035716769 |

| | | | |
|---|---|---|---|
| Ttc37 | 1826.73379 | 0.207093873 | 0.035825827 |
| Zbtb20 | 15163.4824 | -0.253722327 | 0.035827042 |
| Ttc7b | 4723.0748 | 0.255733725 | 0.035853324 |
| Ppp1r1a | 1177.43363 | 0.241426059 | 0.035875924 |
| Raph1 | 2418.34614 | 0.287485009 | 0.035875924 |
| Apcdd1 | 162.515351 | -0.420761517 | 0.03589525 |
| Etfa | 1591.76815 | -0.259329883 | 0.03589525 |
| Gpx8 | 685.245375 | -0.268886245 | 0.035984126 |
| Dennd2a | 1160.24922 | -0.248207786 | 0.036003772 |
| LOC100361 | 8390.37072 | -0.177670726 | 0.036055635 |
| Ccdc107 | 1184.4957 | -0.211129755 | 0.03606564 |
| Gid8 | 1500.49732 | -0.190887534 | 0.036264856 |
| LOC108352 | 311.611092 | -0.382301039 | 0.03631026 |
| Hic2 | 152.317146 | 0.343776729 | 0.03631026 |
| Dixdc1 | 3011.35848 | 0.266236486 | 0.036344636 |
| LOC102550 | 199.522261 | -0.345062315 | 0.036418742 |
| Ring1 | 960.514507 | -0.184473622 | 0.036498255 |
| Dcxr | 527.338956 | -0.313733679 | 0.036549285 |
| Rnf39 | 126.4295 | 0.423997079 | 0.036549285 |
| Cpm | 199.280743 | -0.424082499 | 0.036564368 |
| Crebbp | 12620.6582 | 0.135913663 | 0.036564368 |
| Trpc6 | 181.999686 | 0.402480825 | 0.036564368 |
| Dact3 | 4569.92265 | 0.286158799 | 0.036576385 |
| Fam156b | 129.878671 | 0.328011517 | 0.036576385 |
| Lpp | 2629.29023 | -0.288975521 | 0.036592718 |
| Ccdc34 | 460.718635 | -0.287490896 | 0.036592718 |
| Rbm18 | 1879.04053 | 0.1502993 | 0.036600474 |
| Chrm3 | 1139.59239 | 0.385947941 | 0.036664644 |
| Purg | 4220.19914 | 0.206488581 | 0.03667161 |
| Adck4 | 856.332032 | 0.182013824 | 0.036712491 |
| Kremen1 | 403.053868 | 0.25289994 | 0.036712491 |
| Wnk4 | 117.765796 | 0.336204107 | 0.036712491 |
| Tcf7 | 49.1601119 | -0.387251506 | 0.036716161 |
| Nr4a2 | 1738.97227 | 0.426795091 | 0.03671904 |
| Pafah2 | 687.704921 | -0.229320507 | 0.036727547 |
| Cyp2j10 | 12.7182658 | -0.510349137 | 0.036779936 |

| | | | |
|---|---|---|---|
| Elk4 | 1616.91705 | -0.185570698 | 0.036779936 |
| Eif4a2 | 14201.2936 | 0.154807468 | 0.036779936 |
| Adap1 | 2118.55889 | 0.321291582 | 0.036811534 |
| Peak1 | 4688.48856 | 0.216454668 | 0.036819369 |
| Zdhhc5 | 2560.44091 | 0.227007869 | 0.036858215 |
| Sft2d1 | 420.919127 | -0.235605807 | 0.036934986 |
| Git1 | 9366.32027 | 0.273276135 | 0.036934986 |
| Plppr1 | 381.447771 | 0.326598075 | 0.036934986 |
| Dmd | 6822.53831 | -0.251156386 | 0.036943784 |
| Arf5 | 2886.76637 | 0.199465865 | 0.037013217 |
| Uap1 | 1290.58987 | 0.285601402 | 0.03702177 |
| Nlk | 2030.32106 | 0.330577104 | 0.037049494 |
| Fancg | 86.3803263 | -0.343456399 | 0.037134382 |
| Pitpnm1 | 8233.01987 | 0.206706015 | 0.037228092 |
| Pard3b | 627.592904 | -0.253226843 | 0.037296627 |
| Pex5 | 107.351886 | 0.367495814 | 0.037296627 |
| Coro7 | 1431.84177 | 0.214781823 | 0.037508392 |
| Xrcc3 | 13.5420262 | -0.426514259 | 0.037515411 |
| Tmbim6 | 9572.34434 | -0.223667938 | 0.037515411 |
| Palm2 | 1002.10133 | 0.246962582 | 0.037515411 |
| Spcs3 | 322.943576 | 0.260201701 | 0.037515411 |
| Trim9 | 4847.39825 | 0.273110545 | 0.037515411 |
| Fpgs | 204.200881 | -0.312717053 | 0.037533876 |
| Ptk7 | 768.928124 | -0.307846557 | 0.037672334 |
| Pqlc1 | 441.165318 | 0.292046856 | 0.037686285 |
| Pecr | 551.065829 | -0.23779677 | 0.037703855 |
| Hs3st3b1 | 93.0703484 | -0.420301026 | 0.037730847 |
| Usp19 | 5429.57483 | 0.146230203 | 0.037730847 |
| Il34 | 1283.46378 | 0.335051483 | 0.037741109 |
| Hmg20b | 736.864631 | -0.247392854 | 0.03776176 |
| Slc45a1 | 1898.05449 | 0.276785637 | 0.03776176 |
| Gabra5 | 1718.06131 | 0.404047559 | 0.037775274 |
| Timeless | 50.6187319 | -0.40814646 | 0.03788835 |
| RGD131058 | 744.932862 | 0.390523741 | 0.037910326 |
| Sv2b | 9911.89854 | 0.41326394 | 0.037910326 |
| Panx3 | 14.6602465 | -0.438559744 | 0.037964032 |

| | | | |
|---|---|---|---|
| Rusc1 | 5545.60606 | 0.256099297 | 0.038050624 |
| Egfr | 4425.51921 | -0.280253194 | 0.038092848 |
| LOC108352 | 477.614354 | 0.329452475 | 0.038120358 |
| rnf141 | 3365.87674 | -0.182150627 | 0.038123577 |
| Cemip | 444.646665 | 0.331414046 | 0.038163932 |
| Pdcd6 | 1378.25524 | -0.152377737 | 0.038209754 |
| Rptor | 5697.51376 | 0.134966434 | 0.038209754 |
| Lin7a | 123.268645 | 0.394009352 | 0.038209754 |
| Pgm2l1 | 12142.6062 | 0.309969006 | 0.038238138 |
| Tmem35 | 2138.85787 | 0.329141603 | 0.038317256 |
| Got2 | 8326.05047 | 0.246059301 | 0.038356506 |
| Slc2a10 | 966.273524 | -0.311041943 | 0.03838338 |
| Etfb | 2023.06399 | -0.220089065 | 0.03839362 |
| Spryd3 | 3958.22854 | 0.237636026 | 0.03839362 |
| Rdh13 | 592.216219 | 0.27463107 | 0.038399316 |
| Fstl4 | 836.122159 | 0.275375287 | 0.038399316 |
| Map7d2 | 2801.4331 | 0.275022409 | 0.03840307 |
| Pbld1 | 53.074164 | -0.42793097 | 0.038442033 |
| Pde9a | 861.661853 | -0.227130203 | 0.038442033 |
| Lama5 | 1514.21971 | -0.25051514 | 0.038471893 |
| Ctnna2 | 5725.22575 | 0.205383297 | 0.038565621 |
| Tom1l1 | 622.024378 | -0.259007773 | 0.038568557 |
| Pdzd11 | 752.871866 | -0.192486797 | 0.038568557 |
| LOC100910 | 327.789599 | 0.262149551 | 0.038575409 |
| Obsl1 | 496.391817 | -0.312649237 | 0.038596285 |
| Maml1 | 608.58927 | -0.264780955 | 0.038596285 |
| Enpp5 | 9886.90761 | 0.143603155 | 0.038596285 |
| Evc | 904.866663 | -0.257479201 | 0.038605776 |
| Tram2 | 46.5919079 | -0.415653555 | 0.038668454 |
| Slc35f6 | 2097.40792 | -0.215526746 | 0.038696304 |
| Sypl1 | 1440.41715 | -0.272168391 | 0.038707856 |
| Fnbp4 | 1005.35413 | -0.244804368 | 0.038724319 |
| Adal | 188.767379 | 0.284768775 | 0.038734071 |
| Stam2 | 1167.04241 | -0.245451251 | 0.038758076 |
| Psma7 | 3041.10315 | -0.208292312 | 0.038760885 |
| Chd1 | 1606.50444 | -0.151724828 | 0.038760885 |

| | | | |
|---|---|---|---|
| LOC100912 | 76.0795989 | 0.39557255 | 0.038766301 |
| Hbp1 | 763.709974 | -0.269946121 | 0.03881332 |
| H2afy | 3304.3102 | -0.178186196 | 0.038824817 |
| Uaca | 410.054989 | -0.357950976 | 0.038825389 |
| Prune2 | 4835.42569 | 0.243086024 | 0.038923358 |
| Cecr6 | 1183.67815 | 0.40370336 | 0.038935634 |
| Klhl11 | 4319.38002 | 0.160517506 | 0.038992033 |
| Synj2 | 1629.53078 | 0.383431268 | 0.039099688 |
| Mgmt | 785.543326 | -0.348411248 | 0.039115502 |
| Rnf145 | 2242.26716 | 0.196298543 | 0.039134328 |
| Mapk8 | 3385.0847 | 0.307890993 | 0.039134328 |
| Adcyap1 | 57.2649544 | 0.433169776 | 0.039160529 |
| Sdhc | 5691.10403 | -0.188111689 | 0.039218449 |
| Atp6v0c | 26865.3846 | 0.196647634 | 0.039249309 |
| Fkrp | 2093.16931 | 0.231222234 | 0.039269287 |
| Cecr2 | 946.070235 | -0.288278891 | 0.039285658 |
| Yod1 | 87.9441118 | 0.363988945 | 0.039418057 |
| RGD130953 | 1293.91448 | -0.247631737 | 0.039484693 |
| Vcpip1 | 5203.41188 | 0.126604835 | 0.039695025 |
| Tlr9 | 10.0489122 | -0.473540712 | 0.039756256 |
| Cartpt | 356.031375 | 0.380372406 | 0.039785089 |
| Trim3 | 3006.23518 | 0.22234398 | 0.039805807 |
| Ccna2 | 14.9861465 | -0.439299019 | 0.039809376 |
| LOC108349 | 477.415128 | 0.413840528 | 0.039847252 |
| Kcnk9 | 2072.27055 | 0.258579005 | 0.03989693 |
| Prlhr | 140.639319 | 0.381937595 | 0.03990009 |
| Atg9a | 6013.21733 | 0.140058767 | 0.040107296 |
| Atxn7l2 | 315.381913 | 0.242814181 | 0.040134648 |
| Gnaz | 3546.28003 | 0.273391208 | 0.040150887 |
| Rbm38 | 622.842512 | -0.317838474 | 0.040278317 |
| Fbxo25 | 1259.23783 | 0.211022358 | 0.040278317 |
| Cdc34 | 803.15783 | 0.265410627 | 0.040300947 |
| Atp10d | 168.272382 | -0.358948082 | 0.040341288 |
| Slc4a2 | 2294.63344 | -0.246762626 | 0.040342799 |
| Leprot | 767.049718 | -0.252908996 | 0.040386941 |
| Rab3gap2 | 4504.14707 | 0.149246929 | 0.040402957 |

| | | | |
|---|---|---|---|
| St8sia4 | 162.269687 | 0.356144442 | 0.040408159 |
| Ensa | 5774.10769 | 0.244919669 | 0.040420971 |
| Disp2 | 22133.9979 | 0.265291749 | 0.040420971 |
| LOC100909 | 34.6764518 | -0.376931752 | 0.040433836 |
| Rpl36a | 769.105784 | -0.246560866 | 0.040558379 |
| RGD156454 | 1323.53099 | -0.184226737 | 0.040558379 |
| Naca | 4874.61945 | -0.148814131 | 0.040684591 |
| Bfar | 1363.44466 | -0.158578687 | 0.040763677 |
| Trim47 | 607.718321 | -0.381902919 | 0.040856176 |
| LOC108352 | 11.1410795 | -0.456513471 | 0.040865558 |
| MGC95208 | 231.791116 | -0.322611643 | 0.040878252 |
| Kmt2b | 2041.48352 | 0.157885002 | 0.040881206 |
| Grm7 | 1415.39621 | 0.275651556 | 0.040896272 |
| Smim19 | 877.569975 | -0.22275497 | 0.040903398 |
| Tnfrsf11a | 190.969892 | -0.371921209 | 0.040907276 |
| Rpl22 | 554.882857 | -0.248081909 | 0.040907276 |
| Nol7 | 1239.48154 | -0.188914498 | 0.040907276 |
| Lrp1b | 3137.70743 | 0.244273211 | 0.040907276 |
| LOC108352 | 18.9160455 | -0.422410587 | 0.040938779 |
| Uap1l1 | 931.034122 | -0.282988065 | 0.040938779 |
| Sec61a1 | 3658.5819 | -0.218173729 | 0.040938779 |
| Osbpl10 | 1052.60917 | 0.303990907 | 0.040938779 |
| Tacc2 | 2947.93835 | 0.206493034 | 0.040991633 |
| Cep63 | 509.802412 | -0.228626663 | 0.041045329 |
| Map6 | 4739.57047 | 0.270525426 | 0.041061068 |
| Aff4 | 16365.0439 | 0.14642435 | 0.04118618 |
| Lipm | 4.7432704 | 0.520515174 | 0.041217524 |
| Zbtb38 | 4301.41594 | 0.147671786 | 0.041228481 |
| Tesmin | 10.9800456 | -0.427805311 | 0.041230474 |
| Tnfaip8l2 | 11.1857811 | -0.424833235 | 0.041232154 |
| Itpa | 652.614771 | 0.217952804 | 0.041232154 |
| Flt4 | 28.5374033 | 0.412415461 | 0.041284474 |
| Gpkow | 1919.15215 | -0.180510768 | 0.041296031 |
| Alg2 | 5591.78448 | 0.169409792 | 0.041373371 |
| Rnft2 | 3436.32583 | 0.208424324 | 0.041423122 |
| Map2k4 | 1217.10119 | 0.27413696 | 0.041423122 |

| | | | |
|---|---|---|---|
| Magt1 | 839.857352 | -0.274175428 | 0.04144867 |
| Aftph | 3107.40467 | 0.140241293 | 0.041485321 |
| LOC108351 | 10.0303264 | -0.423729984 | 0.041518613 |
| Slc41a3 | 272.621581 | -0.310092433 | 0.041518613 |
| Pcsk4 | 99.3560229 | 0.362146721 | 0.041518613 |
| Nme2 | 3506.08598 | -0.196687991 | 0.041537445 |
| Crebl2 | 2295.8817 | 0.153374831 | 0.041599164 |
| Sv2a | 18109.1117 | 0.265939923 | 0.041599164 |
| Vps26b | 7055.80015 | 0.130170614 | 0.041642417 |
| Prrg2 | 59.652376 | -0.392535289 | 0.041675139 |
| B4galt5 | 1039.29565 | -0.341419784 | 0.041745467 |
| Papd5 | 2905.27424 | 0.167460573 | 0.041802945 |
| Wipf2 | 2639.85705 | 0.19427303 | 0.041802945 |
| Dpp10 | 2948.54655 | 0.303480659 | 0.041802945 |
| Znhit3 | 324.116448 | 0.345337841 | 0.04188822 |
| Flrt2 | 3795.76198 | 0.162292391 | 0.041943696 |
| Ptgr1 | 8.06536009 | -0.468879961 | 0.041965285 |
| Rab3c | 3722.40971 | 0.274673513 | 0.041965285 |
| Icoslg | 563.361532 | -0.317903157 | 0.042057783 |
| Stau2 | 2560.55373 | 0.211948614 | 0.04207126 |
| Lrrc47 | 2293.75795 | 0.13313724 | 0.042089524 |
| Pgd | 1879.00251 | -0.172584087 | 0.042121085 |
| Tgif2 | 78.4584757 | -0.397579679 | 0.042137148 |
| Rgs8 | 3558.49268 | 0.197933005 | 0.042137148 |
| Foxo6 | 495.896811 | 0.311300161 | 0.042154953 |
| Aifm2 | 1009.0213 | -0.25436524 | 0.042232611 |
| St3gal1 | 1157.61476 | 0.341754589 | 0.042232611 |
| Arhgap44 | 9110.82283 | 0.306415668 | 0.042251231 |
| Papss1 | 4054.12199 | -0.192572823 | 0.042297711 |
| Serinc1 | 27638.2124 | 0.166058008 | 0.042440644 |
| Xpo7 | 4557.0324 | 0.147897083 | 0.042467716 |
| Gtf2h5 | 1153.52758 | -0.18206137 | 0.042488573 |
| Tmem151b | 3798.11717 | 0.269369373 | 0.042499222 |
| Nfkb1 | 1108.16852 | -0.232595674 | 0.042550544 |
| Tmem59l | 5919.60872 | 0.265311928 | 0.042550544 |
| Serp2 | 676.268865 | 0.287398236 | 0.042550544 |

| | | | |
|---|---|---|---|
| Setd1b | 2680.03926 | 0.136638984 | 0.042572794 |
| Chga | 6658.94383 | 0.276487454 | 0.042582335 |
| Mthfr | 513.301825 | 0.242567097 | 0.042589463 |
| Slc29a3 | 476.226487 | -0.30960754 | 0.042689641 |
| Ccdc61 | 223.639467 | -0.24698256 | 0.042689641 |
| Mapt | 22056.5244 | 0.171462703 | 0.042706527 |
| Chmp2a | 2505.85905 | -0.178745375 | 0.042896594 |
| Rnasel | 506.169739 | -0.375656827 | 0.042928424 |
| Kdm3b | 3263.88188 | -0.185133072 | 0.042931395 |
| LOC103689 | 376.071959 | 0.22780713 | 0.042931395 |
| Tectb | 12.2527162 | -0.407396753 | 0.042967825 |
| Zeb2 | 8260.37561 | -0.203200828 | 0.043036163 |
| Ptger1 | 6.99902302 | 0.458471498 | 0.043139536 |
| Taco1 | 233.625415 | 0.239182983 | 0.043188676 |
| LOC102551 | 1404.99904 | 0.282503002 | 0.043188676 |
| Mien1 | 589.498713 | -0.237549204 | 0.043205695 |
| P2ry4 | 37.712303 | -0.396732528 | 0.043398428 |
| Cd320 | 515.469655 | -0.250156447 | 0.043437105 |
| Thsd7a | 2283.7682 | 0.188212413 | 0.043437105 |
| Tmem51 | 928.866903 | -0.291700381 | 0.043484519 |
| Sgk2 | 11.2372234 | -0.450078238 | 0.043499499 |
| Tp53i3 | 345.71342 | -0.280511897 | 0.043499499 |
| Atp6v1c1 | 5721.28706 | 0.237090592 | 0.043499499 |
| NEWGENE_ | 2760.63767 | 0.204876192 | 0.043541813 |
| Dock3 | 12314.879 | 0.309301838 | 0.043541813 |
| Cyth4 | 32.1901927 | -0.404639758 | 0.043545437 |
| Nfatc3 | 793.392313 | -0.254437115 | 0.043577489 |
| Ttpal | 1702.32955 | 0.278172656 | 0.043577489 |
| Tnfsf10 | 15.3770561 | -0.499945832 | 0.043615096 |
| Ppp1r14b | 291.211376 | -0.248581608 | 0.04368018 |
| Mkrn3 | 36.4332048 | -0.395848173 | 0.043687624 |
| Dyrk1b | 1505.19095 | 0.158704396 | 0.043695213 |
| Endou | 59.3734212 | -0.350873631 | 0.043753238 |
| Nfkb2 | 219.073882 | -0.384304198 | 0.04378903 |
| Mdfic | 433.836887 | -0.315684626 | 0.04378903 |
| Egln3 | 1090.76745 | -0.287209636 | 0.04378903 |

| | | | |
|---|---|---|---|
| LOC100912 | 293.843873 | -0.262513803 | 0.04378903 |
| Fcho1 | 924.515092 | 0.255303902 | 0.04378903 |
| Dock11 | 861.164836 | -0.227790421 | 0.043794714 |
| Tmtc1 | 5198.83318 | 0.291445002 | 0.043809019 |
| H2afv | 932.688074 | -0.277145161 | 0.043844078 |
| Jak2 | 2244.06445 | -0.269832332 | 0.043844078 |
| Man2b2 | 769.418823 | -0.229932812 | 0.043879959 |
| Hey1 | 1116.31989 | -0.262336531 | 0.043889622 |
| Agk | 960.056694 | 0.21086482 | 0.043903242 |
| Srcin1 | 11200.5583 | 0.329882272 | 0.043908545 |
| Zbtb49 | 169.200466 | 0.268914159 | 0.044025391 |
| Usp5 | 2247.03379 | 0.167996239 | 0.044082378 |
| Cnksr3 | 725.11183 | -0.301254094 | 0.044219481 |
| Gpld1 | 2458.82812 | -0.264496421 | 0.044239144 |
| Tcf15 | 25.5086502 | 0.398630155 | 0.044548988 |
| Garem1 | 442.602567 | 0.299031143 | 0.044608344 |
| Kcne5 | 275.268928 | -0.346323823 | 0.044618545 |
| LOC100360 | 127.305267 | 0.293737577 | 0.044618545 |
| Asic2 | 2011.69007 | 0.226492183 | 0.044756183 |
| Adam12 | 180.607589 | -0.340645731 | 0.044785063 |
| Vstm2b | 1326.99892 | 0.350224172 | 0.044858771 |
| Fam212b | 1926.89461 | 0.361929769 | 0.044888492 |
| LOC103694 | 74.2521451 | -0.370218013 | 0.044958778 |
| Acin1 | 4724.33516 | 0.160720171 | 0.044958778 |
| LOC102553 | 47.8050245 | 0.38168155 | 0.044975357 |
| Mrps11 | 570.495089 | -0.17115296 | 0.045033746 |
| Zfp52 | 88.7361127 | -0.345799138 | 0.045063806 |
| Olfml1 | 3414.67373 | -0.268039364 | 0.045063806 |
| Zfp395 | 883.764168 | -0.259780815 | 0.045063806 |
| Ccdc90b | 663.366991 | -0.245723327 | 0.045063806 |
| Mpz | 22.168167 | 0.392791656 | 0.045063806 |
| Csf2rb | 5.83340393 | -0.498167475 | 0.045070753 |
| Tssc4 | 557.259868 | -0.219104317 | 0.045090176 |
| Ccng1 | 8144.97209 | -0.27421328 | 0.04509314 |
| Msrb1 | 625.533828 | 0.250960359 | 0.045193213 |
| Fbxl19 | 2201.38857 | 0.25296737 | 0.045250669 |

| | | | |
|---|---|---|---|
| Emc2 | 1806.25944 | -0.150818951 | 0.04528839 |
| Fxyd3 | 39.3146753 | -0.38391678 | 0.045297778 |
| Ywhag | 46783.9087 | 0.243562974 | 0.045297778 |
| LOC103691 | 7.74116633 | -0.417575082 | 0.045375356 |
| Vgll4 | 1285.31798 | -0.232472444 | 0.045410101 |
| Rhbdd1 | 715.471974 | -0.312711844 | 0.045436108 |
| Mrps6 | 668.919053 | -0.257139827 | 0.04553358 |
| Tmcc2 | 3750.20818 | 0.247091475 | 0.04553358 |
| Pxdc1 | 290.226602 | -0.326656006 | 0.045606409 |
| Cadps | 4734.76019 | 0.301389715 | 0.045641515 |
| Rnf185 | 1478.0902 | 0.183865322 | 0.045649821 |
| Zfp423 | 5356.35239 | -0.232975457 | 0.045664133 |
| Pard6a | 305.644803 | 0.268271166 | 0.045694249 |
| Anln | 243.461068 | -0.40147316 | 0.045853035 |
| Elac2 | 1486.0867 | 0.173822451 | 0.045859878 |
| Mrpl3 | 1928.62559 | -0.158015157 | 0.045896182 |
| RGD156213 | 2467.7478 | -0.15811232 | 0.045927534 |
| Naglu | 274.172689 | -0.301778723 | 0.045986878 |
| Tle6 | 7.95729664 | -0.42652129 | 0.046018538 |
| Pald1 | 340.742551 | -0.340925409 | 0.046018538 |
| Clasp1 | 4022.42358 | 0.154175201 | 0.046072726 |
| Dut | 465.330229 | 0.290481828 | 0.046132531 |
| Nusap1 | 13.9953217 | -0.420688002 | 0.046272185 |
| Rmdn1 | 599.401992 | -0.251209316 | 0.046272185 |
| Nfkbia | 617.448385 | -0.362618747 | 0.046417686 |
| Bend6 | 1105.01411 | 0.309282786 | 0.046417686 |
| Kif26a | 443.584376 | -0.212753912 | 0.046469808 |
| Mrps30 | 882.834845 | 0.192947109 | 0.046469808 |
| Stxbp5 | 5881.46384 | 0.271041873 | 0.046469808 |
| Lpgat1 | 6725.4045 | 0.26021956 | 0.046530349 |
| Add3 | 20193.0805 | -0.218713907 | 0.046565022 |
| Rtf1 | 3469.34264 | 0.165382637 | 0.046588428 |
| Wdr33 | 2150.4631 | 0.147097906 | 0.046629938 |
| LOC100912 | 154.695639 | -0.304684667 | 0.046697261 |
| Wdr5 | 614.019723 | 0.197457797 | 0.046727933 |
| Pigt | 5320.79975 | -0.120656561 | 0.046774835 |

| | | | |
|---|---|---|---|
| Rimklb | 1376.67708 | -0.234373848 | 0.046778517 |
| Trim68 | 388.116065 | -0.228775811 | 0.046778517 |
| RGD130918 | 3131.03772 | 0.146467147 | 0.046835826 |
| Cnih3 | 153.156701 | 0.389183085 | 0.046835826 |
| LOC102554 | 26.1575563 | -0.388192824 | 0.046923082 |
| LOC100912 | 111.341853 | -0.317167353 | 0.046923082 |
| Traf2 | 461.373222 | -0.297160816 | 0.046923082 |
| Rbm34 | 989.450995 | -0.16992091 | 0.046923082 |
| Pld3 | 10982.1521 | 0.228961129 | 0.046923082 |
| Casp3 | 1109.22852 | -0.270664947 | 0.047072979 |
| Btf3 | 3094.61804 | -0.197176069 | 0.047072979 |
| Abcg4 | 2207.15929 | 0.237245542 | 0.047206095 |
| Spred3 | 1334.85358 | 0.209547309 | 0.047263366 |
| Chst8 | 135.334168 | 0.316067399 | 0.047292856 |
| Hn1l | 247.394361 | -0.359877912 | 0.047341116 |
| Trmt2a | 578.220757 | -0.206241232 | 0.047390281 |
| Fam69b | 1371.9868 | 0.249220813 | 0.047390281 |
| Ahnak | 3389.11064 | -0.386150397 | 0.047536948 |
| Hexa | 1892.27061 | -0.172997448 | 0.047536948 |
| Ncor1 | 8486.82174 | 0.159548555 | 0.047603746 |
| Eml5 | 869.946646 | 0.192000343 | 0.047619343 |
| Cep295 | 335.504615 | 0.261954118 | 0.047701344 |
| Fam83d | 16.6048494 | -0.397900297 | 0.047767797 |
| Qrich2 | 6.69555985 | -0.453757303 | 0.047854215 |
| Nr1h2 | 1794.87104 | -0.198673036 | 0.047854215 |
| Ptprm | 3864.05334 | 0.140855056 | 0.047854215 |
| Efemp1 | 1122.22815 | -0.29532692 | 0.047948468 |
| Brd4 | 7295.839 | 0.15652759 | 0.047948468 |
| Asmt | 6.30786094 | 0.431098519 | 0.047992407 |
| Cdkl2 | 1446.19307 | 0.317004898 | 0.048139677 |
| Sox8 | 4473.02385 | -0.244369712 | 0.04814223 |
| Fhod3 | 3069.52888 | 0.312316748 | 0.048144759 |
| Ankrd28 | 3485.66646 | -0.190150502 | 0.048183697 |
| Cpne8 | 893.341883 | -0.298514461 | 0.04828007 |
| Dusp18 | 4194.09742 | -0.268764318 | 0.048349155 |
| Vps25 | 985.999803 | -0.163035032 | 0.048353559 |

| | | | |
|---|---|---|---|
| Hist1h2bh | 1587.6386 | -0.243901451 | 0.048386496 |
| Mocs2 | 1580.55272 | -0.198949602 | 0.048386496 |
| Pip4k2c | 1505.35204 | 0.27550414 | 0.048386496 |
| Cdh2 | 13302.9281 | -0.141178217 | 0.048389531 |
| Arl8a | 8580.89706 | -0.157835102 | 0.048708678 |
| Arrdc3 | 969.747608 | -0.292409941 | 0.048732733 |
| LOC100911 | 5.90146531 | -0.478735165 | 0.048758418 |
| Zfp131 | 1211.26853 | 0.158626541 | 0.048946038 |
| Kctd4 | 722.648872 | 0.28476313 | 0.049069711 |
| Vps39 | 3579.89886 | 0.130111356 | 0.049218903 |
| Klhdc3 | 4448.47777 | 0.192835252 | 0.049226808 |
| LOC316820 | 76.3673105 | -0.313310483 | 0.049267123 |
| Ppp1cc | 7507.09732 | 0.154637692 | 0.049267123 |
| Plscr3 | 388.380619 | -0.271245212 | 0.0493617 |
| Glul | 162252.084 | -0.228030123 | 0.04945323 |
| Bace2 | 141.248974 | -0.366606633 | 0.049571932 |
| Ss18l1 | 555.849763 | 0.22900915 | 0.049571932 |
| Lzts2 | 1700.73111 | -0.207713361 | 0.049704199 |
| Whsc1 | 4411.10968 | 0.159857881 | 0.049708309 |
| Tns1 | 4662.85825 | -0.249838838 | 0.04971817 |
| Snap47 | 900.570566 | 0.255913184 | 0.04971817 |
| Erp29 | 1929.34801 | 0.111062978 | 0.049757089 |
| Anapc10 | 625.885522 | -0.225587207 | 0.049759844 |
| Cdh9 | 621.387166 | 0.34170521 | 0.049774752 |
| Anp32e | 6616.36176 | -0.158213667 | 0.049784082 |
| Baalc | 7575.57837 | -0.175237607 | 0.049841524 |
| Mier1 | 2257.84131 | -0.18491549 | 0.04996129 |
| Arrb1 | 24772.3564 | 0.157432003 | 0.04996129 |
| Mafb | 703.08562 | 0.312773499 | 0.049969104 |

| Gene | baseMean | log2FoldChange | adjusted p-value |
|---|---|---|---|
| Apol3 | 64.7870671 | -3.3025258 | 0.01901468 |
| Fgr | 35.87201 | -3.01243746 | 0.01901468 |
| Casp4 | 238.590496 | -1.9553397 | 0.04852069 |
| Hpd | 84.1016083 | 1.066806972 | 0.04852069 |
| Irf1 | 1126.89521 | -2.02259695 | 0.04852069 |
| Klk6 | 198.537556 | -2.04030645 | 0.04852069 |
| Nlrc5 | 371.143833 | -2.0025819 | 0.04852069 |
| Sec14l4 | 20.3061946 | -2.33712411 | 0.04852069 |
