## Supplementary material for "Investigating cocaine- and abstinence-induced effects on astrocyte gene expression in the nucleus accumbens": Exp 2_significant DEGs_supplementary data

| Gene | baseMean | log2FoldChange | adjusted p-value |
|---|---|---|---|
| Fosb | 361.8514835 | 1.425500044 | 7.69E-08 |
| Slc4a11 | 143.1922978 | 1.412162003 | 4.24E-07 |
| Ripk4 | 44.65504897 | 1.369077262 | 7.78E-05 |
| Gpr6 | 440.7409934 | 1.351973654 | 2.41E-08 |
| Krt77 | 36.53058009 | 1.350914689 | 0.00058095 |
| Ptpn7 | 23.53600203 | 1.280029641 | 0.001288941 |
| Bhlhe23 | 39.6585233 | 1.278866334 | 0.000547921 |
| Wfs1 | 3962.317004 | 1.135102106 | 4.52E-10 |
| Itk | 47.5648701 | 1.005080514 | 0.001439153 |
| Gpr88 | 3692.090795 | 0.995317072 | 2.74E-10 |
| Npas4 | 362.011499 | 0.968501703 | 0.001288941 |
| Cartpt | 282.3164318 | 0.962501618 | 0.000135463 |
| Iqgap3 | 175.4441161 | 0.962417072 | 4.64E-06 |
| Sh3rf2 | 180.7510921 | 0.95848799 | 1.75E-05 |
| Penk | 5862.017377 | 0.932227632 | 6.53E-12 |
| Rgs9 | 1292.779767 | 0.922613141 | 5.16E-10 |
| Nexn | 356.922281 | 0.922200053 | 5.40E-05 |
| Egr4 | 804.1976775 | 0.917751657 | 5.93E-05 |
| P2ry1 | 309.6280162 | 0.859071824 | 5.09E-06 |
| Gng7 | 3688.405217 | 0.851889516 | 1.78E-14 |
| Tac3 | 504.7993661 | 0.844946202 | 1.55E-05 |
| Prkch | 314.4142843 | 0.832999325 | 1.74E-05 |
| Akap5 | 3549.165687 | 0.830194231 | 9.44E-09 |
| Adora2a | 1059.212289 | 0.820272177 | 4.76E-08 |
| Slc26a10 | 19.55224478 | 0.813941844 | 0.007179741 |
| Scn4b | 2014.67136 | 0.811908972 | 2.58E-06 |
| Calcr | 385.1664502 | 0.805228884 | 0.005633552 |
| Krt71 | 179.7581875 | 0.797704937 | 0.000138318 |
| Lamp5 | 659.1269984 | 0.779041652 | 7.73E-07 |
| Pde1c | 440.626785 | 0.772847949 | 1.58E-06 |
| Otof | 1233.681645 | 0.768974733 | 0.000498523 |
| Ptpn5 | 2961.43227 | 0.764146604 | 5.79E-15 |
| Foxp2 | 1693.100371 | 0.764096294 | 8.00E-08 |
| Pde10a | 3128.744671 | 0.756027189 | 3.15E-07 |
| Kcnj2 | 121.1224427 | 0.750181409 | 0.001172036 |

| | | | |
|---|---|---|---|
| Gnal | 4382.720362 | 0.745210017 | 3.87E-07 |
| Lppr1 | 296.7316327 | 0.734781656 | 0.000249744 |
| Drd1 | 2460.609413 | 0.732321348 | 2.78E-06 |
| Rxrg | 513.5306764 | 0.731119854 | 2.64E-05 |
| Syndig1 | 553.8217192 | 0.710597229 | 1.51E-06 |
| Ebf1 | 85.02903459 | 0.70196253 | 0.001479987 |
| Filip1 | 795.8498352 | 0.698298521 | 0.000266975 |
| Cpne5 | 4955.728076 | 0.683049311 | 9.84E-12 |
| Dclk3 | 1279.65023 | 0.68228114 | 7.89E-06 |
| Pthlh | 77.04819435 | 0.667648229 | 0.002762787 |
| Tec | 110.6742099 | 0.666628975 | 0.000790124 |
| Pdyn | 2314.972886 | 0.6655283 | 1.05E-05 |
| Dlk1 | 1070.675033 | 0.662766537 | 0.004580222 |
| Kcna5 | 324.3681168 | 0.652540989 | 1.96E-06 |
| Dlx5 | 475.2319091 | 0.643835062 | 5.95E-05 |
| Snai1 | 11.82066188 | 0.639724774 | 0.014518947 |
| Gpr52 | 295.6522703 | 0.639080982 | 3.69E-05 |
| Pde7b | 1256.379792 | 0.636514129 | 8.00E-08 |
| Kcnh4 | 195.5494691 | 0.627845656 | 0.000439314 |
| Rasd2 | 4370.532188 | 0.625908967 | 2.06E-08 |
| Klf5 | 914.7900961 | 0.625103779 | 8.85E-06 |
| Drd2 | 1655.698278 | 0.620184124 | 2.73E-06 |
| Lingo3 | 630.4617561 | 0.612840601 | 3.70E-06 |
| Olfml2a | 40.09550527 | 0.610914436 | 0.008941216 |
| Sp9 | 514.027378 | 0.609832089 | 1.12E-08 |
| Rem2 | 209.4541881 | 0.608351723 | 0.002140696 |
| Rasgrp2 | 660.1175317 | 0.600886468 | 2.41E-10 |
| Opn3 | 396.5479666 | 0.598252034 | 3.45E-06 |
| Pde3a | 740.8545686 | 0.596070969 | 4.85E-06 |
| Pde1b | 4099.810897 | 0.584891592 | 6.66E-09 |
| Lpl | 1505.027139 | 0.583656837 | 0.000377731 |
| Myh7 | 242.4345364 | 0.582839929 | 0.00073841 |
| Pdzd2 | 2168.888335 | 0.582311249 | 0.000662168 |
| Sv2c | 2210.141846 | 0.581879875 | 5.22E-06 |
| Unc13c | 740.7607189 | 0.573857125 | 0.000991409 |
| Necab2 | 1834.509526 | 0.565417767 | 2.87E-07 |

| | | | |
|---|---|---|---|
| Kit | 3144.941767 | 0.563091758 | 8.24E-06 |
| Cdc42ep3 | 349.6067523 | 0.559804181 | 0.000681056 |
| Bcl11b | 5738.545239 | 0.559415476 | 2.19E-05 |
| Adcy5 | 9082.114656 | 0.556957696 | 2.50E-06 |
| Cacna1h | 1270.033219 | 0.549674115 | 0.000443011 |
| Gucy1a3 | 1899.635282 | 0.549426597 | 1.28E-06 |
| Sertad4 | 541.6449117 | 0.542556723 | 3.41E-05 |
| Il1rapl2 | 364.7330538 | 0.542127825 | 0.000173713 |
| Prkg1 | 121.1345134 | 0.539534229 | 0.004996134 |
| Foxp1 | 1725.540346 | 0.528125636 | 0.000175822 |
| Wipf3 | 4144.753322 | 0.52738856 | 1.05E-05 |
| Lzts3 | 2519.475395 | 0.523321716 | 9.19E-08 |
| Chrm4 | 1015.902001 | 0.522491019 | 0.000225225 |
| Wnk4 | 130.5097941 | 0.520749657 | 0.002440353 |
| Kl | 85.70817809 | 0.519237231 | 0.013242625 |
| Arpp21 | 8234.651489 | 0.515908185 | 8.18E-05 |
| Zfp503 | 334.4780086 | 0.511303194 | 0.003892886 |
| Prkg2 | 267.189987 | 0.509884999 | 0.000547921 |
| Meis2 | 5020.62155 | 0.495549235 | 4.87E-09 |
| Asb2 | 102.6090581 | 0.494208409 | 0.006664891 |
| Acvr1c | 167.7972018 | 0.493987434 | 0.002656757 |
| Hpca | 6647.641839 | 0.493002836 | 0.000359237 |
| Synpr | 2853.159898 | 0.492894605 | 1.64E-07 |
| Actn2 | 953.5597086 | 0.491694251 | 0.002190992 |
| Lmo7 | 1735.253239 | 0.491373028 | 0.00127058 |
| Kcns2 | 241.8978562 | 0.486744772 | 0.006922797 |
| Rab40b | 707.7123691 | 0.484743325 | 0.001288941 |
| Syndig1l | 492.9232635 | 0.48398858 | 0.000250533 |
| Ppp1r1b | 12849.45472 | 0.477849867 | 3.62E-05 |
| Trerf1 | 1151.851265 | 0.475248733 | 5.23E-06 |
| Htr1b | 271.3883885 | 0.474826984 | 0.003547552 |
| Ppp3ca | 13759.26458 | 0.472922471 | 8.14E-05 |
| B3gnt2 | 658.5211906 | 0.472716509 | 1.28E-06 |
| RGD1564664 | 1201.637324 | 0.455417008 | 4.08E-06 |
| Junb | 1525.00745 | 0.454220725 | 0.011149816 |
| Asic4 | 1085.854724 | 0.452236745 | 0.001250121 |

| | | | |
|---|---|---|---|
| Egr2 | 208.3089698 | 0.451180741 | 0.024441419 |
| Ppp1r9a | 5252.033452 | 0.444143854 | 0.000371162 |
| Chst15 | 388.2960473 | 0.443348534 | 0.004818988 |
| Tiam1 | 4432.125071 | 0.440736149 | 3.66E-05 |
| Doc2b | 510.3187311 | 0.440173686 | 0.001915782 |
| Pou3f1 | 1020.556314 | 0.439806701 | 0.006627314 |
| Gpr83 | 1074.029308 | 0.432249477 | 6.04E-05 |
| Mast3 | 5755.988864 | 0.430825311 | 0.002122955 |
| Cyld | 2840.238405 | 0.430426786 | 2.94E-09 |
| Kitlg | 480.8570585 | 0.428687145 | 0.001288941 |
| Gpr149 | 287.1385802 | 0.42812531 | 0.009315766 |
| Ncdn | 38472.0715 | 0.427404865 | 4.30E-10 |
| Fstl4 | 602.0896375 | 0.4182616 | 0.003094077 |
| Txlng | 233.5152561 | 0.417897351 | 0.000205591 |
| Cntn3 | 470.752797 | 0.416781016 | 0.002233627 |
| Samd12 | 134.026194 | 0.416732657 | 0.008957101 |
| Fras1 | 1141.007123 | 0.415812874 | 0.002329487 |
| Crip3 | 9.313828631 | 0.415788004 | 0.032603398 |
| Rarb | 962.3814887 | 0.415561634 | 0.001101101 |
| Ppp4r4 | 433.5174745 | 0.415282665 | 0.001054149 |
| Sptb | 5510.750692 | 0.41526066 | 0.000490086 |
| Igfbp4 | 1526.980126 | 0.410151917 | 0.002602523 |
| Mycn | 239.4104249 | 0.409072676 | 0.003550219 |
| Erbb4 | 3173.64337 | 0.407195019 | 7.41E-09 |
| Map2k1 | 6640.415635 | 0.406924245 | 1.12E-06 |
| RGD1311739 | 1744.629014 | 0.406725018 | 5.39E-05 |
| Fbxl16 | 3707.583559 | 0.405472075 | 0.001231423 |
| Egr3 | 697.7009269 | 0.398610416 | 0.021557802 |
| Ptchd1 | 308.239854 | 0.395090124 | 0.004078446 |
| Rnf144b | 387.3649307 | 0.393750234 | 0.001513 |
| Rragd | 673.4786561 | 0.389656193 | 0.000386956 |
| Actn1 | 2500.685878 | 0.387831845 | 0.000503861 |
| Epha4 | 6512.508245 | 0.381838984 | 5.74E-05 |
| Syt10 | 478.4317984 | 0.381767956 | 0.012374565 |
| Ankrd55 | 279.1412545 | 0.380479232 | 0.01343357 |
| Rgs2 | 449.5986831 | 0.379467479 | 0.00333904 |

| | | | |
|---|---|---|---|
| Crtc1 | 2891.981055 | 0.377042749 | 3.69E-06 |
| Gpr176 | 513.2134578 | 0.371949783 | 0.00138013 |
| Frem3 | 100.6885492 | 0.371556461 | 0.021294141 |
| Htr2c | 3571.271147 | 0.371011579 | 0.004141515 |
| Cacna1c | 2405.318727 | 0.370630672 | 0.001187652 |
| Dgkb | 9484.213167 | 0.369601803 | 0.003797014 |
| Zfp483 | 3306.93467 | 0.365697411 | 0.000200044 |
| Begain | 2319.821104 | 0.364924523 | 0.000179332 |
| Crabp1 | 1176.952628 | 0.364898158 | 0.008941216 |
| Gad1 | 19063.36109 | 0.364844542 | 0.000681056 |
| Ago2 | 2824.45661 | 0.362113337 | 3.88E-05 |
| Galnt13 | 672.5262929 | 0.362039599 | 0.003699299 |
| Cacnb4 | 3454.949447 | 0.361519526 | 0.000652398 |
| C2cd2l | 6536.597542 | 0.36132835 | 0.0002469 |
| Elmod1 | 4820.480998 | 0.360556615 | 0.000106518 |
| Mtus2 | 205.9264079 | 0.359426988 | 0.004311259 |
| Lrrk2 | 2983.365092 | 0.358429715 | 0.002167426 |
| LOC100125362 | 4464.312302 | 0.358340836 | 0.000441904 |
| Abcc5 | 2902.305545 | 0.358072402 | 0.000129162 |
| Camk4 | 846.2195121 | 0.358038978 | 0.007338459 |
| Oprm1 | 658.4202902 | 0.358025372 | 0.004576093 |
| RGD1309779 | 58.47335605 | 0.356798881 | 0.019265285 |
| Sytl5 | 310.9248264 | 0.35555513 | 0.007744014 |
| Adamts3 | 509.5222158 | 0.354537538 | 0.007682251 |
| Slc2a13 | 7433.145343 | 0.354220026 | 1.95E-05 |
| Acer2 | 47.35441621 | 0.353951993 | 0.029088595 |
| Lrrc10b | 334.7533522 | 0.353367939 | 0.009306671 |
| Man1c1 | 1303.912027 | 0.3516634 | 2.64E-05 |
| Nkx2-1 | 429.6306998 | 0.349792336 | 0.014596433 |
| Zfp575 | 404.6630034 | 0.3487401 | 0.003309345 |
| Flrt3 | 1288.251502 | 0.347886888 | 0.001491742 |
| Pxdn | 251.1942145 | 0.345725341 | 0.019341006 |
| Gpr173 | 113.8205336 | 0.344810205 | 0.008995403 |
| Zfp57 | 322.7171161 | 0.344071276 | 0.002578923 |
| Pcdh8 | 2955.577775 | 0.343943327 | 0.00035076 |
| Gpr101 | 946.6827241 | 0.343910489 | 0.026339895 |

| | | | |
|---|---|---|---|
| Runx1t1 | 1377.167156 | 0.339920003 | 0.000249744 |
| Col14a1 | 115.8598169 | 0.339168699 | 0.028642697 |
| Man1a1 | 824.5609165 | 0.336690946 | 0.003091319 |
| Lrrc4c | 1917.838826 | 0.335760541 | 0.000114633 |
| Tac1 | 1029.543161 | 0.334886004 | 0.019294918 |
| Slc25a25 | 1655.412143 | 0.334779765 | 0.001447462 |
| Rap1gap | 9157.730239 | 0.334274447 | 1.91E-05 |
| Scn3a | 3256.676625 | 0.33361269 | 0.000328877 |
| Htr6 | 124.8280998 | 0.332021201 | 0.021109252 |
| Kctd8 | 480.8480409 | 0.330886604 | 0.013763992 |
| Jph4 | 3728.516887 | 0.33075276 | 0.003453503 |
| Cacnb2 | 1537.541359 | 0.329938099 | 0.000106518 |
| Tmem164 | 1348.407671 | 0.329283615 | 4.71E-05 |
| Camkk2 | 1352.051574 | 0.32833298 | 0.006960456 |
| Plxdc1 | 315.9379147 | 0.328020223 | 0.023697231 |
| Prrt2 | 3483.827254 | 0.326319777 | 0.002140696 |
| Hs6st2 | 1758.874293 | 0.326207494 | 0.000175319 |
| Trim66 | 892.4176885 | 0.323734123 | 0.000598239 |
| Kcnq2 | 6053.880849 | 0.321993563 | 0.001115714 |
| Dact2 | 830.6093251 | 0.32172696 | 0.000920901 |
| Cdk17 | 5280.469632 | 0.318597184 | 0.000498523 |
| Dnajb4 | 1074.45324 | 0.313232766 | 4.11E-05 |
| Tomm70a | 5381.255247 | 0.313216857 | 8.14E-05 |
| Kcna4 | 1258.438944 | 0.312788051 | 0.005290761 |
| Cpne6 | 1475.871419 | 0.312325261 | 0.007179741 |
| Wscd2 | 565.0684435 | 0.309531739 | 0.003117664 |
| Prkar2b | 2796.458808 | 0.309094621 | 4.18E-05 |
| Ablim2 | 1613.396806 | 0.307990059 | 6.03E-05 |
| Tle1 | 605.6993446 | 0.307921217 | 0.004007865 |
| Kcnab1 | 4760.611347 | 0.306906229 | 0.015749257 |
| Lypd1 | 2649.649068 | 0.303385204 | 0.003021024 |
| Gucy1b3 | 5489.256602 | 0.300529504 | 5.09E-06 |
| Psd | 6006.039369 | 0.299410681 | 0.012128629 |
| Slc9a5 | 428.9459319 | 0.297156489 | 0.010645495 |
| Phactr1 | 7164.961389 | 0.29650712 | 0.013391727 |
| Fkbp1a | 7589.237552 | 0.29625303 | 0.002656757 |

| | | | |
|---|---|---|---|
| Thsd7a | 1338.64745 | 0.29319729 | 0.001324144 |
| Jph3 | 7116.891693 | 0.293036009 | 0.0007495 |
| Kcnh5 | 446.4772838 | 0.292545474 | 0.033662171 |
| Fam102b | 1415.985297 | 0.291972352 | 0.008758482 |
| Ajap1 | 913.1561449 | 0.290336525 | 0.001238723 |
| Ablim3 | 683.141554 | 0.287592319 | 0.000861024 |
| Kcnip2 | 2032.534758 | 0.286405444 | 0.001171586 |
| Rbfox2 | 2049.24637 | 0.285099704 | 1.28E-05 |
| St18 | 557.3545678 | 0.284249357 | 0.02316479 |
| Grm5 | 4703.834607 | 0.283360493 | 0.002183788 |
| Kcnt1 | 1079.94717 | 0.28272568 | 0.00273455 |
| Nfx1 | 2370.53298 | 0.28244646 | 7.69E-08 |
| Phyhip | 10432.63686 | 0.280399571 | 0.01962778 |
| Gabrd | 501.2601206 | 0.278892128 | 0.040557349 |
| Rapgef5 | 2791.898987 | 0.276857108 | 0.002171855 |
| Smad3 | 1574.125646 | 0.276459399 | 0.004126857 |
| Rbfox1 | 6488.222247 | 0.275680621 | 0.019207251 |
| Nsg2 | 25229.31992 | 0.27543347 | 0.000156493 |
| RGD1563441 | 174.9640067 | 0.275373524 | 0.023239844 |
| Il10ra | 79.03241906 | 0.272673506 | 0.036015604 |
| Zfhx2 | 2459.257642 | 0.27259789 | 0.014830148 |
| Mef2d | 3839.655417 | 0.272303192 | 0.000863605 |
| Stc1 | 130.8640726 | 0.271342561 | 0.04443232 |
| Stox1 | 47.59382763 | 0.270830373 | 0.047748655 |
| Gria4 | 3334.99479 | 0.270815663 | 0.001516437 |
| Dhx35 | 447.7316242 | 0.270087503 | 0.006074981 |
| Klhl29 | 2317.392661 | 0.269003678 | 0.001187652 |
| Dlgap3 | 8347.265491 | 0.268561809 | 0.00691669 |
| Tex15 | 257.4680485 | 0.268401351 | 0.023587703 |
| Pdxp | 3506.174037 | 0.266455921 | 0.002656757 |
| Cacna2d3 | 4207.313942 | 0.26534532 | 0.00127058 |
| Pianp | 4829.912086 | 0.265051979 | 0.00127862 |
| Shank3 | 5227.139696 | 0.263919475 | 0.012128629 |
| Necab1 | 930.2600701 | 0.262741277 | 0.0305677 |
| Diaph1 | 1742.852887 | 0.262097886 | 0.002914511 |
| Tasp1 | 316.907582 | 0.261091468 | 0.018522563 |

| | | | |
|---|---|---|---|
| Fbxo41 | 3351.723453 | 0.259933853 | 0.002429263 |
| Arid4a | 1959.468231 | 0.257451636 | 0.00249232 |
| Zdhhc14 | 1138.058457 | 0.257328833 | 0.0084108 |
| Shisa7 | 4079.53129 | 0.256996482 | 0.010460634 |
| Slc24a4 | 235.5612437 | 0.256536408 | 0.046473668 |
| Sptbn4 | 5580.857413 | 0.256012671 | 0.008959981 |
| Zfp189 | 419.9390402 | 0.255771436 | 0.011893324 |
| Rybp | 1623.279209 | 0.253998608 | 0.002859736 |
| Slc12a5 | 16698.60919 | 0.252950902 | 0.005389283 |
| Slc35d3 | 699.6527435 | 0.252652609 | 0.030227168 |
| Cep295 | 261.2642565 | 0.251892154 | 0.039563908 |
| Dlg2 | 6290.830135 | 0.251075316 | 0.000645134 |
| Plekha5 | 1435.557174 | 0.250285729 | 0.002726555 |
| Trank1 | 3621.744768 | 0.249935857 | 0.024996271 |
| LRRTM1 | 1579.888408 | 0.249647983 | 0.001937757 |
| Rbm3 | 271.9303994 | 0.248373326 | 0.040478955 |
| Klf16 | 1723.600363 | 0.248291297 | 0.013132091 |
| Gabra4 | 1405.540428 | 0.247793005 | 0.003853418 |
| Cep41 | 323.249581 | 0.247239999 | 0.023657879 |
| Rapgef6 | 957.2703309 | 0.246798967 | 0.014011967 |
| Dapk1 | 4173.343172 | 0.245696261 | 0.002686209 |
| Gtpbp6 | 474.5915357 | 0.245527785 | 0.008398037 |
| Zfp280d | 1535.509615 | 0.245509634 | 0.001172036 |
| Grb10 | 2498.698367 | 0.245488282 | 0.035076854 |
| Vstm5 | 166.8239076 | 0.243848661 | 0.047655565 |
| Atp6ap1l | 333.5030067 | 0.243607074 | 0.047930523 |
| Syt4 | 7170.281079 | 0.243516544 | 0.004840734 |
| Grm4 | 1183.492015 | 0.242782683 | 0.040370181 |
| Nrxn3 | 9205.580118 | 0.24253103 | 0.00042908 |
| Ncoa5 | 2185.846261 | 0.242088013 | 0.000125917 |
| Pde8b | 2371.384063 | 0.237086981 | 0.005354322 |
| Samd14 | 1283.122592 | 0.236669503 | 0.024046946 |
| Cplx2 | 5065.699862 | 0.236106583 | 0.016371399 |
| Syt6 | 524.32761 | 0.235910744 | 0.049518515 |
| Pdpk1 | 2081.33563 | 0.235781585 | 0.000891074 |
| Cacnb3 | 3022.015453 | 0.234591709 | 0.011331309 |

| | | | |
|---|---|---|---|
| Rgs8 | 2864.826111 | 0.234043367 | 0.021866318 |
| Senp2 | 1432.693619 | 0.232981922 | 0.000370278 |
| Gria1 | 6767.768493 | 0.232421954 | 0.008667063 |
| Ppp1r2 | 5152.227819 | 0.232332992 | 0.000182812 |
| Rasgrf1 | 23921.54751 | 0.231909401 | 0.013312043 |
| Prkacb | 5158.31622 | 0.231720327 | 0.000700845 |
| Pnmal2 | 21043.17493 | 0.231427322 | 0.00086505 |
| Fbxo34 | 1479.058752 | 0.230741711 | 0.015934986 |
| Mbnl1 | 742.6808564 | 0.230159604 | 0.005757218 |
| Tmem158 | 942.0412048 | 0.229440317 | 0.034346593 |
| Ccdc85a | 580.7354562 | 0.229018455 | 0.015934986 |
| Stard13 | 716.519939 | 0.228486393 | 0.012011762 |
| Sipa1l1 | 8397.216829 | 0.22846626 | 0.014396978 |
| Peli3 | 518.0294181 | 0.228404739 | 0.018967014 |
| Mn1 | 3152.18817 | 0.227850566 | 0.003269266 |
| Camkk1 | 2642.452762 | 0.22747605 | 0.007321059 |
| Lppr4 | 6724.95128 | 0.227420887 | 0.02495681 |
| Stk32c | 3120.77321 | 0.22738221 | 0.028642697 |
| Tmem199 | 1105.143515 | 0.226234095 | 0.007321059 |
| Cacnb1 | 3424.208353 | 0.22402749 | 0.008318848 |
| Rims3 | 849.2891719 | 0.223906133 | 0.020022959 |
| Dlx2 | 317.7644714 | 0.223647066 | 0.033187246 |
| Dusp8 | 2779.8112 | 0.223160345 | 0.003061377 |
| Rhobtb2 | 3124.310062 | 0.222907806 | 0.001673135 |
| Ogfod1 | 1528.478683 | 0.22286227 | 0.003130929 |
| Ccdc64 | 680.3025258 | 0.222664709 | 0.018448171 |
| Efnb3 | 3778.706362 | 0.222605624 | 0.01917485 |
| Clvs2 | 286.1684588 | 0.221485522 | 0.025959706 |
| Pkia | 2411.30785 | 0.221160135 | 0.010494278 |
| Ypel2 | 881.7954708 | 0.220846108 | 0.039563908 |
| Map3k13 | 867.9413398 | 0.220431067 | 0.02257536 |
| Mypop | 1205.447633 | 0.220313377 | 0.007321059 |
| Irf2bp1 | 2086.528737 | 0.219727144 | 0.000199309 |
| Fam110b | 1095.80139 | 0.219340775 | 0.020045874 |
| Jakmip1 | 4793.434136 | 0.219077188 | 0.000306425 |
| Epb41l1 | 24200.34524 | 0.218958024 | 0.001765052 |

| | | | |
|---|---|---|---|
| Dos | 10418.42934 | 0.218816653 | 0.00254582 |
| Fam84a | 3423.763023 | 0.218648387 | 0.006280511 |
| Inafm1 | 399.110315 | 0.217054082 | 0.030534974 |
| Spock3 | 3465.979301 | 0.217003077 | 0.02597668 |
| Celf5 | 8385.921957 | 0.216817866 | 0.038251147 |
| Cyfip2 | 35728.84141 | 0.216431057 | 0.030939193 |
| Ppfia4 | 2443.128871 | 0.215900051 | 0.012906207 |
| Nyap1 | 3225.221896 | 0.215731201 | 3.91E-05 |
| Scn2b | 2274.139489 | 0.215130162 | 0.007962852 |
| Arhgap26 | 2444.079308 | 0.215092521 | 0.007774758 |
| Bag4 | 1420.970931 | 0.214788537 | 0.003424369 |
| Plekha8 | 454.1908934 | 0.213822735 | 0.024868187 |
| Zfp280b | 694.6085155 | 0.213643367 | 0.015664411 |
| Csrnp3 | 1556.920859 | 0.213485502 | 0.011416653 |
| Nsg1 | 5030.243571 | 0.213348435 | 0.025340738 |
| Mpped2 | 671.9781324 | 0.213340176 | 0.016157062 |
| Josd1 | 1127.149307 | 0.211836947 | 0.025626751 |
| Elk1 | 907.1609105 | 0.211507598 | 0.013604395 |
| Mafb | 571.071754 | 0.210933763 | 0.048328808 |
| Gripap1 | 5463.788491 | 0.210741928 | 0.004747813 |
| Slc7a14 | 3790.217104 | 0.210625901 | 0.006461345 |
| Arpc1a | 3475.36321 | 0.21018987 | 0.003144871 |
| Stx1b | 8383.06184 | 0.209784092 | 0.003021024 |
| Ica1 | 940.1693483 | 0.209099703 | 0.012260755 |
| Cdk12 | 894.3380466 | 0.207483101 | 0.019790375 |
| Srf | 1921.547894 | 0.207201282 | 0.027936472 |
| Dmtf1 | 2378.288853 | 0.207189649 | 0.00168695 |
| Arhgef9 | 6964.49217 | 0.206499064 | 0.001737079 |
| Nptn | 9318.334023 | 0.206474698 | 0.007179741 |
| Rps6ka2 | 2464.172646 | 0.206341476 | 0.047010595 |
| Fam168b | 9147.138477 | 0.20624083 | 1.39E-09 |
| Pja1 | 9136.067423 | 0.206193701 | 0.001088701 |
| Zfp286a | 227.5073138 | 0.205468716 | 0.039107224 |
| Ptk2b | 13028.04227 | 0.205185001 | 0.028642697 |
| Lrp12 | 1832.034417 | 0.204844557 | 0.029504912 |
| Gabrg3 | 659.8926841 | 0.20471686 | 0.02412873 |

| | | | |
|---|---|---|---|
| Srrm3 | 1790.181204 | 0.204640841 | 0.014828499 |
| Dbndd1 | 536.3296535 | 0.203261299 | 0.025959706 |
| Mesdc1 | 745.5546655 | 0.203252578 | 0.014396859 |
| Neurl1 | 612.5792736 | 0.201581923 | 0.036993966 |
| Map3k3 | 490.5464554 | 0.201355262 | 0.011468738 |
| Ncs1 | 5254.151603 | 0.201288584 | 0.018967014 |
| Tenm4 | 7630.404206 | 0.200913248 | 0.017707568 |
| Chchd4 | 474.1503501 | 0.200517754 | 0.011062592 |
| Tut1 | 706.0320679 | 0.200338705 | 0.029603008 |
| Ank3 | 15181.78252 | 0.199253368 | 0.025626751 |
| Rai1 | 4160.274879 | 0.199123805 | 0.007179741 |
| Ankrd10 | 266.565014 | 0.198790388 | 0.03935445 |
| Rgs7bp | 2857.56348 | 0.198683707 | 0.033332899 |
| Zbtb7a | 3444.651613 | 0.197629563 | 0.000709547 |
| Slc25a30 | 252.5693031 | 0.197338558 | 0.043944822 |
| Bbs4 | 1105.497574 | 0.196921119 | 0.015387293 |
| Mark2 | 2335.00794 | 0.1967548 | 0.027328328 |
| Dgki | 2053.73678 | 0.196380915 | 0.0305677 |
| Sgsm2 | 3507.214212 | 0.196020299 | 0.011431734 |
| Dcp1a | 492.8643484 | 0.195799386 | 0.024955868 |
| Tcf20 | 6921.630501 | 0.195742493 | 0.000249744 |
| Celf3 | 2039.185176 | 0.194492499 | 0.039751587 |
| Nxph1 | 2600.492781 | 0.193938063 | 0.021453203 |
| Pip4k2b | 2755.115582 | 0.193608097 | 0.01309335 |
| Shc2 | 985.4434376 | 0.19360059 | 0.039107224 |
| Dlg4 | 16820.21778 | 0.19357177 | 0.021290194 |
| Fbxo33 | 1183.44157 | 0.192764157 | 0.016545145 |
| Ip6k2 | 1478.081133 | 0.192755025 | 0.028642697 |
| Mnt | 2316.434191 | 0.192735468 | 0.013604395 |
| Camk2b | 21029.65565 | 0.192471796 | 0.044212852 |
| L1cam | 8751.112057 | 0.192254616 | 0.002405782 |
| Ankrd34a | 2740.757881 | 0.191905572 | 0.024732514 |
| Sh2b3 | 321.5724733 | 0.191588716 | 0.047709312 |
| Mink1 | 14447.24766 | 0.19145556 | 0.011416653 |
| Pdp1 | 1392.642657 | 0.191432783 | 0.03624213 |
| Crlf2 | 290.4499113 | 0.191332735 | 0.046052169 |

| | | | |
|---|---|---|---|
| Matk | 1460.228422 | 0.190851419 | 0.034346593 |
| Zfp354c | 391.8609962 | 0.190189626 | 0.026080098 |
| Dnajc5 | 2074.122483 | 0.190148966 | 0.002533331 |
| Slc4a3 | 4043.153247 | 0.190006256 | 0.024644826 |
| Cep290 | 1140.705095 | 0.189561976 | 0.04098609 |
| Ccdc92 | 2212.53027 | 0.189280199 | 0.013391727 |
| Evl | 3993.901136 | 0.189208737 | 0.004674719 |
| Tbc1d8 | 607.5096784 | 0.188894116 | 0.041420953 |
| Rps6ka5 | 1871.717325 | 0.188452307 | 0.035475756 |
| Cbx7 | 1507.068326 | 0.188178073 | 0.017125829 |
| Caly | 7849.407282 | 0.188086811 | 0.037495754 |
| Mgat4a | 1180.592533 | 0.187973976 | 0.02352241 |
| Jade2 | 2195.827539 | 0.187875611 | 0.003071915 |
| Plxna2 | 8165.894746 | 0.187849106 | 0.012151485 |
| Ndel1 | 2467.328782 | 0.187687466 | 0.001088701 |
| Kctd1 | 3060.634327 | 0.187510914 | 0.008941216 |
| Cbfa2t2 | 1163.04409 | 0.187288881 | 0.001238723 |
| Atmin | 3119.885191 | 0.187067817 | 0.012854347 |
| Ppm1h | 1966.501839 | 0.18597259 | 0.009593172 |
| Nalcn | 5223.705955 | 0.185898475 | 0.021453203 |
| Dip2c | 4345.832635 | 0.185670416 | 0.005389283 |
| Chgb | 13911.88822 | 0.185490458 | 0.024289931 |
| Nol4 | 1144.509154 | 0.185286065 | 0.033297082 |
| Ppp1r12c | 3266.782646 | 0.184608533 | 0.003021024 |
| Ppp1r12b | 6506.195403 | 0.184267044 | 0.027936472 |
| Ppp1r13b | 2576.640319 | 0.183486036 | 0.030149379 |
| Gga3 | 1575.674452 | 0.183419069 | 0.025222591 |
| Gaa | 18256.96993 | 0.182479982 | 0.015089783 |
| Gnb5 | 1051.514654 | 0.181762158 | 0.026142931 |
| Gpsm1 | 2993.49318 | 0.181760988 | 0.035533691 |
| Dmtn | 4865.598454 | 0.181306722 | 0.026427274 |
| Hsph1 | 11546.70461 | 0.179358039 | 0.02203306 |
| Trim46 | 1005.155031 | 0.179254567 | 0.047930523 |
| Scg2 | 11816.06306 | 0.178977689 | 0.040005788 |
| Tenm2 | 4299.394515 | 0.178896786 | 0.046195801 |
| Fam220a | 706.5654435 | 0.178319492 | 0.01904588 |

| | | | |
|---|---|---|---|
| Agk | 881.4728719 | 0.17826263 | 0.02597668 |
| Jade1 | 1471.015597 | 0.178055069 | 0.02316479 |
| Sin3b | 1979.746093 | 0.177258075 | 0.00042908 |
| Spire2 | 696.0995284 | 0.176939615 | 0.048946802 |
| Mdm4 | 388.5385661 | 0.176934599 | 0.035612549 |
| Nmnat2 | 1908.568147 | 0.176638146 | 0.024305403 |
| Lpin2 | 2847.483338 | 0.176475447 | 0.006792146 |
| Tomm20 | 3702.878053 | 0.176467385 | 0.02333945 |
| Ndufaf6 | 254.8883681 | 0.176265269 | 0.045313934 |
| Mecp2 | 1473.476539 | 0.175762967 | 0.0065778 |
| Nckipsd | 2005.403461 | 0.175726942 | 0.012309943 |
| Celf4 | 18132.53643 | 0.174767834 | 0.02316479 |
| Sbk1 | 2338.985186 | 0.174717755 | 0.030056562 |
| Camk1d | 4782.979929 | 0.174499077 | 0.013631117 |
| Prkar1b | 10222.57175 | 0.174129681 | 0.018391312 |
| Slmap | 3825.825202 | 0.17388315 | 0.006792146 |
| Mapre2 | 9439.167747 | 0.173781608 | 6.72E-06 |
| Pip5k1a | 1855.755668 | 0.17369441 | 0.047930523 |
| Ctdspl | 734.8566498 | 0.173075718 | 0.02808155 |
| Ppil4 | 1199.807059 | 0.173020811 | 0.048298394 |
| Tmem130 | 32804.92439 | 0.172897805 | 0.029649038 |
| Bex1 | 1147.813961 | 0.172840762 | 0.018967014 |
| Lancl1 | 2407.92001 | 0.172679187 | 0.036563165 |
| Dclk2 | 2649.174805 | 0.172436956 | 0.009961162 |
| Pnmal1 | 2369.014579 | 0.172213649 | 0.04447809 |
| Zfp638 | 2478.224753 | 0.170674615 | 0.029088595 |
| Celf1 | 1834.905733 | 0.170655958 | 0.001798516 |
| Chtf8 | 1974.61872 | 0.170273863 | 0.008512679 |
| Mtmr7 | 1233.610858 | 0.169851589 | 0.0270738 |
| Agfg2 | 1397.807004 | 0.169748175 | 0.01309335 |
| Stim2 | 2011.964373 | 0.169162054 | 0.026899107 |
| Syngr3 | 2091.785605 | 0.169053872 | 0.016535737 |
| Ppip5k1 | 2871.731212 | 0.168710413 | 0.021429355 |
| Abcg1 | 1954.147524 | 0.167591487 | 0.049853094 |
| Prrc2c | 13878.21044 | 0.167451688 | 0.012660145 |
| Dlg3 | 5057.363877 | 0.167143827 | 0.043768839 |

| | | | |
|---|---|---|---|
| Irs2 | 4728.010264 | 0.166834198 | 0.041763806 |
| Grip1 | 1499.894035 | 0.166525349 | 0.024868187 |
| Kcnd2 | 4122.699706 | 0.166480774 | 0.013158351 |
| Fbxo42 | 960.5441339 | 0.166473819 | 0.027680092 |
| Mbnl2 | 6966.370302 | 0.166393955 | 0.009935143 |
| Mmp24 | 1217.597792 | 0.165980645 | 0.022324352 |
| Fam13b | 4434.109404 | 0.165847771 | 0.006588153 |
| Fbxl15 | 458.12678 | 0.165682694 | 0.032937004 |
| Lgi1 | 1648.202492 | 0.165032394 | 0.045313934 |
| Zfp292 | 2904.150936 | 0.164936194 | 0.046820305 |
| Map3k12 | 2830.50464 | 0.164704303 | 0.026279332 |
| Osbpl8 | 5261.800456 | 0.164079907 | 0.021866318 |
| Rps6ka4 | 1477.033552 | 0.164021445 | 0.025702779 |
| Arhgef25 | 1063.305297 | 0.163475645 | 0.049767442 |
| Pbx2 | 1798.611706 | 0.163472267 | 0.017049106 |
| Micu3 | 1020.958317 | 0.161964231 | 0.046224399 |
| Atg13 | 2715.157921 | 0.161752685 | 0.007679935 |
| Sez6l2 | 13349.79941 | 0.161010423 | 0.026658536 |
| Cbx6 | 6305.286351 | 0.160981691 | 0.030035765 |
| Usp22 | 4504.22679 | 0.160910201 | 0.015326828 |
| Phf20 | 2171.250056 | 0.160560533 | 0.008957101 |
| Mapk1 | 16675.37837 | 0.158573482 | 0.04472828 |
| Gtpbp3 | 357.5655424 | 0.158270823 | 0.049853094 |
| Zcchc8 | 487.6388971 | 0.158177214 | 0.049934825 |
| Prpf3 | 688.8695947 | 0.157941741 | 0.038715746 |
| Zfp622 | 748.6935662 | 0.157664621 | 0.041763806 |
| Dzip1l | 1030.386089 | 0.157348393 | 0.040089072 |
| Stx16 | 1192.768717 | 0.156871549 | 0.009344149 |
| Nupl1 | 1312.679685 | 0.155743897 | 0.018133601 |
| Atg14 | 644.6439426 | 0.154332739 | 0.045329382 |
| Mepce | 1225.656741 | 0.154195588 | 0.006074981 |
| Srsf2 | 3274.423548 | 0.153036662 | 0.01430933 |
| Ctnnbip1 | 1021.115448 | 0.152961574 | 0.040005788 |
| Pdzd4 | 2507.702254 | 0.152939267 | 0.038627667 |
| Cep120 | 1318.377684 | 0.152896359 | 0.041541729 |
| Abhd8 | 5275.993168 | 0.152856487 | 0.01783629 |

| | | | |
|---|---|---|---|
| Chpf | 4575.372284 | 0.151836152 | 0.000784596 |
| Bmyc | 726.1679568 | 0.151549476 | 0.047748655 |
| Rc3h2 | 3595.654485 | 0.151370187 | 0.005468026 |
| Usp11 | 6778.010172 | 0.151020375 | 0.041763806 |
| Tulp4 | 7763.649447 | 0.150840001 | 0.032024688 |
| Pja2 | 29734.47418 | 0.150392556 | 0.009935143 |
| Hipk3 | 3536.255591 | 0.149987625 | 0.040743144 |
| Slitrk5 | 2331.857335 | 0.149569424 | 0.045633405 |
| Prepl | 10521.80452 | 0.149467173 | 0.040524536 |
| Dynll2 | 13908.28363 | 0.148154981 | 0.026187002 |
| Hdac11 | 6621.141168 | 0.147994986 | 0.040005788 |
| Mrpl38 | 1257.86055 | 0.147608642 | 0.04472828 |
| Dyrk1a | 2908.854535 | 0.147510549 | 0.004230621 |
| Ezh1 | 2374.402769 | 0.147120499 | 0.023064351 |
| Prkca | 5667.237106 | 0.14695902 | 0.028642697 |
| Crebl2 | 2137.485398 | 0.146735203 | 0.030437403 |
| Epc1 | 1241.176016 | 0.146292864 | 0.039895654 |
| Otud4 | 1254.630034 | 0.146137214 | 0.04047026 |
| Adar | 5951.566023 | 0.145863719 | 0.024305403 |
| Aqr | 2141.869857 | 0.145596897 | 0.021869478 |
| Armcx3 | 3676.927785 | 0.145239162 | 0.005892714 |
| Rnf185 | 1276.193004 | 0.144410628 | 0.023004125 |
| Ptprm | 3113.66856 | 0.14399977 | 0.032538953 |
| Alcam | 7072.802246 | 0.14393825 | 0.028687486 |
| Pom121 | 5577.033296 | 0.143841053 | 0.043598327 |
| Ubqln2 | 13857.60675 | 0.143775942 | 0.001407415 |
| Nr3c2 | 1986.769389 | 0.143712056 | 0.02203306 |
| Slk | 3971.984531 | 0.142887126 | 0.027771811 |
| Larp6 | 1087.359758 | 0.142687529 | 0.041420953 |
| Elmsan1 | 1213.448231 | 0.142157458 | 0.046820305 |
| Mau2 | 2573.308522 | 0.140557154 | 0.037753985 |
| Dpysl5 | 4174.656992 | 0.139982155 | 0.046820305 |
| Cds2 | 4810.936702 | 0.139958585 | 0.009851005 |
| Armc8 | 1708.899674 | 0.139294009 | 0.03624213 |
| Tmem57 | 2642.705532 | 0.139045611 | 0.02850463 |
| Mib2 | 4036.549965 | 0.13900159 | 0.016395152 |

| | | | |
|---|---|---|---|
| Spin1 | 1932.270096 | 0.13895948 | 0.040230345 |
| Aarsd1 | 2085.014203 | 0.138504873 | 0.004710761 |
| Stim1 | 3288.039456 | 0.137545598 | 0.03781136 |
| Tspyl4 | 8974.287634 | 0.13643305 | 0.037364468 |
| Rufy3 | 4635.526721 | 0.136416916 | 0.031259999 |
| Stat5b | 2004.548765 | 0.135948698 | 0.049682839 |
| Stk24 | 1037.151783 | 0.135354724 | 0.033656689 |
| Matr3 | 16284.76894 | 0.135126344 | 0.007472342 |
| Ubp1 | 1027.415369 | 0.134662265 | 0.049853094 |
| Bag5 | 1654.436452 | 0.134659612 | 0.016899408 |
| Sf3a1 | 4105.701104 | 0.134168394 | 0.030149379 |
| Brd4 | 8153.925798 | 0.133021349 | 0.013604395 |
| Nelfa | 977.975778 | 0.132247684 | 0.049185358 |
| Maged1 | 13779.45804 | 0.13190147 | 0.028658911 |
| Ccnk | 1542.397742 | 0.131794717 | 0.025318371 |
| Rnf111 | 2164.474281 | 0.131255845 | 0.018133601 |
| Sf1 | 6851.896678 | 0.13036729 | 0.019595701 |
| Pds5b | 7749.411411 | 0.130046899 | 0.035256586 |
| Akap17a | 2616.96547 | 0.129223351 | 0.00792777 |
| Pogz | 3284.329339 | 0.128047637 | 0.046820305 |
| Btrc | 1986.882995 | 0.127783509 | 0.047930523 |
| Zfp644 | 3055.808838 | 0.126765245 | 0.029088595 |
| Med24 | 2105.563261 | 0.126345441 | 0.021294141 |
| Ddi2 | 2000.288157 | 0.125800363 | 0.049853094 |
| Ldb1 | 2096.664093 | 0.124338734 | 0.011431734 |
| Hmg20a | 2599.951049 | 0.122935971 | 0.03977682 |
| Taok2 | 4290.235812 | 0.122715993 | 0.026883646 |
| Rnf44 | 2004.359883 | 0.122657027 | 0.043768839 |
| Zfr | 14066.33323 | 0.122387753 | 0.012692539 |
| Bcl7b | 1544.829189 | 0.121428422 | 0.04098609 |
| Gltscr1l | 1910.548889 | 0.120370055 | 0.044284634 |
| Rnf187 | 12793.54993 | 0.119986049 | 0.017741036 |
| Tubg1 | 1684.927262 | 0.119190178 | 0.047280998 |
| Mtch1 | 13577.32724 | 0.117990773 | 0.033797488 |
| Zfp532 | 1956.998039 | 0.117865157 | 0.013604395 |
| Camsap2 | 11571.44825 | 0.116991129 | 0.016578751 |

| | | | |
|---|---|---|---|
| Scamp1 | 6589.517321 | 0.115450089 | 0.032657682 |
| Cct6a | 3555.054061 | 0.114406997 | 0.043811263 |
| Nolc1 | 6455.09137 | 0.112973538 | 0.005256768 |
| Xpo7 | 4332.666928 | 0.11247702 | 0.028160868 |
| Khdrbs1 | 3973.905271 | 0.111608396 | 0.025285931 |
| Slc22a17 | 25239.06431 | 0.109993262 | 0.049853094 |
| Rxrb | 1567.469546 | 0.109905695 | 0.026847461 |
| Stox2 | 6726.187263 | 0.109434709 | 0.040267915 |
| Grb2 | 6518.269622 | 0.109111185 | 0.012584608 |
| Wbp11 | 3467.493049 | 0.108541023 | 0.036563165 |
| Rabl6 | 6030.84249 | 0.108289918 | 0.021102121 |
| Plekhb2 | 13104.00931 | 0.107806228 | 0.049049073 |
| Ddx5 | 14448.11159 | 0.095609127 | 0.029088595 |
| Ipo9 | 3313.573676 | 0.092426294 | 0.049853094 |
| Akr1a1 | 6422.395649 | 0.091874698 | 0.037426345 |
| Klhl24 | 4579.91487 | -0.089965935 | 0.045313934 |
| Abi1 | 4314.037623 | -0.101149534 | 0.043156126 |
| Flot2 | 4646.83651 | -0.115845672 | 0.045241927 |
| Adam10 | 3750.07747 | -0.116723126 | 0.049139611 |
| Tspan3 | 15260.59565 | -0.117785746 | 0.027680092 |
| Arl2bp | 3787.176864 | -0.118985462 | 0.005269885 |
| Coasy | 1048.966328 | -0.119501919 | 0.049853094 |
| Rab5c | 7771.99537 | -0.121309325 | 0.045980518 |
| Cadm4 | 7016.37041 | -0.122294403 | 0.019790375 |
| Rpl19 | 7575.504741 | -0.124246943 | 0.046898757 |
| Tmem14c | 1311.94197 | -0.124652673 | 0.039563908 |
| Tex264 | 2004.399405 | -0.127716354 | 0.015387293 |
| Spg20 | 2104.050382 | -0.129502513 | 0.043295222 |
| Eif2s3y | 1560.018451 | -0.131212082 | 0.034728001 |
| Fam134c | 1441.850204 | -0.132705307 | 0.030332396 |
| Cttnbp2nl | 1334.221682 | -0.132715227 | 0.043581726 |
| Mettl9 | 1578.121993 | -0.133787936 | 0.036608462 |
| Ndufb2 | 2172.034462 | -0.134246139 | 0.037495754 |
| Snap29 | 1780.76125 | -0.134993761 | 0.036116276 |
| Mfsd1 | 1849.026298 | -0.135598104 | 0.047378151 |
| Comt | 1540.132458 | -0.135631154 | 0.023697231 |

| | | | |
|---|---|---|---|
| Odc1 | 2774.941562 | -0.137725831 | 0.039895654 |
| Baalc | 6653.292689 | -0.141145579 | 0.030239799 |
| Arhgap5 | 13637.60361 | -0.141979005 | 0.04098609 |
| Dusp3 | 8376.058987 | -0.142549032 | 0.027928671 |
| Ccs | 1037.016313 | -0.143751136 | 0.041392312 |
| Slc48a1 | 5591.874872 | -0.14456952 | 0.044540338 |
| Pon2 | 6350.686277 | -0.145532466 | 0.044540338 |
| Ndst1 | 4286.743053 | -0.145694355 | 0.015861679 |
| Ifnar1 | 3016.082334 | -0.14625134 | 0.008444339 |
| Ybx1 | 3535.160113 | -0.146334404 | 0.032921965 |
| Sgcb | 6830.693206 | -0.146666939 | 0.016314268 |
| Ccni | 3396.925246 | -0.147802874 | 0.013233116 |
| Sp1 | 2723.582459 | -0.148972925 | 0.047049688 |
| Lzic | 718.4871488 | -0.149420327 | 0.037470055 |
| Ccdc90b | 628.8626393 | -0.149884599 | 0.047748655 |
| Scpep1 | 1094.446853 | -0.150378242 | 0.036673106 |
| Rasa3 | 2911.141844 | -0.151998593 | 0.029088595 |
| Lhpp | 3260.215485 | -0.152390237 | 0.036893303 |
| Nav2 | 3215.412591 | -0.152437271 | 0.046898757 |
| Wbp1l | 2397.554511 | -0.152883841 | 0.024247381 |
| Dennd5a | 20085.25863 | -0.153114187 | 0.006424206 |
| Nadk | 2921.741327 | -0.153843145 | 0.023919501 |
| Glo1 | 3036.000025 | -0.153996932 | 0.007179741 |
| March8 | 2014.943587 | -0.154209326 | 0.047748655 |
| Pias3 | 985.8373867 | -0.154426703 | 0.041551339 |
| Ppp2r5a | 3418.010306 | -0.15511467 | 0.024732514 |
| Fgfr2 | 9387.871523 | -0.158545045 | 0.037790755 |
| RT1-Bb | 7.413816749 | -0.159668291 | 0.03624213 |
| LOC306766 | 3888.754214 | -0.159755107 | 0.005081827 |
| Pmm1 | 5771.303837 | -0.161217939 | 0.02316479 |
| Hsd17b4 | 4036.144422 | -0.161251005 | 0.047748655 |
| Idh1 | 1984.405879 | -0.161767506 | 0.046820305 |
| Omg | 8044.351917 | -0.162068875 | 0.004338437 |
| Gca | 1313.034436 | -0.162092735 | 0.042720483 |
| Anxa5 | 4185.059511 | -0.162706896 | 0.047025702 |
| Fasn | 23585.99901 | -0.163027516 | 0.03624213 |

| | | | |
|---|---|---|---|
| Pgap2 | 450.9577825 | -0.163179549 | 0.041420953 |
| Sdhc | 5251.041388 | -0.16442523 | 0.012007103 |
| Pibf1 | 409.3091221 | -0.164762168 | 0.046446029 |
| Szrd1 | 2400.305527 | -0.165712618 | 0.021451971 |
| Cmtm5 | 2424.656975 | -0.165755464 | 0.040685122 |
| RGD1308428 | 1128.344699 | -0.166946468 | 0.025524355 |
| Pomt1 | 980.5948492 | -0.168041618 | 0.041763806 |
| Ctnna1 | 7844.7921 | -0.16871558 | 0.048946802 |
| RGD1304587 | 644.9769766 | -0.169662056 | 0.026296383 |
| Ntrk2 | 100591.7334 | -0.169946131 | 0.028160868 |
| Rgma | 12291.19053 | -0.171103243 | 0.030247411 |
| Golga7 | 2052.268562 | -0.172273235 | 0.016370996 |
| LOC691807 | 2831.431651 | -0.173518769 | 0.015664411 |
| Daglb | 991.1638364 | -0.173640985 | 0.025675565 |
| Tspan6 | 1042.567413 | -0.17404308 | 0.035174782 |
| Gpm6b | 34619.7633 | -0.174191676 | 0.034431659 |
| Suclg2 | 5934.718522 | -0.174769123 | 0.049853094 |
| Hadh | 1846.911824 | -0.176366578 | 0.047010595 |
| Sptlc1 | 1604.675755 | -0.177464034 | 0.004502516 |
| Rap1a | 1368.419439 | -0.177656144 | 0.013406415 |
| Pnpla2 | 1695.973968 | -0.177734619 | 0.040780546 |
| Ddah1 | 6562.746892 | -0.177931279 | 0.007343924 |
| Nsmce1 | 593.3685298 | -0.179009457 | 0.037710881 |
| Dcakd | 2747.875338 | -0.180838944 | 0.049853094 |
| Mlc1 | 47147.37233 | -0.181401505 | 0.037495754 |
| Nudcd2 | 732.855236 | -0.182187877 | 0.035405596 |
| Slc7a5 | 2968.883137 | -0.182457257 | 0.0305677 |
| Slc27a1 | 12414.14315 | -0.182719863 | 0.040005788 |
| Slc35b2 | 882.4598592 | -0.183700688 | 0.019341006 |
| Gpam | 10861.18577 | -0.184539995 | 0.041420953 |
| Pccb | 1198.959005 | -0.184666372 | 0.024441419 |
| Ssfa2 | 4936.849146 | -0.18488689 | 0.026051468 |
| Ubr7 | 2509.744331 | -0.185046798 | 0.032017326 |
| Fuca1 | 1800.955774 | -0.18563109 | 0.02734063 |
| S100a13 | 2051.88377 | -0.18590522 | 0.043790963 |
| Ptprz1 | 32744.90204 | -0.185981354 | 0.030243022 |

| | | | |
|---|---|---|---|
| Necap2 | 1251.678027 | -0.186261608 | 0.031925301 |
| Shc1 | 453.0227992 | -0.186566183 | 0.033855648 |
| Cc2d1b | 1016.815401 | -0.186794759 | 0.010651208 |
| Ctnnd1 | 3929.051393 | -0.186969883 | 0.01163911 |
| Idi1 | 1129.246905 | -0.187524313 | 0.047378151 |
| Krcc1 | 869.1659847 | -0.189329245 | 0.032340128 |
| Pir | 898.6893593 | -0.189958898 | 0.02677863 |
| Tp53inp2 | 6582.197961 | -0.189964977 | 0.037495754 |
| Wscd1 | 3782.178407 | -0.191379542 | 0.043645727 |
| Cd302 | 848.196751 | -0.1915799 | 0.045538379 |
| Chmp1b | 1297.225055 | -0.191826134 | 0.006260517 |
| Dnase2 | 1006.967241 | -0.192119261 | 0.037426345 |
| Plcd1 | 1996.567441 | -0.192540342 | 0.041986704 |
| Peli2 | 2010.813333 | -0.193904216 | 0.030906475 |
| Pttg1ip | 5237.679594 | -0.194547667 | 0.011431734 |
| Tvp23b | 1827.31169 | -0.195144809 | 0.026723413 |
| Smo | 1016.472986 | -0.195853699 | 0.035669741 |
| Fam63a | 590.8465173 | -0.196456814 | 0.027680092 |
| Nek9 | 3001.769741 | -0.196608531 | 0.005197913 |
| Nln | 1827.248646 | -0.196936582 | 0.002361064 |
| Npc2 | 2081.004334 | -0.196953932 | 0.020905332 |
| Tex261 | 590.0947977 | -0.196959717 | 0.021506687 |
| Me1 | 3112.196151 | -0.197008975 | 0.003961308 |
| Amotl2 | 1716.783038 | -0.1970835 | 0.035755922 |
| Ppp2r1b | 464.4559077 | -0.197408815 | 0.025245651 |
| Emc2 | 2336.03553 | -0.197628619 | 0.000269768 |
| Fam107a | 70795.49882 | -0.197902765 | 0.025277544 |
| Sox9 | 17051.75019 | -0.19792846 | 0.025490898 |
| Fabp5 | 2548.396952 | -0.198413413 | 0.026279332 |
| Zfp703 | 3770.230301 | -0.198629515 | 0.014512246 |
| Nfatc1 | 884.7850598 | -0.1987035 | 0.041457518 |
| Hip1 | 2762.791326 | -0.198840124 | 0.043768839 |
| Bax | 594.908267 | -0.199076167 | 0.043598327 |
| Nadk2 | 4703.038573 | -0.200015464 | 0.020164209 |
| Arhgef19 | 1839.71245 | -0.200084678 | 0.036608462 |
| Egln3 | 967.401963 | -0.20059189 | 0.046898757 |

| | | | |
|---|---|---|---|
| Fkbp9 | 1490.992205 | -0.200844662 | 0.035076854 |
| Irf2 | 459.5162649 | -0.200871637 | 0.025222591 |
| Acss2 | 2500.707362 | -0.201323618 | 0.038627667 |
| Tspan14 | 479.6866279 | -0.201325806 | 0.043811263 |
| Sirt2 | 5950.083223 | -0.201907621 | 0.003142987 |
| Myl12a | 832.4890543 | -0.202688874 | 0.021804087 |
| Nfia | 5332.166331 | -0.20338294 | 0.007081559 |
| Pdgfc | 614.8920782 | -0.203433668 | 0.041863225 |
| Sox4 | 1301.097899 | -0.203525579 | 0.026427274 |
| Scamp2 | 793.9273297 | -0.204005563 | 0.023919501 |
| Insig1 | 2761.309109 | -0.204317745 | 0.047748655 |
| Slc12a2 | 3683.255614 | -0.205897886 | 0.026128501 |
| Stx17 | 288.6458202 | -0.20594645 | 0.040267915 |
| Kif13a | 1737.486241 | -0.206319132 | 0.018679046 |
| Sh3bgrl | 7003.245083 | -0.207113902 | 0.005354322 |
| Leprot | 941.9138989 | -0.207282555 | 0.01624203 |
| Gng5 | 627.346271 | -0.207705555 | 0.019705203 |
| Fnbp1 | 5184.67989 | -0.208274784 | 0.015711943 |
| Smim15 | 1106.944431 | -0.208414509 | 0.004502516 |
| Slc13a3 | 822.2823919 | -0.210029368 | 0.041420953 |
| Nop9 | 356.8538944 | -0.210168157 | 0.012450893 |
| Gcdh | 2185.032535 | -0.210625649 | 0.005293068 |
| Scd2 | 346650.6894 | -0.211248625 | 0.018941502 |
| Prelp | 3249.869636 | -0.211485029 | 0.030239799 |
| Ppp1r3c | 6445.630784 | -0.211822783 | 0.025626751 |
| Hey2 | 1129.808021 | -0.212379589 | 0.026296383 |
| Rb1 | 2004.763292 | -0.212566828 | 0.000660142 |
| Lamp2 | 5006.670922 | -0.212717139 | 0.013391727 |
| Itgb5 | 4287.66776 | -0.214265694 | 0.023919501 |
| Gpr37 | 4629.437518 | -0.214277634 | 0.030239799 |
| Oplah | 1669.965824 | -0.214357633 | 0.018391312 |
| Steap3 | 1249.043917 | -0.21484181 | 0.033218991 |
| Rnaset2 | 844.834176 | -0.21656363 | 0.025971012 |
| Hspa2 | 690.995404 | -0.217277384 | 0.028173129 |
| Cyp20a1 | 781.8574386 | -0.217428126 | 0.023249185 |
| Fgfr1 | 5969.190722 | -0.217812728 | 0.00998537 |

| | | | |
|---|---|---|---|
| Glis2 | 659.1759463 | -0.21795878 | 0.024305403 |
| Snx33 | 837.3913042 | -0.218346451 | 0.024289931 |
| Traf7 | 1874.442683 | -0.218856667 | 0.011665085 |
| Spsb2 | 316.9837448 | -0.21891381 | 0.024666609 |
| Lpcat3 | 1169.410963 | -0.219787694 | 0.021102121 |
| Fkbp10 | 601.5341517 | -0.21991688 | 0.041392312 |
| Bphl | 260.741182 | -0.220773075 | 0.033332899 |
| Rsu1 | 1431.143086 | -0.221671681 | 0.016794415 |
| Zbtb7b | 426.4755902 | -0.221895087 | 0.027807083 |
| Fzd10 | 824.192127 | -0.222607647 | 0.049468179 |
| Akt2 | 1869.074763 | -0.223225478 | 0.016176111 |
| Mapre1 | 3120.842357 | -0.223632084 | 0.001238723 |
| Plekhg1 | 1090.489254 | -0.223765901 | 0.023657879 |
| Lhx2 | 4273.862141 | -0.223768887 | 0.014280117 |
| Hipk2 | 11027.6631 | -0.223836023 | 0.003299244 |
| Cd24 | 4797.038552 | -0.224300304 | 0.021109252 |
| Pbxip1 | 4816.096643 | -0.225171749 | 0.025524355 |
| Lace1 | 262.3661367 | -0.225293003 | 0.021804087 |
| Hyal1 | 952.2654872 | -0.225345782 | 0.033218991 |
| Rnpepl1 | 760.0185604 | -0.225522341 | 0.015877077 |
| Ankrd40 | 5654.286264 | -0.225928428 | 0.000154816 |
| Mvp | 804.9140651 | -0.227209686 | 0.039089228 |
| Efemp2 | 262.3415731 | -0.227691453 | 0.042260289 |
| Sepp1 | 23589.14999 | -0.229231661 | 0.005915258 |
| Cers4 | 4771.910983 | -0.229672189 | 0.003117664 |
| Slc38a3 | 6305.619903 | -0.229774849 | 0.026427274 |
| Tlcd1 | 777.4757024 | -0.230083016 | 0.034748924 |
| Efs | 1254.263748 | -0.231651691 | 0.015521472 |
| St3gal4 | 1796.28603 | -0.231753244 | 0.005197913 |
| Tnfsf13 | 768.3705644 | -0.231848332 | 0.044457556 |
| Ostf1 | 613.4687068 | -0.231974016 | 0.028642697 |
| Ost4 | 1178.679179 | -0.232060291 | 0.005854913 |
| Nqo1 | 826.0860032 | -0.232472894 | 0.014828499 |
| Cd151 | 1999.025687 | -0.232580262 | 0.005551777 |
| Trim5 | 552.5492326 | -0.233285608 | 0.031455808 |
| Mmgt2 | 454.7329198 | -0.233714047 | 0.032603398 |

| | | | |
|---|---|---|---|
| Pdzrn3 | 3452.549846 | -0.234481409 | 0.008562766 |
| Tspan12 | 4150.498507 | -0.23588085 | 0.00169016 |
| Gas6 | 6808.805433 | -0.235994619 | 0.00768371 |
| Phyhd1 | 791.6651055 | -0.236783998 | 0.040053617 |
| Slc12a4 | 1195.692404 | -0.236978947 | 0.021109252 |
| Fgfbp3 | 459.8629817 | -0.237079982 | 0.040005788 |
| Hist1h4b | 480.3773093 | -0.237921569 | 0.021840342 |
| Dclre1a | 542.8106985 | -0.238510799 | 0.006986576 |
| Adprhl2 | 1584.571898 | -0.238564231 | 0.001219453 |
| Prtfdc1 | 1360.622003 | -0.239150907 | 0.010257155 |
| Glb1l | 560.1058979 | -0.240353635 | 0.023919501 |
| Ddt | 2936.373952 | -0.241422499 | 0.000551695 |
| Adam17 | 1167.5571 | -0.241904614 | 0.015003021 |
| Pxn | 666.4978772 | -0.242566086 | 0.008612054 |
| Ddr1 | 5222.758705 | -0.242921888 | 0.006737939 |
| Ddx58 | 792.9196473 | -0.243956948 | 0.043768839 |
| Stat3 | 4932.472732 | -0.244045191 | 0.017372677 |
| Fdft1 | 4094.899908 | -0.244174488 | 0.004797976 |
| Tnfsf12 | 665.0546806 | -0.244247956 | 0.009935143 |
| Rela | 1125.469064 | -0.244972137 | 0.004812428 |
| Ctf1 | 275.1187479 | -0.245383387 | 0.037826755 |
| Aldh4a1 | 2839.120006 | -0.245855023 | 0.016265375 |
| Dusp15 | 909.9039557 | -0.246555088 | 0.004230621 |
| Scarb2 | 14539.05615 | -0.247203428 | 0.002996508 |
| Rab31 | 4565.230189 | -0.247776726 | 0.001187652 |
| Hexa | 1672.008087 | -0.247925115 | 0.000891074 |
| Hmgcs1 | 18167.35163 | -0.248440643 | 0.018439386 |
| Acyp1 | 296.7185585 | -0.248445312 | 0.016111417 |
| Npr1 | 518.3472762 | -0.249654818 | 0.041763806 |
| Gna12 | 5740.610811 | -0.24987905 | 0.002808251 |
| Faim | 318.8494674 | -0.250836612 | 0.024819902 |
| Mettl25 | 201.0111086 | -0.251147353 | 0.026427274 |
| Kctd15 | 858.3104509 | -0.253813966 | 0.003091319 |
| Hps1 | 329.1839035 | -0.254307434 | 0.022593647 |
| Sqle | 1987.925876 | -0.254705004 | 0.004639876 |
| Prkcq | 345.2731549 | -0.255311619 | 0.030701628 |

| | | | |
|---|---|---|---|
| Cyp51 | 10907.34693 | -0.255314442 | 0.013604395 |
| Wipi1 | 1756.628596 | -0.25558541 | 0.002925922 |
| Syngr2 | 301.8311734 | -0.256307551 | 0.028642697 |
| Eif4ebp1 | 374.9106966 | -0.256764196 | 0.019896181 |
| Cd81 | 35875.91538 | -0.259106759 | 0.002559176 |
| Ccl7 | 8.096424466 | -0.262491438 | 0.042572538 |
| Cst3 | 77756.20775 | -0.262812266 | 0.003886942 |
| Erbb2 | 414.9434148 | -0.263006528 | 0.019438382 |
| Mt3 | 12892.89868 | -0.263044044 | 0.00059525 |
| Nod1 | 659.5054084 | -0.264785043 | 0.018441434 |
| Acot1 | 91.84113092 | -0.265142806 | 0.048946802 |
| Htra3 | 351.8561831 | -0.265231546 | 0.041392312 |
| Map4k4 | 7280.255639 | -0.265905232 | 0.002167426 |
| Vwa5a | 1136.963632 | -0.266572888 | 0.018391312 |
| B3gnt7 | 82.84139934 | -0.26768881 | 0.048946802 |
| Sardh | 1477.656285 | -0.268185797 | 0.010084317 |
| Gab1 | 2908.948192 | -0.268367355 | 0.00897419 |
| Nkx2-2 | 248.3163541 | -0.268775822 | 0.03624213 |
| Ppap2a | 742.9139466 | -0.269339466 | 0.005492063 |
| Pik3ip1 | 1126.463336 | -0.269549566 | 0.007861133 |
| LOC497899 | 62.67660692 | -0.270852694 | 0.037677064 |
| Eif2ak2 | 1866.312346 | -0.27093278 | 0.010114653 |
| Tp53 | 881.858575 | -0.272260639 | 0.001327326 |
| Nt5dc2 | 817.1926577 | -0.272583203 | 0.008512679 |
| Vimp | 1411.182123 | -0.27305551 | 0.000503609 |
| Abcd1 | 692.3068878 | -0.273987986 | 0.000978906 |
| Mocos | 53.68415742 | -0.27430851 | 0.047748655 |
| Snapc2 | 1872.280622 | -0.275269002 | 0.002183788 |
| Tnfrsf19 | 1786.143001 | -0.276873599 | 0.006178546 |
| Adamtsl4 | 96.10590708 | -0.278502955 | 0.046357563 |
| Hs3st3b1 | 93.46065893 | -0.278721254 | 0.047303365 |
| Pdlim4 | 6135.242103 | -0.279148453 | 0.011062592 |
| Gsta1 | 11748.85822 | -0.279288781 | 0.005290761 |
| Etv4 | 152.8723165 | -0.280055483 | 0.046614855 |
| Vamp3 | 1693.123414 | -0.280505003 | 0.001253554 |
| Mpzl1 | 676.1786884 | -0.280703812 | 0.001479987 |

| | | | |
|---|---|---|---|
| Rhog | 570.6699713 | -0.281254812 | 0.025340738 |
| Il10rb | 123.9957083 | -0.281662141 | 0.028841436 |
| Ltbr | 1277.382552 | -0.281721871 | 0.006436607 |
| Tmem209 | 417.5581591 | -0.282442674 | 0.005877535 |
| Nes | 292.329148 | -0.283265388 | 0.028173129 |
| Psmb8 | 482.2250051 | -0.2842477 | 0.044957874 |
| Pros1 | 502.8959342 | -0.284616963 | 0.016427633 |
| Evi2a | 333.3219161 | -0.284733405 | 0.045538379 |
| Fzd9 | 1309.28206 | -0.285018754 | 0.007798981 |
| Nfe2l2 | 2151.335637 | -0.285385821 | 0.003021024 |
| Bub1b | 16.80370171 | -0.286762395 | 0.046898757 |
| Igtp | 521.9803348 | -0.288545656 | 0.046898757 |
| Psmb9 | 359.767857 | -0.289268346 | 0.045313934 |
| Ctsh | 1916.217058 | -0.289320454 | 0.004154241 |
| Tmem256 | 322.0077223 | -0.289833027 | 0.01217504 |
| Slc14a1 | 2342.8251 | -0.290897084 | 0.005293068 |
| Ldb3 | 63.36332626 | -0.291397303 | 0.040960018 |
| Fa2h | 272.2415409 | -0.291693184 | 0.046224399 |
| Tubb2b | 10383.66966 | -0.291781051 | 0.003192776 |
| RT1-M3-1 | 637.7112478 | -0.292452773 | 0.002036429 |
| Itgal | 18.37968615 | -0.293192337 | 0.049185358 |
| Dapk2 | 44.47909999 | -0.294042628 | 0.043156126 |
| Carhsp1 | 1504.833419 | -0.294905935 | 0.011128752 |
| Hapln3 | 85.79036335 | -0.295654482 | 0.040041859 |
| Mcee | 756.586444 | -0.296414584 | 0.001444994 |
| Plekhd1 | 1550.335482 | -0.296667678 | 0.015551272 |
| Tp53i3 | 256.4649932 | -0.296721554 | 0.011330147 |
| Irf9 | 419.3603313 | -0.297554932 | 0.036116276 |
| Mgmt | 775.9208661 | -0.297991121 | 0.009935143 |
| Fabp7 | 2244.127471 | -0.298019304 | 0.037495754 |
| Adamts1 | 215.9494407 | -0.298341552 | 0.033147482 |
| Nlrp1a | 13.85000409 | -0.298625781 | 0.048946802 |
| Rras2 | 606.4352286 | -0.298813284 | 0.003986341 |
| Ldlr | 1253.280841 | -0.29991521 | 0.009315766 |
| Npepl1 | 633.7396604 | -0.303361289 | 0.002561742 |
| Dock6 | 161.1902934 | -0.30458507 | 0.018967014 |

| | | | |
|---|---|---|---|
| Csrp1 | 12173.6077 | -0.304779449 | 0.001187652 |
| Sh3bp4 | 963.0744342 | -0.305457591 | 0.008882668 |
| Cfb | 136.6528607 | -0.30626891 | 0.049267125 |
| Opalin | 623.527176 | -0.306841085 | 0.04091115 |
| Ctdsp1 | 1791.493611 | -0.307792889 | 0.001700173 |
| Nuf2 | 33.06244867 | -0.308031375 | 0.036460639 |
| Bst2 | 82.70013448 | -0.308157433 | 0.046898757 |
| Lrp10 | 2675.843689 | -0.310187062 | 0.005389283 |
| Antxr1 | 375.6356454 | -0.312926132 | 0.029627131 |
| Prr11 | 16.69694343 | -0.314192718 | 0.041420953 |
| Slc50a1 | 232.8387899 | -0.315187272 | 0.014011967 |
| RGD1311756 | 661.6110016 | -0.315477313 | 0.00652612 |
| Clec2g | 13.72667958 | -0.316418168 | 0.047720469 |
| Gpr37l1 | 28173.26571 | -0.319707482 | 0.001728876 |
| Aamdc | 206.4935803 | -0.320084482 | 0.009593172 |
| Lpar1 | 479.3004865 | -0.320170112 | 0.035426139 |
| Spsb1 | 487.6551728 | -0.321850043 | 0.008612054 |
| Sh3glb1 | 2584.293992 | -0.322984248 | 9.16E-05 |
| Atg16l2 | 114.3149779 | -0.324563088 | 0.024305403 |
| RGD1561157 | 58.39030811 | -0.326874003 | 0.033783071 |
| Irgm | 527.7767198 | -0.32698087 | 0.035174782 |
| Gng11 | 229.3665698 | -0.327429495 | 0.023116594 |
| Aen | 1278.027665 | -0.328259277 | 0.005664875 |
| Btbd7 | 1218.286243 | -0.328532209 | 0.000700845 |
| Slc39a1 | 3969.692471 | -0.32904927 | 0.000813073 |
| Pmp22 | 739.851665 | -0.329219939 | 0.014798387 |
| Litaf | 888.0556787 | -0.330239932 | 0.008834317 |
| Plekha4 | 62.4572659 | -0.331199547 | 0.034620038 |
| Cd180 | 30.20865525 | -0.331902471 | 0.03624213 |
| Sh3bgr | 248.6095746 | -0.332006242 | 0.014354793 |
| Espl1 | 28.71449089 | -0.332728199 | 0.03889564 |
| Hn1l | 222.4187172 | -0.334160646 | 0.015267639 |
| Pla2g16 | 2463.479733 | -0.335045138 | 0.001187652 |
| Loxl4 | 285.9576425 | -0.336362782 | 0.025846377 |
| Cx3cr1 | 49.90448384 | -0.337150444 | 0.033856222 |
| Arpc1b | 81.07301692 | -0.337152289 | 0.031259999 |

| | | | |
|---|---|---|---|
| Hrsp12 | 1921.524766 | -0.338893141 | 0.002612321 |
| RT1-Db1 | 39.67182752 | -0.339850382 | 0.037495754 |
| Ugt1a6 | 153.1697023 | -0.340878858 | 0.02734063 |
| Kif1c | 4192.612331 | -0.341122015 | 0.000103699 |
| Fxyd1 | 2698.905565 | -0.343124577 | 0.004107635 |
| Cd63 | 2704.85794 | -0.343875112 | 0.000206593 |
| Lgi4 | 8258.406055 | -0.344774255 | 0.000249744 |
| Irf1 | 472.3585257 | -0.346001767 | 0.032183586 |
| Sspn | 2183.601902 | -0.346148613 | 0.000255979 |
| Ninj2 | 26.8239543 | -0.347350478 | 0.040148834 |
| Nkd1 | 948.5821572 | -0.347950633 | 0.005197913 |
| Galnt6 | 140.0366627 | -0.348400164 | 0.033351065 |
| Art3 | 320.5656015 | -0.349967682 | 0.030939193 |
| Itpkb | 3044.175357 | -0.350489718 | 0.001208891 |
| Ptpn6 | 22.14698247 | -0.351057454 | 0.031911265 |
| Axl | 3179.484088 | -0.353133244 | 0.002046554 |
| LOC100909539 | 37.75285804 | -0.356550453 | 0.027680092 |
| Ssc5d | 428.3819873 | -0.358968973 | 0.023506325 |
| Ezh2 | 76.61519606 | -0.35961498 | 0.018391312 |
| Jam3 | 1274.942713 | -0.362255301 | 0.000507187 |
| Hist1h1d | 1535.36029 | -0.364759162 | 0.000100892 |
| Rras | 635.4314261 | -0.365016693 | 0.004059497 |
| Col5a1 | 47.43147295 | -0.366484633 | 0.025626751 |
| Has2 | 8.465684225 | -0.366487251 | 0.040148834 |
| Msn | 543.1538519 | -0.36693242 | 0.016314268 |
| Cpxm1 | 149.5441767 | -0.368014557 | 0.01753286 |
| Nat6 | 1969.902647 | -0.368174366 | 0.000700845 |
| Cnn3 | 11782.52097 | -0.370476926 | 0.002944693 |
| Angptl1 | 14.28554564 | -0.370888679 | 0.037513104 |
| Klhl6 | 6.45962549 | -0.372105578 | 0.039187048 |
| Lipt1 | 64.19405082 | -0.372773282 | 0.019581267 |
| Lhfpl2 | 1674.534665 | -0.374858062 | 0.00073841 |
| Tgif1 | 141.4663232 | -0.374858504 | 0.016899408 |
| Hes6 | 901.7931827 | -0.374980498 | 0.000225225 |
| Tspan2 | 1214.395667 | -0.376175984 | 0.005197913 |
| Myo9b | 1900.970247 | -0.37623756 | 1.32E-05 |

| | | | |
|---|---|---|---|
| Casp3 | 1145.606211 | -0.377272766 | 0.00031265 |
| Ptgds | 431.6811991 | -0.378984096 | 0.031925301 |
| S1pr3 | 668.1052735 | -0.381488017 | 0.029649038 |
| Flnc | 1787.574605 | -0.382202421 | 0.011438844 |
| Stk10 | 262.5928627 | -0.382347364 | 0.001676216 |
| Cenpf | 40.87962341 | -0.383656148 | 0.027680092 |
| Evc | 766.5800332 | -0.384852644 | 1.04E-06 |
| Ppap2c | 55.41317114 | -0.386096057 | 0.021109252 |
| Jak3 | 209.7068175 | -0.386180998 | 0.012607018 |
| Nedd1 | 239.8728777 | -0.386875028 | 0.001285849 |
| Trip6 | 222.4276306 | -0.390358147 | 0.008762506 |
| Abtb2 | 390.773635 | -0.390977146 | 0.01343357 |
| Mettl4 | 110.9266633 | -0.391858647 | 0.008384842 |
| Mcam | 271.9130899 | -0.391968588 | 0.026427274 |
| Mid1ip1 | 5774.53622 | -0.39348663 | 5.97E-05 |
| Abi3 | 9.451860893 | -0.394962071 | 0.036416567 |
| Gjc2 | 348.932392 | -0.39520737 | 0.030243022 |
| Gtse1 | 677.3664389 | -0.395597125 | 0.006055963 |
| Pth1r | 297.8968999 | -0.396836978 | 0.007409432 |
| Tspo | 174.037152 | -0.397043039 | 0.023667629 |
| Hist1h2bh | 2677.542708 | -0.398643827 | 1.50E-05 |
| Ddit4l2 | 62.59755887 | -0.399159126 | 0.016578751 |
| Acoxl | 4.640937993 | -0.404622187 | 0.033574363 |
| Ppp1r36 | 616.030075 | -0.405182505 | 0.002602523 |
| Gfap | 82659.47166 | -0.408312326 | 0.023149561 |
| Ifih1 | 392.7557854 | -0.408544784 | 0.006128374 |
| Tnfrsf1a | 869.8141703 | -0.410848855 | 0.000684352 |
| Irf7 | 177.6167389 | -0.412099633 | 0.026133776 |
| Ifi27l2b | 36.95406627 | -0.413119453 | 0.028160868 |
| Tp53inp1 | 223.4011143 | -0.414869306 | 0.007321059 |
| Pstpip2 | 75.10619947 | -0.419854083 | 0.016314268 |
| Itpripl2 | 62.36426524 | -0.423309177 | 0.011006391 |
| Mal | 1167.556022 | -0.424512412 | 0.023252548 |
| Traf4 | 344.5691728 | -0.424739712 | 0.004674719 |
| Eln | 273.5725628 | -0.429962313 | 0.003971172 |
| Msmo1 | 5046.888377 | -0.430250996 | 5.99E-05 |

| | | | |
|---|---|---|---|
| Rhoc | 1064.423487 | -0.432806112 | 0.000498523 |
| Tmem176b | 866.0587741 | -0.43641099 | 0.009124945 |
| Slc44a1 | 2513.324765 | -0.43900826 | 0.004190523 |
| Sesn2 | 1349.767074 | -0.443101741 | 0.000726162 |
| Tmem140 | 52.92623149 | -0.443124931 | 0.019129061 |
| Dhcr7 | 1089.346459 | -0.445008642 | 4.49E-05 |
| Clic1 | 623.2728679 | -0.445380317 | 0.001917212 |
| Capn6 | 62.10289866 | -0.448137421 | 0.019795905 |
| Dbi | 5804.038134 | -0.448485942 | 1.12E-05 |
| Cd99 | 411.2074425 | -0.449725701 | 0.001700173 |
| Tagln2 | 209.572136 | -0.449947171 | 0.022155441 |
| RT1-T24-4 | 332.9866622 | -0.450123136 | 0.009214414 |
| Frmd8 | 1288.375719 | -0.450456026 | 2.58E-06 |
| Scnn1a | 130.8892682 | -0.451817568 | 0.011486192 |
| Cd53 | 29.31397271 | -0.465429114 | 0.021793872 |
| Il1rap | 420.4735518 | -0.465925729 | 0.001407415 |
| Elovl1 | 342.9942567 | -0.466319484 | 0.002878747 |
| Tor3a | 442.8835884 | -0.466395287 | 2.22E-05 |
| Abhd4 | 11286.13924 | -0.468748345 | 2.23E-06 |
| Il1r1 | 18.94679035 | -0.469453197 | 0.027680092 |
| Cxcl9 | 96.30732192 | -0.472245263 | 0.016395152 |
| Plp1 | 12769.57229 | -0.477619193 | 0.019790375 |
| Casp4 | 162.2969613 | -0.479438776 | 0.014396978 |
| Lpcat2 | 69.43246267 | -0.479959274 | 0.013501275 |
| Cnp | 7709.926503 | -0.480048692 | 0.005934592 |
| Rbm43 | 116.0170907 | -0.480790707 | 0.004428997 |
| Ifi27 | 2899.841378 | -0.484007576 | 0.006565133 |
| Fgl2 | 350.6114085 | -0.486693439 | 0.004141515 |
| Dapp1 | 39.33194442 | -0.500320602 | 0.015521472 |
| Abca8a | 820.8891901 | -0.500321478 | 0.013628951 |
| Parp14 | 727.3988232 | -0.503045617 | 0.010645539 |
| Rps27l | 754.3182657 | -0.504094874 | 0.000249744 |
| Csf1 | 1353.315249 | -0.509718736 | 0.001868329 |
| Fyb | 30.4627279 | -0.514918467 | 0.020189274 |
| Mtbp | 53.50450301 | -0.51518719 | 0.00849184 |
| Sp110 | 128.8750053 | -0.515669833 | 0.006513242 |

| | | | |
|---|---|---|---|
| Cp | 1313.988663 | -0.517323603 | 0.016370996 |
| Sulf2 | 6138.498987 | -0.523942335 | 1.46E-06 |
| Ppfibp1 | 266.6488229 | -0.534968221 | 0.000293897 |
| Nckap1l | 23.75282459 | -0.535582283 | 0.020970738 |
| Gltp | 759.5495566 | -0.536511216 | 4.26E-05 |
| Uba7 | 96.81122359 | -0.537012042 | 0.016858701 |
| Parp9 | 461.0214173 | -0.540038111 | 0.002762787 |
| Phldb1 | 1664.013856 | -0.540830962 | 0.000357347 |
| Tap1 | 837.8446153 | -0.548873825 | 0.014396859 |
| Fam83h | 175.1244488 | -0.551319549 | 0.000680072 |
| Pbld1 | 51.0945686 | -0.554986824 | 0.016370996 |
| Scd1 | 1562.086303 | -0.559723507 | 0.000230695 |
| Ccng1 | 10133.45714 | -0.561428604 | 1.74E-07 |
| Ckap2 | 73.95254083 | -0.561964567 | 0.009178759 |
| Fcer1g | 28.10389691 | -0.567302154 | 0.015719664 |
| Pik3ap1 | 12.08033237 | -0.568647455 | 0.020991808 |
| Atf3 | 24.1250456 | -0.569965234 | 0.021290194 |
| Lpar4 | 398.4475835 | -0.572148478 | 0.000140594 |
| Igfbp2 | 205.9170566 | -0.572456775 | 0.018627354 |
| Col16a1 | 841.3049237 | -0.575477635 | 0.002140696 |
| Sapcd2 | 23.01975723 | -0.577957529 | 0.014274859 |
| Chst7 | 214.9631663 | -0.581057119 | 0.001324144 |
| Cndp1 | 28.04915895 | -0.581404793 | 0.012625964 |
| Apbb1ip | 73.60933608 | -0.583193179 | 0.004861903 |
| Icam1 | 182.8345173 | -0.585383741 | 0.002656757 |
| Mybl1 | 266.8497999 | -0.590370348 | 9.10E-05 |
| Tnfaip8l2 | 8.82233051 | -0.590764482 | 0.02175952 |
| Hmox1 | 477.2556122 | -0.597223262 | 0.003269266 |
| Plxnb3 | 591.4798738 | -0.599030443 | 0.001288941 |
| Creb5 | 233.2058428 | -0.610044272 | 0.000168417 |
| Afap1l2 | 493.762997 | -0.61034595 | 0.001276383 |
| Slc39a12 | 35.11170442 | -0.610981398 | 0.004593389 |
| Sat1 | 2477.251845 | -0.611576803 | 1.91E-05 |
| Phlda3 | 2907.027466 | -0.612669151 | 2.35E-06 |
| Mmp19 | 26.68943432 | -0.612718367 | 0.01947731 |
| Adamts4 | 212.0055621 | -0.621722248 | 0.01174786 |

| | | | |
|---|---|---|---|
| Tmem176a | 253.5486234 | -0.621839021 | 0.004428997 |
| Fas | 421.6733578 | -0.624566485 | 0.000114633 |
| Slfn13 | 58.25029285 | -0.625631678 | 0.011119647 |
| Slc27a3 | 810.5252823 | -0.63342859 | 2.86E-05 |
| C1r | 1431.916971 | -0.636318841 | 0.006162965 |
| Ddr2 | 326.0566843 | -0.638048924 | 0.001288941 |
| Bgn | 384.8191134 | -0.639518666 | 0.001407415 |
| Myo1f | 10.0905573 | -0.645652547 | 0.018898778 |
| Shisa2 | 130.0884064 | -0.649068361 | 0.003984864 |
| Trim25 | 159.7712734 | -0.657927718 | 0.00151922 |
| Mag | 1649.993673 | -0.659226315 | 0.011250162 |
| Fcgr2b | 7.961674412 | -0.664199685 | 0.01687174 |
| Fam167a | 248.998344 | -0.682455445 | 0.000441904 |
| Mt2A | 1590.425522 | -0.6853231 | 0.000179332 |
| Ephx1 | 4112.309551 | -0.688176943 | 1.04E-07 |
| P2ry6 | 11.97141036 | -0.691436644 | 0.014396978 |
| Fermt3 | 44.32264378 | -0.693854371 | 0.007321059 |
| Rsad2 | 467.7145128 | -0.707565338 | 0.014125329 |
| Tshr | 191.4292831 | -0.721503895 | 0.005915258 |
| Sfrp2 | 168.2914911 | -0.729424801 | 0.011128752 |
| Cldn11 | 1499.759943 | -0.741089743 | 0.010543901 |
| Dmrtb1 | 9.385722267 | -0.751295232 | 0.015711943 |
| Slc27a6 | 19.39994421 | -0.751393442 | 0.01174786 |
| Vcan | 592.783212 | -0.773213953 | 8.18E-05 |
| Ifitm3 | 147.5282011 | -0.781325308 | 0.013233116 |
| Cdk6 | 54.12938918 | -0.784141518 | 0.007338459 |
| Bcas1 | 1224.886675 | -0.786328882 | 6.37E-05 |
| Fn1 | 217.779601 | -0.794937524 | 5.74E-05 |
| Serinc5 | 1913.92908 | -0.812336398 | 1.57E-07 |
| Cdkn1a | 2806.646878 | -0.821091515 | 0.000105202 |
| Mx1 | 235.1915312 | -0.836675881 | 0.013391727 |
| Enpp6 | 34.60236515 | -0.842241015 | 0.009999001 |
| Blnk | 15.76471571 | -0.861909475 | 0.011362677 |
| Cpm | 218.4631179 | -0.862532755 | 0.001805848 |
| Ascl3 | 9.517780177 | -0.878962782 | 0.010494278 |
| Mog | 525.3683414 | -0.884796673 | 0.005551777 |

| | | | |
|---|---|---|---|
| RT1-Ba | 29.63190245 | -0.89246381 | 0.007308942 |
| Capg | 150.5138036 | -0.894329041 | 0.008869174 |
| C1s | 1004.124088 | -0.919828565 | 0.002237094 |
| Tubb6 | 165.3425705 | -0.927523382 | 0.005089195 |
| Top2a | 47.45441091 | -0.945268112 | 0.00691669 |
| C1rl | 38.12957049 | -0.946436148 | 0.005525371 |
| Bcl3 | 46.16682157 | -0.96447445 | 0.008758482 |
| Pllp | 423.2745054 | -0.966952012 | 0.001516437 |
| MGC105567 | 68.41163706 | -0.983525219 | 0.007338459 |
| Gbp5 | 437.0001691 | -0.992361907 | 0.005664875 |
| Ppic | 74.78665036 | -1.012450567 | 0.000231657 |
| Kazald1 | 9.980896919 | -1.012458184 | 0.007744014 |
| Sema3d | 399.1903164 | -1.014117146 | 0.000277124 |
| Rtp4 | 158.6066423 | -1.044060841 | 0.000948928 |
| Aspg | 68.05182961 | -1.057483834 | 0.004818988 |
| Nusap1 | 11.91858286 | -1.065623168 | 0.007672234 |
| Serping1 | 1972.464505 | -1.084073517 | 0.006257689 |
| Pdlim2 | 69.64443656 | -1.135847208 | 3.19E-05 |
| LOC100910973 | 298.5221864 | -1.160191756 | 0.004082193 |
| Gpnmb | 751.3300052 | -1.165600055 | 0.000443011 |
| MGC108823 | 555.5814526 | -1.184585026 | 0.006260517 |
| Csf1r | 101.6269058 | -1.218139294 | 0.000128235 |
| Abcb1b | 225.4696266 | -1.225638842 | 4.11E-05 |
| Isg15 | 182.6332451 | -1.246078133 | 0.004456658 |
| Mmp2 | 274.5889951 | -1.275165003 | 7.19E-05 |
| Timp1 | 53.75333992 | -1.314132847 | 0.003797014 |
| Col1a2 | 117.3632677 | -1.345980738 | 5.93E-05 |
| C4a | 929.708177 | -1.397072719 | 0.002778773 |
| Chst3 | 17.43999588 | -1.494175015 | 0.003063532 |
| Osmr | 43.44387584 | -1.519487541 | 0.002877356 |
| Cav2 | 153.7741035 | -1.539295259 | 1.04E-07 |
| Mpeg1 | 101.5736678 | -1.568781001 | 6.76E-05 |
| Olig2 | 606.1417983 | -1.583647901 | 1.24E-12 |
| C1qc | 28.7981149 | -1.593387838 | 0.00169016 |
| Gbp2 | 833.7330989 | -1.643867747 | 0.001811262 |
| Ifit3 | 71.33921826 | -1.652262351 | 0.003432328 |

| | | | |
|---|---|---|---|
| Calcrl | 100.4629871 | -1.671020497 | 7.08E-08 |
| Apol3 | 20.75104388 | -1.719119124 | 0.003269266 |
| Cxcl11 | 11.58116095 | -1.736137368 | 0.002762787 |
| Sox10 | 635.6735299 | -1.795637679 | 1.04E-08 |
| Erbb3 | 319.9841722 | -1.796213384 | 5.96E-10 |
| Cd74 | 204.7571086 | -1.828830925 | 0.002516154 |
| Bmp4 | 47.98336117 | -1.870570659 | 6.14E-06 |
| Lcn2 | 68.49004995 | -1.871829547 | 0.002617839 |
| Olig1 | 1529.229499 | -1.880378756 | 2.67E-21 |
| Plek | 14.73041719 | -1.957588344 | 0.001323415 |
| Matn4 | 42.16246145 | -1.987513922 | 0.000344829 |
| Ctss | 133.0650958 | -1.999069228 | 3.90E-07 |
| Slc2a5 | 12.96004967 | -2.000790396 | 0.000320981 |
| Slamf8 | 36.33508678 | -2.002749563 | 0.000249744 |
| Apol9a | 42.14713388 | -2.015770399 | 0.002125131 |
| Cfh | 70.46291433 | -2.108313076 | 6.05E-05 |
| RGD1309362 | 502.1726712 | -2.134762524 | 0.000991409 |
| C3 | 406.5144916 | -2.199849014 | 0.000616825 |
| Arhgap31 | 222.8017751 | -2.262554934 | 2.69E-18 |
| Anxa3 | 78.42925651 | -2.263577662 | 5.31E-08 |
| Itgb2 | 25.08091154 | -2.28356546 | 0.000359237 |
| C1qb | 45.86784839 | -2.362455347 | 1.05E-05 |
| C1qa | 56.85532742 | -2.39121642 | 7.73E-07 |
| Itgam | 40.20327333 | -2.403233019 | 9.73E-05 |
| Cav1 | 196.5938146 | -2.407815518 | 3.49E-19 |
| Pnlip | 31.23089454 | -2.431650964 | 1.45E-07 |
| RT1-DMb | 21.04219302 | -2.520939949 | 6.57E-05 |
| Adgre1 | 20.77759917 | -2.580663879 | 5.31E-05 |
| Neu4 | 77.4446178 | -2.583153547 | 1.20E-14 |
| Cxcl10 | 223.4037959 | -2.605597103 | 0.000660142 |
| Ccl2 | 17.62441728 | -2.62877559 | 0.000726162 |
| Pld4 | 33.06495369 | -2.643403411 | 4.85E-06 |
| Cspg4 | 768.3465006 | -2.728601575 | 8.28E-41 |
| Col5a3 | 438.4709126 | -2.767642134 | 3.49E-19 |
| Ifi47 | 101.4097503 | -2.984629583 | 0.000269768 |
| Chst5 | 30.45888523 | -3.06465726 | 7.40E-10 |

| | | | |
|---|---|---|---|
| Ccnd1 | 344.3165307 | -3.116776442 | 1.66E-28 |
| Fcrl2 | 26.96153006 | -3.502689308 | 1.04E-07 |
| RGD1311892 | 21.50503207 | -3.568691929 | 5.96E-10 |
| Pdgfra | 1182.078546 | -3.575286571 | 8.28E-41 |
| Fam89a | 33.19435762 | -3.594384099 | 9.93E-14 |
| Gpr17 | 177.4145954 | -4.084783099 | 1.82E-43 |

| Gene | baseMean | log2FoldChange | adjusted p-value |
|---|---|---|---|
| Myh9l1 | 144.6701119 | 23.81830111 | 3.23E-13 |
| Egr2 | 208.3089698 | 2.108544744 | 3.08E-06 |
| Npas4 | 362.011499 | 2.037243141 | 3.05E-10 |
| Arc | 2731.514471 | 1.939018901 | 9.22E-08 |
| Nr4a1 | 2042.958823 | 1.740833785 | 3.60E-07 |
| Egr4 | 804.1976775 | 1.676648988 | 1.40E-13 |
| Dusp1 | 739.5727634 | 1.502870465 | 1.36E-11 |
| Fosb | 361.8514835 | 1.499411721 | 9.15E-10 |
| Egr1 | 3992.245488 | 1.421618371 | 6.47E-07 |
| Mypn | 35.94440795 | 1.288841225 | 3.54E-05 |
| Tac1 | 1029.543161 | 1.18784906 | 1.09E-10 |
| Junb | 1525.00745 | 1.181000672 | 1.24E-07 |
| Scn4b | 2014.67136 | 1.134635559 | 2.37E-12 |
| Tgm3 | 5.732101102 | 1.118447486 | 0.002790095 |
| Lamp5 | 659.1269984 | 1.11483698 | 4.19E-14 |
| Gpr6 | 440.7409934 | 1.101682703 | 9.81E-08 |
| Ankrd33b | 310.0086934 | 1.054852246 | 2.14E-06 |
| Gpr88 | 3692.090795 | 1.051769247 | 4.94E-13 |
| Egr3 | 697.7009269 | 1.038240562 | 2.23E-05 |
| Hpca | 6647.641839 | 1.005351731 | 7.10E-15 |
| Rasl10a | 209.9237614 | 1.003509788 | 1.10E-05 |
| Drd1 | 2460.609413 | 0.98870371 | 5.62E-12 |
| Pdyn | 2314.972886 | 0.983151172 | 1.14E-12 |
| Cyr61 | 52.8309587 | 0.977827607 | 0.000596735 |
| Slc4a11 | 143.1922978 | 0.959476665 | 6.89E-06 |
| Rem2 | 209.4541881 | 0.940598999 | 7.01E-07 |
| Otof | 1233.681645 | 0.909272401 | 2.55E-06 |
| Mei1 | 16.83055069 | 0.903892983 | 0.000401291 |
| Pde10a | 3128.744671 | 0.893431924 | 3.05E-11 |
| Vipr1 | 182.9855448 | 0.857444698 | 0.000143524 |
| Sh3rf2 | 180.7510921 | 0.857341867 | 2.70E-06 |
| Iqgap3 | 175.4441161 | 0.831027533 | 1.90E-06 |
| Rasd2 | 4370.532188 | 0.825523789 | 4.88E-16 |
| Kcnq5 | 2178.91997 | 0.818634595 | 3.31E-07 |
| Tac3 | 504.7993661 | 0.815152222 | 9.66E-07 |

| | | | |
|---|---|---|---|
| Adra1d | 116.7405174 | 0.814226764 | 0.000442477 |
| Penk | 5862.017377 | 0.811376134 | 7.40E-11 |
| Akap5 | 3549.165687 | 0.807548703 | 8.11E-10 |
| Adamts2 | 133.8960095 | 0.801835697 | 5.01E-05 |
| Lmo7 | 1735.253239 | 0.793276354 | 2.30E-08 |
| Gpr22 | 923.6328424 | 0.787721623 | 2.02E-06 |
| Lpl | 1505.027139 | 0.786175891 | 9.22E-08 |
| Camkk2 | 1352.051574 | 0.763381205 | 1.25E-09 |
| Cnksr2 | 4805.903656 | 0.761812307 | 2.04E-07 |
| Actn2 | 953.5597086 | 0.760521501 | 3.99E-07 |
| Rbfox1 | 6488.222247 | 0.759182439 | 1.08E-08 |
| Foxp2 | 1693.100371 | 0.755235212 | 4.08E-09 |
| Meox2 | 21.17538583 | 0.752891631 | 0.000919167 |
| Kcnj4 | 1864.314857 | 0.746942322 | 7.13E-05 |
| Rgs9 | 1292.779767 | 0.746203704 | 2.02E-08 |
| Pcp4l1 | 2618.913209 | 0.744478213 | 3.84E-05 |
| Ppp3ca | 13759.26458 | 0.743886856 | 1.11E-11 |
| Pnma1 | 87.61624907 | 0.74154444 | 4.47E-05 |
| Camk4 | 846.2195121 | 0.739298463 | 7.35E-08 |
| Bcl11b | 5738.545239 | 0.731411797 | 7.07E-10 |
| Ephx4 | 336.8414177 | 0.72827323 | 6.30E-05 |
| Asb2 | 102.6090581 | 0.728128152 | 3.72E-05 |
| Cpne5 | 4955.728076 | 0.727663927 | 7.10E-15 |
| Chrm4 | 1015.902001 | 0.725242892 | 1.38E-08 |
| Kcnab1 | 4760.611347 | 0.724808012 | 2.10E-07 |
| Chn1 | 9877.653785 | 0.723982168 | 3.16E-07 |
| Nkiras1 | 2031.171229 | 0.723509448 | 8.48E-08 |
| Kcnh4 | 195.5494691 | 0.723303343 | 3.13E-06 |
| Atp2b1 | 16574.70991 | 0.721640756 | 2.88E-08 |
| Arl4d | 404.017775 | 0.719229753 | 1.93E-05 |
| Necab1 | 930.2600701 | 0.713767509 | 1.12E-06 |
| Hrh3 | 6304.74388 | 0.706707883 | 0.000174491 |
| Cobl | 2385.810692 | 0.701683336 | 3.78E-05 |
| Prkcg | 6127.999555 | 0.701457803 | 9.58E-05 |
| Neto1 | 2814.144474 | 0.697932647 | 1.04E-10 |
| Fbxl16 | 3707.583559 | 0.696694178 | 4.00E-09 |

| | | | |
|---|---|---|---|
| Ptpn5 | 2961.43227 | 0.695399525 | 5.66E-14 |
| Adora2a | 1059.212289 | 0.693285249 | 1.77E-07 |
| Klhl40 | 28.94022396 | 0.690508665 | 0.00273196 |
| Filip1 | 795.8498352 | 0.690313552 | 1.84E-05 |
| Arhgef3 | 2565.163735 | 0.689422781 | 1.54E-11 |
| Wipf3 | 4144.753322 | 0.688992671 | 1.95E-10 |
| Pde1b | 4099.810897 | 0.687554532 | 9.92E-14 |
| Plxdc1 | 315.9379147 | 0.687206344 | 4.38E-05 |
| Pcdha12 | 202.5483124 | 0.687117022 | 0.000186166 |
| LOC100125362 | 4464.312302 | 0.686128748 | 1.06E-12 |
| Wnt10a | 63.21704375 | 0.684601565 | 0.004898757 |
| Nab2 | 660.3000428 | 0.683325109 | 3.08E-06 |
| F12 | 61.90119326 | 0.682109558 | 0.001414533 |
| St8sia5 | 491.1456659 | 0.68126479 | 3.04E-05 |
| Fhl2 | 172.5691735 | 0.679955099 | 0.00031594 |
| Itk | 47.5648701 | 0.678814409 | 0.001215384 |
| Olfml2a | 40.09550527 | 0.677578376 | 0.001414178 |
| Phyhip | 10432.63686 | 0.677450157 | 4.58E-07 |
| Wnk4 | 130.5097941 | 0.674616845 | 1.57E-05 |
| Spata2L | 1064.428666 | 0.674366229 | 7.75E-06 |
| Wfs1 | 3962.317004 | 0.673883051 | 7.04E-06 |
| Chst15 | 388.2960473 | 0.673031898 | 7.88E-06 |
| Itpka | 1454.365021 | 0.670111299 | 0.000444085 |
| Nox1 | 9.178741404 | 0.668466694 | 0.006632602 |
| Syt10 | 478.4317984 | 0.668253039 | 2.74E-05 |
| Rbfox3 | 1749.396537 | 0.667756291 | 9.22E-08 |
| Sult2b1 | 50.74988065 | 0.667599003 | 0.000352638 |
| Lzts3 | 2519.475395 | 0.66673511 | 9.92E-14 |
| Gabrd | 501.2601206 | 0.666154593 | 0.000204636 |
| Lzts1 | 4425.499905 | 0.661149845 | 3.74E-05 |
| Asic4 | 1085.854724 | 0.660660741 | 3.16E-07 |
| Grin2a | 3875.913346 | 0.660469403 | 2.43E-07 |
| Foxp1 | 1725.540346 | 0.660079488 | 1.48E-07 |
| Cacna2d1 | 2992.394182 | 0.657230223 | 1.80E-05 |
| Itpr1 | 18050.86261 | 0.653480633 | 0.000197383 |
| Adcy5 | 9082.114656 | 0.653463696 | 1.02E-09 |

| | | | |
|---|---|---|---|
| Cdc42ep3 | 349.6067523 | 0.653096294 | 6.51E-06 |
| Grasp | 423.3759378 | 0.649002478 | 0.00034315 |
| Cyfip2 | 35728.84141 | 0.645243523 | 2.42E-08 |
| Prkcb | 11392.10926 | 0.644119147 | 3.15E-06 |
| Gng7 | 3688.405217 | 0.64226777 | 9.49E-10 |
| Rtn4rl2 | 311.8880704 | 0.641743108 | 0.003193465 |
| Pcsk2 | 5014.415662 | 0.637948898 | 1.88E-06 |
| Atp6ap1l | 333.5030067 | 0.637907654 | 8.68E-05 |
| Trpv6 | 2.63460037 | 0.635558848 | 0.007870292 |
| Tmem158 | 942.0412048 | 0.634945142 | 7.63E-07 |
| Fos | 772.4876676 | 0.633953785 | 0.000306623 |
| Plk2 | 5928.913533 | 0.63199117 | 6.49E-05 |
| Camk2b | 21029.65565 | 0.631831087 | 2.01E-08 |
| Smpdl3b | 214.9437671 | 0.630599157 | 3.24E-05 |
| Rab40b | 707.7123691 | 0.629390905 | 4.04E-06 |
| Grm4 | 1183.492015 | 0.628278451 | 1.88E-05 |
| Baiap2 | 2676.559735 | 0.62675439 | 0.000192625 |
| Synpr | 2853.159898 | 0.625803832 | 4.94E-13 |
| Sptssb | 111.7403348 | 0.624745707 | 0.000625534 |
| Clspn | 42.97953266 | 0.624090092 | 0.002350764 |
| Htr1b | 271.3883885 | 0.623398346 | 3.58E-05 |
| Paqr9 | 831.9801669 | 0.622552585 | 2.93E-07 |
| Camk2a | 9952.28621 | 0.622051975 | 0.000308642 |
| Mast3 | 5755.988864 | 0.621341047 | 2.03E-06 |
| Vcsa1 | 1.974150951 | 0.62083823 | 0.008198141 |
| Arpp21 | 8234.651489 | 0.62042415 | 1.32E-07 |
| Rgs7bp | 2857.56348 | 0.620364667 | 3.92E-09 |
| Cacnb4 | 3454.949447 | 0.617378746 | 9.15E-10 |
| Camkv | 9066.030183 | 0.616518477 | 0.000149879 |
| RGD1564053 | 655.3753645 | 0.614688454 | 0.000224302 |
| Rgs14 | 1323.72166 | 0.614324533 | 0.000841011 |
| Tgfb3 | 118.3142369 | 0.611911623 | 0.000899649 |
| Nell2 | 10449.06259 | 0.610753154 | 8.46E-05 |
| Kcns2 | 241.8978562 | 0.610619742 | 0.000339194 |
| Epha7 | 1289.708386 | 0.610176201 | 0.000179286 |
| Wasf1 | 6387.14192 | 0.609799314 | 2.59E-06 |

| | | | |
|---|---|---|---|
| Nrip3 | 4428.754177 | 0.609665882 | 3.02E-06 |
| Tomm70a | 5381.255247 | 0.608363606 | 4.49E-16 |
| Kcna5 | 324.3681168 | 0.608351364 | 5.26E-07 |
| Agap2 | 25273.51571 | 0.608215476 | 2.23E-05 |
| Zfp365 | 4674.928452 | 0.60667289 | 3.56E-05 |
| Elmod1 | 4820.480998 | 0.60628014 | 4.31E-12 |
| Nt5dc3 | 1355.943175 | 0.605546158 | 1.80E-05 |
| Dusp5 | 83.4923023 | 0.602566371 | 0.003250778 |
| Ngef | 9054.739399 | 0.602498488 | 0.000112453 |
| Rsph6a | 47.48245875 | 0.602248321 | 0.000761855 |
| Crip3 | 9.313828631 | 0.602061017 | 0.008851239 |
| Pcdha7 | 286.1242017 | 0.60198553 | 3.85E-05 |
| Lin7b | 295.4588529 | 0.598428759 | 1.49E-05 |
| Synpo | 3323.626664 | 0.597512015 | 0.0007671 |
| Brinp2 | 2530.238248 | 0.596381942 | 0.000419923 |
| Nrgn | 7913.210254 | 0.596357195 | 0.001410214 |
| Sema3e | 1695.146287 | 0.594927398 | 0.000504195 |
| Rasgrp1 | 15547.33434 | 0.593915314 | 0.001358472 |
| Kcna1 | 1418.315824 | 0.591083112 | 0.000185068 |
| B3gnt2 | 658.5211906 | 0.590025128 | 3.48E-11 |
| Ccsap | 686.5411567 | 0.588682383 | 3.75E-05 |
| Phactr1 | 7164.961389 | 0.588032038 | 3.78E-06 |
| Pde2a | 7721.057815 | 0.58773285 | 4.14E-05 |
| Zfp189 | 419.9390402 | 0.587624879 | 5.19E-08 |
| Ypel2 | 881.7954708 | 0.587543054 | 4.47E-06 |
| Dmkn | 183.8904233 | 0.58739731 | 0.000256999 |
| Lingo3 | 630.4617561 | 0.586916049 | 5.85E-07 |
| Acvr1c | 167.7972018 | 0.58658824 | 7.45E-05 |
| Tmem41a | 644.4186826 | 0.586527761 | 3.62E-06 |
| Adamts3 | 509.5222158 | 0.585879423 | 1.17E-05 |
| Btbd3 | 1953.347438 | 0.584875076 | 1.37E-05 |
| Rxrg | 513.5306764 | 0.584601525 | 5.75E-05 |
| Mchr1 | 300.1029726 | 0.584452841 | 0.000308642 |
| Me3 | 1050.478336 | 0.583724911 | 1.57E-07 |
| Dlgap3 | 8347.265491 | 0.58280674 | 1.13E-08 |
| Cartpt | 282.3164318 | 0.582285285 | 0.001199925 |

| | | | |
|---|---|---|---|
| Ociad2 | 817.5755002 | 0.582152334 | 6.74E-05 |
| Jph4 | 3728.516887 | 0.581532982 | 1.98E-07 |
| Pak6 | 1920.796796 | 0.581426717 | 1.31E-05 |
| Gpr176 | 513.2134578 | 0.581367192 | 1.49E-07 |
| Cabp1 | 1223.691709 | 0.580984542 | 0.001983535 |
| Klf5 | 914.7900961 | 0.580818766 | 2.56E-06 |
| Homer1 | 3116.405844 | 0.579455628 | 1.94E-05 |
| Rbm24 | 311.2640982 | 0.579208565 | 3.18E-05 |
| Dpf1 | 1786.976031 | 0.579118881 | 1.22E-05 |
| Gpr52 | 295.6522703 | 0.57846164 | 1.28E-05 |
| Prkce | 6819.924697 | 0.578152401 | 8.10E-05 |
| Slc24a2 | 18918.65161 | 0.578012579 | 0.000376834 |
| Trim46 | 1005.155031 | 0.576523814 | 3.99E-08 |
| Neurl1 | 612.5792736 | 0.576443909 | 2.08E-07 |
| Syndig1l | 492.9232635 | 0.574782893 | 1.26E-06 |
| C1qtnf4 | 4616.141374 | 0.572701667 | 9.42E-05 |
| Shisa8 | 335.2597743 | 0.572442396 | 0.003527511 |
| Pvalb | 959.2398038 | 0.571658476 | 0.001453268 |
| Scrt2 | 189.7509082 | 0.571401764 | 5.55E-05 |
| Kcnk4 | 101.8104906 | 0.570569419 | 0.001215384 |
| C2cd2l | 6536.597542 | 0.570507723 | 9.49E-10 |
| Rnf150 | 1010.121082 | 0.569319156 | 2.67E-06 |
| Slit3 | 1851.472698 | 0.568958388 | 0.000625779 |
| Tex30 | 59.92324693 | 0.568546735 | 0.001658992 |
| Ccdc28a | 348.9692023 | 0.568362189 | 3.56E-05 |
| Cntn3 | 470.752797 | 0.568096488 | 8.12E-06 |
| Unc5a | 1313.188097 | 0.568047718 | 2.72E-05 |
| Hs6st3 | 583.0616438 | 0.567341617 | 0.000271264 |
| Fam184b | 256.5774273 | 0.565577909 | 2.93E-05 |
| Igfbp4 | 1526.980126 | 0.565497776 | 9.81E-06 |
| Lrrk2 | 2983.365092 | 0.565166229 | 5.18E-07 |
| Rab15 | 3839.825747 | 0.564901599 | 9.90E-05 |
| Pgbd5 | 1892.299772 | 0.564765937 | 1.64E-05 |
| Snap25 | 55739.23787 | 0.564574847 | 4.23E-06 |
| Kcng2 | 177.8011108 | 0.564487813 | 0.000446175 |
| Lpcat4 | 2035.263187 | 0.564309512 | 8.59E-08 |

| | | | |
|---|---|---|---|
| Them6 | 772.1520529 | 0.564241694 | 1.97E-05 |
| Nr4a2 | 1528.79325 | 0.564172932 | 0.006515386 |
| Rbp4 | 867.4984385 | 0.563165077 | 1.49E-06 |
| Stxbp5l | 2311.762837 | 0.562340438 | 4.12E-06 |
| Icam5 | 4234.595705 | 0.562055989 | 0.000867579 |
| Aspdh | 7.646066562 | 0.562048857 | 0.012718395 |
| LOC365985 | 4128.846766 | 0.561272917 | 0.000581169 |
| Kcnh7 | 1166.850863 | 0.560902122 | 0.001897506 |
| Pcp4 | 4956.296168 | 0.559659729 | 9.96E-05 |
| Sept5 | 12860.09435 | 0.557516029 | 0.000319381 |
| Ets2 | 1893.465471 | 0.55651816 | 1.00E-04 |
| Nr4a3 | 1949.741033 | 0.556201759 | 0.002203946 |
| Ptpn7 | 23.53600203 | 0.554517022 | 0.006541234 |
| Osbp2 | 1769.834243 | 0.553506063 | 1.33E-06 |
| Car7 | 123.5345472 | 0.553433658 | 0.005490502 |
| Chrm1 | 1527.394165 | 0.552741806 | 0.002376254 |
| Gria3 | 5871.813628 | 0.552187714 | 0.0009061 |
| Celf5 | 8385.921957 | 0.551724819 | 7.71E-06 |
| Rnf144b | 387.3649307 | 0.551130519 | 2.34E-06 |
| Vstm5 | 166.8239076 | 0.550913948 | 0.000492059 |
| Mapk10 | 7421.416707 | 0.550680927 | 6.91E-07 |
| Slc1a1 | 1185.28467 | 0.550451477 | 8.01E-07 |
| Lppr4 | 6724.95128 | 0.55021875 | 1.11E-06 |
| Cacna1e | 6691.607503 | 0.549222867 | 0.000436238 |
| Actn1 | 2500.685878 | 0.54858193 | 1.51E-07 |
| Brinp1 | 1953.770056 | 0.548070772 | 1.05E-05 |
| B4galnt1 | 1884.876455 | 0.546277629 | 1.78E-07 |
| Rgs2 | 449.5986831 | 0.544747992 | 1.17E-05 |
| Rtn4r | 404.4611752 | 0.544665091 | 0.004859086 |
| Ppp3r1 | 12868.93689 | 0.543096548 | 2.20E-07 |
| Snca | 4029.951954 | 0.542734796 | 0.001644076 |
| Dact2 | 830.6093251 | 0.542258865 | 7.90E-09 |
| Ppp2r2c | 3151.594588 | 0.541628287 | 5.14E-08 |
| Syngap1 | 8854.815604 | 0.540981267 | 4.22E-05 |
| Myt1l | 6671.716802 | 0.540874622 | 3.54E-06 |
| Itpr3 | 123.9712554 | 0.540715399 | 0.000841011 |

| | | | |
|---|---|---|---|
| Stk32c | 3120.77321 | 0.540350975 | 4.70E-06 |
| Rapgef5 | 2791.898987 | 0.540199493 | 1.35E-09 |
| Kcnh1 | 1848.962045 | 0.540161014 | 0.000347896 |
| Kcna4 | 1258.438944 | 0.539692596 | 1.56E-06 |
| Pdp1 | 1392.642657 | 0.539440384 | 1.47E-07 |
| Zfyve28 | 813.3477631 | 0.538833846 | 2.26E-05 |
| Cacnb2 | 1537.541359 | 0.538422679 | 2.85E-11 |
| Adra2c | 361.7077492 | 0.537383937 | 0.000152508 |
| Slc24a4 | 235.5612437 | 0.536939245 | 0.001308074 |
| Adrb1 | 414.4639453 | 0.536318856 | 8.88E-05 |
| Kcnq3 | 3989.365636 | 0.535705457 | 9.06E-10 |
| Rbm3 | 271.9303994 | 0.535284525 | 0.000324456 |
| Mtus2 | 205.9264079 | 0.534773934 | 1.32E-05 |
| Tbc1d8 | 607.5096784 | 0.534385453 | 4.54E-07 |
| Gnal | 4382.720362 | 0.533723489 | 3.02E-05 |
| LOC689986 | 2517.304954 | 0.533721603 | 4.39E-06 |
| Kctd16 | 348.0764504 | 0.533411539 | 1.99E-05 |
| Rin1 | 953.3915682 | 0.533298713 | 0.000954473 |
| Il34 | 1087.977786 | 0.532954305 | 0.000187842 |
| Fmnl1 | 2422.777956 | 0.532923139 | 3.00E-05 |
| Entpd3 | 551.0415989 | 0.532916009 | 0.000352725 |
| Dgat2 | 899.8983918 | 0.531925839 | 0.000118169 |
| Kcnf1 | 2714.619367 | 0.531014072 | 0.000431798 |
| Fam131b | 3674.809188 | 0.530623061 | 4.82E-05 |
| Prrt2 | 3483.827254 | 0.530253031 | 3.06E-07 |
| Dmtn | 4865.598454 | 0.52970867 | 9.50E-10 |
| Caln1 | 984.4286982 | 0.529540412 | 7.52E-05 |
| Psd | 6006.039369 | 0.529519551 | 2.25E-05 |
| Dgkz | 8568.064666 | 0.527972216 | 0.000257505 |
| Entpd7 | 118.4316803 | 0.527810997 | 0.000417699 |
| Tpd52l1 | 355.7690555 | 0.527695271 | 0.001318499 |
| Pex5l | 1300.499436 | 0.526071413 | 0.000192625 |
| Nkx3-1 | 14.09965172 | 0.5253894 | 0.012551471 |
| Rprml | 563.0839023 | 0.523828543 | 0.002521161 |
| Lrp12 | 1832.034417 | 0.522459174 | 6.36E-07 |
| Lrrc73 | 586.133954 | 0.522374306 | 2.31E-06 |

| | | | |
|---|---|---|---|
| Krt71 | 179.7581875 | 0.520874742 | 0.001418423 |
| Cpne6 | 1475.871419 | 0.520584612 | 9.86E-06 |
| Dgkg | 1766.043182 | 0.520225965 | 6.73E-06 |
| Ankrd34a | 2740.757881 | 0.520208122 | 1.21E-08 |
| Clec1a | 111.2224015 | 0.519889163 | 0.008875859 |
| Gas7 | 17987.72684 | 0.519377254 | 0.001394243 |
| Pik3cd | 424.5466664 | 0.519068543 | 0.000741777 |
| Prmt8 | 831.6093903 | 0.518930093 | 0.000270498 |
| Kcnj3 | 973.6030349 | 0.518915316 | 0.00351618 |
| Actr3b | 551.656246 | 0.518447812 | 0.000155747 |
| Fam102b | 1415.985297 | 0.518422463 | 6.02E-06 |
| Kcnh3 | 1732.590901 | 0.518371379 | 0.006817002 |
| Anks1b | 4475.084025 | 0.518106927 | 1.04E-05 |
| Gpr173 | 113.8205336 | 0.517639565 | 0.000104103 |
| Pcdhac2 | 3144.609846 | 0.517308073 | 4.55E-08 |
| Diras2 | 3899.631782 | 0.517153971 | 0.004323463 |
| Zdhhc14 | 1138.058457 | 0.517119469 | 2.81E-07 |
| Ppp4r4 | 433.5174745 | 0.516718206 | 9.60E-06 |
| Nptn | 9318.334023 | 0.516544467 | 1.53E-11 |
| Ydjc | 420.4265211 | 0.516497077 | 6.82E-05 |
| Kcnq2 | 6053.880849 | 0.515979827 | 6.65E-08 |
| Cnih3 | 146.9952285 | 0.515767019 | 0.007817863 |
| C1ql3 | 537.0397615 | 0.51574649 | 0.00547698 |
| Fkbp1a | 7589.237552 | 0.515738184 | 1.28E-07 |
| Calb1 | 1969.77555 | 0.514025039 | 0.00230929 |
| Ptgs2 | 104.6965175 | 0.514018182 | 0.005226394 |
| Pgr | 304.6324674 | 0.514009343 | 1.92E-05 |
| Syt12 | 606.4350727 | 0.513564233 | 3.74E-05 |
| Hspa12a | 9945.960901 | 0.513522346 | 2.10E-06 |
| Per1 | 2170.396359 | 0.513338719 | 0.00040738 |
| Jph3 | 7116.891693 | 0.512849927 | 1.29E-09 |
| Tpm1 | 3214.545355 | 0.512824693 | 7.80E-05 |
| Slc6a5 | 99.49338693 | 0.512268116 | 0.006200406 |
| Nek2 | 27.87748557 | 0.511985646 | 0.007407006 |
| Gria2 | 16401.27495 | 0.511608138 | 3.01E-07 |
| Phospho1 | 430.543996 | 0.511303522 | 0.002872451 |

| | | | |
|---|---|---|---|
| Tiam1 | 4432.125071 | 0.510988689 | 1.77E-07 |
| Map2k1 | 6640.415635 | 0.510929848 | 6.31E-11 |
| Enc1 | 13858.31557 | 0.510725866 | 0.00192837 |
| Dlg4 | 16820.21778 | 0.509845805 | 8.88E-09 |
| Palmd | 110.1625549 | 0.509349114 | 0.003160205 |
| Cacnb1 | 3424.208353 | 0.509267594 | 4.09E-09 |
| March11 | 71.2208775 | 0.509190029 | 0.002835569 |
| Myo5b | 3223.928536 | 0.509127134 | 0.002083283 |
| Scn8a | 10061.3986 | 0.508859399 | 9.65E-05 |
| Scn3b | 6616.481067 | 0.508468077 | 1.95E-06 |
| Cdk5r1 | 2291.014862 | 0.508300622 | 6.82E-08 |
| Mmp17 | 2389.512687 | 0.508236755 | 0.002589762 |
| Napb | 2504.298013 | 0.508150156 | 3.32E-05 |
| Cdk5r2 | 4518.63001 | 0.507831025 | 9.00E-05 |
| Cacnb3 | 3022.015453 | 0.507356949 | 1.47E-07 |
| Dclk1 | 27295.59628 | 0.507227786 | 0.000840025 |
| Ctxn1 | 6692.887699 | 0.50719912 | 0.000708477 |
| Gabra4 | 1405.540428 | 0.506874863 | 3.44E-09 |
| Kifc2 | 4444.203539 | 0.506546214 | 0.000155142 |
| Lin7a | 143.827923 | 0.506192596 | 0.000402077 |
| Dlk2 | 295.0080002 | 0.506160948 | 0.000154675 |
| Fam163b | 1645.261475 | 0.506151544 | 0.000442488 |
| Fam189b | 1262.404335 | 0.505974952 | 5.08E-06 |
| Crtac1 | 3085.791689 | 0.50589165 | 0.000135813 |
| Gabrb3 | 4024.853612 | 0.505454611 | 6.33E-06 |
| Lrrc7 | 4173.802616 | 0.505372723 | 1.78E-05 |
| Wnt10b | 22.10895302 | 0.504367551 | 0.009712406 |
| Nsf | 25122.98973 | 0.503895566 | 1.51E-07 |
| Fkbp5 | 1505.021012 | 0.503005631 | 9.78E-05 |
| Pcdha9 | 324.3553069 | 0.502539302 | 0.00281999 |
| Opcml | 2394.734889 | 0.501922309 | 0.002082636 |
| Cx3cl1 | 15435.14467 | 0.5018006 | 6.07E-05 |
| Camkk1 | 2642.452762 | 0.500751657 | 6.97E-09 |
| Klf16 | 1723.600363 | 0.500712636 | 2.08E-06 |
| Mef2a | 3642.517496 | 0.500537544 | 2.71E-05 |
| Per2 | 1520.278832 | 0.500127948 | 4.28E-05 |

| | | | |
|---|---|---|---|
| Dok6 | 1153.078366 | 0.499950474 | 2.19E-05 |
| Prkcz | 3339.447346 | 0.499483024 | 2.67E-06 |
| Ncdn | 38472.0715 | 0.499443092 | 9.15E-15 |
| Neu2 | 142.846867 | 0.499349389 | 0.007522331 |
| Tspan17 | 598.8617601 | 0.498828843 | 0.002612537 |
| Tex15 | 257.4680485 | 0.497930253 | 0.000168225 |
| Ptprj | 4870.73615 | 0.497502835 | 1.88E-06 |
| Rims1 | 6622.163002 | 0.496724689 | 0.000195093 |
| Arf3 | 26293.65129 | 0.496355038 | 6.13E-06 |
| LOC100910827 | 7184.564546 | 0.49563751 | 0.000142916 |
| Bcl11a | 2261.544589 | 0.495577099 | 0.000115728 |
| Lmtk2 | 9569.575349 | 0.494705036 | 7.90E-06 |
| Trerf1 | 1151.851265 | 0.494090699 | 2.39E-07 |
| Shc2 | 985.4434376 | 0.493794352 | 3.69E-06 |
| Zfhx2 | 2459.257642 | 0.493081174 | 3.53E-05 |
| Shank3 | 5227.139696 | 0.492886506 | 8.14E-06 |
| Hecw2 | 2179.093896 | 0.492312244 | 6.28E-05 |
| Pip5k1c | 9422.9986 | 0.491917346 | 4.16E-06 |
| Mctp2 | 65.2715631 | 0.491610631 | 0.009971191 |
| Dclk3 | 1279.65023 | 0.490907857 | 0.000175334 |
| Clec11a | 221.0985788 | 0.49028255 | 0.001866798 |
| Mical2 | 3162.899639 | 0.490208605 | 0.002069632 |
| Cntnap1 | 8939.666826 | 0.489866691 | 2.89E-07 |
| Fam81a | 1911.881377 | 0.489295939 | 0.004426108 |
| Lrrtm3 | 1801.207154 | 0.48903923 | 0.000102876 |
| Kcnt1 | 1079.94717 | 0.489025341 | 1.76E-07 |
| Nuak1 | 871.4413778 | 0.488973042 | 0.00094255 |
| Lrrc8b | 1193.778494 | 0.48866804 | 6.98E-05 |
| Cpne4 | 2129.648459 | 0.488034069 | 0.001281402 |
| Kcnip3 | 557.5978107 | 0.487806254 | 0.004387059 |
| Cnih2 | 3653.167348 | 0.487700991 | 6.34E-05 |
| Adcy1 | 5287.685135 | 0.487327424 | 0.002813424 |
| Cntn4 | 1728.901815 | 0.486905239 | 0.000113781 |
| Tmod1 | 726.8258051 | 0.486254128 | 0.000571969 |
| Shisa7 | 4079.53129 | 0.485626205 | 3.28E-06 |
| Dlgap2 | 2029.045578 | 0.485537821 | 0.000397486 |

| | | | |
|---|---|---|---|
| Atp2b2 | 24202.96758 | 0.485379004 | 4.77E-06 |
| Doc2b | 510.3187311 | 0.484500824 | 0.000189839 |
| Hcn1 | 1845.177638 | 0.484163036 | 0.000432929 |
| Golga7b | 422.1677102 | 0.48408434 | 0.002398872 |
| Sidt1 | 1294.646965 | 0.484026407 | 0.005636274 |
| Ddn | 11873.53023 | 0.483918928 | 0.003256305 |
| Cacna1d | 1412.356979 | 0.483677018 | 9.13E-06 |
| Pdzd2 | 2168.888335 | 0.482851891 | 0.000880091 |
| Rimbp2 | 4997.333776 | 0.482681038 | 0.000660179 |
| Mllt11 | 6706.919632 | 0.481605018 | 7.80E-07 |
| Dusp14 | 690.2613533 | 0.481399836 | 0.000296832 |
| Sorbs2 | 4779.111079 | 0.481122792 | 0.000310766 |
| Ppapdc2 | 962.4494822 | 0.481121942 | 0.000403668 |
| Ccne1 | 306.6417031 | 0.48049382 | 0.001593661 |
| Cdk17 | 5280.469632 | 0.480417258 | 5.19E-08 |
| Slc4a10 | 5553.764891 | 0.479850932 | 4.82E-05 |
| Shank2 | 5823.546287 | 0.479605435 | 2.63E-05 |
| Pde7b | 1256.379792 | 0.478949245 | 7.16E-06 |
| Fst | 10.31991761 | 0.478607184 | 0.017846658 |
| Kcnj9 | 1100.3095 | 0.477851264 | 0.001648985 |
| Ahr | 165.5648149 | 0.477639151 | 0.001806033 |
| Syn2 | 11164.41214 | 0.477561257 | 0.000276696 |
| Abcc8 | 540.8692807 | 0.477317192 | 0.000403668 |
| Ptk2b | 13028.04227 | 0.476747256 | 4.49E-06 |
| Rab3a | 18023.69153 | 0.476536469 | 4.85E-07 |
| Dgki | 2053.73678 | 0.476355203 | 2.09E-06 |
| Cbln2 | 693.9441455 | 0.475913823 | 0.007131455 |
| Mark2 | 2335.00794 | 0.47566004 | 9.71E-07 |
| Nrn1 | 737.4367657 | 0.475643768 | 0.012150339 |
| Frmpd4 | 4174.382252 | 0.475335972 | 1.80E-05 |
| B4galt6 | 5211.536574 | 0.474385367 | 4.83E-06 |
| Mmd | 4118.048742 | 0.474334098 | 0.000405326 |
| Rap1gap2 | 2973.712976 | 0.474224532 | 0.001171999 |
| Rps6ka2 | 2464.172646 | 0.474211903 | 0.000159031 |
| Orai2 | 1609.160539 | 0.473584337 | 0.000849828 |
| Dgkb | 9484.213167 | 0.473256336 | 0.000116419 |

| | | | |
|---|---|---|---|
| Grin2b | 9287.764671 | 0.472978024 | 0.000185068 |
| Pthlh | 77.04819435 | 0.472758488 | 0.007041999 |
| Cplx2 | 5065.699862 | 0.472584473 | 6.94E-06 |
| Kcnj2 | 121.1224427 | 0.47250651 | 0.00638963 |
| Lgi1 | 1648.202492 | 0.472043155 | 1.64E-07 |
| Limd2 | 502.3583391 | 0.471073909 | 0.000323009 |
| Wnt9b | 127.0460373 | 0.470689506 | 0.0131232 |
| Sh3bp1 | 408.4218814 | 0.470323729 | 0.000763393 |
| Cap2 | 4535.401776 | 0.469974143 | 4.70E-05 |
| Nppa | 105.1783013 | 0.469163057 | 0.014338011 |
| Cacng3 | 633.4451982 | 0.468715575 | 5.36E-05 |
| Tbpl1 | 604.882079 | 0.468325131 | 2.95E-06 |
| Xkr7 | 259.1218333 | 0.468236422 | 0.00978249 |
| Cntn5 | 246.6946873 | 0.468156753 | 0.002704887 |
| Sept3 | 2372.047263 | 0.466582215 | 7.00E-07 |
| Cacna1c | 2405.318727 | 0.466495101 | 1.46E-05 |
| Csnk1g1 | 243.2207545 | 0.466368792 | 0.001049459 |
| Shank1 | 23262.83338 | 0.46521137 | 0.000314143 |
| Nos1ap | 2105.13283 | 0.465102516 | 0.000515658 |
| Rnf112 | 2309.940283 | 0.465086717 | 2.76E-05 |
| Slc6a17 | 17380.90273 | 0.464986471 | 3.97E-05 |
| Wbscr22 | 295.2949673 | 0.464449599 | 7.23E-05 |
| Lyst | 2209.872026 | 0.463773276 | 7.10E-05 |
| Gls | 8185.597967 | 0.463570778 | 6.18E-06 |
| Uck2 | 516.570178 | 0.462714929 | 0.000468891 |
| Gal3st3 | 772.6081137 | 0.462562249 | 7.63E-06 |
| Bend6 | 1238.341635 | 0.462386912 | 7.75E-05 |
| Cntnap5b | 388.3690169 | 0.462001096 | 0.004960025 |
| Lancl1 | 2407.92001 | 0.461687814 | 2.24E-07 |
| Dlg3 | 5057.363877 | 0.461641814 | 3.64E-07 |
| Mgst3 | 947.1427657 | 0.461630031 | 0.000273224 |
| Lppr1 | 296.7316327 | 0.46145022 | 0.003663646 |
| Rarb | 962.3814887 | 0.46093858 | 8.25E-05 |
| Neurl1b | 360.5294051 | 0.460826369 | 0.002985367 |
| Phlda1 | 1333.955221 | 0.460590418 | 0.000100913 |
| Rnf208 | 1681.375113 | 0.459890854 | 1.16E-07 |

| | | | |
|---|---|---|---|
| Syt4 | 7170.281079 | 0.459880724 | 1.33E-07 |
| Ube2ql1 | 5057.510743 | 0.45958669 | 0.000405328 |
| Chm | 681.7263354 | 0.459369924 | 1.34E-06 |
| Ppp1r9a | 5252.033452 | 0.458901231 | 4.94E-05 |
| Cacna1g | 3169.882935 | 0.458630723 | 0.000108076 |
| Ppp3cb | 6937.573796 | 0.458519792 | 4.29E-05 |
| Ndrg4 | 57815.4121 | 0.458138026 | 3.72E-07 |
| Slitrk4 | 1432.993047 | 0.458086995 | 3.42E-06 |
| Pcdha3 | 438.4859025 | 0.458073293 | 0.00152285 |
| Slc35f1 | 5241.885247 | 0.458021453 | 5.93E-07 |
| Xkr4 | 2154.341432 | 0.457971533 | 0.00021015 |
| Lrg1 | 12.81364818 | 0.457817833 | 0.019063743 |
| Usp31 | 2719.59613 | 0.457772249 | 1.68E-06 |
| Noct | 1332.974678 | 0.45771904 | 5.04E-06 |
| Stxbp5 | 3839.927514 | 0.457654602 | 1.73E-05 |
| Fbp2 | 3.075999313 | 0.457174745 | 0.015786916 |
| Nexn | 356.922281 | 0.457156043 | 0.005912907 |
| Lypd1 | 2649.649068 | 0.457060744 | 6.18E-06 |
| B3gat1 | 3264.285077 | 0.456227678 | 0.000353009 |
| Bdnf | 266.2326796 | 0.456215095 | 0.016923864 |
| Scn2b | 2274.139489 | 0.455841896 | 3.25E-08 |
| Mal2 | 2506.797286 | 0.45572758 | 0.010788985 |
| LOC684871 | 33.93883976 | 0.455149828 | 0.012186494 |
| Grm5 | 4703.834607 | 0.45496906 | 6.25E-07 |
| Kcnc3 | 3413.554141 | 0.454894351 | 0.000411735 |
| Fam49a | 2929.503321 | 0.454794157 | 7.23E-05 |
| Atp1a1 | 17062.30251 | 0.454575844 | 0.000397322 |
| Kcnb1 | 3176.191908 | 0.454544601 | 0.000460983 |
| Scn2a | 18054.78328 | 0.453988397 | 7.80E-05 |
| Olfm1 | 12980.76413 | 0.453947177 | 0.000970463 |
| Nedd4l | 4018.294229 | 0.453607137 | 1.31E-05 |
| Best3 | 10.32162706 | 0.453261769 | 0.020413499 |
| Arhgap20 | 1971.47105 | 0.452521125 | 4.31E-06 |
| Kalrn | 7581.705524 | 0.451921764 | 0.004433611 |
| Rab6b | 26997.76202 | 0.451270687 | 8.29E-07 |
| Slc35d3 | 699.6527435 | 0.450975657 | 0.000820231 |

| | | | |
|---|---|---|---|
| Ncs1 | 5254.151603 | 0.450960887 | 6.03E-07 |
| Prkar2b | 2796.458808 | 0.450831031 | 4.75E-10 |
| Arhgef25 | 1063.305297 | 0.450534313 | 1.02E-06 |
| Sptbn2 | 18376.42686 | 0.450354004 | 0.009469222 |
| Acot4 | 96.69043165 | 0.449471153 | 0.008667542 |
| Rapgef2 | 7225.640019 | 0.449213951 | 4.33E-07 |
| Pcdha5 | 372.2576022 | 0.449011566 | 0.001395812 |
| Gpr61 | 502.7675555 | 0.448912024 | 2.83E-05 |
| Hsph1 | 11546.70461 | 0.448808014 | 3.46E-08 |
| Plekha5 | 1435.557174 | 0.44878548 | 6.38E-08 |
| Add2 | 4511.036479 | 0.448168976 | 2.54E-06 |
| Galnt14 | 1017.84107 | 0.448083217 | 0.005602464 |
| Cacng8 | 1971.458707 | 0.447480082 | 0.000361539 |
| St6gal2 | 470.579669 | 0.446443213 | 0.009193694 |
| Kitlg | 480.8570585 | 0.446278632 | 0.000253827 |
| Dapk1 | 4173.343172 | 0.44611129 | 4.21E-08 |
| Rims2 | 3333.588706 | 0.445915993 | 9.48E-05 |
| Agtpbp1 | 7214.203267 | 0.445621972 | 6.12E-05 |
| Tasp1 | 316.907582 | 0.445398748 | 0.000209464 |
| Kctd13 | 1654.365188 | 0.444925448 | 0.000171835 |
| Sik1 | 538.9062658 | 0.444242902 | 0.001885195 |
| Kcnab2 | 3882.793853 | 0.444142374 | 3.48E-05 |
| Celf3 | 2039.185176 | 0.444070645 | 3.71E-05 |
| Ncald | 9258.003676 | 0.443583847 | 0.006963741 |
| Drd2 | 1655.698278 | 0.443249458 | 0.000136369 |
| Cacng2 | 938.8362358 | 0.442794465 | 1.06E-06 |
| Nlk | 3104.862591 | 0.442639384 | 0.000101701 |
| Nefl | 9362.630158 | 0.442614417 | 0.00136636 |
| Atl1 | 3923.394844 | 0.442534656 | 4.49E-06 |
| Jdp2 | 324.36569 | 0.442183497 | 0.002116167 |
| Dcn | 12.33981021 | 0.442162223 | 0.020807307 |
| Opn3 | 396.5479666 | 0.441927227 | 9.63E-05 |
| Lrp1b | 2985.697349 | 0.441681159 | 0.000272237 |
| Cacna1a | 6229.791091 | 0.441569216 | 0.000102876 |
| Kcnk2 | 1060.751044 | 0.440636848 | 3.32E-05 |
| Fbxo41 | 3351.723453 | 0.440403894 | 2.08E-07 |

| | | | |
|---|---|---|---|
| Hpcal4 | 1941.582591 | 0.440045641 | 0.009457365 |
| Gria1 | 6767.768493 | 0.439951676 | 1.28E-06 |
| Slc25a22 | 4460.672923 | 0.439663845 | 0.002432611 |
| Inpp5j | 1623.425107 | 0.438973177 | 0.000137674 |
| Slitrk3 | 1509.410976 | 0.438711331 | 2.18E-09 |
| Gls2 | 709.0363434 | 0.43839414 | 0.000371268 |
| Adap1 | 2458.322013 | 0.438150546 | 0.000127442 |
| Sipa1l1 | 8397.216829 | 0.43806893 | 8.32E-06 |
| Mat2b | 2701.38744 | 0.437840785 | 0.00125523 |
| Kazn | 3434.2765 | 0.437650723 | 2.70E-05 |
| Il1rapl1 | 240.0865079 | 0.437056715 | 0.002813424 |
| Ryr2 | 6827.514647 | 0.437045891 | 0.000403668 |
| Mapk8 | 4161.551441 | 0.436643819 | 0.001053184 |
| Ywhah | 27498.36832 | 0.436587059 | 9.77E-05 |
| Pgm2l1 | 7336.400167 | 0.436582924 | 0.000592812 |
| Josd1 | 1127.149307 | 0.436489198 | 2.70E-05 |
| Snurf | 15937.8838 | 0.436004244 | 1.57E-07 |
| Igf1 | 180.6137607 | 0.435961842 | 0.006963741 |
| Dnajb5 | 2368.390911 | 0.435597442 | 4.46E-05 |
| Mpp3 | 1205.098969 | 0.435372553 | 3.53E-05 |
| Syndig1 | 553.8217192 | 0.434659421 | 0.000638603 |
| Stambpl1 | 669.9683396 | 0.434565955 | 0.000148927 |
| Aak1 | 17130.67296 | 0.434412587 | 9.28E-07 |
| Grin1 | 18248.42951 | 0.434328368 | 6.94E-06 |
| St8sia3 | 1748.75832 | 0.433479193 | 0.000376756 |
| Gins3 | 92.00674769 | 0.433105791 | 0.00638576 |
| Mthfd1l | 758.7294416 | 0.432434658 | 0.000110928 |
| Asb1 | 787.1959484 | 0.432410787 | 1.01E-05 |
| Olfm3 | 1961.307554 | 0.431901488 | 2.23E-05 |
| Fam126b | 728.6192572 | 0.431465516 | 2.33E-05 |
| Ctsk | 31.90867812 | 0.431064874 | 0.018830345 |
| Calm1 | 33961.12454 | 0.431007513 | 4.50E-06 |
| Pcdha6 | 442.1757022 | 0.430986367 | 0.002549284 |
| Runx1t1 | 1377.167156 | 0.430953452 | 1.08E-06 |
| Cckbr | 658.225142 | 0.430166839 | 0.01670553 |
| Atp6v1g2 | 9310.4147 | 0.430013087 | 2.09E-06 |

| | | | |
|---|---|---|---|
| Vsnl1 | 13767.91356 | 0.429423911 | 2.74E-05 |
| Brsk1 | 6945.928936 | 0.429136349 | 1.86E-05 |
| Panx1 | 649.2275597 | 0.429086559 | 0.000185121 |
| Gpr63 | 174.0695988 | 0.429071227 | 0.005347779 |
| Rnf165 | 861.8103505 | 0.428443038 | 0.001368313 |
| Dnm1 | 28544.60733 | 0.42824082 | 9.30E-06 |
| Btg2 | 576.9494009 | 0.428220901 | 0.003110546 |
| Thy1 | 13277.01905 | 0.42780059 | 0.000186501 |
| RGD1564664 | 1201.637324 | 0.427795581 | 2.47E-06 |
| Oprk1 | 104.4977167 | 0.427782183 | 0.009955071 |
| Gpr12 | 346.1149319 | 0.427594939 | 0.000282699 |
| Erbb4 | 3173.64337 | 0.427226785 | 1.95E-10 |
| Zdhhc23 | 213.1447948 | 0.42696929 | 0.008308536 |
| Gucy1b3 | 5489.256602 | 0.426828292 | 1.53E-11 |
| Sstr4 | 214.3837073 | 0.426327707 | 0.005422926 |
| Fosl2 | 179.7100472 | 0.425837332 | 0.017534581 |
| Snap91 | 9534.954723 | 0.42562244 | 3.74E-05 |
| Kcnh5 | 446.4772838 | 0.425424695 | 0.008909046 |
| Dok4 | 320.4849951 | 0.425389872 | 0.002458007 |
| Shisa6 | 355.2228779 | 0.425201386 | 0.012768029 |
| Acsl4 | 3548.646086 | 0.424534404 | 7.83E-05 |
| Chgb | 13911.88822 | 0.42432774 | 1.01E-06 |
| Cps1 | 41.31804706 | 0.424164175 | 0.020811156 |
| Pde1a | 3542.166041 | 0.424138436 | 0.007815464 |
| Plcxd3 | 1186.009144 | 0.424038821 | 0.004387059 |
| Matk | 1460.228422 | 0.423662247 | 2.20E-05 |
| Hs6st2 | 1758.874293 | 0.423145257 | 3.72E-07 |
| Cyld | 2840.238405 | 0.422913033 | 9.49E-10 |
| Adam19 | 387.4307195 | 0.422739672 | 0.000310766 |
| Abcc5 | 2902.305545 | 0.422726449 | 1.77E-06 |
| Akap6 | 12699.44583 | 0.42264963 | 0.000239796 |
| Mtmr7 | 1233.610858 | 0.422480943 | 1.32E-07 |
| RGD1311739 | 1744.629014 | 0.422056955 | 6.02E-06 |
| Kcnn1 | 242.600633 | 0.420927307 | 0.009232376 |
| Cbfa2t3 | 1012.581338 | 0.420882486 | 0.000271796 |
| Mpped1 | 5169.547379 | 0.420779396 | 0.009082755 |

| | | | |
|---|---|---|---|
| Celf4 | 18132.53643 | 0.420405801 | 1.24E-07 |
| Adarb1 | 7247.710689 | 0.420213364 | 0.001251548 |
| Clvs1 | 1187.20804 | 0.420106962 | 5.59E-06 |
| Necab2 | 1834.509526 | 0.420104179 | 2.63E-05 |
| Pmepa1 | 403.754529 | 0.419797059 | 4.12E-05 |
| Col18a1 | 146.6931644 | 0.419425904 | 0.0234476 |
| Atp6v1b2 | 16603.76194 | 0.419084871 | 1.86E-05 |
| Fam110b | 1095.80139 | 0.41904249 | 3.50E-05 |
| Pip4k2c | 1407.995943 | 0.419031286 | 0.000181488 |
| Il10ra | 79.03241906 | 0.418721338 | 0.006765111 |
| Htatip2 | 123.3261254 | 0.418454331 | 0.01024695 |
| Dffa | 623.7924269 | 0.418433536 | 0.00018426 |
| Cdkl2 | 678.9027173 | 0.418391601 | 0.00258672 |
| Sh3gl2 | 4360.03441 | 0.418343307 | 0.001609267 |
| Gfod1 | 696.5510251 | 0.418144418 | 0.003284137 |
| Dlgap1 | 12883.62187 | 0.417968814 | 0.0024962 |
| Akap1 | 1531.743015 | 0.417948943 | 3.94E-07 |
| Pdpk1 | 2081.33563 | 0.417917281 | 1.29E-09 |
| Kl | 85.70817809 | 0.417865382 | 0.021734435 |
| Syt7 | 1785.491414 | 0.417213905 | 5.08E-05 |
| Fam65b | 1679.830535 | 0.41693405 | 0.000818392 |
| Fbxo34 | 1479.058752 | 0.4168811 | 3.99E-05 |
| Gga3 | 1575.674452 | 0.416856867 | 1.39E-06 |
| Trhde | 940.8934356 | 0.416797846 | 0.000862299 |
| Camta2 | 7059.731487 | 0.41657041 | 1.96E-07 |
| Mapk1 | 16675.37837 | 0.416254134 | 9.28E-07 |
| Plag1 | 104.5416349 | 0.41594948 | 0.006120222 |
| Cttnbp2 | 2929.006317 | 0.415418331 | 0.000381941 |
| Stxbp1 | 28315.58082 | 0.4153494 | 9.70E-06 |
| Ugcg | 4073.212994 | 0.4150333 | 1.94E-06 |
| Tub | 1452.680276 | 0.414552269 | 4.15E-06 |
| Pcdha4 | 476.6518951 | 0.41415746 | 0.003931994 |
| Myh7 | 242.4345364 | 0.414127842 | 0.004452384 |
| Myo5a | 15789.92465 | 0.414015468 | 0.000165067 |
| Plcl2 | 2242.087559 | 0.413976146 | 7.80E-05 |
| R3hdm4 | 2817.487118 | 0.413649897 | 0.000959262 |

| | | | |
|---|---|---|---|
| Gabrg2 | 1773.366376 | 0.413628387 | 2.07E-05 |
| Dlg2 | 6290.830135 | 0.413402542 | 7.90E-09 |
| Syt1 | 25678.66141 | 0.41330148 | 5.17E-05 |
| Cort | 135.9356239 | 0.413236502 | 0.022208263 |
| Arhgef9 | 6964.49217 | 0.413153706 | 9.44E-11 |
| Vstm2b | 1517.106299 | 0.412967607 | 0.004868903 |
| Srrm3 | 1790.181204 | 0.412671051 | 2.18E-06 |
| Ehbp1l1 | 643.9876314 | 0.411793454 | 0.000743101 |
| Vwa5b2 | 379.4059941 | 0.411715078 | 0.002345947 |
| Prr7 | 422.0022505 | 0.411713633 | 0.001442095 |
| Slc2a3 | 1723.864552 | 0.411505622 | 0.000127468 |
| Kcnj6 | 4455.326605 | 0.411492998 | 0.018821957 |
| Ptprk | 898.2125718 | 0.411369518 | 0.002927108 |
| Klhl29 | 2317.392661 | 0.410963576 | 4.04E-07 |
| Fhod3 | 3358.712874 | 0.410684392 | 0.000893991 |
| Kdm4d | 51.44846139 | 0.410587791 | 0.016892639 |
| RGD1565819 | 665.4387263 | 0.410239228 | 0.012081535 |
| Kcnv1 | 1007.402165 | 0.41017104 | 0.023523193 |
| Lrtm2 | 1188.170835 | 0.409995595 | 4.88E-05 |
| Nalcn | 5223.705955 | 0.409935158 | 1.14E-06 |
| Cdh9 | 569.7403542 | 0.409731289 | 0.011425258 |
| Ppip5k1 | 2871.731212 | 0.409323043 | 4.36E-08 |
| Fam65a | 4070.391804 | 0.408985863 | 0.000120163 |
| Myh6 | 35.33839095 | 0.408778413 | 0.025829894 |
| Sult4a1 | 8953.288063 | 0.408483431 | 8.27E-06 |
| Btbd10 | 1620.329638 | 0.408186835 | 1.23E-05 |
| Rasgrp2 | 660.1175317 | 0.408135327 | 3.49E-06 |
| Nrg2 | 275.8378025 | 0.408128743 | 0.000467677 |
| Csrnp3 | 1556.920859 | 0.408100305 | 2.54E-06 |
| Mme | 287.9340273 | 0.407988971 | 0.006950792 |
| Scg5 | 8973.324883 | 0.40785541 | 2.95E-06 |
| Necab3 | 391.6507543 | 0.407428905 | 0.013983597 |
| Maf | 484.0998138 | 0.407289032 | 0.011333201 |
| Tmem25 | 856.0402015 | 0.407166565 | 0.000757764 |
| Crtc1 | 2891.981055 | 0.406644003 | 1.41E-07 |
| Dcbld2 | 678.4517307 | 0.406311033 | 0.000156798 |

| | | | |
|---|---|---|---|
| Dnajc6 | 12597.62792 | 0.406267205 | 7.79E-05 |
| Slc12a5 | 16698.60919 | 0.406218409 | 9.73E-06 |
| Dusp8 | 2779.8112 | 0.406149453 | 5.37E-08 |
| Ldlrad4 | 391.4797645 | 0.405954307 | 0.00423437 |
| Ppfia2 | 2800.792203 | 0.405549197 | 1.62E-06 |
| Nav3 | 3743.14924 | 0.405224251 | 0.003796032 |
| Kcnab3 | 475.2221422 | 0.405213569 | 0.016987813 |
| RGD1304884 | 10881.9307 | 0.405160248 | 4.42E-06 |
| Sncb | 15124.44238 | 0.404611069 | 8.71E-05 |
| Htr2a | 516.3394387 | 0.404201925 | 0.021395048 |
| Epcam | 31.56400382 | 0.403995334 | 0.02536944 |
| Zbtb11 | 2118.1638 | 0.403890833 | 7.01E-06 |
| Lingo1 | 5294.231542 | 0.40379952 | 0.018590663 |
| Slc36a1 | 1405.661902 | 0.403797485 | 0.001013191 |
| Wnt9a | 247.8998265 | 0.403653547 | 0.02406917 |
| Car11 | 1484.361787 | 0.403495949 | 0.000302271 |
| Syt3 | 1206.328162 | 0.403315541 | 4.09E-05 |
| Ppfia3 | 5480.516846 | 0.40304114 | 0.000124616 |
| Pcgf3 | 644.8365459 | 0.402067554 | 2.59E-06 |
| Acta1 | 149.578658 | 0.401855426 | 0.01992462 |
| Zfp280b | 694.6085155 | 0.401712788 | 1.39E-05 |
| Ppp1r12b | 6506.195403 | 0.401605804 | 7.54E-06 |
| Kcnk6 | 18.96085031 | 0.401565551 | 0.027512553 |
| Fgf16 | 3.594742415 | 0.401426399 | 0.022583944 |
| Vamp2 | 40352.36265 | 0.401359121 | 5.79E-06 |
| Slc8a1 | 9661.268105 | 0.401354895 | 0.001764185 |
| Hace1 | 1295.376566 | 0.401110548 | 0.000271886 |
| Slc16a7 | 306.4273668 | 0.401106327 | 0.005246804 |
| Syn1 | 21440.41948 | 0.40082398 | 0.000308642 |
| Mef2d | 3839.655417 | 0.400526073 | 5.05E-07 |
| Ip6k2 | 1478.081133 | 0.400029728 | 3.01E-05 |
| Tanc2 | 6977.388491 | 0.399843882 | 9.62E-06 |
| Fam212b | 2128.099603 | 0.399498925 | 0.021815788 |
| Mfsd4 | 399.2449189 | 0.399498557 | 0.018439216 |
| Grem1 | 160.5981528 | 0.398866134 | 0.027542862 |
| Dopey2 | 2617.558281 | 0.398825654 | 8.25E-05 |

| | | | |
|---|---|---|---|
| Begain | 2319.821104 | 0.398649962 | 1.36E-05 |
| Rph3a | 8263.511759 | 0.398555268 | 0.001375122 |
| LOC100912071 | 167.8871289 | 0.398521933 | 0.00959119 |
| Ism1 | 51.4856241 | 0.398400127 | 0.028056057 |
| Man1a1 | 824.5609165 | 0.39816259 | 0.000317929 |
| Erc2 | 2855.622217 | 0.398085947 | 0.011885047 |
| Sgtb | 2459.294687 | 0.397888336 | 9.13E-06 |
| Tnnc2 | 328.8618117 | 0.397822777 | 0.015937576 |
| Ntng1 | 579.7359526 | 0.397704424 | 0.00607444 |
| Hif3a | 89.51129811 | 0.39743974 | 0.028621306 |
| Prkar1b | 10222.57175 | 0.397112133 | 1.23E-07 |
| Gpr3 | 75.75283489 | 0.397101243 | 0.026526406 |
| Kif5c | 19340.74563 | 0.396974315 | 0.000105043 |
| Rassf5 | 1181.720356 | 0.396973998 | 0.000489821 |
| Pygl | 140.2160086 | 0.396896146 | 0.00607444 |
| Gpr85 | 648.9876667 | 0.396744957 | 0.001861534 |
| Impdh1 | 961.816471 | 0.396534165 | 0.00031617 |
| Epb41l1 | 24200.34524 | 0.395781724 | 8.01E-09 |
| Bsn | 27122.00408 | 0.395773551 | 0.024905504 |
| Dusp2 | 48.02204652 | 0.395711473 | 0.028787875 |
| Klhdc8a | 1008.091973 | 0.395534532 | 0.000153933 |
| Gpr26 | 330.9322396 | 0.395453397 | 0.010292415 |
| Map9 | 2921.778132 | 0.39533051 | 4.94E-05 |
| Lynx1 | 6640.967448 | 0.394939544 | 0.000568072 |
| N4bp3 | 707.8317745 | 0.394859853 | 0.010655846 |
| Adam23 | 4626.20334 | 0.3944128 | 6.91E-06 |
| Rgs4 | 3657.083912 | 0.394378891 | 0.023824043 |
| Slitrk5 | 2331.857335 | 0.394242808 | 5.34E-07 |
| Gdap2 | 746.6545102 | 0.393879121 | 1.84E-05 |
| Clvs2 | 286.1684588 | 0.393481939 | 0.000351559 |
| Ggt7 | 3065.03007 | 0.393259707 | 3.09E-05 |
| Atg16l1 | 1023.874289 | 0.393116115 | 7.22E-06 |
| Gda | 3061.484824 | 0.393001521 | 0.016190163 |
| Nsmf | 7889.476236 | 0.392899727 | 0.015292635 |
| Dusp4 | 184.4651065 | 0.392728718 | 0.01237592 |
| Grm7 | 1009.720374 | 0.392475667 | 6.24E-05 |

| | | | |
|---|---|---|---|
| Unc45b | 32.71049787 | 0.392419807 | 0.029424024 |
| Ankrd34c | 278.108079 | 0.392296527 | 0.004186676 |
| Fbxl15 | 458.12678 | 0.392208536 | 1.42E-06 |
| Spns2 | 702.9917961 | 0.392168683 | 0.000553803 |
| Tbc1d9 | 4529.306073 | 0.392110596 | 1.54E-06 |
| Chrnb2 | 1890.07321 | 0.391832266 | 1.05E-06 |
| Thbs1 | 384.6425477 | 0.391830396 | 0.029651626 |
| Pi4ka | 13579.43838 | 0.391781227 | 6.43E-06 |
| Zbtb16 | 1009.806106 | 0.391758509 | 0.007229442 |
| Stac2 | 1238.982239 | 0.39170574 | 0.006600703 |
| Pask | 133.9804736 | 0.39164772 | 0.010565099 |
| Prkg1 | 121.1345134 | 0.390943308 | 0.017526882 |
| Pdxp | 3506.174037 | 0.390864542 | 8.60E-06 |
| Atpaf1 | 1197.383802 | 0.390760774 | 2.81E-05 |
| Camk2n1 | 1683.695295 | 0.390671991 | 0.00196365 |
| Ensa | 6556.711041 | 0.390629358 | 0.001254391 |
| Micu3 | 1020.958317 | 0.390585824 | 9.29E-06 |
| Synj2 | 1800.589623 | 0.38989071 | 0.026978539 |
| Ndufaf6 | 254.8883681 | 0.389806564 | 7.03E-05 |
| Serpini1 | 5792.009715 | 0.389680919 | 0.020838537 |
| Rundc3a | 5386.769767 | 0.389548301 | 1.62E-05 |
| Rab26 | 779.3191525 | 0.389446279 | 0.012837454 |
| Eef1a2 | 24216.74857 | 0.389391409 | 6.73E-07 |
| Chac1 | 439.4326612 | 0.389043118 | 9.60E-05 |
| Adcy3 | 2905.075783 | 0.388844164 | 0.000172585 |
| Nptxr | 16618.48552 | 0.388386385 | 0.023823843 |
| Atcay | 6489.525244 | 0.388344909 | 5.96E-06 |
| Lrrc4c | 1917.838826 | 0.388299313 | 2.84E-06 |
| Dixdc1 | 2058.102201 | 0.38797812 | 0.000342452 |
| Rab3c | 3558.846244 | 0.386829387 | 1.91E-05 |
| Rpusd1 | 1056.885833 | 0.38682492 | 2.38E-06 |
| Unc13a | 5456.167253 | 0.386726418 | 0.003718108 |
| Fam57b | 837.358775 | 0.386713449 | 0.00071017 |
| Stx1b | 8383.06184 | 0.386653923 | 2.97E-08 |
| Doc2a | 566.9573027 | 0.386610011 | 0.026548864 |
| Sstr1 | 819.0482359 | 0.386468931 | 0.013421627 |

| | | | |
|---|---|---|---|
| Map2 | 45076.98642 | 0.386140465 | 0.001381774 |
| Habp4 | 3736.155113 | 0.386134232 | 1.30E-05 |
| Cacna1i | 2772.387257 | 0.386111962 | 0.003278267 |
| Stard4 | 525.4810447 | 0.385588125 | 0.010566642 |
| Npy1r | 236.7229017 | 0.385466039 | 0.015927298 |
| Ppp1r7 | 3184.879703 | 0.384931005 | 7.29E-05 |
| Slc16a14 | 178.806648 | 0.384923685 | 0.011304375 |
| Rpp25 | 380.3264051 | 0.384787164 | 0.001914822 |
| Lpgat1 | 3755.462751 | 0.384389829 | 3.36E-06 |
| Tmem63c | 1107.136138 | 0.384311484 | 0.000880091 |
| Gcnt2 | 199.4302676 | 0.383936582 | 0.021533213 |
| Slc6a15 | 2000.374173 | 0.38373076 | 1.15E-05 |
| Dlx5 | 475.2319091 | 0.383159165 | 0.004525379 |
| Tmeff2 | 1425.431146 | 0.383135241 | 6.76E-05 |
| Ppp1r1b | 12849.45472 | 0.382941802 | 0.000230931 |
| Snph | 6986.600936 | 0.382716983 | 2.57E-06 |
| St3gal5 | 2739.889706 | 0.382445992 | 0.001976012 |
| Wdr37 | 1652.719776 | 0.38227294 | 4.07E-05 |
| Mafg | 1168.429337 | 0.382232763 | 1.98E-05 |
| Slc8a2 | 7334.349483 | 0.38210447 | 0.011563061 |
| Nsg2 | 25229.31992 | 0.381750356 | 6.74E-08 |
| Sema6b | 4052.215526 | 0.381424799 | 2.47E-05 |
| Ttc9b | 1753.98553 | 0.381301373 | 0.021420588 |
| Miat | 1362.982756 | 0.381106874 | 0.003952957 |
| Rbfox2 | 2049.24637 | 0.381027031 | 1.47E-09 |
| Gnaz | 1342.726663 | 0.380545699 | 0.000133002 |
| Ank3 | 15181.78252 | 0.379835072 | 8.20E-05 |
| Ppp1r13b | 2576.640319 | 0.379271181 | 3.15E-05 |
| Tsc22d1 | 10641.77019 | 0.379258224 | 3.97E-08 |
| Map1b | 100364.3537 | 0.379042656 | 0.000985354 |
| Cipc | 2537.10754 | 0.378869063 | 8.78E-08 |
| Pip5k1a | 1855.755668 | 0.378318803 | 0.000137054 |
| Sema4f | 1527.039491 | 0.378193377 | 6.24E-05 |
| Tcap | 60.04183579 | 0.377903174 | 0.033393719 |
| Cacna2d3 | 4207.313942 | 0.377191388 | 2.85E-06 |
| Jph1 | 180.3263118 | 0.377114261 | 0.026812743 |

| | | | |
|---|---|---|---|
| Slc30a3 | 480.6033797 | 0.377023013 | 0.03404123 |
| Scrt1 | 1278.054849 | 0.376927549 | 0.015900006 |
| Pcdhga2 | 1642.597375 | 0.37689449 | 0.000723934 |
| Atp6v1c1 | 4569.822095 | 0.376628866 | 9.56E-06 |
| Htr2c | 3571.271147 | 0.376581861 | 0.002456352 |
| Cds1 | 891.3665171 | 0.37648849 | 0.000207618 |
| Sertad4 | 541.6449117 | 0.376244791 | 0.001057389 |
| Rsrp1 | 2569.806839 | 0.375811061 | 0.011723743 |
| Wdr7 | 6802.41092 | 0.375651025 | 1.86E-05 |
| Kcnk12 | 215.375492 | 0.375578859 | 0.024817024 |
| Dock9 | 3613.231505 | 0.375573124 | 0.002050956 |
| Cbr3 | 216.2662927 | 0.375337626 | 0.01323774 |
| Gpcpd1 | 974.5758325 | 0.375187808 | 0.000128471 |
| Lrfn2 | 646.3553386 | 0.375034306 | 0.008275526 |
| Eno2 | 28884.23637 | 0.37497231 | 1.41E-07 |
| Spock3 | 3465.979301 | 0.374874335 | 0.000489681 |
| Fgf13 | 1710.457148 | 0.374732842 | 7.68E-05 |
| Evl | 3993.901136 | 0.374703366 | 1.13E-08 |
| Radil | 450.5076316 | 0.374235209 | 0.000532401 |
| Capn12 | 30.67583707 | 0.373779944 | 0.036331155 |
| Robo2 | 4754.32813 | 0.373742518 | 0.004158527 |
| Egfl7 | 75.62314992 | 0.373653641 | 0.028455238 |
| Syt5 | 2274.888506 | 0.373219371 | 0.007966606 |
| Osbpl8 | 5261.800456 | 0.37310275 | 2.53E-07 |
| Atp1b1 | 38418.75259 | 0.372421368 | 0.000332024 |
| Smap2 | 7327.355582 | 0.371801154 | 2.03E-05 |
| Parp6 | 1555.57077 | 0.371745813 | 0.000202309 |
| Slc16a5 | 1.730267335 | 0.371340399 | 0.0194991 |
| RGD1305733 | 6471.067534 | 0.371196562 | 4.41E-05 |
| Nptx2 | 1206.600005 | 0.371054127 | 0.036892743 |
| Oxr1 | 12267.54507 | 0.370948549 | 0.001806033 |
| Sh3bgrl3 | 1886.804373 | 0.370940078 | 0.00137814 |
| Srf | 1921.547894 | 0.370897162 | 0.000363725 |
| Prkch | 314.4142843 | 0.370415238 | 0.013988765 |
| Sv2c | 2210.141846 | 0.370311999 | 0.001030091 |
| Plekha8 | 454.1908934 | 0.370003118 | 0.000387782 |

| | | | |
|---|---|---|---|
| Ttpal | 1479.20902 | 0.369764336 | 1.75E-05 |
| Tlk1 | 2841.734155 | 0.369471354 | 3.91E-05 |
| Map3k13 | 867.9413398 | 0.369381731 | 0.000419923 |
| Dlgap4 | 5701.785036 | 0.369241714 | 0.000139644 |
| Kctd17 | 1454.066287 | 0.368925612 | 1.37E-05 |
| Ccdc132 | 1870.072 | 0.368424843 | 7.72E-06 |
| Ptpru | 1362.1352 | 0.36816174 | 0.015292635 |
| Dusp7 | 1716.750031 | 0.367965366 | 0.00138938 |
| Cacna1b | 4920.412511 | 0.367960177 | 0.000308709 |
| Myh8 | 84.11867935 | 0.36773609 | 0.037924697 |
| Dcaf6 | 2742.455037 | 0.367536072 | 0.000731578 |
| Meis2 | 5020.62155 | 0.367365448 | 3.68E-06 |
| Ttl | 1004.925806 | 0.367001852 | 0.000902908 |
| Ccdc85a | 580.7354562 | 0.366650277 | 0.000274614 |
| Sestd1 | 1203.590595 | 0.366592328 | 0.000569364 |
| Pcdhb21 | 212.3040649 | 0.366402586 | 0.008588691 |
| Cadps | 5889.517384 | 0.36637316 | 0.00022102 |
| LOC690806 | 3177.618144 | 0.366336783 | 4.22E-06 |
| Acap3 | 2076.190362 | 0.366282577 | 0.006706917 |
| St8sia1 | 442.7807931 | 0.366222221 | 0.00439152 |
| Thrb | 3087.0989 | 0.366093798 | 0.004279776 |
| Cnnm1 | 2388.56101 | 0.365518945 | 0.000168225 |
| Smyd2 | 1356.046773 | 0.365495183 | 6.91E-05 |
| Gpr155 | 1279.601672 | 0.365403951 | 0.00032079 |
| Lrrtm2 | 2797.8136 | 0.365150892 | 4.33E-05 |
| Lhfpl5 | 198.8339843 | 0.365133359 | 0.009343788 |
| Grm8 | 376.746908 | 0.364979367 | 0.007946768 |
| Kifap3 | 7988.485986 | 0.363899086 | 2.19E-06 |
| Tmem132d | 869.7664381 | 0.363731917 | 0.032404777 |
| Synj1 | 14040.3733 | 0.362997478 | 2.50E-05 |
| Pak1 | 5287.878576 | 0.362759491 | 0.000179941 |
| Rell2 | 1624.443988 | 0.362658445 | 0.000463846 |
| Sh3bp5 | 830.3008992 | 0.362564681 | 3.81E-05 |
| Cbx6 | 6305.286351 | 0.361973923 | 2.35E-06 |
| Itga11 | 166.5573587 | 0.361820668 | 0.034731898 |
| B4galnt4 | 1750.255479 | 0.361733553 | 0.000626031 |

| | | | |
|---|---|---|---|
| Man1c1 | 1303.912027 | 0.36146702 | 5.05E-06 |
| Prickle1 | 2553.166981 | 0.361205526 | 0.023523193 |
| Mafb | 571.071754 | 0.361159617 | 0.005522485 |
| Allc | 16.00089458 | 0.360853822 | 0.038136158 |
| Tmem132b | 2484.703255 | 0.360773561 | 0.003436524 |
| Rbm41 | 66.55744828 | 0.360721722 | 0.032709024 |
| Celf2 | 5070.260392 | 0.360102392 | 0.000125222 |
| Tcea2 | 1032.223702 | 0.360092378 | 9.98E-06 |
| Tppp | 3123.338379 | 0.360047501 | 0.000187819 |
| Sptb | 5510.750692 | 0.359883025 | 0.00098682 |
| Serpine1 | 58.74063 | 0.359788883 | 0.036227797 |
| Pacsin1 | 6943.889528 | 0.359769977 | 8.12E-06 |
| Ier3 | 57.71757776 | 0.359694201 | 0.025390626 |
| Fabp3 | 516.0079148 | 0.359667756 | 0.00404385 |
| Sez6l2 | 13349.79941 | 0.35960452 | 1.33E-06 |
| Zfp575 | 404.6630034 | 0.359283791 | 0.00171295 |
| Pclo | 19246.00289 | 0.359086314 | 0.035784906 |
| Pde4d | 1309.0425 | 0.358845022 | 7.05E-07 |
| Kcnj11 | 732.9838754 | 0.358822953 | 0.00068522 |
| Tmem200a | 501.9574088 | 0.35878088 | 0.008747455 |
| Runx2 | 56.50343506 | 0.358720741 | 0.041957792 |
| Tmem35 | 2010.825068 | 0.358363457 | 0.001316812 |
| Wnt16 | 28.22066137 | 0.357663515 | 0.0425212 |
| Slc2a13 | 7433.145343 | 0.357577332 | 5.63E-06 |
| Arid4a | 1959.468231 | 0.357444798 | 2.12E-05 |
| Map3k12 | 2830.50464 | 0.357378903 | 2.70E-06 |
| Raph1 | 3202.624635 | 0.357149768 | 9.62E-05 |
| Thbd | 27.78457262 | 0.357007278 | 0.043429145 |
| Ap1s3 | 42.95807816 | 0.356959965 | 0.041280268 |
| Ajap1 | 913.1561449 | 0.356952629 | 4.22E-05 |
| Tigar | 647.3993915 | 0.356791905 | 0.000222439 |
| Dbn1 | 3728.272953 | 0.356760999 | 0.002477481 |
| Mcts1 | 385.4628995 | 0.356676913 | 0.001626107 |
| Chst9 | 14.57143311 | 0.356473682 | 0.041346311 |
| Kctd8 | 480.8480409 | 0.356430682 | 0.010153739 |
| Large | 4434.20101 | 0.356377383 | 0.000513831 |

| | | | |
|---|---|---|---|
| Napg | 4445.664442 | 0.356019161 | 0.000100468 |
| Trib1 | 325.8793794 | 0.355941297 | 0.029347973 |
| Afap1l1 | 84.28967872 | 0.355923909 | 0.032069991 |
| Sbk1 | 2338.985186 | 0.355914887 | 3.07E-05 |
| Ola1 | 3111.637612 | 0.355394367 | 9.60E-06 |
| Mapkbp1 | 1063.213085 | 0.355119483 | 0.000185068 |
| Lrrn3 | 1696.364826 | 0.354774151 | 0.000840224 |
| Ckmt1b | 5730.10842 | 0.354639866 | 3.39E-05 |
| Nxph1 | 2600.492781 | 0.354535818 | 6.32E-05 |
| Msl3 | 464.9803923 | 0.35452181 | 0.000831376 |
| Mtmr12 | 292.4136236 | 0.354482922 | 0.00387717 |
| RGD1310852 | 249.6639713 | 0.354272377 | 0.004112093 |
| Madd | 6932.916151 | 0.353213187 | 1.78E-05 |
| Scn1b | 2962.839926 | 0.352853385 | 0.011837977 |
| Adamtsl3 | 120.3057574 | 0.352632426 | 0.039468079 |
| Pdzd4 | 2507.702254 | 0.352581503 | 4.21E-06 |
| Arel1 | 5268.460552 | 0.352489274 | 5.88E-07 |
| Faah | 867.078562 | 0.352415933 | 0.004499548 |
| Plcb1 | 9387.740254 | 0.352014472 | 0.000121478 |
| Sms | 1801.018194 | 0.351658215 | 0.000302767 |
| Gpr27 | 256.2777746 | 0.35160716 | 0.024137529 |
| Mgat4a | 1180.592533 | 0.351597364 | 5.45E-05 |
| Glcci1 | 752.0529768 | 0.351515941 | 0.000288334 |
| Zfp385b | 1249.040486 | 0.351267977 | 0.000594793 |
| Faxc | 669.1223716 | 0.35121544 | 0.000147144 |
| Tacr2 | 12.31920716 | 0.351075984 | 0.039695508 |
| Cadm3 | 5358.497176 | 0.35069018 | 2.98E-05 |
| Bok | 544.3447324 | 0.350664962 | 0.005431207 |
| Dusp6 | 784.1536723 | 0.35027194 | 0.019874837 |
| Exog | 373.5057852 | 0.350170361 | 0.00140131 |
| Mycbp2 | 15071.00945 | 0.350112012 | 0.00057811 |
| Sh3pxd2a | 866.9499794 | 0.350070035 | 0.002007763 |
| Zwint | 86293.88948 | 0.34999666 | 4.67E-08 |
| Prickle2 | 7979.4164 | 0.349235657 | 0.000112453 |
| Kcnk1 | 2097.863189 | 0.349060159 | 8.36E-05 |
| Actl6b | 1296.109536 | 0.34887995 | 0.001781317 |

| | | | |
|---|---|---|---|
| Mapre3 | 5403.271956 | 0.348872146 | 2.47E-06 |
| Ssx2ip | 1822.481534 | 0.348827918 | 1.28E-07 |
| Npdc1 | 3534.93419 | 0.348758668 | 0.000119076 |
| Nat8l | 8026.738881 | 0.348710657 | 0.000326518 |
| Rasl10b | 1045.501619 | 0.348502171 | 0.000700095 |
| Coro1a | 1110.300659 | 0.348464858 | 0.01761778 |
| Sp9 | 514.027378 | 0.348273417 | 0.000375704 |
| Slc9a3r2 | 1069.749734 | 0.347952987 | 8.18E-05 |
| Uchl1 | 25708.84002 | 0.347870351 | 0.000183716 |
| Neurl2 | 99.69396643 | 0.347762761 | 0.018882978 |
| Sstr3 | 265.9436927 | 0.347752325 | 0.036569268 |
| Gucy1a3 | 1899.635282 | 0.34773086 | 0.000648389 |
| Pnmal1 | 2369.014579 | 0.347653666 | 0.000249278 |
| Gripap1 | 5463.788491 | 0.34731638 | 2.77E-06 |
| Chrm3 | 1419.587056 | 0.347252866 | 0.022665227 |
| Zfp280d | 1535.509615 | 0.347145943 | 2.40E-06 |
| R3hdm1 | 5659.451811 | 0.347008574 | 0.000708828 |
| Nanp | 282.5399166 | 0.346555927 | 2.10E-05 |
| LRRTM1 | 1579.888408 | 0.34652671 | 1.23E-05 |
| Suv39h2 | 188.1841654 | 0.345999509 | 0.017022057 |
| Bag4 | 1420.970931 | 0.345353032 | 1.94E-06 |
| Zfp2 | 429.7930085 | 0.345267208 | 0.002121224 |
| Akap11 | 12538.20237 | 0.345194705 | 0.000271886 |
| Zmiz2 | 8880.8541 | 0.345084748 | 1.38E-05 |
| Pcdha10 | 212.6844431 | 0.345070046 | 0.029377912 |
| Adra2b | 34.10982899 | 0.34499636 | 0.048928606 |
| Lgr5 | 328.8983641 | 0.344946824 | 0.01759164 |
| Clec2l | 472.9751519 | 0.34479976 | 0.021219509 |
| Klf10 | 483.5434477 | 0.344771704 | 0.049542627 |
| Lppr5 | 1596.210911 | 0.344644902 | 0.002717811 |
| Cplx1 | 18649.11711 | 0.344339183 | 0.007018818 |
| Oxtr | 50.39344506 | 0.344009847 | 0.049870953 |
| Ksr1 | 1190.606102 | 0.343969523 | 0.003457968 |
| Ipcef1 | 661.2120174 | 0.343897304 | 0.048423818 |
| Tyro3 | 6394.135624 | 0.343585111 | 0.006597375 |
| Tsc22d3 | 2645.500888 | 0.343473934 | 0.03390701 |

| | | | |
|---|---|---|---|
| Atp6ap2 | 3401.020402 | 0.343262708 | 2.15E-06 |
| Tspan5 | 3227.739688 | 0.343169374 | 0.003518594 |
| Agk | 881.4728719 | 0.343090515 | 4.34E-05 |
| Syngr3 | 2091.785605 | 0.343032755 | 1.25E-06 |
| Azin2 | 374.0916155 | 0.34290823 | 0.000536787 |
| Abracl | 145.5194364 | 0.342764134 | 0.037442833 |
| Camk1 | 1890.877215 | 0.342505255 | 0.000676016 |
| Mbnl1 | 742.6808564 | 0.34238715 | 4.45E-05 |
| Fbxl19 | 869.9048198 | 0.341638686 | 0.000444941 |
| Ttc19 | 1816.32602 | 0.341516256 | 0.000714909 |
| Jakmip2 | 1603.742267 | 0.341145426 | 0.000106124 |
| Myadml2 | 91.12817743 | 0.340903828 | 0.034353393 |
| Cend1 | 10584.34323 | 0.340824781 | 0.000406438 |
| Slc25a14 | 1003.80316 | 0.340735832 | 0.00320407 |
| Mink1 | 14447.24766 | 0.340272387 | 8.25E-06 |
| Apba1 | 7697.172224 | 0.340178556 | 0.000238911 |
| Incenp | 504.7612335 | 0.340100999 | 0.002080046 |
| Myh10 | 17370.46161 | 0.339852762 | 0.001668107 |
| Slit2 | 744.1304709 | 0.339775985 | 0.006213156 |
| Vwc2l | 153.8460436 | 0.339580084 | 0.018938358 |
| Rita1 | 584.2989572 | 0.339403207 | 0.001383471 |
| Igfbp3 | 140.2061881 | 0.339329529 | 0.035050145 |
| Moap1 | 1116.687565 | 0.33894896 | 0.004401296 |
| Abca5 | 1007.482459 | 0.338917834 | 0.001089151 |
| Ppp2r3b | 151.4462984 | 0.338571811 | 0.019108047 |
| Wscd2 | 565.0684435 | 0.338529153 | 0.000996879 |
| Gpr162 | 3558.822378 | 0.3378582 | 0.000143119 |
| Eif5a2 | 2409.020706 | 0.337572319 | 7.89E-05 |
| Grik5 | 6310.0616 | 0.337345245 | 1.45E-06 |
| Ywhaz | 33758.57184 | 0.337103799 | 2.67E-06 |
| Slc7a4 | 922.6372483 | 0.336988138 | 0.002615491 |
| Jakmip1 | 4793.434136 | 0.336807149 | 9.09E-09 |
| Lppr2 | 4866.791909 | 0.336657361 | 0.000357927 |
| Rangap1 | 7407.886831 | 0.336557376 | 8.94E-06 |
| Drd5 | 102.2346574 | 0.336322587 | 0.02486987 |
| Tmem179 | 2639.431587 | 0.336272107 | 0.000224796 |

| | | | |
|---|---|---|---|
| Cep85 | 396.4473643 | 0.335752409 | 0.000732947 |
| Inpp4a | 7871.474571 | 0.335514221 | 1.43E-05 |
| Rap1gap | 9157.730239 | 0.335325633 | 7.22E-06 |
| Dus1l | 424.7072341 | 0.335011327 | 0.000631055 |
| Fscn1 | 4511.497054 | 0.334876494 | 0.007334104 |
| Ptprn2 | 13867.52058 | 0.334293162 | 1.92E-05 |
| Senp2 | 1432.693619 | 0.334245393 | 1.34E-07 |
| Pcdhac1 | 131.5016598 | 0.334049197 | 0.040106217 |
| Mapk9 | 6227.552192 | 0.334021347 | 7.32E-06 |
| Armcx2 | 3329.562399 | 0.333835757 | 2.70E-05 |
| Plxna2 | 8165.894746 | 0.333824547 | 9.75E-06 |
| Adgrl1 | 19697.94421 | 0.333301091 | 1.33E-06 |
| Fam160b2 | 1215.590897 | 0.333244132 | 4.82E-05 |
| Csrnp1 | 193.1364105 | 0.332930709 | 0.031052073 |
| Ttc3 | 24271.14866 | 0.33281593 | 1.61E-05 |
| Sytl5 | 310.9248264 | 0.332767993 | 0.011445678 |
| Dlx2 | 317.7644714 | 0.332552373 | 0.005648352 |
| LOC102548847 | 192.8084333 | 0.332319287 | 0.026152029 |
| Pianp | 4829.912086 | 0.332254705 | 3.69E-05 |
| Spata2 | 1344.968641 | 0.332122476 | 0.000362504 |
| Coro7 | 1671.153156 | 0.332065628 | 0.00021015 |
| Ablim2 | 1613.396806 | 0.331827521 | 6.93E-06 |
| Tomm20 | 3702.878053 | 0.331708359 | 3.67E-05 |
| Wdr47 | 4739.840219 | 0.331126776 | 0.000208738 |
| Coro6 | 229.5188054 | 0.331046331 | 0.03125143 |
| Map2k4 | 1206.759668 | 0.330772016 | 6.64E-05 |
| Mturn | 861.488323 | 0.330622322 | 0.000348118 |
| Tuba4a | 9162.441909 | 0.330569609 | 0.000402658 |
| Tmem60 | 311.292542 | 0.330305199 | 0.011069004 |
| Tp53i11 | 2223.247368 | 0.330218933 | 0.041580553 |
| Celsr2 | 19145.5639 | 0.330210217 | 0.002005475 |
| Gtpbp6 | 474.5915357 | 0.329733303 | 0.000526899 |
| Rybp | 1623.279209 | 0.329664702 | 9.42E-05 |
| Ccdc32 | 1499.003614 | 0.329594802 | 0.000100468 |
| Dtnb | 1228.163849 | 0.329442785 | 0.000101182 |
| Map6 | 5101.64587 | 0.32943214 | 1.34E-05 |

| | | | |
|---|---|---|---|
| Slitrk1 | 3345.705167 | 0.329415005 | 0.002755177 |
| Pcdha2 | 321.41944 | 0.329409807 | 0.029736819 |
| Mknk2 | 1242.882081 | 0.329143126 | 5.63E-05 |
| Tuba8 | 553.7512434 | 0.328948368 | 0.008831004 |
| Cdc42se2 | 2214.543933 | 0.328920084 | 6.66E-05 |
| Ltk | 86.49395041 | 0.328663498 | 0.035717586 |
| Gdap1l1 | 1194.29766 | 0.328584382 | 0.001120009 |
| Spryd3 | 4249.837652 | 0.328529346 | 2.43E-05 |
| Tmem59l | 6338.720311 | 0.328492932 | 2.87E-05 |
| Fbxo9 | 2478.836016 | 0.328311487 | 3.25E-05 |
| Pard6a | 294.9048538 | 0.327295478 | 0.001206424 |
| Fcho1 | 939.885644 | 0.327116771 | 0.001325039 |
| Ccdc64 | 680.3025258 | 0.327054651 | 0.001170519 |
| Slc1a6 | 104.1812512 | 0.326930493 | 0.049707331 |
| Pja2 | 29734.47418 | 0.32690959 | 6.61E-09 |
| Slc25a25 | 1655.412143 | 0.32662038 | 0.001247377 |
| Lrrc24 | 909.404014 | 0.326345322 | 0.000567055 |
| Neurl4 | 3511.633049 | 0.326316147 | 3.94E-05 |
| Sars2 | 361.3524505 | 0.326265078 | 0.006382305 |
| Bex1 | 1147.813961 | 0.325498673 | 1.30E-05 |
| Stmn2 | 8313.756419 | 0.325446925 | 0.000115668 |
| Slc7a8 | 2643.685138 | 0.325365942 | 0.012540404 |
| Chrm2 | 762.8406555 | 0.325339315 | 0.001165143 |
| Siah2 | 336.3728422 | 0.325334721 | 0.010868574 |
| Gtdc1 | 379.9495491 | 0.325121576 | 0.016388142 |
| Acsl5 | 794.0606696 | 0.324308959 | 0.006710052 |
| Tmcc2 | 3307.478353 | 0.324103864 | 9.23E-05 |
| Rasgrf1 | 23921.54751 | 0.324050499 | 0.000943073 |
| Arhgef2 | 2822.642771 | 0.323755567 | 0.008975671 |
| Cdyl2 | 429.6163243 | 0.32372401 | 0.000830984 |
| Slc2a6 | 514.2791242 | 0.323624558 | 0.007960743 |
| Car12 | 426.5996534 | 0.323601925 | 0.034285819 |
| Prkaa2 | 543.1524061 | 0.323530279 | 0.011945474 |
| Kcnma1 | 4031.461182 | 0.323351491 | 0.001577074 |
| Kit | 3144.941767 | 0.323267081 | 0.003774345 |
| Ppp2r5b | 3790.42939 | 0.323143512 | 3.92E-05 |

| | | | |
|---|---|---|---|
| Mtf2 | 525.8255671 | 0.323056272 | 0.002507095 |
| Cdk5 | 1372.386953 | 0.322944768 | 0.001414533 |
| Calm2 | 21342.88944 | 0.322878237 | 2.27E-05 |
| Cops7b | 490.3769173 | 0.322735187 | 0.000652834 |
| Oprd1 | 192.4684229 | 0.322565879 | 0.047560323 |
| Slc45a1 | 2074.501044 | 0.322559227 | 0.000508867 |
| Susd4 | 1725.270321 | 0.322130017 | 0.00027262 |
| Atp1a3 | 102261.1516 | 0.321966518 | 0.001477497 |
| Ppm1e | 4734.481038 | 0.321791771 | 0.009693554 |
| Rgs8 | 2864.826111 | 0.321713015 | 0.003885805 |
| Ocm2 | 14.72636427 | 0.321490216 | 0.04951807 |
| Wdr13 | 1565.616195 | 0.321256817 | 0.002359125 |
| Ier5l | 614.960269 | 0.321202881 | 0.009241851 |
| Hrk | 983.2835845 | 0.321168419 | 0.023352229 |
| Ing5 | 256.9023799 | 0.32079524 | 0.003670575 |
| Fsd1 | 318.8432368 | 0.32075704 | 0.005715007 |
| Zfand5 | 2925.964189 | 0.320754325 | 3.44E-06 |
| Serp2 | 347.642761 | 0.320615703 | 0.019016441 |
| Gdap1 | 2457.726993 | 0.320451336 | 0.002505606 |
| RGD1305587 | 510.1874085 | 0.320420845 | 0.005165547 |
| Mex3b | 614.324713 | 0.320333126 | 0.021713001 |
| Cebpa | 132.5581929 | 0.319897168 | 0.028570555 |
| Itpa | 754.3801363 | 0.319710308 | 0.004243956 |
| Ubald1 | 2706.453819 | 0.319668965 | 2.19E-05 |
| Klc1 | 21914.58607 | 0.319463398 | 0.000124426 |
| Grip1 | 1499.894035 | 0.319420902 | 2.59E-05 |
| Map3k10 | 3225.148357 | 0.318962324 | 0.001414533 |
| Cited2 | 2147.438409 | 0.318950286 | 0.000424753 |
| Strn4 | 4567.040232 | 0.318836576 | 0.001747592 |
| Scn3a | 3256.676625 | 0.318835868 | 0.00031594 |
| Srcin1 | 10704.73443 | 0.318822475 | 0.023603946 |
| Tec | 110.6742099 | 0.318766061 | 0.039214393 |
| Man2a1 | 1130.621011 | 0.317813008 | 0.001394243 |
| Ppapdc3 | 452.1718967 | 0.317556126 | 0.002091477 |
| Plcxd2 | 2226.764258 | 0.317418333 | 0.016730629 |
| Ptchd1 | 308.239854 | 0.317347529 | 0.014050678 |

| | | | |
|---|---|---|---|
| Khdrbs2 | 272.5762193 | 0.31712999 | 0.032852101 |
| Acvr1b | 1763.596257 | 0.316834273 | 3.70E-05 |
| Map2k6 | 444.6747378 | 0.316751098 | 0.007407006 |
| Efnb2 | 512.1258622 | 0.316736653 | 0.011877535 |
| Tenm2 | 4299.394515 | 0.316452586 | 0.001826382 |
| Rundc3b | 1264.465535 | 0.316017403 | 0.001272809 |
| LOC102549726 | 244.3072044 | 0.315956198 | 0.030979794 |
| Numbl | 1372.155754 | 0.315521725 | 0.004693495 |
| Cmas | 2470.583712 | 0.315420229 | 4.47E-05 |
| Csmd1 | 3681.417415 | 0.314588505 | 0.007407006 |
| Gucy1a2 | 1255.648739 | 0.314498025 | 0.000250651 |
| Mmp24 | 1217.597792 | 0.31405235 | 1.97E-05 |
| Dlx1 | 804.7499259 | 0.313914235 | 0.005709065 |
| Stk24 | 1037.151783 | 0.313902864 | 6.25E-07 |
| Ap1s1 | 2116.337513 | 0.313818203 | 0.000103894 |
| Nyap1 | 3225.221896 | 0.313772462 | 4.75E-10 |
| Phf20 | 2171.250056 | 0.313738283 | 1.61E-07 |
| RGD1305014 | 234.2034022 | 0.313735857 | 0.013128675 |
| Atxn1 | 3575.680065 | 0.313656422 | 0.000750944 |
| Git1 | 9480.578716 | 0.313471893 | 0.000594793 |
| Syt17 | 1497.393942 | 0.313437704 | 0.008659298 |
| Pip4k2b | 2755.115582 | 0.313383023 | 7.57E-05 |
| Lysmd1 | 471.4989417 | 0.313282975 | 0.002529975 |
| Cnrip1 | 2027.412935 | 0.31298958 | 0.011827153 |
| Ndel1 | 2467.328782 | 0.312635961 | 1.36E-08 |
| Nap1l2 | 2385.805198 | 0.312582757 | 0.000428128 |
| Syngr1 | 3395.281663 | 0.312415265 | 0.002524964 |
| Crabp1 | 1176.952628 | 0.312209679 | 0.022338058 |
| Plk3 | 253.5804244 | 0.312118492 | 0.039174151 |
| Ttbk1 | 2525.858012 | 0.311872913 | 0.004198735 |
| Gmnn | 81.32129968 | 0.311806148 | 0.047932298 |
| Fbxo33 | 1183.44157 | 0.311724455 | 0.000174821 |
| Tmem181 | 1008.334204 | 0.31165964 | 0.000245882 |
| Clstn1 | 44934.83922 | 0.311477447 | 0.000131116 |
| Sptbn4 | 5580.857413 | 0.311377761 | 0.002036487 |
| Dnajb4 | 1074.45324 | 0.31116327 | 2.14E-05 |

| | | | |
|---|---|---|---|
| RGD1563441 | 174.9640067 | 0.311044554 | 0.019927587 |
| Vwc2 | 444.0766582 | 0.311020295 | 0.017418257 |
| Galnt16 | 4945.630888 | 0.310989584 | 0.00258672 |
| Ankrd12 | 4406.818336 | 0.310677459 | 0.000296832 |
| RGD1307704 | 392.8685689 | 0.310403688 | 0.002448831 |
| Rgs17 | 2423.035141 | 0.310374391 | 8.91E-05 |
| Zfp180 | 1449.119035 | 0.310298138 | 0.000165067 |
| Ezh1 | 2374.402769 | 0.310245201 | 1.12E-06 |
| Basp1 | 9661.058365 | 0.309865161 | 0.028061735 |
| Elavl3 | 2703.580675 | 0.30961501 | 2.37E-06 |
| Ubxn2b | 651.5927958 | 0.309594703 | 0.000658404 |
| Kif5a | 21367.7122 | 0.309559971 | 0.000174114 |
| Map1a | 72196.52273 | 0.309109818 | 0.008416584 |
| Rtn4rl1 | 1834.072452 | 0.309036417 | 0.001524652 |
| Iqsec2 | 4747.817622 | 0.308899096 | 0.003787107 |
| Gng3 | 3940.343598 | 0.308720719 | 0.001066549 |
| Srr | 839.8670542 | 0.308719335 | 0.003064225 |
| Agap3 | 4573.992244 | 0.308701812 | 3.97E-05 |
| Ddi2 | 2000.288157 | 0.308571616 | 1.28E-06 |
| Gnb5 | 1051.514654 | 0.308560742 | 0.00034315 |
| Apitd1 | 214.1198742 | 0.308522521 | 0.010686841 |
| Hras | 1513.008427 | 0.308460707 | 0.004093876 |
| Aplp1 | 23095.42386 | 0.308448752 | 0.001510612 |
| Atmin | 3119.885191 | 0.308410232 | 4.68E-05 |
| Ablim3 | 683.141554 | 0.308272374 | 0.000230095 |
| Mbtps2 | 257.6199202 | 0.307988738 | 0.003449774 |
| Panx2 | 2459.975022 | 0.30797031 | 0.000526257 |
| Grik3 | 3702.813964 | 0.307706541 | 0.025740813 |
| Dync1i1 | 4012.965791 | 0.307582173 | 0.000586761 |
| Rexo1 | 2567.229426 | 0.307271731 | 9.18E-06 |
| Rock2 | 10288.88556 | 0.307153714 | 0.003537371 |
| Syp | 11652.64826 | 0.307052453 | 0.000766621 |
| Pten | 2203.908392 | 0.307042684 | 0.000468684 |
| Tyr | 3.558038473 | 0.306712653 | 0.038780433 |
| Ap2m1 | 13989.5697 | 0.306705361 | 0.000134238 |
| Rnf126 | 1170.346973 | 0.306522889 | 0.004579824 |

| | | | |
|---|---|---|---|
| Pik3cb | 970.3758949 | 0.306432463 | 0.000245882 |
| Galnt13 | 672.5262929 | 0.306418005 | 0.01038453 |
| Snap47 | 887.2345111 | 0.306333293 | 0.004433611 |
| Tusc2 | 1476.350291 | 0.306330252 | 0.007091055 |
| Gsk3a | 4888.596692 | 0.30612173 | 1.82E-06 |
| Ube2q1 | 2919.141744 | 0.305941741 | 3.20E-07 |
| Reep1 | 4035.3643 | 0.305888271 | 2.37E-05 |
| Crmp1 | 5346.635714 | 0.305827353 | 0.000249229 |
| Cacna1h | 1270.033219 | 0.30575857 | 0.021481122 |
| Pcyox1l | 969.770116 | 0.305384851 | 4.92E-05 |
| Fbxo31 | 2161.126636 | 0.305208863 | 3.39E-05 |
| Tsc22d2 | 1224.232386 | 0.304984639 | 0.000280326 |
| Tbk1 | 993.9186758 | 0.30493363 | 0.000180995 |
| Rab3b | 2365.217453 | 0.304376328 | 0.002523867 |
| Gtf2h1 | 905.6381734 | 0.304069399 | 0.000533884 |
| Slmap | 3825.825202 | 0.303939586 | 1.30E-06 |
| Pitpnm2 | 11881.02908 | 0.303625138 | 0.000181367 |
| Gpr158 | 8102.670548 | 0.303486745 | 0.001196554 |
| Coq2 | 1233.240361 | 0.303228289 | 0.000338454 |
| Kcnc2 | 2810.657474 | 0.303226534 | 0.013085346 |
| Slc37a1 | 179.5919657 | 0.303158247 | 0.037966418 |
| Lrrc10b | 334.7533522 | 0.303075414 | 0.023707852 |
| Smad3 | 1574.125646 | 0.30300541 | 0.001637594 |
| Avpi1 | 739.3288829 | 0.30287652 | 0.008987589 |
| Stk25 | 4900.893243 | 0.302702523 | 1.00E-04 |
| Ap1ar | 1052.098684 | 0.302610891 | 7.71E-05 |
| Frmd6 | 601.9535909 | 0.302568259 | 0.011837977 |
| Tmeff1 | 2219.564336 | 0.302504032 | 0.001875659 |
| Mnt | 2316.434191 | 0.301973564 | 0.000146508 |
| Tmem68 | 645.05191 | 0.301897167 | 0.001673847 |
| Svop | 2395.387146 | 0.301844404 | 5.26E-05 |
| Ttc7b | 3814.88569 | 0.301798451 | 6.76E-05 |
| Eogt | 279.8474112 | 0.301662844 | 0.011821446 |
| Gopc | 2538.933169 | 0.301112774 | 4.66E-05 |
| Nfx1 | 2370.53298 | 0.30100235 | 4.27E-09 |
| Hprt1 | 1480.749857 | 0.300996072 | 0.002671234 |

| | | | |
|---|---|---|---|
| Zdhhc8 | 1845.83112 | 0.300919419 | 0.00016609 |
| Ccser1 | 205.6272485 | 0.300906629 | 0.010728816 |
| Got2 | 8109.096299 | 0.300704014 | 3.81E-05 |
| Unc13c | 740.7607189 | 0.300541543 | 0.036586093 |
| Map3k9 | 2294.074235 | 0.300207539 | 0.005252079 |
| Ica1 | 940.1693483 | 0.29997941 | 0.000455805 |
| Pkia | 2411.30785 | 0.299695207 | 0.000728717 |
| Atp6v1e1 | 7401.505806 | 0.299599505 | 1.71E-05 |
| Fgf12 | 818.0944012 | 0.299435698 | 0.001467342 |
| Ppme1 | 5417.414743 | 0.299345673 | 0.000249859 |
| Arf5 | 2694.380843 | 0.299222153 | 0.001170519 |
| Gpr83 | 1074.029308 | 0.299087508 | 0.002402127 |
| Elk1 | 907.1609105 | 0.299082679 | 0.000741777 |
| RGD1307443 | 1871.550575 | 0.298580956 | 0.000738182 |
| Zc2hc1a | 856.9199975 | 0.298557162 | 0.000239776 |
| Thoc5 | 1074.049653 | 0.297171621 | 0.000701177 |
| Pik3r2 | 4937.006163 | 0.297041385 | 0.001893464 |
| Nck2 | 1411.149926 | 0.296939759 | 0.018503907 |
| Rap1gds1 | 11412.62786 | 0.296919356 | 0.019893903 |
| Prdm2 | 3937.038096 | 0.296837644 | 0.00010395 |
| Ppp1r9b | 26488.64256 | 0.296797459 | 0.000158821 |
| Fat3 | 5208.775287 | 0.296706558 | 0.014301518 |
| Ina | 8143.744427 | 0.296648226 | 0.005240097 |
| Ap2a2 | 10851.52355 | 0.296611253 | 0.003490773 |
| Amph | 4682.46621 | 0.296451817 | 0.000433686 |
| Lrrc8c | 854.6600454 | 0.296367236 | 0.002362066 |
| Trim66 | 892.4176885 | 0.295786199 | 0.001020361 |
| Fam84a | 3423.763023 | 0.295652614 | 0.000233324 |
| Sphkap | 3223.290837 | 0.294878891 | 0.010029862 |
| Syt13 | 2235.018126 | 0.294741488 | 0.000448977 |
| Ypel3 | 1739.80854 | 0.294059054 | 0.001579099 |
| Mapk8ip3 | 12712.5635 | 0.294023778 | 0.00022102 |
| Tmem151a | 3636.087697 | 0.293730558 | 0.011612852 |
| Socs5 | 2624.05125 | 0.293480349 | 8.37E-05 |
| Ccdc92 | 2212.53027 | 0.293380251 | 0.000155171 |
| Gspt2 | 1052.49499 | 0.293079353 | 0.003325565 |

| | | | |
|---|---|---|---|
| Cdkl1 | 224.1550038 | 0.292948067 | 0.026712689 |
| Hecw1 | 1475.458035 | 0.292665278 | 0.008245717 |
| Rasal2 | 1340.058649 | 0.292573197 | 0.008070844 |
| Arfip2 | 1368.810178 | 0.29255338 | 0.000757456 |
| Ywhag | 16872.31426 | 0.292506413 | 5.72E-05 |
| Rnf14 | 14500.68952 | 0.292473289 | 5.32E-05 |
| Elavl2 | 3378.360715 | 0.292465732 | 0.02854608 |
| Klhdc3 | 4347.39979 | 0.29230869 | 7.79E-05 |
| B4galt3 | 880.0082361 | 0.291859872 | 0.000533844 |
| Dcp2 | 663.6599473 | 0.291514988 | 0.000880367 |
| Rtn1 | 30865.25437 | 0.291504374 | 6.92E-05 |
| Pbx2 | 1798.611706 | 0.291374882 | 1.96E-05 |
| Sv2a | 13597.76567 | 0.291071826 | 0.000833391 |
| Wsb2 | 2924.607994 | 0.291034793 | 0.000739935 |
| Atg13 | 2715.157921 | 0.291008025 | 7.88E-07 |
| Dpp10 | 1936.874118 | 0.290788524 | 0.003836416 |
| Mbnl2 | 6966.370302 | 0.289903105 | 4.47E-06 |
| Tbc1d24 | 1960.836661 | 0.289545982 | 5.02E-05 |
| Cdc34 | 1009.918259 | 0.289366548 | 0.005606791 |
| Stmn1 | 10277.79252 | 0.289234309 | 0.010101498 |
| Spin1 | 1932.270096 | 0.289224469 | 2.16E-05 |
| Lysmd2 | 941.6948125 | 0.289037196 | 0.0003514 |
| Exoc6 | 1005.025688 | 0.289001493 | 0.00874504 |
| Bmyc | 726.1679568 | 0.288951162 | 0.000382551 |
| Bcat1 | 44185.35312 | 0.288936867 | 0.000142873 |
| Stau2 | 2692.683949 | 0.288745707 | 1.78E-06 |
| Npas2 | 2845.817443 | 0.288582564 | 0.011729399 |
| Cryl1 | 278.634097 | 0.288410942 | 0.020205078 |
| Atp2b3 | 4785.894674 | 0.288304543 | 0.000927255 |
| Htr3a | 175.7799184 | 0.288300808 | 0.029986379 |
| Uhmk1 | 5790.648666 | 0.288285246 | 0.000209556 |
| Gadd45g | 292.2525977 | 0.287867881 | 0.042364428 |
| Phtf2 | 205.3919383 | 0.287846832 | 0.036675908 |
| Ss18l1 | 499.6453022 | 0.287486741 | 0.000145715 |
| Snx10 | 2287.705942 | 0.286900618 | 0.001788056 |
| Rapgef6 | 957.2703309 | 0.286802766 | 0.006755238 |

| | | | |
|---|---|---|---|
| RGD1305455 | 1730.151742 | 0.286747273 | 0.002480053 |
| Stmn3 | 7556.424868 | 0.286612221 | 0.001020166 |
| Faim2 | 8432.197522 | 0.286478462 | 0.000145887 |
| Dscaml1 | 2764.28276 | 0.286193258 | 0.002849644 |
| Zfp667 | 517.8125177 | 0.286131198 | 0.016352527 |
| Faap20 | 181.2572836 | 0.285596125 | 0.011384197 |
| Prkacb | 5158.31622 | 0.285583115 | 1.65E-05 |
| Kcnk3 | 373.1292124 | 0.285381206 | 0.021389539 |
| Ap5b1 | 436.3195596 | 0.285253795 | 0.008599641 |
| Dsel | 527.3356834 | 0.284818193 | 0.005079949 |
| Ogfrl1 | 1532.527334 | 0.284715801 | 0.004847825 |
| Ptprn | 14282.33288 | 0.284527377 | 0.002376254 |
| RGD1307461 | 168.7958404 | 0.284470806 | 0.041476297 |
| Gabarapl1 | 13051.92449 | 0.284141836 | 1.68E-05 |
| Lrfn1 | 1241.813564 | 0.283698326 | 0.003495986 |
| Kcna3 | 628.8630277 | 0.283573554 | 0.002417368 |
| Kcna2 | 2618.264916 | 0.283231781 | 0.014106315 |
| Fam117b | 3680.043574 | 0.282932561 | 1.64E-05 |
| Ppp1r1a | 1217.311971 | 0.28290628 | 0.001629454 |
| Mpped2 | 671.9781324 | 0.282873109 | 0.002361057 |
| Dnajc5 | 2074.122483 | 0.282443257 | 3.48E-06 |
| Trak2 | 5209.211879 | 0.282400836 | 0.000890536 |
| Gnai1 | 3864.744072 | 0.282384291 | 0.001221725 |
| Ppargc1b | 665.1209454 | 0.282311638 | 0.004721613 |
| Kctd1 | 3060.634327 | 0.281943645 | 7.72E-05 |
| Gabra3 | 1253.994334 | 0.281828052 | 0.02090229 |
| Pcbp3 | 1107.106404 | 0.281647285 | 0.000950829 |
| Ypel4 | 881.8557742 | 0.281508494 | 0.01306602 |
| Cnr1 | 7172.105319 | 0.280853213 | 0.016157506 |
| Dnajc14 | 1579.194777 | 0.280516725 | 0.000571693 |
| Mageb18 | 1.828116395 | 0.280399373 | 0.035050145 |
| Armcx3 | 3676.927785 | 0.280328923 | 2.36E-08 |
| Limk1 | 2689.07367 | 0.280042338 | 0.003617689 |
| Necap1 | 7202.601701 | 0.279927837 | 9.00E-06 |
| Mgat3 | 2891.65739 | 0.27982356 | 2.38E-05 |
| Spock1 | 4228.268722 | 0.27953654 | 0.006602324 |

| | | | |
|---|---|---|---|
| Clip1 | 2153.363013 | 0.279515428 | 0.011688362 |
| Insig2 | 186.1421238 | 0.278945561 | 0.022535123 |
| MAST1 | 4538.15901 | 0.278941983 | 0.000308897 |
| Cyp4x1 | 187.9269791 | 0.278668413 | 0.038704631 |
| Ap2a1 | 7761.238026 | 0.278450733 | 6.99E-05 |
| Kctd4 | 595.1903939 | 0.277501794 | 0.008975671 |
| Hist3h2ba | 484.0756959 | 0.277478897 | 0.035152458 |
| Nifk | 679.0915017 | 0.277331329 | 0.001642067 |
| Kcnj12 | 485.6418572 | 0.277157082 | 0.023559854 |
| Fam160a2 | 1679.230442 | 0.277106323 | 0.004863417 |
| Fbxo25 | 1370.618493 | 0.276992814 | 2.23E-05 |
| Ube2q2l | 263.0305508 | 0.276970266 | 0.03257977 |
| Ppm1h | 1966.501839 | 0.276953358 | 0.000105043 |
| Nup210 | 1063.522923 | 0.276938703 | 0.003814058 |
| Camsap3 | 1155.528423 | 0.276930242 | 0.00107362 |
| Mark1 | 2727.250576 | 0.276863212 | 9.23E-05 |
| App | 27295.81026 | 0.276424944 | 3.09E-05 |
| Hmgcll1 | 666.3448397 | 0.276221875 | 0.011730398 |
| Cux2 | 1250.772829 | 0.276162949 | 0.020547333 |
| Fstl4 | 602.0896375 | 0.275967464 | 0.032836097 |
| Epc2 | 1715.062086 | 0.275955104 | 0.000405328 |
| Clip3 | 15360.47154 | 0.27590048 | 4.87E-06 |
| Cyth2 | 2668.620325 | 0.275891373 | 0.010409711 |
| Spry4 | 687.8199824 | 0.275339551 | 0.006406779 |
| Ubxn6 | 828.2709812 | 0.275165285 | 0.002486981 |
| Ssbp4 | 1792.110599 | 0.275134809 | 0.001163975 |
| Klf9 | 5950.773467 | 0.275127105 | 0.002837284 |
| Adck3 | 751.3081045 | 0.275095047 | 0.003382328 |
| Rab11fip2 | 901.4795296 | 0.27495813 | 0.006477527 |
| Srxn1 | 1474.313699 | 0.274927824 | 0.009915989 |
| Map7d2 | 3345.922481 | 0.274717143 | 0.001508159 |
| LOC100125364 | 465.6014967 | 0.274668686 | 0.001642443 |
| Myo16 | 1926.741594 | 0.274660931 | 0.010420729 |
| Ccny | 1198.12956 | 0.274602633 | 8.26E-05 |
| R3hdm2 | 5495.696489 | 0.274584152 | 0.002755393 |
| Naa25 | 760.7425254 | 0.274456023 | 0.004091538 |

| | | | |
|---|---|---|---|
| Spred3 | 324.5644465 | 0.274100224 | 0.010669276 |
| Mfap3 | 328.7966484 | 0.273944171 | 0.018309193 |
| L1cam | 8751.112057 | 0.273919735 | 7.90E-06 |
| Pcmt1 | 4038.060284 | 0.273877011 | 4.54E-06 |
| Uhrf1bp1l | 6101.104573 | 0.273624067 | 0.001290024 |
| Papd7 | 1808.965675 | 0.273620176 | 0.000732947 |
| RGD1562079 | 924.7017324 | 0.273568015 | 0.000104103 |
| Ube2b | 2503.887268 | 0.273367938 | 0.001027841 |
| Kif3c | 8788.393708 | 0.273197156 | 4.94E-05 |
| Got1 | 8932.406173 | 0.272920509 | 0.004668556 |
| Pwwp2b | 402.9020544 | 0.272853704 | 0.022040795 |
| Pom121 | 5577.033296 | 0.272802381 | 0.000188526 |
| Dact3 | 4568.731357 | 0.272558326 | 0.002060539 |
| Vezt | 1601.535924 | 0.272374038 | 5.37E-05 |
| Zfp956 | 170.8317995 | 0.272247279 | 0.041572552 |
| Arpc1a | 3475.36321 | 0.272187849 | 1.00E-04 |
| Mlx | 1017.833566 | 0.272025633 | 0.000287459 |
| Cnot7 | 1367.140723 | 0.270820091 | 1.12E-05 |
| Adsl | 1189.180502 | 0.270737685 | 0.005566437 |
| Pfn2 | 10138.19489 | 0.270710692 | 6.15E-06 |
| Nufip1 | 557.3770853 | 0.27025089 | 0.000481565 |
| Gsr | 926.5789457 | 0.269821412 | 0.001648985 |
| Scamp5 | 3026.737026 | 0.269577625 | 0.000489973 |
| Bcl2l2 | 3986.488782 | 0.26954597 | 0.00026837 |
| Ctps1 | 2186.854745 | 0.269500158 | 6.47E-05 |
| Nmnat2 | 1908.568147 | 0.2694882 | 0.000903606 |
| Brpf3 | 994.9226514 | 0.269208451 | 0.005602846 |
| Unc5c | 1732.754201 | 0.269188513 | 0.001051709 |
| Astn1 | 14542.02739 | 0.269085764 | 0.014389301 |
| Gpr45 | 234.1740059 | 0.268640628 | 0.044192906 |
| H2afy2 | 683.2142629 | 0.268506716 | 0.004292859 |
| N4bp2l1 | 421.1630882 | 0.26831214 | 0.014942816 |
| Epha5 | 4260.297468 | 0.268186831 | 0.009221237 |
| Tubb3 | 14846.56723 | 0.26799676 | 0.000879365 |
| Fkrp | 1778.568848 | 0.267821609 | 0.001611576 |
| Palm | 5652.660793 | 0.267460497 | 0.00127953 |

| | | | |
|---|---|---|---|
| Rgs7 | 2017.957122 | 0.267229923 | 0.004512802 |
| Tpp2 | 2525.25242 | 0.266803699 | 9.93E-06 |
| Ldha | 8701.521858 | 0.266790014 | 0.02232975 |
| Trim2 | 14299.29666 | 0.266605741 | 1.75E-05 |
| Gsto1 | 1540.497429 | 0.266577219 | 0.001192039 |
| Sbno1 | 3752.533011 | 0.266415288 | 0.002952419 |
| Camsap1 | 6132.19665 | 0.266412163 | 3.36E-05 |
| Arhgap39 | 1700.471014 | 0.266099504 | 0.000251998 |
| Zfp483 | 3306.93467 | 0.265905845 | 0.003925806 |
| Tssc1 | 715.2970612 | 0.265874108 | 0.015266453 |
| Rims3 | 849.2891719 | 0.265667426 | 0.009970136 |
| Trim3 | 2497.556644 | 0.265203062 | 0.00125523 |
| Abca7 | 724.5700216 | 0.264577794 | 0.035691833 |
| Pptc7 | 3126.519895 | 0.26455209 | 0.002510697 |
| Kcnc1 | 5593.235637 | 0.264490464 | 0.03248044 |
| Arl6ip4 | 1451.057305 | 0.264379817 | 0.000866518 |
| Rtn2 | 1108.631883 | 0.264330989 | 0.005067566 |
| Cdc25b | 338.4934169 | 0.264329082 | 0.033357584 |
| Ndrg3 | 10232.72117 | 0.263611761 | 0.002584573 |
| Klhl8 | 2040.970537 | 0.263559476 | 0.000140532 |
| Trpm2 | 590.2716251 | 0.263431139 | 0.023083039 |
| Naa20 | 726.7088239 | 0.263256634 | 0.00262524 |
| Ptprm | 3113.66856 | 0.263092978 | 9.55E-05 |
| Dck | 430.5980397 | 0.263056108 | 0.019136951 |
| Dzip1l | 1030.386089 | 0.262859768 | 0.001057389 |
| Tusc3 | 4647.111377 | 0.262791364 | 0.003121979 |
| Pkm | 38139.73465 | 0.2626911 | 1.46E-05 |
| Mdh1 | 14226.42185 | 0.262672117 | 0.000204097 |
| Lrp11 | 6788.342539 | 0.262597186 | 0.000186178 |
| Tango2 | 1132.511198 | 0.262502333 | 0.005280474 |
| RGD1307235 | 2764.427559 | 0.262012912 | 0.001626107 |
| Ndfip2 | 2804.865487 | 0.261577685 | 0.000615194 |
| Jakmip3 | 764.5503559 | 0.261533951 | 0.003232016 |
| Dynll1 | 4283.049964 | 0.261414599 | 0.022599097 |
| Sez6 | 18051.98155 | 0.261127114 | 0.001506709 |
| Ap2s1 | 2204.222098 | 0.26090615 | 0.010149865 |

| | | | |
|---|---|---|---|
| Slc39a10 | 5074.311329 | 0.260868548 | 0.000522314 |
| Atp6v1a | 21296.10281 | 0.260756152 | 0.000161487 |
| Sort1 | 25055.45557 | 0.260600935 | 0.012883968 |
| Lrrc20 | 313.7078032 | 0.260579586 | 0.019545794 |
| Ly6h | 3222.067489 | 0.260473431 | 0.009480013 |
| Tbcc | 426.1489965 | 0.259671598 | 0.015565982 |
| Tomm34 | 2711.495097 | 0.259309385 | 0.000304541 |
| Vgf | 8502.952996 | 0.259307625 | 0.007061091 |
| Slc9a5 | 428.9459319 | 0.259179313 | 0.028052808 |
| Fn3k | 530.5031974 | 0.259150953 | 0.020378366 |
| Wasl | 5089.980956 | 0.259150853 | 7.58E-05 |
| Chchd6 | 1101.689466 | 0.259120587 | 0.011260585 |
| Ehd3 | 2841.40512 | 0.258256914 | 0.001524191 |
| Sf3a3 | 1184.837263 | 0.258228573 | 0.001643119 |
| Bicd1 | 2080.933512 | 0.257928764 | 0.000180507 |
| Pi4k2a | 1350.449517 | 0.257782429 | 0.001993439 |
| Nrxn3 | 9205.580118 | 0.257554578 | 0.000108706 |
| Gabbr2 | 12377.29593 | 0.257167864 | 0.014761318 |
| Arfgef2 | 2940.975944 | 0.256972479 | 0.000302271 |
| RGD1307554 | 790.5634895 | 0.256781798 | 0.001199925 |
| Dnajc2 | 1293.816097 | 0.256543305 | 0.008255152 |
| Atp13a3 | 123.5462763 | 0.256324698 | 0.045949989 |
| Prdm11 | 482.762597 | 0.256302859 | 0.022098947 |
| RGD1306502 | 1140.750491 | 0.255966852 | 0.002431783 |
| Gap43 | 10973.6049 | 0.255565887 | 0.011640947 |
| Nol4 | 1144.509154 | 0.255521581 | 0.006862861 |
| Adck1 | 261.5069414 | 0.254443872 | 0.015240775 |
| Pin1 | 2486.666882 | 0.254196119 | 0.000750972 |
| Rnf219 | 453.8169059 | 0.254084729 | 0.003985309 |
| Bnip3l | 4445.635701 | 0.253798992 | 4.03E-05 |
| Ing2 | 530.9298727 | 0.253411441 | 0.008303093 |
| Eef1e1 | 558.8569273 | 0.253310813 | 0.012112711 |
| Tram1l1 | 986.3423857 | 0.253239782 | 0.00330102 |
| RGD1559747 | 434.3586126 | 0.253191376 | 0.042835941 |
| Lmo4 | 3247.184688 | 0.25299875 | 0.016763909 |
| Extl2 | 1550.35156 | 0.252985602 | 0.000986792 |

| | | | |
|---|---|---|---|
| Taf5l | 397.1845257 | 0.252681203 | 0.011998927 |
| Sept6 | 1262.167543 | 0.252593093 | 0.001853535 |
| RGD621098 | 588.0470732 | 0.252096882 | 0.001268238 |
| Dohh | 1239.435294 | 0.252079566 | 0.005501953 |
| March9 | 449.3083165 | 0.251649992 | 0.018603318 |
| Spock2 | 7196.205338 | 0.251576305 | 0.000319381 |
| Hipk3 | 3536.255591 | 0.251352948 | 0.000900898 |
| Zbtb7a | 3444.651613 | 0.251106609 | 7.32E-06 |
| Ppp1cc | 6849.086691 | 0.251105393 | 1.16E-07 |
| Rufy3 | 4635.526721 | 0.250990992 | 5.55E-05 |
| Fry | 15613.77987 | 0.250980887 | 0.000508867 |
| Stim2 | 2011.964373 | 0.250630472 | 0.001506526 |
| Tmem160 | 935.3929829 | 0.25047886 | 0.023716728 |
| Zfp472 | 112.1973162 | 0.249923533 | 0.049158361 |
| Tef | 8433.124243 | 0.249800584 | 0.000241792 |
| Deaf1 | 750.4698222 | 0.249765768 | 0.004847825 |
| Zbed4 | 390.9748498 | 0.249737658 | 0.026356061 |
| Napa | 4049.347403 | 0.249653613 | 2.44E-05 |
| Rnf8 | 776.5897575 | 0.249410708 | 0.000466715 |
| Chd5 | 16189.50003 | 0.249265207 | 0.003293512 |
| Atp6v1h | 1646.307229 | 0.248866068 | 0.002529203 |
| Sclt1 | 419.492383 | 0.248857241 | 0.007244624 |
| Mesdc1 | 745.5546655 | 0.248658074 | 0.003537088 |
| Ankrd13d | 1319.064658 | 0.248609915 | 0.002359851 |
| Zdhhc5 | 1548.410883 | 0.248586831 | 0.003193465 |
| Mtrf1l | 281.4695639 | 0.248558485 | 0.023428763 |
| Calm3 | 9119.138713 | 0.248525994 | 0.000453076 |
| Rnft2 | 3540.554377 | 0.248504705 | 0.000178118 |
| Ppfia4 | 2443.128871 | 0.248462191 | 0.005471061 |
| Map4k2 | 807.83683 | 0.2481739 | 0.010838397 |
| Zfp133 | 436.3594295 | 0.248138601 | 0.028957545 |
| Ints7 | 901.5901984 | 0.248095101 | 0.007376732 |
| Gnl3 | 486.6704594 | 0.248003812 | 0.014355472 |
| Aff3 | 2325.8773 | 0.247929657 | 0.007614368 |
| Kcnb2 | 835.9645437 | 0.24788274 | 0.020661792 |
| Dync1li1 | 1674.068556 | 0.247749712 | 0.000470404 |

| | | | |
|---|---|---|---|
| Cyp46a1 | 5744.472912 | 0.247710807 | 0.001716607 |
| Lnp | 553.5663508 | 0.247568403 | 0.025551202 |
| Dos | 10418.42934 | 0.247549484 | 0.000474341 |
| Snapc5 | 887.5686906 | 0.247504553 | 0.000985354 |
| Zfp612 | 2056.649927 | 0.247216622 | 0.002309455 |
| Mcart1 | 1137.961894 | 0.24697638 | 0.000762658 |
| Srpk2 | 7368.921512 | 0.246820465 | 2.96E-07 |
| Dbndd1 | 536.3296535 | 0.246796317 | 0.012186494 |
| Rai1 | 4160.274879 | 0.246740346 | 0.00079829 |
| Pak7 | 793.5580803 | 0.246715499 | 0.045949989 |
| Sgsm2 | 3507.214212 | 0.246131664 | 0.001627209 |
| Pwp2 | 642.3579879 | 0.246017814 | 0.002203946 |
| Ppp1r12c | 3266.782646 | 0.245862808 | 4.28E-05 |
| Acta2 | 1.71218709 | 0.245781227 | 0.048486563 |
| Cd200 | 4207.031693 | 0.245451997 | 0.001499894 |
| Ywhab | 26732.08313 | 0.245290111 | 6.21E-05 |
| Ppm1l | 6611.168259 | 0.244808851 | 0.00666608 |
| Akt3 | 1747.783784 | 0.244462556 | 0.00087558 |
| Snx7 | 624.7553214 | 0.244184007 | 0.041346311 |
| Alcam | 7072.802246 | 0.243702603 | 0.00018998 |
| Glrb | 2344.196665 | 0.243461277 | 0.005420525 |
| Dhcr24 | 2450.650621 | 0.243180203 | 0.031390995 |
| Sepw1 | 2556.028648 | 0.243172838 | 0.014103617 |
| Atp6v0e2 | 2578.069723 | 0.242888287 | 0.001477497 |
| Mapk8ip1 | 11997.60798 | 0.242742673 | 0.000291427 |
| Mn1 | 3152.18817 | 0.242580998 | 0.00151134 |
| RGD1566265 | 533.5114856 | 0.242378743 | 0.015062779 |
| Armc10 | 808.5327589 | 0.242091411 | 0.010961628 |
| mrpl11 | 430.585289 | 0.242060179 | 0.008531479 |
| Adam11 | 2550.506363 | 0.241843614 | 0.016749792 |
| Rims4 | 305.2012292 | 0.241279045 | 0.047603579 |
| Acot7 | 6198.289602 | 0.241249872 | 0.001934682 |
| Zfyve9 | 4042.927321 | 0.240992512 | 6.49E-05 |
| Abhd8 | 5275.993168 | 0.2409496 | 0.000137421 |
| Ntmt1 | 352.4250667 | 0.240596573 | 0.014355472 |
| Bzw2 | 623.3634989 | 0.240393594 | 0.0068838 |

| | | | |
|---|---|---|---|
| Lipa | 1176.357977 | 0.240272867 | 0.003103241 |
| Snupn | 304.8225642 | 0.240253825 | 0.008070844 |
| Gria4 | 3334.99479 | 0.24024035 | 0.004007406 |
| Kif3b | 3191.88509 | 0.2401892 | 2.67E-05 |
| Pcdh9 | 5738.644062 | 0.239814367 | 0.028147469 |
| Pop5 | 374.5694652 | 0.239642109 | 0.034242731 |
| Strbp | 3845.219359 | 0.239461819 | 0.003129479 |
| Tesk1 | 1683.445436 | 0.238762545 | 0.001165579 |
| Acvr2a | 967.9992164 | 0.238761241 | 0.020378366 |
| Dmtf1 | 2378.288853 | 0.238760589 | 0.000177885 |
| Tyw5 | 232.3214207 | 0.238694388 | 0.048145681 |
| RGD1565498 | 290.6261825 | 0.238496802 | 0.032769116 |
| Wipf2 | 2025.235943 | 0.238272723 | 5.44E-05 |
| Mmp16 | 1601.296717 | 0.238212829 | 0.041580553 |
| Kcnd1 | 357.1597269 | 0.238104488 | 0.034116427 |
| St6galnac6 | 1201.368721 | 0.238067211 | 0.000644201 |
| Etv3 | 386.1438352 | 0.23758255 | 0.01833544 |
| Sdr39u1 | 706.077212 | 0.23747776 | 0.003962353 |
| Alkbh6 | 350.3590093 | 0.237341288 | 0.02194135 |
| Sobp | 2466.524708 | 0.237169214 | 0.014856178 |
| Morc3 | 786.3744725 | 0.236989723 | 0.017451702 |
| Dip2c | 4345.832635 | 0.236725583 | 0.000278043 |
| Dctn1 | 16623.16183 | 0.236503344 | 0.002813321 |
| Scg2 | 11816.06306 | 0.236323155 | 0.013360795 |
| Dpp6 | 8745.929779 | 0.236292375 | 0.005678888 |
| Ttc33 | 999.7464016 | 0.236290919 | 0.004057149 |
| Camk1d | 4782.979929 | 0.236278724 | 0.000789755 |
| Chchd4 | 474.1503501 | 0.236146896 | 0.002879401 |
| Jade2 | 2195.827539 | 0.235867704 | 0.000121964 |
| Zfp382 | 252.2176929 | 0.235649434 | 0.038035103 |
| Timm10 | 269.9583959 | 0.235168061 | 0.032400015 |
| Mtmr1 | 1756.334821 | 0.23505615 | 0.004388374 |
| Tenm3 | 15287.74017 | 0.234834305 | 0.041607351 |
| Tmem132a | 7837.715314 | 0.234520366 | 0.042134473 |
| Abhd17a | 2065.622634 | 0.234516872 | 0.00163179 |
| Tmem38a | 1218.159993 | 0.234356014 | 0.004555797 |

| | | | |
|---|---|---|---|
| Fdps | 2787.926627 | 0.234065562 | 0.024194578 |
| Ercc6 | 920.9440548 | 0.233993686 | 0.007170798 |
| Rcor2 | 588.9431633 | 0.233877455 | 0.0303609 |
| Mrpl54 | 873.9383278 | 0.233624443 | 0.002589762 |
| Timm8b | 1571.733685 | 0.233623321 | 0.015335953 |
| Man1a2 | 2515.775087 | 0.23345602 | 2.14E-05 |
| Thra | 11606.41436 | 0.233380339 | 0.010728816 |
| Sgsm3 | 1948.024811 | 0.23315325 | 0.000302085 |
| Wdr24 | 836.011779 | 0.233065059 | 0.005049339 |
| Nudt4 | 2784.36596 | 0.233015629 | 0.006632602 |
| Rasa1 | 2533.026912 | 0.232984848 | 0.000458326 |
| Mkrn1 | 838.6857406 | 0.23278724 | 0.000808968 |
| Usp33 | 3136.717779 | 0.232608168 | 0.000745657 |
| Maml3 | 298.8953166 | 0.232605653 | 0.035152458 |
| Tspyl4 | 8974.287634 | 0.23257431 | 0.000338898 |
| Zfp593 | 372.1149887 | 0.232328277 | 0.026705849 |
| Lcmt1 | 1166.102899 | 0.23214464 | 0.000602581 |
| Atf2 | 3395.988627 | 0.231992242 | 0.000378291 |
| Rps6kc1 | 1851.403061 | 0.231991125 | 0.000807738 |
| Cry2 | 5805.029497 | 0.231771561 | 0.001224132 |
| Rfng | 495.6780617 | 0.23175058 | 0.002118933 |
| Mgat5b | 1403.736943 | 0.231714965 | 0.022455063 |
| Tmem169 | 402.3399961 | 0.231368052 | 0.01339566 |
| Fbxo21 | 5287.937977 | 0.23122861 | 0.000120232 |
| Cyb561 | 1040.144525 | 0.231219717 | 0.029347973 |
| Slc8a3 | 1390.121939 | 0.230871411 | 0.010253665 |
| Ap2b1 | 15359.50914 | 0.230813423 | 0.000317517 |
| RGD1560108 | 769.4931235 | 0.230637908 | 0.014103617 |
| Prelid1 | 2517.21651 | 0.230630067 | 0.00124056 |
| Lrrtm4 | 452.3896748 | 0.230552599 | 0.029081623 |
| Them4 | 643.5245457 | 0.230295429 | 0.01249185 |
| Trim23 | 2286.144517 | 0.230186311 | 0.004972936 |
| Ulk1 | 2890.071648 | 0.230146086 | 1.64E-05 |
| Ppp2r2a | 3841.039453 | 0.230030926 | 0.003334037 |
| Slc35c1 | 711.662486 | 0.229847982 | 0.024791848 |
| Tmem183a | 929.9230026 | 0.229449738 | 0.000201158 |

| | | | |
|---|---|---|---|
| Wdr54 | 754.4994403 | 0.229003612 | 0.038243258 |
| Spire2 | 696.0995284 | 0.228511432 | 0.024273236 |
| Dexi | 1045.884198 | 0.228493717 | 0.011326605 |
| Fam13b | 4434.109404 | 0.228351452 | 0.000104103 |
| Fnip2 | 706.8006602 | 0.228259572 | 0.025518733 |
| Pld3 | 10527.85969 | 0.227659447 | 0.001304229 |
| Trim32 | 4124.723362 | 0.227563642 | 0.000181488 |
| Itpkc | 443.4151376 | 0.227525357 | 0.022292807 |
| Foxj3 | 3721.48124 | 0.227198576 | 0.000329521 |
| Actr2 | 4492.472123 | 0.226908967 | 3.06E-05 |
| Iqsec3 | 9502.457571 | 0.226896481 | 0.019410057 |
| Smpd3 | 4592.197108 | 0.226780903 | 0.001997153 |
| Celsr3 | 2080.002172 | 0.226702468 | 0.022265139 |
| Azin1 | 5130.351332 | 0.226596001 | 0.000732826 |
| Cyb5r4 | 541.7273502 | 0.226311603 | 0.026234398 |
| Mrps30 | 875.9904024 | 0.226245265 | 0.001061625 |
| Mfsd6 | 5808.720831 | 0.226155805 | 0.007542447 |
| Tcerg1 | 3350.127797 | 0.225845317 | 0.001971899 |
| RGD1562618 | 2542.097932 | 0.225409616 | 0.015062159 |
| Nap1l3 | 2367.383189 | 0.225368438 | 0.001361226 |
| Dnajb1 | 1606.045034 | 0.225256818 | 0.015762421 |
| Msl1 | 2822.358427 | 0.225178649 | 0.000195774 |
| Atp6v0c | 24005.34649 | 0.225115533 | 0.00036069 |
| Herc3 | 4933.588415 | 0.225061413 | 0.000158821 |
| Pim2 | 1241.476907 | 0.225001723 | 0.010171222 |
| Gars | 3851.457111 | 0.224911341 | 0.001090269 |
| Smarcd1 | 3115.164431 | 0.224822568 | 0.000750972 |
| Clcn4 | 5114.310366 | 0.224581321 | 0.001573222 |
| Mier2 | 630.3151497 | 0.224546049 | 0.005577018 |
| Pgam5 | 778.5849458 | 0.224384648 | 0.006137228 |
| Arhgef7 | 3460.238604 | 0.224366918 | 0.002228134 |
| Papolg | 599.4976242 | 0.224297666 | 0.018383561 |
| Fstl5 | 2238.107024 | 0.224216674 | 0.044192906 |
| Arrb2 | 955.7754762 | 0.224154109 | 0.00071455 |
| Atpif1 | 3060.029959 | 0.224121692 | 0.010711232 |
| Asic2 | 2142.992071 | 0.223818421 | 0.00982347 |

| | | | |
|---|---|---|---|
| Usp13 | 1682.11135 | 0.22374153 | 0.00109664 |
| Zswim3 | 458.645924 | 0.22367001 | 0.01480513 |
| Smyd5 | 472.9707098 | 0.223597063 | 0.028388887 |
| Ttll1 | 1033.47302 | 0.223562004 | 0.002821459 |
| Smurf1 | 1970.644877 | 0.223501788 | 0.014274113 |
| Rnf26 | 985.8649348 | 0.223491834 | 0.006706917 |
| Rhobtb2 | 3124.310062 | 0.223479726 | 0.001166624 |
| Mtmr4 | 3952.080468 | 0.22343185 | 0.002067744 |
| RGD1309188 | 2859.027793 | 0.223405901 | 7.38E-05 |
| Rab3ip | 405.2453516 | 0.223179441 | 0.022342754 |
| Cdk14 | 4234.546068 | 0.223129828 | 0.001826382 |
| Ube2j1 | 999.2286738 | 0.223056728 | 0.000924654 |
| Becn1 | 2919.696212 | 0.222989123 | 0.000643477 |
| Mrps34 | 1138.580919 | 0.222967578 | 0.004668556 |
| Isca1 | 4206.475654 | 0.222937757 | 0.00012803 |
| Kcnd2 | 4122.699706 | 0.222797397 | 0.000714909 |
| Mmaa | 343.7686525 | 0.222681662 | 0.010171222 |
| Akap8l | 1037.959237 | 0.222555956 | 0.004863417 |
| Apbb1 | 8477.600845 | 0.222201463 | 0.008660728 |
| Uqcc2 | 855.9781786 | 0.222127218 | 0.007403216 |
| Rabif | 698.935269 | 0.221978147 | 0.015565982 |
| Strn | 3446.232944 | 0.22133239 | 0.003719026 |
| Msto1 | 514.6625705 | 0.221184397 | 0.024791848 |
| Armc8 | 1708.899674 | 0.221071009 | 0.000807738 |
| Prkag2 | 1167.694656 | 0.220873601 | 0.028326192 |
| Dcaf12 | 2174.360251 | 0.220775811 | 0.021734435 |
| Eml2 | 4460.322475 | 0.22074834 | 0.000739935 |
| Nceh1 | 3078.681915 | 0.220708834 | 0.001885195 |
| ST7 | 422.4956422 | 0.220551783 | 0.03476064 |
| Gcc2 | 3570.627576 | 0.220352795 | 0.039116775 |
| Ogfod1 | 1528.478683 | 0.220340678 | 0.002969217 |
| Lmbr1 | 920.4346711 | 0.220333159 | 0.003078036 |
| Tubgcp5 | 702.8150639 | 0.220323318 | 0.006320827 |
| Arpc4 | 4407.733502 | 0.220322475 | 0.001371288 |
| Tmem260 | 1106.519456 | 0.220242724 | 0.005047001 |
| Ube2k | 3931.738519 | 0.219987229 | 1.40E-05 |

| | | | |
|---|---|---|---|
| Actr1b | 8200.61077 | 0.2198579 | 0.000109291 |
| Dhx57 | 1628.453967 | 0.219618961 | 0.004232362 |
| Stk39 | 2002.020759 | 0.219587254 | 0.009034189 |
| Tspan13 | 2799.396172 | 0.219448585 | 0.00136991 |
| Twf2 | 481.9621796 | 0.219435612 | 0.046138112 |
| Cbx7 | 1507.068326 | 0.219374042 | 0.006540692 |
| Pgam1 | 14722.83279 | 0.219337662 | 0.000304541 |
| Ndufa5 | 987.7359866 | 0.21916017 | 0.016959761 |
| Gmeb1 | 330.4872286 | 0.219057349 | 0.025210033 |
| Rcor1 | 684.0751067 | 0.218626328 | 0.006950121 |
| Fen1 | 326.3746124 | 0.218558821 | 0.045139158 |
| Fyttd1 | 1707.439882 | 0.218147552 | 0.013224588 |
| Tcf20 | 6921.630501 | 0.217849076 | 1.92E-05 |
| Fam168b | 9147.138477 | 0.217790891 | 4.74E-11 |
| Lrfn5 | 936.9472377 | 0.217763673 | 0.045631608 |
| Msh3 | 350.8195515 | 0.217665804 | 0.035268421 |
| Creld1 | 1988.769506 | 0.217632218 | 0.000314692 |
| Cyth3 | 1000.765789 | 0.217622514 | 0.014087389 |
| Gad1 | 19063.36109 | 0.217454674 | 0.028786824 |
| Etnk1 | 9247.990231 | 0.217033176 | 6.00E-05 |
| Epha4 | 6512.508245 | 0.216756117 | 0.013543385 |
| Dda1 | 1729.486524 | 0.216684533 | 0.000446175 |
| RGD1311899 | 7734.242684 | 0.216555033 | 0.000938391 |
| Gpd1l | 1487.974073 | 0.216464415 | 0.008228415 |
| Asns | 1655.196027 | 0.216398545 | 0.007193277 |
| Gmcl1 | 873.8345685 | 0.216279093 | 0.018268271 |
| Prps1 | 3521.308198 | 0.216242524 | 0.001508777 |
| Mettl13 | 700.8309721 | 0.216189881 | 0.0099772 |
| Ints2 | 655.3231667 | 0.21615287 | 0.034483641 |
| Carm1 | 2790.340369 | 0.216060571 | 0.000822724 |
| Zfyve27 | 2076.892308 | 0.215611686 | 0.008592167 |
| Kras | 675.5301772 | 0.215561876 | 0.014326581 |
| Eml5 | 614.8973571 | 0.215110993 | 0.046823404 |
| Csrnp2 | 1164.888206 | 0.215025524 | 0.016782289 |
| B3galt1 | 1075.484696 | 0.214881853 | 0.003020495 |
| Hps3 | 582.1514402 | 0.214777303 | 0.018882978 |

| | | | |
|---|---|---|---|
| Dyrk1b | 1248.854111 | 0.214348797 | 0.000119664 |
| Nphp4 | 437.5757143 | 0.214131309 | 0.02522534 |
| Hn1 | 994.9462362 | 0.214107438 | 0.022859062 |
| Lmbrd2 | 2607.983388 | 0.213866088 | 0.007960743 |
| Pygo2 | 1591.851849 | 0.213823891 | 0.001488923 |
| Ssh2 | 1335.655136 | 0.213748003 | 0.019107627 |
| Usp14 | 4389.964147 | 0.213650472 | 0.000472085 |
| Gorasp2 | 3264.054289 | 0.213641732 | 0.000374138 |
| Acot9 | 810.992468 | 0.213585778 | 0.005648352 |
| Rnf44 | 2004.359883 | 0.213573981 | 0.000280326 |
| Pde4a | 1923.605301 | 0.213492138 | 0.031781318 |
| Gale | 288.1014038 | 0.213397016 | 0.028880221 |
| Mrpl17 | 406.5839447 | 0.213298881 | 0.028199717 |
| Fam193b | 370.9307986 | 0.213282234 | 0.029737113 |
| Ddx55 | 387.7725831 | 0.2129646 | 0.015565982 |
| Slc9a1 | 2258.25943 | 0.212895318 | 0.003348448 |
| LOC100365289 | 619.6089215 | 0.212577694 | 0.012615717 |
| Ube4b | 5411.295007 | 0.212248261 | 0.000743626 |
| Brms1l | 1558.468336 | 0.212173655 | 0.000553886 |
| Carf | 361.0916708 | 0.212118924 | 0.045894711 |
| Hmox2 | 2457.007536 | 0.211205878 | 0.004693495 |
| Opa1 | 4367.884277 | 0.211090234 | 0.000151281 |
| Tpd52 | 362.1171594 | 0.21082163 | 0.048190413 |
| Cep290 | 1140.705095 | 0.210681999 | 0.041020515 |
| Capn15 | 1357.265025 | 0.210279508 | 0.021368972 |
| Rab6a | 17056.78481 | 0.210150324 | 0.002764571 |
| Pde4dip | 8825.373113 | 0.210033059 | 0.001276088 |
| Eif4enif1 | 1708.401803 | 0.209981188 | 0.005826824 |
| Tmem151b | 2624.975052 | 0.209791166 | 0.012743907 |
| Sfxn3 | 2145.094785 | 0.209215101 | 0.00611085 |
| Tbl1xr1 | 4750.066843 | 0.209027037 | 0.000241895 |
| Atp5g1 | 1526.602983 | 0.209015276 | 0.004323685 |
| Ublcp1 | 1574.813295 | 0.209014133 | 0.006561183 |
| Prune | 765.9150161 | 0.208707455 | 0.031752121 |
| Tiprl | 979.5365699 | 0.208649856 | 0.005498012 |
| Cmss1 | 362.3075647 | 0.208585132 | 0.044177034 |

| | | | |
|---|---|---|---|
| Iffo1 | 1012.114889 | 0.208311975 | 0.007346534 |
| Chpf | 4575.372284 | 0.208293237 | 9.75E-07 |
| Rnf19b | 498.3155994 | 0.208261878 | 0.018847522 |
| RGD1563072 | 896.645288 | 0.207987874 | 0.012768029 |
| Pwwp2a | 556.1298429 | 0.207939745 | 0.023759929 |
| Eif4a2 | 13228.59536 | 0.207844978 | 0.000544843 |
| Ndufa4 | 3551.759484 | 0.20775305 | 0.000302767 |
| Rprd1a | 4107.191064 | 0.207650384 | 2.16E-07 |
| Pnmal2 | 21043.17493 | 0.207323309 | 0.001935512 |
| Nipa1 | 991.1839626 | 0.207318406 | 0.008184981 |
| Ddx19b | 1148.149077 | 0.207240436 | 0.001494138 |
| Nus1 | 2175.002413 | 0.207184041 | 0.001411045 |
| Ogt | 4443.054037 | 0.206918507 | 0.007562529 |
| Spred2 | 1232.026267 | 0.206910167 | 0.006818606 |
| Trappc2l | 867.8543319 | 0.206747082 | 0.007061091 |
| Gatsl2 | 9754.255126 | 0.206525397 | 0.000387782 |
| Rrm2b | 499.4925239 | 0.206466348 | 0.004614319 |
| Fam73b | 2548.016108 | 0.205749514 | 0.00010346 |
| Atxn3 | 410.1227805 | 0.205680956 | 0.039293284 |
| Snn | 3568.235165 | 0.205673431 | 0.019652107 |
| Dyrk2 | 2150.125742 | 0.205643047 | 0.046209511 |
| Cyb5r1 | 1025.683271 | 0.205527135 | 0.037723974 |
| Gltscr2 | 1534.764784 | 0.205482664 | 0.002929144 |
| Ndufaf5 | 811.5261436 | 0.20545792 | 0.013943469 |
| Slc12a6 | 4003.954295 | 0.205330548 | 0.000287459 |
| Pds5b | 7749.411411 | 0.205276722 | 0.000605978 |
| Urb1 | 702.9362531 | 0.205257348 | 0.036551361 |
| Irgq | 7131.569024 | 0.205204555 | 4.11E-05 |
| Dynll2 | 13908.28363 | 0.205176912 | 0.0018302 |
| Htt | 7231.213745 | 0.205144204 | 0.010357265 |
| Rnmtl1 | 261.8957428 | 0.204991713 | 0.049542627 |
| Coa3 | 1058.037968 | 0.204986414 | 0.010155019 |
| Clip4 | 614.1644408 | 0.204741305 | 0.040433401 |
| Slc30a4 | 2549.31128 | 0.204723626 | 0.00239052 |
| RGD1563986 | 729.6346879 | 0.204692392 | 0.028880221 |
| Adgrb2 | 13139.2254 | 0.204547326 | 0.008624426 |

| | | | |
|---|---|---|---|
| Ankh | 4216.222521 | 0.204155335 | 0.003117345 |
| Scamp1 | 6589.517321 | 0.203864397 | 5.75E-05 |
| Lancl2 | 1453.523388 | 0.203529469 | 0.00343417 |
| Smg9 | 654.703111 | 0.203492639 | 0.016388142 |
| Pacs1 | 3167.370121 | 0.202957463 | 0.016990993 |
| Ppp1ca | 4836.616965 | 0.202956687 | 0.007156638 |
| Rnf6 | 3528.844396 | 0.202864525 | 0.000489973 |
| Kcna6 | 4890.439814 | 0.202672872 | 0.023568933 |
| Diaph1 | 1742.852887 | 0.20254747 | 0.018707905 |
| Kctd3 | 2622.602867 | 0.202478561 | 0.001408071 |
| Lrfn4 | 677.8583822 | 0.201895032 | 0.028804276 |
| Pitpnm1 | 7117.81941 | 0.20182012 | 0.002350532 |
| Cdh13 | 9267.272864 | 0.201783024 | 0.027822389 |
| Mak16 | 538.0146087 | 0.201697349 | 0.01796069 |
| Cnot1 | 5803.872747 | 0.201309184 | 9.80E-05 |
| Cap1 | 7711.667143 | 0.201290322 | 0.006154176 |
| Dynlt3 | 2722.423837 | 0.201153429 | 0.018181884 |
| Gpr137 | 1671.872324 | 0.200645458 | 0.005385166 |
| Ppp1r2 | 5152.227819 | 0.200514659 | 0.000739935 |
| Csnk1g3 | 1984.59405 | 0.20008735 | 0.003193465 |
| Ccdc174 | 491.3030612 | 0.200000292 | 0.033289315 |
| Usp22 | 4504.22679 | 0.199949743 | 0.002032363 |
| Btrc | 1986.882995 | 0.199949566 | 0.001665415 |
| Acsl1 | 1277.935636 | 0.199787689 | 0.008453468 |
| Cnnm2 | 1463.373529 | 0.199652804 | 0.01364181 |
| Dtd1 | 954.4165293 | 0.199535518 | 0.005722311 |
| Zfp281 | 992.0659716 | 0.199460501 | 0.013941943 |
| Spg7 | 1393.109939 | 0.199363458 | 0.001027672 |
| Atp6v0a1 | 18998.16182 | 0.199344474 | 0.002805549 |
| Fam134a | 6139.650672 | 0.199257439 | 3.34E-05 |
| Wdr44 | 980.0023494 | 0.199206704 | 0.002431391 |
| Mypop | 1205.447633 | 0.199020346 | 0.015229121 |
| Snx30 | 1064.853866 | 0.199003003 | 0.015027858 |
| Tnfrsf21 | 3103.336021 | 0.198897024 | 0.01445176 |
| Ppp1r37 | 3628.084641 | 0.198635258 | 0.002390954 |
| Mrps2 | 1604.526567 | 0.198630157 | 0.001668857 |

| | | | |
|---|---|---|---|
| Tm2d2 | 1522.234442 | 0.198325156 | 0.001670795 |
| Dusp26 | 1526.893373 | 0.19813041 | 0.010724593 |
| Gramd1a | 1638.835145 | 0.197848715 | 0.017846658 |
| Chst1 | 5786.733574 | 0.197795601 | 0.001622167 |
| Fbxo11 | 4489.375774 | 0.197778173 | 0.000807384 |
| Tarsl2 | 751.4830349 | 0.197613601 | 0.038961336 |
| Zfp251 | 766.5389527 | 0.197477534 | 0.019635839 |
| Rtn3 | 34020.92339 | 0.197253651 | 0.000203179 |
| Hdac11 | 6621.141168 | 0.197183393 | 0.007186984 |
| Sugp2 | 3035.917108 | 0.196768152 | 0.002407846 |
| Bap1 | 5295.910918 | 0.196563174 | 0.007542447 |
| Aftph | 2879.054151 | 0.196489617 | 1.98E-07 |
| Fam69b | 1388.420664 | 0.196432519 | 0.01827172 |
| Ctnna2 | 5400.368161 | 0.196346933 | 0.000688449 |
| Slc4a1ap | 1578.002272 | 0.196292715 | 0.007110107 |
| Abl2 | 1547.858677 | 0.196145165 | 0.000295888 |
| Vps45 | 1382.651616 | 0.196009091 | 0.000351879 |
| Atp6v1d | 7950.462657 | 0.195997525 | 0.001267362 |
| Wdr74 | 341.5049759 | 0.195885929 | 0.046647585 |
| Sema4g | 1430.646565 | 0.195564276 | 0.027470381 |
| Mapre2 | 9439.167747 | 0.195536263 | 1.22E-07 |
| Zdhhc21 | 780.3282322 | 0.195265665 | 0.038089619 |
| Pafah1b1 | 20814.79089 | 0.194892431 | 0.000552708 |
| Jmjd6 | 371.5779512 | 0.194737219 | 0.04001138 |
| Enpp5 | 8835.824767 | 0.194113924 | 7.33E-07 |
| Spats2 | 1212.177429 | 0.194112768 | 0.010040959 |
| Trim9 | 4401.499134 | 0.193993089 | 0.046876486 |
| Mapt | 24782.31246 | 0.193155321 | 2.62E-05 |
| Casp9 | 915.0900649 | 0.193009604 | 0.004456536 |
| Baz2a | 2324.984924 | 0.192938504 | 0.006120222 |
| Pdxk | 1207.586204 | 0.192924557 | 0.01918716 |
| Ncoa6 | 4396.059471 | 0.192909023 | 0.002529843 |
| Polrmt | 903.2477486 | 0.192792572 | 0.015083176 |
| Lrrc49 | 1875.911096 | 0.192767404 | 0.001787253 |
| Serinc1 | 23631.1076 | 0.192522648 | 0.001812609 |
| Mfsd12 | 1015.667145 | 0.19250732 | 0.024846095 |

| | | | |
|---|---|---|---|
| Map1s | 2949.080984 | 0.19239351 | 0.005240097 |
| Kcnip2 | 2032.534758 | 0.192009395 | 0.022716434 |
| Tulp4 | 7763.649447 | 0.191973464 | 0.006660123 |
| Lhfpl4 | 725.7504569 | 0.19188671 | 0.010434904 |
| Grb2 | 6518.269622 | 0.191712159 | 1.49E-06 |
| Reep5 | 12253.51891 | 0.19153823 | 0.001599256 |
| Bin1 | 4536.765075 | 0.190991985 | 0.022749792 |
| Dnajc16 | 1593.290056 | 0.190793024 | 0.009991179 |
| Mphosph8 | 2592.900731 | 0.190616304 | 0.004436933 |
| Tmem30a | 6054.598165 | 0.190507727 | 0.000177647 |
| Fam76a | 1278.902289 | 0.190501035 | 0.042409248 |
| Irf2bpl | 3089.764758 | 0.190066443 | 0.016352527 |
| Map1lc3a | 4421.479737 | 0.190065902 | 0.003722618 |
| Cinp | 600.180506 | 0.18978633 | 0.044436976 |
| Disp2 | 22706.58824 | 0.189566607 | 0.023358607 |
| Ahdc1 | 4872.301282 | 0.189462516 | 0.008453936 |
| Tmem229b | 831.0804589 | 0.189085192 | 0.049542627 |
| Rnf123 | 3226.97496 | 0.188877455 | 0.004093876 |
| Rtkn | 2172.075337 | 0.188830841 | 0.015870919 |
| March6 | 11086.75189 | 0.188795651 | 0.000220266 |
| Parp1 | 2644.598994 | 0.18861192 | 0.001221725 |
| Nlgn2 | 10590.37516 | 0.1882734 | 0.000155747 |
| Armt1 | 669.7703522 | 0.188235248 | 0.010077968 |
| Nomo1 | 6640.933229 | 0.188232334 | 0.024399304 |
| Prkca | 5667.237106 | 0.188038765 | 0.004668556 |
| Bclaf1 | 4188.488163 | 0.188019918 | 0.025551202 |
| Rhbdl3 | 963.2538867 | 0.187786443 | 0.028130953 |
| Ophn1 | 1141.243994 | 0.187706661 | 0.03191949 |
| Zfp638 | 2478.224753 | 0.187590279 | 0.020223816 |
| Triqk | 341.8155296 | 0.187589312 | 0.048925225 |
| Ubn1 | 1969.738165 | 0.187448665 | 0.009584967 |
| Slc30a5 | 1280.217241 | 0.187402029 | 0.001508777 |
| Fkbp3 | 3428.539965 | 0.187300748 | 0.006825507 |
| Zmynd19 | 552.4091946 | 0.187180024 | 0.03349982 |
| Atp6v1f | 3060.208962 | 0.187121332 | 0.020309011 |
| Ralgapa1 | 3548.200269 | 0.186836679 | 0.011559278 |

| | | | |
|---|---|---|---|
| Metap1 | 1503.313395 | 0.186634699 | 0.002517418 |
| Mapk14 | 1944.552805 | 0.186596482 | 0.002582768 |
| Tnpo2 | 5548.637907 | 0.186593125 | 0.00069639 |
| Vprbp | 1519.897682 | 0.186536964 | 0.012546984 |
| Utp18 | 744.5757584 | 0.18643785 | 0.025584714 |
| Sptlc2 | 1431.216188 | 0.186361298 | 0.005976526 |
| Maz | 1269.102831 | 0.186359064 | 0.013898362 |
| Sh3glb2 | 3379.675072 | 0.186307307 | 0.031979674 |
| Anapc7 | 1050.432194 | 0.186142954 | 0.034958215 |
| Adgrb3 | 5330.177955 | 0.186065619 | 0.000878822 |
| Tsc1 | 1691.514064 | 0.186042999 | 0.00388751 |
| Denr | 455.1595989 | 0.185755384 | 0.033251434 |
| Ncam2 | 4696.092489 | 0.185695841 | 0.036721495 |
| Usp32 | 8272.503322 | 0.185439818 | 4.51E-05 |
| Peo1 | 1218.178231 | 0.185304422 | 0.002263206 |
| Cdipt | 3501.123342 | 0.185194358 | 0.001806163 |
| Vps29 | 1855.363553 | 0.185096992 | 0.010785378 |
| Amigo1 | 1248.858311 | 0.185052745 | 0.044712624 |
| Dagla | 4546.734868 | 0.184992799 | 0.015416695 |
| Gtpbp2 | 724.438792 | 0.184896694 | 0.020150866 |
| Camsap2 | 11571.44825 | 0.184615228 | 4.21E-05 |
| Rnf185 | 1276.193004 | 0.184162305 | 0.002629115 |
| Adrbk1 | 3794.186435 | 0.184123036 | 0.012424718 |
| Cenpc | 636.0835802 | 0.183972448 | 0.034952535 |
| Mark4 | 4025.506234 | 0.183785115 | 0.011188562 |
| Nipsnap1 | 3138.437704 | 0.183370899 | 0.000421923 |
| Ppm1a | 2623.804024 | 0.183328055 | 0.000985354 |
| Ppp2r2b | 4041.037771 | 0.183247139 | 0.012589323 |
| Tmem11 | 826.8585736 | 0.182995863 | 0.009363804 |
| Pafah1b2 | 4599.636598 | 0.182985865 | 0.000173206 |
| Apbb3 | 606.3173077 | 0.1824266 | 0.037202958 |
| Prdx5 | 3156.457212 | 0.182422411 | 0.001471664 |
| Zfp292 | 2904.150936 | 0.181939527 | 0.040516177 |
| Arfgef1 | 4928.716415 | 0.181922551 | 0.000776891 |
| Prepl | 10521.80452 | 0.181867819 | 0.014418621 |
| RGD1306941 | 894.80616 | 0.181745772 | 0.021677558 |

| | | | |
|---|---|---|---|
| Armc1 | 1748.797068 | 0.181013672 | 0.001786827 |
| Kiaa0408 | 3571.251309 | 0.180636814 | 0.000338454 |
| Guk1 | 2235.34093 | 0.18060362 | 0.041972194 |
| Yipf4 | 1040.414844 | 0.180525974 | 0.032679872 |
| Brd4 | 8153.925798 | 0.180504449 | 0.000359747 |
| Mcm3ap | 2555.729452 | 0.180348151 | 0.004521245 |
| Clstn3 | 14756.93221 | 0.18002793 | 0.011688362 |
| Srpk1 | 1572.850981 | 0.179821722 | 0.005357739 |
| Gdi1 | 23122.2481 | 0.179516811 | 0.001467613 |
| Aes | 6912.019825 | 0.179500477 | 0.001935512 |
| Trim37 | 5972.502025 | 0.178881224 | 3.32E-05 |
| Ocrl | 1670.509264 | 0.178807714 | 0.037096606 |
| Flrt2 | 1967.518118 | 0.178746268 | 0.011622698 |
| Asb6 | 659.7715371 | 0.178572155 | 0.028887399 |
| Cul2 | 1626.120021 | 0.178487059 | 0.007551724 |
| Mfsd11 | 785.0170104 | 0.178238866 | 0.013869122 |
| Irs2 | 4728.010264 | 0.178124659 | 0.040404112 |
| Proser1 | 912.0133824 | 0.177970756 | 0.008360219 |
| Papd5 | 2048.19582 | 0.177557318 | 0.001609267 |
| Uqcr11 | 3566.626453 | 0.177542373 | 0.016367712 |
| Phospho2 | 615.8955798 | 0.176879497 | 0.023279185 |
| Ddx25 | 958.0608219 | 0.176711937 | 0.022220245 |
| Fam49b | 3246.795299 | 0.176619235 | 0.002774461 |
| Shc3 | 2809.128038 | 0.176372603 | 0.038470369 |
| Polr1a | 1015.651252 | 0.176274463 | 0.039307832 |
| Zfp523 | 1054.238494 | 0.176155886 | 0.040566434 |
| Foxk2 | 2778.331666 | 0.176078442 | 0.003182171 |
| Gba | 3620.962838 | 0.175997919 | 0.01039212 |
| Stx16 | 1192.768717 | 0.175988487 | 0.002255168 |
| Thsd7a | 1338.64745 | 0.175549718 | 0.043184844 |
| Matr3 | 16284.76894 | 0.175081698 | 0.000204097 |
| Slc25a23 | 2074.081531 | 0.174919459 | 0.011856678 |
| Otub1 | 6447.628076 | 0.174630646 | 0.002519449 |
| Slk | 3971.984531 | 0.174498455 | 0.005945241 |
| Rnf11l1 | 2774.154077 | 0.174313214 | 0.004798663 |
| Thoc3 | 1014.115047 | 0.17410641 | 0.028703089 |

| | | | |
|---|---|---|---|
| Fam174a | 687.4241433 | 0.174010008 | 0.046103631 |
| Crbn | 1684.994438 | 0.173725665 | 0.007373971 |
| Vopp1 | 3392.005387 | 0.173615219 | 0.020475024 |
| Rnf145 | 2362.926722 | 0.173532094 | 0.014845903 |
| Trappc6b | 1153.947532 | 0.173424844 | 0.010230085 |
| Prkaca | 8267.592449 | 0.173325746 | 9.94E-05 |
| Tdg | 674.8527528 | 0.173284742 | 0.044884491 |
| Dgke | 1674.843285 | 0.173204719 | 0.034542238 |
| Zfp346 | 980.0897702 | 0.173177962 | 0.012388826 |
| Shoc2 | 1823.946523 | 0.173032041 | 0.001938589 |
| Med9 | 774.0559368 | 0.173003711 | 0.027973087 |
| Lgalsl | 2890.328057 | 0.172917871 | 0.006292511 |
| Prmt5 | 2076.48018 | 0.172835471 | 0.012023079 |
| Rfxap | 1105.871477 | 0.17273111 | 0.027383348 |
| Usp5 | 7733.16852 | 0.171818188 | 0.002320693 |
| Gpi | 21390.10991 | 0.17162504 | 0.002480053 |
| Tatdn2 | 1720.03372 | 0.171618052 | 0.00431713 |
| Entpd4 | 1388.151119 | 0.171397362 | 0.026153523 |
| Med1 | 2359.571159 | 0.170822385 | 0.005310992 |
| Usp15 | 2024.04306 | 0.170515604 | 0.000403614 |
| Dgcr14 | 589.1663939 | 0.17048557 | 0.023702895 |
| Pdk3 | 1203.226457 | 0.170373916 | 0.018538805 |
| Zbtb38 | 2604.874971 | 0.170289339 | 0.007807209 |
| Ppp1r12a | 3631.183174 | 0.170254201 | 0.00168027 |
| Gtf3c1 | 7755.521567 | 0.169996491 | 0.008579217 |
| Nedd8 | 2221.786619 | 0.16996025 | 0.049194163 |
| Sh3bp5l | 1275.629302 | 0.169840186 | 0.014552916 |
| Tbc1d10b | 3312.819139 | 0.169716533 | 0.00200133 |
| Sbf1 | 8944.917765 | 0.16967302 | 0.041635388 |
| Zc3h15 | 3826.039139 | 0.169659222 | 0.003110546 |
| Ube3b | 2828.329077 | 0.169533842 | 0.047379645 |
| Igsf8 | 3869.812526 | 0.169514754 | 0.018326777 |
| Lypla2 | 1351.305875 | 0.169408097 | 0.01380877 |
| Sucla2 | 4824.806339 | 0.169305682 | 0.004589557 |
| Tmem199 | 1105.143515 | 0.168253448 | 0.043844729 |
| Epn1 | 6518.075484 | 0.168151367 | 0.008189185 |

| | | | |
|---|---|---|---|
| Pex14 | 1297.059131 | 0.168140132 | 0.00761303 |
| Dpp9 | 2060.992856 | 0.168123974 | 0.007813569 |
| Ube2i | 1480.177628 | 0.167957922 | 0.016401421 |
| Fnip1 | 2568.972916 | 0.167211332 | 0.019842018 |
| Casp8ap2 | 934.4647539 | 0.167001094 | 0.018792624 |
| Fbxw2 | 1270.931524 | 0.166882442 | 0.04320649 |
| Uba2 | 1281.047815 | 0.166611593 | 0.009221237 |
| Cpeb4 | 4061.285764 | 0.166325893 | 0.017831084 |
| Cops8 | 1639.458787 | 0.166235462 | 0.003179893 |
| Ighmbp2 | 808.6922358 | 0.166053812 | 0.012074205 |
| Chp1 | 5699.733772 | 0.165854782 | 0.000113621 |
| Larp7 | 899.5958161 | 0.16576273 | 0.018847522 |
| Crebzf | 1277.951638 | 0.165498434 | 0.043684034 |
| Nupl1 | 1312.679685 | 0.165376231 | 0.009627444 |
| Narf | 2314.713003 | 0.165307881 | 0.012746678 |
| Pak3 | 3029.871412 | 0.165261578 | 0.02835069 |
| Ago2 | 2824.45661 | 0.165158407 | 0.041370607 |
| Naa38 | 1344.305445 | 0.165081668 | 0.025082747 |
| Dyrk1a | 2908.854535 | 0.165055941 | 0.000596735 |
| Egln2 | 2144.924538 | 0.164941704 | 0.002432611 |
| Fkbp8 | 9098.798733 | 0.164598233 | 0.013868489 |
| Atp13a2 | 7498.10625 | 0.164585201 | 0.009767043 |
| Ubl4a | 1045.754321 | 0.164506986 | 0.046993649 |
| Pknox2 | 1255.834521 | 0.164397243 | 0.025855867 |
| Arf1 | 4796.54919 | 0.164320052 | 0.02841553 |
| Mtch1 | 13577.32724 | 0.164309277 | 0.001611511 |
| Zfp955a | 1573.47445 | 0.164057898 | 0.012992104 |
| Prrt1 | 5794.181967 | 0.164042997 | 0.041739702 |
| Ralgapb | 6608.737685 | 0.163685072 | 0.003582725 |
| Fam219a | 3190.529115 | 0.163464213 | 0.001410555 |
| Ap3m2 | 1497.00886 | 0.163384166 | 0.016401421 |
| Grk6 | 1330.700704 | 0.163244249 | 0.019180985 |
| Fbll1 | 1463.963315 | 0.163057198 | 0.047393411 |
| Kctd10 | 1594.751079 | 0.16304608 | 0.002944628 |
| Zdhhc17 | 2822.822377 | 0.163041992 | 0.000238279 |
| Rc3h2 | 3595.654485 | 0.16243151 | 0.001488923 |

| | | | |
|---|---|---|---|
| Psmc6 | 2551.243419 | 0.16225858 | 0.000910537 |
| Cdc27 | 2178.43717 | 0.162202424 | 0.002582768 |
| Bop1 | 1150.59781 | 0.1621654 | 0.017418257 |
| Pfdn6 | 1136.118059 | 0.16174085 | 0.041453687 |
| Foxj2 | 2047.359645 | 0.161604248 | 0.035996998 |
| Iars | 2989.943945 | 0.16112918 | 0.008353743 |
| Slc25a46 | 2403.319016 | 0.160974526 | 0.021417743 |
| Ube2n | 3164.903769 | 0.160903814 | 0.001396443 |
| Slc22a17 | 25239.06431 | 0.160844546 | 0.002213779 |
| Col4a3bp | 3485.575772 | 0.160483484 | 0.006530331 |
| Bace1 | 2135.159121 | 0.160383209 | 0.023918345 |
| Tubb5 | 20390.39505 | 0.160259853 | 0.00431713 |
| Fkbpl | 495.219176 | 0.159859629 | 0.025139265 |
| Nfyb | 870.3185459 | 0.159341846 | 0.04492952 |
| Mcf2l | 3494.102953 | 0.159025155 | 0.035419657 |
| Kat2a | 2330.720179 | 0.158959688 | 0.020040925 |
| Megf9 | 4253.022403 | 0.158775007 | 0.020955753 |
| Larp4b | 5088.251408 | 0.158698977 | 0.00087558 |
| RGD1310429 | 1866.717924 | 0.158694369 | 0.022474742 |
| Peli1 | 1312.972755 | 0.158366203 | 0.030057934 |
| Rtn4 | 17230.10271 | 0.158120195 | 0.039236585 |
| Nr3c2 | 1986.769389 | 0.158105475 | 0.00866383 |
| Thumpd1 | 2141.828528 | 0.157590065 | 0.004170474 |
| Tnpo3 | 2540.93263 | 0.157554526 | 0.005580794 |
| Nop58 | 1037.474071 | 0.15738886 | 0.035152458 |
| Slc7a14 | 3790.217104 | 0.157231941 | 0.037646631 |
| Dync1li2 | 10053.48991 | 0.156950897 | 4.53E-05 |
| Tmem178b | 7115.525289 | 0.156944442 | 0.027399532 |
| Rnf19a | 1504.513737 | 0.156780921 | 0.025310298 |
| Sec23a | 3052.855615 | 0.156726541 | 0.0050527 |
| Znrf2 | 768.8359898 | 0.156713274 | 0.038470369 |
| Hint1 | 2762.123526 | 0.156672548 | 0.011579831 |
| Zfp46 | 1716.87552 | 0.156583733 | 0.01109108 |
| Vps53 | 2204.279819 | 0.156571735 | 0.003531611 |
| Stx6 | 2998.557846 | 0.156211384 | 0.005858461 |
| H2afz | 1387.947585 | 0.156116567 | 0.033103206 |

| | | | |
|---|---|---|---|
| Abi2 | 5996.445011 | 0.156047196 | 0.020251314 |
| Alg2 | 5786.154785 | 0.155972403 | 0.011759848 |
| Vps41 | 7781.02206 | 0.155938156 | 0.010171222 |
| Gba2 | 1525.754936 | 0.155412578 | 0.020413499 |
| Zfp644 | 3055.808838 | 0.155334417 | 0.00473992 |
| Usp45 | 777.0309794 | 0.155028584 | 0.034121336 |
| Coq6 | 645.3480753 | 0.155025117 | 0.03164229 |
| Aagab | 738.2775668 | 0.1549532 | 0.042112327 |
| Ppat | 887.0005682 | 0.15458995 | 0.033366286 |
| Ctnnbip1 | 1021.115448 | 0.154314071 | 0.0406727 |
| Slc35e1 | 2249.315869 | 0.154125741 | 0.00313526 |
| Usp11 | 6778.010172 | 0.153834163 | 0.041392548 |
| Foxk1 | 1560.721874 | 0.153383179 | 0.048678397 |
| Capza2 | 4879.740001 | 0.153054118 | 0.012462908 |
| Cds2 | 4810.936702 | 0.15302766 | 0.002529975 |
| Dpysl5 | 4174.656992 | 0.152929746 | 0.030038631 |
| Ncoa5 | 2185.846261 | 0.152755328 | 0.010566642 |
| Hdlbp | 9906.942421 | 0.152480637 | 0.007018818 |
| Rnf187 | 12793.54993 | 0.152476707 | 0.001061432 |
| Mrpl37 | 1577.775669 | 0.152126427 | 0.022814445 |
| Mecp2 | 1473.476539 | 0.150667813 | 0.01480513 |
| Ube3a | 4843.795747 | 0.150599202 | 0.01374973 |
| Dlc1 | 2671.341426 | 0.150282168 | 0.012424718 |
| Tspyl1 | 6603.495551 | 0.150259503 | 0.000595057 |
| Usp20 | 3514.615464 | 0.1499195 | 0.017578624 |
| Ylpm1 | 8087.629389 | 0.149696039 | 0.019635839 |
| Myo9a | 5901.738636 | 0.149642897 | 0.019990009 |
| Rnf10 | 7219.449344 | 0.149630647 | 0.000237874 |
| Nup155 | 1073.18146 | 0.149453559 | 0.041273672 |
| Pja1 | 9136.067423 | 0.149401963 | 0.012248151 |
| Ubp1 | 1027.415369 | 0.149400951 | 0.028503172 |
| Dcun1d4 | 2917.67393 | 0.149232594 | 0.030437422 |
| Hagh | 1585.533275 | 0.148616169 | 0.006088536 |
| Ndfip1 | 11516.13772 | 0.148380042 | 0.002628368 |
| Mib2 | 4036.549965 | 0.148211944 | 0.006753348 |
| Camk2g | 3269.940157 | 0.148163131 | 0.049992114 |

| | | | |
|---|---|---|---|
| Sin3b | 1979.746093 | 0.147700556 | 0.001648985 |
| Sars | 4475.456889 | 0.147232156 | 0.012266372 |
| Rtf1 | 3029.465921 | 0.14695812 | 0.009386977 |
| Btbd6 | 1390.024296 | 0.146846149 | 0.017418257 |
| Mpp2 | 3166.483052 | 0.146487991 | 0.022161681 |
| Kif1b | 38718.64451 | 0.146131984 | 0.003736508 |
| Coro1c | 3682.639183 | 0.145997805 | 0.011885047 |
| Asnsd1 | 1228.439057 | 0.145809628 | 0.036227797 |
| Ehmt2 | 5190.245042 | 0.14528017 | 0.010083973 |
| Tsr2 | 3982.112658 | 0.144874872 | 0.011013158 |
| LOC499339 | 1156.556808 | 0.144792613 | 0.015292635 |
| Dnajc11 | 1805.335415 | 0.144608658 | 0.00255033 |
| Osbpl2 | 2315.444501 | 0.144512341 | 0.009612677 |
| Cul3 | 6060.266571 | 0.14437536 | 0.001624094 |
| Ubap2 | 2478.087563 | 0.144176235 | 0.011592397 |
| Impa1 | 1851.797134 | 0.143895436 | 0.012914729 |
| Med15 | 1876.41981 | 0.143875648 | 0.049638157 |
| Tmem57 | 2642.705532 | 0.143643041 | 0.019136951 |
| Bcorl1 | 1739.853585 | 0.143188002 | 0.048979886 |
| Cox6a1 | 9109.737082 | 0.142905204 | 0.013085346 |
| Ash1l | 12466.02759 | 0.142824183 | 0.029813689 |
| Gltscr1l | 1910.548889 | 0.142373207 | 0.012080043 |
| Rabggtb | 1710.757943 | 0.142297002 | 0.024387541 |
| Trappc13 | 1216.30312 | 0.142272606 | 0.027029014 |
| Csnk1g2 | 2187.388404 | 0.141992192 | 0.044661744 |
| Ankrd50 | 2091.644209 | 0.14196759 | 0.045897385 |
| Ggnbp2 | 2605.605745 | 0.141807927 | 0.005841193 |
| Smcr8 | 1550.560578 | 0.141774155 | 0.031326729 |
| Prrc2c | 13878.21044 | 0.141648417 | 0.028703089 |
| Bcl7b | 1544.829189 | 0.141645272 | 0.011974822 |
| Nckipsd | 2005.403461 | 0.141224027 | 0.03829135 |
| Pitpna | 7523.881919 | 0.141216317 | 0.00423773 |
| Pnma2 | 2949.60188 | 0.140710704 | 0.0266191 |
| Aqr | 2141.869857 | 0.140491601 | 0.020967204 |
| Usp7 | 5768.962025 | 0.140468218 | 0.000448644 |
| Zfp532 | 1956.998039 | 0.140462885 | 0.001199925 |

| | | | |
|---|---|---|---|
| Strn3 | 2712.587289 | 0.140280424 | 0.012586839 |
| Safb | 4954.509862 | 0.140135137 | 0.010293887 |
| Eif4b | 8188.153914 | 0.139304037 | 0.00026199 |
| Gpm6a | 27503.73915 | 0.138914098 | 0.038954746 |
| Taok2 | 4290.235812 | 0.138687309 | 0.007473644 |
| Dis3l2 | 1092.433185 | 0.138641007 | 0.048302914 |
| Klhdc2 | 3133.980555 | 0.138511102 | 0.007238781 |
| Marf1 | 4599.858988 | 0.138427632 | 0.011896706 |
| Vmp1 | 2018.532062 | 0.138133565 | 0.024698136 |
| Aldoa | 49859.36452 | 0.137993753 | 0.007777568 |
| Atp6ap1 | 9182.496168 | 0.137736633 | 0.002203946 |
| Gdpd1 | 836.8784747 | 0.137715683 | 0.042134473 |
| Mrps9 | 1054.483567 | 0.137444453 | 0.047807061 |
| Yme1l1 | 3089.309995 | 0.137431173 | 0.019903157 |
| Tmem55b | 1721.099393 | 0.137381865 | 0.029140072 |
| Med24 | 2105.563261 | 0.137243215 | 0.007018818 |
| Hcfc1r1 | 2898.741453 | 0.137237706 | 0.020251314 |
| Aarsd1 | 2085.014203 | 0.136245554 | 0.002581764 |
| Cog2 | 1117.541186 | 0.136242573 | 0.035297035 |
| Tnks2 | 7741.36416 | 0.136104481 | 0.017830482 |
| Cldn12 | 1630.685849 | 0.135931826 | 0.03057947 |
| Fam20c | 2631.503316 | 0.135876098 | 0.038954746 |
| Ap1b1 | 5771.888009 | 0.135447574 | 0.030563962 |
| Rrp1 | 1216.549502 | 0.134788289 | 0.048190413 |
| Sirt5 | 2268.809843 | 0.134315889 | 0.009356894 |
| Tomm40 | 1419.92749 | 0.134241222 | 0.02965463 |
| St3gal2 | 2223.460512 | 0.133945709 | 0.014696249 |
| Wdtc1 | 2528.732495 | 0.133658965 | 0.012331186 |
| Ptpn4 | 5660.49562 | 0.133584933 | 0.038265829 |
| Scoc | 3353.201344 | 0.133256064 | 0.021015044 |
| Mboat7 | 5445.80329 | 0.133107321 | 0.037039996 |
| Ing3 | 663.850165 | 0.133050805 | 0.026374392 |
| Rbm18 | 1695.195281 | 0.131749597 | 0.041507997 |
| Chmp7 | 3378.170458 | 0.131732856 | 0.011885047 |
| Dnaja2 | 5404.429226 | 0.131501167 | 0.000554569 |
| Hsp90ab1 | 59638.81647 | 0.131444437 | 0.005651074 |

| | | | |
|---|---|---|---|
| Kansl2 | 1873.054487 | 0.130769189 | 0.009495248 |
| Ranbp2 | 9132.786903 | 0.130065946 | 0.020519229 |
| Kpna3 | 2146.856242 | 0.129741656 | 0.014326581 |
| Gramd1b | 3019.234825 | 0.129322717 | 0.04597216 |
| Wrnip1 | 1662.397003 | 0.129316898 | 0.042185843 |
| Clock | 3192.576058 | 0.12909673 | 0.04113225 |
| RGD1308601 | 1564.13294 | 0.128032217 | 0.025055493 |
| Fam8a1 | 3324.620291 | 0.127427378 | 0.033439949 |
| Rab2a | 2424.557785 | 0.127397704 | 0.022814445 |
| Gatad2b | 3212.55738 | 0.127312007 | 0.015201896 |
| Stub1 | 4831.386671 | 0.12723256 | 0.022090765 |
| Pam | 10120.08279 | 0.127177596 | 0.03212032 |
| Gsk3b | 4354.292804 | 0.127150512 | 0.00124056 |
| Golga3 | 3199.273816 | 0.12639573 | 0.02194135 |
| Agap1 | 4974.836378 | 0.126196858 | 0.029140072 |
| Ubr1 | 3342.946977 | 0.125864252 | 0.013374456 |
| Ddx24 | 5556.771001 | 0.125651435 | 0.022338058 |
| Ppp2r1a | 16668.29828 | 0.125630412 | 0.001897506 |
| Trip12 | 7977.097466 | 0.124394896 | 0.013865872 |
| Cpsf2 | 1952.51518 | 0.124370477 | 0.04597216 |
| Supt6h | 10075.27863 | 0.123573252 | 0.024233422 |
| Hmgb3 | 1457.594922 | 0.123028734 | 0.042926805 |
| Ngfrap1 | 1380.239804 | 0.123019376 | 0.034666977 |
| Fxr2 | 4271.891297 | 0.122416476 | 0.010785378 |
| Rogdi | 5013.94234 | 0.12232878 | 0.004499548 |
| Myl12b | 7493.585723 | 0.121886564 | 0.01739845 |
| RGD1305938 | 1859.000875 | 0.121831835 | 0.044712624 |
| Ube2z | 4002.751865 | 0.121675089 | 0.020740663 |
| Pum2 | 7155.055534 | 0.121562347 | 0.012634703 |
| Ube2s | 1401.19653 | 0.121277901 | 0.019823737 |
| Scaf1 | 8603.547654 | 0.121172203 | 0.039601016 |
| Bsg | 14284.68965 | 0.121108723 | 0.020431381 |
| Rabl6 | 6030.84249 | 0.121029957 | 0.00420574 |
| Akr1a1 | 6422.395649 | 0.120457153 | 0.001865521 |
| Celf1 | 1834.905733 | 0.120030209 | 0.018653255 |
| Abr | 20310.39821 | 0.118847346 | 0.008987589 |

| | | | |
|---|---|---|---|
| Kpna1 | 1941.201749 | 0.11877397 | 0.041081416 |
| Gnl1 | 3046.199508 | 0.1185384 | 0.03431286 |
| Eif4a1 | 6950.793901 | 0.118483218 | 0.039617536 |
| Vdac1 | 14241.91244 | 0.118443461 | 0.01445176 |
| Nckap1 | 20231.46071 | 0.118237416 | 0.011554723 |
| Dlat | 3688.909025 | 0.11777056 | 0.043602297 |
| Atp6v0d1 | 10648.81347 | 0.117668657 | 0.021824132 |
| Ypel5 | 3281.782964 | 0.117646014 | 0.017261564 |
| Pex6 | 2145.63128 | 0.117253283 | 0.047379645 |
| Smc1a | 4963.271685 | 0.117129285 | 0.028199717 |
| Dnajc7 | 3453.701317 | 0.117064095 | 0.01877848 |
| Pspc1 | 1577.495916 | 0.116547674 | 0.018720564 |
| Rnf111 | 2164.474281 | 0.116097368 | 0.024571822 |
| Rxrb | 1567.469546 | 0.115296384 | 0.010480008 |
| Acin1 | 4234.834675 | 0.114581731 | 0.040542479 |
| Cbfa2t2 | 1163.04409 | 0.113976606 | 0.035691833 |
| Ubqln2 | 13857.60675 | 0.113897844 | 0.006176954 |
| Ctbp1 | 6743.922988 | 0.113787499 | 0.00564783 |
| Morf4l2 | 5504.920446 | 0.113604483 | 0.037940202 |
| Rabgef1 | 917.8181487 | 0.113519812 | 0.045988168 |
| Itfg1 | 9848.398279 | 0.112849084 | 0.004088175 |
| Clta | 5531.040282 | 0.111857148 | 0.04320649 |
| Nicn1 | 2624.542898 | 0.111832256 | 0.017953844 |
| Akap10 | 1184.222014 | 0.111433754 | 0.0204231 |
| Pcyox1 | 6203.820068 | 0.110833281 | 0.035831811 |
| Ube2m | 3476.135635 | 0.110674335 | 0.042844524 |
| Psmc3 | 4406.604149 | 0.110643642 | 0.022814445 |
| Sec16a | 5353.334429 | 0.110600462 | 0.039142856 |
| Eif1b | 2331.211763 | 0.110088037 | 0.029081623 |
| Ccar2 | 4288.169253 | 0.109759556 | 0.009061877 |
| Oaz1 | 6898.682974 | 0.109548366 | 0.029315736 |
| Supt16h | 4149.672763 | 0.109008022 | 0.036588153 |
| Dnm1l | 4748.608055 | 0.108720603 | 0.035448946 |
| Ikbkap | 3472.978411 | 0.108311467 | 0.047458949 |
| Dhx30 | 5049.981288 | 0.108264622 | 0.031030807 |
| Cltc | 42214.55306 | 0.107811673 | 0.021528689 |

| | | | |
|---|---|---|---|
| Sae1 | 2105.357821 | 0.107147352 | 0.046200201 |
| Ralbp1 | 2979.687874 | 0.106885563 | 0.022098947 |
| Rad23b | 10796.16704 | 0.105830635 | 0.013330302 |
| Vcpip1 | 4403.706114 | 0.105613891 | 0.008851239 |
| Trap1 | 2765.831551 | 0.105155398 | 0.034951661 |
| Map7d1 | 12837.15845 | 0.105141635 | 0.046562614 |
| Setd5 | 5256.132284 | 0.105079222 | 0.037991562 |
| Cmip | 9119.532746 | 0.104337667 | 0.040067873 |
| Mgea5 | 4404.087139 | 0.104194552 | 0.046438468 |
| Lnpep | 4014.774695 | 0.103999676 | 0.049542627 |
| Ociad1 | 4661.780291 | 0.10346371 | 0.020468398 |
| Nolc1 | 6455.09137 | 0.103289289 | 0.004668556 |
| Wbp11 | 3467.493049 | 0.102809868 | 0.030421623 |
| Gnb1 | 11078.43036 | 0.102488715 | 0.046876486 |
| Ankrd17 | 10209.87864 | 0.102472881 | 0.016923864 |
| Zfp148 | 3621.998721 | 0.102305501 | 0.020811156 |
| Gnao1 | 20031.21751 | 0.100687196 | 0.021674811 |
| Plekhb2 | 13104.00931 | 0.100060283 | 0.048460537 |
| Hspa4 | 11047.19096 | 0.099828196 | 0.009936102 |
| Dpysl2 | 49379.89693 | 0.099669254 | 0.027772022 |
| Nrd1 | 4724.915226 | 0.099397245 | 0.022352602 |
| Qrich1 | 3124.744019 | 0.098630203 | 0.021817139 |
| Slc25a44 | 2491.140813 | 0.098306468 | 0.046533954 |
| Arid1a | 10943.28182 | 0.098141166 | 0.039578437 |
| Agpat1 | 6268.130865 | 0.097646412 | 0.042917775 |
| LOC294154 | 4307.923169 | 0.097245464 | 0.04600367 |
| Cops7a | 4384.610783 | 0.096513037 | 0.041352147 |
| Akap17a | 2616.96547 | 0.095193781 | 0.033901581 |
| Cct8 | 5562.80722 | 0.094548278 | 0.03431286 |
| Tollip | 4999.053045 | 0.094295107 | 0.023515462 |
| Cab39 | 6312.722949 | 0.094094459 | 0.037261255 |
| Psmc5 | 4237.458743 | 0.09396872 | 0.039713537 |
| LOC680039 | 5909.053742 | 0.093285427 | 0.047172927 |
| Rhot1 | 2109.432353 | 0.091874205 | 0.039447253 |
| Scrn1 | 5557.34746 | 0.090921512 | 0.041151961 |
| Supt5h | 8184.582867 | 0.089473778 | 0.023832172 |

| | | | |
|---|---|---|---|
| Ppp2ca | 9765.52275 | 0.082649268 | 0.028545163 |
| Prkci | 3984.649274 | 0.068129384 | 0.04367931 |
| Hnrnpk | 12010.42735 | -0.078006448 | 0.037442833 |
| Tm9sf4 | 2485.48995 | -0.081646742 | 0.036129737 |
| Usp47 | 6110.291095 | -0.08207655 | 0.03889408 |
| Atp5f1 | 8099.951377 | -0.083320297 | 0.034185779 |
| Rab5b | 9461.851611 | -0.084960069 | 0.009588529 |
| Eif4g2 | 34846.72451 | -0.086230744 | 0.023273372 |
| Cd2bp2 | 4324.596903 | -0.088045284 | 0.028262453 |
| Mtmr3 | 4744.381382 | -0.088564768 | 0.021296418 |
| Lonp1 | 4489.733879 | -0.089321783 | 0.040712055 |
| Gtf2i | 9612.125716 | -0.093435021 | 0.04113225 |
| Tmed2 | 4437.308886 | -0.096520366 | 0.044496656 |
| Arl6ip1 | 7315.735627 | -0.096521538 | 0.030343028 |
| Gdi2 | 8845.511182 | -0.097342227 | 0.024344919 |
| Yipf5 | 1702.824202 | -0.097664861 | 0.042798976 |
| Trim35 | 11480.34267 | -0.09877019 | 0.045271423 |
| Bsdc1 | 3840.718266 | -0.100169503 | 0.044884491 |
| Dbnl | 2735.219948 | -0.101017187 | 0.040278052 |
| Ik | 4357.725247 | -0.102531388 | 0.022275482 |
| Khsrp | 3837.455707 | -0.102764892 | 0.045563563 |
| Dgcr2 | 2976.592623 | -0.103622045 | 0.041572552 |
| Api5 | 3731.428172 | -0.103731793 | 0.043391181 |
| Tprg1l | 7635.108023 | -0.103797443 | 0.049293769 |
| M6pr | 3899.043308 | -0.104986885 | 0.032680079 |
| Ccpg1 | 3331.545586 | -0.105493169 | 0.037175181 |
| Pa2g4 | 2645.212301 | -0.105583438 | 0.047807061 |
| Copz1 | 2685.301362 | -0.107776538 | 0.032320803 |
| Nbr1 | 10260.6274 | -0.107828221 | 0.002458007 |
| Chst10 | 2756.963359 | -0.108693932 | 0.015762421 |
| Raf1 | 2353.941038 | -0.10918516 | 0.022629852 |
| Pcif1 | 1306.856711 | -0.110082407 | 0.034958215 |
| Rbbp4 | 3337.29621 | -0.11014662 | 0.047560323 |
| Ncstn | 2754.439211 | -0.110663013 | 0.015015996 |
| Ndufa12 | 1564.108621 | -0.111327146 | 0.04113225 |
| Ndufs7 | 3393.323787 | -0.111564188 | 0.032074311 |

| | | | |
|---|---|---|---|
| Lars | 3366.677647 | -0.112813171 | 0.026807275 |
| Tbc1d5 | 2188.675236 | -0.113641355 | 0.028261868 |
| Zhx1 | 5502.964396 | -0.113666357 | 0.046844801 |
| Ehd1 | 2660.8997 | -0.114737887 | 0.02711779 |
| Caskin1 | 14895.88242 | -0.115137802 | 0.038961336 |
| Spag9 | 24037.29765 | -0.115653235 | 0.038054617 |
| Ankib1 | 3041.927848 | -0.115904176 | 0.022814445 |
| Stoml2 | 1402.158952 | -0.11636525 | 0.046526272 |
| Raly | 2722.458663 | -0.116443819 | 0.029865442 |
| Gpr107 | 2689.099536 | -0.117287129 | 0.022696413 |
| Klhl7 | 3601.06881 | -0.117531712 | 0.041476297 |
| Srprb | 1145.623895 | -0.118735711 | 0.044884491 |
| Btf3 | 2735.090827 | -0.118755031 | 0.039536825 |
| Ssr2 | 1241.593984 | -0.118959713 | 0.040736401 |
| Phc2 | 2304.37908 | -0.119090256 | 0.035528714 |
| Ptpra | 10685.01938 | -0.119774759 | 0.000940172 |
| Abl1 | 3154.320764 | -0.119880501 | 0.04414195 |
| Tbc1d9b | 7094.753411 | -0.120169462 | 0.015292635 |
| Tmem33 | 3218.382865 | -0.120306787 | 0.03826335 |
| Smc3 | 4303.300809 | -0.120581958 | 0.0466356 |
| Fam134c | 1441.850204 | -0.121272741 | 0.037613641 |
| Pdxdc1 | 1452.781232 | -0.121616899 | 0.037065247 |
| Arsb | 5030.335731 | -0.121656118 | 0.033312481 |
| Chmp5 | 3864.100007 | -0.121828826 | 0.032178039 |
| Fam120a | 11293.07282 | -0.122336406 | 0.0194991 |
| Mvb12b | 2685.018558 | -0.122587717 | 0.026202364 |
| Adcy2 | 9863.466857 | -0.123082665 | 0.048483173 |
| Dad1 | 2424.519684 | -0.123720435 | 0.04456294 |
| Rbmxrtl | 1603.184155 | -0.123931058 | 0.043557067 |
| Eif5 | 11586.8702 | -0.124037378 | 0.041226845 |
| Polr2e | 2012.53979 | -0.125109846 | 0.03293963 |
| Manbal | 1854.32451 | -0.125142385 | 0.036877741 |
| Mmgt1 | 1922.445671 | -0.125170667 | 0.035909843 |
| Snx27 | 8033.324035 | -0.125680343 | 0.014092697 |
| Plbd2 | 2500.63248 | -0.125799562 | 0.016935195 |
| Ube2l3 | 5923.499315 | -0.126448785 | 0.001951763 |

| | | | |
|---|---|---|---|
| Sdcbp | 7931.291043 | -0.126601204 | 0.039420784 |
| Gosr2 | 2136.853551 | -0.126889862 | 0.00610728 |
| Ptdss2 | 3216.441129 | -0.127554898 | 0.005514161 |
| Sod1 | 7068.421078 | -0.127945874 | 0.018414334 |
| Kdelr1 | 1994.652111 | -0.128600316 | 0.026875324 |
| Oxa1l | 2092.666723 | -0.128910701 | 0.047303885 |
| Emc7 | 1665.026802 | -0.129729333 | 0.035784906 |
| Sar1b | 1562.827549 | -0.130778739 | 0.029315736 |
| Smpd1 | 4373.383655 | -0.130904324 | 0.031753084 |
| Acp2 | 2088.94024 | -0.131553 | 0.010149865 |
| Sec22b | 1647.261937 | -0.131692589 | 0.024150179 |
| Rab7a | 13666.79389 | -0.131991479 | 0.00205016 |
| Hmgb1 | 4371.824571 | -0.132633808 | 0.036786456 |
| Cpsf3 | 1392.177816 | -0.133062554 | 0.036877741 |
| Rnf115 | 997.7499035 | -0.133728731 | 0.030044814 |
| Pdcd11 | 1628.126694 | -0.133884554 | 0.018586785 |
| Rpl18 | 2604.42889 | -0.133948295 | 0.026526406 |
| Emc2 | 2336.03553 | -0.133959783 | 0.008599641 |
| Tab1 | 1452.966795 | -0.134258023 | 0.048101698 |
| Smu1 | 2074.948885 | -0.134266807 | 0.043328117 |
| Crk | 1787.148256 | -0.13516705 | 0.044496656 |
| St13 | 5728.259665 | -0.135432993 | 0.004333928 |
| Fam172a | 1832.008348 | -0.135658174 | 0.039276863 |
| Zfp91 | 5852.234145 | -0.135961889 | 0.00310529 |
| Hnrnpul1 | 7470.351514 | -0.136337185 | 0.004333928 |
| Casc4 | 8931.777081 | -0.136368624 | 0.006706917 |
| Hm13 | 1978.362189 | -0.136918388 | 0.009995523 |
| Pigq | 2394.747651 | -0.138209896 | 0.00797097 |
| Uqcrfs1 | 4609.161229 | -0.138576478 | 0.014384357 |
| Oxsr1 | 1748.384859 | -0.138609472 | 0.041575325 |
| Pitpnb | 1842.879479 | -0.138977196 | 0.023807337 |
| Pole3 | 1773.148861 | -0.139257235 | 0.027364332 |
| Wdr1 | 10084.15111 | -0.13992315 | 0.024823012 |
| Rcbtb1 | 1623.855869 | -0.1399748 | 0.024791848 |
| Tbrg1 | 849.333059 | -0.140015846 | 0.046123203 |
| Flot2 | 4646.83651 | -0.140115399 | 0.009994799 |

| | | | |
|---|---|---|---|
| Cisd2 | 2552.831806 | -0.140364456 | 0.012424718 |
| Eef1d | 2078.516492 | -0.140597447 | 0.027419387 |
| Psmd9 | 664.0163589 | -0.141230315 | 0.031583562 |
| Ssr1 | 4049.628105 | -0.141323474 | 0.00200133 |
| Tex264 | 2004.399405 | -0.141376416 | 0.003759605 |
| Cnpy3 | 2600.621771 | -0.141576817 | 0.008081341 |
| Atad1 | 4365.139189 | -0.141949019 | 0.000402658 |
| Phf2 | 3109.957368 | -0.142007143 | 0.030220464 |
| Ifnar1 | 3016.082334 | -0.142091258 | 0.006285872 |
| Pigt | 4359.089396 | -0.142161079 | 0.007865177 |
| Ctnnbl1 | 1456.130477 | -0.1426707 | 0.04901362 |
| RGD1303003 | 3908.541893 | -0.142996097 | 0.039468079 |
| Wbp1l | 2397.554511 | -0.143114703 | 0.029954087 |
| Stat5b | 2004.548765 | -0.143629847 | 0.037272121 |
| Eprs | 5380.980864 | -0.143926038 | 0.003747201 |
| Mib1 | 11499.47389 | -0.144119542 | 0.035152458 |
| RGD1310553 | 1665.10224 | -0.144510079 | 0.016923864 |
| Wrb | 2003.680572 | -0.144531626 | 0.010447814 |
| Phf10 | 873.5729653 | -0.145099614 | 0.036327113 |
| Trim8 | 8856.763663 | -0.145245993 | 0.018373197 |
| Tmem55a | 2409.61614 | -0.145323671 | 0.027470381 |
| Dirc2 | 3587.273348 | -0.145656397 | 0.000923027 |
| Fam19a5 | 3726.335477 | -0.14570133 | 0.029733867 |
| Coro1b | 2757.820388 | -0.145873101 | 0.036864065 |
| Zc3h7b | 11382.47468 | -0.146361939 | 0.002672174 |
| Ganab | 7579.240951 | -0.146614473 | 0.00959119 |
| RGD1310352 | 2911.252436 | -0.14680475 | 0.005567047 |
| Ptp4a2 | 1235.133662 | -0.146920448 | 0.014133421 |
| Paics | 2951.862003 | -0.147230223 | 0.014362647 |
| Arf4 | 2990.743579 | -0.147306123 | 0.001673071 |
| Gbas | 1669.862128 | -0.147700354 | 0.018534551 |
| Tmub1 | 888.1966526 | -0.14774847 | 0.036134134 |
| Adh5 | 1677.879898 | -0.148416908 | 0.026126138 |
| Stt3b | 3600.678967 | -0.149148255 | 0.003525287 |
| Lasp1 | 7102.775094 | -0.149275675 | 0.008491113 |
| RGD1311783 | 1729.320552 | -0.149553537 | 0.033888079 |

| | | | |
|---|---|---|---|
| Gtf3c2 | 1407.074268 | -0.149922326 | 0.021957416 |
| Canx | 32425.47224 | -0.150050076 | 0.00152972 |
| Lgmn | 2257.310629 | -0.150330898 | 0.016159296 |
| Pgls | 847.6207109 | -0.15069918 | 0.02854608 |
| Psmf1 | 1199.818488 | -0.15078275 | 0.048588401 |
| Drap1 | 2459.544009 | -0.151218636 | 0.008710887 |
| Tmem59 | 6485.144277 | -0.152177398 | 0.001114274 |
| Ddx39b | 3788.398802 | -0.152314516 | 0.002644861 |
| Serf2 | 2704.796458 | -0.152440513 | 0.037850918 |
| Rrm1 | 1350.023504 | -0.1530354 | 0.039236585 |
| Gclc | 3660.984161 | -0.153877741 | 0.033289315 |
| Ctsb | 22477.57714 | -0.153908848 | 0.004955639 |
| Emc3 | 4034.541676 | -0.155007228 | 0.027364332 |
| Ext2 | 1408.495163 | -0.155048608 | 0.040115943 |
| Chchd3 | 2471.474 | -0.155108851 | 0.008070171 |
| Hacd3 | 10066.50412 | -0.155219995 | 0.011483501 |
| Ebna1bp2 | 1415.833285 | -0.155382042 | 0.020433225 |
| Gtf2b | 711.6312622 | -0.155947075 | 0.032074311 |
| Pdk2 | 7609.377124 | -0.156521016 | 0.015664807 |
| Hmgn2 | 2383.668077 | -0.156559707 | 0.031308575 |
| Sdccag3 | 1083.527663 | -0.156775602 | 0.02073198 |
| Slc25a39 | 1357.775595 | -0.157222752 | 0.017196199 |
| Serp1 | 2877.103827 | -0.157334272 | 0.013479162 |
| LOC361635 | 2036.085242 | -0.15779248 | 0.026379536 |
| Sec63 | 6036.761606 | -0.158411569 | 0.038461703 |
| Slc25a17 | 2343.51861 | -0.158488941 | 0.005286915 |
| Lsamp | 5575.800103 | -0.158587017 | 0.038548495 |
| Alkbh5 | 1914.866363 | -0.158981255 | 0.01471667 |
| Chst11 | 1563.221221 | -0.159180616 | 0.039204078 |
| Cdh2 | 12596.35484 | -0.159344307 | 0.005887146 |
| Setdb1 | 1772.050993 | -0.159736608 | 0.006001839 |
| Bpgm | 3092.336942 | -0.15984698 | 0.001865075 |
| Tmco3 | 1700.3468 | -0.159880756 | 0.018953696 |
| Hsp90b1 | 17180.00054 | -0.159972762 | 0.011876738 |
| Por | 8905.657928 | -0.15998744 | 0.046728685 |
| RGD1562136 | 2236.427718 | -0.160102062 | 0.002670189 |

| | | | |
|---|---|---|---|
| Erp44 | 957.0134374 | -0.160317368 | 0.019166144 |
| Hars2 | 1283.481873 | -0.160375949 | 0.023287604 |
| Dvl3 | 2147.936063 | -0.160524638 | 0.009043704 |
| Cadm4 | 7016.37041 | -0.1606983 | 0.000949009 |
| Adcy6 | 2365.03044 | -0.161140711 | 0.013589627 |
| Hiatl1 | 1832.552656 | -0.161159096 | 0.007279215 |
| Tle3 | 1989.568944 | -0.161570081 | 0.039526414 |
| Mrpl4 | 1447.711952 | -0.161675411 | 0.00183866 |
| Armc5 | 1185.627703 | -0.161793145 | 0.027269423 |
| Osbpl9 | 1698.157875 | -0.162194311 | 0.046103442 |
| Txndc5 | 2303.059434 | -0.162389284 | 0.008225891 |
| RGD1562987 | 2229.565573 | -0.162731804 | 0.000615768 |
| Tax1bp1 | 10599.75467 | -0.162830275 | 0.010357265 |
| Aurkaip1 | 851.5447201 | -0.163148009 | 0.044739293 |
| Pigu | 618.3680304 | -0.163598424 | 0.035152458 |
| Lats1 | 3891.647872 | -0.163631809 | 0.00937931 |
| Ndst1 | 4286.743053 | -0.163804652 | 0.004419211 |
| Lzts2 | 1505.450275 | -0.163820493 | 0.041266106 |
| Cmtr1 | 4235.230765 | -0.163997279 | 0.000547016 |
| Cnih4 | 551.2633545 | -0.164088814 | 0.034718813 |
| Yae1d1 | 1008.02652 | -0.16422553 | 0.017022057 |
| Mtfr1l | 1544.412448 | -0.164663487 | 0.007099134 |
| Prpf18 | 3627.35846 | -0.164749791 | 0.011555382 |
| Prpf40a | 1957.574131 | -0.164884766 | 0.015952109 |
| Sbds | 3480.152277 | -0.164896945 | 0.023124567 |
| Mlec | 3052.840682 | -0.165075838 | 0.012317827 |
| Mpnd | 1353.589045 | -0.165261123 | 0.031442156 |
| Slc35e4 | 1749.634853 | -0.165625725 | 0.044107401 |
| Xpo4 | 1039.349682 | -0.165644619 | 0.045262958 |
| Pogk | 1537.666316 | -0.165787258 | 0.031519977 |
| Nlgn1 | 2010.660494 | -0.165925829 | 0.035008103 |
| Amn1 | 467.2535433 | -0.166103716 | 0.049933859 |
| Pigz | 1564.417881 | -0.166731988 | 0.009179195 |
| Cldnd1 | 2964.617527 | -0.166996903 | 0.006989664 |
| Ccs | 1037.016313 | -0.167083369 | 0.017830482 |
| Rad21 | 6178.956092 | -0.167426059 | 0.001525361 |

| | | | |
|---|---|---|---|
| Csgalnact1 | 1522.561058 | -0.167888904 | 0.044664311 |
| Gid8 | 1330.002805 | -0.167954732 | 0.028199717 |
| Pmm1 | 5771.303837 | -0.168094439 | 0.016815765 |
| Ints4 | 1861.210099 | -0.168179512 | 0.019808899 |
| Ptcd1 | 643.4331346 | -0.168190545 | 0.031398302 |
| Sema6d | 7764.982939 | -0.168193003 | 0.040678354 |
| Ncam1 | 21677.24882 | -0.168703946 | 0.003609139 |
| Mrpl3 | 1987.680373 | -0.168906848 | 0.000396691 |
| Pcyt1a | 1411.642021 | -0.168999189 | 0.043331589 |
| Prnp | 27881.83945 | -0.169611793 | 0.009480013 |
| Ckb | 73526.81195 | -0.169684384 | 0.033623613 |
| Ppfia1 | 2458.890941 | -0.169763981 | 0.013573688 |
| Smarce1 | 2450.075383 | -0.169833846 | 0.000986792 |
| Pcdhga7 | 3532.508129 | -0.169936127 | 0.018658576 |
| Timmdc1 | 1308.345545 | -0.170365777 | 0.026723797 |
| Tfg | 1859.637731 | -0.170649848 | 0.007334104 |
| Bloc1s6 | 567.0050648 | -0.170656398 | 0.04342208 |
| Extl3 | 4109.937676 | -0.171344671 | 7.76E-05 |
| Oat | 5896.253515 | -0.171509938 | 0.011379239 |
| Diablo | 824.7959255 | -0.17153272 | 0.026055773 |
| Metap2 | 2337.848134 | -0.171546637 | 0.000953118 |
| Mta2 | 1645.165845 | -0.171758671 | 0.018943723 |
| Zdhhc20 | 711.049251 | -0.171847807 | 0.044884491 |
| Sptlc1 | 1604.675755 | -0.171916006 | 0.00385804 |
| Tcea1 | 2320.346457 | -0.171998106 | 0.001687318 |
| Limch1 | 5210.061682 | -0.172790952 | 0.009480013 |
| Degs1 | 2817.556753 | -0.173228264 | 0.003836416 |
| Vegfb | 1405.127416 | -0.173432851 | 0.020625632 |
| Ap3b1 | 701.5128299 | -0.173928766 | 0.044436976 |
| Fbxo30 | 928.8704644 | -0.17394374 | 0.036675908 |
| Agpat3 | 16230.14201 | -0.173994888 | 0.004847825 |
| Mapk1ip1l | 1242.386669 | -0.174154608 | 0.004693495 |
| Zfp422 | 881.5319874 | -0.174194422 | 0.033029838 |
| Rnf114 | 1343.724968 | -0.174392385 | 0.023887598 |
| Gca | 1313.034436 | -0.174411833 | 0.037324102 |
| Pank3 | 8725.688883 | -0.174499589 | 0.033366286 |

| | | | |
|---|---|---|---|
| RGD1310127 | 1414.025494 | -0.174630991 | 0.004261302 |
| Mre11a | 626.0456292 | -0.175008259 | 0.046793877 |
| Tfam | 862.4499659 | -0.175032093 | 0.010045575 |
| Galnt18 | 1317.720593 | -0.175193807 | 0.043003276 |
| Nudt3 | 13295.6923 | -0.175339504 | 0.02119585 |
| Nr3c1 | 6145.585807 | -0.175477959 | 0.006137228 |
| Mboat2 | 2053.837205 | -0.175904603 | 0.04997837 |
| Pacsin3 | 1940.119432 | -0.175947825 | 0.045729828 |
| Map2k2 | 1925.774068 | -0.175983727 | 0.005185706 |
| Paip1 | 2876.441139 | -0.176224996 | 0.001049857 |
| Sc5d | 5443.460376 | -0.176225309 | 0.008591609 |
| Rcan1 | 1715.821376 | -0.176334665 | 0.010718937 |
| Asah1 | 1854.939334 | -0.176471901 | 0.00453125 |
| Spcs2 | 3005.40749 | -0.176607636 | 0.003367331 |
| Surf4 | 3077.593935 | -0.176768586 | 0.003029645 |
| Sema4d | 1855.744514 | -0.176773439 | 0.007562792 |
| Cdk5rap3 | 1562.1419 | -0.176875936 | 0.023370403 |
| Me2 | 1037.667303 | -0.176895011 | 0.036786456 |
| Lrrfip2 | 655.9832888 | -0.17711918 | 0.043376548 |
| Dpysl3 | 1739.206898 | -0.17726485 | 0.029788128 |
| Repin1 | 604.3351377 | -0.177320601 | 0.038023716 |
| Poc5 | 452.9816672 | -0.177530798 | 0.031781318 |
| Dpy19l3 | 10560.54212 | -0.177705563 | 0.008960495 |
| Snrpc | 792.0597886 | -0.178113895 | 0.018586785 |
| Thyn1 | 1218.98568 | -0.178479197 | 0.019780651 |
| Sec24a | 2047.831528 | -0.178958417 | 0.004790477 |
| Lamtor4 | 1071.906817 | -0.179153027 | 0.04367931 |
| Nbn | 1188.96882 | -0.179164413 | 0.019794127 |
| Txndc15 | 3299.553769 | -0.179706817 | 3.72E-05 |
| Gfm1 | 2052.11559 | -0.180443363 | 0.032972478 |
| Gns | 2090.551533 | -0.180599137 | 0.004158527 |
| Ccdc22 | 719.3744542 | -0.181172231 | 0.027470381 |
| Ppp4r3b | 3250.958397 | -0.181288338 | 0.000914809 |
| Abcd3 | 4240.03019 | -0.181874433 | 0.000994931 |
| Man2c1 | 1391.158766 | -0.181883403 | 0.010569014 |
| Mdm2 | 1065.389936 | -0.182047008 | 0.04408643 |

| | | | |
|---|---|---|---|
| Rcn2 | 8367.096623 | -0.182090633 | 0.01293307 |
| Zcchc2 | 759.9910041 | -0.182134274 | 0.015722854 |
| Bcap31 | 2083.406913 | -0.182503071 | 0.00062425 |
| Anapc1 | 3423.758022 | -0.182555854 | 0.000628987 |
| Ilk | 2709.353624 | -0.182747881 | 0.000648389 |
| Selt | 5513.83359 | -0.182766143 | 0.005074428 |
| Slc39a7 | 2992.375801 | -0.18286322 | 0.006021508 |
| Cad | 1428.398412 | -0.183755066 | 0.025136791 |
| RGD1562218 | 1248.363035 | -0.183916315 | 0.017536673 |
| Mettl9 | 1578.121993 | -0.184681381 | 0.003118582 |
| Fbxo3 | 3590.83357 | -0.184706636 | 0.006676478 |
| Gtf2a1 | 1320.204105 | -0.184735281 | 0.006085849 |
| Ppp2r5a | 3418.010306 | -0.185218215 | 0.006755238 |
| Mphosph6 | 670.1261061 | -0.185488648 | 0.023244979 |
| Otulin | 1632.00685 | -0.185721388 | 0.015786916 |
| Gabarap | 3631.992663 | -0.18612253 | 0.000493137 |
| Etfb | 2125.940739 | -0.186492448 | 0.024778181 |
| Pir | 898.6893593 | -0.187217275 | 0.037822119 |
| Bckdk | 1485.856185 | -0.18726026 | 0.010149865 |
| Bbx | 1329.36198 | -0.187325367 | 0.025305059 |
| Stx4 | 590.4591716 | -0.187787035 | 0.015248909 |
| Aff1 | 1083.191677 | -0.187889976 | 0.016990993 |
| Trafd1 | 960.1383445 | -0.188952813 | 0.035013629 |
| Rexo4 | 2317.680883 | -0.189168638 | 0.000824341 |
| Lrrc58 | 2718.130245 | -0.189817132 | 0.027591598 |
| Adk | 3575.893816 | -0.189845866 | 0.025136791 |
| Hipk1 | 11464.41911 | -0.190049605 | 0.000910537 |
| Tm9sf1 | 640.572645 | -0.190060906 | 0.017245163 |
| Crip2 | 4766.578766 | -0.190791755 | 0.010592764 |
| Elk4 | 1586.241263 | -0.191054134 | 0.00431713 |
| Fyn | 3307.107672 | -0.191843003 | 0.003293512 |
| Fer | 648.3964967 | -0.192185883 | 0.025310298 |
| Pigk | 1924.197995 | -0.192199328 | 0.004981837 |
| Rhoa | 3104.523084 | -0.192417336 | 0.003850644 |
| Adam9 | 3102.077746 | -0.192923736 | 0.009664435 |
| Coasy | 1048.966328 | -0.193218626 | 0.001013191 |

| | | | |
|---|---|---|---|
| Ctnnd2 | 24719.05627 | -0.193256333 | 0.001806583 |
| Zbtb1 | 524.7639245 | -0.193397555 | 0.01827172 |
| Ap5m1 | 544.6610468 | -0.193403637 | 0.044838641 |
| Rab18 | 5260.943467 | -0.193556862 | 6.22E-05 |
| Gfra1 | 2618.944387 | -0.193582508 | 0.048422609 |
| Sec61a2 | 2521.575269 | -0.194150991 | 0.000525706 |
| Banf1 | 1496.740084 | -0.194265513 | 0.008118041 |
| Otud7b | 2174.972774 | -0.194387874 | 0.035831811 |
| RGD1311847 | 426.2725292 | -0.194791722 | 0.044679068 |
| Ankrd40 | 5654.286264 | -0.1954732 | 0.000596313 |
| Dnajc3 | 1651.154728 | -0.195483843 | 0.00938133 |
| Map3k1 | 561.636624 | -0.195524301 | 0.047184927 |
| Ssr3 | 4833.597363 | -0.195627346 | 0.002471472 |
| Txndc12 | 1075.86282 | -0.195806328 | 0.003552161 |
| Snx14 | 1254.073991 | -0.196221162 | 0.013155632 |
| Gpr75 | 687.9381391 | -0.196391184 | 0.039526414 |
| Rbl2 | 2665.518008 | -0.196604077 | 0.008629629 |
| Eif4ebp2 | 1052.42357 | -0.197036763 | 0.003672475 |
| Slitrk2 | 2188.502951 | -0.197094351 | 0.026223186 |
| Ccdc91 | 1520.856523 | -0.1971166 | 0.006120222 |
| Tmem218 | 663.0015163 | -0.197213376 | 0.027178755 |
| Abcd1 | 692.3068878 | -0.197765543 | 0.012558645 |
| Fchsd2 | 2697.132866 | -0.197988538 | 0.001321258 |
| Desi2 | 1104.238298 | -0.198088431 | 0.021388376 |
| Src | 2685.276903 | -0.198236939 | 0.003092332 |
| Gtf2e1 | 1159.414543 | -0.198273027 | 0.030057613 |
| Pik3ip1 | 1126.463336 | -0.198761319 | 0.049408312 |
| Kif5b | 7663.032684 | -0.198775357 | 0.002053329 |
| Nr1h2 | 1344.427361 | -0.198982805 | 0.011885047 |
| Kpna2 | 1234.362412 | -0.199129424 | 0.025015386 |
| Calr | 20361.88224 | -0.199325602 | 0.002062182 |
| Rnd2 | 2637.512776 | -0.199346901 | 0.031066082 |
| Fbxo8 | 648.4938073 | -0.19953111 | 0.014084793 |
| Fam50a | 897.8277891 | -0.199944305 | 0.007326108 |
| Sox4 | 1301.097899 | -0.200003546 | 0.04102509 |
| Aph1b | 546.6413349 | -0.200089169 | 0.037723974 |

| | | | |
|---|---|---|---|
| Snapin | 2036.160645 | -0.200500171 | 0.005074428 |
| Mrpl27 | 890.7759169 | -0.200688004 | 0.007147386 |
| Samd5 | 1137.544202 | -0.200704971 | 0.027973962 |
| Apeh | 1499.550745 | -0.200925108 | 0.005365667 |
| Efcab14 | 3109.064379 | -0.20092559 | 0.000840224 |
| Nubp2 | 579.7542802 | -0.201236602 | 0.043481662 |
| Tmem104 | 564.2021782 | -0.201382898 | 0.043184844 |
| Rxra | 923.8575307 | -0.201582033 | 0.025159782 |
| Nlrx1 | 525.1424019 | -0.201672075 | 0.028148272 |
| Fundc1 | 924.3770363 | -0.201752802 | 0.005639331 |
| Cdo1 | 2133.914551 | -0.201845781 | 0.03269278 |
| Plekhm2 | 3578.231061 | -0.201863362 | 0.000701546 |
| Sec11a | 505.9829282 | -0.202108618 | 0.031077233 |
| Aadat | 1615.945701 | -0.202141422 | 0.023704809 |
| Saraf | 6765.348366 | -0.202212853 | 0.00874504 |
| Mxd4 | 758.6219396 | -0.202860437 | 0.029737113 |
| Ckap4 | 1057.272909 | -0.202879601 | 0.016840496 |
| Smurf2 | 2721.474219 | -0.202907328 | 0.004320218 |
| Nudt12 | 737.7960326 | -0.203207842 | 0.032680079 |
| Dpf2 | 2270.598879 | -0.203274305 | 0.002899504 |
| Prrc1 | 510.9558378 | -0.203675902 | 0.025693797 |
| Ncoa4 | 4429.948081 | -0.203757501 | 0.001477849 |
| Taf13 | 1976.017736 | -0.204225518 | 0.045397692 |
| Arl13b | 451.4827759 | -0.20459732 | 0.042642888 |
| Thumpd3 | 1043.980866 | -0.204757001 | 0.000840224 |
| Pou3f3 | 4308.469824 | -0.204776535 | 0.022098947 |
| Kifc3 | 2684.728624 | -0.205548192 | 0.011495122 |
| Dhx32 | 1580.158656 | -0.205716315 | 0.004186493 |
| Pc | 5790.061857 | -0.205806313 | 0.004824343 |
| Zfp192 | 1046.261735 | -0.205810755 | 0.019929563 |
| Ints12 | 273.5973435 | -0.206253629 | 0.047932298 |
| Anks1a | 2315.968742 | -0.206523539 | 0.022188676 |
| Cecr5 | 456.3049594 | -0.206755765 | 0.032744227 |
| Parn | 415.3681427 | -0.206898039 | 0.043019063 |
| Siae | 907.2636583 | -0.207128227 | 0.016268643 |
| Farp1 | 4265.608658 | -0.207177778 | 0.024056503 |

| | | | |
|---|---|---|---|
| Pfas | 1823.557768 | -0.207437762 | 0.016318827 |
| Lhfp | 1856.213273 | -0.208264941 | 0.002283391 |
| Atp11a | 3029.321118 | -0.208431475 | 0.004266779 |
| Smarcc1 | 3613.142156 | -0.208578587 | 0.000546569 |
| Znfx1 | 2381.918519 | -0.209227285 | 0.01576116 |
| Asf1a | 767.4365453 | -0.20949774 | 0.040202328 |
| Wnk1 | 15243.07977 | -0.209516923 | 0.000463817 |
| Atic | 2827.565773 | -0.20957355 | 0.006026271 |
| Stard7 | 1527.396398 | -0.209679281 | 0.003901574 |
| Trove2 | 3074.592573 | -0.209933932 | 0.001963548 |
| Cd59 | 3526.155946 | -0.210212212 | 0.012080043 |
| Phkb | 3376.142191 | -0.210278655 | 0.007812363 |
| Snx9 | 520.7883726 | -0.210385718 | 0.034320272 |
| Mrpl40 | 888.5480528 | -0.210555081 | 0.008113052 |
| Akr7a2 | 1052.580848 | -0.210710111 | 0.002847374 |
| Foxn3 | 2516.445418 | -0.210736149 | 0.012836853 |
| Ttll12 | 864.3661566 | -0.211086605 | 0.00910708 |
| Lig3 | 776.1041936 | -0.211125623 | 0.004976223 |
| Rras2 | 606.4352286 | -0.211260543 | 0.036569268 |
| Rfx5 | 1634.991382 | -0.211540076 | 0.02342589 |
| Ccdc12 | 767.4174312 | -0.211728997 | 0.007465382 |
| Rab10 | 8226.692972 | -0.212097169 | 0.000157789 |
| Ddt | 2936.373952 | -0.212242037 | 0.001609267 |
| Taf15 | 1308.652707 | -0.212881157 | 0.008245717 |
| Cbfb | 676.4504712 | -0.212892094 | 0.029733867 |
| Pgd | 1654.864918 | -0.213245464 | 0.001609267 |
| Itga6 | 5588.780548 | -0.213265255 | 0.009329349 |
| Comt | 1540.132458 | -0.213351305 | 0.000206311 |
| Ccdc93 | 825.6243492 | -0.213528506 | 0.021481122 |
| Vkorc1 | 297.1967517 | -0.213577977 | 0.034070947 |
| RGD1564541 | 1484.03635 | -0.213918927 | 0.002522882 |
| Mavs | 776.2638514 | -0.214034491 | 0.007815464 |
| Fgd6 | 1543.153647 | -0.214094178 | 0.048355987 |
| Grm3 | 15312.47247 | -0.214275036 | 0.014260868 |
| Rufy1 | 1240.335093 | -0.214411398 | 0.007147386 |
| Elmod2 | 656.9997166 | -0.214786804 | 0.037787265 |

| | | | |
|---|---|---|---|
| Rpe | 796.1686798 | -0.214893338 | 0.007522331 |
| Spire1 | 12279.67031 | -0.214992328 | 0.00281999 |
| Bbs1 | 2409.167273 | -0.21538375 | 0.004579824 |
| Als2 | 2218.389636 | -0.21546912 | 0.001857613 |
| Mlycd | 1392.163686 | -0.215688755 | 0.00778778 |
| Lat2 | 2.127161336 | -0.215758664 | 0.045397692 |
| Pik3c2a | 3477.557295 | -0.215829694 | 0.000498757 |
| Kdelr2 | 1219.80704 | -0.215983797 | 0.000686858 |
| Sppl2b | 688.2366554 | -0.216400824 | 0.018854913 |
| Ccnc | 1311.644423 | -0.216539461 | 0.001987385 |
| Hadha | 6537.574348 | -0.216642324 | 0.003589118 |
| Tgfbr1 | 609.2701475 | -0.21672362 | 0.029095706 |
| Galc | 592.9243333 | -0.216960614 | 0.042409248 |
| Anp32e | 2086.418832 | -0.21700144 | 0.010054737 |
| Slc48a1 | 5591.874872 | -0.217069135 | 0.003250649 |
| Pink1 | 5988.247927 | -0.217141337 | 0.000985354 |
| Spon1 | 6114.663241 | -0.217144795 | 0.020267766 |
| RGD1565685 | 371.6010924 | -0.217368049 | 0.041081416 |
| Timm17b | 424.7991131 | -0.217438706 | 0.029733867 |
| Afap1 | 1238.393074 | -0.217531463 | 0.04413695 |
| Wdr35l | 2765.889079 | -0.217577891 | 0.000187767 |
| Mipep | 1602.170631 | -0.217805592 | 0.01927369 |
| Stam2 | 973.9114624 | -0.217852408 | 0.000838748 |
| Rab8a | 1903.555497 | -0.217907973 | 0.000145887 |
| Tom1 | 1768.095014 | -0.218002409 | 0.005841193 |
| Hspa5 | 19670.66355 | -0.218191969 | 0.028769237 |
| Tmem189 | 1436.194321 | -0.21822856 | 0.020060668 |
| Crkl | 2853.664785 | -0.218261288 | 0.000508867 |
| Gdf11 | 3456.918161 | -0.218809105 | 0.016069956 |
| Sgcb | 6830.693206 | -0.218982169 | 0.000204097 |
| Rab9a | 1139.297788 | -0.219426012 | 0.012711726 |
| Mpv17l | 230.9947112 | -0.219499262 | 0.040816319 |
| Sowahc | 503.5007474 | -0.219571502 | 0.031979674 |
| Tmbim6 | 5573.220199 | -0.219605926 | 0.000225486 |
| Sncaip | 783.1538823 | -0.219618831 | 0.022991808 |
| Ptar1 | 1236.529685 | -0.219626222 | 0.024399304 |

| | | | |
|---|---|---|---|
| Snx18 | 1508.89609 | -0.219765092 | 0.00997349 |
| Gmpr2 | 423.3859664 | -0.220530247 | 0.025408163 |
| Cachd1 | 1995.766102 | -0.220555579 | 0.025874937 |
| Stag2 | 4340.536963 | -0.22063064 | 0.001825611 |
| Tmem180 | 643.6530133 | -0.220897892 | 0.013074117 |
| Flii | 4626.584312 | -0.221128247 | 0.000552441 |
| Tmem186 | 721.5852897 | -0.22198083 | 0.002309063 |
| Sfr1 | 3033.552326 | -0.222195117 | 0.003230384 |
| H2afy | 3129.676434 | -0.222294696 | 0.000301594 |
| Hexim2 | 410.3051779 | -0.222733526 | 0.013615035 |
| Zeb1 | 5660.042829 | -0.222736433 | 0.002651621 |
| Tnfaip1 | 3266.790167 | -0.222779333 | 0.000464956 |
| Rnf180 | 1803.209186 | -0.223154639 | 0.004517356 |
| Sec14l2 | 3300.680214 | -0.223190841 | 0.047379645 |
| Tor1aip1 | 2073.174668 | -0.22321546 | 0.003449692 |
| Galm | 997.074287 | -0.223246097 | 0.011060847 |
| Mesdc2 | 3827.677095 | -0.22331635 | 0.000187533 |
| Kdelc2 | 925.9502719 | -0.223477795 | 0.004170474 |
| Pnpt1 | 814.3807024 | -0.223560191 | 0.007401927 |
| Pcca | 1186.902506 | -0.224174902 | 0.022814445 |
| Cgrrf1 | 531.8145403 | -0.224614947 | 0.005128574 |
| RGD1562865 | 376.6814635 | -0.224776823 | 0.044466614 |
| Acot13 | 663.9730629 | -0.22479526 | 0.009193977 |
| Plekhf2 | 1300.543419 | -0.22483075 | 0.016680097 |
| Anxa5 | 4185.059511 | -0.22533943 | 0.011046521 |
| Tpst2 | 1058.38647 | -0.22544902 | 0.015762421 |
| Hdac8 | 762.0937003 | -0.225499507 | 0.018786958 |
| Fzd9 | 1309.28206 | -0.225607164 | 0.03600114 |
| Sgce | 857.9579659 | -0.225737958 | 0.010786879 |
| Fjx1 | 11567.62663 | -0.22596937 | 0.025902587 |
| Acox1 | 3952.498882 | -0.226135158 | 0.021479639 |
| Ppm1f | 1618.756279 | -0.226243657 | 0.014942816 |
| Dnajb2 | 2337.43398 | -0.22647296 | 0.001410214 |
| Sypl1 | 718.7058629 | -0.226924504 | 0.012522344 |
| Lactb | 701.6816016 | -0.227334214 | 0.023228201 |
| Adi1 | 1090.73666 | -0.227398415 | 0.037241065 |

| | | | |
|---|---|---|---|
| Rbpj | 2097.937924 | -0.227552622 | 0.00377394 |
| Tcp11l2 | 716.9599879 | -0.227728784 | 0.010033766 |
| G6pd | 3286.158546 | -0.227746312 | 0.000255 |
| Pdia6 | 4645.057925 | -0.228003254 | 0.000406438 |
| Cadm1 | 7559.023374 | -0.228122113 | 0.013085346 |
| Cotl1 | 2899.676496 | -0.228149646 | 0.002396593 |
| Nfkb1 | 950.8675601 | -0.228180914 | 0.002480053 |
| Zfp703 | 3770.230301 | -0.228188289 | 0.006004671 |
| LOC306766 | 3888.754214 | -0.228212368 | 2.85E-05 |
| Elf2 | 951.8918329 | -0.228303689 | 0.005955317 |
| Arsa | 1087.38061 | -0.228333566 | 0.013023457 |
| Zfp516 | 1077.315515 | -0.228492439 | 0.010401317 |
| Efnb1 | 635.9826914 | -0.228588269 | 0.009531657 |
| Tkfc | 292.1795504 | -0.228722489 | 0.04301741 |
| Gstz1 | 1201.455538 | -0.228736522 | 0.02197366 |
| Polr2d | 360.0822951 | -0.229387222 | 0.010040959 |
| Ddx19a | 1577.368922 | -0.229459749 | 0.005490502 |
| P4ha1 | 2743.714661 | -0.229619462 | 0.003850644 |
| Dfna5 | 489.5573499 | -0.229815438 | 0.035036358 |
| Npc1 | 3094.80739 | -0.229838199 | 0.013596789 |
| Eftud1 | 732.403976 | -0.230580381 | 0.007260114 |
| Haus7 | 248.4565375 | -0.230600046 | 0.024075743 |
| Slc35f5 | 559.5096111 | -0.230731723 | 0.024336128 |
| Ctns | 469.294074 | -0.230809068 | 0.03948544 |
| Plxdc2 | 2077.360368 | -0.230876412 | 0.01271823 |
| Ntrk2 | 100591.7334 | -0.230894241 | 0.004083159 |
| Tmem256 | 322.0077223 | -0.231253086 | 0.049653482 |
| Edem2 | 562.9973523 | -0.231446144 | 0.013596029 |
| Nek4 | 530.5498306 | -0.231474036 | 0.019622887 |
| Ggcx | 714.7302314 | -0.231986739 | 0.047932298 |
| Zfp652 | 1211.403286 | -0.232084365 | 0.007427309 |
| Pepd | 1710.697225 | -0.232439745 | 0.00959119 |
| Aprt | 334.3649053 | -0.232585004 | 0.020827167 |
| Gabpa | 1631.128607 | -0.232645247 | 0.012688321 |
| Cpped1 | 548.7104047 | -0.232810436 | 0.007777568 |
| Prps2 | 503.1744557 | -0.232922811 | 0.011911137 |

| | | | |
|---|---|---|---|
| Afg3l1 | 922.2890548 | -0.232947628 | 0.00985844 |
| Cd164 | 4497.791494 | -0.233021334 | 0.001663181 |
| Dhx40 | 1840.050803 | -0.23338798 | 0.000668553 |
| Krt32 | 1.592790269 | -0.23341886 | 0.03590352 |
| Ak3 | 1294.360838 | -0.233482142 | 0.003374293 |
| Gtf2h5 | 936.4876324 | -0.233915121 | 0.001198913 |
| Tceal8 | 1773.374485 | -0.235151131 | 0.000986789 |
| Aldh3a2 | 2698.701386 | -0.235183381 | 0.001166624 |
| Pcna | 730.1514497 | -0.235273769 | 0.002170434 |
| Nln | 1827.248646 | -0.235349716 | 0.000163847 |
| Ubxn4 | 4237.095375 | -0.235475595 | 4.95E-05 |
| Egfr | 2247.436002 | -0.23562115 | 0.011998927 |
| Chsy1 | 1468.343441 | -0.235852197 | 0.002100895 |
| Apln | 4240.071365 | -0.23609697 | 0.022219191 |
| Tgfb2 | 3374.652264 | -0.236231187 | 0.008453936 |
| Pdia3 | 9583.435325 | -0.236240669 | 1.00E-04 |
| Igfbp7 | 4491.984156 | -0.236327058 | 0.01704767 |
| Vat1 | 2351.037107 | -0.237081445 | 0.049901099 |
| Igsf11 | 6455.087989 | -0.23708267 | 0.005590012 |
| Tspan3 | 15260.59565 | -0.237287042 | 2.67E-06 |
| Pkn2 | 2878.467676 | -0.237425004 | 0.00139305 |
| Syde1 | 358.5467622 | -0.237620443 | 0.039536825 |
| Gyg1 | 1133.348361 | -0.237745766 | 0.003942336 |
| Wdr91 | 1811.532089 | -0.237807007 | 0.003860256 |
| Wwox | 364.380612 | -0.237831777 | 0.027822389 |
| Ssh3 | 387.4889202 | -0.237867946 | 0.016401421 |
| RGD1309534 | 1302.547964 | -0.238003688 | 0.00978304 |
| Mcl1 | 850.2391292 | -0.238042265 | 0.030957322 |
| Xrcc5 | 930.5309444 | -0.238518899 | 0.006089266 |
| Tor1aip2 | 2825.521028 | -0.238534388 | 5.29E-05 |
| Mrpl45 | 1667.238043 | -0.238543446 | 0.00125523 |
| Stx17 | 288.6458202 | -0.23911482 | 0.036682471 |
| Tcaim | 507.8590532 | -0.239395107 | 0.021368156 |
| Pkd2 | 1546.300932 | -0.239838011 | 0.004772804 |
| Nfix | 4220.97308 | -0.239857129 | 0.003602802 |
| Prkx | 385.263653 | -0.240091637 | 0.013669571 |

| | | | |
|---|---|---|---|
| Ccdc28b | 440.3459636 | -0.240275023 | 0.04746648 |
| Tab2 | 3288.707847 | -0.240509425 | 0.024959966 |
| Ppib | 1848.775574 | -0.240521797 | 0.000809649 |
| Cd320 | 431.2456404 | -0.240686583 | 0.033027925 |
| Epas1 | 12351.099 | -0.240844452 | 0.038996381 |
| Ift27 | 527.8434248 | -0.240933863 | 0.034743329 |
| Eda | 487.3927987 | -0.2410509 | 0.040433381 |
| Adprhl2 | 1584.571898 | -0.241164912 | 0.000744575 |
| Prg4 | 2.279858487 | -0.241213024 | 0.042137113 |
| Rbms2 | 245.9307368 | -0.241474161 | 0.044679068 |
| Stag1 | 1565.554671 | -0.24168105 | 0.004470584 |
| Pigh | 476.769582 | -0.241806366 | 0.004433611 |
| Golim4 | 1453.730986 | -0.241815294 | 0.008210233 |
| Nasp | 384.1294077 | -0.241845459 | 0.04743897 |
| Man2b1 | 861.2561735 | -0.242093347 | 0.009135145 |
| Nampt | 1690.641717 | -0.242095762 | 0.000992958 |
| LOC689574 | 778.589819 | -0.24217657 | 0.015588367 |
| C1qtnf5 | 2506.25789 | -0.242178018 | 0.012883416 |
| Txnrd2 | 432.0407193 | -0.242307552 | 0.020646145 |
| Med12 | 1683.753165 | -0.242365403 | 0.001611675 |
| LOC100910558 | 808.9791022 | -0.24285856 | 0.037347212 |
| Sts | 965.5772227 | -0.242996906 | 0.013423843 |
| Il33 | 1718.265174 | -0.243213441 | 0.043482609 |
| Raver2 | 1836.299985 | -0.243516917 | 0.024150179 |
| Cnppd1 | 899.5092783 | -0.243582478 | 0.003121979 |
| Tmlhe | 314.5492155 | -0.243774843 | 0.039571108 |
| Hsd17b11 | 1604.237741 | -0.244151689 | 0.016935195 |
| RGD1311345 | 922.2823295 | -0.244307304 | 0.003809049 |
| Slc25a20 | 1557.665218 | -0.244403037 | 0.0008745 |
| Hacl1 | 580.1218202 | -0.244650695 | 0.01573839 |
| Tmem9b | 1136.598099 | -0.244653401 | 0.00052861 |
| Tsc22d4 | 4186.952456 | -0.244966381 | 0.008070844 |
| Ubr7 | 2509.744331 | -0.24497661 | 0.008629629 |
| Paqr8 | 8477.357447 | -0.24510002 | 0.0436009 |
| Ctbs | 907.3501419 | -0.245447686 | 0.011855808 |
| Rhoq | 1656.420518 | -0.24547726 | 0.011689283 |

| | | | |
|---|---|---|---|
| Arhgef12 | 12388.01455 | -0.245496308 | 0.00115063 |
| Dbt | 2131.386445 | -0.245740761 | 0.011079136 |
| Ddah1 | 6562.746892 | -0.246085853 | 0.000154675 |
| Msi1 | 1472.405197 | -0.246181271 | 0.024977981 |
| Evi5 | 6438.931763 | -0.246281106 | 0.000254846 |
| Tdp1 | 176.2829386 | -0.24650666 | 0.03494165 |
| Hsd17b10 | 1704.44615 | -0.246660685 | 0.003925806 |
| Commd8 | 1874.035114 | -0.247231504 | 0.001027841 |
| Akap13 | 1237.153552 | -0.247528503 | 0.004400085 |
| Prkab1 | 304.0400012 | -0.247541581 | 0.034745834 |
| Dstn | 4150.461338 | -0.247568164 | 6.76E-05 |
| Ermp1 | 3502.083429 | -0.24771787 | 0.00114526 |
| Svil | 557.0310874 | -0.248264954 | 0.046278844 |
| Mfge8 | 22687.48429 | -0.24836863 | 0.0220393 |
| Shisa5 | 1240.302774 | -0.248412915 | 0.003406683 |
| Frmd4a | 1483.502864 | -0.2485942 | 0.00139912 |
| Pcbp4 | 4344.817269 | -0.248665305 | 3.32E-05 |
| Nedd9 | 1360.225489 | -0.249134926 | 0.010171222 |
| Rasa3 | 2911.141844 | -0.249219726 | 0.000382551 |
| Papss1 | 3703.700135 | -0.249413225 | 0.000464229 |
| Sft2d1 | 425.093424 | -0.249434686 | 0.021762019 |
| Bmpr1b | 1278.735338 | -0.24979949 | 0.016812998 |
| Dazap2 | 7028.290914 | -0.250070061 | 0.000586359 |
| Hip1 | 2762.791326 | -0.250142684 | 0.027822389 |
| Pex2 | 520.7229536 | -0.25018706 | 0.008026708 |
| Slc3a2 | 11742.95562 | -0.250339832 | 0.001471985 |
| Sall2 | 5730.84432 | -0.250343206 | 0.005047001 |
| Odc1 | 2774.941562 | -0.250570851 | 0.000182259 |
| Nr1d1 | 6469.944276 | -0.250743767 | 0.038879276 |
| Smim15 | 1106.944431 | -0.251030775 | 0.000501795 |
| Smox | 2452.237285 | -0.251111665 | 0.021066122 |
| Cflar | 667.8078667 | -0.251161652 | 0.010506342 |
| S100a11 | 1.532609178 | -0.251328058 | 0.044697817 |
| Tex261 | 590.0947977 | -0.251399489 | 0.005309755 |
| Agpat5 | 2461.314586 | -0.251806604 | 0.00124056 |
| Atg10 | 192.2226399 | -0.252394779 | 0.028147469 |

| | | | |
|---|---|---|---|
| Usp6nl | 1831.919085 | -0.252821526 | 0.004292494 |
| Tmed7 | 4197.94066 | -0.252895214 | 8.65E-06 |
| Zadh2 | 728.0923266 | -0.253073463 | 0.013726493 |
| Ppp1r14c | 425.498202 | -0.253197468 | 0.031512209 |
| Rpn2 | 5319.950002 | -0.253411846 | 0.00011505 |
| Chka | 973.298529 | -0.253465449 | 0.006166648 |
| Pamr1 | 1375.441996 | -0.253751736 | 0.036992401 |
| Wwc2 | 2920.168382 | -0.253933716 | 4.94E-05 |
| Aida | 1804.874926 | -0.254223081 | 0.000310766 |
| Acp6 | 398.1789702 | -0.25440867 | 0.005639331 |
| Hnrnpf | 5628.82102 | -0.254542417 | 0.00018714 |
| Ano6 | 449.3431526 | -0.255203766 | 0.011445678 |
| Comtd1 | 928.6899844 | -0.255268642 | 0.015349985 |
| Snta1 | 2433.22374 | -0.255393758 | 0.003787107 |
| RGD1359158 | 382.1188391 | -0.255436527 | 0.010228755 |
| Cat | 5162.036936 | -0.255663939 | 0.003720329 |
| Lsm6 | 549.6482018 | -0.25607364 | 0.0096283 |
| Rab5c | 7771.99537 | -0.256102215 | 1.48E-05 |
| Cdk2ap2 | 554.8099743 | -0.25614466 | 0.014133421 |
| RGD1359634 | 462.8544497 | -0.256150073 | 0.038224629 |
| Fgf2 | 873.5788175 | -0.256415989 | 0.017741295 |
| Ugp2 | 3870.044361 | -0.256549326 | 0.000113781 |
| Gnpda2 | 694.2663143 | -0.257230435 | 0.011218166 |
| Pex7 | 351.2139015 | -0.257319028 | 0.030907759 |
| Acadsb | 774.9690337 | -0.257598807 | 0.008909046 |
| Npr2 | 1446.797466 | -0.257709428 | 0.004158527 |
| Lzic | 718.4871488 | -0.257916831 | 0.000411388 |
| Wars2 | 185.65985 | -0.258338227 | 0.038148756 |
| Rap1a | 1368.419439 | -0.258490058 | 0.000310661 |
| RGD1565616 | 8762.288061 | -0.258542122 | 0.012082906 |
| Poglut1 | 1061.009542 | -0.25864899 | 0.007186984 |
| Myd88 | 419.2927046 | -0.258661853 | 0.034145908 |
| Sptssa | 803.6263522 | -0.258780672 | 0.009780825 |
| Cdyl | 519.9557407 | -0.259161179 | 0.020444539 |
| Coq9 | 1733.525522 | -0.259161577 | 0.003798409 |
| Cpq | 2126.936952 | -0.259166415 | 0.007622412 |

| | | | |
|---|---|---|---|
| Ctsz | 740.1815012 | -0.259514453 | 0.005285706 |
| Tmem38b | 443.4340126 | -0.259552048 | 0.004732147 |
| Hadh | 1846.911824 | -0.259585609 | 0.009434246 |
| Arhgap5 | 13637.60361 | -0.260125844 | 0.000224724 |
| RGD1566052 | 413.3756952 | -0.260317948 | 0.024303465 |
| Nop9 | 356.8538944 | -0.26046425 | 0.002359125 |
| Aga | 600.0344842 | -0.260960893 | 0.004442836 |
| Creg1 | 1861.567745 | -0.26098564 | 0.006673009 |
| Hey1 | 686.1204532 | -0.2610099 | 0.009936102 |
| Hexb | 2079.298198 | -0.261214043 | 0.00124056 |
| Casp3 | 1145.606211 | -0.261341151 | 0.007334104 |
| Wdr60 | 676.8204352 | -0.261422217 | 0.010055277 |
| Dpagt1 | 862.3499205 | -0.261438861 | 0.001675481 |
| Abhd1 | 591.7760325 | -0.261984821 | 0.022798006 |
| Mfsd1 | 1849.026298 | -0.262025018 | 0.000154758 |
| Eif2b1 | 611.3391225 | -0.26217016 | 0.003725603 |
| Arhgap12 | 3136.609985 | -0.262175276 | 0.001010509 |
| Mospd3 | 578.3309758 | -0.262442868 | 0.005138945 |
| Xylb | 395.780358 | -0.26276818 | 0.014513152 |
| Skap2 | 340.6236722 | -0.262852088 | 0.041823163 |
| Twf1 | 2391.549493 | -0.262893444 | 0.000522468 |
| Orai1 | 359.6333166 | -0.262949506 | 0.022098947 |
| Sesn3 | 11367.26694 | -0.263207917 | 0.000195713 |
| Rab21 | 4418.974738 | -0.263253479 | 4.89E-05 |
| Dennd5a | 20085.25863 | -0.263326897 | 1.03E-06 |
| Spg20 | 2104.050382 | -0.263404937 | 3.17E-05 |
| Apba3 | 426.7246137 | -0.263443041 | 0.027627193 |
| Nsmaf | 365.631663 | -0.263518359 | 0.008065625 |
| Ankrd13a | 1913.215596 | -0.263608083 | 0.001083534 |
| Vimp | 1411.182123 | -0.263622962 | 0.000508867 |
| Lix1l | 2567.455616 | -0.264162873 | 0.001310359 |
| Hsd17b12 | 3280.793845 | -0.264500607 | 0.001145431 |
| Tfrc | 3415.346994 | -0.264898587 | 0.044712624 |
| Wasf2 | 340.2015925 | -0.264933656 | 0.024225986 |
| Gulp1 | 378.749139 | -0.265295979 | 0.027853232 |
| Bet1 | 804.9482846 | -0.265404946 | 0.003510205 |

| | | | |
|---|---|---|---|
| Xiap | 1342.702557 | -0.265708568 | 0.000121478 |
| Mapkapk2 | 804.8396813 | -0.265774702 | 0.002585171 |
| Nnt | 4717.329683 | -0.265838525 | 0.002209887 |
| Cdk2ap1 | 1077.054654 | -0.2660044 | 0.000858273 |
| Endog | 497.887552 | -0.266133108 | 0.018030055 |
| Abat | 11052.55879 | -0.266440049 | 0.011007771 |
| Tead3 | 246.5013039 | -0.266783385 | 0.044712624 |
| Mfsd10 | 376.1427368 | -0.267086133 | 0.044343468 |
| Lrrc48 | 487.5160636 | -0.267274389 | 0.022161681 |
| Tspan14 | 479.6866279 | -0.267784512 | 0.020646145 |
| Fbxw8 | 1758.085792 | -0.267815012 | 0.00099841 |
| Pex10 | 394.5488017 | -0.267895365 | 0.002072079 |
| Sbf2 | 3136.858658 | -0.268176463 | 0.000331874 |
| Oxnad1 | 536.7472551 | -0.268437162 | 0.008507574 |
| Ccdc90b | 628.8626393 | -0.268639172 | 0.000750972 |
| H2afv | 875.2476889 | -0.269196373 | 0.004711663 |
| Wdr53 | 160.4944837 | -0.269309756 | 0.013731436 |
| Hspb11 | 212.1985244 | -0.269507598 | 0.030166364 |
| Mad2l2 | 138.3065816 | -0.269531342 | 0.046325694 |
| Zdhhc4 | 209.4880056 | -0.269553911 | 0.044145704 |
| Sec61a1 | 3283.305405 | -0.269677241 | 0.000419804 |
| Lxn | 1621.676283 | -0.269825608 | 0.009913523 |
| G6pc3 | 1251.622588 | -0.269870141 | 0.003925831 |
| Decr1 | 1193.448046 | -0.270031465 | 0.015952109 |
| Dync2li1 | 535.5217558 | -0.270725228 | 0.011657243 |
| Ccng1 | 10133.45714 | -0.270738794 | 0.005533694 |
| Cyp4f4 | 947.7918771 | -0.270808206 | 0.047083583 |
| Sdhc | 5251.041388 | -0.270835627 | 2.49E-05 |
| Tspan2 | 1214.395667 | -0.270939828 | 0.033251434 |
| Klf3 | 1768.570103 | -0.271021508 | 0.002717851 |
| Glmp | 1077.03151 | -0.271259446 | 0.002044671 |
| Ahcyl1 | 20251.53033 | -0.271862624 | 0.008240881 |
| Ergic3 | 2955.145905 | -0.272025898 | 6.37E-06 |
| Nudcd2 | 732.855236 | -0.272201938 | 0.00387717 |
| Cep112 | 193.5800009 | -0.272688303 | 0.021525858 |
| Dusp15 | 909.9039557 | -0.272768053 | 0.001500778 |

| | | | |
|---|---|---|---|
| Anapc10 | 306.2002177 | -0.272814017 | 0.001581032 |
| Swap70 | 2775.493586 | -0.272991801 | 0.015256368 |
| Nr2e1 | 1215.349436 | -0.2732181 | 0.004416048 |
| Oas1f | 1.907228701 | -0.273389038 | 0.026189836 |
| Cxcl14 | 7703.976533 | -0.273397112 | 0.007873894 |
| Stk40 | 1167.745526 | -0.273812933 | 0.012839377 |
| Zfp608 | 1777.653986 | -0.273863225 | 0.004948263 |
| Aldoc | 96558.38122 | -0.27389329 | 0.010224245 |
| Prkra | 772.7194159 | -0.273992333 | 0.006292511 |
| Bckdhb | 1328.931443 | -0.274179138 | 0.001049749 |
| Ccbl2 | 274.4387298 | -0.274867264 | 0.015029172 |
| Adhfe1 | 3572.146163 | -0.274969263 | 0.020483051 |
| P4hb | 7149.465165 | -0.275026094 | 4.29E-05 |
| Tyw1 | 1044.925187 | -0.275697043 | 0.001020166 |
| Spg21 | 836.7152782 | -0.275698873 | 0.000294885 |
| Hbp1 | 776.4400481 | -0.275766504 | 0.006839097 |
| Tdrd7 | 2177.842434 | -0.275874228 | 0.00118017 |
| Dhrs3 | 972.2021322 | -0.275989171 | 0.020776778 |
| Arid5a | 532.6984949 | -0.276084085 | 0.011440402 |
| Iqgap1 | 496.791951 | -0.276101991 | 0.005467138 |
| Muc15 | 2.227228144 | -0.276318548 | 0.020089853 |
| Tmem134 | 217.0840562 | -0.276374357 | 0.031908682 |
| Macrod1 | 353.801335 | -0.276427582 | 0.009684717 |
| Emc8 | 900.9811039 | -0.276467421 | 0.004112849 |
| Mdm1 | 403.6634717 | -0.276745182 | 0.024965612 |
| Mageh1 | 1241.128286 | -0.276762076 | 0.000259237 |
| Mt3 | 12892.89868 | -0.276768204 | 0.000193354 |
| Atp6v0a2 | 1235.719311 | -0.276940632 | 0.000186891 |
| Ldhd | 284.9397197 | -0.277104527 | 0.022933556 |
| Dpp7 | 1722.592323 | -0.277261412 | 4.14E-05 |
| P3h1 | 288.5649924 | -0.277274325 | 0.031268784 |
| Nadk | 2921.741327 | -0.277338957 | 4.35E-05 |
| Shpk | 370.7372046 | -0.277468762 | 0.014805683 |
| Nudt5 | 346.8752594 | -0.277576447 | 0.012143946 |
| LOC691807 | 2831.431651 | -0.277658844 | 0.000114678 |
| Kctd18 | 426.5432509 | -0.278321734 | 0.014298011 |

| | | | |
|---|---|---|---|
| Ide | 2268.735739 | -0.278535055 | 0.001203399 |
| St3gal4 | 1796.28603 | -0.278696643 | 0.000808968 |
| RGD1308428 | 1128.344699 | -0.27888897 | 0.000260516 |
| Myh11 | 551.6238395 | -0.278962252 | 0.049048928 |
| Rgl2 | 482.0046124 | -0.27898595 | 0.004899976 |
| Aldh7a1 | 5207.734117 | -0.27902742 | 0.023043151 |
| Tbcel | 900.466028 | -0.279193384 | 0.002341478 |
| Golga7 | 2052.268562 | -0.279233805 | 0.000105578 |
| Xpnpep3 | 570.4433314 | -0.279427961 | 0.002835569 |
| Aifm2 | 735.9400151 | -0.279519135 | 0.017728094 |
| Olfml1 | 3351.630332 | -0.279713112 | 0.012709084 |
| Fut10 | 176.5779489 | -0.279738186 | 0.044239617 |
| Hist1h1d | 1535.36029 | -0.27974678 | 0.00147506 |
| Slc25a1 | 2338.779878 | -0.279765294 | 0.001733326 |
| Sox8 | 4112.758431 | -0.279836705 | 0.001321266 |
| RGD1306001 | 127.4601272 | -0.279866078 | 0.0412796 |
| Chmp1b | 1297.225055 | -0.280005687 | 4.93E-05 |
| Slc50a1 | 232.8387899 | -0.280075181 | 0.032852267 |
| Lgr4 | 2629.962279 | -0.280183641 | 0.00109008 |
| Cbwd1 | 138.6866912 | -0.28018858 | 0.031809197 |
| H3f3b | 6516.870845 | -0.280203894 | 4.91E-05 |
| Echdc1 | 2014.914305 | -0.28041051 | 0.001254391 |
| Cdon | 496.6989649 | -0.280548046 | 0.01907998 |
| Cdkn1b | 1575.784518 | -0.280795544 | 0.000469054 |
| Naga | 1524.126946 | -0.28088961 | 0.004902714 |
| Pycr2 | 847.3585728 | -0.281019014 | 0.000294937 |
| Tns1 | 2334.311111 | -0.281389374 | 0.009511576 |
| Dpy19l4 | 1316.329968 | -0.281497891 | 0.003724962 |
| Il11ra1 | 1397.615958 | -0.281548562 | 0.003525287 |
| Gpr146 | 659.5995336 | -0.28244947 | 0.014458901 |
| Tfeb | 444.4976581 | -0.282618143 | 0.036242859 |
| Pdia4 | 2282.657662 | -0.282688373 | 0.002589987 |
| Hmbs | 719.808409 | -0.28269137 | 0.002587471 |
| Ttf2 | 203.6413885 | -0.282776071 | 0.028199717 |
| Col11a2 | 7315.227604 | -0.28298073 | 0.024514893 |
| Efcab1 | 425.9461808 | -0.282997364 | 0.037940202 |

| | | | |
|---|---|---|---|
| Zmpste24 | 1342.391552 | -0.283349733 | 0.002702754 |
| Lca5 | 638.7054691 | -0.283558727 | 0.005555961 |
| Rmdn1 | 601.5712727 | -0.283594126 | 0.015900006 |
| Fam3a | 437.1080252 | -0.283749389 | 0.003718108 |
| Atraid | 679.5640511 | -0.283950451 | 0.005079949 |
| Hddc3 | 878.372107 | -0.283984704 | 0.002552263 |
| Clcc1 | 2031.957358 | -0.284095369 | 0.003850644 |
| Atf1 | 427.3134283 | -0.284347384 | 0.002699573 |
| Pomt2 | 806.5557004 | -0.284592971 | 0.004264865 |
| Map4 | 16708.36334 | -0.284596293 | 9.00E-05 |
| Leprot | 941.9138989 | -0.284832163 | 0.001486999 |
| Slc2a1 | 6164.848063 | -0.284873035 | 0.007625273 |
| Cnnm3 | 1993.593531 | -0.284900183 | 0.00088711 |
| Odf2l | 215.2439478 | -0.284904481 | 0.011563061 |
| Lmnb1 | 347.718248 | -0.285094934 | 0.014458901 |
| Man2b2 | 1042.089632 | -0.285121296 | 0.001114318 |
| Gstm5 | 1615.807058 | -0.28522451 | 0.009575648 |
| Ptpn21 | 384.2612451 | -0.285346874 | 0.005595247 |
| Sh3bgrl | 7003.245083 | -0.285711632 | 0.000108464 |
| Akr1b10 | 137.89367 | -0.285922151 | 0.038470369 |
| Hexa | 1672.008087 | -0.286296355 | 7.54E-05 |
| Trps1 | 4674.772237 | -0.286397628 | 0.002201765 |
| Lhx2 | 4273.862141 | -0.286445984 | 0.002693928 |
| Ppp1r14b | 357.107468 | -0.286655108 | 0.009347348 |
| Cep89 | 351.14186 | -0.28698625 | 0.011770957 |
| Ccdc181 | 3379.926712 | -0.287137633 | 0.000965237 |
| Kctd11 | 101.2311042 | -0.28717446 | 0.049496072 |
| Lamp1 | 17200.62165 | -0.287784258 | 5.27E-05 |
| Ece1 | 3107.629842 | -0.28794183 | 0.003420564 |
| Manba | 835.0400628 | -0.287995194 | 0.005687357 |
| Zfp518a | 474.1570184 | -0.288147421 | 0.004512802 |
| Slc41a1 | 9175.873309 | -0.288152279 | 0.000338385 |
| RT1-Db1 | 39.67182752 | -0.288479863 | 0.047485036 |
| Ugdh | 578.6105825 | -0.288694995 | 0.001174011 |
| Homer3 | 395.8725911 | -0.288817389 | 0.011885047 |
| Prdx6 | 4508.947378 | -0.289068058 | 0.004578723 |

| | | | |
|---|---|---|---|
| Prex2 | 39321.65292 | -0.289686027 | 0.006640202 |
| Gli4 | 155.6594235 | -0.289765018 | 0.036470781 |
| Tmed5 | 1273.427111 | -0.290133223 | 0.000975283 |
| Ghr | 340.5189215 | -0.290453175 | 0.023432209 |
| Samhd1 | 393.5918394 | -0.290460459 | 0.002410757 |
| Dtwd2 | 115.3053339 | -0.290633703 | 0.045749172 |
| Gys1 | 2369.433981 | -0.290639652 | 0.005636274 |
| Nhlrc2 | 1028.643909 | -0.290795576 | 0.000325832 |
| March3 | 273.2977353 | -0.29086207 | 0.01745001 |
| Dars2 | 634.0943668 | -0.291091359 | 0.009913523 |
| Hmgn5b | 1058.601537 | -0.291108227 | 0.035717586 |
| Creb3l2 | 817.5286101 | -0.29153176 | 0.000576173 |
| Meis1 | 507.5488735 | -0.29172567 | 0.018439404 |
| Usp53 | 5121.239607 | -0.291798692 | 0.005507542 |
| Hmgn1 | 2915.478391 | -0.29196314 | 0.000588446 |
| Tmed10 | 3368.618384 | -0.292108299 | 4.00E-05 |
| Rp2 | 381.1778834 | -0.292720433 | 0.008681037 |
| Slc25a24 | 209.4409483 | -0.292904867 | 0.020807307 |
| Cyp2d4 | 1908.498657 | -0.293087354 | 0.038819866 |
| Hnrnph3 | 1888.851093 | -0.293201643 | 0.000938572 |
| Zfp438 | 395.8088716 | -0.293341292 | 0.012616283 |
| Evi2b | 2.765157428 | -0.293548874 | 0.037067128 |
| Nphp3 | 185.2818997 | -0.29372617 | 0.015256368 |
| Usp16 | 284.3954022 | -0.293818473 | 0.0119014 |
| Isy1 | 593.7094834 | -0.293994813 | 0.008393423 |
| Snap23 | 472.9566524 | -0.294339448 | 0.018326777 |
| Synm | 8112.455711 | -0.294444157 | 0.015565389 |
| Hps1 | 329.1839035 | -0.294482383 | 0.01528732 |
| Tspan33 | 1137.115502 | -0.294617591 | 0.001687318 |
| Galnt10 | 366.1753144 | -0.294661002 | 0.02643844 |
| Fzd10 | 824.192127 | -0.294711582 | 0.0335379 |
| Kctd5 | 948.9564661 | -0.294816173 | 0.000234264 |
| Lix1 | 7773.40573 | -0.295267114 | 0.00439152 |
| Tjp1 | 8444.812301 | -0.295581079 | 0.002920298 |
| Upp1 | 261.3489112 | -0.295614659 | 0.022583944 |
| Fam126a | 333.7462842 | -0.296077089 | 0.006804616 |

| | | | |
|---|---|---|---|
| Slc13a4 | 360.9933229 | -0.296266965 | 0.019982802 |
| Eef2k | 1901.064614 | -0.296284067 | 0.003490773 |
| Fchsd1 | 404.4158106 | -0.296480111 | 0.007796815 |
| Bmpr1a | 5014.569093 | -0.296538696 | 0.002582768 |
| Ilvbl | 895.4591129 | -0.296629543 | 0.002297468 |
| Idh2 | 6325.804573 | -0.296637523 | 0.008222036 |
| Galnt4 | 305.1357188 | -0.296735691 | 0.020330244 |
| Pld5 | 512.9258726 | -0.296777562 | 0.002560623 |
| Gabrg1 | 5499.902427 | -0.296878282 | 0.008140714 |
| Snx24 | 857.9614257 | -0.297607376 | 0.001358716 |
| Nedd1 | 239.8728777 | -0.297616473 | 0.007654493 |
| Josd2 | 378.5630128 | -0.297694131 | 0.010933972 |
| Shmt2 | 1200.374775 | -0.29795077 | 0.000185565 |
| Agl | 7039.47129 | -0.297981977 | 0.001394274 |
| Aldh5a1 | 18021.77382 | -0.298010006 | 0.000953118 |
| Mynn | 348.2948481 | -0.298253539 | 0.002301046 |
| Sufu | 447.033985 | -0.298791382 | 0.008139391 |
| Samd4a | 1347.084929 | -0.299133974 | 0.001611576 |
| Mocs1 | 251.4054521 | -0.299195277 | 0.033177595 |
| Echdc2 | 180.4696708 | -0.299234315 | 0.024340559 |
| Soat1 | 861.7657073 | -0.29940728 | 0.000663504 |
| Lrrc42 | 937.2286874 | -0.29946843 | 8.36E-05 |
| Mccc2 | 1720.730095 | -0.299586612 | 0.000184718 |
| Slc31a1 | 675.1949721 | -0.299903834 | 0.000643305 |
| Fbxw17 | 139.8387014 | -0.300337651 | 0.040433381 |
| Vipr2 | 333.1561831 | -0.300585065 | 0.013575402 |
| Cpt1a | 5650.307675 | -0.30090958 | 0.006071056 |
| Pi15 | 3.165705748 | -0.301067058 | 0.044144655 |
| Bloc1s5 | 977.891378 | -0.30116494 | 8.41E-05 |
| Mapre1 | 3120.842357 | -0.301605903 | 7.22E-06 |
| Tmem229a | 12892.47156 | -0.301736699 | 0.001396443 |
| Dock7 | 3596.059909 | -0.302123942 | 0.000738182 |
| Xkr8 | 179.9996595 | -0.302156929 | 0.022219191 |
| Tprn | 495.8440272 | -0.302169467 | 0.035184714 |
| Zcchc9 | 469.8340787 | -0.302179032 | 0.008541625 |
| Limk2 | 2604.559387 | -0.302206533 | 9.52E-05 |

| | | | |
|---|---|---|---|
| Unc119b | 1865.583068 | -0.302422094 | 0.000233712 |
| RT1-DMa | 234.7644155 | -0.302484261 | 0.00761303 |
| Slfn3 | 7.268729399 | -0.303059545 | 0.039854541 |
| Casc1 | 361.876886 | -0.303480972 | 0.018579337 |
| Hspb8 | 2785.567342 | -0.303542927 | 0.008678502 |
| Rassf8 | 405.8956824 | -0.303861348 | 0.015229178 |
| Mical1 | 326.8621608 | -0.304078195 | 0.006950792 |
| RGD1309808 | 3.29947623 | -0.304094296 | 0.039601016 |
| Xrcc6 | 1086.594258 | -0.305726637 | 0.0007671 |
| Phlpp1 | 13694.93358 | -0.305974984 | 0.000185192 |
| Tmc5 | 179.802349 | -0.306080414 | 0.01952029 |
| Pdcl | 1068.610766 | -0.306127047 | 1.43E-05 |
| Nxn | 1180.680705 | -0.306199676 | 0.009469222 |
| Dnm2 | 2634.409256 | -0.306330647 | 0.000393514 |
| Nek9 | 3001.769741 | -0.307133114 | 9.29E-06 |
| Ciita | 7.166136261 | -0.307148191 | 0.042706363 |
| Abhd14a | 376.3257465 | -0.307619826 | 0.016824371 |
| Pold1 | 153.7535386 | -0.307639999 | 0.027495278 |
| Hsd17b8 | 370.0985443 | -0.307690366 | 0.004757554 |
| Add3 | 17624.77184 | -0.307842267 | 0.001030091 |
| Sdc2 | 4191.530616 | -0.308042705 | 0.001319942 |
| Ptn | 10542.35611 | -0.308188588 | 0.000889959 |
| Pyroxd2 | 1086.412517 | -0.308350252 | 0.005373791 |
| Zkscan3 | 789.4841091 | -0.308350923 | 7.57E-05 |
| Lman1 | 1535.223001 | -0.308371028 | 0.005347779 |
| Scp2 | 10807.53534 | -0.308497505 | 2.84E-06 |
| Slc39a6 | 2395.628016 | -0.308724969 | 0.00057811 |
| Rb1 | 2004.763292 | -0.308740674 | 2.79E-07 |
| Mmel1 | 246.3104053 | -0.308829509 | 0.041589449 |
| Cmbl | 814.1849947 | -0.308929436 | 0.008863252 |
| Mfsd9 | 246.2275287 | -0.30903867 | 0.016247463 |
| Mthfd1 | 2729.898824 | -0.309044686 | 0.000361764 |
| Nnat | 12484.35954 | -0.309137105 | 0.044884491 |
| Tspan31 | 1181.395139 | -0.309619008 | 1.44E-05 |
| Nt5c1a | 701.1832533 | -0.309723033 | 0.00272057 |
| Qser1 | 1676.909655 | -0.309985578 | 0.000295469 |

| | | | |
|---|---|---|---|
| Saxo2 | 18.15530987 | -0.310041626 | 0.048474509 |
| Zfand3 | 3506.048634 | -0.310167275 | 1.65E-05 |
| Maged2 | 1867.969372 | -0.310259156 | 0.000574693 |
| Pon2 | 6350.686277 | -0.311037772 | 3.32E-05 |
| Wdr5b | 78.76310884 | -0.3113301 | 0.038470369 |
| S100b | 104607.3363 | -0.311698407 | 0.000455775 |
| Pygb | 29197.54581 | -0.311753221 | 0.004847825 |
| Gatm | 11695.28465 | -0.311764727 | 0.005433125 |
| Pdlim3 | 137.2981925 | -0.311883454 | 0.032291704 |
| Urod | 1613.379928 | -0.312033658 | 1.62E-05 |
| Mapkapk3 | 1176.78339 | -0.312053344 | 0.002516783 |
| Abhd5 | 285.539115 | -0.312252633 | 0.012425382 |
| Zfp347 | 147.9034602 | -0.312495687 | 0.031268784 |
| Lrrn1 | 2183.741271 | -0.312656717 | 0.005258036 |
| Isg20l2 | 236.9734112 | -0.312792382 | 0.007823216 |
| Cdk4 | 1158.799327 | -0.313220472 | 1.47E-06 |
| Sox7 | 9.071139069 | -0.313418537 | 0.047440318 |
| Nacc2 | 9707.817824 | -0.313419249 | 0.001229954 |
| Asun | 941.1690676 | -0.313448484 | 0.000838162 |
| Homez | 183.1910637 | -0.31364095 | 0.02522534 |
| Slc13a3 | 822.2823919 | -0.314135617 | 0.009498774 |
| Prrg1 | 277.4434098 | -0.314298614 | 0.011419725 |
| Smc4 | 699.2618736 | -0.314622676 | 0.000329023 |
| Cyfip1 | 4839.462969 | -0.314643086 | 0.000163673 |
| Vav1 | 6.915978178 | -0.314737424 | 0.049964755 |
| Ltbp3 | 3714.929705 | -0.314883659 | 0.000887467 |
| Eci2 | 1920.40612 | -0.315088758 | 0.002447781 |
| Cuedc1 | 1050.347999 | -0.31519257 | 0.000293664 |
| Rnh1 | 1698.987442 | -0.315239949 | 0.000191757 |
| Marcksl1 | 3236.097091 | -0.315285607 | 0.000945738 |
| Ephb3 | 1479.528171 | -0.315441044 | 0.003204694 |
| Ppfibp1 | 266.6488229 | -0.315606613 | 0.012927362 |
| Hibadh | 4746.084536 | -0.315707593 | 0.000179941 |
| Bcl7c | 189.3871918 | -0.315812489 | 0.023824043 |
| Gcdh | 2185.032535 | -0.316034123 | 2.66E-05 |
| Gtf3c6 | 550.430076 | -0.31616899 | 0.000931959 |

| | | | |
|---|---|---|---|
| Rnpepl1 | 760.0185604 | -0.316377998 | 0.001313851 |
| Ppp1r18 | 501.2890571 | -0.316420125 | 0.00765977 |
| Cers4 | 4771.910983 | -0.316854543 | 3.83E-05 |
| Ift81 | 1470.806804 | -0.317143859 | 0.001422611 |
| Sirt2 | 5950.083223 | -0.317249306 | 2.29E-06 |
| Arhgap11a | 98.56001846 | -0.317637794 | 0.042677002 |
| Rnf182 | 235.4521795 | -0.318034395 | 0.017953844 |
| Bckdha | 2353.587592 | -0.318185241 | 7.04E-05 |
| Acy3 | 544.6555451 | -0.318320321 | 0.003726911 |
| Slc29a4 | 936.9275851 | -0.318530653 | 0.030247125 |
| Ikbkb | 923.5995712 | -0.318555414 | 0.001292932 |
| Ush1c | 10.20889089 | -0.318697206 | 0.047311351 |
| Foxo1 | 5127.698472 | -0.318743344 | 0.002355386 |
| Smim20 | 608.521763 | -0.319064868 | 0.001819901 |
| Pgm2 | 493.296978 | -0.319639156 | 0.001261987 |
| Pgm1 | 7500.694658 | -0.320062566 | 0.000690822 |
| Suox | 960.6804851 | -0.320411692 | 0.000405767 |
| Ttc30b | 1110.989438 | -0.320715886 | 0.000302271 |
| Ccl19 | 14.04829775 | -0.320828592 | 0.039468079 |
| Nrf1 | 366.7064591 | -0.321271203 | 0.002409578 |
| Bnip2 | 718.9044744 | -0.321340157 | 0.003726911 |
| Pgm3 | 1044.74085 | -0.321415526 | 0.00126911 |
| Uap1l1 | 864.2317788 | -0.321529075 | 0.001044258 |
| Ybx1 | 3535.160113 | -0.321533162 | 3.32E-06 |
| Cnpy4 | 457.1627639 | -0.321879075 | 0.006005848 |
| Entpd1 | 593.3491746 | -0.321897939 | 0.004101305 |
| Aox1 | 1585.818715 | -0.322263077 | 0.008557968 |
| Vasp | 251.0373477 | -0.322311035 | 0.007912705 |
| Ptprf | 12515.07631 | -0.32244865 | 0.000114635 |
| Bphl | 260.741182 | -0.322452939 | 0.00640438 |
| Adamts9 | 255.3332002 | -0.322922521 | 0.016423768 |
| Ivd | 4673.971903 | -0.32294086 | 3.54E-05 |
| Cdh20 | 1232.444791 | -0.322995979 | 0.00796532 |
| Zfp191 | 1087.121235 | -0.323085932 | 0.000143119 |
| Slc7a2 | 2172.615112 | -0.323116719 | 0.004359616 |
| Triobp | 1028.299901 | -0.323380322 | 0.001338248 |

| | | | |
|---|---|---|---|
| Fzd5 | 404.7898622 | -0.323707768 | 0.033647662 |
| Slc22a5 | 452.9620632 | -0.323802647 | 0.004860029 |
| Gadd45a | 479.7684096 | -0.3238372 | 0.009943868 |
| Stk3 | 243.8728356 | -0.32392165 | 0.003947042 |
| Acbd5 | 2623.467371 | -0.323929659 | 0.000596735 |
| Mcee | 756.586444 | -0.324091925 | 0.000341805 |
| Cdk5rap2 | 1004.61931 | -0.324124467 | 0.005385166 |
| Dhcr7 | 1089.346459 | -0.324247232 | 0.00107576 |
| Nrbp2 | 3661.836529 | -0.324766699 | 0.000507732 |
| Thnsl2 | 543.1978275 | -0.324802947 | 0.004673849 |
| Enah | 9312.333747 | -0.325349803 | 0.000448243 |
| Hk2 | 161.2774685 | -0.325504689 | 0.027438401 |
| Qtrt1 | 128.468938 | -0.325550356 | 0.02194135 |
| Pea15 | 23445.62453 | -0.325715993 | 0.000107422 |
| Nfatc4 | 26.11071286 | -0.325961801 | 0.049923719 |
| Sox6 | 1630.630387 | -0.326070141 | 0.00053661 |
| Fcgr3a | 3.086445647 | -0.326236446 | 0.032074311 |
| Slc25a34 | 1256.387219 | -0.326241917 | 0.008678502 |
| Olfml2b | 344.6889814 | -0.326340867 | 0.005047001 |
| Idua | 611.7155219 | -0.326389854 | 0.004036889 |
| Lims1 | 484.3475684 | -0.326489819 | 0.006176954 |
| Mcc | 572.6196674 | -0.326583414 | 0.000823464 |
| Dusp22 | 454.2587829 | -0.326590196 | 0.00531492 |
| Hibch | 901.7419253 | -0.326736788 | 0.000619967 |
| E2f6 | 1637.36154 | -0.326949352 | 0.000469121 |
| Tnf | 1.762524257 | -0.326963432 | 0.019823737 |
| Ak4 | 1059.272201 | -0.327301072 | 0.003619204 |
| Smad5 | 1280.886302 | -0.327471792 | 0.000117452 |
| Tpp1 | 5429.14923 | -0.327496886 | 0.001697834 |
| Lpp | 394.5505997 | -0.328069616 | 0.00827811 |
| Hist1h2bh | 2677.542708 | -0.328129364 | 0.000132876 |
| Crybb1 | 189.8334814 | -0.328581052 | 0.02477514 |
| Cdc42ep4 | 7830.025073 | -0.328750058 | 0.001145431 |
| Ybx1-ps3 | 226.5945407 | -0.328955457 | 0.035963315 |
| Lama5 | 1311.856469 | -0.328955668 | 0.002519449 |
| Etfdh | 2241.25 | -0.329093883 | 0.000337688 |

| | | | |
|---|---|---|---|
| Rasa2 | 742.459277 | -0.329142614 | 5.00E-05 |
| Scara3 | 6553.652922 | -0.32943042 | 0.000596735 |
| Ttc30a | 77.01275803 | -0.329632941 | 0.025512861 |
| Acadl | 2873.465535 | -0.329713312 | 0.000557854 |
| Glyctk | 235.9889577 | -0.32990902 | 0.011998927 |
| Morn2 | 82.6535249 | -0.330239028 | 0.04296069 |
| Tmem43 | 1246.328405 | -0.330681653 | 6.53E-05 |
| Epb4.1l5 | 706.6881334 | -0.331074092 | 0.007259346 |
| Rab34 | 626.6469839 | -0.331362752 | 0.003204694 |
| Tpcn1 | 3785.442625 | -0.331598556 | 0.000183749 |
| Ryk | 713.1958242 | -0.331760248 | 0.000405255 |
| Crispld2 | 332.612338 | -0.331976275 | 0.004097155 |
| Lig1 | 734.7810456 | -0.332205598 | 0.002858114 |
| Phf19 | 64.57171449 | -0.332255756 | 0.038148756 |
| Plekhd1 | 1550.335482 | -0.332289374 | 0.010093494 |
| Rgs20 | 1471.675502 | -0.332344911 | 0.000353009 |
| Bcat2 | 269.9858126 | -0.33250103 | 0.019540605 |
| Hdac1 | 1138.58862 | -0.332524593 | 0.000591328 |
| Fam63a | 590.8465173 | -0.332704782 | 0.000576173 |
| Rorb | 4058.099634 | -0.332800043 | 5.94E-05 |
| Tvp23b | 1827.31169 | -0.333066844 | 0.00046407 |
| Vezf1 | 2851.488766 | -0.333169758 | 6.93E-05 |
| Calu | 4572.498741 | -0.333198219 | 1.30E-05 |
| Pdzrn3 | 3452.549846 | -0.333369261 | 0.000256999 |
| Gsta3 | 2.052993853 | -0.333396755 | 0.026870085 |
| Pccb | 1198.959005 | -0.333441382 | 0.000112123 |
| Sgk2 | 8.318590847 | -0.333757401 | 0.04597216 |
| Tram1 | 2221.651845 | -0.333882821 | 3.07E-06 |
| Gareml | 2654.379069 | -0.334060995 | 0.001165579 |
| Ccdc34 | 570.423595 | -0.334220193 | 0.005554656 |
| Nfia | 5332.166331 | -0.334254462 | 9.56E-06 |
| Psph | 723.0068132 | -0.334377567 | 0.000230461 |
| Misp | 4.638902535 | -0.334737832 | 0.033894787 |
| Ttc29 | 51.34108482 | -0.335584103 | 0.049445548 |
| Mcmbp | 1143.186846 | -0.335593377 | 3.53E-05 |
| Fut2 | 146.2167197 | -0.335731784 | 0.024327402 |

| | | | |
|---|---|---|---|
| Id4 | 9750.266173 | -0.335878586 | 0.00065761 |
| Plbd1 | 2.640780161 | -0.335964125 | 0.023707852 |
| Wdfy2 | 839.2254739 | -0.336087049 | 6.73E-05 |
| Ccpg1os | 222.6421595 | -0.33617721 | 0.015633038 |
| Kcnn4 | 69.8363374 | -0.336297289 | 0.042269342 |
| Trim67 | 111.214359 | -0.33644557 | 0.042823959 |
| Sepsecs | 548.7232344 | -0.336530126 | 0.000432354 |
| Fkbp14 | 464.0528205 | -0.33669655 | 0.010293887 |
| Nat1 | 87.97455916 | -0.336837668 | 0.034140641 |
| Fsip1 | 18.9048961 | -0.336947784 | 0.048672348 |
| Sqrdl | 262.1169806 | -0.33696998 | 0.016023915 |
| Hgf | 363.4949723 | -0.337212008 | 0.043093801 |
| Dusp19 | 441.9279549 | -0.337285607 | 0.001394274 |
| Il6st | 8283.342165 | -0.337304622 | 6.21E-06 |
| Rbl1 | 73.9620898 | -0.337746777 | 0.047815091 |
| Bcan | 46309.23645 | -0.337755476 | 0.005722311 |
| Slc1a2 | 94395.90528 | -0.337763628 | 0.000587052 |
| Acad11 | 1274.922716 | -0.33799542 | 0.00164972 |
| Trim47 | 424.6117663 | -0.338105396 | 0.030153488 |
| Agtrap | 898.2247554 | -0.338182073 | 3.70E-05 |
| Akt2 | 1869.074763 | -0.338285967 | 0.000554897 |
| Hsdl2 | 3129.41808 | -0.33855738 | 0.000230931 |
| Abhd3 | 11877.38417 | -0.338679128 | 0.000115356 |
| Fam129b | 5336.068757 | -0.338695212 | 0.000550425 |
| Asb4 | 11.51335147 | -0.338847498 | 0.039713537 |
| Tmem209 | 417.5581591 | -0.338949534 | 0.001033968 |
| Spata33 | 95.15822198 | -0.339080357 | 0.019893903 |
| Sorcs2 | 8215.832468 | -0.339277313 | 0.000995061 |
| Efemp1 | 882.4870263 | -0.339692242 | 0.006963741 |
| Lrp1 | 87542.76526 | -0.339935327 | 1.32E-05 |
| Echs1 | 3029.696237 | -0.340044392 | 0.000134598 |
| Dmd | 2603.784403 | -0.340119095 | 0.000581158 |
| Pi4k2b | 179.8432069 | -0.340279279 | 0.015396381 |
| Ston2 | 9614.032882 | -0.340511671 | 0.000751679 |
| Nmnat3 | 248.6797209 | -0.34067365 | 0.021368156 |
| Acad10 | 786.5963769 | -0.34095936 | 0.002275189 |

| | | | |
|---|---|---|---|
| Cyb5a | 1025.11467 | -0.34101528 | 0.005232594 |
| Tmpo | 1361.760039 | -0.341271448 | 0.000220235 |
| Gnai2 | 17810.93708 | -0.341337917 | 5.55E-05 |
| LOC688553 | 43.98132326 | -0.341374755 | 0.042409248 |
| Rbbp9 | 702.1551265 | -0.341492691 | 0.001025881 |
| Ostf1 | 613.4687068 | -0.341683914 | 0.004357857 |
| Snx5 | 2689.351102 | -0.341698774 | 2.35E-06 |
| Galr1 | 95.14737703 | -0.34218344 | 0.042409248 |
| Ggact | 322.9650715 | -0.342366584 | 0.003808499 |
| Mt1a | 223.097735 | -0.342582444 | 0.043729577 |
| Rbks | 296.2629058 | -0.342691499 | 0.003566259 |
| Ccdc153 | 231.9715651 | -0.342754725 | 0.042003049 |
| Csad | 1980.724015 | -0.342786997 | 0.001735392 |
| Mccc1 | 918.8240026 | -0.343348539 | 0.001820217 |
| Slc35f6 | 2060.175849 | -0.343498132 | 1.47E-05 |
| As3mt | 507.014296 | -0.343616851 | 0.001555482 |
| Bhlhe40 | 3357.065039 | -0.343831505 | 0.001865075 |
| LOC100364673 | 3.728824191 | -0.343868792 | 0.03696638 |
| Ctf1 | 275.1187479 | -0.343944312 | 0.013751834 |
| Rock1 | 2158.854624 | -0.343988296 | 0.002028217 |
| Gcsh | 3043.94606 | -0.344001393 | 4.26E-06 |
| Mob1a | 1339.649562 | -0.344743898 | 0.000576173 |
| Tmem129 | 1195.65044 | -0.34477887 | 3.30E-05 |
| Pex12 | 908.2417706 | -0.345250797 | 0.001311643 |
| Wdpcp | 244.4145362 | -0.345632548 | 0.010385901 |
| Pcdhgb7 | 3010.513897 | -0.345734504 | 0.001711373 |
| Nlrp3 | 14.98482104 | -0.345861417 | 0.044884491 |
| Erlin2 | 2338.668758 | -0.346075807 | 4.35E-05 |
| Gprc5b | 13512.12528 | -0.34716141 | 2.34E-05 |
| Nfatc3 | 578.4345503 | -0.347311336 | 0.001110041 |
| Cers2 | 812.266381 | -0.347311722 | 0.003850644 |
| Plod3 | 1388.083586 | -0.347465508 | 0.000234433 |
| Car5b | 286.3152958 | -0.347526373 | 0.02006339 |
| Hadhb | 5908.633887 | -0.347812303 | 3.53E-05 |
| Lhpp | 3260.215485 | -0.347913089 | 3.79E-06 |
| Btd | 706.2440744 | -0.347918736 | 0.001222282 |

| | | | |
|---|---|---|---|
| Hmgcs2 | 1486.9111 | -0.348158201 | 0.044508981 |
| Sh2d4b | 5.978623163 | -0.348327267 | 0.039468079 |
| Rnf13 | 2707.439563 | -0.348361328 | 1.94E-05 |
| P2rx2 | 22.42161244 | -0.348363393 | 0.046728685 |
| Gstt1 | 199.8133628 | -0.34878455 | 0.011060847 |
| Hes6 | 901.7931827 | -0.34884446 | 0.000246085 |
| RGD1563200 | 8.324478337 | -0.348869603 | 0.032312495 |
| S100a13 | 2051.88377 | -0.349191113 | 0.000903613 |
| Gpm6b | 34619.7633 | -0.349584927 | 7.58E-05 |
| Tjp3 | 113.3826029 | -0.349678278 | 0.043027099 |
| Ifitm2 | 49.83006656 | -0.349738073 | 0.046417234 |
| Fuz | 435.864729 | -0.349876659 | 0.003680004 |
| Tmem47 | 14859.1099 | -0.350144331 | 6.87E-05 |
| Itm2b | 27191.46409 | -0.350181186 | 8.42E-07 |
| Morn1 | 234.0551762 | -0.350329733 | 0.008678462 |
| Actl6a | 228.8814013 | -0.350810714 | 0.006672507 |
| Phkg1 | 209.8299986 | -0.351017945 | 0.026032667 |
| Rgcc | 2208.913424 | -0.351166521 | 0.000618466 |
| Dbndd2 | 4128.147462 | -0.351286588 | 0.008135499 |
| Zcchc24 | 10934.12277 | -0.351301225 | 7.26E-05 |
| S100a1 | 152.4479255 | -0.351386668 | 0.017034672 |
| Pign | 259.5120815 | -0.351486365 | 0.008529338 |
| Rnf144a | 1118.350814 | -0.351724517 | 0.001189698 |
| Prdx4 | 631.6195221 | -0.351729004 | 0.001444602 |
| Adgrg1 | 19953.7063 | -0.351925296 | 0.00140131 |
| Zfp217 | 87.81548671 | -0.351933095 | 0.0406727 |
| Pdgfrb | 2708.620284 | -0.352043041 | 0.001988952 |
| Slc26a1 | 285.1943242 | -0.352211325 | 0.00647735 |
| Idh1 | 1984.405879 | -0.352257718 | 6.72E-05 |
| Fubp3 | 1602.610461 | -0.352509654 | 0.000287459 |
| Vax1 | 114.648533 | -0.352639518 | 0.031496731 |
| Fgfr1 | 5969.190722 | -0.352991256 | 4.46E-05 |
| RGD1563941 | 18.9227848 | -0.353126212 | 0.044487038 |
| Gnai3 | 2361.693119 | -0.353228711 | 1.64E-05 |
| Hrsp12 | 1921.524766 | -0.353368058 | 0.001159916 |
| Slc5a11 | 61.1247949 | -0.353736809 | 0.040433381 |

| | | | |
|---|---|---|---|
| Sept10 | 265.0485289 | -0.354428058 | 0.004336596 |
| Stat2 | 1955.949098 | -0.355201649 | 0.006744468 |
| C1rl | 38.12957049 | -0.355320565 | 0.04310017 |
| Cldn10 | 2647.279521 | -0.355391108 | 0.000415334 |
| Slc1a3 | 106358.9699 | -0.355442539 | 0.000432354 |
| Pnrc2 | 2452.398375 | -0.35554024 | 0.000122062 |
| Abhd14b | 1338.505647 | -0.355608192 | 0.000156798 |
| Ctnnd1 | 3929.051393 | -0.355783096 | 1.79E-06 |
| Hsd11b1 | 2037.292352 | -0.355982359 | 0.003985309 |
| Usp51 | 44.53025215 | -0.356200516 | 0.036228063 |
| Casp8 | 7.227117953 | -0.356269283 | 0.037272121 |
| Ccdc173 | 45.71218867 | -0.356464236 | 0.043184844 |
| Arsk | 565.4318613 | -0.356562461 | 0.001052026 |
| Gatsl3 | 99.04186364 | -0.356725061 | 0.038216103 |
| Pcdhgc3 | 38078.3817 | -0.356810674 | 3.39E-05 |
| Ppdpf | 1320.738561 | -0.356922323 | 9.76E-05 |
| Rsph10b | 331.5359528 | -0.357311651 | 0.042533846 |
| RT1-CE14 | 35.40680738 | -0.357328594 | 0.039420784 |
| Slc46a1 | 307.6761507 | -0.357339727 | 0.000469054 |
| Daw1 | 68.39340183 | -0.357353483 | 0.037339971 |
| Kif26a | 364.9934752 | -0.357437309 | 0.005044796 |
| Kazald1 | 9.980896919 | -0.357812533 | 0.039617536 |
| Gng5 | 627.346271 | -0.35821699 | 0.000151302 |
| Fez2 | 674.180829 | -0.35853613 | 0.000492059 |
| Ppfibp2 | 510.2518042 | -0.358689112 | 0.001295659 |
| Prcp | 410.2565461 | -0.358770828 | 0.004838615 |
| Ccl7 | 8.096424466 | -0.35886563 | 0.026054781 |
| Sppl2a | 856.1196912 | -0.358919423 | 0.000166845 |
| Chp2 | 40.66572477 | -0.358939097 | 0.042112327 |
| Spag1 | 305.0813482 | -0.359021196 | 0.010961208 |
| Irak4 | 157.6712309 | -0.359039829 | 0.016014021 |
| Dars | 1427.144194 | -0.359246302 | 3.53E-05 |
| Atp1a2 | 204784.3388 | -0.359367155 | 0.003655795 |
| Tcf12 | 1714.158002 | -0.359369411 | 4.77E-06 |
| Arhgef16 | 256.9352258 | -0.359388947 | 0.026003468 |
| Traf7 | 1874.442683 | -0.359573223 | 5.67E-05 |

| | | | |
|---|---|---|---|
| Szrd1 | 2400.305527 | -0.359631322 | 7.88E-07 |
| Fuom | 351.0452982 | -0.35965342 | 0.002535214 |
| Appl2 | 4373.586996 | -0.359719868 | 0.001114409 |
| Gli2 | 524.7383631 | -0.35972142 | 0.003827122 |
| Tyms | 368.8431723 | -0.359787415 | 0.000347896 |
| Pofut1 | 280.7621367 | -0.360201159 | 0.0019215 |
| Tcf3 | 741.1219672 | -0.360308885 | 0.000224724 |
| Mycbp | 187.6140934 | -0.360313432 | 0.018793859 |
| Krt36 | 4.678129486 | -0.36033185 | 0.030902467 |
| Pttg1 | 16.7007507 | -0.360507267 | 0.039658906 |
| Rassf6 | 19.10155522 | -0.360565207 | 0.038840647 |
| Myl9 | 131.5297252 | -0.360703618 | 0.021296418 |
| Stk38 | 517.004915 | -0.360869346 | 0.000254846 |
| Stard8 | 958.7171757 | -0.361128858 | 0.000351222 |
| Tnpo1 | 5045.348078 | -0.361145813 | 5.67E-05 |
| Nadk2 | 4703.038573 | -0.36115408 | 6.92E-05 |
| Fam107a | 70795.49882 | -0.36137468 | 0.000145887 |
| Kctd15 | 858.3104509 | -0.361400586 | 2.16E-05 |
| Nes | 292.329148 | -0.361516761 | 0.013285891 |
| Vwf | 64.10889472 | -0.361797043 | 0.041160099 |
| Ninj2 | 26.8239543 | -0.361802514 | 0.037627349 |
| Cspg5 | 24903.45784 | -0.361936155 | 1.49E-05 |
| Rdm1 | 181.0526754 | -0.362015811 | 0.003299618 |
| Chpt1 | 1555.072714 | -0.362094304 | 0.000468891 |
| Spc25 | 21.86830681 | -0.362170591 | 0.041081416 |
| Dnase1l1 | 147.15116 | -0.362319762 | 0.008725978 |
| Hist1h4b | 480.3773093 | -0.362543584 | 0.001380051 |
| Serpine2 | 15861.23118 | -0.362625782 | 0.00073562 |
| Parva | 354.7213319 | -0.362661044 | 0.003674012 |
| Asl | 667.3873725 | -0.36283114 | 9.49E-06 |
| Nit2 | 909.4243632 | -0.363053179 | 0.000308642 |
| Tp53inp1 | 223.4011143 | -0.363320825 | 0.012923605 |
| Fam199x | 165.0204365 | -0.363358373 | 0.009606499 |
| Glb1 | 889.63693 | -0.363409915 | 0.000849668 |
| Mns1 | 295.0559001 | -0.364019095 | 0.038188123 |
| Alkbh3 | 850.2483735 | -0.364067494 | 0.000320745 |

| | | | |
|---|---|---|---|
| Ttyh1 | 71774.93464 | -0.36407496 | 0.000295892 |
| Abcb4 | 340.62073 | -0.364121949 | 0.001374178 |
| Mapk12 | 307.8690019 | -0.36454475 | 0.000772661 |
| Npc2 | 2081.004334 | -0.364576938 | 4.72E-05 |
| Ahnak | 2403.067686 | -0.364855408 | 0.028638588 |
| Tecta | 194.3048745 | -0.364977701 | 0.031254329 |
| Cryz | 348.3710252 | -0.365029611 | 0.005060903 |
| Vangl1 | 404.9882031 | -0.365141546 | 0.003129479 |
| Marveld1 | 130.4266097 | -0.365231546 | 0.020267766 |
| Gpx8 | 694.5496553 | -0.365418233 | 0.000397486 |
| Kcnj10 | 4783.315697 | -0.365698333 | 0.001696806 |
| Nckap1l | 23.75282459 | -0.365959844 | 0.037272121 |
| Mfap4 | 66.62037159 | -0.366213591 | 0.035043934 |
| Slc27a6 | 19.39994421 | -0.366543871 | 0.037214587 |
| Dcxr | 484.8964375 | -0.366569858 | 0.001816265 |
| Dysf | 45.08355698 | -0.366601566 | 0.039214393 |
| Ccdc8 | 563.5743852 | -0.366682289 | 0.00136991 |
| Crtap | 37.48022438 | -0.366746521 | 0.036435097 |
| F2r | 4827.928003 | -0.36675832 | 4.69E-05 |
| Prdx1 | 3241.931047 | -0.36678119 | 7.05E-06 |
| Arfip1 | 506.5223226 | -0.36680182 | 8.51E-05 |
| Cyp4f6 | 351.253897 | -0.366805297 | 0.014845903 |
| Dag1 | 14104.86076 | -0.366818319 | 8.73E-07 |
| Rnd3 | 1484.867647 | -0.367058597 | 0.000536919 |
| Ntsr2 | 19237.3237 | -0.367139847 | 0.000756062 |
| Pcdh15 | 192.3158828 | -0.367872143 | 0.015052782 |
| Gna13 | 2272.141934 | -0.367972836 | 8.37E-05 |
| Acadm | 2342.819925 | -0.368165659 | 0.000408254 |
| Ccr5 | 17.16083871 | -0.369131593 | 0.034969901 |
| Scarb2 | 14539.05615 | -0.369244105 | 8.04E-06 |
| Vim | 2065.142253 | -0.36928236 | 0.036651766 |
| Tnfaip8 | 310.4134588 | -0.369448457 | 0.010272006 |
| LOC501110 | 925.0511185 | -0.36972553 | 0.000283503 |
| Pgpep1 | 712.2117835 | -0.369846047 | 0.000444941 |
| Adpgk | 766.9935457 | -0.369911928 | 0.005297894 |
| Nkain4 | 1015.514653 | -0.37015232 | 0.000192625 |

| | | | |
|---|---|---|---|
| Ugt8 | 705.2233727 | -0.370261322 | 0.037643265 |
| Foxo4 | 997.4927159 | -0.370400006 | 1.58E-05 |
| Igsf1 | 15691.72123 | -0.370460578 | 0.013128539 |
| Creb3l1 | 263.2578653 | -0.370672884 | 0.025082747 |
| Sod3 | 3028.840446 | -0.370753484 | 0.001324779 |
| Cyp4f17 | 387.5684973 | -0.371117167 | 0.010565155 |
| Dnah1 | 374.8406766 | -0.371150551 | 0.034718813 |
| Rnf215 | 749.9217438 | -0.371364519 | 0.000343829 |
| Pfkfb3 | 1627.055507 | -0.371475146 | 0.000777536 |
| Atp7b | 1293.224135 | -0.371607637 | 0.003149157 |
| Lrp4 | 12963.76719 | -0.371984986 | 0.001207964 |
| Acss3 | 222.6904118 | -0.372691892 | 0.021996257 |
| Elovl2 | 6226.313337 | -0.372731124 | 0.000804749 |
| Krcc1 | 869.1659847 | -0.372756668 | 0.000116805 |
| Aass | 1334.25293 | -0.373086001 | 0.004112093 |
| Gldc | 4472.112319 | -0.373307857 | 0.000683368 |
| Syde2 | 560.5002854 | -0.373314686 | 0.000777536 |
| Stbd1 | 300.754274 | -0.373475707 | 0.007259346 |
| Tmtc2 | 1303.050047 | -0.373971423 | 0.000837892 |
| Knstrn | 7.35894918 | -0.373986689 | 0.031496731 |
| Rhbdd1 | 157.7251396 | -0.37426351 | 0.012670274 |
| Pragmin | 1102.970697 | -0.374688781 | 1.91E-05 |
| Mgmt | 775.9208661 | -0.374736728 | 0.00161893 |
| Dhrs4 | 539.4593488 | -0.37480995 | 0.001826517 |
| Tmem51 | 767.201858 | -0.374960138 | 0.001123339 |
| Evc2 | 330.8388429 | -0.375684213 | 0.003842088 |
| Laptm4a | 17497.91121 | -0.375723285 | 6.13E-05 |
| Renbp | 455.236964 | -0.376146627 | 0.000830689 |
| Enho | 3279.600897 | -0.376253471 | 1.17E-05 |
| Gli3 | 4389.376334 | -0.376511068 | 0.00013422 |
| Pmp22 | 739.851665 | -0.37653204 | 0.007346534 |
| Prrx1 | 857.5482181 | -0.376865229 | 0.001726845 |
| Rdx | 5135.405441 | -0.377009471 | 9.81E-06 |
| Qk | 16200.84212 | -0.377095561 | 2.57E-05 |
| Spata1 | 11.55197792 | -0.377207066 | 0.033439949 |
| Fam181b | 2552.939065 | -0.377342729 | 0.001644076 |

| | | | |
|---|---|---|---|
| Sfxn5 | 12096.97834 | -0.377450829 | 0.000206311 |
| Cpe | 172620.0819 | -0.377622209 | 1.35E-05 |
| Uhrf1 | 53.1144127 | -0.377676483 | 0.032905522 |
| Scrn2 | 248.9833463 | -0.377708445 | 0.003901574 |
| Dhtkd1 | 2063.835 | -0.378131023 | 0.000187224 |
| Fgfr3 | 10251.77886 | -0.37841055 | 0.000348327 |
| Col6a2 | 52.94161645 | -0.378457846 | 0.034450065 |
| S1pr1 | 23183.90297 | -0.378535079 | 0.000115339 |
| Mdh1b | 54.60343833 | -0.378736004 | 0.025446431 |
| Fads2 | 10344.38817 | -0.378833586 | 0.000135813 |
| Sdsl | 423.8483164 | -0.378853415 | 0.005841193 |
| Yif1a | 613.2039686 | -0.378902644 | 7.29E-05 |
| Col9a2 | 102.3567557 | -0.378994667 | 0.020320492 |
| RGD1565002 | 2758.301957 | -0.379004954 | 1.17E-05 |
| Ppap2b | 50531.47532 | -0.379189929 | 7.95E-05 |
| Fabp5 | 2548.396952 | -0.379376564 | 8.81E-05 |
| Aco1 | 2340.077642 | -0.379557046 | 7.83E-05 |
| RGD1305464 | 777.3087283 | -0.37960189 | 0.000490616 |
| Ybx3 | 528.785864 | -0.379647369 | 0.001717755 |
| Nme4 | 57.30031536 | -0.379774783 | 0.030247125 |
| Aifm3 | 4524.334238 | -0.380005104 | 0.002203946 |
| Snapc2 | 1872.280622 | -0.380007743 | 1.74E-05 |
| Clec2g | 13.72667958 | -0.380183116 | 0.031267903 |
| LOC288978 | 26.39961035 | -0.380484613 | 0.033768027 |
| Tspan12 | 4150.498507 | -0.380673933 | 2.53E-07 |
| Metrn | 1213.497177 | -0.381168907 | 2.68E-05 |
| Calml4 | 37.95295549 | -0.381290185 | 0.031979674 |
| Acads | 1009.129639 | -0.381357427 | 0.001581032 |
| Rhpn1 | 556.8295342 | -0.381906656 | 0.000878282 |
| Slc9a3r1 | 5707.830867 | -0.381959425 | 0.000722389 |
| Fam60a | 48.17006376 | -0.382016539 | 0.033029566 |
| Fat1 | 15641.28737 | -0.382083733 | 4.88E-06 |
| Gstt3 | 3240.84078 | -0.382271734 | 8.94E-05 |
| Anxa1 | 128.375859 | -0.382567213 | 0.031677979 |
| Ddr2 | 326.0566843 | -0.382603708 | 0.01576116 |
| Lmcd1 | 1315.681668 | -0.382840275 | 0.001581032 |

| | | | |
|---|---|---|---|
| Fah | 327.952372 | -0.383361629 | 0.012493308 |
| Pxdc1 | 194.1982026 | -0.38336887 | 0.020791071 |
| Zfyve21 | 574.0640566 | -0.383955688 | 0.000207213 |
| Scd2 | 346650.6894 | -0.384317973 | 5.54E-05 |
| Crot | 1357.328837 | -0.384656272 | 0.0001812 |
| Plscr4 | 242.9775301 | -0.38478051 | 0.017643283 |
| Pygm | 9257.611794 | -0.384820283 | 0.001395814 |
| Dnase2 | 1006.967241 | -0.384853655 | 0.000200117 |
| Dhrs1 | 359.7163656 | -0.384967798 | 0.000230931 |
| Fnta | 2684.029243 | -0.385012437 | 1.34E-07 |
| Heyl | 4774.826673 | -0.385222417 | 0.000723934 |
| Fut4 | 78.20495073 | -0.385786963 | 0.028630406 |
| Jam2 | 6401.229628 | -0.385894185 | 9.58E-05 |
| Trim65 | 219.1807035 | -0.385980311 | 0.002691991 |
| Hcrt | 16.89573621 | -0.38639464 | 0.029357602 |
| Mcm2 | 362.5051546 | -0.386529899 | 0.002355386 |
| Lca5l | 17.57040001 | -0.386556921 | 0.030453358 |
| Nfe2l2 | 2151.335637 | -0.387037292 | 4.90E-05 |
| Rab31 | 4565.230189 | -0.387311842 | 2.09E-07 |
| Casq2 | 124.4332335 | -0.387504258 | 0.021067014 |
| Slc12a9 | 777.7624071 | -0.387654151 | 5.78E-05 |
| Zbtb7c | 230.1802483 | -0.387875869 | 0.011648139 |
| Cml5 | 259.3311474 | -0.388001324 | 0.005074428 |
| Rhpn2 | 1076.593574 | -0.38803332 | 0.000316507 |
| Cd180 | 30.20865525 | -0.388170918 | 0.031456782 |
| Aldh2 | 9208.087345 | -0.388743502 | 6.66E-05 |
| RGD1306233 | 151.5561694 | -0.389176732 | 0.019805021 |
| Klhl5 | 9698.295517 | -0.38928007 | 3.23E-05 |
| Troap | 5.593127266 | -0.390062134 | 0.028199717 |
| Thsd1 | 370.3283196 | -0.39010024 | 0.010171222 |
| Prkd2 | 222.87828 | -0.390168737 | 0.000878282 |
| Ascl1 | 178.0338984 | -0.390589107 | 0.025550947 |
| Stk36 | 221.4670856 | -0.390765038 | 0.013128539 |
| Megf10 | 1991.079003 | -0.390856423 | 0.00329538 |
| Pak4 | 677.0842714 | -0.390925126 | 9.42E-07 |
| Inppl1 | 4411.705763 | -0.391041333 | 1.41E-05 |

| | | | |
|---|---|---|---|
| Kif11 | 39.99416543 | -0.391047182 | 0.028494727 |
| Gpc5 | 5775.268384 | -0.391176343 | 0.000402658 |
| Sepp1 | 23589.14999 | -0.39123527 | 3.19E-06 |
| Fancl | 208.7614492 | -0.391470449 | 0.005897282 |
| Prr18 | 70.46651866 | -0.391560713 | 0.025624278 |
| Zfyve16 | 753.608518 | -0.391677407 | 1.08E-05 |
| Myh14 | 5092.371617 | -0.391681585 | 1.21E-05 |
| Ddah2 | 4005.360987 | -0.391745849 | 6.41E-05 |
| Spsb2 | 316.9837448 | -0.392033017 | 0.000230745 |
| Gpc3 | 273.5332301 | -0.392473545 | 0.025986166 |
| Acaa2 | 1330.977722 | -0.392494273 | 0.000843972 |
| Gins1 | 19.5471652 | -0.392531393 | 0.029993023 |
| Ptgr2 | 3955.463372 | -0.393067948 | 0.000222439 |
| Mag | 1649.993673 | -0.393208605 | 0.029860606 |
| Myrf | 1343.902879 | -0.393254371 | 0.029116188 |
| LOC100910802 | 40.58801339 | -0.393287193 | 0.029109341 |
| Ncan | 23838.32895 | -0.393601241 | 0.000896569 |
| Atf7 | 401.2859129 | -0.393652384 | 0.001920802 |
| St5 | 1812.727768 | -0.393781261 | 6.26E-06 |
| Abcd4 | 145.2950364 | -0.394022557 | 0.002568099 |
| Ppp1r3c | 6445.630784 | -0.394054109 | 0.000151507 |
| Evi2a | 333.3219161 | -0.394169847 | 0.028630406 |
| Gpld1 | 1867.925403 | -0.394184569 | 0.001343019 |
| Cyp4f5 | 481.0050729 | -0.394260249 | 0.001687461 |
| Pld2 | 2939.418729 | -0.394336223 | 0.00075972 |
| Sox12 | 349.4286677 | -0.394451819 | 0.000619251 |
| Plekhb1 | 12761.84171 | -0.394496225 | 7.20E-06 |
| Rela | 1125.469064 | -0.394637304 | 6.15E-06 |
| Pld1 | 2090.642955 | -0.394657195 | 0.000125189 |
| Usp40 | 704.8921822 | -0.394666907 | 0.000137456 |
| Mmd2 | 16910.4005 | -0.395063532 | 2.17E-05 |
| Slc7a5 | 2968.883137 | -0.39516262 | 1.37E-05 |
| Gal3st4 | 458.8895979 | -0.395235935 | 0.00272762 |
| Bax | 594.908267 | -0.395236658 | 0.000609415 |
| Sh3bp4 | 963.0744342 | -0.396137212 | 0.000878282 |
| Slc4a2 | 1899.78806 | -0.396620946 | 1.36E-05 |

| | | | |
|---|---|---|---|
| Plekhh1 | 692.4896066 | -0.39671517 | 0.01739845 |
| Oplah | 1669.965824 | -0.396812655 | 3.72E-05 |
| Tmem123 | 416.665556 | -0.397207621 | 0.012735421 |
| Plagl1 | 193.5604367 | -0.397433661 | 0.028630406 |
| Fzd2 | 3137.246217 | -0.397527407 | 0.000233712 |
| Tmem159 | 59.77577185 | -0.39754073 | 0.023259789 |
| Chst7 | 214.9631663 | -0.39766621 | 0.008557968 |
| Zfp474 | 22.07558895 | -0.397699616 | 0.027631103 |
| Myo9b | 1900.970247 | -0.397765009 | 1.01E-06 |
| Apoe | 374076.2724 | -0.398136158 | 0.000187533 |
| Prex1 | 12375.84969 | -0.398182754 | 4.82E-06 |
| Chi3l1 | 10917.5846 | -0.398208546 | 0.000527927 |
| Sfrp2 | 168.2914911 | -0.398251082 | 0.028307058 |
| Sh3tc1 | 141.3268284 | -0.398388558 | 0.013331932 |
| March8 | 2014.943587 | -0.398559159 | 1.78E-06 |
| Ccdc39 | 505.096093 | -0.398749548 | 0.009632571 |
| Ccdc67 | 12.87524138 | -0.398911489 | 0.027405441 |
| Cdt1 | 42.59610033 | -0.39894226 | 0.026019941 |
| Siglec15 | 42.32944654 | -0.399051649 | 0.026526406 |
| Wipi1 | 1756.628596 | -0.3992808 | 2.85E-06 |
| Efhb | 23.79559861 | -0.399350719 | 0.027761709 |
| Pak2 | 2058.265613 | -0.399365828 | 6.07E-06 |
| Sorbs3 | 2800.974238 | -0.399449807 | 2.85E-06 |
| Lfng | 1866.679594 | -0.399523642 | 0.001632582 |
| Lrrc46 | 104.8456298 | -0.399601392 | 0.027698992 |
| Erap1 | 1211.763992 | -0.399611196 | 0.000498757 |
| Sds | 75.81277944 | -0.399725506 | 0.021963602 |
| Smpdl3a | 1105.239165 | -0.399901184 | 0.000558971 |
| Pvrl3 | 761.7652985 | -0.400202458 | 0.000725396 |
| Asrgl1 | 8848.273349 | -0.400308536 | 1.76E-05 |
| Zfp395 | 854.9594113 | -0.400389854 | 0.000174821 |
| Casp6 | 79.96747464 | -0.400598472 | 0.020010797 |
| Mcm5 | 27.81076056 | -0.400923491 | 0.027070781 |
| Rarg | 340.8848672 | -0.400960565 | 0.000566823 |
| Zfp423 | 3948.33327 | -0.401579546 | 3.42E-05 |
| Zhx3 | 5802.522693 | -0.401729776 | 0.002767481 |

| | | | |
|---|---|---|---|
| Phka1 | 3286.447163 | -0.401811683 | 0.001724088 |
| Ptger2 | 49.84309021 | -0.40207473 | 0.026882892 |
| Tctn2 | 325.2349224 | -0.402890555 | 0.003406042 |
| Ccdc151 | 49.63854072 | -0.402999167 | 0.02718902 |
| Cercam | 91.31421799 | -0.403047041 | 0.014092697 |
| Tmem107 | 262.1152408 | -0.403238542 | 0.007401927 |
| Mak | 35.90832103 | -0.403267144 | 0.027025842 |
| Agmo | 59.5843993 | -0.404398336 | 0.025829894 |
| Ap3m1 | 1530.486474 | -0.404671125 | 0.000138614 |
| RGD1562726 | 161.8313924 | -0.404861805 | 0.017900943 |
| Paqr6 | 464.0482494 | -0.404930007 | 0.008909046 |
| Cuedc2 | 192.8808601 | -0.40517496 | 0.006660123 |
| B4galt1 | 49.03876939 | -0.405455377 | 0.024791848 |
| Cyp20a1 | 781.8574386 | -0.405556508 | 9.80E-05 |
| Sox13 | 734.5677016 | -0.405740947 | 0.000301588 |
| Ifi30 | 287.3235609 | -0.405842462 | 0.004491482 |
| Notch1 | 10361.15923 | -0.406024123 | 0.00012169 |
| Rsph1 | 139.8107757 | -0.406254616 | 0.026059621 |
| Wnt7b | 1233.605571 | -0.406555887 | 0.000176068 |
| Ptk7 | 561.3215987 | -0.406663939 | 0.000131133 |
| Ppap2a | 742.9139466 | -0.407256756 | 3.07E-05 |
| Amotl2 | 1716.783038 | -0.407469641 | 0.000110565 |
| Rsph9 | 172.2381293 | -0.407756547 | 0.020509223 |
| Pxn | 666.4978772 | -0.407926351 | 1.67E-05 |
| Sardh | 1477.656285 | -0.408152635 | 0.000160608 |
| Zfp496 | 649.6456004 | -0.408207972 | 4.13E-06 |
| Tmem120a | 489.6051141 | -0.408756759 | 0.001099829 |
| Aldh6a1 | 10260.25185 | -0.40881622 | 0.000126759 |
| Slc15a3 | 14.40591572 | -0.409081917 | 0.025802102 |
| Btbd7 | 1218.286243 | -0.409156675 | 1.04E-05 |
| Eci1 | 891.4141095 | -0.409364227 | 0.000125238 |
| Glis2 | 659.1759463 | -0.409725095 | 0.000107238 |
| Gna12 | 5740.610811 | -0.410081465 | 8.11E-07 |
| Aldh9a1 | 4156.102476 | -0.410100524 | 1.41E-05 |
| Lima1 | 1004.21112 | -0.410141175 | 0.000763393 |
| Ptpn13 | 2714.035316 | -0.410210365 | 0.000133993 |

| | | | |
|---|---|---|---|
| Car8 | 1331.030883 | -0.41025876 | 0.000701546 |
| Ndrg2 | 135232.729 | -0.410523906 | 4.69E-05 |
| Tor3a | 442.8835884 | -0.411036606 | 3.21E-05 |
| Glud1 | 40168.52143 | -0.411079384 | 0.000292673 |
| Timeless | 49.73666388 | -0.411127617 | 0.020461489 |
| Aldh1l1 | 2703.226747 | -0.41113675 | 0.001096119 |
| Yap1 | 1222.771118 | -0.411159676 | 0.000127238 |
| Bmf | 160.3864482 | -0.411237187 | 0.018890611 |
| Fuca1 | 1800.955774 | -0.411465931 | 4.67E-06 |
| Dusp18 | 2259.010425 | -0.411549901 | 2.55E-06 |
| Rnase4 | 5888.370089 | -0.41163711 | 6.06E-06 |
| Slc27a1 | 12414.14315 | -0.411746266 | 3.30E-05 |
| RGD1560608 | 16.5871031 | -0.411901737 | 0.025284946 |
| Stxbp3 | 1359.72472 | -0.412045642 | 4.29E-05 |
| Tp53i3 | 256.4649932 | -0.412440779 | 0.000659045 |
| Slfn2 | 9.584119567 | -0.412451603 | 0.025502514 |
| Ankfy1 | 4732.173466 | -0.412762948 | 3.26E-09 |
| Stard6 | 103.3671264 | -0.413039448 | 0.011103981 |
| Pou3f2 | 3585.370291 | -0.413074666 | 1.27E-05 |
| LOC100151767 | 555.8839576 | -0.413393471 | 1.62E-05 |
| Ech1 | 2518.713969 | -0.413460387 | 7.88E-07 |
| Sept2 | 8925.02829 | -0.413537339 | 3.49E-06 |
| Ctdsp1 | 1791.493611 | -0.413786225 | 1.49E-05 |
| Cenpj | 371.2301851 | -0.413944901 | 0.001422611 |
| Acadvl | 2617.53926 | -0.41407173 | 5.13E-05 |
| Acsbg1 | 30720.82865 | -0.414453277 | 0.000246085 |
| Blnk | 15.76471571 | -0.41449924 | 0.024882813 |
| Itpripl2 | 62.36426524 | -0.41476474 | 0.009511576 |
| Ptbp1 | 2185.581837 | -0.414793395 | 0.000123121 |
| Sfrp1 | 1815.304046 | -0.414850353 | 0.019716019 |
| C4b | 18.814218 | -0.415077389 | 0.025051737 |
| Rgs3 | 1857.525695 | -0.415081201 | 1.22E-05 |
| Polm | 291.0766259 | -0.415160586 | 0.00020808 |
| Zfp36l1 | 3428.511111 | -0.415922503 | 4.38E-05 |
| Magt1 | 659.8636509 | -0.416044515 | 0.000170694 |
| Hipk2 | 11027.6631 | -0.416468928 | 3.97E-08 |

| | | | |
|---|---|---|---|
| Scrib | 2667.05886 | -0.416800285 | 1.21E-07 |
| Afp | 24.8788919 | -0.416860882 | 0.024056503 |
| Frem1 | 306.9182793 | -0.416929509 | 0.01118086 |
| Smpd2 | 298.5983889 | -0.416963602 | 0.000364311 |
| Cdk2 | 84.38033841 | -0.416976239 | 0.013589627 |
| Frem2 | 1675.098343 | -0.417115842 | 3.53E-05 |
| Plod1 | 1709.731283 | -0.417161853 | 0.000366941 |
| Cd151 | 1999.025687 | -0.417246714 | 8.11E-07 |
| Esr1 | 55.85089516 | -0.417503437 | 0.017622271 |
| Tinagl1 | 11.85305164 | -0.417546292 | 0.024399304 |
| Creld2 | 427.7560332 | -0.417931766 | 0.002307581 |
| Papss2 | 3135.708794 | -0.417935198 | 0.000365861 |
| Lrrc23 | 278.5630279 | -0.417979212 | 0.012080043 |
| Top2a | 47.45441091 | -0.417998835 | 0.023832172 |
| Ptrf | 114.9093674 | -0.418076697 | 0.02030254 |
| Pih1d1 | 209.3606348 | -0.418418914 | 0.001897855 |
| Sucnr1 | 58.60810302 | -0.418430652 | 0.017143229 |
| Lrrc8a | 8316.165133 | -0.418546025 | 3.49E-05 |
| Lrrc16a | 3728.804667 | -0.4185817 | 1.92E-05 |
| Mansc1 | 342.4638211 | -0.418832897 | 0.000398218 |
| Kif13a | 1737.486241 | -0.41920073 | 5.81E-06 |
| Ss18 | 1603.495393 | -0.419413878 | 9.54E-06 |
| Gstm1 | 15132.16384 | -0.41954664 | 0.000467229 |
| Fermt2 | 7819.046597 | -0.419867006 | 6.54E-05 |
| Glul | 141104.3549 | -0.41997463 | 3.95E-05 |
| Syngr2 | 301.8311734 | -0.420049549 | 0.001739637 |
| Elovl5 | 1801.614456 | -0.420104671 | 1.70E-05 |
| Ndp | 1529.688905 | -0.420158139 | 2.16E-05 |
| Prom1 | 19.47145178 | -0.420534918 | 0.023566641 |
| Mdfi | 6.280517483 | -0.420828085 | 0.023643916 |
| Car13 | 61.54410913 | -0.420921796 | 0.018439746 |
| Spidr | 166.7257861 | -0.421197002 | 0.001268238 |
| Nuf2 | 33.06244867 | -0.42120565 | 0.016878704 |
| Psd2 | 13728.92845 | -0.42133456 | 5.27E-05 |
| Tead1 | 612.884916 | -0.421536807 | 0.000164034 |
| Slc8b1 | 380.2376759 | -0.421808005 | 0.000250847 |

| | | | |
|---|---|---|---|
| Gkap1 | 1344.281076 | -0.421823587 | 1.64E-07 |
| Mybl1 | 266.8497999 | -0.421940649 | 0.001078256 |
| Zfp219 | 1741.27167 | -0.422563833 | 3.61E-07 |
| Flnb | 1333.321807 | -0.422632881 | 1.82E-05 |
| Podxl | 333.31798 | -0.422682699 | 0.006796697 |
| Cpt2 | 900.0251867 | -0.42280029 | 0.0006106 |
| Shc1 | 453.0227992 | -0.422882913 | 1.10E-05 |
| Etfa | 1618.227789 | -0.422942624 | 1.28E-07 |
| RT1-M3-1 | 637.7112478 | -0.422973102 | 4.69E-06 |
| Tcirg1 | 500.0247185 | -0.423000565 | 0.000481403 |
| Nqo1 | 826.0860032 | -0.423115873 | 2.71E-05 |
| Slco1c1 | 3635.682807 | -0.423161241 | 0.000453393 |
| Vhl | 2658.226249 | -0.423321987 | 4.59E-07 |
| Cyp7b1 | 4386.282378 | -0.423889671 | 4.22E-05 |
| Galnt1 | 3047.517246 | -0.423965601 | 5.83E-07 |
| Gm2a | 3428.279821 | -0.424174784 | 9.23E-05 |
| Csrp1 | 12173.6077 | -0.424612193 | 3.15E-06 |
| Cpm | 218.4631179 | -0.425161236 | 0.01833544 |
| Mtmr10 | 2101.176312 | -0.425174865 | 1.68E-06 |
| Trip10 | 442.6337796 | -0.425336967 | 0.001073558 |
| Ramp2 | 148.965194 | -0.425358383 | 0.006524116 |
| Ctnna1 | 7844.7921 | -0.425710656 | 8.94E-06 |
| Capsl | 113.7013664 | -0.425866704 | 0.022814445 |
| Cox4i2 | 24.17275009 | -0.425901633 | 0.02023835 |
| Lmo1 | 398.4845349 | -0.425984857 | 0.004088175 |
| Fgfr2 | 9387.871523 | -0.426072608 | 1.16E-07 |
| Tril | 25800.9044 | -0.426669199 | 3.83E-05 |
| Mtmr11 | 258.6030884 | -0.426680194 | 0.00159979 |
| Prtfdc1 | 1360.622003 | -0.427274781 | 9.30E-06 |
| Cdc42bpg | 621.6915 | -0.427403014 | 0.000555246 |
| Slc44a1 | 2513.324765 | -0.427584857 | 0.002582529 |
| Tst | 6155.072637 | -0.428121695 | 1.56E-05 |
| Reep3 | 3060.940017 | -0.42818251 | 3.05E-06 |
| Enpp6 | 34.60236515 | -0.428248138 | 0.022220245 |
| Rassf2 | 2928.941765 | -0.428466794 | 4.58E-05 |
| Sesn2 | 1349.767074 | -0.428667167 | 0.000310766 |

| | | | |
|---|---|---|---|
| Fyco1 | 1167.188363 | -0.428964686 | 3.15E-05 |
| Pskh1 | 68.19930852 | -0.428969329 | 0.008725978 |
| Sft2d2 | 408.4931485 | -0.429420763 | 0.00388751 |
| Txnip | 1888.587997 | -0.429498242 | 0.014663328 |
| Rilp | 152.8931872 | -0.429507722 | 0.008373181 |
| Ssfa2 | 4936.849146 | -0.42983173 | 1.12E-06 |
| Apoc4 | 39.27280925 | -0.429995717 | 0.01864702 |
| Tp53 | 881.858575 | -0.430107973 | 2.20E-07 |
| Ccrl2 | 6.919953053 | -0.430824842 | 0.02294165 |
| Ttll10 | 15.3254725 | -0.431561218 | 0.022815865 |
| Piezo1 | 255.6077099 | -0.431808525 | 0.002249202 |
| Sdc4 | 13336.61149 | -0.432267472 | 3.98E-06 |
| Acsm3 | 152.5314757 | -0.432328482 | 0.018707905 |
| Slc44a2 | 2186.883957 | -0.432541822 | 0.000146306 |
| Tshr | 191.4292831 | -0.433081731 | 0.018787693 |
| Sp1 | 2723.582459 | -0.433287416 | 4.46E-08 |
| Rrm2 | 36.9734951 | -0.433761128 | 0.018218096 |
| Wdr34 | 516.6575192 | -0.434556681 | 4.49E-05 |
| Ctbp2 | 715.3755562 | -0.434568641 | 0.000142192 |
| Rhog | 570.6699713 | -0.43458641 | 0.002084487 |
| Stxbp4 | 506.8697477 | -0.434693174 | 0.000384356 |
| Ppic | 74.78665036 | -0.434951892 | 0.013085346 |
| Bmp1 | 1405.401422 | -0.435111604 | 0.000184699 |
| Hsd17b4 | 4036.144422 | -0.435238155 | 1.03E-06 |
| Tnfsf12 | 665.0546806 | -0.435259217 | 9.13E-06 |
| Irx2 | 5.592961108 | -0.435438025 | 0.016522085 |
| Gas2l2 | 56.12609247 | -0.436057318 | 0.021586672 |
| Tcn2 | 1740.052526 | -0.436475043 | 0.000428909 |
| Zfp641 | 225.8370718 | -0.436686749 | 0.000410402 |
| Hoga1 | 369.0171913 | -0.437038044 | 0.001301906 |
| Acsf2 | 6068.15246 | -0.437119562 | 3.07E-05 |
| Cd22 | 4.966389167 | -0.437199353 | 0.020113818 |
| Dusp16 | 285.708591 | -0.438061467 | 3.22E-05 |
| Bcl2 | 356.7839392 | -0.438691079 | 7.95E-05 |
| Folh1 | 3546.052712 | -0.439146178 | 0.000185068 |
| Gja1 | 89969.78872 | -0.439146222 | 9.93E-06 |

| | | | |
|---|---|---|---|
| RGD1311756 | 661.6110016 | -0.439160752 | 0.000156547 |
| Ppp1r36 | 616.030075 | -0.439390404 | 0.000455138 |
| Ifi27 | 2899.841378 | -0.439499665 | 0.006804616 |
| Gamt | 2050.961722 | -0.439844783 | 1.62E-05 |
| Hes5 | 696.6405974 | -0.440100801 | 0.001577128 |
| Tk1 | 369.6705809 | -0.440211624 | 0.00293646 |
| Hn1l | 222.4187172 | -0.440340413 | 0.002083283 |
| Psat1 | 14462.94705 | -0.440440083 | 2.23E-05 |
| Afmid | 833.0202684 | -0.440798682 | 0.000869597 |
| C4a | 929.708177 | -0.440994096 | 0.02023835 |
| Has2 | 8.465684225 | -0.441542145 | 0.021713001 |
| Tmem198b | 200.0885558 | -0.441550791 | 0.00028409 |
| Ezh2 | 76.61519606 | -0.44211596 | 0.005193249 |
| RGD1559896 | 20189.25433 | -0.442208807 | 1.76E-05 |
| Itpr2 | 3573.302694 | -0.442299518 | 0.000129342 |
| Rreb1 | 844.4967374 | -0.44246568 | 2.56E-05 |
| Cecr2 | 521.5293075 | -0.442551877 | 2.71E-05 |
| RGD1560672 | 184.1520066 | -0.443107261 | 0.015437338 |
| Paqr7 | 2887.590797 | -0.44319809 | 1.18E-05 |
| Pvrl2 | 411.0021133 | -0.443287716 | 1.52E-05 |
| Rfx2 | 449.9943479 | -0.443360675 | 0.003193465 |
| Sntb1 | 2520.325215 | -0.443515221 | 9.21E-06 |
| Sgpl1 | 803.7625561 | -0.443527596 | 1.03E-06 |
| Zbtb20 | 13117.50435 | -0.443581872 | 6.25E-07 |
| Pttg1ip | 5237.679594 | -0.443796831 | 1.27E-08 |
| Fnbp1 | 5184.67989 | -0.443920634 | 8.49E-07 |
| Sult1d1 | 1478.658388 | -0.444110927 | 0.000611912 |
| Rasgrp3 | 138.1027666 | -0.444161266 | 0.016627016 |
| Glb1l | 560.1058979 | -0.444455041 | 0.000156798 |
| Nrarp | 1419.119315 | -0.444637028 | 0.000388042 |
| Mettl20 | 136.4122248 | -0.445160484 | 0.005648352 |
| Cbs | 9077.637931 | -0.445511552 | 0.000183267 |
| Abcc4 | 587.3317986 | -0.445779939 | 7.52E-05 |
| Lamb2 | 3267.872226 | -0.446002874 | 0.000178118 |
| Kank2 | 1383.546829 | -0.446425058 | 4.07E-06 |
| Zhx2 | 2336.145303 | -0.446655166 | 1.68E-06 |

| | | | |
|---|---|---|---|
| Acss1 | 6995.944821 | -0.447368597 | 2.30E-05 |
| Lnc016 | 231.883173 | -0.447590412 | 0.018721323 |
| Slc25a21 | 90.82706696 | -0.447612676 | 0.009606499 |
| Ezr | 11152.30683 | -0.447744454 | 3.65E-05 |
| Mob3a | 548.3645384 | -0.447793968 | 7.88E-07 |
| Psme2 | 694.3547442 | -0.447851505 | 0.000338385 |
| Cyp2j3 | 1988.947992 | -0.447943608 | 0.000201158 |
| Il17rd | 357.4589312 | -0.448071729 | 0.000200187 |
| Gusb | 290.5177464 | -0.448256965 | 0.001030091 |
| Suclg2 | 5934.718522 | -0.448327786 | 1.10E-05 |
| Lpar4 | 398.4475835 | -0.44833751 | 0.000514865 |
| Phgdh | 10279.75072 | -0.448468199 | 3.11E-05 |
| Dhrs11 | 192.9042924 | -0.448510751 | 0.006387688 |
| Timp3 | 3692.671158 | -0.449134285 | 3.92E-06 |
| Il10rb | 123.9957083 | -0.449164294 | 0.002341243 |
| Mlc1 | 47147.37233 | -0.449407902 | 2.95E-06 |
| Clu | 93581.02888 | -0.449470051 | 2.23E-05 |
| Guca2b | 3.215747883 | -0.450394862 | 0.01759164 |
| Evc | 766.5800332 | -0.45071973 | 1.38E-09 |
| Gsta4 | 653.6883489 | -0.45090002 | 0.00053661 |
| Tsku | 250.2577005 | -0.451214726 | 0.001764536 |
| Limd1 | 772.6407384 | -0.452030598 | 2.08E-05 |
| Susd3 | 5.846717663 | -0.452382006 | 0.020461489 |
| Itgb1 | 7238.609621 | -0.452503841 | 7.03E-06 |
| Jak3 | 209.7068175 | -0.452515337 | 0.003672475 |
| Rmrp | 1539.362057 | -0.452586765 | 0.020164923 |
| Slc4a5 | 19.17895616 | -0.452587686 | 0.0159132 |
| Maob | 9216.594951 | -0.453064451 | 4.85E-05 |
| Phyh | 2694.727476 | -0.453200707 | 7.85E-06 |
| LOC361346 | 131.402065 | -0.453255128 | 0.002781383 |
| Serpinf1 | 36.4424994 | -0.453344838 | 0.018426047 |
| Tnfrsf12a | 57.0404391 | -0.453488153 | 0.009955071 |
| Lrguk | 88.06738368 | -0.454169576 | 0.014552916 |
| Dock1 | 5442.139043 | -0.454200376 | 5.91E-07 |
| Atp1b2 | 66243.06351 | -0.454834042 | 4.04E-06 |
| Sdc1 | 54.1647732 | -0.454837926 | 0.009579433 |

| | | | |
|---|---|---|---|
| Myo10 | 11883.09162 | -0.455166435 | 3.05E-06 |
| Wipf1 | 1040.781575 | -0.455329211 | 0.0002548 |
| Plxnb2 | 9688.060073 | -0.455469938 | 4.23E-07 |
| Lrrc36 | 24.14195028 | -0.455762198 | 0.016935195 |
| Hapln3 | 85.79036335 | -0.456174106 | 0.011188562 |
| Mpp6 | 2667.109708 | -0.456195839 | 0.001769254 |
| Isoc1 | 1844.971947 | -0.456260195 | 0.000554569 |
| Plce1 | 945.1458233 | -0.456504662 | 0.000943548 |
| Cmtm5 | 2424.656975 | -0.456650007 | 1.75E-07 |
| Nsmce1 | 593.3685298 | -0.456650538 | 1.29E-06 |
| Grhpr | 1045.117133 | -0.456789502 | 5.89E-07 |
| Gal3st1 | 79.23346023 | -0.457001837 | 0.008659298 |
| Tmcc3 | 1880.75727 | -0.457037744 | 3.39E-05 |
| Tmem173 | 4.331622857 | -0.457317401 | 0.019597532 |
| Trpm3 | 6601.835157 | -0.458132994 | 2.07E-05 |
| Cd24 | 4797.038552 | -0.459071616 | 1.30E-05 |
| Cml1 | 173.9223444 | -0.459319601 | 0.003583445 |
| Mfap2 | 69.32023766 | -0.459741596 | 0.010525674 |
| Traf3ip2 | 63.11228337 | -0.459788427 | 0.009730327 |
| Tcf7l1 | 710.6896565 | -0.459846775 | 9.98E-06 |
| Mmp15 | 4733.454929 | -0.460585309 | 2.36E-05 |
| Mybl2 | 6.68003239 | -0.460843983 | 0.0194991 |
| LOC654482 | 132.4910439 | -0.460998751 | 0.005808675 |
| RT1-Da | 54.40235816 | -0.461521803 | 0.017887966 |
| Pdgfc | 614.8920782 | -0.461721635 | 8.25E-05 |
| Cd81 | 35875.91538 | -0.461747723 | 6.27E-08 |
| Rlbp1 | 1247.81714 | -0.463294681 | 0.00104872 |
| Cd302 | 848.196751 | -0.46386572 | 3.30E-05 |
| Crip1 | 127.6415647 | -0.463962619 | 0.014582776 |
| Sp5 | 52.98250034 | -0.464338062 | 0.013523263 |
| Ddr1 | 5222.758705 | -0.464531183 | 3.66E-07 |
| Bfsp1 | 30.04791009 | -0.464657381 | 0.013388437 |
| Hey2 | 1129.808021 | -0.464690575 | 1.05E-05 |
| Cib1 | 654.8187355 | -0.464794896 | 5.29E-05 |
| Cldn11 | 1499.759943 | -0.465646381 | 0.016047251 |
| Ppp2r1b | 464.4559077 | -0.4661707 | 7.88E-07 |

| | | | |
|---|---|---|---|
| Zfp36l2 | 7738.71304 | -0.466460108 | 0.000102754 |
| Ctsh | 1916.217058 | -0.466727157 | 3.64E-06 |
| Sh3glb1 | 2584.293992 | -0.467179109 | 3.49E-09 |
| Ednra | 409.7282976 | -0.467250402 | 0.011850784 |
| Rcbtb2 | 620.7126204 | -0.467516033 | 1.75E-05 |
| Fxyd3 | 45.7439467 | -0.467708289 | 0.008910998 |
| Rbp1 | 1199.875763 | -0.467933484 | 0.003500737 |
| Kif1c | 4192.612331 | -0.468041921 | 2.43E-08 |
| Steap3 | 1249.043917 | -0.468089362 | 4.85E-05 |
| Atf5 | 1160.061549 | -0.468256967 | 3.42E-06 |
| Spata6 | 450.3055264 | -0.468284375 | 9.76E-05 |
| S100a16 | 6491.291795 | -0.468585151 | 8.21E-06 |
| Flna | 1587.588121 | -0.468766009 | 0.008393423 |
| Fbln1 | 2879.837856 | -0.468994067 | 0.000103964 |
| Eno4 | 67.41993271 | -0.46939813 | 0.013374456 |
| Lamp2 | 5006.670922 | -0.469661752 | 1.47E-07 |
| Ampd3 | 4620.931032 | -0.470360468 | 2.81E-08 |
| Fgfbp3 | 459.8629817 | -0.470518591 | 0.000714909 |
| Efs | 1254.263748 | -0.470785478 | 3.51E-06 |
| Mif4gd | 221.6977143 | -0.471400952 | 8.06E-05 |
| Entpd2 | 3782.770292 | -0.471454163 | 3.73E-05 |
| Agt | 7366.181505 | -0.472472976 | 0.012679829 |
| Timp4 | 1308.209388 | -0.47289481 | 0.000896775 |
| Lmf2 | 1235.538582 | -0.473115034 | 1.67E-08 |
| Cdc14a | 461.1565018 | -0.473377167 | 0.00025656 |
| Mis18bp1 | 13.69831658 | -0.473723783 | 0.016247463 |
| Col11a1 | 252.2275329 | -0.474017149 | 0.013085346 |
| Cenpf | 40.87962341 | -0.474077699 | 0.013038173 |
| Slc13a5 | 1405.558555 | -0.474083802 | 0.00028311 |
| Tp73 | 151.0995917 | -0.474228587 | 0.016247463 |
| Mdk | 311.7035593 | -0.474964136 | 0.001288571 |
| Vgll4 | 882.3103187 | -0.475140168 | 1.72E-06 |
| Elf1 | 252.2870211 | -0.47520754 | 0.000317929 |
| Ttc7a | 846.2384416 | -0.475234313 | 6.13E-05 |
| Slc4a4 | 17889.79408 | -0.475548584 | 0.000173516 |
| Slc43a3 | 152.3049725 | -0.47580473 | 0.004108651 |

| | | | |
|---|---|---|---|
| Rad9b | 62.52505888 | -0.47623429 | 0.009758112 |
| Sall1 | 2240.412708 | -0.476459535 | 4.19E-07 |
| Arhgef1 | 921.9915428 | -0.476545259 | 2.81E-08 |
| Arhgef40 | 1527.458082 | -0.476577271 | 1.42E-06 |
| Tlr3 | 350.2462776 | -0.477253046 | 0.000353773 |
| Llgl1 | 4590.297729 | -0.477378238 | 2.08E-07 |
| Fosl1 | 8.473597968 | -0.477645561 | 0.018067553 |
| Tspan6 | 1042.567413 | -0.477732261 | 8.59E-08 |
| Cand2 | 1104.701081 | -0.478014966 | 4.07E-05 |
| Kif13b | 1013.558127 | -0.478782518 | 4.09E-05 |
| Cald1 | 207.9218882 | -0.479116886 | 0.001044913 |
| Apol3 | 20.75104388 | -0.479643138 | 0.018072988 |
| Parvb | 4701.995662 | -0.480395877 | 7.40E-06 |
| Stat3 | 4932.472732 | -0.480670352 | 1.34E-05 |
| Lrp10 | 2675.843689 | -0.480823626 | 1.64E-05 |
| Tmem117 | 355.8937654 | -0.48095057 | 0.000623578 |
| Slc16a1 | 2439.756575 | -0.481202499 | 2.80E-05 |
| Cpne3 | 417.1599451 | -0.481754908 | 8.61E-05 |
| Scnn1a | 130.8892682 | -0.481895787 | 0.005087116 |
| Gng12 | 409.281791 | -0.481963425 | 3.12E-05 |
| Folr1 | 38.76012421 | -0.482285732 | 0.016812998 |
| Nfkb2 | 170.2381366 | -0.482464901 | 0.002473752 |
| Kcnk10 | 172.5817679 | -0.483078962 | 0.001324581 |
| Frmd8 | 1288.375719 | -0.483476548 | 5.05E-08 |
| Fkbp10 | 601.5341517 | -0.483617954 | 0.000179012 |
| Cd82 | 1237.43112 | -0.484384198 | 3.32E-05 |
| RGD1565033 | 405.042309 | -0.484972594 | 0.000169523 |
| Lrrc2 | 478.2482605 | -0.485003826 | 5.32E-05 |
| Dchs1 | 5576.58735 | -0.485223927 | 1.20E-05 |
| Sox9 | 17051.75019 | -0.485934705 | 3.90E-07 |
| Col4a1 | 227.4618417 | -0.486353884 | 0.000244894 |
| Sspn | 2183.601902 | -0.486582359 | 7.39E-08 |
| Plekhg1 | 1090.489254 | -0.48672731 | 7.07E-06 |
| Bcam | 600.0782458 | -0.486985107 | 0.007716967 |
| Mpst | 705.8209695 | -0.487215034 | 4.65E-06 |
| Ubxn10 | 93.09273953 | -0.487465147 | 0.012471676 |

| | | | |
|---|---|---|---|
| Mgst1 | 1089.43695 | -0.487584655 | 0.000512256 |
| Arhgap18 | 470.1646112 | -0.488703792 | 0.000116026 |
| Bgn | 384.8191134 | -0.488707071 | 0.00281999 |
| Nuak2 | 320.2440533 | -0.488974338 | 0.000216909 |
| Tspan15 | 217.8890507 | -0.488977087 | 0.001797855 |
| Abtb2 | 390.773635 | -0.489132703 | 0.002301046 |
| Casp7 | 277.3674039 | -0.490294913 | 4.40E-05 |
| Map4k4 | 7280.255639 | -0.490340722 | 1.09E-08 |
| Tmem179b | 68.74412905 | -0.490556155 | 0.003718108 |
| Mthfs | 368.2062173 | -0.490735756 | 6.01E-05 |
| Pdlim5 | 2688.381059 | -0.49108053 | 3.70E-05 |
| Cxcr4 | 105.1404602 | -0.491080957 | 0.008692577 |
| Gstk1 | 594.2607328 | -0.491206497 | 1.45E-06 |
| Sh3bgr | 248.6095746 | -0.491352557 | 0.00052861 |
| Gpr165 | 1527.460957 | -0.491525087 | 0.011103981 |
| Wwtr1 | 1031.796903 | -0.492170747 | 5.07E-05 |
| Slc7a11 | 17094.3284 | -0.493398642 | 1.11E-06 |
| Abcc3 | 59.36812111 | -0.493887452 | 0.010086871 |
| Socs3 | 30.49947283 | -0.494268604 | 0.015229178 |
| Dnaaf5 | 580.9926687 | -0.494359899 | 1.45E-05 |
| Vcl | 3592.756027 | -0.494588185 | 7.25E-05 |
| Cyba | 157.6699057 | -0.494935118 | 0.006618299 |
| Pard3b | 568.3047953 | -0.495231504 | 4.28E-05 |
| Slc39a12 | 35.11170442 | -0.495635101 | 0.005635915 |
| Srsf9 | 331.7428723 | -0.495681551 | 4.19E-05 |
| Rft1 | 353.6478465 | -0.496166768 | 4.06E-07 |
| Ptprz1 | 32744.90204 | -0.496360559 | 1.10E-07 |
| Scamp2 | 793.9273297 | -0.496396431 | 3.49E-07 |
| P2rx6 | 178.7389722 | -0.49652441 | 0.008557968 |
| Tjp2 | 3166.849355 | -0.496653404 | 7.97E-05 |
| Amt | 869.5873217 | -0.496702833 | 4.31E-06 |
| Elovl1 | 342.9942567 | -0.496725927 | 0.000456997 |
| Car2 | 2551.179414 | -0.497223005 | 0.000114082 |
| Atf3 | 24.1250456 | -0.497625313 | 0.014458901 |
| Notch3 | 5283.967277 | -0.498913552 | 0.000191891 |
| Gsn | 4124.927984 | -0.499569801 | 0.008760806 |

| | | | |
|---|---|---|---|
| Tmbim1 | 1136.66556 | -0.500222655 | 0.000320745 |
| Cmpk2 | 1416.576265 | -0.502023794 | 0.000182695 |
| Acy1 | 1640.67699 | -0.502207827 | 2.07E-05 |
| Tax1bp3 | 209.8908399 | -0.502272313 | 8.39E-05 |
| Tep1 | 1276.93868 | -0.502439622 | 3.60E-07 |
| Rps27l | 754.3182657 | -0.502560887 | 3.71E-05 |
| Wscd1 | 3782.178407 | -0.502923853 | 4.52E-06 |
| Nat6 | 1969.902647 | -0.502932255 | 9.75E-07 |
| Aqp4 | 106463.7046 | -0.503022367 | 2.98E-05 |
| Carhsp1 | 1504.833419 | -0.503158742 | 3.06E-05 |
| Adamtsl4 | 96.10590708 | -0.503274968 | 0.007170798 |
| Rgma | 12291.19053 | -0.503394014 | 1.48E-09 |
| Cd99 | 411.2074425 | -0.503585172 | 0.000108891 |
| Angpt1 | 123.590294 | -0.503725657 | 0.001050824 |
| Plcd1 | 1996.567441 | -0.503760197 | 3.99E-06 |
| Mtss1l | 28476.84832 | -0.503889615 | 1.12E-07 |
| Gpam | 10861.18577 | -0.504749828 | 8.70E-07 |
| LOC691909 | 136.8260503 | -0.504751472 | 0.002689621 |
| Snx33 | 837.3913042 | -0.505124841 | 1.90E-06 |
| Itpkb | 3044.175357 | -0.50664849 | 9.46E-07 |
| Pnpla7 | 2580.091953 | -0.506774905 | 9.29E-06 |
| Ckap2 | 73.95254083 | -0.506804333 | 0.006963741 |
| Fa2h | 272.2415409 | -0.50691908 | 0.010783726 |
| Pih1d2 | 73.79089382 | -0.507010758 | 0.008629629 |
| Lrp5 | 1248.511408 | -0.507576766 | 3.22E-05 |
| Pbxip1 | 4816.096643 | -0.507782237 | 5.94E-06 |
| Erbb2 | 414.9434148 | -0.50801611 | 3.48E-05 |
| Prelp | 3249.869636 | -0.508046322 | 3.79E-06 |
| Slc12a2 | 3683.255614 | -0.508310328 | 5.71E-07 |
| Fkbp9 | 1490.992205 | -0.508518316 | 2.40E-06 |
| Lrrc66 | 210.9131336 | -0.508711356 | 0.00406627 |
| Atp13a4 | 1025.856825 | -0.508990136 | 0.000250651 |
| Gsta1 | 11748.85822 | -0.509199909 | 5.05E-07 |
| Adamts4 | 212.0055621 | -0.509582816 | 0.009913523 |
| Notch2 | 6599.437697 | -0.509789868 | 3.75E-07 |
| Rsu1 | 1431.143086 | -0.509807672 | 2.38E-07 |

| | | | |
|---|---|---|---|
| Gabre | 112.0778506 | -0.509846103 | 0.00849962 |
| Npepl1 | 633.7396604 | -0.510744024 | 2.53E-07 |
| Tns3 | 11999.73117 | -0.510808487 | 3.51E-06 |
| Arhgap17 | 306.1390639 | -0.511200791 | 1.46E-05 |
| Vangl2 | 731.6511745 | -0.511286832 | 1.92E-05 |
| Rras | 635.4314261 | -0.51151599 | 3.17E-05 |
| Col27a1 | 192.900801 | -0.511521232 | 8.27E-05 |
| Sumf2 | 562.864275 | -0.512586365 | 1.03E-06 |
| Tead2 | 39.44131859 | -0.513076306 | 0.009111841 |
| Cst3 | 77756.20775 | -0.513233094 | 2.01E-08 |
| Lrrk1 | 819.1421294 | -0.513409451 | 8.72E-06 |
| Serpinb9 | 479.0063995 | -0.513416859 | 2.30E-05 |
| Smo | 1016.472986 | -0.513818 | 9.81E-07 |
| Pald1 | 309.3259631 | -0.514386166 | 0.00052842 |
| Cyp4v3 | 1849.653861 | -0.515434954 | 2.63E-06 |
| Dynlt1 | 209.617856 | -0.515663193 | 0.000169074 |
| Myl12a | 832.4890543 | -0.516057788 | 4.76E-08 |
| Nek6 | 1675.441323 | -0.516266861 | 1.21E-06 |
| Fam19a4 | 73.04020892 | -0.517356342 | 0.010856956 |
| Vwa5b1 | 209.4966782 | -0.517660923 | 0.009231605 |
| Ifit2 | 179.6104371 | -0.518496868 | 0.010149865 |
| Tapbp | 914.6091208 | -0.518548599 | 4.30E-05 |
| Ribc1 | 177.0059885 | -0.519104075 | 0.000351327 |
| Olfml3 | 350.8873701 | -0.519502693 | 8.96E-05 |
| Tubb2b | 10383.66966 | -0.52002731 | 1.38E-07 |
| Necap2 | 1251.678027 | -0.520304135 | 4.46E-08 |
| RGD1306739 | 1266.967405 | -0.520580705 | 6.25E-07 |
| Ripk1 | 548.1394859 | -0.520782453 | 1.47E-06 |
| Tpm4 | 189.3038256 | -0.520792613 | 0.000508867 |
| Lpar1 | 479.3004865 | -0.520804691 | 0.00601342 |
| Crb2 | 179.6688853 | -0.520985858 | 0.004958142 |
| Gstt2 | 290.1709858 | -0.521024636 | 9.30E-06 |
| Ptch1 | 5230.57579 | -0.521842963 | 8.61E-05 |
| Ssc5d | 428.3819873 | -0.522235846 | 0.003073519 |
| Unc5b | 2086.75982 | -0.522643749 | 8.84E-07 |
| Sox2 | 6581.346377 | -0.524160968 | 8.78E-07 |

| | | | |
|---|---|---|---|
| Trim34 | 261.8564576 | -0.524619212 | 0.000890691 |
| Nfatc1 | 884.7850598 | -0.525685347 | 3.65E-06 |
| Sipa1 | 84.01795011 | -0.525934367 | 0.003959025 |
| Itga7 | 1008.481077 | -0.526709656 | 0.00021015 |
| Clic1 | 623.2728679 | -0.527320102 | 5.63E-05 |
| Tgm2 | 315.1025078 | -0.527458614 | 0.011885047 |
| Rom1 | 268.2717937 | -0.527928284 | 0.000117954 |
| Ptpn22 | 40.72531127 | -0.527934097 | 0.00601342 |
| Dap | 462.2241594 | -0.52803475 | 1.60E-06 |
| Col5a2 | 1360.245996 | -0.528652363 | 0.000403614 |
| P2rx7 | 431.5688834 | -0.5287059 | 0.000329521 |
| Rnaset2 | 844.834176 | -0.529163899 | 9.17E-07 |
| Gbx2 | 188.9600686 | -0.529274486 | 0.005926295 |
| Acot2 | 787.7921546 | -0.529375279 | 8.06E-06 |
| Pnp | 5267.107154 | -0.530533437 | 6.46E-07 |
| Fam167a | 248.998344 | -0.530690437 | 0.00079829 |
| Ptprc | 12.817485 | -0.531283716 | 0.011924599 |
| Gli1 | 532.5602408 | -0.531569541 | 0.001572427 |
| Cnn3 | 11782.52097 | -0.531793364 | 8.81E-06 |
| Tmod3 | 618.8195737 | -0.532072419 | 1.37E-05 |
| Cd63 | 2704.85794 | -0.532649681 | 1.43E-09 |
| RGD1309028 | 46.3349256 | -0.532758656 | 0.006921751 |
| Sp140 | 29.81562652 | -0.532830486 | 0.009751217 |
| Kcng4 | 1452.794843 | -0.532929871 | 0.001513505 |
| Gucy2e | 5.005669487 | -0.533164676 | 0.014260868 |
| Il13ra1 | 373.8143624 | -0.533436485 | 0.002173133 |
| Mcm3 | 42.54832909 | -0.533512414 | 0.00562611 |
| Sec23b | 1546.317768 | -0.533910079 | 2.53E-07 |
| Grin2c | 7065.786017 | -0.534041233 | 2.62E-06 |
| Efhd1 | 3604.348329 | -0.534254226 | 2.28E-07 |
| Hps5 | 470.0809236 | -0.534714694 | 4.77E-06 |
| Dapp1 | 39.33194442 | -0.534844825 | 0.006516876 |
| Itgb8 | 2933.323295 | -0.535365463 | 9.23E-08 |
| Cyp2t1 | 48.83645346 | -0.53722963 | 0.003889956 |
| Ggta1 | 294.3107629 | -0.538413542 | 0.002143443 |
| Vamp3 | 1693.123414 | -0.539082434 | 1.95E-10 |

| | | | |
|---|---|---|---|
| Srebf1 | 12221.83697 | -0.539671944 | 2.04E-07 |
| Stk10 | 262.5928627 | -0.540252976 | 2.67E-06 |
| Myo1e | 821.2089491 | -0.540611824 | 4.32E-06 |
| Ston1 | 628.4699007 | -0.540837748 | 3.18E-05 |
| Pth1r | 297.8968999 | -0.541994197 | 0.000183668 |
| Slc6a11 | 9124.794931 | -0.542325925 | 0.000406438 |
| Prkd3 | 1942.757007 | -0.542797171 | 4.91E-08 |
| Lgi4 | 8258.406055 | -0.543927864 | 1.29E-09 |
| Mov10 | 177.6883894 | -0.544437538 | 0.000446907 |
| Gpr37l1 | 28173.26571 | -0.544653009 | 4.86E-08 |
| Vstm4 | 260.4229436 | -0.544763782 | 0.00024771 |
| Efemp2 | 262.3415731 | -0.545094818 | 6.57E-05 |
| Sec14l4 | 10.85150671 | -0.545542576 | 0.012333687 |
| Lrrc51 | 45.57830279 | -0.545753704 | 0.008588662 |
| Gpr37 | 4629.437518 | -0.545768009 | 1.11E-06 |
| Caskin2 | 3354.258733 | -0.546866965 | 2.70E-06 |
| Spef2 | 92.95359835 | -0.547343965 | 0.007684397 |
| Lap3 | 2192.750632 | -0.547428082 | 6.13E-06 |
| Slc2a10 | 840.58994 | -0.54783326 | 4.99E-07 |
| Mxra8 | 667.3918658 | -0.548801427 | 0.000296799 |
| Hells | 75.84583028 | -0.549070897 | 0.000840224 |
| Tspan18 | 300.2911695 | -0.55125618 | 0.006164732 |
| Cdc20 | 33.79716384 | -0.551366823 | 0.004589557 |
| Lyn | 158.9453914 | -0.552007167 | 0.00272762 |
| Crlf1 | 3172.444268 | -0.552382616 | 0.003754327 |
| Slc39a1 | 3969.692471 | -0.553453226 | 4.95E-09 |
| Adgre5 | 162.2082802 | -0.553561632 | 0.000996966 |
| Clic4 | 756.8439883 | -0.554769081 | 9.66E-07 |
| Ephx1 | 4112.309551 | -0.554775851 | 1.70E-06 |
| Prodh | 3383.721825 | -0.555143568 | 9.91E-06 |
| Otx2 | 158.8597997 | -0.555577705 | 0.002432611 |
| Nod1 | 659.5054084 | -0.556980671 | 5.08E-06 |
| Ankrd44 | 270.3151937 | -0.557326322 | 3.35E-06 |
| Plek | 14.73041719 | -0.557645038 | 0.010409711 |
| Sat1 | 2477.251845 | -0.558104091 | 7.63E-06 |
| Trip6 | 222.4276306 | -0.558150129 | 0.000151032 |

| | | | |
|---|---|---|---|
| Itgb5 | 4287.66776 | -0.558331069 | 8.30E-08 |
| Slc1a4 | 2781.384411 | -0.558401307 | 7.81E-08 |
| Arpc1b | 81.07301692 | -0.559464109 | 0.003520431 |
| Efhc1 | 310.4977609 | -0.560008683 | 0.001117776 |
| Plin2 | 602.0647883 | -0.56045798 | 4.88E-06 |
| Stom | 86.70890207 | -0.56060622 | 0.004057149 |
| Tapbpl | 741.979078 | -0.561001867 | 9.17E-07 |
| Dnai1 | 192.4414507 | -0.561694637 | 0.002646891 |
| Serpina11 | 28.56262858 | -0.562005073 | 0.00379248 |
| RGD1311251 | 64.20379471 | -0.562359329 | 0.003295064 |
| Slc12a4 | 1195.692404 | -0.562725011 | 6.18E-07 |
| Arhgef19 | 1839.71245 | -0.562923406 | 2.76E-07 |
| Lhfpl2 | 1674.534665 | -0.563265206 | 8.30E-08 |
| Apobec1 | 68.26064289 | -0.563466937 | 0.002582768 |
| Gab1 | 2908.948192 | -0.56349863 | 1.33E-07 |
| Akna | 951.9472931 | -0.565905022 | 3.01E-06 |
| Plxnb3 | 591.4798738 | -0.566668765 | 0.000341805 |
| Stat5a | 255.6668511 | -0.566881943 | 1.67E-05 |
| Tcf7l2 | 1037.394766 | -0.567841067 | 1.03E-06 |
| Stx2 | 98.06130709 | -0.568255604 | 0.000493807 |
| Ube2l6 | 157.7831105 | -0.56994475 | 0.003457968 |
| Nt5dc2 | 817.1926577 | -0.570089041 | 1.16E-07 |
| Ednrb | 7514.002115 | -0.570810127 | 5.02E-06 |
| Tmem81 | 37.32684472 | -0.571294179 | 0.005044796 |
| Dcakd | 2747.875338 | -0.571637511 | 1.23E-07 |
| Mvp | 804.9140651 | -0.572281511 | 1.40E-05 |
| Slc38a3 | 6305.619903 | -0.57375892 | 9.51E-07 |
| Pla2g16 | 2463.479733 | -0.574749619 | 8.05E-09 |
| Crh | 71.47552566 | -0.575048936 | 0.005726193 |
| Gfap | 82659.47166 | -0.575075266 | 0.004083088 |
| Pla2g4a | 65.42568689 | -0.579127949 | 0.002203946 |
| Fxyd1 | 2698.905565 | -0.579137738 | 9.00E-07 |
| Rhoc | 1064.423487 | -0.580653029 | 3.57E-07 |
| Adam17 | 1167.5571 | -0.580957061 | 4.80E-08 |
| Slc29a3 | 339.0336765 | -0.581195145 | 3.17E-05 |
| Hcls1 | 43.3018634 | -0.582709361 | 0.001687461 |

| | | | |
|---|---|---|---|
| Lpcat3 | 1169.410963 | -0.582854459 | 1.77E-08 |
| Rest | 476.5493183 | -0.583776299 | 1.15E-07 |
| Ltbr | 1277.382552 | -0.58379922 | 3.49E-08 |
| Cldn9 | 185.5408474 | -0.584069561 | 0.002549755 |
| Dbi | 5804.038134 | -0.584325213 | 5.47E-10 |
| Axl | 3179.484088 | -0.584660806 | 1.29E-07 |
| Gtse1 | 677.3664389 | -0.584933779 | 3.17E-05 |
| Icam1 | 182.8345173 | -0.58503819 | 0.000463817 |
| Slc12a7 | 396.5411469 | -0.586420884 | 0.000146024 |
| Ptafr | 22.80997111 | -0.586625777 | 0.007243474 |
| Nkd1 | 948.5821572 | -0.586995042 | 1.90E-06 |
| Ttc23 | 169.3527895 | -0.58710927 | 1.37E-05 |
| Plcb3 | 1246.076114 | -0.588440873 | 5.85E-07 |
| Rarres1 | 69.63037653 | -0.588715845 | 0.000596735 |
| Ifih1 | 392.7557854 | -0.589959528 | 4.22E-05 |
| Tnfrsf19 | 1786.143001 | -0.590448254 | 1.10E-08 |
| Ptpn6 | 22.14698247 | -0.590649024 | 0.004387059 |
| Naprt1 | 1067.361287 | -0.591149316 | 7.33E-06 |
| Cdk6 | 54.12938918 | -0.591156445 | 0.005193249 |
| Jam3 | 1274.942713 | -0.592058925 | 1.97E-09 |
| Plcd4 | 2268.450675 | -0.592230349 | 2.97E-06 |
| Zmynd10 | 280.2767343 | -0.592592699 | 0.006371893 |
| Ncf1 | 25.8900789 | -0.59277906 | 0.007602751 |
| Tekt3 | 56.68665576 | -0.593277126 | 0.005490502 |
| Tmco4 | 397.7580907 | -0.594852912 | 3.42E-05 |
| Ccdc114 | 90.93646077 | -0.595157863 | 0.005365667 |
| Gng11 | 229.3665698 | -0.59535281 | 0.000231118 |
| Ikbip | 476.3631333 | -0.595406128 | 3.20E-06 |
| RT1-A2 | 1708.759972 | -0.595576418 | 0.004274465 |
| Pdlim4 | 6135.242103 | -0.595594576 | 2.48E-07 |
| Sash3 | 7.395496272 | -0.59606795 | 0.009996942 |
| Nde1 | 367.0132278 | -0.596677027 | 2.77E-06 |
| Pros1 | 502.8959342 | -0.596844293 | 3.53E-06 |
| Cpxm1 | 149.5441767 | -0.599582415 | 0.000324456 |
| F11r | 12.18441365 | -0.601895194 | 0.009480013 |
| Cd276 | 465.8795861 | -0.603173796 | 3.23E-09 |

| | | | |
|---|---|---|---|
| RT1-CE5 | 757.76987 | -0.603482649 | 0.002432611 |
| Cdkn1a | 2806.646878 | -0.604094136 | 0.000289423 |
| Slc14a1 | 2342.8251 | -0.604884796 | 1.10E-08 |
| Serpinh1 | 3598.049122 | -0.604893196 | 5.54E-07 |
| Msn | 543.1538519 | -0.60521748 | 0.000207879 |
| Loxl3 | 531.4070261 | -0.605315013 | 1.64E-06 |
| Gsap | 7.862617166 | -0.605548344 | 0.010811911 |
| Cdc42ep1 | 472.3432546 | -0.6057492 | 9.82E-05 |
| Gpx7 | 264.2990072 | -0.606142093 | 1.24E-06 |
| Shc4 | 130.158149 | -0.606280942 | 0.000855507 |
| Igsf10 | 414.8261764 | -0.606467411 | 0.000428616 |
| Psme1 | 1695.035776 | -0.606833452 | 4.92E-07 |
| Mid1ip1 | 5774.53622 | -0.608400881 | 2.47E-11 |
| Litaf | 888.0556787 | -0.609296778 | 2.48E-06 |
| Smoc1 | 297.4489578 | -0.610547297 | 9.65E-05 |
| Abhd4 | 11286.13924 | -0.610909362 | 1.66E-11 |
| Alpl | 765.8623206 | -0.611043355 | 6.75E-06 |
| Tln1 | 3478.754569 | -0.611844975 | 3.12E-09 |
| Cd9 | 1293.456428 | -0.613542919 | 0.002432611 |
| Slc22a4 | 178.9148979 | -0.613970992 | 0.000754563 |
| Mrc2 | 263.5176079 | -0.614301535 | 0.000181089 |
| Slc1a5 | 29.43183764 | -0.61442599 | 0.003602802 |
| Pltp | 2917.38795 | -0.61516166 | 7.01E-06 |
| Slc25a18 | 4037.308355 | -0.615731853 | 2.47E-05 |
| Slc20a2 | 3079.051621 | -0.615908992 | 1.54E-09 |
| Hmgb2 | 349.4388006 | -0.616085865 | 9.57E-06 |
| Fas | 421.6733578 | -0.618721137 | 8.73E-06 |
| Ly86 | 4.747183415 | -0.619253762 | 0.0096283 |
| Mog | 525.3683414 | -0.619423368 | 0.004385796 |
| Adamts1 | 215.9494407 | -0.619525027 | 0.000314555 |
| Aqp9 | 268.858745 | -0.619690796 | 1.23E-05 |
| Tppp3 | 718.5027601 | -0.620774787 | 0.001990617 |
| Ehd2 | 373.5378461 | -0.621040666 | 0.000175464 |
| Slc27a3 | 810.5252823 | -0.624270734 | 2.03E-06 |
| Oasl | 10.71377021 | -0.626330947 | 0.010082659 |
| Abca8a | 820.8891901 | -0.626606763 | 0.001993439 |

| | | | |
|---|---|---|---|
| Opalin | 623.527176 | -0.628003667 | 0.003231294 |
| Lrrc9 | 149.5389425 | -0.628501357 | 0.000446907 |
| Gltp | 759.5495566 | -0.62881349 | 7.66E-08 |
| Plat | 2003.135763 | -0.629277681 | 2.70E-06 |
| Abca1 | 8705.968999 | -0.630125217 | 2.59E-10 |
| Adamts12 | 141.6701027 | -0.630498399 | 9.85E-05 |
| Pdpn | 2316.953251 | -0.630688931 | 3.17E-07 |
| Wnt5a | 202.024105 | -0.632535941 | 2.26E-05 |
| Padi2 | 3759.320409 | -0.632964671 | 0.00027975 |
| Nkx6-2 | 463.7458238 | -0.634536589 | 0.000143341 |
| Nmi | 133.4046105 | -0.635390645 | 8.14E-05 |
| Eif2ak2 | 1866.312346 | -0.635760014 | 9.42E-09 |
| Snx22 | 8.379291546 | -0.635789841 | 0.009820684 |
| Matn1 | 3.295010873 | -0.635900645 | 0.007430668 |
| Heph | 722.6242761 | -0.636307251 | 0.000970705 |
| MGC112715 | 1757.412647 | -0.637524404 | 1.18E-06 |
| B3gnt7 | 82.84139934 | -0.640940088 | 0.001094828 |
| Angptl4 | 229.4005035 | -0.642295195 | 0.000344064 |
| Lcat | 355.0731866 | -0.644782348 | 0.000233778 |
| Slc7a10 | 2380.727897 | -0.645536909 | 3.99E-08 |
| Vamp5 | 6.219364538 | -0.645687192 | 0.007663117 |
| Aldh3a1 | 66.59454992 | -0.646141572 | 0.001112425 |
| Itih3 | 25901.08574 | -0.647586502 | 0.000102876 |
| Col1a2 | 117.3632677 | -0.64805952 | 0.001267362 |
| Phlda3 | 2907.027466 | -0.648145859 | 2.59E-08 |
| Flnc | 1787.574605 | -0.653641911 | 2.86E-05 |
| Osgin1 | 89.72881912 | -0.654396947 | 0.000100046 |
| Ccdc80 | 830.4966556 | -0.655098912 | 9.29E-06 |
| Dock8 | 58.90067189 | -0.65690088 | 0.000668553 |
| Aldh4a1 | 2839.120006 | -0.658556511 | 2.88E-09 |
| RT1-A3 | 193.6382678 | -0.659433422 | 0.006252184 |
| Cdh19 | 28.97744184 | -0.659846427 | 0.003144001 |
| Gpnmb | 751.3300052 | -0.661672347 | 0.00138814 |
| LOC498368 | 283.4045504 | -0.664097595 | 7.45E-07 |
| Uaca | 316.5937942 | -0.66958882 | 2.77E-06 |
| Vcam1 | 7432.72066 | -0.669885061 | 1.27E-09 |

| | | | |
|---|---|---|---|
| Mmp14 | 899.182165 | -0.670008182 | 0.000396194 |
| Trim5 | 552.5492326 | -0.670312997 | 1.48E-07 |
| Kank1 | 1308.560601 | -0.670763129 | 2.70E-06 |
| RT1-S3 | 1850.745017 | -0.671265256 | 0.000904076 |
| RT1-CE15 | 136.0580007 | -0.671338663 | 0.00123185 |
| Creb5 | 233.2058428 | -0.673001002 | 1.56E-06 |
| Dnali1 | 482.4964949 | -0.674017793 | 0.000182068 |
| Kif19 | 346.2422072 | -0.674163315 | 0.000126759 |
| Nkx2-2 | 248.3163541 | -0.675552347 | 2.32E-05 |
| Cnp | 7709.926503 | -0.675553131 | 5.13E-05 |
| Zc3hav1 | 426.098974 | -0.67601074 | 2.80E-05 |
| Slc2a12 | 184.766296 | -0.679588211 | 0.000466715 |
| Rab13 | 549.7925402 | -0.680481429 | 1.68E-06 |
| Adgrg2 | 79.91797478 | -0.683522388 | 0.000425124 |
| Themis2 | 12.34742897 | -0.686502799 | 0.006023329 |
| Capn6 | 62.10289866 | -0.687823148 | 0.001634726 |
| Oas1i | 49.38918096 | -0.688540064 | 0.003910145 |
| Gsdmd | 287.5811716 | -0.690300974 | 1.54E-06 |
| Qprt | 43.75517384 | -0.692849036 | 0.001706387 |
| Igf2 | 61.99556079 | -0.696557465 | 0.001353329 |
| Tmem100 | 1789.388725 | -0.69707884 | 2.96E-07 |
| Zfp36 | 135.7635018 | -0.697524864 | 0.000346905 |
| Apbb1ip | 73.60933608 | -0.699316754 | 0.000202309 |
| Npr1 | 518.3472762 | -0.700919293 | 8.60E-06 |
| Eln | 273.5725628 | -0.701209518 | 1.08E-06 |
| Enpp2 | 1970.739579 | -0.701215325 | 6.30E-06 |
| Havcr2 | 13.25216883 | -0.702233561 | 0.004357363 |
| Plac8 | 10.99899382 | -0.702966064 | 0.007139698 |
| Hyal1 | 952.2654872 | -0.703984944 | 1.94E-08 |
| Eva1b | 65.10168811 | -0.704168084 | 0.001962671 |
| Ifi35 | 172.4024348 | -0.707105272 | 2.33E-05 |
| Tmem98 | 245.8181761 | -0.710843181 | 8.71E-06 |
| Ubd | 14.54165072 | -0.71145348 | 0.003822556 |
| Mt2A | 1590.425522 | -0.713239485 | 4.82E-06 |
| RGD1561157 | 58.39030811 | -0.714027103 | 0.000642835 |
| Slfn13 | 58.25029285 | -0.717407149 | 0.001702948 |

| | | | |
|---|---|---|---|
| Slc6a9 | 5242.229922 | -0.719343921 | 1.90E-08 |
| P2ry12 | 14.32121742 | -0.720272983 | 0.005046021 |
| Fgfrl1 | 1259.112285 | -0.722168793 | 2.06E-08 |
| Muc1 | 74.94000593 | -0.724489602 | 0.000306986 |
| Ngb | 51.18957299 | -0.728561006 | 0.00200133 |
| Cd38 | 960.4335231 | -0.728615207 | 1.39E-09 |
| Htra3 | 351.8561831 | -0.730803372 | 2.14E-05 |
| Ppap2c | 55.41317114 | -0.737035146 | 0.000159514 |
| Antxr1 | 375.6356454 | -0.738206817 | 4.22E-05 |
| Echdc3 | 28.38991688 | -0.741500612 | 0.000840224 |
| Rab7b | 489.1800107 | -0.744267432 | 2.89E-07 |
| Plekhg2 | 241.035227 | -0.744351306 | 1.98E-07 |
| Col16a1 | 841.3049237 | -0.745068745 | 9.98E-06 |
| Bin2 | 28.1040295 | -0.747330904 | 0.0018302 |
| Colec12 | 70.30996482 | -0.751017925 | 0.000290388 |
| Phyhd1 | 791.6651055 | -0.756908811 | 1.87E-07 |
| Tnfrsf1b | 9.15475911 | -0.759144792 | 0.006445555 |
| Phldb1 | 1664.013856 | -0.760078739 | 2.68E-08 |
| Tlr9 | 10.09101327 | -0.761628225 | 0.004942284 |
| Tap2 | 856.1777457 | -0.762308906 | 1.31E-06 |
| Ttc12 | 328.2415809 | -0.764446803 | 1.20E-06 |
| Cebpd | 620.4084434 | -0.7697268 | 0.00050187 |
| Ddx58 | 792.9196473 | -0.773174444 | 1.14E-06 |
| Rph3al | 350.8497068 | -0.774616899 | 4.92E-07 |
| Fgr | 21.83819288 | -0.775481975 | 0.003894215 |
| Sp110 | 128.8750053 | -0.775801312 | 2.92E-05 |
| Csf1 | 1353.315249 | -0.778401926 | 3.15E-07 |
| Tsnaxip1 | 32.19067105 | -0.778502464 | 0.003863417 |
| Cd68 | 9.460486402 | -0.779265605 | 0.003910145 |
| Thbs2 | 66.27029145 | -0.780904949 | 0.000484629 |
| Foxj1 | 841.0797301 | -0.783526631 | 8.93E-05 |
| RT1-CE16 | 166.4563101 | -0.792280847 | 0.000340929 |
| Traf4 | 344.5691728 | -0.795459514 | 1.00E-07 |
| Cmtm3 | 34.99103037 | -0.800096371 | 0.00106731 |
| Sparc | 37598.57016 | -0.804696687 | 7.67E-06 |
| Tmem176b | 866.0587741 | -0.806002805 | 4.04E-06 |

| | | | |
|---|---|---|---|
| Pdlim2 | 69.64443656 | -0.812175634 | 5.12E-05 |
| Myo1f | 10.0905573 | -0.816609583 | 0.003910145 |
| Cp | 1313.988663 | -0.826457109 | 0.000922482 |
| RT1-DMb | 21.04219302 | -0.826506202 | 0.003193465 |
| Tubb6 | 165.3425705 | -0.827957371 | 0.001247377 |
| Cx3cr1 | 49.90448384 | -0.828624924 | 0.000536919 |
| Vwa5a | 1136.963632 | -0.829283686 | 6.16E-11 |
| Bcl3 | 46.16682157 | -0.830726863 | 0.003302544 |
| Gdpd2 | 458.958981 | -0.830961794 | 1.58E-06 |
| B2m | 25830.76251 | -0.835533094 | 4.92E-05 |
| Tmem255b | 30.31663576 | -0.835590473 | 0.000527929 |
| Gpr84 | 8.531970423 | -0.835883885 | 0.005310992 |
| Bcas1 | 1224.886675 | -0.847411162 | 6.23E-07 |
| Cdkn1c | 77.02681506 | -0.848263283 | 0.000174122 |
| Tspan11 | 187.4227901 | -0.851607093 | 7.42E-06 |
| Clic2 | 12.24134607 | -0.858329654 | 0.002600481 |
| Fam161a | 64.92881347 | -0.862081424 | 0.000311247 |
| Tekt1 | 264.4701627 | -0.863192285 | 1.51E-05 |
| Tmem176a | 253.5486234 | -0.863656162 | 3.59E-05 |
| Lpcat2 | 69.43246267 | -0.869482195 | 0.000103894 |
| Fabp7 | 2244.127471 | -0.87088314 | 2.46E-05 |
| Bub1b | 16.80370171 | -0.871821223 | 0.000784253 |
| Tnfrsf1a | 869.8141703 | -0.874457027 | 1.43E-14 |
| A2m | 498.5196235 | -0.879732715 | 0.000181367 |
| Serping1 | 1972.464505 | -0.880533091 | 0.002449763 |
| Abcb1b | 225.4696266 | -0.883710503 | 4.66E-05 |
| Trim21 | 250.1432757 | -0.885783613 | 1.15E-06 |
| Tspo | 174.037152 | -0.886797865 | 0.000207173 |
| Irf9 | 419.3603313 | -0.90641857 | 7.01E-06 |
| Mcam | 271.9130899 | -0.907584513 | 0.000467311 |
| Tgif2 | 139.0225665 | -0.920854147 | 1.68E-06 |
| C1ql1 | 386.890871 | -0.923638371 | 9.81E-08 |
| Parp3 | 300.4592867 | -0.924885748 | 3.32E-06 |
| Kcnj16 | 529.2518434 | -0.931920878 | 0.000108706 |
| Dab2 | 132.6452271 | -0.939883373 | 3.99E-06 |
| Fn1 | 217.779601 | -0.939950861 | 8.48E-08 |

| | | | |
|---|---|---|---|
| RT1-CE10 | 247.0515965 | -0.954439501 | 0.00031603 |
| Myo7b | 2.059373851 | -0.958665499 | 0.002552263 |
| RT1-A1 | 680.6519262 | -0.959147666 | 9.32E-05 |
| Grifin | 6.890436046 | -0.961347902 | 0.003719026 |
| Serinc5 | 1913.92908 | -0.976008671 | 6.94E-12 |
| S1pr3 | 668.1052735 | -0.981598428 | 0.000574698 |
| Dhx58 | 201.4920525 | -0.988212677 | 1.93E-05 |
| Rhoj | 41.80089397 | -0.991125418 | 1.24E-05 |
| Rbm43 | 116.0170907 | -0.999304607 | 9.11E-09 |
| Mmp2 | 274.5889951 | -1.00719199 | 3.00E-05 |
| Laptm5 | 37.75812342 | -1.009574361 | 0.00059914 |
| Slco2b1 | 37.91237989 | -1.011269361 | 3.17E-05 |
| Zmym6nb | 45.95079524 | -1.016426002 | 0.000233778 |
| Helz2 | 85.08582648 | -1.037033196 | 5.75E-05 |
| Mx2 | 2426.407938 | -1.047401158 | 0.000343149 |
| Afap1l2 | 493.762997 | -1.056378116 | 1.08E-08 |
| Dnai2 | 116.4933185 | -1.061622235 | 2.50E-05 |
| Parp9 | 461.0214173 | -1.064467345 | 6.96E-09 |
| RT1-CE4 | 133.0785333 | -1.068210427 | 0.000110565 |
| Dtx3l | 701.5481025 | -1.068902037 | 2.03E-06 |
| RT1-T24-4 | 332.9866622 | -1.069739837 | 8.16E-08 |
| Mki67 | 32.7910202 | -1.085122094 | 0.000190831 |
| Pllp | 423.2745054 | -1.113862645 | 3.45E-05 |
| Dock6 | 161.1902934 | -1.116711412 | 1.87E-12 |
| LOC681766 | 16.82028652 | -1.120844851 | 0.001493792 |
| Gsx1 | 4.258495376 | -1.124501744 | 0.002597157 |
| Fxyd5 | 12.81951208 | -1.126288772 | 0.000708477 |
| C2 | 223.8908444 | -1.12965905 | 0.00058538 |
| Lgals3bp | 753.1893042 | -1.130351382 | 1.26E-06 |
| Bst2 | 82.70013448 | -1.155229342 | 0.001129876 |
| Uba7 | 96.81122359 | -1.155654033 | 0.000264615 |
| Acot1 | 91.84113092 | -1.17261076 | 1.44E-06 |
| RT1-N3 | 41.30577063 | -1.181442617 | 0.001146913 |
| Cxcl16 | 167.7752772 | -1.193558883 | 2.37E-06 |
| Trim25 | 159.7712734 | -1.212034101 | 1.13E-08 |
| Sema3d | 399.1903164 | -1.212043045 | 1.33E-06 |

| | | | |
|---|---|---|---|
| Zbp1 | 3.728791403 | -1.223524235 | 0.001200179 |
| Ifi44 | 209.4460464 | -1.230382163 | 7.74E-08 |
| Oas1a | 75.32870661 | -1.233839732 | 0.001090595 |
| Stat1 | 4056.666291 | -1.235085304 | 4.77E-06 |
| LOC308990 | 2.381122746 | -1.250335607 | 0.001674801 |
| Epsti1 | 12.84937651 | -1.256592218 | 0.001154068 |
| Aif1 | 8.292806045 | -1.275862802 | 0.000916578 |
| Calcrl | 100.4629871 | -1.276896938 | 9.32E-07 |
| Vcan | 592.783212 | -1.281644322 | 2.05E-11 |
| Adamts7 | 21.94162133 | -1.294901308 | 0.000255727 |
| C3 | 406.5144916 | -1.29888501 | 0.001070478 |
| Parp14 | 727.3988232 | -1.310326506 | 5.63E-07 |
| C1r | 1431.916971 | -1.319618493 | 2.62E-06 |
| C1s | 1004.124088 | -1.332936956 | 9.62E-06 |
| Psmb8 | 482.2250051 | -1.338344597 | 4.78E-06 |
| Matn4 | 42.16246145 | -1.341970633 | 0.000402708 |
| Timp1 | 53.75333992 | -1.34514842 | 0.000503861 |
| Irx1 | 41.38487083 | -1.401945745 | 6.72E-05 |
| Erbb3 | 319.9841722 | -1.414981788 | 6.18E-08 |
| Rac2 | 8.193147327 | -1.418100047 | 0.001483905 |
| Apol9a | 42.14713388 | -1.422212982 | 0.001474801 |
| Rhbdf2 | 63.78390422 | -1.427368396 | 1.39E-09 |
| Cxcl9 | 96.30732192 | -1.44377091 | 0.000990533 |
| Spi1 | 7.784158665 | -1.448407753 | 0.00147726 |
| Fcer1g | 28.10389691 | -1.462563048 | 9.19E-05 |
| Socs1 | 38.04863651 | -1.466058471 | 1.54E-05 |
| Casp4 | 162.2969613 | -1.469603248 | 8.72E-07 |
| Gbp4 | 21.36716466 | -1.475387348 | 0.000849828 |
| Cfh | 70.46291433 | -1.478273552 | 0.000145448 |
| Tagln2 | 209.572136 | -1.506432704 | 3.39E-05 |
| Psmb10 | 347.1107319 | -1.523452267 | 1.28E-05 |
| MGC105567 | 68.41163706 | -1.547111338 | 0.000243179 |
| Sox10 | 635.6735299 | -1.589026464 | 3.35E-08 |
| Adgre1 | 20.77759917 | -1.598535225 | 0.000245401 |
| Irf1 | 472.3585257 | -1.601241757 | 2.94E-06 |
| Csf1r | 101.6269058 | -1.622436012 | 5.63E-08 |

| | | | |
|---|---|---|---|
| Irgm | 527.7767198 | -1.657542414 | 6.36E-07 |
| Mgp | 8.967814264 | -1.667466332 | 0.000442909 |
| Oas1b | 59.29023618 | -1.685995462 | 0.000527927 |
| P2ry6 | 11.97141036 | -1.686846913 | 0.000241792 |
| Igtp | 521.9803348 | -1.749847552 | 1.50E-05 |
| Tap1 | 837.8446153 | -1.758710352 | 3.18E-06 |
| Csf2rb | 6.582571797 | -1.790816955 | 0.000779398 |
| Chst3 | 17.43999588 | -1.855573422 | 0.000127468 |
| Bmp4 | 47.98336117 | -1.855783728 | 4.22E-07 |
| Psmb9 | 359.767857 | -1.867065418 | 7.75E-07 |
| Irf8 | 41.39491844 | -1.88507751 | 2.21E-05 |
| Pld4 | 33.06495369 | -1.904975727 | 5.21E-05 |
| Olig1 | 1529.229499 | -1.90891987 | 7.51E-24 |
| Cav2 | 153.7741035 | -1.910985444 | 1.09E-12 |
| Pnlip | 31.23089454 | -1.947126742 | 2.29E-06 |
| Rtp4 | 158.6066423 | -1.951524218 | 2.70E-09 |
| Il18bp | 176.1441494 | -1.963495489 | 3.07E-08 |
| Irf7 | 177.6167389 | -1.990377013 | 1.62E-06 |
| Parvg | 7.905878515 | -1.997978424 | 0.000171835 |
| Qrfpr | 18.07836828 | -1.999575506 | 0.000371864 |
| C1qa | 56.85532742 | -1.999719725 | 3.20E-06 |
| Cdh3 | 27.02794979 | -2.003858276 | 6.94E-06 |
| Ifitm3 | 147.5282011 | -2.01698445 | 0.00010395 |
| Cfb | 136.6528607 | -2.074489209 | 0.000192625 |
| Olig2 | 606.1417983 | -2.081192315 | 5.62E-24 |
| Gpr34 | 12.02248222 | -2.091971627 | 9.00E-05 |
| Ccl2 | 17.62441728 | -2.120832213 | 0.000391177 |
| Mpeg1 | 101.5736678 | -2.155559692 | 2.92E-09 |
| Slamf8 | 36.33508678 | -2.162874063 | 5.01E-06 |
| Hsh2d | 5.682909328 | -2.163343619 | 0.000421923 |
| Arhgap31 | 222.8017751 | -2.177077097 | 2.27E-18 |
| Anxa3 | 78.42925651 | -2.20761933 | 8.52E-09 |
| Tmem106a | 11.98982392 | -2.231890764 | 5.27E-05 |
| Oasl2 | 218.6182184 | -2.232226302 | 1.47E-07 |
| Gbp5 | 437.0001691 | -2.279226949 | 1.04E-06 |
| RT1-Ba | 29.63190245 | -2.28340919 | 0.00032813 |

| | | | |
|---|---|---|---|
| Mrap | 5.495310574 | -2.283922864 | 0.000341805 |
| Cav1 | 196.5938146 | -2.297409429 | 4.03E-19 |
| Ctss | 133.0650958 | -2.355975551 | 1.03E-10 |
| Itgam | 40.20327333 | -2.372572008 | 1.17E-05 |
| LOC100910973 | 298.5221864 | -2.373219005 | 7.75E-07 |
| Gbp2 | 833.7330989 | -2.468160956 | 3.63E-06 |
| C1qb | 45.86784839 | -2.484636346 | 2.85E-07 |
| Itgal | 18.37968615 | -2.533278668 | 2.70E-06 |
| MGC108823 | 555.5814526 | -2.557445334 | 1.08E-05 |
| Cd74 | 204.7571086 | -2.567990568 | 5.45E-05 |
| Isg15 | 182.6332451 | -2.59229472 | 1.07E-06 |
| Cspg4 | 768.3465006 | -2.662391121 | 6.99E-42 |
| Lgals9 | 80.3019239 | -2.675257815 | 8.59E-08 |
| Fam89a | 33.19435762 | -2.676794067 | 1.33E-09 |
| Itgb2 | 25.08091154 | -2.728902602 | 1.96E-06 |
| Usp18 | 56.14930203 | -2.825555436 | 6.43E-08 |
| Rsad2 | 467.7145128 | -2.860050552 | 9.09E-08 |
| RGD1309362 | 502.1726712 | -2.907009696 | 2.84E-06 |
| Fcrl2 | 26.96153006 | -3.015226359 | 5.83E-08 |
| C1qc | 28.7981149 | -3.086717717 | 8.52E-09 |
| Col5a3 | 438.4709126 | -3.179478079 | 2.03E-27 |
| Chst5 | 30.45888523 | -3.26709391 | 2.47E-11 |
| Neu4 | 77.4446178 | -3.346666255 | 1.07E-25 |
| Ccnd1 | 344.3165307 | -3.444190753 | 1.20E-37 |
| Mx1 | 235.1915312 | -3.578542113 | 1.64E-07 |
| Ifit3 | 71.33921826 | -3.652438456 | 5.76E-06 |
| Ifi47 | 101.4097503 | -3.672308984 | 1.84E-06 |
| Gpr17 | 177.4145954 | -3.783215153 | 2.44E-42 |
| Pdgfra | 1182.078546 | -3.794307889 | 7.07E-49 |
| RGD1311892 | 21.50503207 | -3.795453543 | 1.12E-12 |
| Cxcl11 | 11.58116095 | -4.103786036 | 4.75E-06 |
| Cxcl10 | 223.4037959 | -4.28401712 | 2.98E-08 |

| Gene | baseMean | log2FoldChange | adjusted p-value |
|---|---|---|---|
| Myh9l1 | 144.6701119 | 25.11521868 | 2.06E-13 |

| Gene | baseMean | log2FoldChange | adjusted p-value |
| --- | --- | --- | --- |
| Mocs3 | 423.71109 | -0.100392327 | 0.023864064 |
