## Supplementary material for "Investigating cocaine- and abstinence-induced effects on astrocyte gene expression in the nucleus accumbens": Experiment 1_Common and Group Specific Significant DEGs

| Gene Name |
| --- |
| Egr2 |
| Npas4 |
| Egr4 |
| Fosb |
| Tac1 |
| Junb |
| Scn4b |
| Lamp5 |
| Gpr6 |
| Gpr88 |
| Egr3 |
| Hpca |
| Drd1 |
| Pdyn |
| Slc4a11 |
| Rem2 |
| Otof |
| Pde10a |
| Sh3rf2 |
| Iqgap3 |
| Rasd2 |
| Tac3 |
| Penk |
| Akap5 |
| Lmo7 |
| Lpl |
| Camkk2 |
| Actn2 |
| Rbfox1 |
| Foxp2 |
| Rgs9 |
| Ppp3ca |
| Camk4 |
| Bcl11b |
| Asb2 |
| Cpne5 |
| Chrm4 |
| Kcnab1 |
| Kcnh4 |
| Necab1 |

Fbxl16
Ptpn5
Adora2a
Filip1
Wipf3
Pde1b
Plxdc1
LOC100125362
Itk
Olfml2a
Phyhip
Wnk4
Wfs1
Chst15
Syt10
Lzts3
Gabrd
Asic4
Foxp1
Adcy5
Cdc42ep3
Cyfip2
Gng7
Atp6ap1l
Tmem158
Camk2b
Rab40b
Grm4
Synpr
Htr1b
Mast3
Arpp21
Rgs7bp
Cacnb4
Kcns2
Tomm70a
Kcna5
Elmod1
Crip3
B3gnt2
Phactr1

Zfp189
Ypel2
Lingo3
Acvr1c
Adamts3
Rxrg
Dlgap3
Cartpt
Jph4
Gpr176
Klf5
Gpr52
Trim46
Neurl1
Syndig1l
C2cd2l
Cntn3
Igfbp4
Lrrk2
Ptpn7
Celf5
Rnf144b
Vstm5
Lppr4
Actn1
Rgs2
Dact2
Stk32c
Rapgef5
Kcna4
Pdp1
Cacnb2
Slc24a4
Rbm3
Mtus2
Tbc1d8
Gnal
Prrt2
Dmtn
Psd
Lrp12

Krt71
Cpne6
Ankrd34a
Fam102b
Gpr173
Zdhhc14
Ppp4r4
Nptn
Kcnq2
Fkbp1a
Jph3
Tiam1
Map2k1
Dlg4
Cacnb1
Cacnb3
Gabra4
Camkk1
Klf16
Ncdn
Tex15
Trerf1
Shc2
Zfhx2
Shank3
Dclk3
Kcnt1
Shisa7
Doc2b
Pdzd2
Cdk17
Pde7b
Ptk2b
Dgki
Mark2
Rps6ka2
Dgkb
Pthlh
Cplx2
Kcnj2
Lgi1

Cacna1c
Lancl1
Dlg3
Lppr1
Rarb
Syt4
Ppp1r9a
Nexn
Lypd1
Scn2b
Grm5
Slc35d3
Ncs1
Prkar2b
Arhgef25
Hsph1
Plekha5
Kitlg
Dapk1
Tasp1
Celf3
Drd2
Opn3
Fbxo41
Gria1
Sipa1l1
Josd1
Syndig1
Runx1t1
RGD1564664
Erbb4
Gucy1b3
Kcnh5
Chgb
Matk
Hs6st2
Cyld
Abcc5
Mtmr7
RGD1311739
Celf4

Necab2
Fam110b
Il10ra
Pdpk1
Kl
Fbxo34
Gga3
Mapk1
Myh7
Dlg2
Arhgef9
Srrm3
Klhl29
Nalcn
Ppip5k1
Rasgrp2
Csrnp3
Crtc1
Slc12a5
Dusp8
Zfp280b
Ppp1r12b
Mef2d
Ip6k2
Begain
Man1a1
Prkar1b
Epb41l1
Slitrk5
Clvs2
Fbxl15
Prkg1
Pdxp
Micu3
Ndufaf6
Lrrc4c
Stx1b
Dlx5
Ppp1r1b
Nsg2
Rbfox2

Ank3
Ppp1r13b
Pip5k1a
Cacna2d3
Htr2c
Sertad4
Spock3
Evl
Osbpl8
Srf
Prkch
Sv2c
Plekha8
Map3k13
Meis2
Ccdc85a
Cbx6
Man1c1
Mafb
Sptb
Sez6l2
Zfp575
Slc2a13
Arid4a
Map3k12
Ajap1
Kctd8
Sbk1
Nxph1
Pdzd4
Mgat4a
Sp9
Gucy1a3
Pnmal1
Gripap1
Zfp280d
LRRTM1
Bag4
Agk
Syngr3
Mbnl1

Mink1
Wscd2
Jakmip1
Rap1gap
Senp2
Plxna2
Sytl5
Dlx2
Pianp
Ablim2
Tomm20
Gtpbp6
Rybp
Ccdc64
Pja2
Slc25a25
Bex1
Rasgrf1
Kit
Rgs8
Grip1
Scn3a
Tec
Ptchd1
Tenm2
Mmp24
Stk24
Nyap1
Phf20
Pip4k2b
Ndel1
Crabp1
Fbxo33
Sptbn4
Dnajb4
RGD1563441
Ezh1
Ddi2
Gnb5
Atmin
Ablim3

Galnt13
Cacna1h
Slmap
Lrrc10b
Smad3
Mnt
Nfx1
Unc13c
Ica1
Pkia
Gpr83
Elk1
Trim66
Fam84a
Ccdc92
Pbx2
Atg13
Mbnl2
Spin1
Bmyc
Rapgef6
Prkacb
Mpped2
Dnajc5
Kctd1
Armcx3
Ppm1h
Fstl4
L1cam
Pom121
Arpc1a
Nmnat2
Zfp483
Rims3
Ptprm
Dzip1l
Slc9a5
Nrxn3
Nol4
Hipk3
Zbtb7a

Rufy3
Stim2
Mesdc1
Ppfia4
Dos
Dbndd1
Rai1
Sgsm2
Ppp1r12c
Alcam
Mn1
Abhd8
Gria4
Dmtf1
Dip2c
Scg2
Camk1d
Chchd4
Jade2
Tspyl4
Spire2
Fam13b
Rhobtb2
Kcnd2
Armc8
Ogfod1
Cbx7
Tcf20
Fam168b
Gad1
Epha4
Rnf44
Cep290
Chpf
Pnmal2
Pds5b
Dynll2
Scamp1
Diaph1
Ppp1r2
Usp22

Btrc
Mypop
Hdac11
Mapre2
Kcnip2
Tulp4
Grb2
Prkca
Zfp638
Camsap2
Rnf185
Zfp292
Prepl
Brd4
Irs2
Stx16
Thsd7a
Matr3
Slk
Tmem199
Nupl1
Ago2
Dyrk1a
Mtch1
Rc3h2
Slc22a17
Nr3c2
Slc7a14
Zfp644
Ctnnbip1
Usp11
Cds2
Dpysl5
Ncoa5
Rnf187
Mecp2
Pja1
Ubp1
Mib2
Sin3b
Tmem57

Gltscr1l
Prrc2c
Bcl7b
Nckipsd
Aqr
Zfp532
Taok2
Med24
Aarsd1
Rabl6
Akr1a1
Celf1
Rnf111
Rxrb
Cbfa2t2
Ubqln2
Nolc1
Wbp11
Plekhb2
Akap17a
Fam134c
Emc2
Flot2
Tex264
Ifnar1
Wbp1l
Stat5b
Cadm4
Ndst1
Ccs
Pmm1
Sptlc1
Gca
Mettl9
Ppp2r5a
Pir
Coasy
Ankrd40
Abcd1
Pik3ip1
Sox4

Rras2
Ddt
Comt
Slc48a1
Sgcb
Anxa5
Fzd9
Zfp703
LOC306766
Ntrk2
Tmem256
Nln
Tspan3
Stx17
Adprhl2
Ubr7
Ddah1
Rasa3
Hip1
Odc1
Smim15
Tex261
Rab5c
Lzic
Rap1a
Hadh
Arhgap5
Nop9
Casp3
Mfsd1
Dennd5a
Spg20
Vimp
Tspan14
Ccdc90b
Ccng1
Sdhc
Tspan2
Nudcd2
Dusp15
Mt3

Nadk
LOC691807
St3gal4
RGD1308428
Golga7
Hist1h1d
Chmp1b
Slc50a1
Leprot
Sh3bgrl
Hexa
Lhx2
RT1-Db1
Hps1
Fzd10
Nedd1
Mapre1
Nek9
Rb1
Pon2
Slc13a3
Ppfibp1
Gcdh
Rnpepl1
Cers4
Sirt2
Ybx1
Bphl
Mcee
Dhcr7
Hist1h2bh
Plekhd1
Fam63a
Tvp23b
Pdzrn3
Pccb
Nfia
Akt2
Tmem209
Ostf1
Ctf1

Lhpp
Hes6
S100a13
Gpm6b
Idh1
Fgfr1
Hrsp12
C1rl
Ctnnd1
Kazald1
Gng5
Ccl7
Traf7
Szrd1
Nadk2
Fam107a
Kctd15
Nes
Ninj2
Hist1h4b
Tp53inp1
Npc2
Nckap1l
Slc27a6
Scarb2
Krcc1
Mgmt
Pmp22
Fabp5
Snapc2
Clec2g
Tspan12
Ddr2
Scd2
Dnase2
Nfe2l2
Rab31
Cd180
Sepp1
Spsb2
Mag

Ppp1r3c
Evi2a
Rela
Slc7a5
Bax
Sh3bp4
Oplah
Chst7
Myo9b
Sfrp2
March8
Wipi1
Cyp20a1
Ppap2a
Amotl2
Pxn
Sardh
Btbd7
Glis2
Gna12
Tor3a
Fuca1
Slc27a1
Tp53i3
Ctdsp1
Blnk
Itpripl2
Hipk2
Cd151
Top2a
Kif13a
Syngr2
Nuf2
Mybl1
Shc1
RT1-M3-1
Nqo1
Csrp1
Cpm
Ctnna1
Fgfr2

Prtfdc1
Slc44a1
Enpp6
Sesn2
Ssfa2
Tp53
Tshr
Sp1
Rhog
Ppic
Hsd17b4
Tnfsf12
RGD1311756
Ppp1r36
Ifi27
Hn1l
C4a
Has2
Ezh2
Pttg1ip
Fnbp1
Glb1l
Suclg2
Lpar4
Il10rb
Mlc1
Evc
Jak3
Hapln3
Cmtm5
Nsmce1
Cd24
Pdgfc
Cd81
Cd302
Ddr1
Hey2
Cldn11
Ppp2r1b
Ctsh
Sh3glb1

Kif1c
Steap3
Lamp2
Fgfbp3
Efs
Cenpf
Tspan6
Apol3
Stat3
Lrp10
Scnn1a
Frmd8
Fkbp10
Sox9
Sspn
Plekhg1
Bgn
Abtb2
Map4k4
Sh3bgr
Slc39a12
Ptprz1
Scamp2
Elovl1
Atf3
Rps27l
Wscd1
Nat6
Carhsp1
Adamtsl4
Rgma
Cd99
Plcd1
Gpam
Snx33
Itpkb
Ckap2
Fa2h
Pbxip1
Erbb2
Prelp

Slc12a2
Fkbp9
Gsta1
Adamts4
Rsu1
Npepl1
Rras
Cst3
Smo
Myl12a
Tubb2b
Necap2
Lpar1
Ssc5d
Nfatc1
Clic1
Rnaset2
Fam167a
Cnn3
Cd63
Dapp1
Vamp3
Stk10
Pth1r
Lgi4
Gpr37l1
Efemp2
Gpr37
Slc39a1
Ephx1
Nod1
Plek
Sat1
Trip6
Itgb5
Arpc1b
Slc12a4
Arhgef19
Lhfpl2
Gab1
Plxnb3

Nt5dc2
Dcakd
Mvp
Slc38a3
Pla2g16
Gfap
Fxyd1
Rhoc
Adam17
Lpcat3
Ltbr
Dbi
Axl
Gtse1
Icam1
Nkd1
Ifih1
Tnfrsf19
Ptpn6
Cdk6
Jam3
Gng11
Pdlim4
Pros1
Cpxm1
Cdkn1a
Slc14a1
Msn
Mid1ip1
Litaf
Abhd4
Fas
Mog
Adamts1
Slc27a3
Abca8a
Opalin
Gltp
Eif2ak2
B3gnt7
Col1a2

Phlda3
Flnc
Aldh4a1
Gpnmb
Trim5
Creb5
Nkx2-2
Cnp
Capn6
Apbb1ip
Npr1
Eln
Hyal1
Mt2A
RGD1561157
Slfn13
Htra3
Ppap2c
Antxr1
Col16a1
Phyhd1
Phldb1
Ddx58
Sp110
Csf1
Traf4
Tmem176b
Pdlim2
Myo1f
Cp
RT1-DMb
Tubb6
Cx3cr1
Vwa5a
Bcl3
Bcas1
Tmem176a
Lpcat2
Fabp7
Bub1b
Tnfrsf1a

Serping1
Abcb1b
Tspo
Irf9
Mcam
Fn1
Serinc5
S1pr3
Rbm43
Mmp2
Afap1l2
Parp9
RT1-T24-4
Pllp
Dock6
Bst2
Uba7
Acot1
Trim25
Sema3d
Calcrl
Vcan
C3
Parp14
C1r
C1s
Psmb8
Matn4
Timp1
Erbb3
Apol9a
Cxcl9
Fcer1g
Casp4
Cfh
Tagln2
MGC105567
Sox10
Adgre1
Irf1
Csf1r

Irgm
P2ry6
Igtp
Tap1
Chst3
Bmp4
Psmb9
Pld4
Olig1
Cav2
Pnlip
Rtp4
Irf7
C1qa
Ifitm3
Cfb
Olig2
Ccl2
Mpeg1
Slamf8
Arhgap31
Anxa3
Gbp5
RT1-Ba
Cav1
Ctss
Itgam
LOC100910973
Gbp2
C1qb
Itgal
MGC108823
Cd74
Isg15
Cspg4
Fam89a
Itgb2
Rsad2
RGD1309362
Fcrl2
C1qc

Col5a3
Chst5
Neu4
Ccnd1
Mx1
Ifit3
Ifi47
Gpr17
Pdgfra
RGD1311892
Cxcl11
Cxcl10

Gene Name
Ripk4
P2ry1
Slc26a10
Calcr
Il1rapl2
Zfp503
Prkg2
Samd12
Frem3
RGD1309779
Pxdn
Pcdh8
Gpr101
Col14a1
Tmem164
Stc1
Trank1
Peli3
Nsg1
Zfp286a
Slc25a30
Bbs4
Dcp1a
Zfp354c
Gaa
Fam220a
Jade1
Mdm4
Agfg2
Fbxo42
Zcchc8
Crebl2
Adar
Ipo9
Tmem14c
Ndufb2
Dusp3
Nav2
Glo1
Omg

Pomt1
Daglb
Slc35b2
Cc2d1b
Insig1
Hmgcs1
Acyp1
Mettl25
Sqle
Etv4
Mpzl1
Ldlr
Prr11
Aamdc
Mettl4
Abi3
Cd53
Pbld1
Pik3ap1
Igfbp2
Cndp1
Ascl3
Osmr
