## Supplementary material for "Investigating cocaine- and abstinence-induced effects on astrocyte gene expression in the nucleus accumbens": Experiment 2_Common and Group Specific Significant DEGs

| Gene Name |
| --- |
| Pdgfra |
| Gpr17 |
| Col5a3 |
| Olig1 |
| Cspg4 |
| Pmel |
| Egr3 |
| Arhgap31 |
| Olig2 |
| Ccnd1 |
| Cav2 |
| Susd5 |
| Pde10a |
| Calcrl |
| Fosb |
| Lzts3 |
| Lmo7 |
| Ncdn |
| Mmp2 |
| Fam89a |
| Necab1 |
| Cav1 |
| Egr4 |
| Crtc1 |
| RGD1310819 |
| Wipf3 |
| Anxa3 |
| Ppp3ca |
| Htr1b |
| Map3k13 |
| LOC100125362 |
| Serinc5 |
| Pllp |
| Per2 |
| Sowaha |
| Chst3 |
| LOC103692025 |
| Sema3d |
| Mtus2 |
| Sf1 |

Scd
Sox10
Lingo3
Adcy5
Psd
Slc20a2
Baiap2
Erbb3
Camk2a
Tiam1
Lamp5
Mast3
Mink1
Lgi1
Bmp4
Cacnb1
Akap5
Mef2d
Actn1
Pde2a
Rasd2
Mctp1
Gria1
Fas
Drd1
Bcl11b
Camk4
Phlda3
Fbxl16
Mid1ip1
Tomm70
Penk
Rph3al
Hist1h1d
Ptk2b
Pdpk1
Phactr1
Lzts1
Mpeg1
Rab40b
Chrm4

Ngef
Pdzd2
Pde1b
Rgs14
Map2k1
Dlgap2
Rbfox1
Phldb1
Kank1
Sipa1l1
Chst15
Camk2b
Tp53inp1
Fkbp5
Atp2b1
Gltp
Matn4
Foxp1
Kcnt1
Gabra4
Itpka
Sat1
Sh2d5
Dbi
Fam167a
Nab2
Rgs2
RGD1311892
Tubb2b
Gpr158
Hpca
Egr2
Afap1l2
Sesn2
Agap2
Camkv
Grasp
Ncoa5
Gpr88
Actn2
Lyst

Tpd52l1
Ppm1h
Arpp21
Aldh4a1
Pitpnm3
Rnf150
Rps27l
Cdk17
Ctss
C2cd2l
Plppr4
Cacnb4
Dll1
Scn4b
Dmtf1
Cdkn1a
Shank3
Snx22
Ash1l
Wfs1
Ptgs2
Pde7b
Thrb
Jph4
C1qa
RGD1562037
Cpne5
Tmem158
Dopey2
Carm1
Josd1
Dact2
Dgki
Gucy1b3
Dbx2
LOC100912459
Icam5
Lrrc7
Shisa7
Ccsap
Kcnab1

Cd38
Cdk5r1
Cyfip2
Cacna1g
Ddah1
RGD1564664
Nkiras1
Kctd13
Man1a1
Adcy9
Dhcr7
B3gnt2
Cnksr2
Abhd4
Kctd1
Grm4
Homer1
Jph3
Tiam2
Trim46
Chn1
Ankrd63
Traf7
Cnp
Rem2
Ptpn5
Slc4a11
Top2a
Rnf144b
Ebpl
Ano3
Actr3b
Cers4
Klhl29
Spata2L
Kcnj4
Dlg4
Asic4
Stk32c
Nt5dc3
Kcnj2

Tubgcp5
Ppp3r1
Braf
Gtse1
Kcnh1
Hdac4
Otof
Nova2
Eva1b
Gabrd
Gpr12
Arel1
Nbea
Kcnq5
Camkk1
Ppdpf
Kcnf1
Tjp2
Cdkl5
Synpo
Dusp8
Gcc2
Slmap
Hyal1
Myo5b
Kctd16
Akap11
Celf5
Zfp189
Syngap1
Frmd8
Gucy1a2
Rasgrp2
Gabrb3
Rapgef4
Itgam
Pros1
Fabp5
Cacna1c
Cnnm1
Gria2

Creb5
Fam189b
Skil
Tsc22d1
Bcr
RGD1311739
Lcor
Fkbp1a
Dcbld2
Csrnp2
Xbp1
Mapk1
Trerf1
Dlg2
Bax
Npepl1
Cacna1e
Syndig1l
Ppp1r13b
Prkcz
Ap3b2
Bahd1
Rgs9
Mchr1
Pak6
Gfod1
Nptn
Fam65b
Deptor
Traf4
Nr4a3
Meis2
Vsir
Fam84a
Hsph1
Grin2b
Lgi4
Ablim2
Egr1
Wasl
Oprk1

Tyms
RGD1566359
Lmtk2
Lrrk2
Dgkh
Sh3rf2
Itpr1
Cbx6
Elf1
Gga3
Gtf3c1
Pdyn
Slc2a13
Tac1
Tnfaip8
Cdh3
Ids
Sgpl1
Tmem198
LOC100910792
Zmat4
Lpl
Ctnnd1
Camkk2
Zfhx2
Slc24a4
Sympk
Fry
Ankrd33b
Akap1
Adam17
RGD1560470
Slc12a5
Ryr2
Strip2
Fnta
Wasf1
Nr4a1
Mark2
Npr1
Neurl4

Klf16
Syne1
Cnst
Mtmr12
Plcd1
Kcnh3
Map4k4
Dgkg
Mkl2
Dlgap3
Cdyl2
Emp2
Cacnb2
Lrp12
LOC689986
Arc
Ppp1r9a
Pcsk2
Micall2
Rdx
Abl2
Mapkbp1
Arf3
Afg3l1
Dlg3
Pitpnm2
Dpf1
RGD1304884
Atp1a1
Ezh1
Syt4
Slc27a3
Slit3
Osbpl8
Hpcal4
Sgip1
Mapre1
Wscd2
Nhsl2
C1ql1
Gpr63

Bhlhe23
Ikbip
Zfyve28
Dip2c
Zdhhc23
Ccdc106
Tanc2
Ephx1
Tvp23b
Trrap
Chrm1
Cobl
Kcns2
Npas4
Lamb1
Rai1
Prrt2
Nudt12
Rrm2
Necap2
Rapgef2
Dnajb5
Kcna4
Neto1
Serac1
Klhl2
Usp7
Prkcg
Pdp1
Celf2
Grin1
Sorbs2
Gnal
Fbxo41
Nsg2
Prkcb
Tmem255b
Elmod1
Vangl2
Tecta
Doc2b

Stxbp5l
Cpne6
Sh3rf3
Plcb1
Btbd8
Caln1
Rab21
Sirt2
Car11
Madd
Pnp
Nedd4l
Scn3b
Mpp3
Ubr7
Ablim3
Slc4a10
Krcc1
Micu3
Zfp385b
Sash1
Lrrc8b
Rap1gap
Arhgef3
Celf1
Scn2b
Ssx2ip
Fuca1
Kmt2d
Add2
Kalrn
Prokr2
Casd1
Ets2
Rasgef1a
Dmxl2
Btbd7
Chpf
Irs2
Pcdh1
Htt

Arhgap32
Paip1
Erbb4
Zfp488
Adamts15
Dusp1
Stk3
Kif11
Rapgef5
Gpr6
B3gnt7
Apobec1
Hmgxb3
Dbn1
Aak1
Dlgap4
Sertad4
Mbnl2
Lcn2
Grm5
Fmnl1
Fubp3
Necab2
Grin2a
Rhoc
Foxp2
Ypel2
Arhgef9
Elk1
Elovl5
Gas7
Pja2
Plxnb3
Irx1
Acss2
Idh1
Pgbd5
Pbxip1
Tspyl1
Anks1b
Mef2a

LOC100910518
Rbfox3
Ranbp2
Rgcc
Cx3cl1
Fam126b
Srf
Prp2l1
Bdnf
Trim32
Gng7
RGD1311345
Calm1
Ago2
Gna12
Klhl3
Fbxo34
Unc79
Pcdha4
Celf4
Lrrtm3
Arfgef2
Ppp4r4
B4galnt1
Lancl1
Glce
LOC108348061
Dmxl1
Kitlg
Dgat2
Hipk3
Fgfbp3
Camsap1
Arhgap20
Arhgap10
Tmem30a
Cacna2d1
Cap2
Pi4ka
Klf5
Dgkb

Fam65a
Cst3
Cacna1i
Kcna5
Sgsm3
Frmpd4
Mkl1
Rnf208
Rc3h2
Hapln3
Rab31
Grip1
Dapk1
Ptprn
Kcnk2
Acvr1c
Fosl2
Gpr176
Fam8a1
Rasgrp1
Ccser1
Nell2
Cdc42ep3
Noct
Ss18
Shf
Wdr7
Gal3st3
Tesc
Plk2
Cntn3
Slc25a13
Synpr
Hes6
Drd2
Mecp2
Slc35f3
Syt5
Fgf13
LOC501038
Man1c1

Soga1
Aldh1l1
Ankrd17
Sik3
Ptprj
Dmtn
Sv2c
Ttc19
Antxr1
Nat6
Cd302
Adamts3
Sgsm2
Per1
Ap1s1
Prkar2b
Dpp7
Rhoa
Akap9
Rida
Actr2
Slitrk5
Usp33
Fam171a2
Zfp831
Gad1
C1qtnf4
Calb1
Ankrd34a
Mmd
LOC102546809
Vcan
Dars
Stambpl1
Map9
Itgb5
Ppp3cb
Zmpste24
Plcl2
Nol6
Cbfa2t3

Arap1
Ctxn1
Scn3a
Csrnp3
LOC102551114
Pde1c
RGD1307100
Maged2
Armcx3
Tpm1
Plxdc1
Clvs2
Tcf20
Ajap1
Aga
Celf3
Bgn
Evc2
Frmd6
Pcsk1
Lmbrd2
Adra2b
Gas1
Ppfia2
LOC365985
Ifi30
Padi2
Actl6b
Ncapd2
Syt12
Arhgef26
Dok6
Efs
Ephx4
Fyn
Gpr37l1
Atl1
Sp9
Zfp280d
LOC102549506
Tmed5

Klf3
Pip5k1a
Reps2
Senp2
Sept2
Sox2
Fam117b
Ankrd45
LOC103693584
Nexn
Stk10
Tcf7l1
Vstm2a
Prickle2
Arhgap6
Sox4
Msmo1
Syt10
Glmp
St8sia3
P4htm
Tyro3
Sox9
Pdzrn3
Plxna2
Diras2
Nfx1
LOC102550954
Opn3
Sumf2
Echdc1
Cacna1h
Wdr17
Slk
Nanos1
Prkch
Tac3
Snap23
Ywhaz
Arhgap33
Glcci1

Nxph1
Rgs7bp
Sugp2
Prdm15
Rprd1a
Aprt
Prr12
Nol4
Arl4d
Plekhd1
Zdhhc14
Atf2
Ppp1r7
Wdr47
Scg2
LOC100911196
Atg16l1
Pdgfb
Cpxm1
Asb2
Ip6k2
Snx5
G6pd
L1cam
Jam3
Macrod2
RGD1310209
Crtac1
Strn4
Vipr1
Kcna1
Ctdsp1
Cyp2j3
Fermt2
Dtnb
Pmp22
Sh3bgrl
Nthl1
Tbc1d8
Mdk
Hmgcs1

Dach1
Sbds
Ttc28
Arpp19
Fdft1
Rcbtb2
Kcnt2
Prrx1
Csrp1
Vps50
Kctd8
Runx1t1
LOC680142
Lrrc73
Pex3
Ube2q2l
LOC102547697
Hs6st2
Entpd2
Pnlip
Sowahc
Clock
Hecw2
Pigk
Rock2
Cited2
Mapre2
Lamp1
Arl15
Coq2
Dnaja2
Tnpo3
Pou3f2
Armcx2
Syt1
Slc7a8
Sytl5
Arpc1a
Rbp4
Hsd17b4
Fam110b

Ppp1r3c
Arid1b
Napa
Iffo1
Kcnv1
Axl
Dusp2
Cyld
Slitrk3
Tcf7l2
Car8
Col9a3
Kank2
Amph
Dusp4
Gprin3
Rbm24
Gpr52
Arse
Trim66
Mtmr7
Rab27b
Cd81
Coch
Slc6a11
Syn2
Lrig1
Brinp1
Wipi1
Hsd17b12
Rnf13
Lyplal1
Cyp51
Slc25a1
Hist1h3a
Nup210
Smpdl3b
Myh14
Pdzrn4
Zfp644
Tet3

Ttll12
Hepacam
Dag1
Pdlim5
Krt71
Fras1
Stx16
Lims1
RGD1561931
Cercam
Rps6kl1
Luc7l
Xrcc6
Ube2w
Ccs
Mat2b
Tmem41a
Pdp2
Scai
Tmc6
Sez6
Zbtb16
Sf3a1
March8
Kif1c
Nfe2l2
Lhpp
Mbnl1
Cat
Rftn2
Sgtb
Serpina3n
Syndig1
RGD1309676
Otud1
Slc35d3
Gripap1
Slc22a5
Syt16
Acbd5
LOC102553814

Ptprn2
Mthfd1
Ginm1
Gpcpd1
Plppr5
Adgra3
Btd
Kcnb1
Sdhb
Asrgl1
Scd2
Ddhd1
Dclk3
Npc2
Epb41l4b
Gpr22
Scarb2
Tomm20
Cherp
Gstk1
Cd2bp2
Paqr9
LOC102552990
Pum2
Sepp1
Tcaf1
Lrrc10b
Ide
Kcnh4
Nit2
Ercc6
Cntnap3b
Smarcd1
Prpf8
Itgav
Scp2
Ttc12
Klf10
Kcnd2
Filip1
Coro1a

Ndel1
Supt6h
Eftud2
Csad
Dpysl3
Neu2
Dusp5
Tmem181
Prrc2c
Mlc1
Gng4
Arid4a
Aldh9a1
Usp6nl
Golm1
Mapk12
Zfp496
Dock1
Ing2
Dync1h1
Rassf10
Itga9
Trim23
Chmp1b
Ccdc17
Cuedc1
Psmc3ip
Ctso
Dcaf1
Ddah2
Fam102b
Sspn
Adora2a
Sik1
Folh1
Snx18
LOC102546421
Prkg2
Cnot7
Fbxw4
Eif4enif1

Meis1
Alcam
Usp22
Gcsh
Reep3
Trim7
Ebf1
Park7
Wnt10a
Ddit4l
Dscr3
Ermp1
Mmp24
Cadm4
Arl13b
Atpaf2
Afap1l1
Iqgap3
Tjp1
Acsbg1
Prkacb
Dusp14
Glis3
Emc7
Cnot1
Tmed3
Fads2
Fbln2
Kit
Pcp4l1
Appl1
Ppp1r1a
Dixdc1
Crebbp
Purg
Kremen1
Pafah2
Plppr1
Dmd
Hmg20b
Fstl4

Obsl1
Enpp5
Sdhc
Cartpt
Grm7
Itpa
Rgs8
Setd1b
Ccdc61
LOC102551716
Thsd7a
NEWGENE_1305560
Dock3
Cyth4
Endou
Agk
Usp5
Kcne5
Olfml1
Ccng1
Add3
Cdh2
Vps39
Glul
Tns1
Anp32e
Mafb

| Gene Name |
|---|
| Aamp |
| Abcd4 |
| Abhd13 |
| Acaa1 |
| Acadvl |
| Acbd4 |
| Acot11 |
| Acot13 |
| Acot3 |
| Acvr2a |
| Adamts12 |
| Agl |
| Ak3 |
| Akap3 |
| Aldh7a1 |
| Amotl2 |
| Amt |
| Ankrd40 |
| Ankrd50 |
| Aox1 |
| Asf1a |
| Atox1 |
| Atp1a2 |
| Atp6ap1l |
| Atraid |
| Atrip |
| Atxn7l1 |
| Bag4 |
| Bckdk |
| Bcor |
| Bmp1 |
| Bmp2 |
| Brox |
| Brpf1 |
| C11H22orf29 |
| Cacna2d3 |
| Camk1 |
| Card10 |
| Caskin2 |
| Casz1 |

Catsper2
Cbr3
Ccdc28b
Ccdc77
Ccdc91
Ccnh
Cd59
Cdc42se2
Cep120
Chl1
Chtf8
Cinp
Cipc
Clcn4
Cln5
Clspn
Cntn5
Col14a1
Crabp1
Cramp1
Crot
Csnk1g1
Csrnp1
Ctsd
Dab2
Ddt
Dhrs11
Dlx5
Dmpk
Dnase2
Dnm2
Dnph1
Ebi3
Edc3
Efnb2
Eif2d
Eif4ebp1
Emd
Enox2
Epc2
Epha4

Erich3
Etv3
Etv4
Exoc6
Eya2
Fads1
Fam124a
Fam129b
Fam13b
Fam184b
Fam189a2
Fam213a
Fam49a
Fam78b
Fasn
Fbxl5
Fbxo22
Fbxo3
Fech
Fhl4
Fos
Frem3
Fst
Fuca2
Fut2
Fut4
Galnt13
Gcnt2
Gltscr1
Glyctk
Gmeb2
Gpr149
Gpr83
Gtf2h1
Gucy1a3
Gusb
Hadha
Hadhb
Has3
Hibadh
Hist1h2bcl1

Hist3h2ba
Hmgn5b
Hsd17b11
Hsdl2
Hspa13
Htr2c
Igfbp4
Il10ra
Il11ra1
Il1rapl2
Inf2
Ints7
Isca2
Islr
Ism1
Itgb8
Jph1
Junb
Kantr
Kcnk13
Kctd12
Kdelc2
Kifc3
Kl
Klf2
Klf7
Lca5
Ldhd
Lgr6
Lmo4
LOC100359498
LOC100909405
LOC100909824
LOC100910732
LOC100911029
LOC100911299
LOC100911483
LOC100911734
LOC102547155
LOC102547715
LOC102550527

LOC102551811
LOC102557119
LOC103691165
LOC108348625
LOC108350745
LOC361635
LOC684270
LOC687399
LOC689561
Lonp2
Lrp5
Lrrc4
Lrrc4c
Lsamp
Lss
Ltbp3
Lypd1
Map3k3
Mapkap1
March4
Matr3
Mavs
Mblac2
Me1
Megf11
Mfsd1
Mgat4a
Mgme1
Mier2
Mmab
Mmp15
Mpv17l2
Mrpl32
Mrpl4
Mrpl45
Mthfs
Myh7
ND6
Ndrg2
Ninl
Nkpd1

Nr3c1
Ntng1
Nudt5
Ofd1
Ogdh
Oplah
Oprm1
Osbpl3
P2ry1
Pbx2
Pccb
Pcdhgb4
Pfkm
Pgm3
Phka1
Phlda1
Phyh
Pik3cg
Pir
Pirt
Pkia
Pla2g2c
Plbd2
Plcxd3
Plekha7
Pmm1
Poglut1
Pogz
Polm
Ppox
Prodh1
Psenen
Psph
Pstk
Ptgr2
Pthlh
Ptpn7
Pygm
Rab34
Rad9a
Rad9b

Rarb
Rasal2
Rcl1
Rcor1
Rft1
Rfx4
Rfx5
RGD1303003
RGD1311756
RGD1565536
RGD1565775
RGD1566265
RGD621098
Rgs10
Rgs6
Rictor
Ripk4
Rnf170
Rnf180
Rnf215
Rock1
Rpp25
Rps6ka5
Rxrg
Rybp
Sardh
Sc5d
Sema6d
Sepsecs
Sept10
Setd4
Shisa4
Slc1a3
Slc31a1
Slc39a7
Slc46a1
Slc7a14
Smad3
Smo
Smoc1
Snx7

Sod3
Son
Sox12
Sox21
Sp4
Spata13
Spock3
Ssbp2
Ssbp4
Ssc5d
Ssfa2
Sstr4
St6galnac4
Stap2
Steap3
Suclg2
Tbc1d9b
Tbcel
Tcaim
Tcf3
Tdg
Tec
Tex15
Tex30
Tfe3
Thnsl2
Timp4
Tkt
Tlcd1
Tll2
Tm2d1
Tmem134
Tmem184b
Tmem241
Tmem242
Tmem38a
Tmod1
Tnfsf12
Tpp1
Trappc2
Trim65

Tshz1
Unc5c
Usp8
Vdac2
Vegfb
Vimp
Vof16
Vps13c
Vps29
Vps36
Vstm5
Wdr46
Wfdc1
Yipf1
Yipf3
Zbtb8a
Zc3h12c
Zcchc14
Zfand3
Zfp281
Zfp292
Zfp532
Zfp536
Zfp704
Zfp786
Zhx2
Zswim6
