## Supplementary material for "Investigating cocaine- and abstinence-induced effects on astrocyte gene expression in the nucleus accumbens": Experiment 1_Pathway Enrichment Analysis

| Source | Term Name |
|---|---|
| REAC | Activation of NMDA receptors and postsynaptic events |
| REAC | Activation of the AP-1 family of transcription factors |
| REAC | Ca-dependent events |
| REAC | Calmodulin induced events |
| REAC | CaM pathway |
| REAC | cGMP effects |
| REAC | DAG and IP3 signaling |
| REAC | Estrogen-stimulated signaling through PRKCZ |
| REAC | G alpha (i) signalling events |
| REAC | G-protein mediated events |
| REAC | MAPK family signaling cascades |
| REAC | MAPK1/MAPK3 signaling |
| REAC | Neurexins and neuroligins |
| REAC | Neuronal System |
| REAC | Neurotransmitter receptors and postsynaptic signal transmission |
| REAC | NGF-stimulated transcription |
| REAC | Nitric oxide stimulates guanylate cyclase |
| REAC | Nuclear Events (kinase and transcription factor activation) |
| REAC | Opioid Signalling |
| REAC | Platelet activation, signaling and aggregation |
| REAC | PLC beta mediated events |
| REAC | Post NMDA receptor activation events |
| REAC | Potassium Channels |
| REAC | Protein-protein interactions at synapses |
| REAC | RAF/MAP kinase cascade |
| REAC | Rap1 signalling |
| REAC | RHO GTPases Activate WASPs and WAVEs |

| | |
|---|---|
| REAC | Signal Transduction |
| REAC | Signaling by NTRKs |
| REAC | Signaling by Receptor Tyrosine Kinases |
| REAC | Signaling by Rho GTPases |
| REAC | Signaling by Rho GTPases, Miro GTPases and RHOBTB3 |
| REAC | Signaling by VEGF |
| REAC | Synaptic adhesion-like molecules |
| REAC | Transmission across Chemical Synapses |
| REAC | Unblocking of NMDA receptors, glutamate binding and activation |
| REAC | VEGFA-VEGFR2 Pathway |
| REAC | Voltage gated Potassium channels |
| REAC | alpha-linolenic (omega3) and linoleic (omega6) acid metabolism |
| REAC | alpha-linolenic acid (ALA) metabolism |
| REAC | Axon guidance |
| REAC | Beta oxidation of palmitoyl-CoA to myristoyl-CoA |
| REAC | Cholesterol biosynthesis |
| REAC | Citric acid cycle (TCA cycle) |
| REAC | ECM proteoglycans |
| REAC | Fatty acid metabolism |
| REAC | Fatty acyl-CoA biosynthesis |
| REAC | Linoleic acid (LA) metabolism |

| | |
|---|---|
| REAC | Metabolism |
| REAC | Metabolism of lipids |
| REAC | Metabolism of steroids |
| REAC | Mitochondrial Fatty Acid Beta-Oxidation |
| REAC | Nervous system development |
| REAC | Peroxisomal protein import |
| REAC | Protein localization |
| REAC | RHO GTPases Activate ROCKs |
| REAC | Semaphorin interactions |
| REAC | Signaling by VEGF |
| REAC | The activation of arylsulfatases |
| REAC | VEGFA-VEGFR2 Pathway |

| Term ID | p-value | Directionality |
|---|---|---|
| REAC:R-RNO-442755 | 5.8715E-06 | Up |
| REAC:R-RNO-450341 | 0.0415 | Up |
| REAC:R-RNO-111996 | 0.0002 | Up |
| REAC:R-RNO-111933 | 0.0009 | Up |
| REAC:R-RNO-111997 | 0.0009 | Up |
| REAC:R-RNO-418457 | 0.0415 | Up |
| REAC:R-RNO-1489509 | 0.0009 | Up |
| REAC:R-RNO-9634635 | 0.0300 | Up |
| REAC:R-RNO-418594 | 0.0039 | Up |
| REAC:R-RNO-112040 | 0.0004 | Up |
| REAC:R-RNO-5683057 | 0.0018 | Up |
| REAC:R-RNO-5684996 | 0.0080 | Up |
| REAC:R-RNO-6794361 | 0.0087 | Up |
| REAC:R-RNO-112316 | 3.7522E-11 | Up |
| REAC:R-RNO-112314 | 0.0001 | Up |
| REAC:R-RNO-9031628 | 0.0171 | Up |
| REAC:R-RNO-392154 | 0.0415 | Up |
| REAC:R-RNO-198725 | 0.0377 | Up |
| REAC:R-RNO-111885 | 0.0065 | Up |
| REAC:R-RNO-76002 | 0.0038 | Up |
| REAC:R-RNO-112043 | 0.0004 | Up |
| REAC:R-RNO-438064 | 0.0340 | Up |
| REAC:R-RNO-1296071 | 0.0011 | Up |
| REAC:R-RNO-6794362 | 4.6174E-05 | Up |
| REAC:R-RNO-5673001 | 0.0065 | Up |
| REAC:R-RNO-392517 | 0.0019 | Up |
| REAC:R-RNO-5663213 | 0.0175 | Up |

| | | |
|---|---|---|
| REAC:R-RNO-162582 | 0.0002 | Up |
| REAC:R-RNO-166520 | 0.0140 | Up |
| REAC:R-RNO-9006934 | 0.0130 | Up |
| REAC:R-RNO-194315 | 0.0106 | Up |
| REAC:R-RNO-9716542 | 0.0106 | Up |
| REAC:R-RNO-194138 | 0.0415 | Up |
| REAC:R-RNO-8849932 | 0.0340 | Up |
| REAC:R-RNO-112315 | 1.6911E-05 | Up |
| REAC:R-RNO-438066 | 0.0003 | Up |
| REAC:R-RNO-4420097 | 0.0298 | Up |
| REAC:R-RNO-1296072 | 1.6911E-05 | Up |
| REAC:R-RNO-2046104 | 0.0103 | Down |
| REAC:R-RNO-2046106 | 0.0103 | Down |
| REAC:R-RNO-422475 | 0.0027 | Down |
| REAC:R-RNO-77305 | 0.0303 | Down |
| REAC:R-RNO-191273 | 0.0027 | Down |
| REAC:R-RNO-71403 | 0.0443 | Down |
| REAC:R-RNO-3000178 | 0.0027 | Down |
| REAC:R-RNO-8978868 | 1.0986E-05 | Down |
| REAC:R-RNO-75105 | 0.0103 | Down |
| REAC:R-RNO-2046105 | 0.0429 | Down |

| | | |
|---|---|---|
| REAC:R-RNO-1430728 | 2.6360E-06 | Down |
| REAC:R-RNO-556833 | 0.0014 | Down |
| REAC:R-RNO-8957322 | 0.0294 | Down |
| REAC:R-RNO-77289 | 0.0294 | Down |
| REAC:R-RNO-9675108 | 0.0027 | Down |
| REAC:R-RNO-9033241 | 0.0294 | Down |
| REAC:R-RNO-9609507 | 0.0294 | Down |
| REAC:R-RNO-5627117 | 0.0303 | Down |
| REAC:R-RNO-373755 | 0.0148 | Down |
| REAC:R-RNO-194138 | 0.0148 | Down |
| REAC:R-RNO-1663150 | 0.0429 | Down |
| REAC:R-RNO-4420097 | 0.0294 | Down |

| Intersection |
| --- |
| Camk4,Gria1,Actn2,Prkar2b,Dlg3,Prkacb,Dlg2,Grin1,Dlg4,Camk1 |
| Mapk1,Fos,Atf2 |
| Pde1b,Camk4,Mapk1,Prkar2b,Prkcg,Prkacb,Adcy9 |
| Pde1b,Camk4,Prkar2b,Prkcg,Prkacb,Adcy9 |
| Pde1b,Camk4,Prkar2b,Prkcg,Prkacb,Adcy9 |
| Pde10a,Pde1b,Prkg2 |
| Pde1b,Camk4,Prkar2b,Prkcg,Prkacb,Adcy9 |
| Mapk1,Prkcz,Pdpk1 |
| Pde1b,Rgs9,Mchr1,Lpl,Camk4,Penk,Rgs14,Htr1b,Mapk1,Prkar2b,Grm4,Lrp12,Oprk1,Prkcg,Rbp4,Prkacb,Adra2b,Opn3,Sstr4,Adcy9,Grm7,Rgs8,Oprm1,Plcb1,Rgs10 |
| Pde1b,Camk4,Mapk1,Prkar2b,Prkcg,Prkacb,Adcy9,Plcb1 |
| Cdc42ep3,Actn2,Cnksr2,Mapk1,Kitlg,Dlg3,Braf,Kit,Prkacb,Rasgrp1,Dusp8,Irs2,Prkg2,Dlg2,Kl,Grin1,Dusp2,Dlg4,Ptpn7,Rasgef1a,Rasal2,Pdgfb,Kalrn |
| Actn2,Cnksr2,Mapk1,Kitlg,Dlg3,Braf,Kit,Rasgrp1,Dusp8,Irs2,Prkg2,Dlg2,Kl,Grin1,Dusp2,Dlg4,Ptpn7,Rasgef1a,Rasal2,Pdgfb |
| Grm5,Homer1,Dlg3,Lrrtm3,Dlg2,Dlg4 |
| Kcnab1,Camk4,Gria1,Actn2,Kcnj2,Cacnb4,Gabra4,Kcnh1,Kcnf1,Prkar2b,Kcnq5,Grm5,Kcnk2,Homer1,Kcna5,Dlg3,Prkcg,Lrrtm3,Prkacb,Gad1,Gabrb3,Il1rapl2,Kcns2,Adcy9,Dlg2,Ppfia2,Slitrk3,Slitrk5,Prkcb,Grin1,Kcnv1,Cacna2d3,Syt1,Dlg4,Kcnh3,Kcna1,Kcnb1,Camk1 |
| Camk4,Gria1,Actn2,Kcnj2,Gabra4,Prkar2b,Dlg3,Prkcg,Prkacb,Gabrb3,Adcy9,Dlg2,Prkcb,Grin1,Dlg4,Camk1 |
| Egr2,Nab2,Srf |
| Pde10a,Pde1b,Prkg2 |
| Egr2,Nab2,Srf,Mapk1,Rps6ka5 |
| Pde1b,Camk4,Mapk1,Prkar2b,Prkcg,Prkacb,Adcy9,Oprm1,Plcb1 |
| Actn1,Actn2,Prkch,Mapk1,Dgkh,Prkcg,Dgki,Pdpk1,Adra2b,Rasgrp1,Ywhaz,Dgkg,Prkcb,Rab27b,Rapgef4,Pik3cg,Pdgfb,P2ry1,Islr |
| Pde1b,Camk4,Mapk1,Prkar2b,Prkcg,Prkacb,Adcy9,Plcb1 |
| Camk4,Prkar2b,Prkacb,Camk1 |
| Kcnab1,Kcnj2,Kcnh1,Kcnf1,Kcnq5,Kcnk2,Kcna5,Kcns2,Kcnv1,Kcnh3,Kcna1,Kcnb1 |
| Gria1,Grm5,Homer1,Dlg3,Lrrtm3,Il1rapl2,Dlg2,Ppfia2,Slitrk3,Slitrk5,Grin1,Dlg4 |
| Actn2,Cnksr2,Mapk1,Kitlg,Dlg3,Braf,Kit,Rasgrp1,Dusp8,Irs2,Prkg2,Dlg2,Kl,Grin1,Dusp2,Dlg4,Ptpn7,Rasgef1a,Rasal2,Pdgfb |
| Rap1gap,Prkacb,Rasgrp1,Ywhaz,Rapgef4 |
| Wipf3,Baiap2,Mapk1,Wasf1,Cyfip2,Actr2 |

| |
|---|
| Pde10a,Ppp3ca,Pde7b,Tiam1,Pde1b,Wipf3,Tiam2,Egr2,Rgs9,Mchr1,Lpl,Camk4,Cdc42ep3,Nab2,Actn2,Baiap2,Penk,Nr4a1,Srf,Lrrk2,Adora2a,Cnksr2,Rgs14,Htr1b,Skil,Fkbp1a,Rgs2,Prkch,Arhgap6,Mapk1,Wasf1,Prkar2b,Dgkh,Grm4,Grm5,Smad3,Ptk2b,Kitlg,Nedd4l,Bcr,Lrp12,Oprk1,Dlg3,Cyfip2,Lamb1,Chrm1,Arhgap20,Prkcg,Rbp4,Drd2,Prkcz,Dgki,Braf,Kit,Pdpk1,Prkacb,Adra2b,Fam13b,Ranbp2,Pip5k1a,Rasgrp1,Dusp8,Ywhaz,Arhgap32,Irs2,Ndel1,Prkg2,Cbx6,Rarb,Opn3,Sstr4,Htr2c,Dgkg,Adcy9,Vps29,Dlg2,Grm7,Actr2,Tec,Ptprj,Fkbp5,Kl,Prkcb,Grin1,Madd,Carm1,Rxrg,Arhgef3,Dusp2,Crabp1,Rgs8,Rapgef4,Fst,Oprm1,Dlg4,Bdnf,Bag4,Ptpn7,Rictor,Fmnl1,Rasgef1a,Pik3cg,Wnt10a,Rasal2,Ofd1,Pdgfb,Iqgap3,Rps6ka5,Kalrn,Scai,Dock3,Arhgap10,P2ry1,Gga3,Arhgap33,Plcb1,Rgs10 |
| Egr2,Nab2,Srf,Mapk1,Braf,Irs2,Bdnf,Rps6ka5,Dock3 |
| Egr2,Nab2,Baiap2,Srf,Mapk1,Wasf1,Ptk2b,Kitlg,Cyfip2,Lamb1,Prkcz,Braf,Kit,Pdpk1,Prkacb,Irs2,Tec,Ptprj,Kl,Prkcb,Bdnf,Rictor,Pdgfb,Rps6ka5,Dock3,Gga3 |
| Tiam1,Wipf3,Tiam2,Baiap2,Srf,Arhgap6,Mapk1,Wasf1,Bcr,Cyfip2,Arhgap20,Prkcz,Pdpk1,Fam13b,Ranbp2,Ywhaz,Arhgap32,Ndel1,Actr2,Prkcb,Arhgef3,Dlg4,Fmnl1,Iqgap3,Kalrn,Scai,Arhgap10,Arhgap33 |
| Tiam1,Wipf3,Tiam2,Baiap2,Srf,Arhgap6,Mapk1,Wasf1,Bcr,Cyfip2,Arhgap20,Prkcz,Pdpk1,Fam13b,Ranbp2,Ywhaz,Arhgap32,Ndel1,Actr2,Prkcb,Arhgef3,Dlg4,Fmnl1,Iqgap3,Kalrn,Scai,Arhgap10,Arhgap33 |
| Baiap2,Wasf1,Ptk2b,Cyfip2,Prkcz,Pdpk1,Prkacb,Prkcb,Rictor |
| Gria1,Dlg3,Grin1,Dlg4 |
| Camk4,Gria1,Actn2,Kcnj2,Cacnb4,Gabra4,Prkar2b,Dlg3,Prkcg,Prkacb,Gad1,Gabrb3,Adcy9,Dlg2,Ppfia2,Prkcb,Grin1,Cacna2d3,Syt1,Dlg4,Camk1 |
| Gria1,Actn2,Dlg3,Dlg2,Grin1,Dlg4 |
| Baiap2,Wasf1,Ptk2b,Cyfip2,Prkcz,Pdpk1,Prkacb,Prkcb,Rictor |
| Kcnab1,Kcnh1,Kcnf1,Kcnq5,Kcna5,Kcns2,Kcnv1,Kcnh3,Kcna1,Kcnb1 |
| Fads2,Elovl5,Fads1,Hsd17b4 |
| Fads2,Elovl5,Fads1,Hsd17b4 |
| Col5a3,Mmp2,Tubb2b,Plxnb3,LOC100910732,Rhoc,Rdx,Itgav,Sema6d,Itga9,Col9a3,Dpysl3,Rhoa,Mapk12,Psenen,Dag1,Fyn,Dnm2 |
| Hadha,Hadhb |
| Msmo1,Cyp51,Dhcr7,Sc5d,Hmgcs1,Fdft1 |
| Sdhb,Ogdh,Suclg2,Sdhc |
| Col5a3,Matn4,Bgn,Itgav,Itga9,Col9a3,Dag1 |
| Scd,Mid1ip1,Acot13,Dbi,Acbd4,Acsbg1,Fads2,Elovl5,Acot11,Fads1,Slc25a1,Fasn,Hsd17b4,Hadha,Phyh,Hadhb,Hsd17b12,Cyp2j3 |
| Scd,Acsbg1,Elovl5,Slc25a1,Fasn,Hsd17b12 |
| Fads2,Elovl5,Fads1 |

| |
|---|
| Scd,Abhd4,Msmo1,Mid1ip1,Cd38,Acot13,Tnfaip8,Acss2,Ephx1,Dbi,Ddhd1,Cers4,Cyp51,Sat1,Dhcr7,Abcd4,Amt,Rida,Aldh1l1,Acbd4,Slc6a11,Acsbg1,Fads2,Bckdk,Mmab,Sc5d,Elovl5,Hmgcs1,Pnp,Acot11,Sdhb,B3gnt7,Fads1,Mthfs,Bgn,Slc25a1,Idh1,Fasn,Asrgl1,Prodh1,Agl,Ogdh,Aldh7a1,Oplah,Hsd17b4,Sgpl1,Nudt5,Wipi1,G6pd,Suclg2,Glyctk,Sumf2,Hibadh,Dnph1,Hadha,Nudt12,Slc46a1,Sdhc,Arse,Hsd17b11,Phyh,Psph,Fdft1,Aprt,Hadhb,Lhpp,Glul,Ddah1,Fech,LOC108348061,Ppox,Gstk1,Isca2,Hsd17b12,Cyp2j3,ND6,Tyms,Gcsh,Dnm2 |
| Scd,Abhd4,Msmo1,Mid1ip1,Acot13,Tnfaip8,Dbi,Ddhd1,Cers4,Cyp51,Dhcr7,Acbd4,Acsbg1,Fads2,Sc5d,Elovl5,Hmgcs1,Acot11,Fads1,Slc25a1,Fasn,Hsd17b4,Sgpl1,Wipi1,Sumf2,Hadha,Arse,Hsd17b11,Phyh,Fdft1,Hadhb,Hsd17b12,Cyp2j3 |
| Msmo1,Cyp51,Dhcr7,Sc5d,Hmgcs1,Hsd17b4,Hsd17b11,Fdft1,Hsd17b12 |
| Acot13,Dbi,Acot11,Hadha,Hadhb |
| Col5a3,Mmp2,Tubb2b,Plxnb3,LOC100910732,Rhoc,Rdx,Itgav,Sema6d,Itga9,Col9a3,Dpysl3,Rhoa,Mapk12,Psenen,Dag1,Fyn,Dnm2 |
| Cat,Idh1,Hsd17b4,Ide,Lonp2,Phyh,Gstk1 |
| Pex3,Ldhd,Cat,Idh1,Hsd17b4,Ide,Lonp2,Phyh,Gstk1 |
| Rhoc,Rhoa |
| Plxnb3,LOC100910732,Rhoc,Sema6d,Dpysl3,Rhoa,Fyn |
| Axl,LOC100910732,Ctnnd1,Itgav,Rhoa,Mapk12,Vegfb,Fyn,Mapkap1 |
| Wipi1,Sumf2,Arse |
| Axl,LOC100910732,Ctnnd1,Itgav,Rhoa,Mapk12,Fyn,Mapkap1 |

| Source | Term Name |
| --- | --- |
| REAC | Acetylcholine Neurotransmitter Release Cycle |
| REAC | Activation of Ca-permeable Kainate Receptor |
| REAC | Activation of kainate receptors upon glutamate binding |
| REAC | Activation of NMDA receptors and postsynaptic events |
| REAC | Amine ligand-binding receptors |
| REAC | Antigen processing: Ubiquitination & Proteasome degradation |
| REAC | Axon guidance |
| REAC | Ca-dependent events |
| REAC | Calmodulin induced events |
| REAC | CaM pathway |
| REAC | Cardiac conduction |
| REAC | CREB1 phosphorylation through the activation of Adenylate Cyclase |
| REAC | DAG and IP3 signaling |
| REAC | DARPP-32 events |
| REAC | Disinhibition of SNARE formation |
| REAC | Dopamine Neurotransmitter Release Cycle |
| REAC | Effects of PIP2 hydrolysis |
| REAC | Estrogen-stimulated signaling through PRKCZ |
| REAC | G alpha (i) signalling events |
| REAC | G-protein mediated events |
| REAC | GABA synthesis, release, reuptake and degradation |
| REAC | Glucagon signaling in metabolic regulation |
| REAC | Glutamate binding, activation of AMPA receptors and synaptic plasticity |
| REAC | Glutamate Neurotransmitter Release Cycle |
| REAC | Golgi Associated Vesicle Biogenesis |

| | |
|---|---|
| REAC | Hemostasis |
| REAC | Insulin receptor recycling |
| REAC | Integration of energy metabolism |
| REAC | Ion channel transport |
| REAC | Ion homeostasis |
| REAC | Ionotropic activity of kainate receptors |
| REAC | LGI-ADAM interactions |
| REAC | MAPK family signaling cascades |
| REAC | Membrane Trafficking |
| REAC | Muscarinic acetylcholine receptors |
| REAC | Muscle contraction |
| REAC | Nervous system development |
| REAC | Neurexins and neuroligins |
| REAC | Neuronal System |
| REAC | Neurotransmitter receptors and postsynaptic signal transmission |
| REAC | Neurotransmitter release cycle |
| REAC | Norepinephrine Neurotransmitter Release Cycle |
| REAC | Opioid Signalling |

| | |
|---|---|
| REAC | Phospholipid metabolism |
| REAC | PI Metabolism |
| REAC | PKA activation |
| REAC | PKA activation in glucagon signalling |
| REAC | PKA-mediated phosphorylation of CREB |
| REAC | Platelet activation, signaling and aggregation |
| REAC | Platelet calcium homeostasis |
| REAC | Platelet homeostasis |
| REAC | PLC beta mediated events |
| REAC | Post NMDA receptor activation events |
| REAC | Potassium Channels |
| REAC | Progressive trimming of alpha-1,2-linked mannose residues from Man9/8/7GlcNAc2 to produce Man5GlcNAc2 |
| REAC | Protein-protein interactions at synapses |
| REAC | Rap1 signalling |
| REAC | Reduction of cytosolic Ca++ levels |
| REAC | Regulation of insulin secretion |
| REAC | Serotonin Neurotransmitter Release Cycle |
| REAC | Signaling by NTRK1 (TRKA) |
| REAC | Signaling by NTRKs |
| REAC | Signaling by Receptor Tyrosine Kinases |
| REAC | Signaling by Rho GTPases |

| | |
|---|---|
| REAC | Signaling by Rho GTPases, Miro GTPases and RHOBTB3 |
| REAC | Synaptic adhesion-like molecules |
| REAC | Synthesis of PI |
| REAC | Synthesis of PIPs at the plasma membrane |
| REAC | Trafficking of AMPA receptors |
| REAC | Trafficking of GluR2-containing AMPA receptors |
| REAC | trans-Golgi Network Vesicle Budding |
| REAC | Transmission across Chemical Synapses |
| REAC | Unblocking of NMDA receptors, glutamate binding and activation |
| REAC | VEGFR2 mediated cell proliferation |
| REAC | Voltage gated Potassium channels |
| REAC | WNT5A-dependent internalization of FZD4 |
| REAC | Antiviral mechanism by IFN-stimulated genes |
| REAC | Apoptosis |
| REAC | Apoptotic factor-mediated response |
| REAC | Axon guidance |
| REAC | C-type lectin receptors (CLRs) |
| REAC | Cell junction organization |
| REAC | Cell-Cell communication |
| REAC | Cell-extracellular matrix interactions |
| REAC | Cellular response to chemical stress |
| REAC | Cyclin D associated events in G1 |
| REAC | Cytokine Signaling in Immune system |

| | |
|---|---|
| REAC | Deactivation of the beta-catenin transactivating complex |
| REAC | Death Receptor Signalling |
| REAC | Detoxification of Reactive Oxygen Species |
| REAC | DEx/H-box helicases activate type I IFN and inflammatory cytokines production |
| REAC | ECM proteoglycans |
| REAC | Extracellular matrix organization |
| REAC | Factors involved in megakaryocyte development and platelet production |
| REAC | G alpha (12/13) signalling events |
| REAC | G1 Phase |
| REAC | Glycosaminoglycan metabolism |
| REAC | GRB2:SOS provides linkage to MAPK signaling for Integrins |
| REAC | Hemostasis |

| | |
|---|---|
| REAC | Immune System |
| REAC | Inflammasomes |
| REAC | Innate Immune System |
| REAC | Integrin signaling |
| REAC | Interleukin-20 family signaling |
| REAC | Lysosomal oligosaccharide catabolism |
| REAC | MAP2K and MAPK activation |
| REAC | MET activates PTK2 signaling |
| REAC | MET promotes cell motility |
| REAC | Mitotic G1 phase and G1/S transition |
| REAC | Nervous system development |

| | |
|---|---|
| REAC | Neutrophil degranulation |
| REAC | Nicotinamide salvaging |
| REAC | Nicotinate metabolism |
| REAC | Nucleotide-binding domain, leucine rich repeat containing receptor (NLR) signaling pathways |
| REAC | p130Cas linkage to MAPK signaling for integrins |
| REAC | p75 NTR receptor-mediated signalling |
| REAC | PECAM1 interactions |
| REAC | Platelet activation, signaling and aggregation |
| REAC | Platelet degranulation |
| REAC | Programmed Cell Death |
| REAC | Regulated Necrosis |
| REAC | Regulation of cytoskeletal remodeling and cell spreading by IPP complex components |
| REAC | Regulation of necroptotic cell death |
| REAC | Regulation of RUNX1 Expression and Activity |
| REAC | Regulation of TNFR1 signaling |
| REAC | Response to elevated platelet cytosolic Ca2+ |
| REAC | RIP-mediated NFkB activation via ZBP1 |
| REAC | RIPK1-mediated regulated necrosis |
| REAC | RUNX3 regulates WNT signaling |
| REAC | Signaling by MET |
| REAC | Signaling by Non-Receptor Tyrosine Kinases |
| REAC | Signaling by PTK6 |

| | |
|---|---|
| REAC | Signaling by Rho GTPases |
| REAC | Signaling by Rho GTPases, Miro GTPases and RHOBTB3 |
| REAC | Signaling by VEGF |
| REAC | SMAC (DIABLO) binds to IAPs |
| REAC | SMAC, XIAP-regulated apoptotic response |
| REAC | SMAC(DIABLO)-mediated dissociation of IAP:caspase complexes |
| REAC | Smooth Muscle Contraction |
| REAC | TNF signaling |
| REAC | TNFR1-induced NFkappaB signaling pathway |
| REAC | TP53 Regulates Transcription of Genes Involved in G1 Cell Cycle Arrest |
| REAC | Transcriptional regulation by RUNX3 |
| REAC | VEGFA-VEGFR2 Pathway |
| REAC | ZBP1(DAI) mediated induction of type I IFNs |

| Term ID | p-value | Directionality |
|---|---|---|
| REAC:R-RNO-264642 | 0.0021 | Up |
| REAC:R-RNO-451308 | 0.0249 | Up |
| REAC:R-RNO-451326 | 0.0249 | Up |
| REAC:R-RNO-442755 | 2.9781E-06 | Up |
| REAC:R-RNO-375280 | 0.0054 | Up |
| REAC:R-RNO-983168 | 0.0325 | Up |
| REAC:R-RNO-422475 | 0.0371 | Up |
| REAC:R-RNO-111996 | 1.4385E-06 | Up |
| REAC:R-RNO-111933 | 5.5991E-06 | Up |
| REAC:R-RNO-111997 | 5.5991E-06 | Up |
| REAC:R-RNO-5576891 | 0.0003 | Up |
| REAC:R-RNO-442720 | 0.0054 | Up |
| REAC:R-RNO-1489509 | 5.5991E-06 | Up |
| REAC:R-RNO-180024 | 0.0051 | Up |
| REAC:R-RNO-114516 | 0.0340 | Up |
| REAC:R-RNO-212676 | 0.0003 | Up |
| REAC:R-RNO-114508 | 0.0051 | Up |
| REAC:R-RNO-9634635 | 0.0131 | Up |
| REAC:R-RNO-418594 | 0.0468 | Up |
| REAC:R-RNO-112040 | 5.5991E-06 | Up |
| REAC:R-RNO-888590 | 0.0001 | Up |
| REAC:R-RNO-163359 | 0.0131 | Up |
| REAC:R-RNO-399721 | 6.9077E-06 | Up |
| REAC:R-RNO-210500 | 0.0188 | Up |
| REAC:R-RNO-432722 | 0.0038 | Up |

| | | |
|---|---|---|
| REAC:R-RNO-109582 | 0.0340 | Up |
| REAC:R-RNO-77387 | 0.0449 | Up |
| REAC:R-RNO-163685 | 0.0012 | Up |
| REAC:R-RNO-983712 | 0.0246 | Up |
| REAC:R-RNO-5578775 | 0.0058 | Up |
| REAC:R-RNO-451306 | 0.0249 | Up |
| REAC:R-RNO-5682910 | 0.0008 | Up |
| REAC:R-RNO-5683057 | 0.0267 | Up |
| REAC:R-RNO-199991 | 0.0155 | Up |
| REAC:R-RNO-390648 | 0.0340 | Up |
| REAC:R-RNO-397014 | 0.0152 | Up |
| REAC:R-RNO-9675108 | 0.0395 | Up |
| REAC:R-RNO-6794361 | 0.0001 | Up |
| REAC:R-RNO-112316 | 1.4360E-27 | Up |
| REAC:R-RNO-112314 | 3.8470E-11 | Up |
| REAC:R-RNO-112310 | 0.0002 | Up |
| REAC:R-RNO-181430 | 0.0037 | Up |
| REAC:R-RNO-111885 | 0.0002 | Up |

| | | |
|---|---|---|
| REAC:R-RNO-1483257 | 3.9774E-05 | Up |
| REAC:R-RNO-1483255 | 0.0001 | Up |
| REAC:R-RNO-163615 | 0.0021 | Up |
| REAC:R-RNO-164378 | 0.0054 | Up |
| REAC:R-RNO-111931 | 0.0021 | Up |
| REAC:R-RNO-76002 | 0.0194 | Up |
| REAC:R-RNO-418360 | 0.0340 | Up |
| REAC:R-RNO-418346 | 0.0024 | Up |
| REAC:R-RNO-112043 | 5.5991E-06 | Up |
| REAC:R-RNO-438064 | 0.0086 | Up |
| REAC:R-RNO-1296071 | 8.7214E-08 | Up |
| REAC:R-RNO-964827 | 0.0118 | Up |
| REAC:R-RNO-6794362 | 1.6063E-05 | Up |
| REAC:R-RNO-392517 | 0.0037 | Up |
| REAC:R-RNO-418359 | 0.0010 | Up |
| REAC:R-RNO-422356 | 0.0075 | Up |
| REAC:R-RNO-181429 | 0.0008 | Up |
| REAC:R-RNO-187037 | 0.0046 | Up |
| REAC:R-RNO-166520 | 0.0026 | Up |
| REAC:R-RNO-9006934 | 0.0106 | Up |
| REAC:R-RNO-194315 | 0.0028 | Up |

| | | |
|---|---|---|
| REAC:R-RNO-9716542 | 0.0028 | Up |
| REAC:R-RNO-8849932 | 0.0473 | Up |
| REAC:R-RNO-1483226 | 0.0054 | Up |
| REAC:R-RNO-1660499 | 0.0005 | Up |
| REAC:R-RNO-399719 | 6.9077E-06 | Up |
| REAC:R-RNO-416993 | 0.0002 | Up |
| REAC:R-RNO-199992 | 0.0045 | Up |
| REAC:R-RNO-112315 | 6.5202E-16 | Up |
| REAC:R-RNO-438066 | 0.0008 | Up |
| REAC:R-RNO-5218921 | 0.0051 | Up |
| REAC:R-RNO-1296072 | 2.9781E-06 | Up |
| REAC:R-RNO-5099900 | 0.0054 | Up |
| ral mechanism by IFN-stimulated g | 0.0430 | Down |
| Apoptosis | 0.0441 | Down |
| Apoptotic factor-mediated respons | 0.0430 | Down |
| Axon guidance | 0.0443 | Down |
| C-type lectin receptors (CLRs) | 0.0409 | Down |
| Cell junction organization | 0.0430 | Down |
| Cell-Cell communication | 0.0315 | Down |
| Cell-extracellular matrix interaction | 0.0161 | Down |
| Cellular response to chemical stres | 0.0315 | Down |
| Cyclin D associated events in G1 | 0.0028 | Down |
| ytokine Signaling in Immune syste | 0.0161 | Down |

| | | |
|---|---|---|
| n of the beta-catenin transactivati | 0.0315 | Down |
| Death Receptor Signalling | 0.0005 | Down |
| oxification of Reactive Oxygen Spe | 0.0315 | Down |
| ctivate type I IFN and inflammator | 0.0141 | Down |
| ECM proteoglycans | 0.0043 | Down |
| Extracellular matrix organization | 0.0387 | Down |
| megakaryocyte development and | 0.0416 | Down |
| G alpha (12/13) signalling events | 0.0491 | Down |
| G1 Phase | 0.0028 | Down |
| Glycosaminoglycan metabolism | 0.0161 | Down |
| rovides linkage to MAPK signaling | 0.0252 | Down |
| Hemostasis | 0.0001 | Down |

| | | |
|---|---|---|
| Immune System | 2.6691E-08 | Down |
| Inflammasomes | 0.0161 | Down |
| Innate Immune System | 4.8643E-09 | Down |
| Integrin signaling | 0.0374 | Down |
| Interleukin-20 family signaling | 0.0430 | Down |
| sosomal oligosaccharide catabolis | 0.0315 | Down |
| MAP2K and MAPK activation | 0.0161 | Down |
| MET activates PTK2 signaling | 0.0441 | Down |
| MET promotes cell motility | 0.0059 | Down |
| litotic G1 phase and G1/S transitio | 0.0161 | Down |
| Nervous system development | 0.0337 | Down |

| | | |
|---|---|---|
| Neutrophil degranulation | 4.8643E-09 | Down |
| Nicotinamide salvaging | 0.0161 | Down |
| Nicotinate metabolism | 0.0374 | Down |
| eucine rich repeat containing recep | 0.0026 | Down |
| as linkage to MAPK signaling for in | 0.0161 | Down |
| 75 NTR receptor-mediated signallin | 0.0161 | Down |
| PECAM1 interactions | 0.0315 | Down |
| let activation, signaling and aggreg | 0.0001 | Down |
| Platelet degranulation | 0.0026 | Down |
| Programmed Cell Death | 0.0315 | Down |
| Regulated Necrosis | 0.0315 | Down |
| remodeling and cell spreading by | 0.0315 | Down |
| Regulation of necroptotic cell death | 0.0315 | Down |
| ation of RUNX1 Expression and A | 0.0315 | Down |
| Regulation of TNFR1 signaling | 0.0371 | Down |
| onse to elevated platelet cytosolic | 0.0023 | Down |
| P-mediated NFkB activation via ZB | 0.0374 | Down |
| RIPK1-mediated regulated necrosis | 0.0315 | Down |
| RUNX3 regulates WNT signaling | 0.0315 | Down |
| Signaling by MET | 0.0315 | Down |
| ling by Non-Receptor Tyrosine Kin | 0.0315 | Down |
| Signaling by PTK6 | 0.0315 | Down |

| | | |
|---|---|---|
| Signaling by Rho GTPases | 0.0315 | Down |
| Rho GTPases, Miro GTPases an | 0.0315 | Down |
| Signaling by VEGF | 0.0315 | Down |
| SMAC (DIABLO) binds to IAPs | 0.0315 | Down |
| C, XIAP-regulated apoptotic resp | 0.0220 | Down |
| -mediated dissociation of IAP:cas | 0.0315 | Down |
| Smooth Muscle Contraction | 0.0289 | Down |
| TNF signaling | 0.0430 | Down |
| 1-induced NFkappaB signaling pa | 0.0374 | Down |
| anscription of Genes Involved in G | 0.0141 | Down |
| ranscriptional regulation by RUNX | 0.0388 | Down |
| VEGFA-VEGFR2 Pathway | 0.0337 | Down |
| (DAI) mediated induction of type I | 0.0485 | Down |

| Intersection |
| --- |
| Ppfia2,Vamp2,Syt1,Snap25,Stxbp1,Rab3a |
| Dlg4,Dlg3,Ncald,Grik5 |
| Dlg4,Dlg3,Ncald,Grik5 |
| Gria1,Camk4,Actn2,Dlg4,Dlg2,Rps6ka2,Dlg3,Grin1,Prkar2b,Prkar2a,Prkar1b,Nefl,Prkacb |
| Htr1b,Chrm1,Hrh3,Drd2,Chrm2,Adra2b,Htr1d,Adra2c,Chrm3 |
| Lmo7,Fbxl16,Rnf144b,Arel1,Hectd1,Klhl2,Fbxo41,Nedd4l,Pja2,Trim32,Klhl3,Asb1,Ube2o,Herc1,Hace1,Herc2,Asb2,Lonrf1,Hecw2,Ltn1,Rnf6,Trip12,Zbtb16,Fbxl18,Rnf126,Tpp2,Dcaf1,Pja1,Ube2k,Fbxw4,Ubr1,Znrf1,Trim9,Klhl11,Cdc34,Fbxl19 |
| Tiam1,Cdk5r1,Mapk1,Grin1,Rap1gap,Kalrn,Irs2,Sptan1,Prkca,Sptbn4,Epha6,Ap2a1,Actr2,Dok6,Plxna2,Efna3,Dnm1,Hsp90ab1,Plxna4,Epha7,Ap2b1,Lypla2,Ap2m1,Hras,Arhgef7,Cdk5,Prkacb,Dscaml1,St8sia4 |
| Camk4,Pde1b,Adcy9,Mapk1,Prkcg,Prkca,Adcy1,Prkar2b,Prkar2a,Prkar1b,Prkacb |
| Camk4,Pde1b,Adcy9,Prkcg,Prkca,Adcy1,Prkar2b,Prkar2a,Prkar1b,Prkacb |
| Camk4,Pde1b,Adcy9,Prkcg,Prkca,Adcy1,Prkar2b,Prkar2a,Prkar1b,Prkacb |
| Atp2b1,Scn4b,Kcnj2,Atp1a1,Kcnk1,Scn3b,Scn2b,Atp2a2,Atp2b2,Kcnk2,Fgf13,Kcnj11,Akap9,Slc8a1,Atp1a3,Atp2b3,Atp2b4,Kcnk4,Atp1b1,Kcnk9 |
| Prkar2b,Prkar2a,Prkar1b,Prkacb |
| Camk4,Pde1b,Adcy9,Prkcg,Prkca,Adcy1,Prkar2b,Prkar2a,Prkar1b,Prkacb |
| Prkar2b,Ppp1r1b,Prkar2a,Prkar1b,Cdk5,Pde4d,Prkacb |
| Prkcg,Prkcb,Prkca |
| Ppfia2,Lin7b,Vamp2,Syt1,Syn1,Snap25,Stxbp1,Rab3a |
| Dgki,Dgkh,Dgkg,Dgka,Dgkz,Prkch,Dagla |
| Pdpk1,Mapk1,Prkcz,Hras |
| Htr1b,Camk4,Penk,Pde1b,Rgs14,Adcy9,Grm4,Mapk1,Rgs9,Mchr1,Oprk1,Lpl,Lrp12,Prkcg,Plcb1,Sstr2,Prkca,Adcy1,Prkar2b,Chrm2,Adra2b,Htr1d,Opn3,Adra2c,Rbp4,Ppp1r1b,Prkar2a,Prkar1b,Lrp8,Rgs17,Sstr3,Cdk5,Pde4d,Prkacb,Gnaz,Grm7,Rgs8 |
| Camk4,Pde1b,Adcy9,Mapk1,Prkcg,Plcb1,Prkca,Adcy1,Prkar2b,Prkar2a,Prkar1b,Prkacb,Gnaz |
| Dnajc5,Gad1,Hspa8,Vamp2,Syt1,Slc32a1,Snap25,Stxbp1,Rab3a |
| Prkar2b,Prkar2a,Prkar1b,Prkacb |
| Gria1,Dlg4,Prkcg,Cacng3,Prkcb,Epb41l1,Prkca,Ap2a1,Ap2b1,Ap2m1,Cacng2 |
| Ppfia2,Vamp2,Syt1,Snap25,Stxbp1,Rab3a |
| Tpd52l1,Ap1s1,Ocrl,Hspa8,Golgb1,Dnajc6,Vamp2,Napa,Arf1,Tpd52,Igf2r,Arrb1 |

| |
|---|
| Pde10a,Actn1,Pdpk1,Pde1b,Atp2b1,Actn2,Dgki,Rapgef4,Mapk1,Dgkh,Dgkg,Dock4,Prkcg,Prkcb,Gp1bb,Atp2a2,Prkca,Dgka,Atp2b2,Rasgrp1,Dgkz,Prkar2b,Mafg,Epcam,Kif21a,Adra2b,Slc8a1,Kif1b,Kifc2,Prkch,Ywhaz,Kif3b,Pdgfb,Kifap3,Kif1a,Adra2c,Slc7a8,Rab27b,Ola1,Dagla,Atp2b3,Atp2b4,Prkar2a,Prkar1b,Brpf3,Lrp8,Tgfb3,Hras,Atp1b1,Pcyox1l,Prkg2,Habp4,Prkacb,Dock3,Arrb1 |
| Atp6v1a,Atp6v1b2,Atp6v1g2,Atp6v0e2,Atp6v0c,Atp6v1c1 |
| Rapgef4,Abcc8,Plcb1,Kcng2,Prkca,Kcnj11,Prkar2b,Cacnb3,Adra2c,Prkag2,Prkar2a,Kcnb1,Prkar1b,Prkacb |
| Atp2b1,Ano3,Atp1a1,Nedd4l,Atp2a2,Atp2b2,Atp9a,Wnk2,Atp1a3,Atp6v1a,Atp6v1b2,Atp2b3,Atp2b4,Ano2,Atp6v1g2,Atp6v0e2,Atp1b1,Wnk3,Wnk4,Atp6v0c,Atp6v1c1,Asic2 |
| Atp2b1,Atp1a1,Atp2a2,Atp2b2,Kcnj11,Slc8a1,Atp1a3,Atp2b3,Atp2b4,Atp1b1 |
| Dlg4,Dlg3,Ncald,Grik5 |
| Lgi1,Dlg4,Cacng3,Stx1b,Adam22,Adam23,Cacng2 |
| Actn2,Cnksr2,Dlg4,Braf,Dusp8,Mapk1,Dlg2,Dlg3,Grin1,Kalrn,Rasgef1a,Irs2,Sptan1,Kitlg,Rasgrp1,Cdc42ep3,Sptbn4,Shc2,Xpo1,Ptpn3,Pdgfb,Dusp2,Spred2,Ksr2,Hras,Prkg2,Nefl,Prkacb,Kit,Abhd17a,Spred3,Ppp1cc,Arrb1 |
| Gria1,Tpd52l1,Clvs1,LOC100910792,Arf3,Sec16a,Sgip1,Ubqln2,Madd,Sptan1,Tsc1,Sptbn4,Ap2a1,Ap1s1,Actr2,Mia3,Ulk1,Clvs2,Kif21a,Chrm2,Ocrl,Reps2,Kif1b,Hspa8,Kifc2,Cnih2,Golgb1,Dnm1,Kif3b,Kifap3,Dnajc6,Kif1a,Vamp2,Synj1,Syt1,Napa,Rab27b,Arfip2,Arf1,Stx16,Sbf1,Trappc13,Ap2b1,Dynll2,Tbc1d16,Rab6b,Cyth2,Ap2m1,Napg,Rab3a,Dynll1,Tpd52,Igf2r,Pafah1b1,Trappc12,Ric1,Wnk4,Arf5,Cnih3,Arrb1 |
| Chrm1,Chrm2,Chrm3 |
| Atp2b1,Actn2,Scn4b,Kcnj2,Atp1a1,Kcnk1,Scn3b,Scn2b,Atp2a2,Atp2b2,Kcnk2,Fgf13,Kcnj11,Akap9,Tpm1,Slc8a1,Atp1a3,Atp2b3,Atp2b4,Kcnk4,Atp1b1,Kcnk9 |
| Tiam1,Cdk5r1,Mapk1,Grin1,Rap1gap,Kalrn,Irs2,Sptan1,Prkca,Sptbn4,Epha6,Ap2a1,Actr2,Dok6,Plxna2,Efna3,Dnm1,Hsp90ab1,Plxna4,Epha7,Ap2b1,Lypla2,Ap2m1,Hras,Arhgef7,Cdk5,Prkacb,Dscaml1,St8sia4 |
| Homer1,Dlg4,Dlg2,Dlg3,Epb41l1,Grm5,Lrrtm3,LRRTM1,Shank2,Dlgap1,Shank1 |
| Gria1,Camk4,Gabra4,Actn2,Cacnb4,Kcnab1,Adcy9,Homer1,Dlg4,Kcnj2,Kcnh1,Kcnq5,Kcnf1,Gabrb3,Dlg2,Kcnh3,Rps6ka2,Dlg3,Dnajc5,Kcns2,Kcnk1,Abcc8,Prkcg,Cacng3,Grin1,Prkcb,Epb41l1,Kcng2,Grm5,Prkca,Adcy1,Lrrtm3,Kcna5,Kcnj3,Kcnk2,Kcnj11,Ap2a1,Prkar2b,Slitrk5,Gad1,Kcnma1,Ppfia2,Chrnb2,LRRTM1,Cacnb3,Hspa8,Shank2,Dlgap1,Lin7b,Kcna1,Kcnab2,Vamp2,Shank1,Syt1,Ncald,Kcnq3,Kcnv1,Syn1,Slitrk3,Panx1,Kcnh7,Kcnn1,Prkar2a,Slc32a1,Kcnb1,Ap2b1,Grik5,Kcnk4,Prkar1b,Ap2m1,Snap25,Stxbp1,Kcnj9,Rab3a,Glrb,Rtn3,Nefl,Prkacb,Cacng2,Gabra5,Kcnk9 |
| Gria1,Camk4,Gabra4,Actn2,Adcy9,Dlg4,Kcnj2,Gabrb3,Dlg2,Rps6ka2,Dlg3,Prkcg,Cacng3,Grin1,Prkcb,Epb41l1,Prkca,Adcy1,Kcnj3,Ap2a1,Prkar2b,Chrnb2,Ncald,Prkar2a,Ap2b1,Grik5,Prkar1b,Ap2m1,Kcnj9,Glrb,Nefl,Prkacb,Cacng2,Gabra5 |
| Dnajc5,Gad1,Ppfia2,Hspa8,Lin7b,Vamp2,Syt1,Syn1,Slc32a1,Snap25,Stxbp1,Rab3a |
| Ppfia2,Vamp2,Syt1,Snap25,Stxbp1,Rab3a |
| Camk4,Pde1b,Adcy9,Mapk1,Prkcg,Plcb1,Prkca,Adcy1,Prkar2b,Ppp1r1b,Prkar2a,Prkar1b,Cdk5,Pde4d,Prkacb,Gnaz |

| |
|---|
| Pitpnm3,Mtmr12,Arf3,Pitpnm2,Plekha5,Osbpl8,Cpne6,Mtmr4,Ddhd2,Mtmr1,Dgat2,Lpin2,Ocrl,Pip5k1a,Lpcat4,Pip4k2b,Synj1,Miga1,Pten,Inpp5j,Mtmr7,Arf1,Sbf1,Pikfyve,Gpd1l,Cds1,Mtmr6,Pi4k2a,Cpne7,Pitpnm1,Osbpl10,Agk,Lpgat1,Pip4k2c |
| Mtmr12,Arf3,Plekha5,Mtmr4,Mtmr1,Ocrl,Pip5k1a,Pip4k2b,Synj1,Pten,Inpp5j,Mtmr7,Arf1,Sbf1,Pikfyve,Mtmr6,Pi4k2a,Pip4k2c |
| Adcy9,Adcy1,Prkar2b,Prkar2a,Prkar1b,Prkacb |
| Prkar2b,Prkar2a,Prkar1b,Prkacb |
| Adcy9,Adcy1,Prkar2b,Prkar2a,Prkar1b,Prkacb |
| Actn1,Pdpk1,Actn2,Dgki,Rapgef4,Mapk1,Dgkh,Dgkg,Prkcg,Prkcb,Gp1bb,Prkca,Dgka,Rasgrp1,Dgkz,Adra2b,Prkch,Ywhaz,Pdgfb,Adra2c,Rab27b,Ola1,Dagla,Brpf3,Tgfb3,Pcyox1l,Habp4,Arrb1 |
| Atp2b1,Atp2a2,Atp2b2,Slc8a1,Atp2b3,Atp2b4 |
| Pde10a,Pde1b,Atp2b1,Atp2a2,Atp2b2,Slc8a1,Atp2b3,Atp2b4,Lrp8,Prkg2 |
| Camk4,Pde1b,Adcy9,Mapk1,Prkcg,Plcb1,Prkca,Adcy1,Prkar2b,Prkar2a,Prkar1b,Prkacb,Gnaz |
| Camk4,Rps6ka2,Prkar2b,Prkar2a,Prkar1b,Prkacb |
| Kcnab1,Kcnj2,Kcnh1,Kcnq5,Kcnf1,Kcnh3,Kcns2,Kcnk1,Abcc8,Kcng2,Kcna5,Kcnj3,Kcnk2,Kcnj11,Kcnma1,Kcna1,Kcnab2,Kcnq3,Kcnv1,Kcnh7,Kcnn1,Kcnb1,Kcnk4,Kcnj9,Kcnk9 |
| Man1a1,Man1c1,Man1a2 |
| Gria1,Homer1,Dlg4,Dlg2,Dlg3,Grin1,Epb41l1,Grm5,Lrrtm3,Slitrk5,Ppfia2,LRRTM1,Shank2,Dlgap1,Shank1,Slitrk3,Rtn3 |
| Rapgef4,Rap1gap,Rasgrp1,Ywhaz,Rap1gap2,Prkacb |
| Atp2b1,Atp2a2,Atp2b2,Slc8a1,Atp2b3,Atp2b4 |
| Rapgef4,Abcc8,Plcb1,Kcng2,Prkca,Kcnj11,Cacnb3,Adra2c,Kcnb1,Prkacb |
| Ppfia2,Vamp2,Syt1,Syn1,Snap25,Stxbp1,Rab3a |
| Nab2,Egr2,Braf,Mapk1,Rps6ka2,Irs2,Srf,Ap2a1,Shc2,Rapgef1,Ap2b1,Ap2m1,Hras |
| Nab2,Egr2,Braf,Mapk1,Rps6ka2,Irs2,Srf,Bdnf,Ap2a1,Shc2,Rapgef1,Ap2b1,Ap2m1,Hras,Dock3 |
| Baiap2,Ptk2b,Pdpk1,Nab2,Egr2,Cyfip2,Braf,Mapk1,Prkcz,Gga3,Wasf1,Rps6ka2,Lamb1,Mtor,Prkcb,Irs2,Prkca,Srf,Ptpru,Bdnf,Kitlg,Ptprj,Ap2a1,Shc2,Rbfox2,Ptpn3,Pdgfb,Rapgef1,Spred2,Atp6v1a,Atp6v1b2,Atp6v1g2,Ap2b1,Ap2m1,Hras,Atp6v0e2,Matk,Ptpn18,Arhgef7,Prkacb,Kit,Atp6v0c,Flrt2,Atp6v1c1,Dock3 |
| Wipf3,Baiap2,Tiam1,Pdpk1,Cyfip2,Tiam2,Dlg4,Bcr,Mapk1,Ppp1r12b,Prkcz,Wasf1,Prkcb,Arhgef3,Kalrn,Arhgap32,Rangap1,Fmnl1,Prkca,Ranbp2,Srf,Arhgap10,Arhgap20,Actr2,Ocrl,Xpo1,Arhgap6,Ywhaz,Arhgap33,Rhot2,Rhobtb2,Mcf2l,Arhgap39,Zwint,Lin7b,Plekhg5,Hsp90ab1,Scai,Clip1,Dynll2,Ophn1,Ndel1,Dynll1,Arhgap26,Arhgef7,Iqgap3,Pafah1b1,Arhgap44,Ywhag,Clasp1,Ppp1cc |

| |
|---|
| Wipf3,Baiap2,Tiam1,Pdpk1,Cyfip2,Tiam2,Dlg4,Bcr,Mapk1,Ppp1r12b,Prkcz,Wasf1,Prkcb,Arhgef3,Kalrn,Arhgap32,Rangap1,Fmnl1,Prkca,Ranbp2,Srf,Arhgap10,Arhgap20,Actr2,Ocrl,Xpo1,Arhgap6,Ywhaz,Arhgap33,Rhot2,Rhobtb2,Mcf2l,Arhgap39,Zwint,Lin7b,Plekhg5,Hsp90ab1,Scai,Clip1,Dynll2,Ophn1,Ndel1,Dynll1,Arhgap26,Arhgef7,Iqgap3,Pafah1b1,Arhgap44,Ywhag,Clasp1,Ppp1cc |
| Gria1,Dlg4,Dlg3,Grin1,Rtn3 |
| Pitpnm3,Pitpnm2,Cds1,Pitpnm1 |
| Plekha5,Mtmr1,Ocrl,Pip5k1a,Pip4k2b,Synj1,Pten,Inpp5j,Arf1,Mtmr6,Pi4k2a,Pip4k2c |
| Gria1,Dlg4,Prkcg,Cacng3,Prkcb,Epb41l1,Prkca,Ap2a1,Ap2b1,Ap2m1,Cacng2 |
| Gria1,Prkcg,Prkcb,Prkca,Ap2a1,Ap2b1,Ap2m1 |
| Tpd52l1,Clvs1,Ap1s1,Clvs2,Ocrl,Hspa8,Golgb1,Dnajc6,Vamp2,Napa,Arf1,Tpd52,Igf2r,Arrb1 |
| Gria1,Camk4,Gabra4,Actn2,Cacnb4,Adcy9,Dlg4,Kcnj2,Gabrb3,Dlg2,Rps6ka2,Dlg3,Dnajc5,Prkcg,Cacng3,Grin1,Prkcb,Epb41l1,Prkca,Adcy1,Kcnj3,Ap2a1,Prkar2b,Gad1,Ppfia2,Chrnb2,Cacnb3,Hspa8,Lin7b,Vamp2,Syt1,Ncald,Syn1,Prkar2a,Slc32a1,Ap2b1,Grik5,Prkar1b,Ap2m1,Snap25,Stxbp1,Kcnj9,Rab3a,Glrb,Nefl,Prkacb,Cacng2,Gabra5 |
| Gria1,Actn2,Dlg4,Dlg2,Dlg3,Grin1,Nefl |
| Pdpk1,Prkcz,Prkcb,Prkca,Hras |
| Kcnab1,Kcnh1,Kcnq5,Kcnf1,Kcnh3,Kcns2,Kcng2,Kcna5,Kcna1,Kcnab2,Kcnq3,Kcnv1,Kcnh7,Kcnb1 |
| Prkcg,Prkcb,Prkca,Ap2a1,Ap2b1,Ap2m1 |
| Isg15,Usp18,Stat1,Ube2l6,Trim25,Uba7,Mapk3,Rnasel |
| Hist1h1d,Tjp2,Ripk1,Casp7,Bid,Bak1,Cflar,Dbnl,Birc2,Mapk3,Ywhae,Diablo,Xiap,Tjp1,Traf2,Casp3 |
| Casp7,Bak1,Mapk3,Diablo,Xiap,Casp3 |
| Col5a3,Mmp2,Tubb2b,Arpc1b,Gab1,Rdx,Ezr,Rhoc,Plxnb3,Myo9b,Rras,Nras,Rhoa,Tln1,Fyn,Tubb6,Lyn,Ptprc,Myl12a,Mapk3,Src,Col9a3,Tyrobp,Dag1,Col5a2,Ephb4,Col9a2,Itgav,Dpysl3,Mapk12,Itga9,Egfr |
| Psmb10,Psmb8,Psme2,Rela,Relb,Nras,Fcer1g,Fyn,Lyn,Src,Pycard,Psmf1,Nfatc1,Tab1,Nfkb1,Nfatc3,Nfkb2,Nfkbia |
| Lims2,Rsu1,Cdh3,Ctnnd1,Ctnna1,Parvb,Pxn,Flna,Fermt2,Nectin2,Lims1,Cdh4,Nectin3 |
| Lims2,Rsu1,Cdh3,Ctnnd1,Ctnna1,Fyn,Parvb,Pxn,Flna,Skap2,Fermt2,Nectin2,Tyrobp,Lims1,Cdh4,Iqgap1,Nectin3 |
| Lims2,Rsu1,Parvb,Pxn,Flna,Fermt2,Lims1 |
| Ncf1,Prdx1,Cyba,Cat,Sod2,Sod1,Gstp1,Gpx8 |
| Ccnd1,Cdkn1a,Rb1,Cdk6,Cdk4,Ppp2r1b,Lyn,Src,Cdkn1c,Ccne2,Cdkn1b,E2f5,Jak2 |
| Isg15,Usp18,Birc3,Stat1,Psmb10,Ube2l6,Irf9,Trim25,Uba7,Psmb8,Psme2,Csf1,Ltbr,Rela,Eda2r,Stat3,Il13ra1,Jak3,Tnfrsf1b,Relb,Nras,Nod1,Stat2,Csk,Ppp2r1b,Ripk2,Fyn,Lyn,Birc2,Mapk3,Psmf1,Il7r,Mapkapk2,Il10rb,Il12rb1,Myd88,Canx,Xiap,Cdkn1b,Tab1,Tnfrsf11a,Nfkb1,Rnasel,Nfkb2,Jak2,Csf2rb,Nfkbia,Traf2,Casp3 |

| |
|---|
| Sox2,Tcf7l1,Sox9,Tcf7l2,Hdac1,Sox13,Xiap,Tcf7 |
| Birc3,Ripk1,Rela,Mag,Arhgef10,Plekhg2,Rhoa,Ripk2,Arhgef26,Tnfaip3,Akap13,Cflar,Birc2,Ywhae,Myd88,Fgd3,Otud7b,Xiap,Tab1,Sppl2a,Arhgef40,Prex1,Arhgef19,Nfkb1,Nfkbia,Traf2,Casp3 |
| Ncf1,Prdx1,Cyba,Cat,Sod2,Sod1,Gstp1,Gpx8 |
| Irf7,Rela,Myd88,Nfkb1,Nfkb2 |
| Col5a3,Matn4,Itga7,Bgn,Sparc,Col9a3,Dag1,Col5a2,Col9a2,Itgav,Itga9 |
| Col5a3,Mmp2,Matn4,Col22a1,Vcam1,Icam1,Mmp14,Adamts4,Lama4,Pcolce,Col16a1,Itgal,Timp1,Loxl4,Col27a1,Itga7,Ddr2,Itgb2,Bgn,Sparc,Jam3,Col9a3,Ppib,Dag1,Col5a2,Vwf,Sdc4,Col9a2,Itgav,Col12a1,Cep295nl,Itga9,Efemp2,Fbln2,Lama5,Adam12 |
| Tubb2b,Kif18b,Dock8,Kif11,Kif13b,Tubb6,Ehd1,Dock6,Dock5,Rad51b,Kifc1,Ehd2,Carmil1,Cenpe,Hdac1,Kif5b,Hmg20b,Pgd,Dock11,Jak2,Kif26a |
| Arhgef10,Rhoc,Plekhg2,Gna12,Rhoa,Arhgef26,Akap13,Fgd3,Arhgef40,Prex1,Arhgef19 |
| Ccnd1,Cdkn1a,Rb1,Cdk6,Cdk4,Ppp2r1b,Lyn,Src,Cdkn1c,Ccne2,Cdkn1b,E2f5,Jak2 |
| Chst5,Slc35b2,Has2,B3gnt7,Glb1l,Chst7,Bgn,Slc26a2,Chsy1,Galns,Sdc4,Chst14,St3gal4,Prelp,Cspg5,Hs3st3b1,B4galt5,Papss1,Naglu,Hexa |
| Apbb1ip,Rap1a,Tln1,Src,Vwf,Rap1b |
| Cd74,Tubb2b,Gng12,Serping1,Lgals3bp,Fgr,Kif18b,Mag,Lamp2,Dock8,Slc7a5,Tf,Itgal,Apbb1ip,Kif11,P2rx7,Hgf,Vcl,Plek,Slc16a1,Timp1,Rap1a,Cd63,Nras,Lhfpl2,Fcer1g,Kif13b,Rhog,Csk,Rhoa,Rarres2,Itgb2,Maged2,Tln1,Sparc,Fyn,Prcp,Tubb6,Ehd1,Lyn,Jam3,Flna,Wdr1,Rac2,Slc7a11,Cd9,P2ry12,Lcp2,Dock6,Mapk3,Src,Dock5,Rad51b,Srgn,Kifc1,Ehd2,Carmil1,Stxbp3,Vwf,Sdc4,Rap1b,Cenpe,Hdac1,Itih3,Cd84,Gnai3,Sod1,Kif5b,Gnai2,Tagln2,Hmg20b,Pde9a,Pgd,Dock11,Jak2,Kif26a |

| |
|---|
| Cd74,RT1-Da,Isg15,Tubb2b,Usp18,Clec2g,Irf7,Birc3,C1qa,C1r,Serping1,Tlr2,Stat1,Traf7,Casp4,Aim2,Dynlt1,Psmb10,Ube2l6,Man2b1,Rnf213,Vcam1,Irf9,Cfh,Arpc1b,Trim25,Uba7,Psmb8,C2,Pld4,Ifitm3,Icam1,Psme2,Ptx3,Csf1,Ltbr,RT1-S3,Fgr,Rnaset2,Rab5c,Rela,Eda2r,Kif18b,Nckap1l,Stat3,Trim21,Gm2a,Gsdmd,Il13ra1,Herc6,Ncf1,Manba,Jak3,Lamp2,Lgals3,Tnfrsf1b,Pnp,Itgal,Kif11,Cnn2,Tcirg1,P2rx7,Lcn2,Dtx3l,Rab18,Idh1,Vcl,RT1-N3,Tapbp,Myo9b,Tom1,Relb,Cst3,Rap1a,Cd63,Nras,Fcer1g,Bst2,Kif13b,Nod1,Stat2,Rhog,Csk,B2m,Chi3l1,Dpp7,Rhoa,Pstpip1,Slc2a5,Ppp2r1b,Itgb2,Ifitm2,RT1-M3-1,Ube2c,Cnpy3,Rnf114,Ripk2,Aga,Gpr84,Golga7,Ifi30,Fyn,Tnfaip3,Tmem63a,Prcp,Tmbim1,Ptafr,Unc93b1,Tubb6,Ctsz,Aprt,Asah1,Dbnl,Cpne3,Galns,Lyn,Dera,Rac2,Birc2,Ptprc,Cd274,Entpd2,Mgst1,Lcp2,Lamp1,Nlrp3,Mapk3,Src,Nectin2,Wasf2,Tlr1,Tyrobp,Stbd1,Cyba,Pycard,Kifc1,Psmf1,Il7r,Mapkapk2,Tlr7,Fgl2,Slc44a2,Xrcc6,Tmc6,Iqgap1,Cat,Il10rb,Arhgap45,Cdc20,Il12rb1,Fcgr3a,Myd88,Rap1b,Cenpe,Npc2,Canx,Rnf115,Nit2,Prdx4,Itgav,Bin2,Cmtm6,Nfatc1,Ctsc,Sipa1,Rab7a,Mapk12,Cep295nl,Nlrp1a,Kif5b,Spsb1,Fbxo6,Anapc13,Xiap,Gstp1,Cdkn1b,Tab1,Rasgrp3,Blnk,Fbxo7,Cd33,Cpped1,Tspan14,Tlr9,Tnfrsf11a,Nme2,Icoslg,Pgd,Nfkb1,Rnasel,Nfatc3,Nfkb2,Jak2,Csf2rb,Nfkbia,Kif26a,Traf2,Casp3,Arl8a,Anapc10 |
| Aim2,P2rx7,Pstpip1,Nlrp3,Pycard,Nlrp1a |
| Irf7,Birc3,C1qa,C1r,Serping1,Tlr2,Casp4,Aim2,Dynlt1,Psmb10,Man2b1,Cfh,Arpc1b,Psmb8,C2,Pld4,Psme2,Ptx3,RT1-S3,Fgr,Rnaset2,Rab5c,Rela,Nckap1l,Trim21,Gm2a,Gsdmd,Ncf1,Manba,Lamp2,Lgals3,Tnfrsf1b,Pnp,Itgal,Cnn2,Tcirg1,P2rx7,Lcn2,Rab18,Idh1,Vcl,RT1-N3,Myo9b,Tom1,Relb,Cst3,Rap1a,Cd63,Nras,Fcer1g,Bst2,Nod1,Rhog,B2m,Chi3l1,Dpp7,Rhoa,Pstpip1,Slc2a5,Ppp2r1b,Itgb2,RT1-M3-1,Cnpy3,Ripk2,Aga,Gpr84,Golga7,Fyn,Tnfaip3,Tmem63a,Prcp,Tmbim1,Ptafr,Unc93b1,Ctsz,Aprt,Asah1,Dbnl,Cpne3,Galns,Lyn,Dera,Rac2,Birc2,Ptprc,Mgst1,Lcp2,Lamp1,Nlrp3,Mapk3,Src,Wasf2,Tlr1,Tyrobp,Stbd1,Cyba,Pycard,Psmf1,Mapkapk2,Tlr7,Fgl2,Slc44a2,Xrcc6,Tmc6,Iqgap1,Cat,Arhgap45,Fcgr3a,Myd88,Rap1b,Npc2,Nit2,Prdx4,Itgav,Bin2,Cmtm6,Nfatc1,Ctsc,Rab7a,Mapk12,Cep295nl,Nlrp1a,Gstp1,Tab1,Cpped1,Tspan14,Tlr9,Nme2,Nfkb1,Nfatc3,Nfkb2,Nfkbia,Arl8a |
| Apbb1ip,Rap1a,Csk,Tln1,Src,Vwf,Rap1b |
| Stat1,Stat3,Jak3,Stat2,Il10rb,Jak2 |
| Man2b1,Manba,Man2b2 |
| Apbb1ip,Vcl,Rap1a,Nras,Csk,Tln1,Mapk3,Src,Iqgap1,Vwf,Rap1b |
| Col5a3,Lama4,Hgf,Col27a1,Src,Col5a2,Lama5 |
| Col5a3,Gab1,Lama4,Hgf,Rap1a,Col27a1,Tns3,Src,Col5a2,Rap1b,Lama5 |
| Ccnd1,Cdkn1a,Rb1,Psmb10,Mcm10,Psmb8,Psme2,Cdk6,Mcm3,Cdk4,Mcm5,Mcm2,Ppp2r1b,Lyn,Src,Psmf1,Cdkn1c,Hdac1,Ccne2,Cdkn1b,E2f5,Ccna2,Jak2 |
| Col5a3,Mmp2,Tubb2b,Arpc1b,Gab1,Rdx,Ezr,Rhoc,Plxnb3,Myo9b,Rras,Nras,Rhoa,Adgrg6,Tln1,Fyn,Tubb6,Lyn,Ptprc,Myl12a,Mapk3,Src,Col9a3,Tyrobp,Dag1,Col5a2,Ephb4,Col9a2,Itgav,Dpysl3,Mapk12,Itga9,Egfr |

| |
|---|
| Tlr2,Dynlt1,Man2b1,Ptx3,RT1-S3,Fgr,Rnaset2,Rab5c,Nckap1l,Gm2a,Gsdmd,Manba,Lamp2,Lgals3,Tnfrsf1b,Pnp,Itgal,Cnn2,Tcirg1,Lcn2,Rab18,Idh1,Vcl,RT1-N3,Tom1,Cst3,Rap1a,Cd63,Nras,Fcer1g,Bst2,Rhog,B2m,Chi3l1,Dpp7,Rhoa,Slc2a5,Itgb2,RT1-M3-1,Aga,Gpr84,Golga7,Tmem63a,Prcp,Tmbim1,Ptafr,Ctsz,Aprt,Asah1,Dbnl,Cpne3,Galns,Dera,Ptprc,Mgst1,Lamp1,Tyrobp,Stbd1,Cyba,Pycard,Fgl2,Slc44a2,Xrcc6,Tmc6,Iqgap1,Cat,Arhgap45,Fcgr3a,Rap1b,Npc2,Nit2,Prdx4,Itgav,Bin2,Cmtm6,Ctsc,Rab7a,Cep295nl,Gstp1,Cpped1,Tspan14,Nme2,Nfkb1,Arl8a |
| Parp14,Parp10,Parp9,Nudt12,Parp8,Nampt,Parp4 |
| Cd38,Parp14,Parp10,Parp9,Nudt12,Parp8,Nampt,Parp4 |
| Birc3,Casp4,Aim2,P2rx7,Nod1,Pstpip1,Ripk2,Tnfaip3,Birc2,Nlrp3,Pycard,Mapk12,Nlrp1a,Tab1 |
| Apbb1ip,Rap1a,Tln1,Src,Vwf,Rap1b |
| Rela,Mag,Arhgef10,Plekhg2,Rhoa,Ripk2,Arhgef26,Akap13,Ywhae,Myd88,Fgd3,Arhgef40,Prex1,Arhgef19,Nfkb1,Nfkbia,Casp3 |
| Fyn,Lyn,Src |
| Gng12,Serping1,Lgals3bp,Lamp2,Tf,Apbb1ip,Hgf,Vcl,Plek,Timp1,Rap1a,Cd63,Lhfpl2,Fcer1g,Rhog,Csk,Rhoa,Rarres2,Maged2,Tln1,Sparc,Fyn,Lyn,Flna,Wdr1,Rac2,Cd9,P2ry12,Lcp2,Mapk3,Src,Srgn,Stxbp3,Vwf,Rap1b,Itih3,Gnai3,Sod1,Gnai2,Tagln2 |
| Serping1,Lgals3bp,Lamp2,Tf,Hgf,Vcl,Plek,Timp1,Cd63,Lhfpl2,Rarres2,Maged2,Tln1,Sparc,Flna,Wdr1,Cd9,Srgn,Vwf,Itih3,Sod1,Tagln2 |
| Hist1h1d,Birc3,Tjp2,Ripk1,Casp7,Mlkl,Bid,Flot2,Bak1,Cflar,Dbnl,Birc2,Mapk3,Ywhae,Diablo,Xiap,Tjp1,Traf2,Casp3 |
| Birc3,Ripk1,Mlkl,Flot2,Cflar,Birc2,Xiap,Traf2 |
| Rsu1,Parvb,Pxn,Lims1 |
| Birc3,Ripk1,Mlkl,Flot2,Cflar,Birc2,Xiap,Traf2 |
| Ccnd1,Cdk6,Pml,Cbfb |
| Birc3,Ripk1,Tnfaip3,Cflar,Birc2,Otud7b,Xiap,Sppl2a,Traf2 |
| Serping1,Lgals3bp,Lamp2,Tf,Hgf,Vcl,Plek,Timp1,Cd63,Lhfpl2,Rarres2,Maged2,Tln1,Sparc,Flna,Wdr1,Cd9,Srgn,Stxbp3,Vwf,Itih3,Sod1,Tagln2 |
| Rela,Myd88,Nfkb1,Nfkb2,Nfkbia |
| Birc3,Ripk1,Mlkl,Flot2,Cflar,Birc2,Xiap,Traf2 |
| Tcf7l1,Tcf7l2,Tcf7 |
| Col5a3,Gab1,Lama4,Stat3,Hgf,Rap1a,Nras,Col27a1,Tns3,Src,Lrig1,Col5a2,Rap1b,Lama5,Stam2 |
| Ccnd1,Gpnmb,Stat3,Nras,Cdk4,Rhoa,Arap1,Pxn,Cdkn1b,Egfr |
| Ccnd1,Gpnmb,Stat3,Nras,Cdk4,Rhoa,Arap1,Pxn,Cdkn1b,Egfr |

| |
|---|
| Arhgap31,Tubb2b,Srgap1,Arpc1b,Arhgap42,Kif14,Nckap1l,Mapre1,Ncf1,Ppp1r14a,Arhgef10,Ctnna1,Rhoc,Plekhg2,Myo9b,Mylk,Rhog,Rhoa,Rhoj,Ppp2r1b,Myh11,Arhgap17,Arap1,Syde1,Arhgef26,Arhgap30,Tubb6,Akap13,Flna,Rac2,Mapk3,Wasf2,Ywhae,Cyba,Nde1,Myh14,Fmnl3,Iqgap1,Arhgap45,Cdc20,Tax1bp3,Cenpe,Fgd3,Arhgap4,Kif5b,Ktn1,Pfn1,Rnf135,Arhgef40,Prex1,Arhgef19,H2afv |
| Arhgap31,Tubb2b,Srgap1,Arpc1b,Arhgap42,Kif14,Nckap1l,Mapre1,Ncf1,Ppp1r14a,Arhgef10,Ctnna1,Rhoc,Plekhg2,Myo9b,Mylk,Rhog,Rhoa,Rhoj,Ppp2r1b,Myh11,Arhgap17,Arap1,Syde1,Arhgef26,Arhgap30,Tubb6,Akap13,Flna,Rac2,Mapk3,Wasf2,Ywhae,Cyba,Nde1,Myh14,Fmnl3,Iqgap1,Arhgap45,Cdc20,Tax1bp3,Cenpe,Fgd3,Arhgap4,Kif5b,Ktn1,Pfn1,Rnf135,Arhgef40,Prex1,Arhgef19,H2afv |
| Ctnnd1,Nckap1l,Ncf1,Ctnna1,Nras,Rhoa,Fyn,Pgf,Pxn,Src,Wasf2,Axl,Cyba,Mapkapk2,Itgav,Mapk12 |
| Casp7,Diablo,Xiap |
| Casp7,Diablo,Xiap,Casp3 |
| Casp7,Diablo,Xiap |
| Vcl,Mylk,Sorbs1,Myh11,Tln1,Pxn,Myl12a,Tpm4,Sorbs3 |
| Birc3,Ripk1,Tnfaip3,Cflar,Birc2,Otud7b,Xiap,Tab1,Sppl2a,Traf2 |
| Birc3,Ripk1,Tnfaip3,Birc2,Otud7b,Xiap,Tab1,Traf2 |
| Cdkn1a,Cdkn1c,Ccne2,Cdkn1b,Ccna2 |
| Ccnd1,Tead3,Psmb10,Psmb8,Psme2,Tp53,Tcf7l1,Src,Tcf7l2,Psmf1,Mdm2,Cbfb,Tcf7 |
| Ctnnd1,Nckap1l,Ncf1,Ctnna1,Nras,Rhoa,Fyn,Pxn,Src,Wasf2,Axl,Cyba,Mapkapk2,Itgav,Mapk12 |
| Rela,Myd88,Nfkb1,Nfkb2,Nfkbia |

| Source |
| --- |
| REAC |
| REAC |
| REAC |
| REAC |
| REAC |
| REAC |
| REAC |
| REAC |
| REAC |
| REAC |
| REAC |

| Term Name |
| --- |
| Metabolism of amino acids and derivatives |
| Phenylalanine and tyrosine metabolism |
| Tyrosine catabolism |
| Fcgamma receptor (FCGR) dependent phagocytosis |
| FCGR activation |
| Immune System |
| Innate Immune System |
| NOD1/2 Signaling Pathway |
| Nucleotide-binding domain, leucine rich repeat containing receptor (NLR) signaling pathways |
| Platelet homeostasis |
| Platelet sensitization by LDL |

| Term ID | p-value | Directionality | Intersection |
|---|---|---|---|
| REAC:R-RNO-71291 | 0.0415 | Up | Hpd |
| REAC:R-RNO-8963691 | 0.0031 | Up | Hpd |
| REAC:R-RNO-8963684 | 0.0031 | Up | Hpd |
| REAC:R-RNO-2029480 | 0.0396 | Down | Fgr |
| REAC:R-RNO-2029481 | 0.0239 | Down | Fgr |
| REAC:R-RNO-168256 | 0.0459 | Down | Fgr,Casp4 |
| REAC:R-RNO-168249 | 0.0239 | Down | Fgr,Casp4 |
| REAC:R-RNO-168638 | 0.0239 | Down | Casp4 |
| REAC:R-RNO-168643 | 0.0239 | Down | Casp4 |
| REAC:R-RNO-418346 | 0.0239 | Down | Fgr |
| REAC:R-RNO-432142 | 0.0092 | Down | Fgr |
