## Supplementary material for "Investigating cocaine- and abstinence-induced effects on astrocyte gene expression in the nucleus accumbens": Experiment 2_Pathway Enrichment Analysis

| Source | Term Name |
|---|---|
| REAC | Activation of NMDA receptors and postsynaptic events |
| REAC | Ca-dependent events |
| REAC | Calmodulin induced events |
| REAC | CaM pathway |
| REAC | CREB1 phosphorylation through the activation of Adenylate Cycl |
| REAC | DAG and IP3 signaling |
| REAC | G alpha (s) signalling events |
| REAC | G-protein mediated events |
| REAC | GPCR downstream signalling |
| REAC | Intracellular signaling by second messengers |
| REAC | MAPK family signaling cascades |
| REAC | Neuronal System |
| REAC | Neurotransmitter receptors and postsynaptic signal transmission |
| REAC | Opioid Signalling |
| REAC | PKA activation in glucagon signalling |
| REAC | PLC beta mediated events |
| REAC | Post NMDA receptor activation events |
| REAC | Presynaptic depolarization and calcium channel opening |
| REAC | Protein-protein interactions at synapses |
| REAC | Signal Transduction |
| REAC | Signaling by GPCR |
| REAC | Signaling by NTRK1 (TRKA) |

| | |
|---|---|
| REAC | Transmission across Chemical Synapses |
| REAC | Cholesterol biosynthesis |
| REAC | Metabolism |

| Term ID | p-value | Directionality |
|---|---|---|
| REAC:R-RNO-442755 | 2.4089E-06 | Up |
| REAC:R-RNO-111996 | 3.3300E-05 | Up |
| REAC:R-RNO-111933 | 0.0005 | Up |
| REAC:R-RNO-111997 | 0.0005 | Up |
| REAC:R-RNO-442720 | 0.0454 | Up |
| REAC:R-RNO-148950 | 0.0005 | Up |
| REAC:R-RNO-418555 | 1.7119E-05 | Up |
| REAC:R-RNO-112040 | 0.0004 | Up |
| REAC:R-RNO-388396 | 0.0009 | Up |
| REAC:R-RNO-900692 | 0.0235 | Up |
| REAC:R-RNO-568305 | 0.0243 | Up |
| REAC:R-RNO-112316 | 1.3215E-08 | Up |
| REAC:R-RNO-112314 | 0.0001 | Up |
| REAC:R-RNO-111885 | 0.0078 | Up |
| REAC:R-RNO-164378 | 0.0454 | Up |
| REAC:R-RNO-112043 | 0.0001 | Up |
| REAC:R-RNO-438064 | 0.0026 | Up |
| REAC:R-RNO-112308 | 0.0053 | Up |
| REAC:R-RNO-679436 | 0.0168 | Up |
| REAC:R-RNO-162582 | 0.0001 | Up |
| REAC:R-RNO-372790 | 0.0036 | Up |
| REAC:R-RNO-187037 | 0.0140 | Up |

| | | |
|---|---|---|
| REAC:R-RNO-112315 | 2.7723E-07 | Up |
| REAC:R-RNO-191273 | 0.0174 | Down |
| REAC:R-RNO-143072 | 0.0465 | Down |

| Intersection |
| --- |
| ACTN2,CAMK4,PRKAR2B,DLG2,GRIA1,PRKACB,RPS6KA2,PRKAR1B,DLG3 |
| PDE1B,CAMK4,PRKAR2B,PRKACB,PRKAR1B,MAPK1,PRKCA |
| PDE1B,CAMK4,PRKAR2B,PRKACB,PRKAR1B,PRKCA |
| PDE1B,CAMK4,PRKAR2B,PRKACB,PRKAR1B,PRKCA |
| PRKAR2B,PRKACB,PRKAR1B |
| PDE1B,CAMK4,PRKAR2B,PRKACB,PRKAR1B,PRKCA |
| PDE10A,PDE7B,PDE3A,PDE1B,PRKAR2B,PRKACB,PRKAR1B |
| PDE1B,CAMK4,PRKAR2B,PRKACB,PRKAR1B,MAPK1,PRKCA |
| PENK,RGS9,P2RY1,PRKCH,PDE10A,PDE7B,OPN3,PDE3A,PDE1B,PPP1R1B,HTR1B,TIAM1,RGS2,HTR2C,CAMK4,PRKAR2B,GRM5,GRM4,PDPK1,RGS8,PRKACB,RPS6KA2,DGKI,PRKAR1B,MAPK1,PRKCA,GRB2 |
| PDE1B,KIT,KL,KITLG,RRAGD,CAMK4,PRKAR2B,PDPK1,PRKACB,PIP4K2B,PRKAR1B,IRS2,CBX6,MAPK1,PRKCA,GRB2 |
| PTPN7,KIT,CDC42EP3,KL,PRKG2,ACTN2,KITLG,SPTB,SPTBN4,DLG2,PRKACB,DUSP8,SHC2,DLG3,IRS2,MAPK1,GRB2 |
| KCNJ2,KCNA5,IL1RAPL2,ACTN2,KCNS2,GAD1,CACNB4,CAMK4,CACNB2,PRKAR2B,KCNAB1,KCNH5,GRM5,CACNA2D3,DLG2,LRRTM1,GABRA4,CACNB3,GRIA1,PRKACB,EPB41L1,RPS6KA2,GABRG3,DNAJC5,PRKAR1B,DLG3,SLITRK5,PRKCA |
| KCNJ2,ACTN2,CAMK4,PRKAR2B,DLG2,GABRA4,GRIA1,PRKACB,EPB41L1,RPS6KA2,GABRG3,PRKAR1B,DLG3,PRKCA |
| PDE1B,PPP1R1B,CAMK4,PRKAR2B,PRKACB,PRKAR1B,MAPK1,PRKCA |
| PRKAR2B,PRKACB,PRKAR1B |
| PDE1B,CAMK4,PRKAR2B,PRKACB,PRKAR1B,MAPK1,PRKCA |
| CAMK4,PRKAR2B,PRKACB,RPS6KA2,PRKAR1B |
| CACNB4,CACNB2,CACNA2D3,CACNB3 |
| IL1RAPL2,GRM5,DLG2,LRRTM1,GRIA1,EPB41L1,DLG3,SLITRK5 |
| PTPN7,IQGAP3,PENK,RGS9,P2RY1,PRKCH,ADORA2A,PDE10A,TEC,PDE7B,DRD2,OPN3,PDE3A,PDE1B,KIT,CDC42EP3,WIPF3,KL,PRKG2,ACTN2,PPP1R1B,HTR1B,PPP3CA,EGR2,TIAM1,KITLG,RARB,SPTB,RRAGD,ACTN1,RGS2,HTR2C,CRABP1,LRRK2,CAMK4,FLRT3,PRKAR2B,TLE1,GRM5,SMAD3,SPTBN4,DLG2,GRM4,PDPK1,RGS8,PRKACB,STARD13,DUSP8,RHOBTB2,CYFIP2,ARHGAP26,SRF,RPS6KA2,PTK2B,DGKI,PIP4K2B,SHC2,MATK,RPS6KA5,NDEL1,PPP1R12B,GGA3,NCKIPSD,PRKAR1B,DLG3,IRS2,FAM13B,CBX6,MAPK1,CTNNBIP1,SLITRK5,DYNLL2,PRKCA,ARMCX3,CCNK,RNF111,BTRC,KHDRBS1,GRB2,DDX5 |
| PENK,RGS9,P2RY1,PRKCH,ADORA2A,PDE10A,PDE7B,DRD2,OPN3,PDE3A,PDE1B,PPP1R1B,HTR1B,TIAM1,RGS2,HTR2C,CAMK4,PRKAR2B,GRM5,GRM4,PDPK1,RGS8,PRKACB,RPS6KA2,DGKI,PRKAR1B,MAPK1,PRKCA,GRB2 |
| EGR2,SRF,RPS6KA2,SHC2,RPS6KA5,IRS2,MAPK1,GRB2 |

| |
|---|
| KCNJ2,ACTN2,GAD1,CACNB4,CAMK4,CACNB2,PRKAR2B,CACNA2D3,DLG2,GABRA4,CACNB3,GRIA1,PRKACB,EPB41L1,RPS6KA2,GABRG3,DNAJC5,PRKAR1B,DLG3,PRKCA |
| FDFT1,HMGCS1,SQLE,CYP51,MSMO1,DHCR7 |
| COASY,COMT,ODC1,NDST1,NADK,GLO1,IDH1,FASN,SDHC,SUCLG2,HADH,SPTLC1,RAP1A,PNPLA2,DDAH1,SLC7A5,NADK2,ACSS2,GNG5,GCDH,PRELP,OPLAH,LPCAT3,CERS4,NQO1,GLB1L,FDFT1,HEXA,HMGCS1,SQLE,CYP51,B3GNT7,ABCD1,MOCOS,HS3ST3B1,GSTA1,FA2H,MCEE,FABP7,GNG11,PLEKHA4,UGT1A6,ITPKB,HAS2,LIPT1,PTGDS,MID1IP1,TSPO,ACOXL,MSMO1,DHCR7,ABHD4,PARP14,GLTP,SCD1,CHST7,TNFAIP8L2,SAT1,BGN,EPHX1,VCAN,ENPP6,NEU4,PLD4,CHST5 |

| Source | Term Name |
|---|---|
| REAC | Neuronal System |
| REAC | Transmission across Chemical Synapses |
| REAC | Neurotransmitter receptors and postsynaptic signal transmission |
| REAC | Potassium Channels |
| REAC | Voltage gated Potassium channels |
| REAC | Protein-protein interactions at synapses |
| REAC | Opioid Signalling |
| REAC | Activation of NMDA receptors and postsynaptic events |
| REAC | PLC beta mediated events |
| REAC | G-protein mediated events |
| REAC | Ca-dependent events |
| REAC | Glutamate binding, activation of AMPA receptors and synaptic plasti |
| REAC | Trafficking of AMPA receptors |
| REAC | Neurexins and neuroligins |

| | |
|---|---|
| REAC | Signal Transduction |
| REAC | DAG and IP3 signaling |
| REAC | Calmodulin induced events |
| REAC | CaM pathway |
| REAC | Unblocking of NMDA receptors, glutamate binding and activation |
| REAC | Trafficking of GluR2-containing AMPA receptors |
| REAC | Cardiac conduction |
| REAC | Signaling by NTRK1 (TRKA) |
| REAC | Integration of energy metabolism |
| REAC | Muscle contraction |
| REAC | Regulation of insulin secretion |
| REAC | Signaling by NTRKs |
| REAC | WNT5A-dependent internalization of FZD4 |
| REAC | DARPP-32 events |

| | |
|---|---|
| REAC | Axon guidance |
| REAC | Nervous system development |
| REAC | GPCR downstream signalling |
| REAC | RHO GTPase Effectors |
| REAC | Dopamine Neurotransmitter Release Cycle |
| REAC | G alpha (s) signalling events |
| REAC | G alpha (i) signalling events |
| REAC | L1CAM interactions |
| REAC | MAPK family signaling cascades |
| REAC | Signaling by Receptor Tyrosine Kinases |
| REAC | Serotonin Neurotransmitter Release Cycle |
| REAC | Amine ligand-binding receptors |
| REAC | Recycling pathway of L1 |
| REAC | Antigen processing: Ubiquitination & Proteasome degradation |

| | |
|---|---|
| REAC | Metabolism |
| REAC | RHOQ GTPase cycle |
| REAC | Neutrophil degranulation |
| REAC | RHO GTPase cycle |

| | |
|---|---|
| REAC | Innate Immune System |
| REAC | Metabolism of carbohydrates |
| REAC | CS/DS degradation |
| REAC | Sphingolipid metabolism |
| REAC | Glycosphingolipid metabolism |
| REAC | Branched-chain amino acid catabolism |
| REAC | mitochondrial fatty acid beta-oxidation of saturated fatty acids |
| REAC | Beta oxidation of lauroyl-CoA to decanoyl-CoA-CoA |

| Term ID | p-value | Directionality |
|---|---|---|
| REAC:R-RNO-112316 | 8.4919E-32 | Up |
| REAC:R-RNO-112315 | 2.8334E-14 | Up |
| REAC:R-RNO-112314 | 1.6822E-10 | Up |
| REAC:R-RNO-1296071 | 8.8382E-10 | Up |
| REAC:R-RNO-1296072 | 4.0869E-09 | Up |
| REAC:R-RNO-6794362 | 2.0180E-07 | Up |
| REAC:R-RNO-111885 | 1.1068E-06 | Up |
| REAC:R-RNO-442755 | 4.1030E-06 | Up |
| REAC:R-RNO-112043 | 5.9554E-06 | Up |
| REAC:R-RNO-112040 | 8.6782E-06 | Up |
| REAC:R-RNO-111996 | 1.0750E-05 | Up |
| REAC:R-RNO-399721 | 1.7910E-05 | Up |
| REAC:R-RNO-399719 | 1.7910E-05 | Up |
| REAC:R-RNO-6794361 | 2.1920E-05 | Up |

| | | |
|---|---|---|
| REAC:R-RNO-162582 | 4.7512E-05 | Up |
| REAC:R-RNO-1489509 | 0.0001 | Up |
| REAC:R-RNO-111933 | 0.0001 | Up |
| REAC:R-RNO-111997 | 0.0001 | Up |
| REAC:R-RNO-438066 | 0.0001 | Up |
| REAC:R-RNO-416993 | 0.0002 | Up |
| REAC:R-RNO-5576891 | 0.0002 | Up |
| REAC:R-RNO-187037 | 0.0002 | Up |
| REAC:R-RNO-163685 | 0.0004 | Up |
| REAC:R-RNO-397014 | 0.0004 | Up |
| REAC:R-RNO-422356 | 0.0004 | Up |
| REAC:R-RNO-166520 | 0.0016 | Up |
| REAC:R-RNO-5099900 | 0.0025 | Up |
| REAC:R-RNO-180024 | 0.0025 | Up |

| | | |
|---|---|---|
| REAC:R-RNO-422475 | 0.0031 | Up |
| REAC:R-RNO-9675108 | 0.0037 | Up |
| REAC:R-RNO-388396 | 0.0056 | Up |
| REAC:R-RNO-195258 | 0.0068 | Up |
| REAC:R-RNO-212676 | 0.0143 | Up |
| REAC:R-RNO-418555 | 0.0152 | Up |
| REAC:R-RNO-418594 | 0.0153 | Up |
| REAC:R-RNO-373760 | 0.0169 | Up |
| REAC:R-RNO-5683057 | 0.0181 | Up |
| REAC:R-RNO-9006934 | 0.0197 | Up |
| REAC:R-RNO-181429 | 0.0299 | Up |
| REAC:R-RNO-375280 | 0.0317 | Up |
| REAC:R-RNO-437239 | 0.0346 | Up |
| REAC:R-RNO-983168 | 0.0410 | Up |

| | | |
|---|---|---|
| REAC:R-RNO-1430728 | 2.9828E-07 | Down |
| REAC:R-RNO-9013406 | 0.0001 | Down |
| REAC:R-RNO-6798695 | 0.0004 | Down |
| REAC:R-RNO-9012999 | 0.0007 | Down |

| | | |
|---|---|---|
| REAC:R-RNO-168249 | 0.0048 | Down |
| REAC:R-RNO-71387 | 0.0094 | Down |
| REAC:R-RNO-2024101 | 0.0125 | Down |
| REAC:R-RNO-428157 | 0.0156 | Down |
| REAC:R-RNO-1660662 | 0.0175 | Down |
| REAC:R-RNO-70895 | 0.0188 | Down |
| REAC:R-RNO-77286 | 0.0221 | Down |
| REAC:R-RNO-77310 | 0.0221 | Down |

| Intersection |
| --- |
| KCNQ5,ACTN2,CAMK4,KCNAB1,PRKCG,PRKCB,CACNB4,KCNS2,KCNA5,LIN7B,KCNA1,HOMER1,KCNK4,SNAP25,KCNG2,KCNH7,SLC1A1,CACNB2,KCNF1,KCNJ3,KCNH3,CACNB3,GABRA4,GABRB3,LRRC7,LRRTM3,ADCY1,SHANK2,KCNJ9,ABCC8,RPS6KA2,KCNJ2,CACNG3,SHANK1,DLG3,GRM5,KCNB1,PRKAR2B,NCALD,CACNG2,NEFL,KCNK2,GRIA1,SLITRK3,GRIN1,CALM1,PANX1,KCNH5,DLGAP1,STXBP1,GABRG2,DLG2,SYT1,KCNV1,PPFIA2,KCNAB3,KCNK6,VAMP2,SYN1,PRKAR1B,EPB41L1,SLITRK5,CHRNB2,CACNA2D3,LRRTM2,KCNJ11,KCNK1,LRRTM1,SLITRK1,KCNMA1,GNG3,PANX2,AP2M1,KCNC2,PRKACB,KCNK3,LRFN1,KCNA3,KCNA2,DNAJC5,GNAI1,AP2A1,KCNC1,AP2S1,GLRB,KCND1,AP2B1,LRRTM4,GAD1,LRFN4,RTN3,NLGN2,PRKCA,CAMK2G,GNB1 |
| ACTN2,CAMK4,PRKCG,PRKCB,CACNB4,LIN7B,SNAP25,SLC1A1,CACNB2,KCNJ3,CACNB3,GABRA4,GABRB3,LRRC7,ADCY1,KCNJ9,RPS6KA2,KCNJ2,CACNG3,DLG3,PRKAR2B,NCALD,CACNG2,NEFL,GRIA1,GRIN1,CALM1,STXBP1,GABRG2,DLG2,SYT1,PPFIA2,VAMP2,SYN1,PRKAR1B,EPB41L1,CHRNB2,CACNA2D3,GNG3,AP2M1,PRKACB,DNAJC5,GNAI1,AP2A1,AP2S1,GLRB,AP2B1,GAD1,PRKCA,CAMK2G,GNB1 |
| ACTN2,CAMK4,PRKCG,PRKCB,KCNJ3,GABRA4,GABRB3,LRRC7,ADCY1,KCNJ9,RPS6KA2,KCNJ2,CACNG3,DLG3,PRKAR2B,NCALD,CACNG2,NEFL,GRIA1,GRIN1,CALM1,GABRG2,DLG2,PRKAR1B,EPB41L1,CHRNB2,GNG3,AP2M1,PRKACB,GNAI1,AP2A1,AP2S1,GLRB,AP2B1,PRKCA,CAMK2G,GNB1 |
| KCNQ5,KCNAB1,KCNS2,KCNA5,KCNA1,KCNK4,KCNG2,KCNH7,KCNF1,KCNJ3,KCNH3,KCNJ9,ABCC8,KCNJ2,KCNB1,KCNK2,KCNH5,KCNV1,KCNAB3,KCNK6,KCNJ11,KCNK1,KCNMA1,GNG3,KCNC2,KCNK3,KCNA3,KCNA2,KCNC1,KCND1,GNB1 |
| KCNQ5,KCNAB1,KCNS2,KCNA5,KCNA1,KCNG2,KCNH7,KCNF1,KCNH3,KCNB1,KCNH5,KCNV1,KCNAB3,KCNC2,KCNA3,KCNA2,KCNC1,KCND1 |
| HOMER1,LRRTM3,SHANK2,SHANK1,DLG3,GRM5,GRIA1,SLITRK3,GRIN1,DLGAP1,DLG2,PPFIA2,EPB41L1,SLITRK5,LRRTM2,LRRTM1,SLITRK1,LRFN1,LRRTM4,LRFN4,RTN3,NLGN2 |
| CAMK4,PRKCG,PDE1B,ADCY1,PRKAR2B,CALM1,PDE1A,MAPK1,PRKAR1B,PPP1R1B,PDE4D,PLCB1,GNG3,PRKACB,GNAI1,PPP1CA,PRKCA,PPP2R1A,GNB1,PPP2CA |
| ACTN2,CAMK4,LRRC7,RPS6KA2,DLG3,PRKAR2B,NEFL,GRIA1,GRIN1,CALM1,DLG2,PRKAR1B,PRKACB,CAMK2G |
| CAMK4,PRKCG,PDE1B,ADCY1,PRKAR2B,CALM1,PDE1A,MAPK1,PRKAR1B,PLCB1,PRKACB,PRKCA |
| CAMK4,PRKCG,PDE1B,ADCY1,PRKAR2B,CALM1,PDE1A,MAPK1,PRKAR1B,PLCB1,PRKACB,GNAI1,PRKCA |
| CAMK4,PRKCG,PDE1B,ADCY1,PRKAR2B,CALM1,PDE1A,MAPK1,PRKAR1B,PRKACB,PRKCA |
| PRKCG,PRKCB,CACNG3,CACNG2,GRIA1,EPB41L1,AP2M1,AP2A1,AP2S1,AP2B1,PRKCA,CAMK2G |
| PRKCG,PRKCB,CACNG3,CACNG2,GRIA1,EPB41L1,AP2M1,AP2A1,AP2S1,AP2B1,PRKCA,CAMK2G |
| HOMER1,LRRTM3,SHANK2,SHANK1,DLG3,GRM5,DLGAP1,DLG2,EPB41L1,LRRTM2,LRRTM1,LRRTM4,NLGN2 |

| |
|---|
| EGR2,NR4A1,PDE10A,VIPR1,IQGAP3,PENK,CNKSR2,ACTN2,RGS9,PPP3CA,CAMK4,HRH3,PRKCG,ADORA2A,ARHGEF3,WIPF3,PDE1B,WNT10A,NAB2,CDC42EP3,CYFIP2,PRKCB,GRM4,BAIAP2,HTR1B,RGS14,TGFB3,LIN7B,MCHR1,PAK6,LRRK2,PTPN7,CHRM1,ACTN1,RGS2,ADRA2C,FMNL1,DGKZ,DGKG,PIK3CD,TIAM1,LRRC7,WNT10B,PRKCZ,PTPRJ,SHC2,ADCY1,PDE7B,FST,PTK2B,DGKI,RPS6KA2,WNT9B,SH3BP1,DLG3,RARB,BDNF,GRM5,ARHGAP20,PRKAR2B,KITLG,DRD2,NLK,NEFL,OPN3,SLITRK3,YWHAH,IGF1,GRIN1,CALM1,ATP6V1G2,OPRK1,SSTR4,PDE1A,MATK,PMEPA1,ATP6V1B2,GFOD1,PDPK1,KL,GGA3,MAPK1,DLG2,DUSP8,HTR2A,LINGO1,PPP1R12B,FGF16,PRKAR1B,DUSP2,RGS4,SLITRK5,GRM7,THBS1,SSTR1,PPP1R1B,GNAZ,SEMA4F,ATP6V1C1,HTR2C,SRF,PRKCH,PTPRU,DUSP7,PAK1,CBX6,SPTB,IER3,PDE4D,WNT16,PLCB1,TACR2,DUSP6,ZWINT,SSTR3,CHRM3,ADRA2B,LGR5,OXTR,KSR1,INCENP,YWHAZ,RANGAP1,SPATA2,TUBA4A,TUBA8,STMN2,CHRM2,ARHGEF2,KIT,PPP2R5B,OPRD1,RGS8,KLC1,TEC,KHDRBS2,ACVR1B,PIP4K2B,NDEL1,CRABP1,SPTBN4,RGS17,GNG3,PTEN,AP2M1,TBK1,SMAD3,NCK2,HECW1,RASAL2,YWHAG,UHMK1,PRKACB,TRAK2,GNAI1,CNR1,ARMCX3,CLIP1,AP2A1,HIST3H2BA,CLIP3,SPRED3,PFN2,TUBB3,RGS7,DYNLL1,AP2S1,ATP6V1A,PPP1CC,ATP6V1H,YWHAB,ATP6V0E2,ZFYVE9,MTMR1,ABHD17A,MKRN1,AP2B1,THEM4,ULK1,FAM13B,DNAJB1,ATP6V0C,USP13,SMURF1,RHOBTB2,MTMR4,PRKAG2,ARPC4,NDUFA5,CARM1,NPHP4,PYGO2,HMOX2,TBL1XR1,SPRED2,DYNLL2,PPP1CA,BTRC,ABL2,ATP6V1D,ZDHHC21,PAFAH1B1,CASP9,GRB2,DISP2,RTKN,PARP1,PRKCA,OPHN1,RALGAPA1,VPS29,DAGLA,CENPC,FLRT2,IRS2,DGKE,SHOC2,PDK3,PPP1R12A,NEDD8,PAK3,PSMC6,MCF2L,RTN4,DYNC1LI2,CTNNBIP1,CAMK2G,NCKIPSD,TNKS2,HSP90AB1,RANBP2,GATAD2B,STUB1,GOLGA3,PPP2R1A,DLAT,RNF111,CTBP1,CLTA,UBE2M,GNB1,GNAO1,CAB39,PSMC5,RHOT1,PPP2CA,PRKCI |
| CAMK4,PRKCG,PDE1B,ADCY1,PRKAR2B,CALM1,PDE1A,PRKAR1B,PRKACB,PRKCA |
| CAMK4,PRKCG,PDE1B,ADCY1,PRKAR2B,CALM1,PDE1A,PRKAR1B,PRKACB,PRKCA |
| CAMK4,PRKCG,PDE1B,ADCY1,PRKAR2B,CALM1,PDE1A,PRKAR1B,PRKACB,PRKCA |
| ACTN2,LRRC7,DLG3,NEFL,GRIA1,GRIN1,CALM1,DLG2,CAMK2G |
| PRKCG,PRKCB,GRIA1,AP2M1,AP2A1,AP2S1,AP2B1,PRKCA |
| KCNA5,KCNK4,CACNB2,ATP2B2,KCNJ2,NPPA,ATP1A1,KCNK2,CALM1,MME,KCNK6,SLC8A1,KCNK12,KCNJ11,KCNK1,ATP2B3,KCNK3,KCND1,SLC8A3,KCNIP2,CAMK2G |
| EGR2,NAB2,SHC2,RPS6KA2,MAPK1,SRF,DUSP7,DUSP6,AP2M1,AP2A1,AP2S1,YWHAB,AP2B1,GRB2,IRS2,PPP2R1A,CLTA,PPP2CA |
| KCNG2,CACNB2,ADRA2C,CACNB3,ABCC8,KCNB1,PRKAR2B,ACSL4,PRKAR1B,KCNJ11,PLCB1,GNG3,KCNC2,PRKACB,GNAI1,PRKAG2,PRKCA,GNB1 |
| ACTN2,KCNA5,KCNK4,CACNB2,TPM1,ATP2B2,KCNJ2,NPPA,ATP1A1,KCNK2,CALM1,MME,ACTA1,KCNK6,SLC8A1,TNNC2,KCNK12,PAK1,KCNJ11,KCNK1,ATP2B3,KCNK3,ACTA2,KCND1,SLC8A3,KCNIP2,CAMK2G,MYL12B |
| KCNG2,CACNB2,ADRA2C,CACNB3,ABCC8,KCNB1,ACSL4,KCNJ11,PLCB1,GNG3,KCNC2,PRKACB,GNAI1,PRKCA,GNB1 |
| EGR2,NAB2,SHC2,RPS6KA2,BDNF,MAPK1,SRF,DUSP7,DUSP6,AP2M1,AP2A1,AP2S1,YWHAB,AP2B1,GRB2,IRS2,PPP2R1A,CLTA,PPP2CA |
| PRKCG,PRKCB,AP2M1,AP2A1,AP2S1,AP2B1,PRKCA,CLTA |
| PRKAR2B,PRKAR1B,PPP1R1B,PDE4D,PRKACB,PPP1CA,PPP2R1A,PPP2CA |

| |
|---|
| SEMA3E,PIK3CD,TIAM1,CDK5R1,DOK6,GRIN1,DNM1,MAPK1,PAK1,SPTB,RAP1GAP,PLXNA2,TUBA4A,TUBA8,EFNB2,SPTBN4,AP2M1,CRMP1,NCK2,DSCAML1,PRKACB,AP2A1,TUBB3,AP2S1,GAP43,AP2B1,ARPC4,EPHA4,GRB2,PRKCA,IRS2,LYPLA2,PAK3,DPYSL5,HSP90AB1,MYL12B,CLTA,DPYSL2 |
| SEMA3E,PIK3CD,TIAM1,CDK5R1,DOK6,GRIN1,DNM1,MAPK1,PAK1,SPTB,RAP1GAP,PLXNA2,TUBA4A,TUBA8,EFNB2,SPTBN4,AP2M1,CRMP1,NCK2,DSCAML1,PRKACB,AP2A1,TUBB3,AP2S1,GAP43,AP2B1,ARPC4,EPHA4,GRB2,PRKCA,IRS2,LYPLA2,PAK3,DPYSL5,HSP90AB1,MYL12B,CLTA,DPYSL2 |
| PDE10A,PENK,RGS9,CAMK4,PRKCG,ARHGEF3,PDE1B,GRM4,HTR1B,RGS14,MCHR1,CHRM1,RGS2,ADRA2C,DGKZ,DGKG,TIAM1,ADCY1,PDE7B,DGKI,RPS6KA2,GRM5,PRKAR2B,OPN3,CALM1,OPRK1,SSTR4,PDE1A,PDPK1,MAPK1,HTR2A,PRKAR1B,RGS4,GRM7,SSTR1,PPP1R1B,GNAZ,HTR2C,PRKCH,PAK1,PDE4D,PLCB1,TACR2,SSTR3,CHRM3,ADRA2B,OXTR,CHRM2,ARHGEF2,OPRD1,RGS8,RGS17,GNG3,PRKACB,GNAI1,CNR1,RGS7,PPP1CA,GRB2,PRKCA,DAGLA,DGKE,MCF2L,PPP2R1A,GNB1,PPP2CA |
| [illegible] IQGAP3,WIPF3,CYFIP2,PRKCB,BAIAP2,LIN7B,FMNL1,PRKCZ,YWHAH,CALM1,PDPK1,MAPK1,PPP1R12B,SRF,PAK1,ZWINT,INCENP,YWHAZ,RANGAP1,TUBA4A,TUBA8,PPP2R5B,KLC1,NDEL1,YWHAG,CLIP1,HIST3H2BA,PFN2,TUBB3,DYNLL1,PPP1CC,YWHAB,ARPC4,DYNLL2,PAFAH1B1,GRB2,RTKN,PRKCA,CENPC,PPP1R12A,PAK3,DYNC1LI2,NCKIPSD,RANBP2,PPP2R1A,PPP2CA |
| LIN7B,SNAP25,STXBP1,SYT1,PPFIA2,VAMP2,SYN1 |
| PDE10A,PDE1B,PDE7B,PRKAR2B,PDE1A,PRKAR1B,PDE4D,PRKACB |
| PENK,RGS9,CAMK4,PRKCG,PDE1B,GRM4,HTR1B,RGS14,MCHR1,ADRA2C,ADCY1,PRKAR2B,OPN3,CALM1,OPRK1,SSTR4,PDE1A,MAPK1,PRKAR1B,RGS4,GRM7,SSTR1,PPP1R1B,PDE4D,PLCB1,SSTR3,ADRA2B,CHRM2,OPRD1,RGS8,RGS17,GNG3,PRKACB,GNAI1,CNR1,RGS7,PPP1CA,PRKCA,PPP2R1A,GNB1,PPP2CA |
| DNM1,MAPK1,PAK1,SPTB,TUBA4A,TUBA8,SPTBN4,AP2M1,AP2A1,TUBB3,AP2S1,GAP43,AP2B1,LYPLA2,CLTA,DPYSL2 |
| CNKSR2,ACTN2,CDC42EP3,PTPN7,LRRC7,SHC2,DLG3,KITLG,NEFL,GRIN1,CALM1,KL,MAPK1,DLG2,DUSP8,FGF16,DUSP2,DUSP7,PAK1,SPTB,DUSP6,KSR1,KIT,PPP2R5B,SPTBN4,RASAL2,PRKACB,SPRED3,PPP1CC,YWHAB,ABHD17A,DNAJB1,SPRED2,GRB2,IRS2,SHOC2,PAK3,PSMC6,CAMK2G,PPP2R1A,PSMC5,PPP2CA |
| EGR2,NAB2,CYFIP2,PRKCB,BAIAP2,PRKCZ,PTPRJ,SHC2,PTK2B,RPS6KA2,BDNF,KITLG,IGF1,CALM1,ATP6V1G2,MATK,ATP6V1B2,PDPK1,KL,GGA3,MAPK1,FGF16,THBS1,ATP6V1C1,SRF,PTPRU,DUSP7,PAK1,DUSP6,KIT,TEC,AP2M1,NCK2,PRKACB,AP2A1,AP2S1,ATP6V1A,ATP6V1H,YWHAB,ATP6V0E2,AP2B1,THEM4,ATP6V0C,SPRED2,ATP6V1D,GRB2,PRKCA,FLRT2,IRS2,PAK3,STUB1,PPP2R1A,CLTA,PPP2CA |
| SNAP25,STXBP1,SYT1,PPFIA2,VAMP2,SYN1 |
| HRH3,HTR1B,CHRM1,ADRA2C,DRD2,HTR2A,HTR2C,CHRM3,ADRA2B,CHRM2 |
| DNM1,MAPK1,TUBA4A,TUBA8,AP2M1,AP2A1,TUBB3,AP2S1,AP2B1,CLTA,DPYSL2 |
| LMO7,ASB2,FBXL16,RNF144B,HECW2,FBXO41,HACE1,FBXL15,ZBTB16,AREL1,FBXL19,PJA2,SIAH2,UBE2Q1,FBXO31,RNF14,CDC34,MKRN1,FBXO21,TRIM32,HERC3,SMURF1,UBE2K,RNF19B,RNF6,BTRC,TRIM9,ANAPC7,TRIM37,ASB6,CUL2,UBE3B,PSMC6,UBE2N,RNF19A,ZNRF2,UBE3A,BTBD6,STUB1,UBR1,UBE2Z,UBE2S,RNF111,UBE2M,LNPEP,PSMC5 |

| |
|---|
| MTMR3,NDUFA12,NDUFS7,ARSB,PTDSS2,UQCRFS1,PITPNB,PAICS,ADH5,PSMF1,RRM1,GCLC,EXT2,HACD3,BPGM,OSBPL9,NDST1,CSGALNACT1,PCYT1A,OAT,SPTLC1,DEGS1,AGPAT3,PANK3,NUDT3,MBOAT2,SC5D,ME2,GNS,CAD,ETFB,BCKDK,COASY,ABCD1,CDO1,NUDT12,ATIC,STARD7,PGD,COMT,VKORC1,RUFY1,RPE,MLYCD,PIK3C2A,HADHA,GALC,GMPR2,TMEM186,ACOT13,ACOX1,ADI1,G6PD,ARSA,TKFC,CHSY1,CD320,NAMPT,TMLHE,HSD17B11,HACL1,DBT,DDAH1,PAPSS1,ODC1,SMOX,AGPAT5,CHKA,ACP6,UGP2,GNPDA2,ACADSB,RAP1A,SPTSSA,HADH,HEXB,HSD17B12,MAPKAPK2,NNT,G6PC3,SDHC,ALDOC,BCKDHB,NADK,SHPK,NUDT5,ALDH7A1,SLC2A1,MAN2B2,GSTM5,AKR1B10,HEXA,MANBA,SAMHD1,GYS1,CYP2D4,SHMT2,AGL,MOCS1,MCCC2,CPT1A,CMBL,NT5C1A,GCDH,CERS4,ACY3,PGM2,IVD,SLC22A5,ACBD5,MCEE,DHCR7,HK2,IDUA,HIBCH,ACADL,GLYCTK,PSPH,PLBD1,NAT1,ACAD11,ABHD3,LRP1,ECHS1,PI4K2B,NMNAT3,GNAI2,RBKS,GCSH,CERS2,CAR5B,HADHB,HMGCS2,GSTT1,PHKG1,IDH1,ABHD14B,ARSK,SLC46A1,GNG5,TYMS,NADK2,CSPG5,CHPT1,GLB1,TNFAIP8,ADPGK,UGT8,CYP4F17,PFKFB3,ACSS3,ELOVL2,GLDC,FADS2,SDSL,NME4,FAH,INPPL1,DDAH2,ABCD4,SLC7A5,OPLAH,CHST7,SDS,AGMO,PTPN13,GLUD1,ALDH1L1,SMPD2,PAPSS2,ELOVL5,CAR13,NQO1,ENPP6,SDC4,HOGA1,ACSF2,PSAT1,HAS2,GLB1L,CBS,ACSS1,SLC25A21,PSME2,CYP2J3,SUCLG2,MAOB,PHYH,SDC1,RBP1,CPNE3,GNG12,MPST,MGST1,BGN,MTHFS,GSTK1,AMT,CAR2,ITPKB,FA2H,PRELP,GSTA1,SUMF2,CYP4V3,PNP,SREBF1,SLC6A11,EPHX1,PRODH,SAT1,LPCAT3,GNG11,MID1IP1,ABHD4,GLTP,B3GNT7,QPRT,CD38,FABP7,TSPO,VCAN,PARP14,PLD4,CHST5,NEU4 |
| RAB7A,ARL13B,SYDE1,ARHGAP5,TFRC,IQGAP1,SNAP23,CDC42EP4,VANGL1,PAK4,PREX1,PAK2,SCRIB,GJA1,FNBP1,STEAP3,ARHGAP17,STOM,CDC42EP1,SLC1A5 |
| RAB5B,GDI2,PA2G4,NCSTN,ARSB,SDCBP,RAB7A,ERP44,TXNDC5,MLEC,DEGS1,GCA,SURF4,GNS,RHOA,DNAJC3,ATP11A,CD59,ARSA,CPPED1,VAT1,XRCC5,S100A11,ANO6,CAT,RAB5C,RAP1A,CTSZ,AGA,CREG1,HEXB,TSPAN14,ALDOC,IQGAP1,DPP7,MANBA,SNAP23,AGL,XRCC6,CYFIP1,PGM2,IDH1,PRCP,DNASE1L1,GLB1,NPC2,NCKAP1L,STBD1,CHI3L1,RT1-M3-1,TCIRG1,RHOG,CTSH,LAMP2,CPNE3,MGST1,TMEM179B,VCL,GSN,TMBIM1,RNASET2,PNP,PTPRC,CD63,ADGRE5,STOM,PTAFR,RT1-S3,GSDMD,TNFRSF1B,FGR,CD68,B2M,GPR84,RT1-A1,BST2,ITGAL,ITGB2 |
| RAB7A,TMEM59,DDX39B,EMC3,ABCD3,BCAP31,RHOA,RND2,ARL13B,FARP1,RRAS2,STAM2,TNFAIP1,TOR1AIP1,SYDE1,ARHGEF12,DBT,ARHGAP5,ARHGAP12,TFRC,WASF2,SWAP70,IQGAP1,PREX2,SNAP23,SHMT2,DOCK7,ADD3,CYFIP1,ARHGAP11A,ACBD5,CDC42EP4,ARHGEF16,STK38,STARD8,VANGL1,NCKAP1L,RND3,GNA13,SYDE2,STBD1,RHPN2,PAK4,PREX1,PAK2,PTPN13,ANKFY1,SCRIB,FERMT2,RHOG,GJA1,FNBP1,STEAP3,PLEKHG1,ARHGAP18,ARHGAP17,VANGL2,TPM4,TMOD3,STOM,ARHGEF19,NCF1,CDC42EP1,SLC1A5,DOCK8,UACA,PLEKHG2,MCAM,RHOJ,DOCK6,EPSTI1,ARHGAP31 |

| |
|---|
| RAB5B,GDI2,PA2G4,RAF1,NCSTN,ABL1,ARSB,SDCBP,RAB7A,TAB1,CRK,CNPY3,PSMF1,HSP90B1,ERP44,TXNDC5,TAX1BP1,MLEC,DEGS1,GCA,SURF4,GNS,FYN,RHOA,DNAJC3,SRC,NLRX1,ATP11A,CD59,MAVS,ARSA,CPPED1,VAT1,XRCC5,TAB2,S100A11,ANO6,CAT,RAB5C,RAP1A,MYD88,CTSZ,AGA,CREG1,HEXB,WASF2,MAPKAPK2,TSPAN14,ALDOC,IQGAP1,DPP7,ATF1,MANBA,SNAP23,AGL,XRCC6,MAPKAPK3,CYFIP1,PGM2,NLRP3,NFATC3,IDH1,CASP8,PRCP,DNASE1L1,GLB1,MAPK12,NPC2,NCKAP1L,STBD1,RELA,CHI3L1,SIGLEC15,PAK2,C4B,RT1-M3-1,TCIRG1,TXNIP,RHOG,PSME2,MYO10,CTSH,LAMP2,CPNE3,NFKB2,MGST1,TMEM179B,VCL,GSN,TMBIM1,NFATC1,RNASET2,PNP,PTPRC,CD63,LYN,ADGRE5,NOD1,ARPC1B,STOM,PTAFR,NCF1,RT1-S3,GSDMD,TNFRSF1B,TLR9,FGR,CD68,B2M,GPR84,SERPING1,TRIM21,RT1- |
| ARSB,EXT2,BPGM,NDST1,CSGALNACT1,GNS,PGD,RPE,G6PD,TKFC,CHSY1,PAPSS1,UGP2,GNPDA2,HEXB,G6PC3,ALDOC,SHPK,SLC2A1,MAN2B2,HEXA,MANBA,GYS1,AGL,PGM2,HK2,IDUA,GLYCTK,RBKS,PHKG1,CSPG5,GLB1,ADPGK,PFKFB3,CHST7,PAPSS2,SDC4,HAS2,GLB1L,SDC1,BGN,PRELP,B3GNT7,VCAN,CHST5 |
| ARSB,HEXB,HEXA,IDUA,CSPG5,BGN,VCAN |
| ARSB,SPTLC1,DEGS1,GALC,ARSA,SPTSSA,HEXB,HEXA,CERS4,CERS2,ARSK,GLB1,UGT8,SMPD2,GLB1L,FA2H,SUMF2,GLTP,NEU4 |
| ARSB,GALC,ARSA,HEXB,HEXA,ARSK,GLB1,UGT8,SMPD2,GLB1L,SUMF2,GLTP,NEU4 |
| BCKDK,DBT,ACADSB,BCKDHB,MCCC2,IVD,HIBCH,ECHS1 |
| HADHA,HADH,ACADL,ECHS1,HADHB |
| HADHA,HADH,ACADL,ECHS1,HADHB |
