## Supplemental Figures for "Investigating cocaine- and abstinence-induced effects on astrocyte gene expression in the nucleus accumbens"

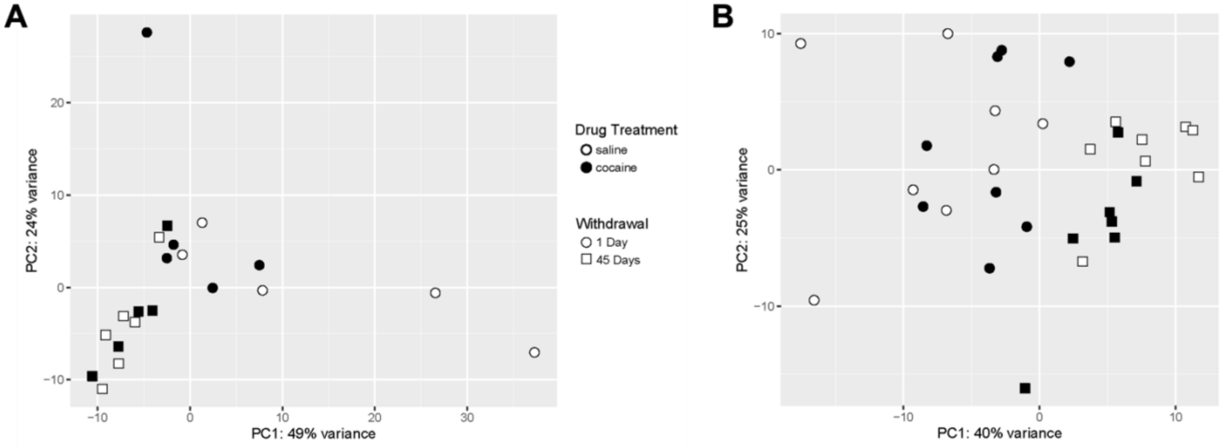


**Supplementary Figure 1**: *PCA plots showing that time had a larger effect on variance in RNA-seq results of both experiments.* PCA plots displaying variance of four experimental groups (SAL-WD1, SAL-WD45, COC-WD1, and COC-WD45), denoted by shape and color. PCA plot of experiment 1 shown on the left (S1A) and experiment 2 shown on the right (S1B).

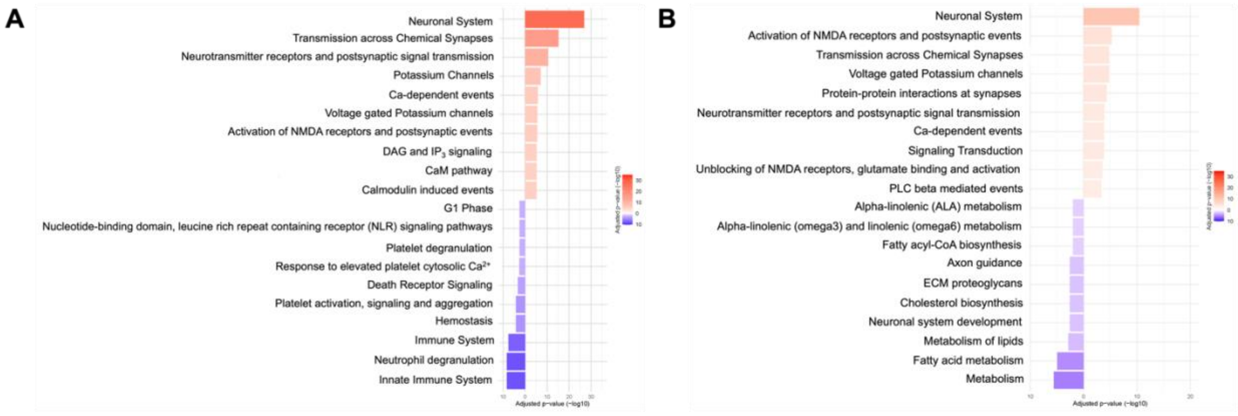


**Supplementary Figure 2**: *Pathway enrichment analyses for saline- and cocaine-treated rats across abstinence (WD45 vs. WD1) in experiment 1*. Pathway enrichment analysis displaying most significant cellular pathways within same drug treatment groups, saline (**S2A**) and cocaine (**S2B**) rats across 45 days of home cage abstinence. Downregulated pathways are shown in blue and upregulated pathways are shown in red (adjusted p-value < 0.05).
